# Human acrocentric chromosome short arm *de novo* mutation and recombination

**DOI:** 10.64898/2025.12.16.694519

**Authors:** Jiadong Lin, F. Kumara Mastrorosa, Michelle D. Noyes, DongAhn Yoo, Arang Rhie, David Porubsky, Kendra Hoekzema, Katherine M. Munson, Nidhi Koundinya, W. Scott Watkins, Lynn B. Jorde, Aaron R. Quinlan, Deborah W. Neklason, Adam M. Phillippy, Evan E. Eichler

## Abstract

The extraordinary repetitive content of human acrocentric short arms has prevented detailed investigations into recombination and *de novo* mutation. Integrating multiple sequencing technologies, we created 156 phased short arms and assessed 107 intergenerational transmissions from 23 samples in a four-generation pedigree. We observed a significant depletion (*P*<0.0001) of p-arm allelic recombination but one ectopic chr13–chr21 recombination breakpoint mediated by a 630 kbp segmental duplication mapping 1.6 Mbp distal to the SST1 array. In contrast, 18 maternal-biased q-arm allelic recombinations are significantly enriched within 5 Mbp of the centromere. Compared to autosomal euchromatin, the overall p-arm *de novo* single-nucleotide variant rate (1.33×10⁻⁷ per base pair per generation) is 10-fold higher, with a significant reduction of C>T but increased C>G and A>C mutations. We hypothesize that acrocentric sequence composition biases and the dearth of allelic recombination contribute to an elevated mutation rate and unique mutational signatures suggestive of mismatch repair defects and oxidative stress-induced DNA lesions.

## INTRODUCTION

Advances in long-read sequencing (LRS) technologies and assembly algorithms have enabled the complete or nearly complete assembly of human genomes^1–6^. The first complete human haploid reference genome, T2T-CHM13, provided one of the first opportunities to characterize genetic and epigenetic variation in previously inaccessible regions of the genome^7–10^. Leveraging LRS technology, both the Human Pangenome Reference Consortium (HPRC) and Human Genome Structural Variation Consortium (HGSVC) have now completed population-level surveys of many complex regions, including centromeres and segmental duplications (SDs)^11–14^. Despite these advances, standard application of LRS and assembly tools still result in typical diploid genome assemblies with 50-100 gaps, and many of the remaining gaps map to the short arm of human acrocentric chromosomes^15^.

The five short arms of human acrocentric chromosomes (SAACs), i.e., chr13, chr14, chr15, chr21, and chr22, harbor multiple layers of repetitive DNA with almost no unique sequence. They contain different types of tandemly repeated sequences, including large tracts of satellite DNA, and greater than 60% of the remaining DNA can be classified as interchromosomal or intrachromosomal SDs^7,10,16,17^. Embedded within this repetitive DNA is a ribosomal DNA (rDNA) cluster on each chromosome that encodes the rRNA scaffolding components of the ribosome. The rDNA clusters interact among nonhomologous chromosomes to form the nucleolar organizer regions (NORs)^7,18,19^. While highly variable between individuals, the five rDNA clusters combined typically encode 200-600 rDNA genes organized in a head-to-tail rDNA configuration that is uniformly transcribed centromerically^7,18^. The basic unit of rDNA repetition is ∼45 kbp in length and consists of a ∼13 kbp “operon” that is processed to produce the 18S, 5.8S, and 28S rRNA, followed by a 32 kbp intergenic spacer region^20^. Each 45 kbp unit is nearly identical at the sequence level leading to these regions being frequently collapsed even if LRS technologies are applied leading to the development of computational tools to estimate copy number^19,21^. The co-localization of NORs from different SAACs brings rDNA and its adjacent proximal and distal sequences in closer proximity when chromosomes synapse and recombine in meiosis. The 3D genomic proximity and highly identical sequence at and flanking the rDNA has been thought to drive ectopic exchanges between nonhomologous SAACs via nonallelic homologous recombination (NAHR), leading to both the formation of natural hybrid short arms in the human population as well as Robertsonian (ROB) translocations^19,22^.

Previous cytogenetic studies have suggested preferential pairing and exchanges, where chr13-chr21 and chr14-chr22 are the most frequently recombined heterologous chromosomes likely due to greater shared SD and satellite content between these particular SAACs^23^. In more recent studies examining incomplete acrocentric DNA contigs from the HPRC, Guarracino and colleagues^19^ identified shared homology domains, termed pseudo-homologous regions (PHRs), centered around a SST1 macrosatellite repeat that showed patterns of positional homology entropy consistent with hotspots of ectopic recombination. LRS and assembly of three ROB patients identified a common breakpoint associated with the SST1 repeat^24^ embedded within the PHR strongly implicating these sequences in ectopic recombination events.

Owing to the complex repetitive content and mutational dynamics of SAACs, most large-scale studies of *de novo* mutation (DNM) and meiotic recombination have excluded these 75 Mbp of human DNA from analysis^25–28^. Indeed, even our own recent long-read-based study of a four-generation, 28-member pedigree, CEPH1463, failed to phase and assemble the SAACs adequately^29^. The goal of this study, then, was to establish a baseline for DNMs and recombination in these regions by the direct comparison of parents and offspring by specifically focusing on assembly of the acrocentric short arm DNA in each generation. Here, we generated additional data, including deep PacBio high-fidelity (HiFi), ultra-long Oxford Nanopore Technologies (UL-ONT), and long-range interaction Hi-C data, to create a high-quality assembly set targeting 23 samples in the pedigree. With nearly complete, fully phased short arms, we examined their sequence composition and variation among individuals in this family. The comparison between offspring and parents enables us to accurately detect DNMs and unambiguously assess patterns of recombination. Compared to other genomic regions, we found unique mutation and recombination signatures on the SAACs, leading to new insights into their genetic instability and properties regarding their inheritance and mutation.

## RESULTS

### Genome sequence and short arm assembly

In addition to the previous HiFi and UL-ONT data^29^, we generated another ∼20× HiFi and R10 UL-ONT data for two G3 samples: NA12879 and NA12886. We produced R10 UL-ONT data (read length > 100 kbp) for 11/12 G4 samples and two G3 spouses (200080 and 200100) (**Figure 1a**). We additionally generated the genome long-range interaction Hi-C data for 23 samples from G2-G4 (**Figure 1a**). The two G1 samples are not included in this study because we only have their cell lines, which might introduce artifacts in variation analysis compared to the primary peripheral lymphocyte DNA. The G4 sample 200105 was unavailable and we were not able to collect additional blood for sequencing. We assembled 23 samples with Verkko2^30^ using an average sequence coverage of 40× HiFi and 30× UL-ONT as well as 30× Hi-C data (**Figure S1a**, **Table S1**). Verkko2 was specifically designed to improve human acrocentric assembly by combining deep LRS with Hi-C data for accurate assemblies of the short arms as well as rare translocations, such as Robertsonians^24,30^. The majority of our assemblies have high contiguity (median AuN 142 Mbp) and the median base quality QV value is 55.37 (**Figure S1a**). For convenience, the G3 sample NA12879 and its offspring are referred to as G4_fam1, and the G3 sample NA12886 and its offspring as G4_fam2.

**Figure 1.**
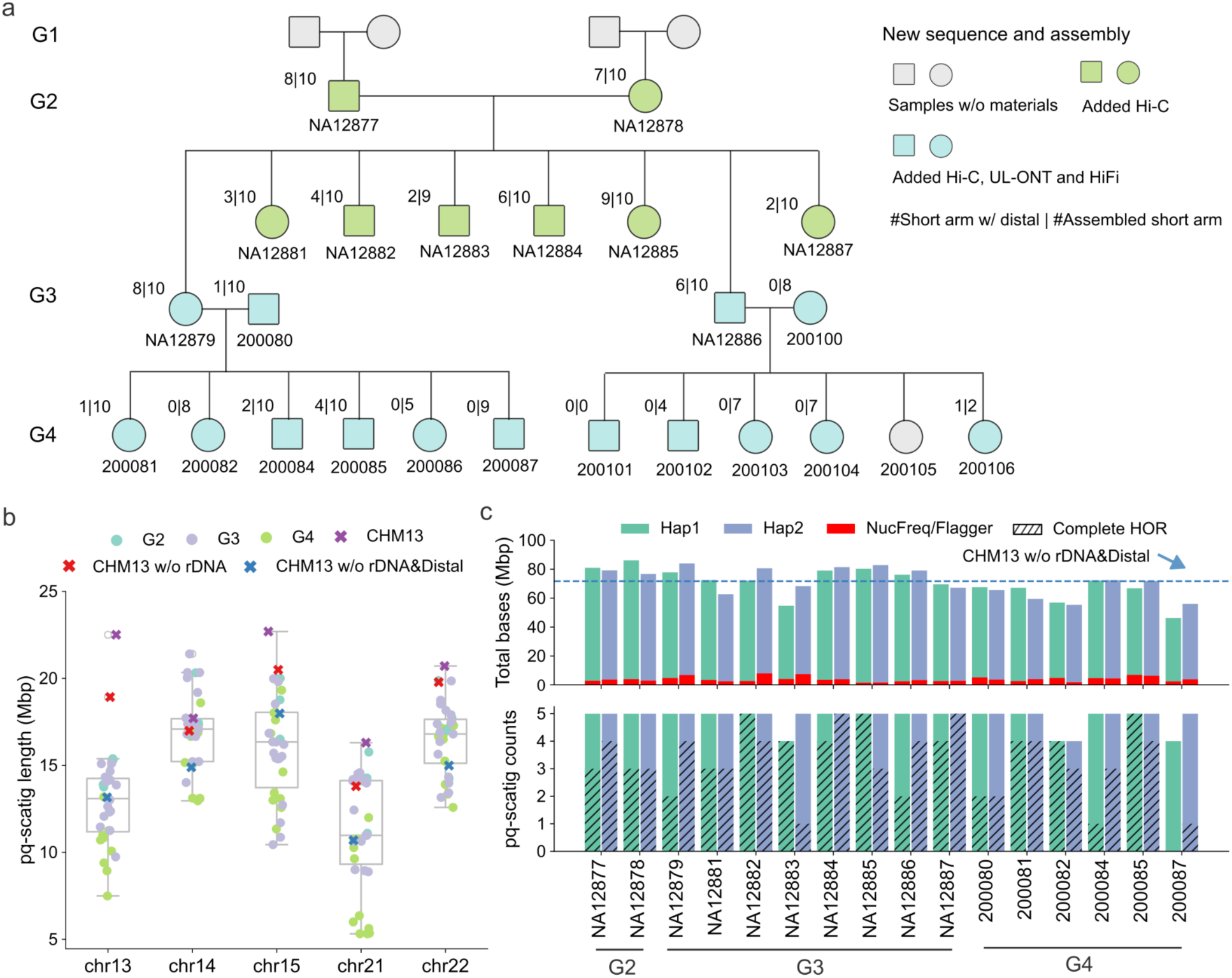
Sequence and assembly of acrocentric short arms from the CEPH1463 family. a) Overview of newly generated sequence data necessary to phase and assemble the acrocentric short arms for individuals in the CEPH1463 pedigree. The fraction for each sample summarizes the number of acrocentric chromosomes (numerator) where distal (band p11) and proximal (band p13) sequences were successfully assembled versus the total number (denominator) of phased assemblies assigned to a chromosome (pq-scatigs). b) Length distribution of obtained pq-scatigs among different chromosomes compared to the expected size based on the T2T-CHM13 reference genome. Each dot represents one pq-scatig. These contigs sum to the length of the p-arm, centromere, and q-arm. T2T-CHM13 (CHM13) sums the length of the short arm, centromere, and 5 Mbp on the q-arm. Based on this length, we then subtract rDNA to get the length of T2T-CHM13 without rDNA (CHM13 w/o rDNA) and subtract both rDNA and distal sequence to get the length of T2T-CHM13 without rDNA and distal sequence (CHM13 w/o rDNA&Distal). c) Summary of pq-scatig counts, total bases, and misassembled bases per haplotype. The misassembled bases are marked by Flagger or NucFreq. The blue dashed line shows the total bases of targeted regions on the T2T-CHM13 five chromosomes, excluding the rDNA and distal sequence.

To identify the short arm assembly for each haplotype, we specifically searched for a single pq-scatig (scaffolded contigs) that spans the p-arm and uniquely aligns to the q-arm of the SAACs based on comparison to the T2T-CHM13 reference genome (**Methods**). The size of targeted regions on T2T-CHM13 are 13.16 Mbp (chr13), 14.89 Mbp (chr14), 17.98 Mbp (chr15), 10.69 Mbp (chr21) and 14.99 Mbp (chr22) excluding rDNA and distal sequence (band p13). We excluded six samples (200086, 200101, 200102, 200103, 200104 and 200106) with less than 30 Mbp sequence per haplotype because these samples were deemed too fragmented for variation discovery or comprehensive recombination analysis on the short arms (**Data S1**). Sample 200100 was also excluded in the analysis because all its offspring assemblies did not pass our QC thresholds.

In total, we generated 156 pq-scatigs (chr13 (n=29), chr14 (n=32), chr15 (n=31), chr21 (n=32) and chr22 (n=32)), representing 97% of the expected haplotypes from 16 samples and consisting of the assembled p-arm, centromere, and q-arm (**Figure 1a, Table S1**). The length of pq-scatigs varies between and within chromosomes, with the median length of 16.9 Mbp for chr14 and 16.5 Mbp for chr22, followed by chr15 (16.1 Mbp), chr13 (13.0 Mbp), and chr21 (11.1 Mbp) (**Figure 1b, Figure S1b-c**). For each sample, across all chromosomes, we obtained an average of 70.9 Mbp of acrocentric sequence per haplotype (around 90% of the size compared to the length of T2T-CHM13 excluding rDNA) (**Figure 1c**). We estimate that 4.8% of the bases are misassembled or collapsed, corresponding to an average of 0.72 Mbp per haplotype per chromosome (**Figure 1c, Methods**). There is no significant difference of misassembled bases between distal sequence and other parts of the pq-scatig (**Figure S1d**).

As expected^19,24^, the large rDNA tandem repeat array (band p12) failed to assemble, representing a predictable collapse region in each acrocentric assembly. Nevertheless, incorporating Hi-C data successfully scaffolded the distal (band p11) and proximal (band p13) sequences in 41% (64/156) of all pq-scatigs (chr13 (n=9), chr14 (n=17), chr15 (n=11), chr21 (n=11) and chr22 (n=16)) (**Table S1**). All partially resolved rDNA tandem arrays as well as other assembled regions defined by NucFreq and Flagger^14^ were annotated and excluded from downstream analyses (**Figure S1e**). With respect to the higher-order repeat (HOR) aSat (*α*-satellite) centromeric DNA, we find that 103 pq-scatigs (12 in G2, 58 in G3, four in 200080 and 29 in G4_fam1) are completely assembled (**Figure 1c, Table S1**). For this family we reported HOR median lengths of 0.6 Mbp for chr13, 1.9 Mbp for chr14, 1.0 Mbp for chr15, 1.1 Mbp for chr21, and 2.4 Mbp for chr22 (**Table S1**). From these complete centromeres, we identified the centromere dip region (CDR), with an average length of 242 kbp, ranging from 50 kbp to 480 kbp (**Table S1**).

### Sequence composition and diversity

Limiting our analyses to 64 short arms (covering five chromosomes from 16 samples) that contained both proximal and distal sequences, we annotated and compared the repeat content of the SAACs. Consistent with the T2T-CHM13 organization, the short arms are largely composed of characteristic blocks of SDs and satellite DNA such as HSat1, HSat3, etc., while their organization and length vary depending on the chromosome (Figure 2a, **Data S1**). For example, chr15 contains significantly longer HSat3 sequence (3.53 Mbp per haplotype, accounting for 68.37% of the short arm) than the other four chromosomes (chr13 (1.08 Mbp), chr14 (1.35 Mbp), chr21 (1.53 Mbp) and chr22 (1.41 Mbp)) (**Figure S2a**). Between chr15 haplotypes, however, HSat3 length variation is considerable, ranging from 1.69 Mbp to 6.78 Mbp (**Figure 2a**). We also noticed a block of diverged HSat3 repeats on chr15 located between the HSat3 and HSat1B array (**Figure S2b**). This satellite block shares ∼93% sequence identity with the HSat3 array but is entirely distinct from the adjacent HSat1B sequence (**Figure S2b**). The methylation levels based on analysis of UL-ONT data from lymphoblastoid cell lines also varies between the chr15 satellite blocks with some haplotypes showing higher methylation at HSat1B (∼75%) compared to HSat3 (∼50%) (**Figure S3a**). Chr13, chr14, and chr21 all harbor the recombinogenic SST1 macrosatellite recently implicated in ROB chromosome formation and a target for hypomethylation in cancer cells^31^. SST1 repeat lengths in this family appear to be bimodally distributed with lengths ranging from 21 kbp to 104 kbp among all 74 correctly assembled SST1 arrays (**Figure S2c**). Even though the SST1 length varies, SST1 CpG methylation levels are consistently high (∼78%) when compared to flanking regions (∼48%) across all three chromosomes (**Figure S3b-d**).

**Figure 2.**
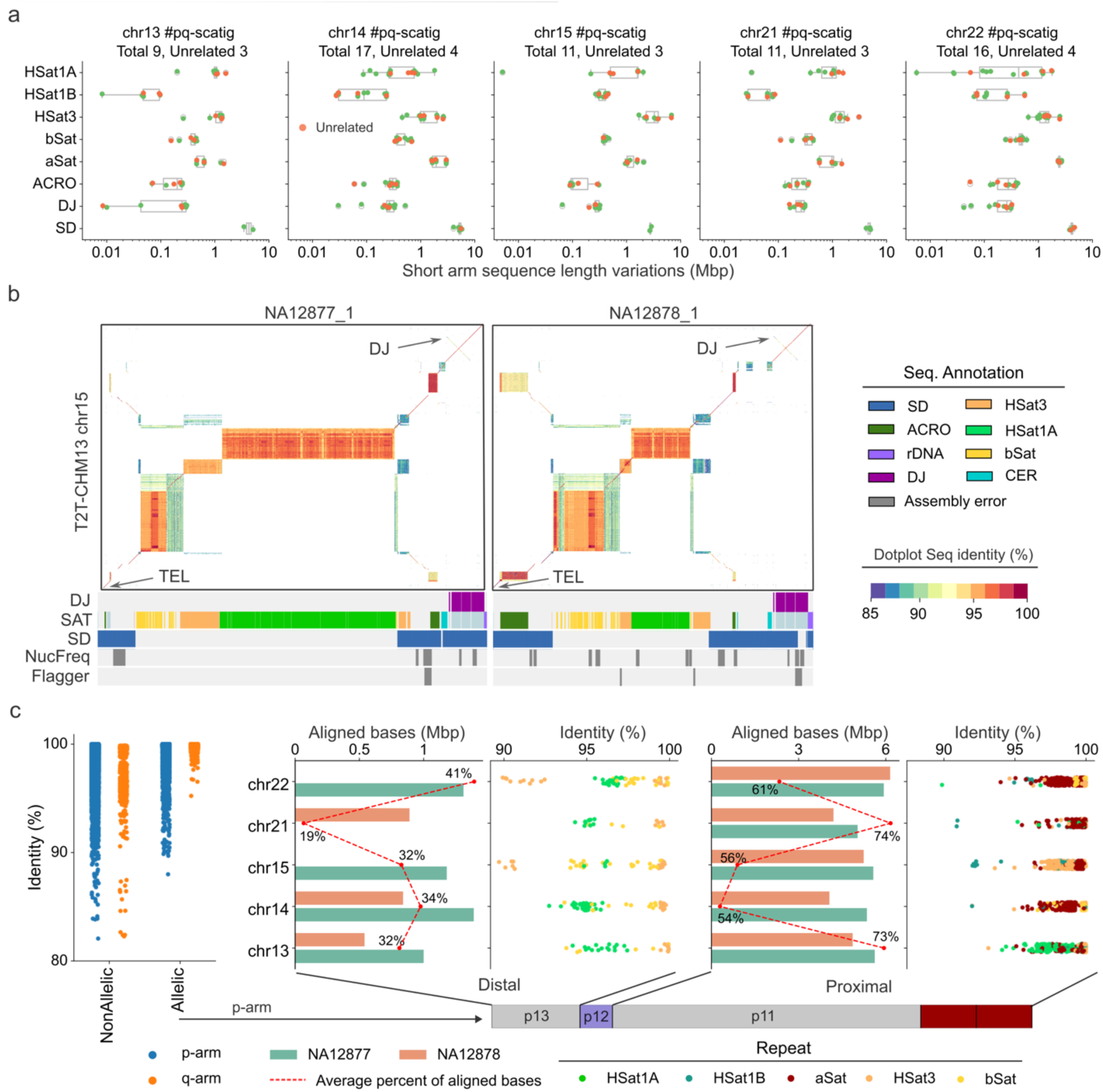
Short arm sequence composition and allelic diversity. a) Length variation of different repeat sequences on the short arm of pq-scatigs with assembled distal and proximal sequences. The orange dots indicate pq-scatigs from three unrelated genomes (G2 NA12877, NA12878 and G3 spouse 200080). b) ModDotPlot^34^ (v0.9.8) comparison of the distal sequence between unrelated samples (G2 NA12877 and NA12878) against the T2T-CHM13 reference genome. The distal junction (DJ) sequence is on the upper right corner. The bottom left corner is the telomere (TEL). The sequence annotations are indicated below each plot with DJ, SD (segmental duplication) and SAT (satellite). c) Summary of sequence identity of 10 kbp binned alignments on p-arm and q-arm between allelic and nonallelic chromosomes of samples NA12877 and NA12878. The percentage of aligned distal and proximal allelic sequence and their sequence identity is further stratified by different satellite repeats. Each dot represents a 10 kbp alignment. The red dashed line is the average of aligned bases on each chromosome among NA12877 and NA12878.

Besides satellite sequences, all SAACs possess one characteristic SD sequence, known as the distal junction (DJ), which forms the distal boundary flanking the rDNA array. DJ is conserved among cell types and shows the chromatin signatures for promoters and active transcription^16,32,33^. The length of the DJ sequence ranges from 297 kbp to 333 kbp among samples in this family (**Figure 2a**). It has been reported that DJ sequences show considerable variation among different SAACs^32,33^. Using a graph-based approach, we constructed a graph from 12 assemblies without errors across DJ sequences and these resolved into 12 distinct “bubbles” with three of these showing nested patterns suggesting extensive structural variation (**Figure S2d**). For example, the chr14 DJ in NA12879 carries a 1,525 bp deletion and three insertions compared to the chr22 DJ sequence in NA12877 (**Figure S2d**). Aside from DJ, we also assessed the variation of the rest of the distal sequence. We note that most of the distal sequence variation is driven by the expansion or contraction of repeat arrays (**Data S1**). For example, the AT-rich 42 bp repeat HSat1A array on chr15 from NA12877 is expanded to a length six times longer than T2T-CHM13, approaching 1.64 Mbp (∼57% of the total distal sequence length) without any assembly errors (**Figure 2b**).

Using the q-arm of each SAAC as an anchor, we compared the extent of allelic versus nonallelic sequence diversity (**Methods**). The alignments revealed a q-arm inversion polymorphism in chr14, chr15, and chr22 within this family (**Figure S4a, Data S1**). Unlike the q-arm, a remarkable feature of the short arms is the high degree of both allelic and nonallelic sequence divergence within the individual genomes (**Figure S4b**). Excluding incorrectly assembled sequences, 65% of the proximal allelic sequences could be aligned and 70% of these aligned bases showed greater than 99% sequence identity (**Figure 2c**). Assessed by different types of human satellite sequence, most of the proximal low-identity allelic alignments (<95%) originated from HSat1B in chr15 and chr21 (**Figure 2c, Figure S4c**). Moreover, the comparison of chr13 and chr21 also revealed both allelic and nonallelic homologous sequences of length ∼3.2 Mbp corresponding to the PHR thought to be important to ectopic recombination (**Figure S4d**). In contrast, only 20-40% of the allelic distal sequence could be aligned, and the aligned portions exhibit even lower allelic identity, with most HSat1 sequences showing only ∼95% identity (**Figure 2c**).

### Short arm transmission of DNA between generations

The completion of short arm assemblies within a family allows for a comprehensive evaluation of their transmission, recombination, and DNM across generations. Using all-vs-all alignments across assemblies from the pedigree and requiring >1 Mbp homology and >99% sequence identity between parents and offspring, we identified a total of 107 transmissions, accounting for 1.1 Gbp and 0.5 Gbp of transmitted base pairs between G2 and G3 and G3 and G4, respectively (**Methods**). This corresponds to an average of 66 Mbp per haplotype, including the p-arm, centromere, and q-arm (**Figure 3a**). Excluding q-arm DNA used to anchor pq-scatigs, we assessed 831 Mbp of short arm transmitted bases, with a median length of 36.8 Mbp bases per haplotype. Across a total of 13 transmissions per haplotype per chromosome (eight in G3 and five in G4_fam1) from G2 to G4_fam1, the most frequently transmitted haplotypes are chr15_h1 (8/13) and chr22_h1 (7/13) from NA12877, and chr13_h2 (6/13), chr14_h2 (9/13), and chr21_h2 (8/13) from NA12878 (**Figure 3b, Data S1**). Using parental haplotypes as a reference, we measured the sequence identity in sliding windows for transmitted SAACs within families. The analysis showed that transmitted segments were virtually identical (99.97%) from one generation to another allowing for the detection of candidate DNMs and recombinant chromosomes (**Figure 3c**).

**Figure 3.**
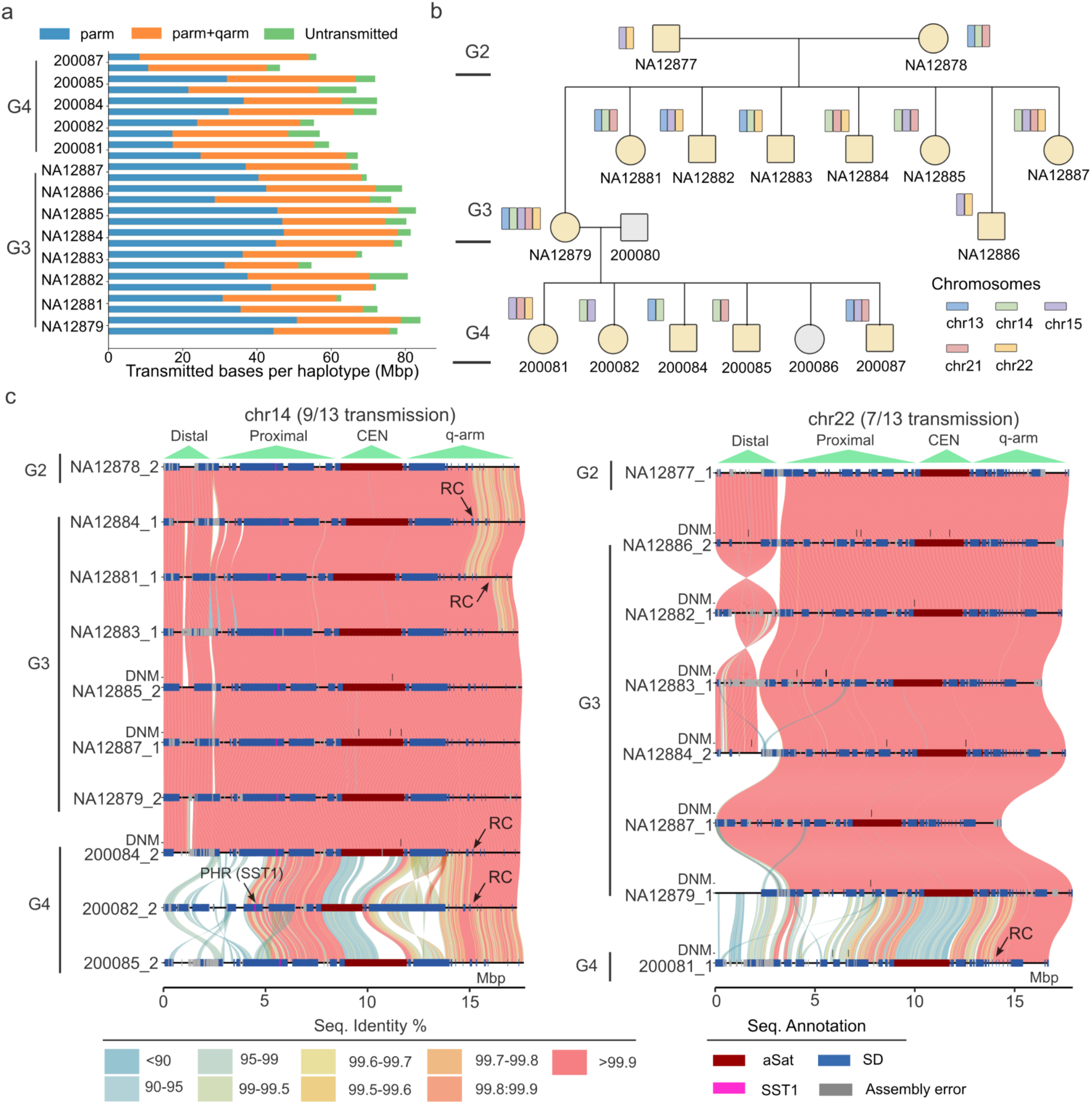
Short arm intergenerational transmissions. a) Summary of the number of transmitted base pairs in this family, stratified by p-arm only (short arm with centromere) and entire pq-scatigs (including 5 Mbp distal to the centromere). b) The most frequently transmitted haplotypes by pedigree sample and chromosome from G2 to G4. c) Stacked SVbyEye^36^ plot depicting the most frequently transmitted chr14 and chr22 haplotypes from G2 to G3 and G4. The heatmap shows the sequence identity (%) based on minimap2 alignment of each contig haplotype to the sequence above. High-identity matches clearly distinguish transmitted (nearly continuous orange pink) from non-transmitted acrocentric (bottom G4). Potential sites of *de novo* mutation (DNM) are indicated by black ticks along with SD annotation (blue bars) and candidate recombinant (RC) chromosomes. Note that the G3 DNMs are detected against the G2 reference and G4 DNMs are detected against the G3 parental haplotype.

These transmissions include a previously reported 515 kbp 14q11 pericentromeric inversion (**Figure S5a**)^29^ and 49 completely sequenced centromeres (32 transmissions from G2 to G3 and 17 from G3 to G4) (**Figure S5b**). The transmitted centromeres allow us to assess intergenerational stability of kinetochore attachment by examining the hypomethylated CDR as a proxy^13^. Overall, the majority (98%) have smaller than 200 kbp CDR shifts in the offspring and 27 out of the 49 transmitted centromeres show CDR shifts smaller than 50 kbp (**Methods**). Further comparisons with unrelated centromeres revealed that the CDR of chr14 (Mann–Whitney–Wilcoxon two-sided test, *P* = 0.026) and chr22 (Mann–Whitney–Wilcoxon two-sided test, *P* = 0.00024) are significantly more stable upon transmission, showing median shifts of 15 kbp and 14 kbp, respectively (**Figure S5c-d**). Chr21 did not show any significant differences between transmitted (average HOR size of 776 kbp) and unrelated haplotypes (average HOR size of 1.07 Mbp) likely due to the smaller and more restricted size of the centromere^13,35^. From four multigenerational transmitted centromeres, we noticed that the position of the CDR could be inherited between G2 and G3 but could also change dramatically when transmitted to G4 (**Data S1**). For example, the chr13 CDR position in G2 and G3 are 3.6 kbp and 8 kbp to the start of the aSat while the position becomes 53 kbp when transmitted to G4 (**Figure S5e**).

### Allelic and ectopic recombination biases

Based on the assembled and transmitted contigs, we mapped all allelic and nonallelic recombination breakpoints within the pq-scatigs containing both short arms and extending 5 Mbp pericentromerically into the q-arm. Recombinant child chromosomes were readily identified based on sequence-identity transitions (**Figure 3c**). By further examining the read alignment pattern and excluding incorrectly phased parental haplotypes, we characterized 19 total recombination events (18 allelic) to a breakpoint resolution of 3 kbp (**Figure 4a, Table S2, Data S1, Methods**). One chr21 recombination event in G3 sample NA12879 was transmitted to samples in G4_fam1 (**Table S2**). Of these recombinations, 74% (14/19) originated from the maternal germline consistent with the maternal bias for meiotic recombination^25,29,37,38^. We mapped the recombination breakpoint to the meiosis-specific histone methyltransferase PRDM9 motif frequently associated with recombination^37^ and found that the distance ranged from 0.6 kbp to 1.1 Mbp (**Data S1**, **Methods**). Not a single allelic recombination was observed on the p-arm, but all 18 allelic meiotic recombination events mapped to the q11.2 regions (**Figure 4a, Figure S6a**). We note that all the three chr15 recombination events on the long arm mapped immediately adjacent to an SD-mediated inversion potentially delimiting a boundary disrupting meiotic synapsis extending centromerically (**Figure 4b**).

**Figure 4.**
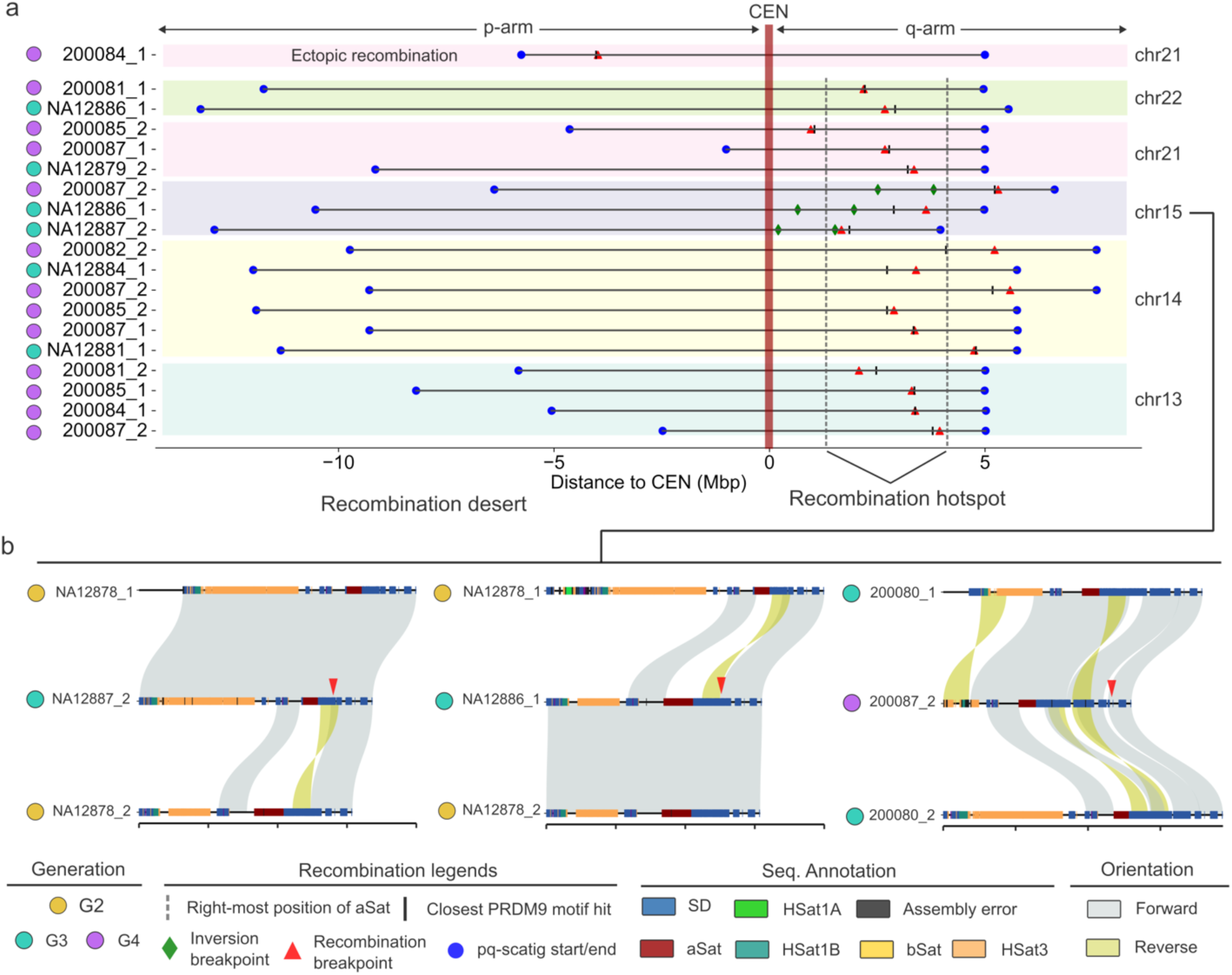
Summary of meiotic recombination events. a) Schematic summarizing the location of 19 meiotic recombination events (breakpoints in red triangles) identified from transmissions for approximately 66 Mbp of DNA per haplotype from chromosomes 13,14,15, 21 and 22. Of the 19 events, 18 clustered on the q-arm (dashed vertical lines) with not a single allelic exchange detected on the p-arm but instead a single ectopic event occurring between chromosomes 13 and 21. Recombination breakpoints are shown with respect to their nearest best PRDM9 motif hit (black tick mark) and the location of pericentromeric inversions (green diamond). b) Three SVbyEye examples of q-arm recombination events (red) are depicted showing parental haplotypes (top and bottom) compared to the recombinant chromosomes (middle) and the location of the pericentromeric inversions at the edges (reversed alignment in yellow).

We tested by simulation whether the pericentromeric q-arm of human acrocentric chromosomes showed a bias for meiotic recombination (**Methods**). Previously, we had constructed a recombination map in this family identifying a total of 532 recombination breakpoints in G3 and 307 in G4_fam1 mapped against T2T-CHM13 euchromatin regions^29^. We note that pericentromeric recombination events were exceedingly rare in our previous analysis of the family. There are, for example, only five q-arm pericentromeric recombinations detected among metacentric chromosomes compared to a total of 18 pericentromeric recombination events in five acrocentric chromosomes. Because this difference might be due to methodological differences for our previous recombination detection approach, we conservatively excluded pericentromeric regions from metacentric chromosomes in the comparison. By randomizing the genome into 5 Mbp segments and controlling for the number of transmissions, we observed a significant 2.3-fold (*P*=0.0045) enrichment of recombination events in the 25 Mbp of acrocentric q-arm pericentromeric DNA assayed when compared to the rest of the euchromatin recombination breakpoints in this family (**Figure S6b**). At a chromosome level, the most significant enrichment was found for chr14 (*P*=0.0014) and chr13 (*P*=0.04). We also applied a similar approach to assess the difference of p-arm recombinations in 38 Mbp (size of T2T-CHM13 proximal sequences) and found significant depletion (*P*<0.0001) of SAAC recombination events (zero compared to an average of 12 events from metacentric chromosome p-arms) (**Figure S6c**).

While no allelic meiotic recombination event was detected on the short arm, we did observe one putative ectopic recombination event between chr13 and chr21. It is predicted to have occurred in the maternal germline (G3-NA12879) resulting in a “hybrid” chr21 and a recombinant chr13 haplotype corresponding to the sample (G4-200084) (**Figure 5a, Figure S7a**). Note that the 200084 chr21 haplotype missed the distal sequence so its alignment on NA12879 chr21 haplotype starts after the rDNA array (**Figure 5a**). To eliminate the possibility of misassembly, we first assessed the transition region for potential collapsed sequence using Flagger and NucFreq, finding no evidence for misassembly (**Figure 5b, Figure S7b**). We further examined the breakpoint by mapping the individual HiFi reads from the child 200084 back to the maternal chr13 and chr21 haplotypes. Consistent with ectopic recombination, the child’s read coverage drops dramatically at the breakpoint on the maternal chr13 reference haplotype with few ambiguous alignments on the q-arm, while we observed strong read coverage signal starting again immediately after the breakpoint for the chr21 reference haplotype (**Figure 5b**). We refined the breakpoint (about 18.7 kbp resolution) to a 630 kbp homologous region that shares 99.53% identity between the maternal chr13 and chr21 haplotypes (**Figure 5c**). Notably, a cluster of high-confidence PRDM9 binding motifs are enriched at the ectopic recombination breakpoint and it is 12 kbp from the best PRDM9 hit. However, this breakpoint is located at 1.6 Mbp proximally from the SST1 array on chr21 where we also found an enrichment of PRDM9 hits (**Figure 5c**). We hypothesize that NAHR between the 630 kbp SD shared between chr13 and chr21 drove the formation of this “hybrid” chr21 p-arm (**Figure 5d, Figure S7a**).

**Figure 5.**
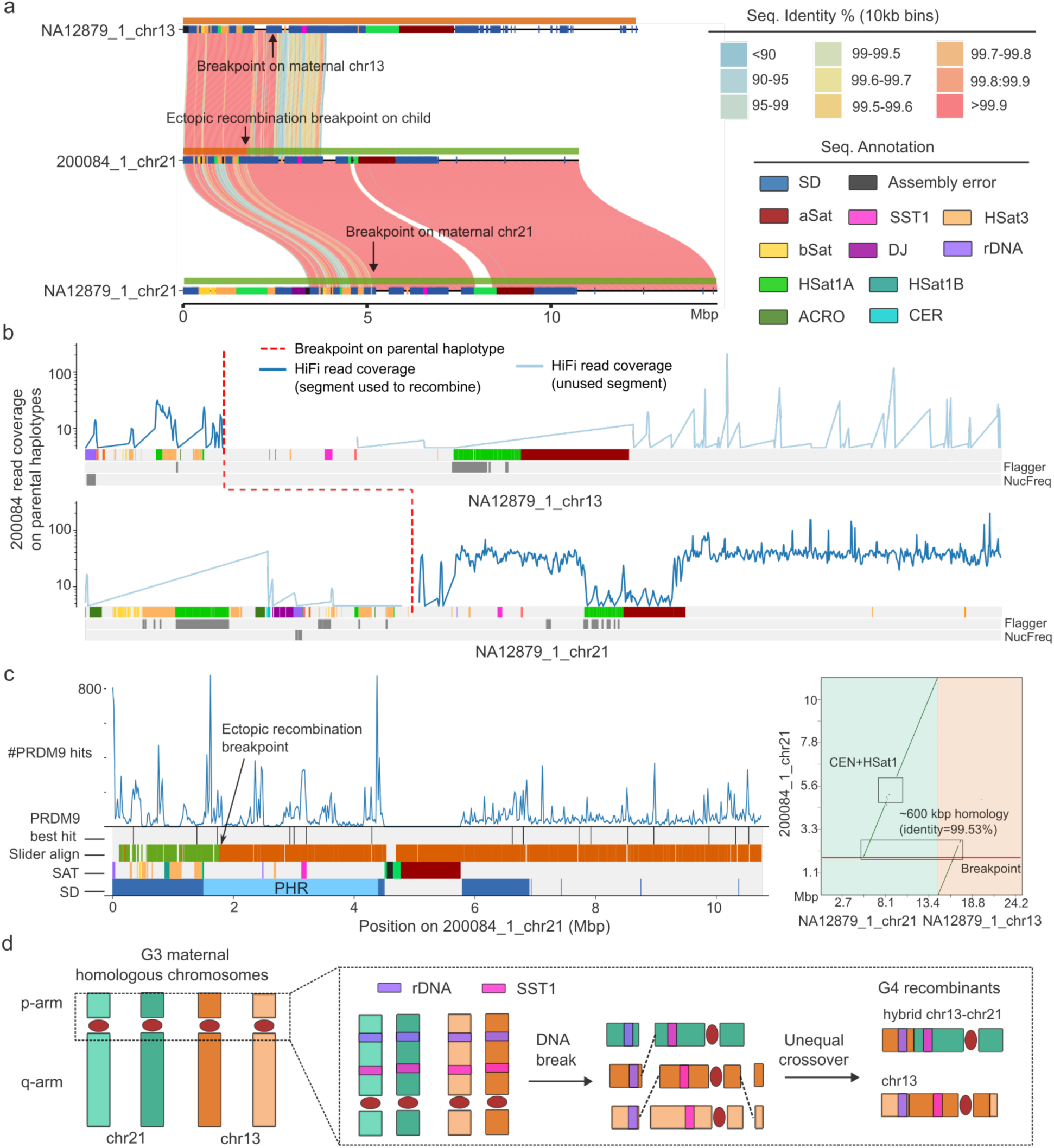
Analysis of an ectopic recombination between chr21 and chr13. a) Chr13-chr21 ectopic recombination shown by SVbyEye based on an all-vs-all analysis of child to parental haplotypes. b) Read coverage plot of a child’s HiFi sequence reads aligned to maternal haplotypes. The dash red line shows the approximate breakpoint on maternal haplotypes identified by 10 kbp sliding window alignments. The y-axis is the read coverage, and the x-axis shows the sequence annotation of maternal haplotypes. c) Annotation of ectopic recombinant chromosome showing (from top to bottom) the distribution of PRDM9 sites, the best PRDM9 hits, perfect alignment of 10 kbp sliding window analysis between maternal and offspring haplotypes, along with SD and satellite annotation, including the location of the SST1 and pseudo-homologous region (PHR). The right panel shows a dot matrix analysis display MashMap alignment between child and maternal sequences. The highly identical homologous SD sequence between maternal chr13 and chr21 likely mediates nonallelic homologous recombination (NAHR). d) Model depicting the chr13-chr21 ectopic recombination by NAHR along with an allelic chr13 recombination on the q-arm in G4 sample 200084.

### De novo mutations

Using the 107 intergenerationally transmitted short arm DNA segments (70 from G2 to G3, 37 from G3 to G4), we next focused on detecting DNMs, including single-nucleotide variants (SNVs) and structural variants (SVs) (**Methods**). Most previous DNM studies excluded these regions because of the extraordinary high-identity repeat content and the challenges associated with mismapping sequence data^29,39^. These limitations were overcome by leveraging the personalized T2T reference sequence (i.e., using parents as reference). We applied a conservative approach to discover DNM, namely all *de novo* variants have to be supported orthogonally by HiFi and UL-ONT in the child but absent from the parental reads (**Methods**). Based on our assessment of an average of 30.7 Mbp (per haplotype) callable sequence on short arm, we identified 103 germline *de novo* SNVs corresponding to 66 SNVs among eight G3 children and 37 SNVs among five G4 children (**Figure 6a, Figure S8, Figure S9, Table S3**).

**Figure 6.**
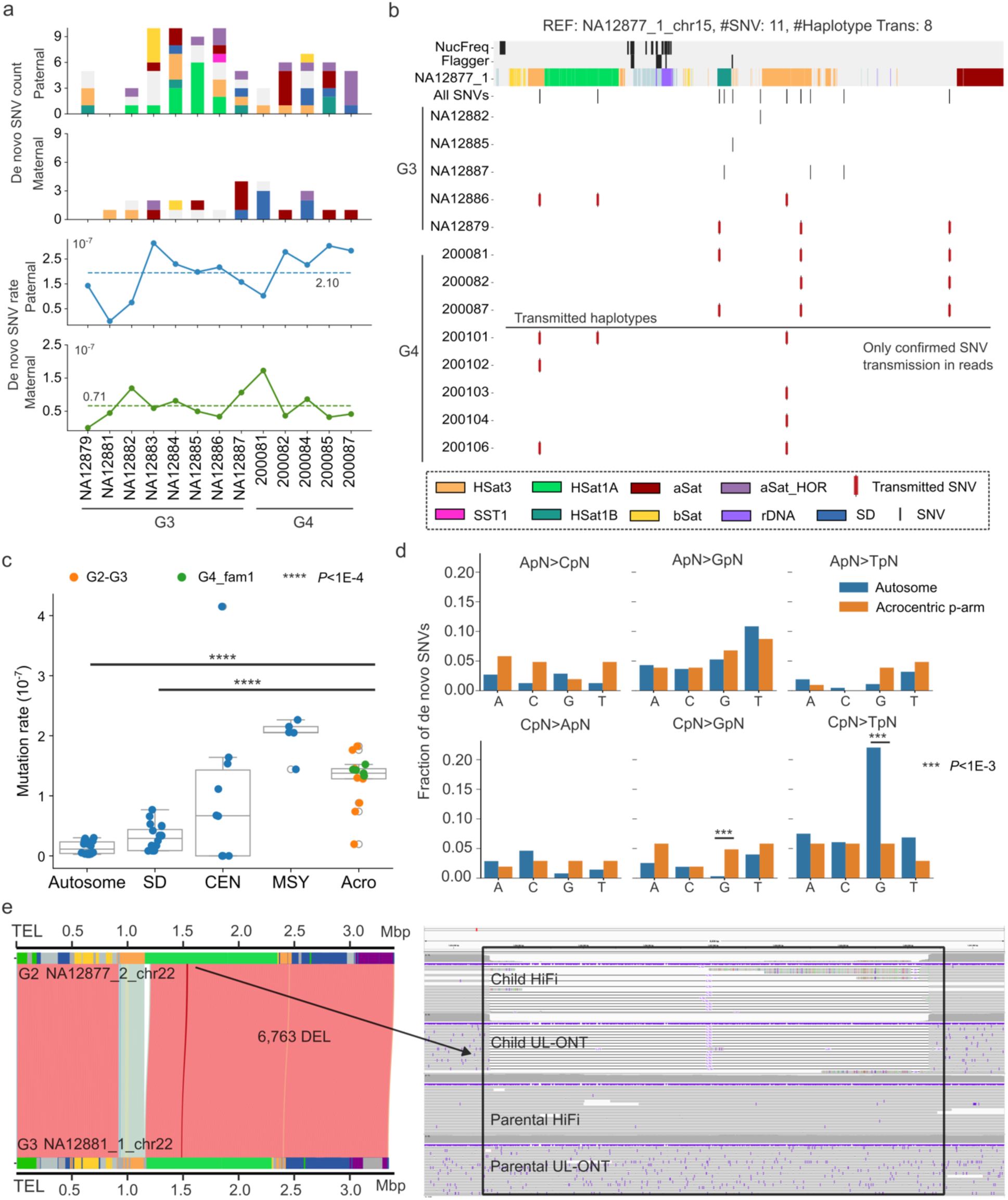
Summary of *de novo* mutations in acrocentric short arms. a) *De novo* single-nucleotide variant (SNV) counts and estimated mutation rate per individual in the pedigree. The two dashed lines show the average paternal (blue) and maternal (green) mutation rates among all children. b) Examples of chr15 *de novo* SNVs (red) arising in G3 and transmitted to G4 based on mapping to the grandfather’s (G2) haplotype as a reference genome. Top tracks indicate the repeat and assembly error annotation of the grandfather, NA12877, reference haplotype from the distal sequence to the centromeric satellite DNA of chr15p. c) Summary of the estimated mutation rate comparing acrocentric short arms (Acro) to autosomal regions (Autosome), segmental duplications (SDs), centromeres (CEN), and male-specific Y-chromosomal region (MSY), including Yq12 heterochromatin from the same pedigree. Note, the centromere here includes higher-order repeat and its flanking monomer regions. Significant differences were calculated based on Mann-Whitney-Wilcoxon two-sided test; \*\*\*\**P*<0.0001. d) Comparison of dinucleotide mutational signatures between autosome chromosomes and acrocentric short arms. A significant difference is determined using two-sided Fisher’s exact test with Benjamini-Hochberg correction; \**P*<0.05, \*\**P*<0.01, \*\*\**P*<0.001, \*\*\*\**P*<0.0001. e) Example of *de novo* deletion mapping within the HSat1A array from the distal sequence. The left SVbyEye plot between parent (NA12877) and child (NA12881) shows the location of this deletion on the distal sequence. The IGV screenshot on the panel shows the alignment of child HiFi/UL-ONT to reference haplotype (NA12877) as well as the parental HiFi/UL-ONT alignment to the reference haplotype. The deletion is only observed in child’s alignments and absent from parents.

While not all regions are completely ascertained or fully resolved at the same level, we observe one to ten SNVs per transmission from all SAACs (**Data S1**). This includes 13 SNVs from distal sequence, which was particularly challenging to assemble without errors. Using transmission to the G4 generation as a form of orthogonal validation, we initially found that 85% (14/16) of the *de novo* SNVs detected in the G3 parental samples, NA12879 and NA12886, were subsequently confirmed as inherited in their G4 children. For example, all six SNVs detected in G3 samples with offspring are inherited to G4, an observation we confirmed using the transmitted G2 father’s chr15 as the reference genome (**Figure 6b**). Subsequent analysis showed that the two *de novo* SNVs identified on chr22 in NA12879 as not transmitted to G4_fam1 were simply the result of an allelic recombination event on the q-arm; as a result, none of the five children in G4_fam1 inherited the segment harboring the two SNVs (**Data S1**).

Considering the 103 detected *de novo* SNVs, 80% mapped to repetitive sequences with only 21 mapping to regions not annotated as an SD or satellite DNA. Among the 70 SNVs mapping to satellite DNA, 13 were assigned to HSat1A, five to HSat1B, 13 to HSat3, six to bSat (*β*-satellite), 18 to monomeric aSat, 14 to HOR aSat, and one to the SST1 repeat (**Figure 6a**).

However, the sample size is too small to claim biological enrichment for the specific satellite sequences, and we note that the HSat1A repeat on chr13 shows as many as four SNVs (one from HSat1A on distal sequence and three from HSat1A adjacent to the centromere) originating from the paternal genome in a single transmission (between NA12877 and NA12885) (**Data S1**).

We assigned the *de novo* SNVs to a parent of origin: 79 paternal and 24 maternal events estimating a paternal and maternal germline mutation rate of 2.10×10^-7^ and 0.71×10^-7^ per base pair per generation, respectively (**Figure 6a**). We observe no significant difference in the mutation rate among different SAACs though the overall mutation rate on the short arm (1.33×10^-7^) is significantly elevated—up to 10-fold higher when compared to autosomal estimates from the same pedigree (1.37×10^-8^, Mann-Whitney-Wilcoxon two-sided test, *P*=1.8×10^-6^) and more comparable to the male-specific Y-chromosomal region (MSY), including the Yq12 heterochromatin region (1.99×10^-7^) where DYZ1/HSat3 and DYZ2/HSat1B repeat arrays predominate (**Figure 6c**). With respect to mutational signatures, we observe a significant depletion of CpG>TpG transitions (two-sided Fisher’s exact test with Benjamini-Hochberg correction, *P*=0.00004) on the short arm when compared to the autosome (**Figure 6d**). In contrast, we observe a significant excess of C>G (two-sided Fisher’s exact test with Benjamini-Hochberg correction, *P*=0.0046) and A>C (two-sided Fisher’s exact test with Benjamini-Hochberg correction, *P*=0.0056) substitutions (**Figure 6d**). Compared to autosomal regions, the A>C mutations on the p-arm are found in regions significantly enriched with AT-rich repeat sequence (Mann-Whitney U two-sided test with Benjamini-Hochberg correction, *P*=0.0083).

In addition to *de novo* SNVs, we also detected eight *de novo* SVs from 13 children in G3 and G4_fam1, including six deletions (DEL) and two insertions (INS) (**Methods**). SV sizes ranged from 135 bp to 6,763 bp (**Table S3**) and all *de novo* SVs mapped to satellite repeats. The three novel chr22 DELs result in stepwise changes of the HOR aSat, i.e., 1,364 bp deletion of one 8-mer in two chr22 transmissions (NA12884 and NA12877, NA12887 and NA12878) and one 680 bp insertion of 4-mer in chr21 (**Figure S10a-c**). Besides the aSat, we identified a 1,407 bp DEL in the chr13 SST1 array of 200081 compared to the father 200080, approximately matching the length of the SST1 consensus unit (**Figure S10d**). There are also two DELs found inside distal sequence, such as a 6,763 *de novo* DEL that deletes the HSat1A sequence on chr22 in NA12881 (**Figure 6e**). Finally, two SVs identified in the G3 individual NA12886—a 680 bp INS affecting the HOR and a 1,794 bp DEL within HSat3—were both confirmed as transmitted to the next G4 generation (**Data S1**).

## DISCUSSION

The acrocentric short arms have long been known to be among the most mutationally dynamic and repeat-rich regions of the human genome^19,20,22,32,33,40^. Consequently, they have been the last regions to be reliably assembled^7^ and are typically excluded from both large-scale and high-resolution maps of human DNM and recombination atlases^25,28,41–43^, including our own recent near-T2T analysis of this family, CEPH1463^29^. Recent algorithmic developments over the last three years focused on resolving this last portion of the human genome by taking advantage of deep LRS data as well as Hi-C data to phase and assemble the majority of short arm acrocentric DNA^7,24,30^. The additional data enabled us to assemble 97% of the acrocentric DNA (not requiring the completeness of rDNA and distal sequence) for 13 transmissions (G2 to G4_fam1). These phased short arms allow us to assess meiotic recombination and discover eight *de novo* SVs and 103 *de novo* SNVs across all five SAACs (none of the *de novo* SVs mapping to the aSat had been previously reported). Based on our analysis of this single family, we make three observations: 1) SAACs are significantly depleted for meiotic crossovers while immediate pericentromeric sequence on the long arm is an apparent hotspot for such recombination events; 2) ectopic recombination is rare—occurring only once in the 13 transmissions we assessed (**Figure 4**); and 3) SAACs mutate ∼10-fold faster than autosomal euchromatin and show distinct mutational signatures (**Figure 6**).

Like other forms of *de novo* SNVs^44–46^, SAAC mutations are predominantly paternal in origin (79 from paternal vs. 24 from maternal germline). In contrast, the eight SVs were equally distributed between the maternal and paternal germlines and mapped exclusively to satellite DNA consistent with mechanisms associated with tandem repeat expansions and contractions of HOR units^29^ (**Figure S10**). While we are still underestimating mutations due to collapsed repeats and unresolved rDNA, our findings suggest an overall high SV mutation rate with four to six SAAC chromosomes structurally changing per parental transmission. Similarly, we estimate that the *de novo* SNV mutation rate is among the highest in the genome comparable to other repeat-rich chromosomal regions including centromeres^29^. The mutation rate on the short arm (1.33×10^-7^), for example, is significantly elevated—up to 10-fold higher than autosomal regions (1.37×10^−8^, Mann-Whitney-Wilcoxon two-sided test, *P*=1.8×10^−6^) but also fourfold higher than that of SD sequences (2.99×10^−8^, Mann-Whitney-Wilcoxon two-sided test, *P*=3.1×10^−6^) based on analysis of individuals from the same family (**Figure 6c**). It is in fact most comparable to the chrY and, in particular, the Yq12 heterochromatin region (1.99×10^−7^), which is enriched for similar satellites and structurally divergent between individuals in the human population^47^.

Notably, SAACs also exhibit unique DNM signatures. For example, we observe one of the most significant depletions in the rate of CpG>TpG transitions (two-sided Fisher’s exact test with Benjamini-Hochberg correction, *P*=0.00004) when compared to the autosomal genome average (**Figure 6d**). Similar depletions have been observed for other repetitive regions of the genome^29,48^ and one potential explanation is that they may result from an excess of double-strand breaks and GC-biased gene conversions^48^. We also observe a significant increase in C>G (two-sided Fisher’s exact test with Benjamini-Hochberg correction, *P*=0.0046) and A>C (two-sided Fisher’s exact test with Benjamini-Hochberg correction, *P*=0.0056) substitutions consistent with signatures of mismatch repair deficiency and oxidative DNA damage–associated transversion, respectively^49–51^. The AT-rich repeat environment on the p-arm might allow for the production of more 8-hydroxyadenine, a byproduct of oxidative damage and, thus, contribute to the higher A>C mutation rate^52^. We note that 14 out of the 103 SAAC *de novo* SNVs had identical matches elsewhere of the same repeat type, including eight in aSat, four in HSat3, one in bSat, and one in SST1. This indicates that these 14 SNVs might originate from interlocus gene conversion events similar to a recent observation for the *DYZ1*/*DYZ2* repeats on the Y chromosome^29^.

A surprising finding was the lack of any evidence of allelic recombination in the p-arm and instead an excess of maternal (75%) recombination events mapping to the q-arm of the SAACs in relative close proximity to the centromere. This apparent recombination desert on the p-arm is consistent with a meiotic recombination study showing that SAACs have fewer MLH1 (crossover-associated protein) foci in human oocytes^53^. We should caution that not all SAAC DNA were fully resolved and that one cannot rule out the possibility of cryptic double recombination events occurring precisely in the p-arm repeat regions that we did not fully assemble (e.g., rDNA and the largest satellite blocks). Nevertheless, given the amount of sequence we surveyed and null distribution constructed from genome-wide crossover events from the same individuals, our results strongly suggest that the p-arm depletion and q-arm excess within 5 Mbp of the centromere is highly unlikely (*P*<0.0001).

One possible model to explain this feature may be that the extreme structural and allelic diversity of homologous p-arms prevents efficient synapsis during the prophase I stage of meiosis^53,54^. Failure of synapsis to occur would serve as a barrier to both standard recombination as well as limit opportunities to repair double-strand breaks and mutated base pairs through sister chromatid and homologous chromosome template repair helping to explain both the elevated mutation rate^55^. Indeed, synapsis of the p-arm without complete synapsis of the q-arm and centromere has rarely been observed for an acrocentric bivalent, which suggests that acrocentric p-arms show limited capability to synapse^56^. This, in turn, could lead to an excess of recombination events immediately adjacent to the centromere (i.e., q-arm excess) or suboptimal pairing between high-identity homologous repeated SDs. A previous study on sperm also showed a noticeable increase in crossover rate on the pericentromeric q-arm of acrocentric chromosomes when compared to other autosomes^43^. With respect to the latter, it is noteworthy that the q-arm of SAACs are known hotspots of NAHR leading to several of the most common genomic disorders, such as 14q11.2^57^, 15q11.2^58^, and 22q11.2 microdeletion syndromes^59,60^. This dearth of allelic recombination might, therefore, indirectly promote NAHR-associated diseases by increasing both allelic and non-allelic recombination events on the long arm. In this study, we found evidence for only a single chr13–chr21 p-arm ectopic recombination event. Contrary to predictions based on a population-level analysis of the HPRC samples^19^ and a more recent study of breakpoints associated with ROB translocations^24^, the breakpoints did not map at or near the SST1 repeat. Instead, our results reveal that the event was most likely the result of an NAHR event involving a massive SD (630 kbp in size) that is shared between maternal chr13 and chr21. Additional multigenerational family studies and high-quality personalized assemblies will be needed to more systematically assess the frequency and properties of SAAC ectopic recombination and to confirm more generally the properties of recombination and DNM we discovered in this family.

## SUPPLEMENTARY INFORMATION

Supplementary Figures: File containing Figures S1-S10.

Table S1: Summaries for sequencing data, assembly quality and pq-scatig annotations.

Table S2: Summary of centromere transmission and recombinations.

Table S3: All detected DNMs.

Table S4: Target sequences used for acrocentric repeat annotation.

Data S1: Figures for all pq-scatig repeat annotation and alignments.

Data S2: All pq-scatig ModDotPlots, NucFreq plots, and centromere validation plots.

Data S3: IGV screenshots for DNMs.

## ACKNOWLEDGEMENTS

We thank Jennifer Gerton, Tamara Potapova, Leo de Lima, and Glennis A. Logsdon for analysis discussions and Tonia Brown for editing and preparation of this manuscript. The following cell lines were obtained from the NIGMS Human Genetic Cell Repository at the Coriell Institute for Medical Research: GM12889, GM12890, GM12891, GM12892, GM12877, GM12878, GM12879, GM12881, GM12882, GM12883, GM12884, GM12885, GM12886 and GM12887. This research was supported, in part, by funding from the National Human Genome Research Institute of the National Institutes of Health (NIH) grants R01HG002385 and R01HG010169 (to E.E.E.) and R35GM118335 (to L.B.J.). Part of this work utilized the computational resources of the NIH HPC Biowulf cluster (https://hpc.nih.gov). This research was supported in part by the Intramural Research Program of the National Institutes of Health (NIH). The contributions of the NIH authors are considered Works of the United States Government. The findings and conclusions presented in this paper are those of the author(s) and do not necessarily reflect the views of the NIH or the U.S. Department of Health and Human Services. E.E.E. is an investigator of the Howard Hughes Medical Institute (HHMI).

This article is subject to HHMI’s Open Access to Publications policy. HHMI lab heads have previously granted a nonexclusive CC BY 4.0 license to the public and a sublicensable license to HHMI in their research articles. Pursuant to those licenses, the author-accepted manuscript of this article can be made freely available under a CC BY 4.0 license immediately upon publication.

## AUTHOR CONTRIBUTIONS

J.L. and E.E.E. conceived the project; D.W.N., L.B.J., and A.R.Q. collected and provided the samples from the pedigree. K.M.M. and K.H., prepared the samples and generated the sequencing data. J.L. and A.R. performed the genome assembly and evaluation. J.L. performed the transmission and recombination analysis. J.L. and F.K.M. did the centromere analysis. J.L. and M.D.N. performed the DNM analysis. J.L. and D.P. performed the inversion analysis. D.Y. annotated the segmental duplications and A.R. annotated the satellite sequences. N.K. organized and uploaded the data. J.L., F.K.M., A.M.P. and E.E.E. drafted the manuscript. All authors read and approved the manuscript.

## DECLARATION OF INTERESTS

E.E.E. is a scientific advisory board (SAB) member of Variant Bio. All other authors declare no competing interests.

## METHODS

### Ethics declarations

Informed consent was obtained from the CEPH/Utah individuals and the University of Utah Institutional Review Board approved the study (University of Utah IRB reference IRB_00065564). This includes consent for open access of research data for 23 members; the remaining five provided informed consent for biobanking with controlled access (see Data availability).

### Genome sequencing

#### Cell lines

We used cell lines collected and created from the previous study^29^. Briefly, cell lines of 10 samples from G2-G3 were obtained from Coriell Institute of Medical Research (CEPH collection). Cell lines for G3 spouses and G4 family members were generated in-house as EBV transformed lymphoblastoid cell lines. These cell lines were then grown up and used for Hi-C and UL-ONT.

#### Long-read sequencing data generation

We used all the data generated in Porubsky, et al.^29^ but produced extra UL-ONT reads for samples in G2 and G3, new UL-ONT for G3 spouses and G4, Hi-C reads for G2, G3, G3 spouses and G4, and PacBio HiFi for G3 parents of G4 and G3 spouses (**Table S1**). Due to lack of a cell line, 200105 from G4 was not included in the data production.

#### PacBio HiFi

DNA used for PacBio HiFi sequencing was extracted from whole blood collected from G3 spouses using the Flexigene system (Qiagen 51206). Additional sequencing libraries were prepared as described in Porubsky, et al., however, sequencing was performed on the PacBio Revio platform on Revio SMRT Cells (PacBio, 102-202-200) using the Revio SPRQ chemistry (PacBio, 102-817-600) with 30 h movies on SMRT Link version 13.3.

#### Ultra-long ONT

Ultra-high molecular weight gDNA was extracted from the lymphoblastoid cell lines using a phenol chloroform protocol^61^. For some samples, DNA was extracted using the NEB Monarch HMW DNA extraction kit for Cells & Blood (#T3050L) following the manufacturer’s protocol with the following exceptions: 6 million cells were used for the starting input with a shaking speed of 600 rpm during the lysis step. DNA was precipitated with 300 uL EEB from ONT.

Libraries were constructed using the Ultra-Long DNA Sequencing Kit V14 (SQK-ULK114). Approximately 40 ug of Phenol Chloroform extracted DNA or two aliquots of NEB Monarch extracted DNA (600uL total) was mixed with FRA enzyme and FDB buffer as described in the protocol and incubated for 10 minutes at RT, followed by a 10-minute heat-inactivation at 75°C. RAP enzyme was mixed with the DNA solution and incubated at RT for 1hr. The final library was eluted in 450 uL EB. 75 uL of library was loaded onto a primed FLO-PRO114M R10.4.1 flow cell for sequencing on the PromethION, with two nuclease washes and reloads after 24 and 48 hours of sequencing.

#### Hi-C data generation

Lymphoblastoid cell lines were used for Hi-C 3D genome mapping. Libraries were prepared using the Dovetail® Micro-C Kit, which uses a micrococcal nuclease (MNase) to produce uniform DNA fragments. The libraries were sequenced on the NovaSeq X with paired-end 150 bp reads up to 30× coverage of valid pairs.

### Genome assembly

Phased genome assemblies were generated using Verkko2^30^ (v2.2.1), which improves repeat resolution and gap closing, and most importantly, introduces proximity-ligation-based haplotype phasing and scaffolding. We used a combination of HiFi, UL-ONT, and Hi-C to create the phased assemblies for 23 samples from G2 (NA12877, NA12878), G3 (NA12879, NA12881, NA12882, NA12883, NA12884, NA12885, NA12886, NA12887), G3 spouses (200080, 200100) and G4 (200081, 200082, 200084, 200085, 200086, 200087, 200101, 200102, 200103, 200104, 200106).

### Assembly evaluation

We applied the same tools and pipelines used by Porubsky, et al.^29^ to evaluate the base quality and the structural accuracy of each phased assembly. We used our in-house evaluation pipeline (https://github.com/EichlerLab/assembly_qc) to calculate the base quality, assembly contiguity, and gene completeness. For base-pair quality, Mery^62^ (v1.0) was used to count the 21-mers from Illumina reads. Merqury^62^ (v1.1) compares the 21-mers against those in the assembled genomes and flags base-pair errors by finding 21-mers uniquely found in the assembly. Compleasm^63^ (v0.2.4, https://github.com/huangnengCSU/compleasm) was used to evaluate the gene completeness in our assembly.

To evaluate structural accuracy, we first aligned sample-specific HiFi reads to their matched phased genome assemblies using minimap2^64^ (v2.28) with parameters ‘-I 10 G -Y -y 100 –eqx - L –cs’. NucFreq (v0.1, https://github.com/mrvollger/NucFreq) and HMM-Flagger (v1.1.0, https://github.com/mobinasri/flagger) were used to identify structural errors based on the read to assembly alignment. NucFreq calculates the nucleotides frequencies to identify collapsed and miss assembly The collapsed assembly were regions with second-highest nucleotide count exceeding five; and misassembly, where all nucleotides were zero. HMM-Flagger uses a hidden Markov model to detect anomalies in the read coverage and classifies the assembly into erroneous, false duplicates, and collapse.

### CpG methylation analysis

To determine the CpG methylation of each pq-scatig, we basecalled raw R10 UL-ONT data with Dorado (v0.7.2, https://github.com/nanoporetech/dorado). We used the methylation previously generated for G2 and G3 samples with R9 UL-ONT data. For each sample, the UL-ONT data was aligned to its own diploid genome assembly using minimap2 (v2.28) with parameters ‘-ax - Y --DM --eqx -y -I 15G’. We then used the Modkit (v0.4.4, https://github.com/nanoporetech/modkit) pileup option with parameters ‘--preset traditional’ to convert modBAM to bedMethyl files. For each haplotype, the methylation value is further averaged in every 10 kbp bins obtained from BEDTools option ‘makewindows -w 10000’.

### Centromere analysis

Centromere location, size, and repeat composition were identified with CenMAP (v0.3.1, https://github.com/logsdon-lab/CenMAP). The workflow first determines the assembly contigs containing the centromeres and uses alignment of HiFi reads to the sample assembly to identify possible sequence errors (e.g., collapses, misjoints, etc.) with NucFlag (https://github.com/logsdon-lab/NucFlag). Also, CenMAP identifies and plots the centromeric repeat composition and higher-order repeat (HOR) organization with RepeatMasker and HumAS-SD (https://github.com/fedorrik/HumAS-HMMER_for_AnVIL), respectively. Centromeric methylation was investigated aligning the ONT sequencing reads containing methylation tags to their respective sample’s genome assembly using minimap2 (v2.28). Centromere dip regions (CDRs) were identified and visualized using CDR-Finder (v1.0.0, https://github.com/EichlerLab/CDR-Finder) with default parameters. The CDR position is estimated as the difference between CDR start and the start position of the aSat (*α*-satellite). For different samples, the CDR shift is calculated as the difference between the CDR positions of two haplotypes.

### Short arm sequence annotation and quality control

#### Identification of short arm pq-scatigs

We applied an approach similar to Guarracino, et al.^19^ for acrocentric ‘pq-scatig’ identification. The assemblies were first aligned to the T2T-CHM13 reference genome with minimap2 (v2.28) parameters ‘-x asm20 --secondary=no -s 25000 -K 8G -c --eqx --cs’. For a single contig, alignments shorter than 100 kbp or exhibiting <90% sequence identity were excluded from further analysis. Using q-arm as an anchor, we additionally required the q-arm alignment should be greater than 1 Mbp and 1 Mbp away from centromere to avoid the alignment ambiguity caused by SDs. Finally, we obtain the ‘pq-scatig’ for each chromosome that covers the p-arm, centromere, and 5 Mbp on the q-arm side.

#### Segmental duplications (SDs)

Repetitive elements within the genome assemblies were masked using a combination of three tools. Tandem Repeats Finder^65^ (v4.1.0) is applied with the command “trf {asm.fa} 2 7 7 80 10 50 2000 -l 30 -h -ngs”. RepeatMasker^66^ (v4.1.5) is run with the options “-s -e ncbi -xsmall - species human {asm.fa}”. In addition, WindowMasker^67^ (v2.2.22) is executed in two stages: first to generate counts “-mk_counts -mem 16384 -smem 2048 -infmt fasta -sformat obinary -in {asm.fa} -out {asm.count}”, followed by the masking step “-infmt fasta -ustat {asm.count} -dust T -outfmt interval -in {asm.fa} -out {asm.interval}”. The resulting repeat annotations from all three tools were merged, and the corresponding BED intervals are used to softmask the assemblies. SDs are subsequently detected using SEDEF^68^ (v1.1) on these repeat-softmasked sequences. Only SDs with >90% sequence identity, lengths exceeding 1 kbp, and <70% satellite DNA content are retained for downstream analyses.

#### Satellite and other repeats

The short arms of the acrocentric chromosomes are enriched for satellite repeats with certain patterns characteristic of certain chromosomes and/or haplotypes^7^. To obtain an overview of the sequence content and confirm sequences belonging to the acrocentric short arms, particularly the distal and proximal sequences from the rDNA, satellite sequences in the assemblies were annotated based on a set of targeted sequence (**Table S4**).

In brief, the target repeat sequences were selected based on the CenSatv2.0 annotation available for T2T-CHM13v2.0 (https://s3-us-west-2.amazonaws.com/human-pangenomics/T2T/CHM13/assemblies/annotation/chm13v2.0_censat_v2.1.bed)^69^. Most were selected from chr13 or chrY, except for HSat1A, HSat3_A5, and HSat3_B3. For these three HSat repeats, the original consensus generated for classifying the satellite array subgroups were used as described by Altemose^70^. The consensus sequences are available as fasta files on https://github.com/altemose/HSatReview/tree/main. The rDNA reference KY962518 was rotated to begin upstream of the 45S TSS, to include the 45S promoter region.

The target sequences were found using minimap2 (v2.28) in each assembly. Because most mappers are designed to find one best match for each query in the reference, using the assembly as the reference would yield only one result for each mapped target satellite. Thus, the reference and query sequences were flipped, mapping the assembly (query) to the targeted repeat sequence (reference) instead. This approach allows a more sensitive search of the target sequence in the assembly. Once all the alignments are collected in a PAF file, the reference and query coordinates were inverted with rustybam (v0.1.33, https://mrvollger.github.io/rustybam) and converted to BED format. Alignment blocks within 500 bp were merged with BEDTools (v2.31.1)^71^ and formatted with designated colors and filtered to be over 2 kbp or 7–8 kbp to exclude excessive alignment matches to LINE elements.

Below are the command lines used to identify the target sequences:

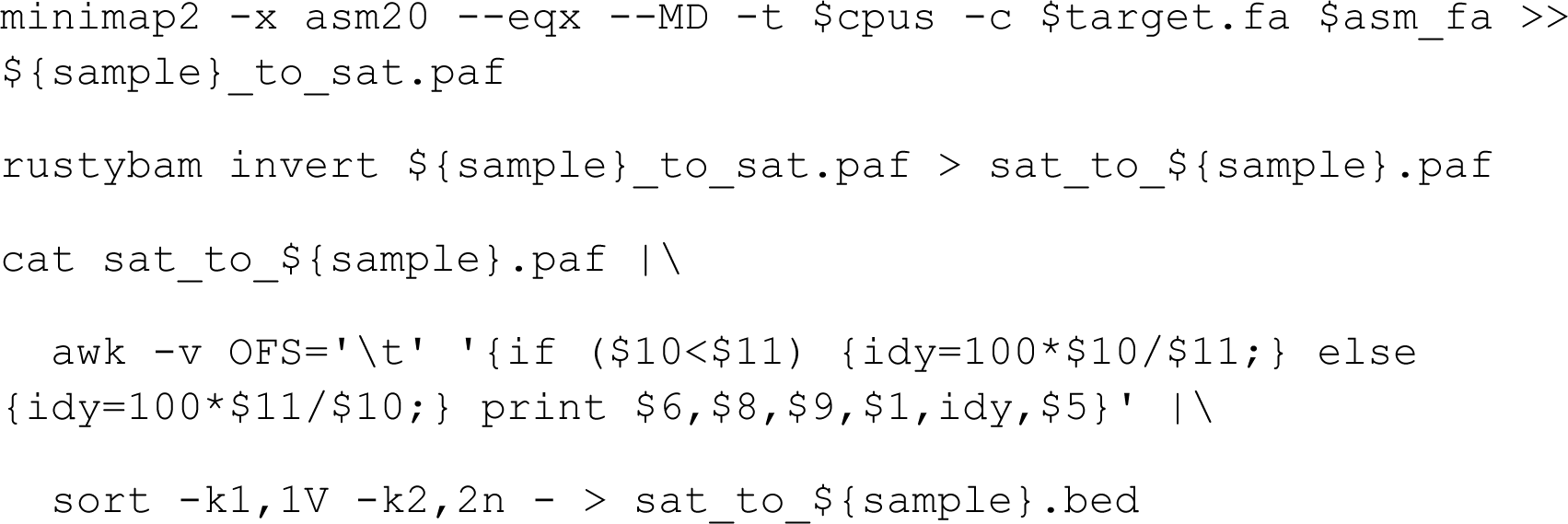

For each target satellite, merging and filtering was applied along with the color assignment. Telomere sequences and gaps were annotated with Seqtk (v1.4, https://github.com/lh3/seqtk) using tel -d5000 and gap -l 1 parameters.

Below are the command lines used to merge and filter satellite annotations:

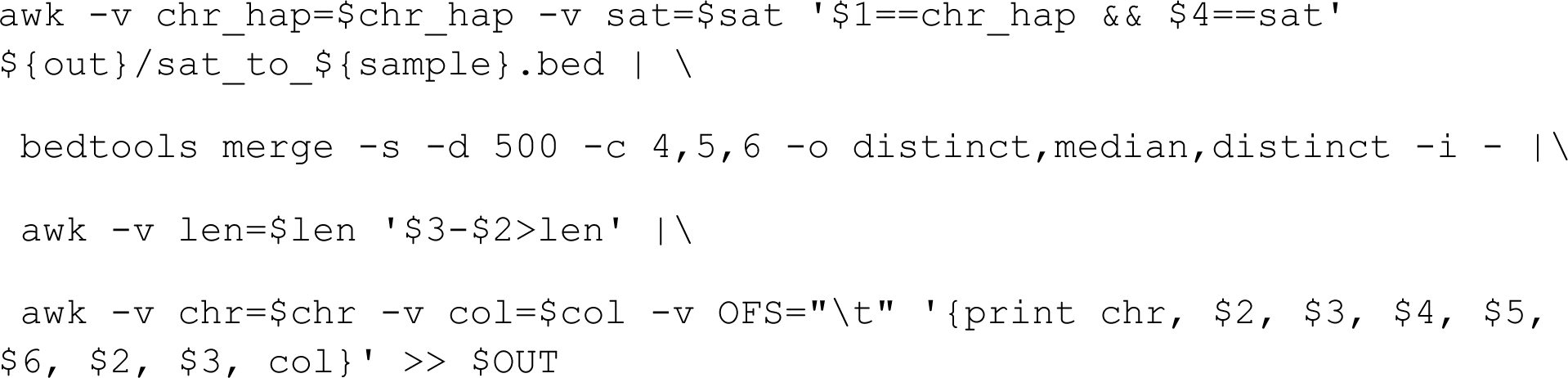

The final BED file was then concatenated and sorted for manual inspection and visualization. The final short arm satellite annotation is available on (https://github.com/Platinum-Pedigree-Consortium/AcroMutRecomb), with the full source code available under https://github.com/arangrhie/Scratch/blob/master/Acro/src/annotate.sh.

#### Evaluation of short arm structure

For each pq-scatigs, we created two plots to help us assess the structural completeness, i.e., determining whether the pq-scatigs contain distal sequence. The sequence dotplot of each ‘pq-scatig’ is created with ModDotPlot^34^ (v0.9.8). The NucFreq plot along with Flagger and satellites sequence annotations, especially for ACRO, distal junction (DJ), and proximal junction (PJ) (**Data S2**). The distal sequence is required to contain the subtelomeric ACRO repeat and the distal junction flank rDNA array.

### Short arm sequence variation analysis

#### Distal junction sequence variation

The distal junction (DJ) sequences without any assembly errors are extracted based on the annotation. The DJ sequence graph is further created with minigraph^72^ (v0.21) options “-cxggs - t16 {sample1}_DJ.fa {sample2}_DJ.fa {sample3}_DJ.fa”. The DJ graph is visualized with Bandage^73^ (v0.8.1). We used the chr22 DJ sequence from G2 sample NA12877 as the reference to show the variations in the DJ graph (**Figure S2d**). The path for each DJ sequence is identified with “-cxasm --call -t16 graph.gfa {sample1}_DJ.fa”.

#### Short arm variation and allelic diversity

To assess the short arm variation, we created the all-vs-all alignment for each chromosome, which included the p-arm, centromere, and q-arm via minimap2 (v2.28) with parameters “-x asm20 -c –eqx -D -P –dual=no {input.multi.fasta} {input.multi.fasta}”. An SVbyEye plot was created to show the variations of single chromosomes within this family (**Figure S2a**).

For the two unrelated individuals (NA12878 and NA12877), we used the same minimap2 settings to create the all-vs-all alignment within individual genomes to assess allelic and nonallelic variation. Alignments located inside incorrectly assembled regions were excluded in the downstream analysis. We used a 10 kbp sliding window aligner with rustybam (v0.1.33) command “liftover –bed {regions.bed} {aln.paf}” and then calculated the sequence identity with “stats –paf {10kb_slider.paf}”. Each 10 kbp alignment was assigned to specific repeat classes based on the SD and satellite annotations. The alignment was assigned as a unique region (SEQ) if outside of the SD and satellite sequence. To avoid overcounting of the bases from overlapping alignments, we merged the 10 kbp slider alignment into alignment blocks with BEDTools (v2.30.0) “merge -c -o collapse”. The option “collapse” keeps the information of all aligned 10 kbp segments inside each block. The average aligned sequence identity was calculated using all alignments inside each block. Alignments were grouped as allelic if the reference and query sequences corresponded to allelic chromosomes within individual genomes; all others were classified as nonallelic.

### Transmission and recombination analysis

#### Identification of parent to children transmitted bases

This four-generation family contains 13 transmissions: eight from G2 to G3 and five from NA12879 and 200080 to their offspring. To identify transmitted bases, wfmash (v0.13.0) was used to align all offspring’s ‘pq-scatig’ to their parents with parameters ‘-s 50k -l 150k -p 90 -n 1 -H 0.001’. For example, all sequences from G3 are aligned to G2 sequences to find the best homologous between offspring and parents. We used segment seed length 50 kbp (-s) and kept one best mapping (-n), requiring homologous regions at least 150 kbp (-l) long and estimated k-mer identity of 90%. Due to the high sequence similarity, we further kept alignment blocks longer than 1 Mbp with sequence identity greater than 99% to avoid counting transmitted bases from misalignments. The transmitted bases were further validated with minimap2 (v2.28) all-vs-all alignment of parameters “-x asm20 -c --eqx -D -P --dual=no” and manually inspected with SVbyEye (https://github.com/daewoooo/SVbyEye).

#### Recombination detection and validation

To identify meiotic recombination, we specifically looked for offspring’s single contig split aligned to its paternal or maternal contigs. We used the same alignment created in identifying transmitted bases. The homologous recombination can be identified if the offspring’s contig is aligned to the maternal or paternal homolog chromosomes. Two steps are further applied to validate each recombination. Firstly, we created all-vs-all alignments for potential recombination events with minimap2 (v2.28) parameters “-x asm20 -c --eqx -D -P --dual=no” and visualized with SVbyEye. Secondly, we aligned the offspring’s HiFi reads to the parental haplotypes that were involved in the recombination and examined the read coverage pattern on parental haplotypes. The coverage was calculated on each parental haplotype by deepTools2 (v3.5.5)^74^ with options “bamCoverage –minMappingQuality 30 –binSize 10000”. We required mapping quality greater than 30 to avoid ambiguous alignments between parental haplotypes. The recombination is correct if the read coverage pattern matches the SVbyEye all-vs-all alignment view. There were two major causes of the false discoveries: 1) the contigs aligned to both maternal and paternal haplotypes due to haplotype switch and 2) the split aligned child contigs to incorrectly scaffolded distal sequence (**Data S1**).

#### Recombination breakpoint refinement

We first used the 10 kbp sliding aligner to locate the approximate recombination breakpoints. This was achieved by rustybam (v0.1.33) with “rb liftover –bed <(bedtools makewindows -w 10000 <(printf “$contig\t0\t$size\n”) {input.paf} | rb stats -q –paf > {output.aligned.stats}” (details at https://github.com/Platinum-Pedigree-Consortium/AcroMutRecomb). The input PAF file is the all-vs-all alignment created for each recombination. By examining the exact 10 kbp segment matches in the output file, the approximate breakpoint on the child’s haplotype is the transition position of alignments against parental haplotypes. To further refine the breakpoint, we used the paralog-specific variants^59,75^ of parental haplotypes. The parental paralog sequences were first obtained by identifying segments on the child’s haplotype that aligned to both parental haplotypes. Specifically, we aligned the child’s sequence to parental sequence with MashMap (v3.1.3) parameter “-s 10000” to obtain the child’s segment that aligned to both parental haplotypes near the approximate breakpoint. This helps us identify the homology sequence between parental haplotypes that potentially mediate the recombination. To identify paralog-specific variants, child and parental sequences were extracted from a 30 kbp flanking region at the approximate breakpoint. Using these sequences, we then created the multiple sequence alignment (MSA) with (v7.525)^76^ default parameters. The breakpoint was refined into an interval where the boundaries correspond to the two nearest paralog-specific variants in the MSA.

#### Test the significance of recombinations

The genome-wide recombination map of this family against T2T-CHM13 is available at https://static-content.springer.com/esm/art%3A10.1038%2Fs41586-025-08922-2/MediaObjects/41586_2025_8922_MOESM10_ESM.xlsx. From this table, we obtained the G3 recombination breakpoint from tab “G3_refined” and G4 recombinations from the “G4” tab.

To assess the significance of the q-arm recombination, we created the null distribution by randomly shuffling five 5 Mbp regions across the T2T-CHM13 genome 5,000 times. This was completed with the BEDTools (v2.30.0) command “bedtools shuffle -excl {excluded.bed} -i {input.bed} -g {t2t_chm13.txt}”. For option “-excl”, it includes the SAAC regions and the 5 Mbp intervals flanking the centromere on both p-arm and q-arm sides. The “{input.bed}” file contains the coordinates of five 5 Mbp pericentromeric regions on the acrocentric chromosomes in T2T-CHM13. We count the number of recombinations in the randomly selected 25 Mbp regions with BEDTools via command “bedtools intersect -c -a {regions}.bed -b {recombination}.bed”. The option “-a” includes the randomly shuffled regions. The last column in the output keeps the number of recombinations used to create the null distribution. The empirical p-value is calculated for the observed 18 q-arm recombinations toward the null distribution. We repeated the above steps for the p-arm significance test but changed the shuffled region size to 36 Mbp (i.e., the size of five SAAC excluding rDNA on T2T-CHM13). Moreover, we only shuffled the regions on the p-arm of metacentric chromosomes.

#### PRDM9 motif and breakpoint association

The PRDM9 motifs are detected with the function FIMO in MEME suite^77^ (v5.5.5). The details can be found in https://github.com/AndreaGuarracino/readcombination/tree/master/prdm9_binding_motifs. We kept the hits of the first 14 PRDM9 motif^78^ and counted the number of occurrences present in 20 kb windows along the recombinant with BEDTools command “bedtools intersect -c -a 20kb_window.bed -b fimo_output.bed > fimo_output_20kb_cnt.bed”. The output is used to create the figure for the number of PRDM9 motif hits. The best hits are obtained from narrow peaks created by FIMO and we measured the distance from the refined recombination breakpoint to its nearest best motif.

### *De novo* mutations

#### Do novo variants calling and validation

We directly detected *de novo* mutations (DNMs) from offspring haplotype to parental haplotype alignment created by wfmash in the transmission analysis section. We used wgatools^79^ (v1.0.0, https://github.com/wjwei-handsome/wgatools) with parameters ‘call -f paf -s -r’ to detect single-nucleotide variant (SNV) and structural variant (SV) from parent-to-offspring transmitted sequences, including SNVs, insertions, deletions, and inversions. All mutations found on the child haplotype are potentially DNMs and are further validated to identify true DNMs.

The candidate DNMs were detected from the haploid-to-haploid ‘pq-scatig’ alignments. To avoid incorrect calls arising from assembly errors, we developed a pipeline to distinguish false discoveries from true DNMs based on HiFi and UL-ONT reads. First of all, all DNMs were required to be outside of the assembly error regions on both the reference sequence (parental haplotype) and the query sequence (child haplotype). For a pq-scatig alignment between parent and offspring (i.e., alignment from a transmission), we align the offspring’s HiFi and UL-ONT reads to its matched parental haplotype and also align each parent’s HiFi and UL-ONT reads to its own haplotype. HiFi reads are aligned with minimap2 (v2.28) parameters “-a -x map-hifi --eqx”. UL-ONT reads are aligned with minimap2 (v2.28) parameters “-a -x map-ont --eqx”. Using these read alignments, a valid *de novo* SNV should be seen in at least three offspring HiFi and UL-ONT reads yet absent from both parental HiFi and UL-ONT reads.

For *de novo* SVs, we run Delly^80^ (v1.3.3) and Sniffles2^81^ (v2.6.1) on both HiFi and UL-ONT read alignments with default parameters using the same alignments described above. We then use the Truvari^82^ (v5.2.0) collapse option with parameters “--pctseq 0.8 --pctsize 0.8 --refdist 1000 --sizemin 50 --sizemax 100000 --gt het -k maxqual --intra” to examine if SVs detected from the contig were also detected by HiFi and UL-ONT reads. Briefly, an SV detected from the contig and read is the same if they share at least 80% allele size and sequence identity similarity, and their breakpoint should be within 1 kbp. We then decode the SV support information from the “SUPP” column in the Truvari integrated output to identity candidate *de novo* SVs. A true *de novo* SV should only be detected by Delly or Sniffles in the child’s HiFi or UL-ONT reads but absent from parents’ HiFi and UL-ONT reads.

We finally viewed the IGV screenshots of each candidate *de novo* SV and SNV to confirm the true DNMs and only report those that pass manual inspections (**Data S3**).

#### Mutation rate calculation

To calculate the *de novo* SNV rate, we first identified callable regions from those transmitted bases without assembly or alignment error. As described in the previous section, transmitted bases are defined as alignments spanning at least 1 Mbp with greater than 99% sequence identity. We then identified misalignment and incorrectly assembled regions from transmitted bases as follows. Because some pq-scatigs miss distal sequences and similar repeat arrays occur on both distal and proximal sides, we applied a 10 kbp sliding aligner to examine misalignments between distal and proximal regions (**Data S1**). Alignment errors were identified as one or multiple 10 kbp windows that have smaller than 99.9% sequence identity between parents and offspring. The sliding window aligner was completed with BEDTools (v2.30.0) and rustybam (v0.1.33). For assembly errors, we used the union of NucFreq and Flagger masked regions as incorrect assembly since NucFreq and Flagger use different methodologies. The unioned regions were obtained via a BEDTools merge option for each pq-scatig. We also excluded regions on the q-arm side that starts from the right-most aSat position. The *de novo* SNV mutation rate for every haplotype was estimated as the number of *de novo* SNVs divided by the total callable bases.

## DATA AVAILABILITY

All underlying data from 28 members of the family are available as part of the AWS Open Data program, European Nucleotide Archive (ENA) or dbGaP. Newly generated sequencing data and assemblies for 19 family members (G2-NA12877, G2-NA12878, G3-NA12879, G3-NA12881, G3-NA12882, G3-NA12885, G3-NA12886, G3-200080, G4-200081, G4-200082, G4-200084, G4-200085, G4-200086, G4-200087, G3-200100, G4-200101, G4-200102, G4-200104 and G4-200106) who provided consent for their data to be publicly accessible for development of new technologies, study of human variation, research on the biology of DNA and study of health and disease are available via the AWS Open Data program (s3://platinum-pedigree-data/) as well as the European Nucleotide Archive (BioProject: PRJEB86317). Sequencing data and assemblies for four family members (G3-NA12883, G3-NA12884, G3-NA12887 and G4-200103) who did not consent for open access are available at dbGaP (phs003793; Platinum Pedigree Consortium LRS).

## CODE AVAILABILITY

Custom code and pipelines used in this study are publicly available at GitHub (https://github.com/Platinum-Pedigree-Consortium/AcroMutRecomb).

## Supplementary Figures for

**Figure S1.**
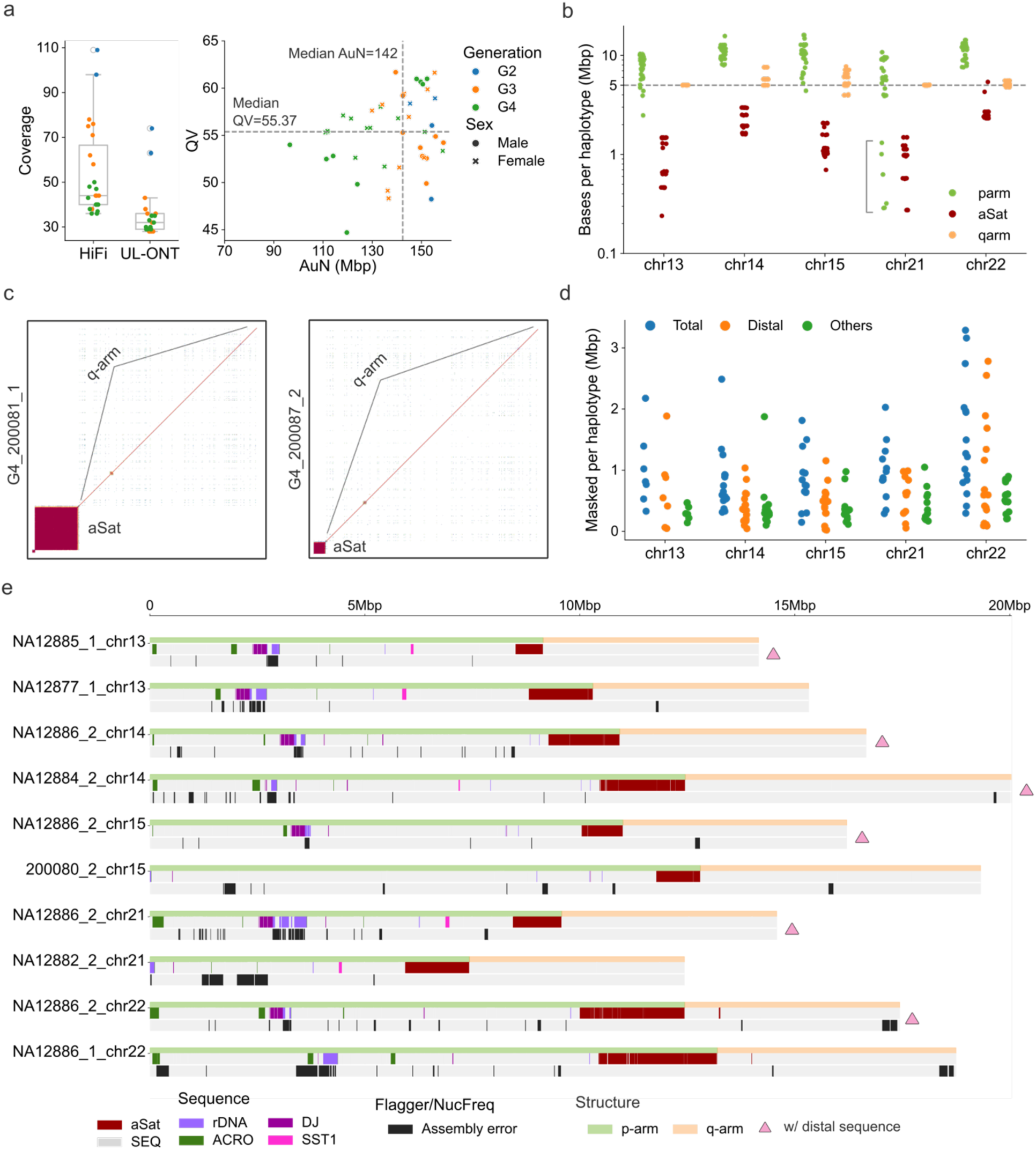
Short arm assembly evaluation. a) Data coverage and the assembly quality (base quality QV and contiguity AuN). The QV is calculated with Merqury (v1.1) based on Illumina short-read data. b) Number of bases per haplotype for p-arm, aSat (*α*-satellite), and q-arm sequences. The six pq-scatigs with shorter p-arms are indicated by the bracket. c) Two examples of the six short p-arms that only contain a limited proportion of pericentromeric sequence on the short arm side. d) NucFreq and Flagger identified incorrect bases per haplotype for all sequences (Total), distal sequence (Distal), and sequence from non-distal regions (Others). e) Examples of structure diagrams comparing pq-scatigs with distal sequence and without distal sequence. The distal sequence is completed if it contains the subtelomeric ACRO repeat (dark green) and the distal junction (DJ; dark purple) flank rDNA array.

**Figure S2.**
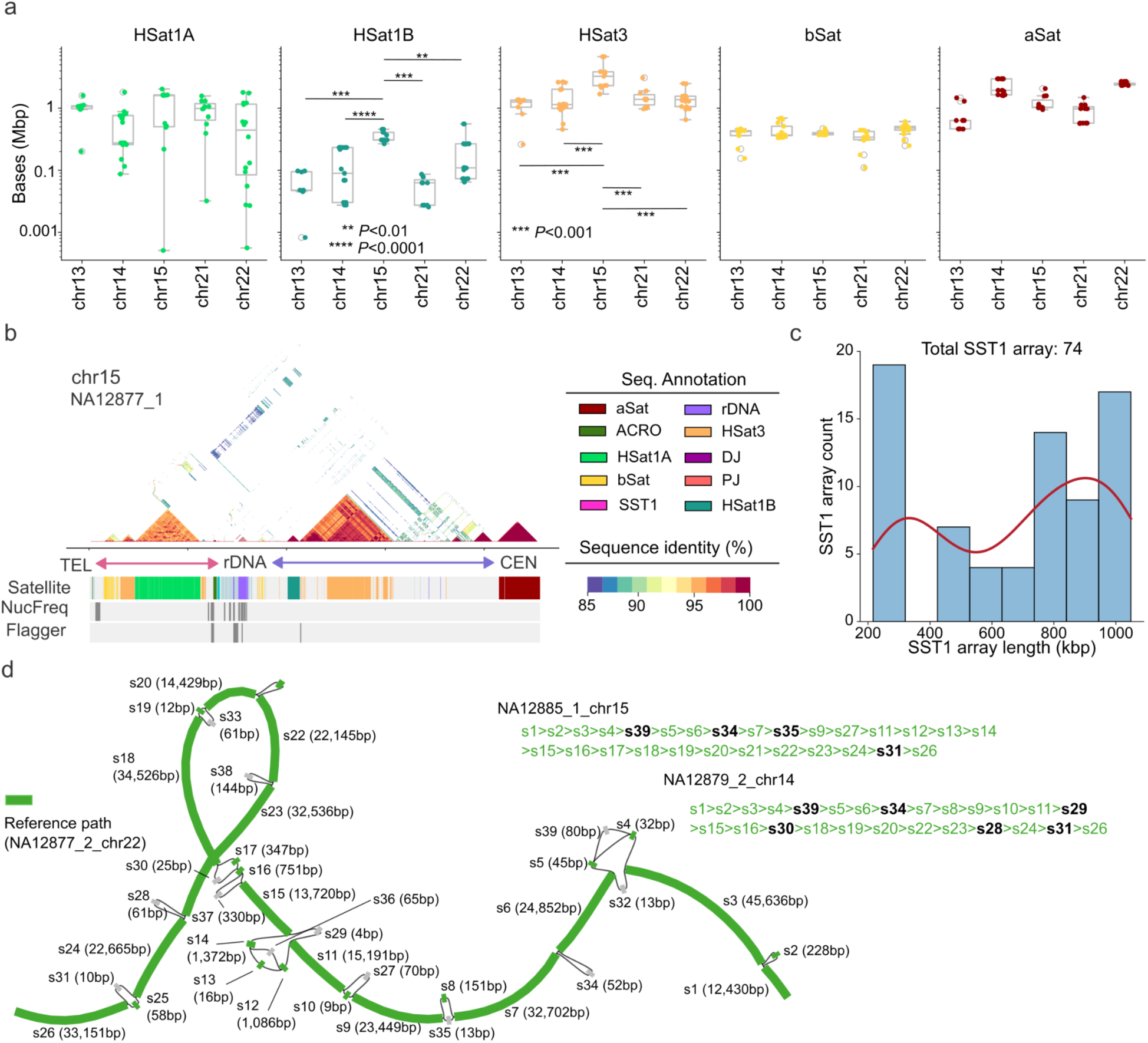
Genetic variations of short arm sequence. a) Comparison of satellite sequence among different chromosomes. A significant difference is determined using Mann-Whitney-Wilcoxon two-sided test; \*\**P*<0.01, \*\*\**P*<0.001, \*\*\*\**P*<0.0001. b) Satellite sequence organization on chr15 with a long HSat3 expansion on the proximal portion of p-arm. The diverged HSat3 sequence between HSat1B and HSat3 shares ∼95% identity with HSat3 repeats but is different from HSat1B. c) Length distribution of all SST1 repeat arrays within this family without assembly error. d) Minigraph (v0.21) using 12 correctly assembled distal junction (DJ) sequences within this family. The two example paths are the chr14 and chr15 DJ sequences from NA12885 and NA12879, respectively, when using chr22 from NA12877 as the reference path. The nodes in green indicate the reference sequence and bolded black indicates variation in the sequence.

**Figure S3.**
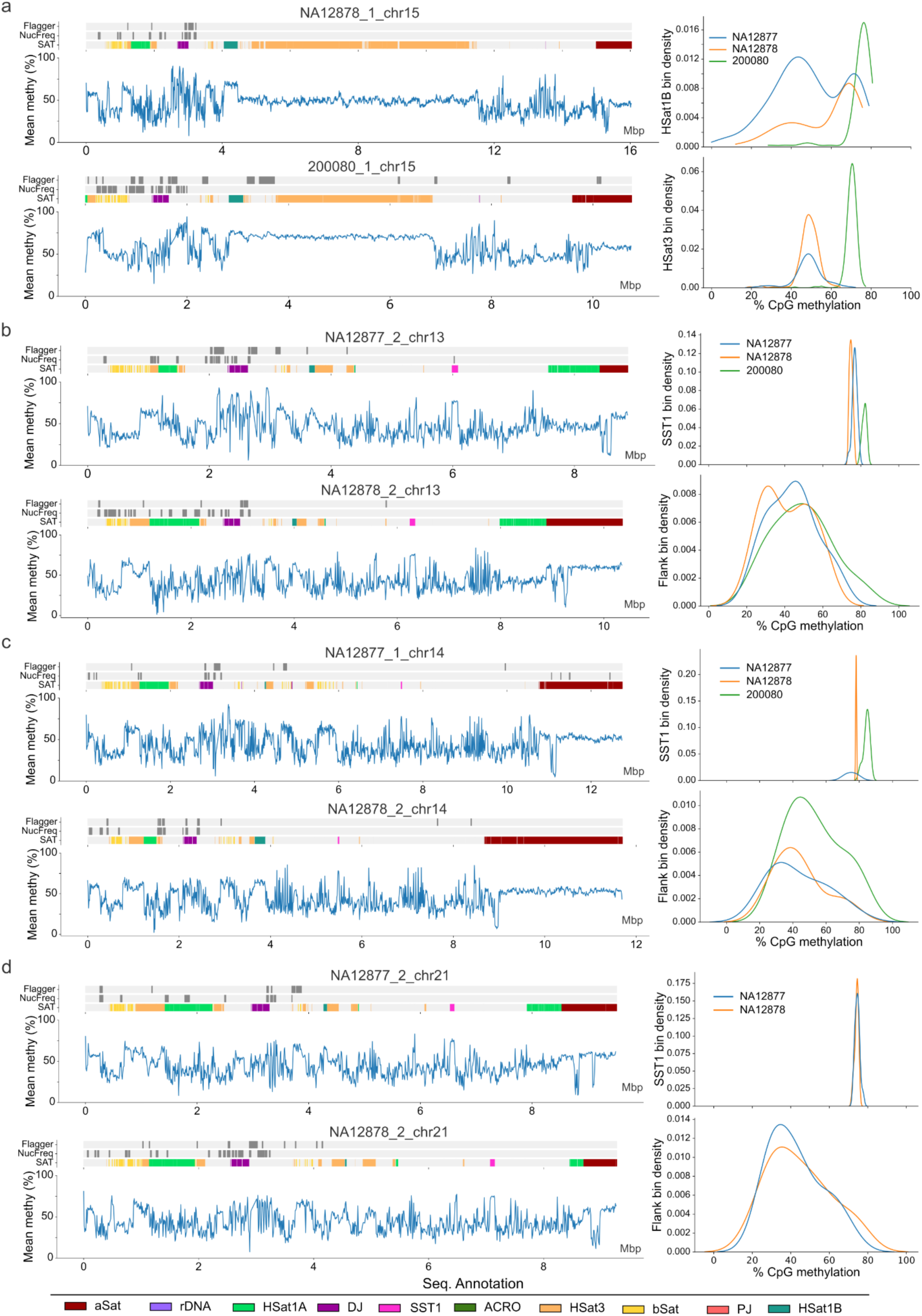
Epigenetic variation of satellite sequence on short arms. The assembly quality, satellite sequence annotation, and methylation of unrelated haplotypes of HSat3 from a) chr15 and SST1 from b) chr13, c) chr14 and d) chr21. a) The right panel shows the density of binned CpG methylation of HSat1B and HSat3 for each haplotype. b-d) The right panels show the density of binned CpG methylation of SST1 and the 50 kbp flanking region for each haplotype.

**Figure S4.**
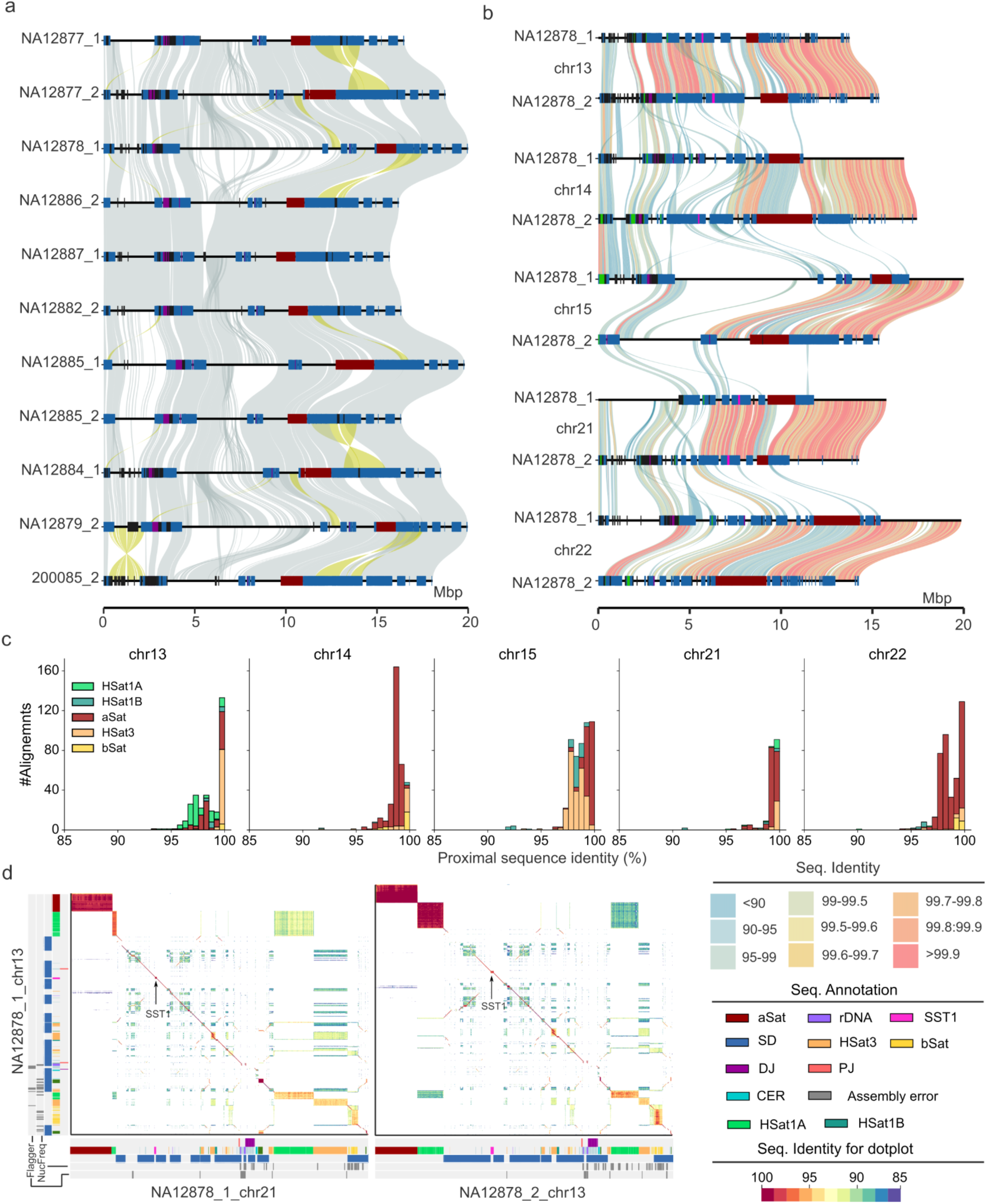
Allelic and nonallelic alignment of acrocentric short arms. a) Comparison of chr15 pq-scatigs containing both distal and proximal sequence assembled. The alignment reveals chr15 q-arm inversion polymorphism (reversed alignment in yellow) within this family. b) SVbyEye plot shows the all-vs-all alignment of all pq-scatigs from NA12878, revealing extensive allelic and nonallelic variations on short arms. c) Allelic sequence identity of 10 kbp binned alignment on short arms. This plot only shows the alignment in HSat1, aSat (*α*-satellite), HSat3, and bSat (*β*-satellite). All alignments inside incorrectly assembled regions are excluded in the evaluation. d) ModDotPlot (v0.9.8) shows the variation between chr13 homologous chromosomes as well as chr13 and chr21 heterologous chromosomes.

**Figure S5.**
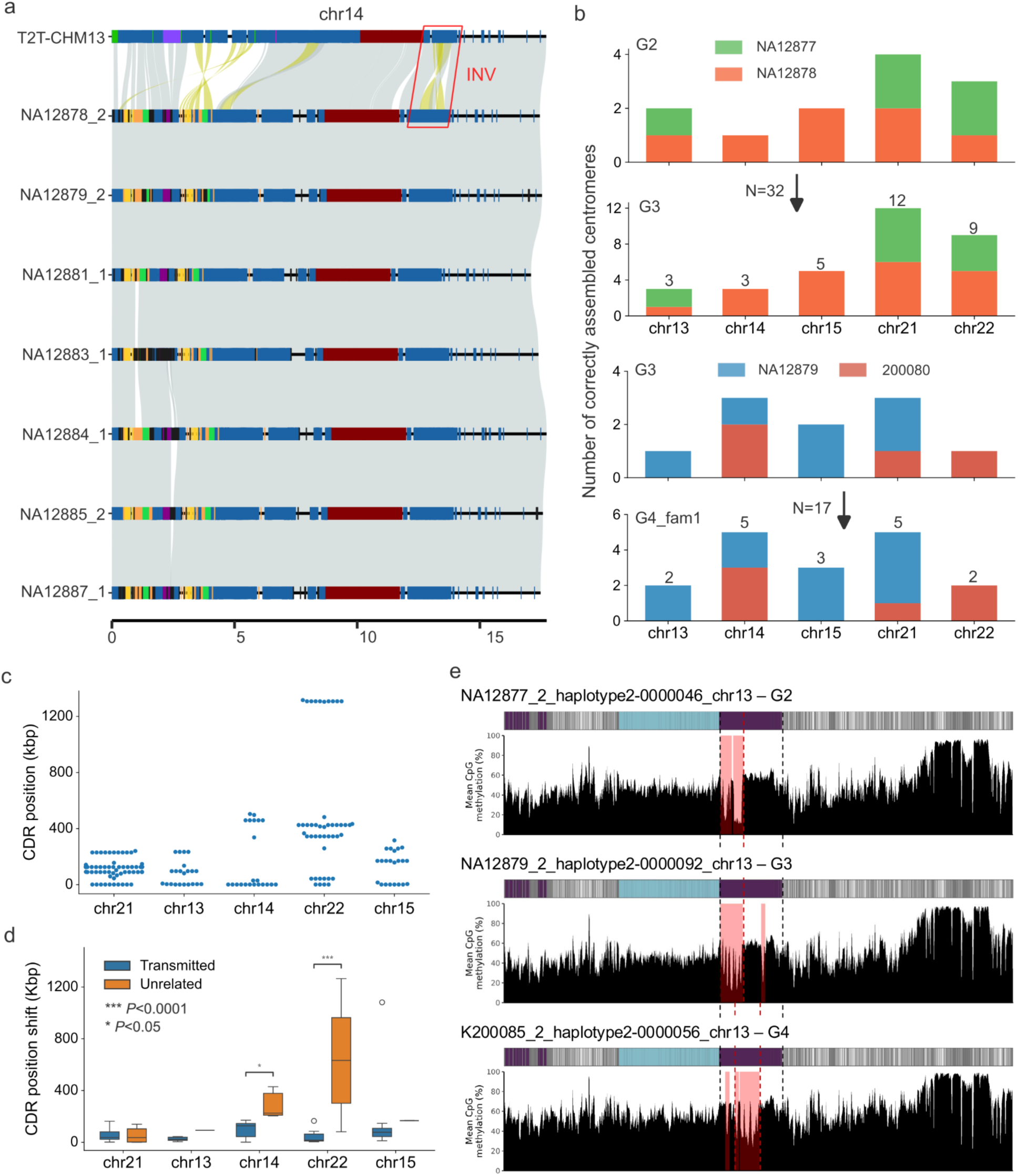
Intergenerational transmission of the short arms. a) Inheritance of an inversion detected against the T2T-CHM13 reference genome. b) Number of accessed intergenerational centromere transmissions from G2 to G3 and G3 to G4_fam1. c) Position of the centromere dip region (CDR), which is measured with respect to the start coordinate of the aSat (*α*-satellite) array. d) Comparison of CDR position shift between transmitted and unrelated individuals. A significant difference is determined using Mann-Whitney-Wilcoxon two-sided test; \**P*<0.05. \*\*\**P*<0.001. Note there are not enough correctly assembled chr13 and chr15 centromeres from unrelated individuals (NA12877, NA12878 and 200080) to make a comparison. e) Example of centromere transmitted from G2 to G4. The CDR positions in G2 and G3 are 3.6 kbp and 8 kbp to the start of the aSat, respectively, while the position becomes 53 kbp when transmitted to G4. The other three centromeres transmitted across two generations are shown in Data S1.

**Figure S6.**
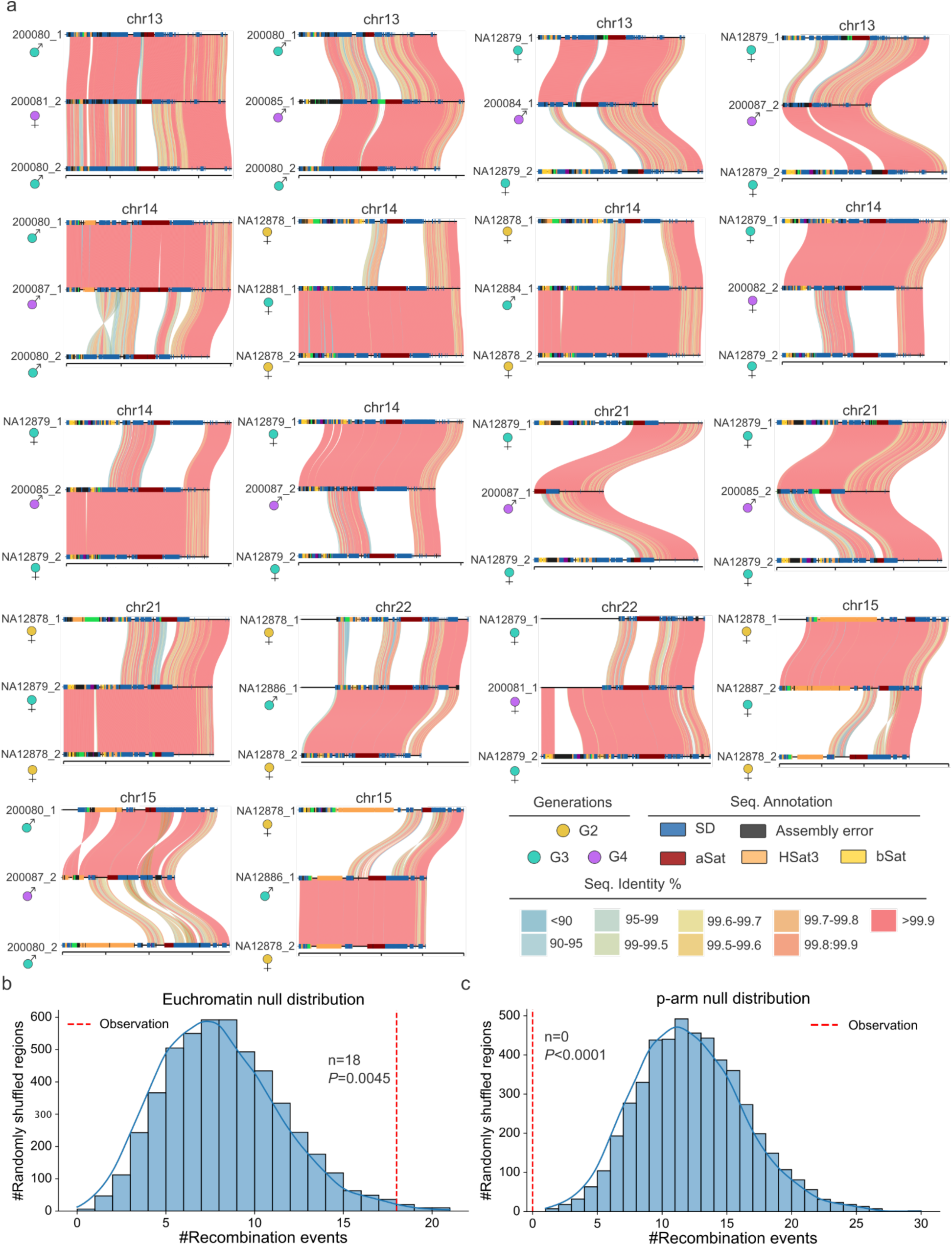
Summary of the 18 allelic recombinations and their comparison to the null distribution. a) The all-vs-all alignment of 18 q-arm pericentromeric recombinations visualized by SVbyEye. In each plot, the haplotype in the middle is the combination product and the parental haplotypes are on top and bottom, respectively. The read-depth profile and breakpoint alignments of each recombination are shown in **Data S1**. b) Euchromatin recombination distribution created by randomly shuffling segments across the genome exclude SAACs, metacentric pericentromeric regions on T2T-CHM13. The red dashed line is the number of observed recombinations on 25 Mbp acrocentric q-arm pericentromeric regions. c) Metacentric chromosome p-arm recombination distribution created by randomly shuffling a 36 Mbp segment on T2T-CHM13 SAAC proximal sequences. The red dashed line is the number of observed recombinations on SAAC regions of 38 Mbp.

**Figure S7.**
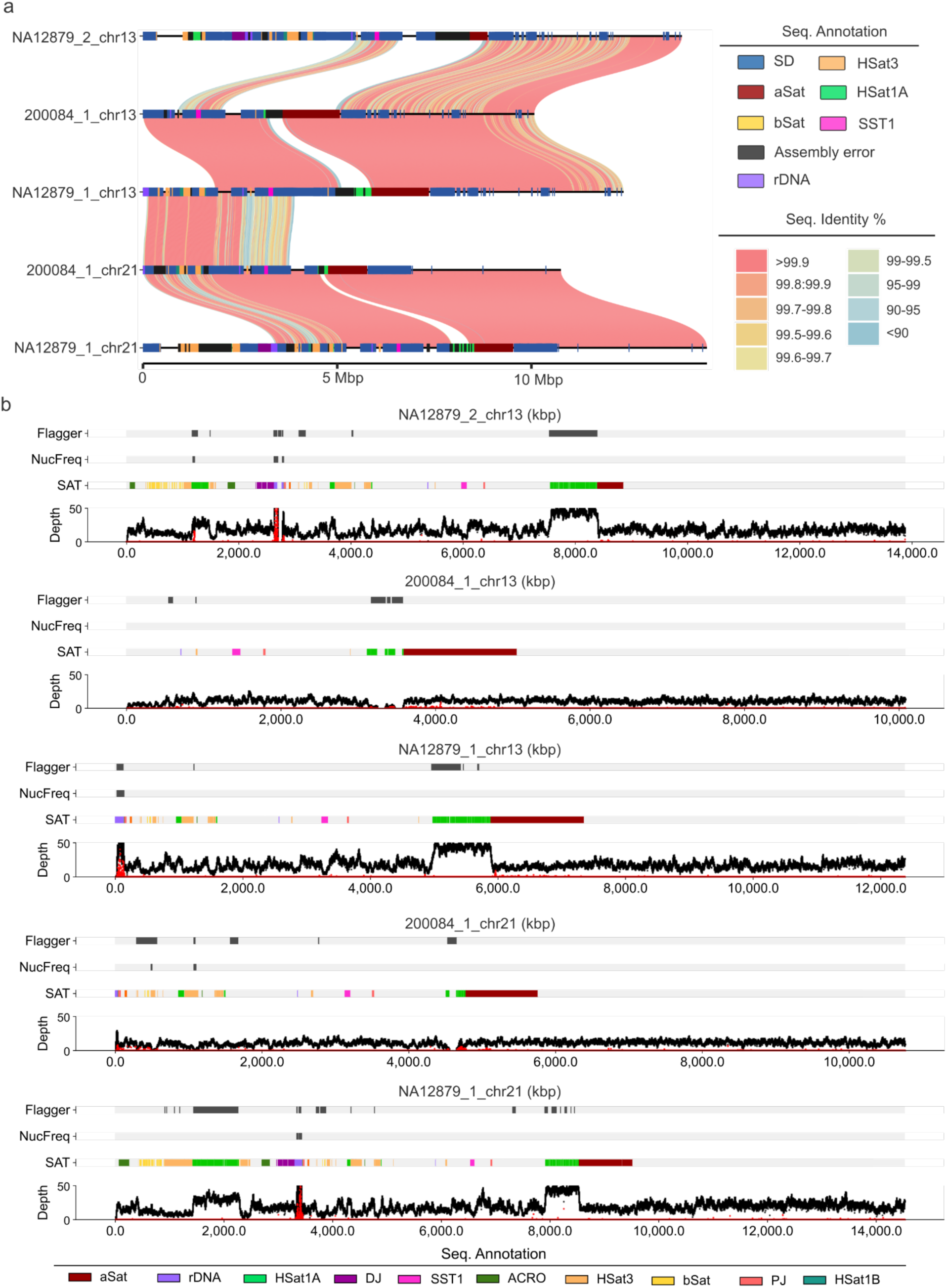
Chr13 and chr21 recombinations in sample 200084. a) SVbyEye plot shows the all-vs-all alignment of the two recombinations in 200084: one chr13-chr21 ectopic recombination and the other homologous recombination on chr13 q-arm. b) Assembly quality assessed by Flagger and NucFreq for the five contigs involved in the two recombinations.

**Figure S8.**
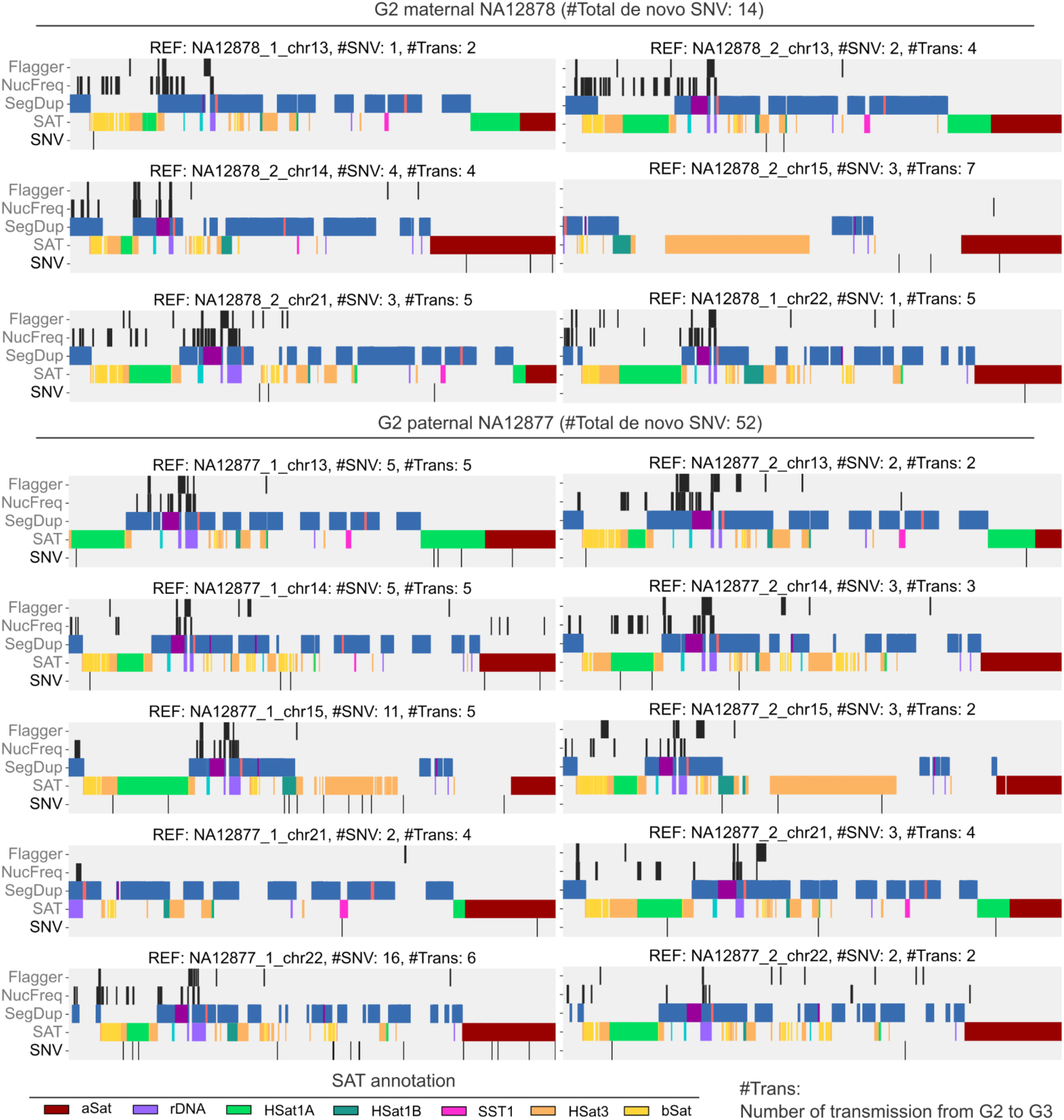
*De novo* SNVs detected in G3 against G2 parents NA12877 and NA12878. Each plot shows the detected variants against one parental haplotype. The reference haplotype name (REF), number of single-nucleotide variants (#SNV), and transmissions (#Trans) are indicated on the top of each plot. The number of transmissions is counted from G2 to eight G3 children (NA12879, NA12881, NA12882, NA12883, NA12884, NA12885, NA12886 and NA12887).

**Figure S9.**
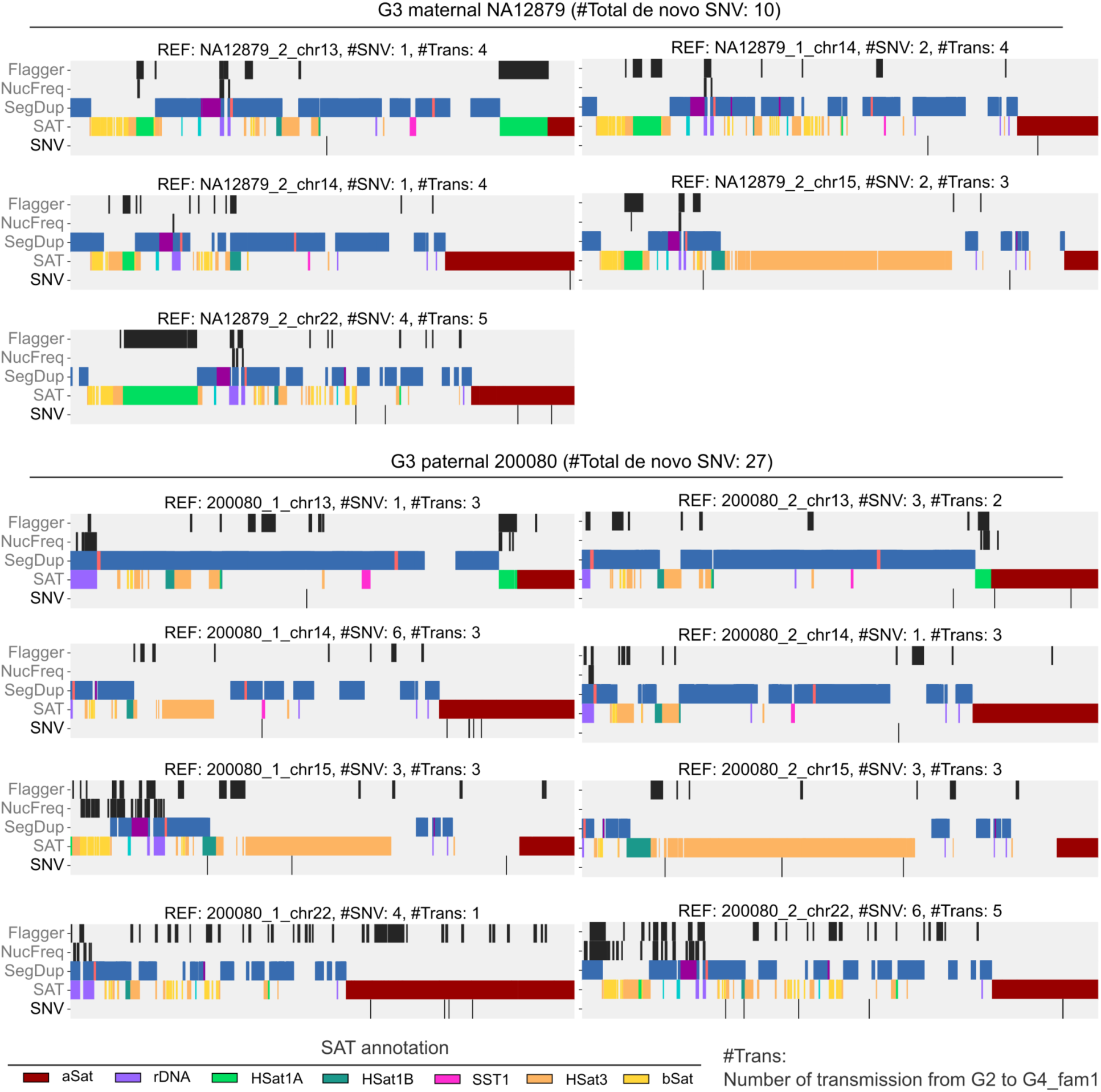
*De novo* SNVs detected in G4_fam1 against G3 parents NA12879 and 200080. Each plot shows the detected variants against one parental haplotype. The reference haplotype name (REF), number of single-nucleotide variants (#SNV), and transmissions (#Trans) are indicated correspondingly. The number of transmissions is counted from G3 (NA12879 and 200080) to five G4_fam1 children (200081, 200083, 200084, 200085 and 200087).

**Figure S10.**
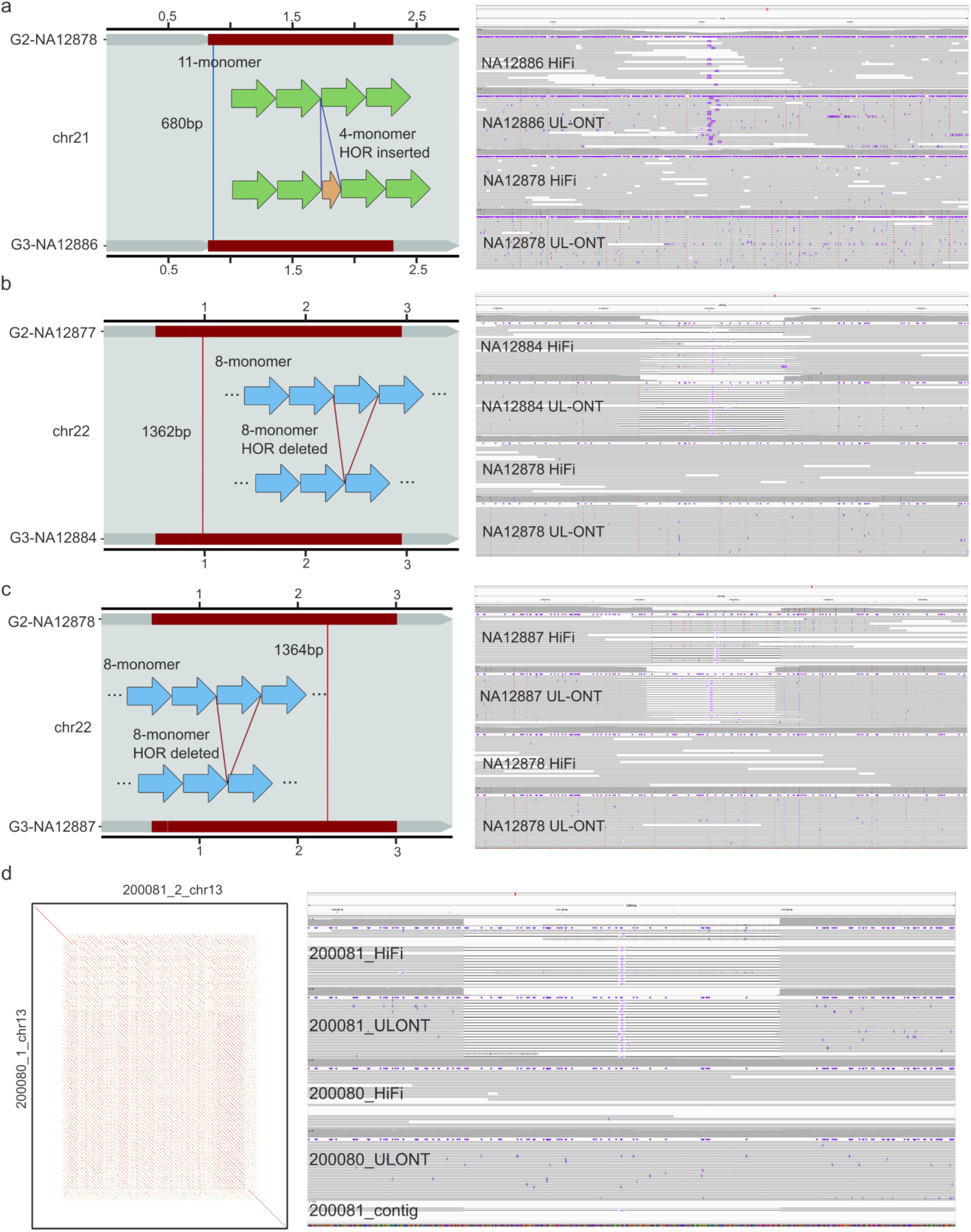
*De novo* SVs detected in HOR and SST1 arrays. a-c) Three *de novo* structural variants (SVs) detected in centromere higher-order repeat (HOR) arrays. The left alignment shows the transmission and location of the SV. The right panel is the IGV screenshot of HiFi and UL-ONT reads aligned to the paternal reference haplotype. d) The left panel shows the difference of the SST1 repeat array with the reads supported (right IGV screenshot) one 1407 bp deletion inside the array.

