## Supplementary material for "Human acrocentric chromosome short arm *de novo* mutation and recombination": Data S1

### Data S1 for:
### Human acrocentric chromosome short arm *de novo* mutation and recombination

#### Figures 1-48

Jiadong Lin[1], F. Kumara Mastrorosa[1], Michelle D. Noyes[1], DongAhn Yoo[1], Arang Rhie[2], David Porubsky[1,3], Kendra Hoekzema[1], Katherine M. Munson[1], Nidhi Koundinya[1], W. Scott Watkins[4], Lynn B. Jorde[4], Aaron R. Quinlan[4], Deborah W. Neklason[5], Adam M. Phillippy[2], Evan E. Eichler[1,6]

**Affiliations**
[1]Department of Genome Sciences, University of Washington School of Medicine, Seattle, WA, USA
[2]Genome Informatics Section, Center for Genomics and Data Science Research, National Human Genome Research Institute, National Institutes of Health, Bethesda, MD, USA
[3]European Molecular Biology Laboratory (EMBL), Genome Biology Unit, Heidelberg, Germany
[4]Department of Human Genetics, University of Utah, Salt Lake City, UT, USA
[5]Department of Internal Medicine, University of Utah, Salt Lake City, UT, USA
[6]Howard Hughes Medical Institute, University of Washington, Seattle, WA, USA

**Corresponding author**
Evan E. Eichler, Ph.D.
Department of Genome Sciences
University of Washington School of Medicine
3720 15th Ave NE, S413C
Box 355065
Seattle, WA 98195-5065


**Abbreviations**
**DJ:** Distal junction
**PJ:** Proximal junction
**aSat:** $\alpha$-satellite
**bSat:** $\beta$-satellite
**SD:** Segmental duplication
**TEL:** Telomere
**CEN:** Centromere
**SAT:** Satellite

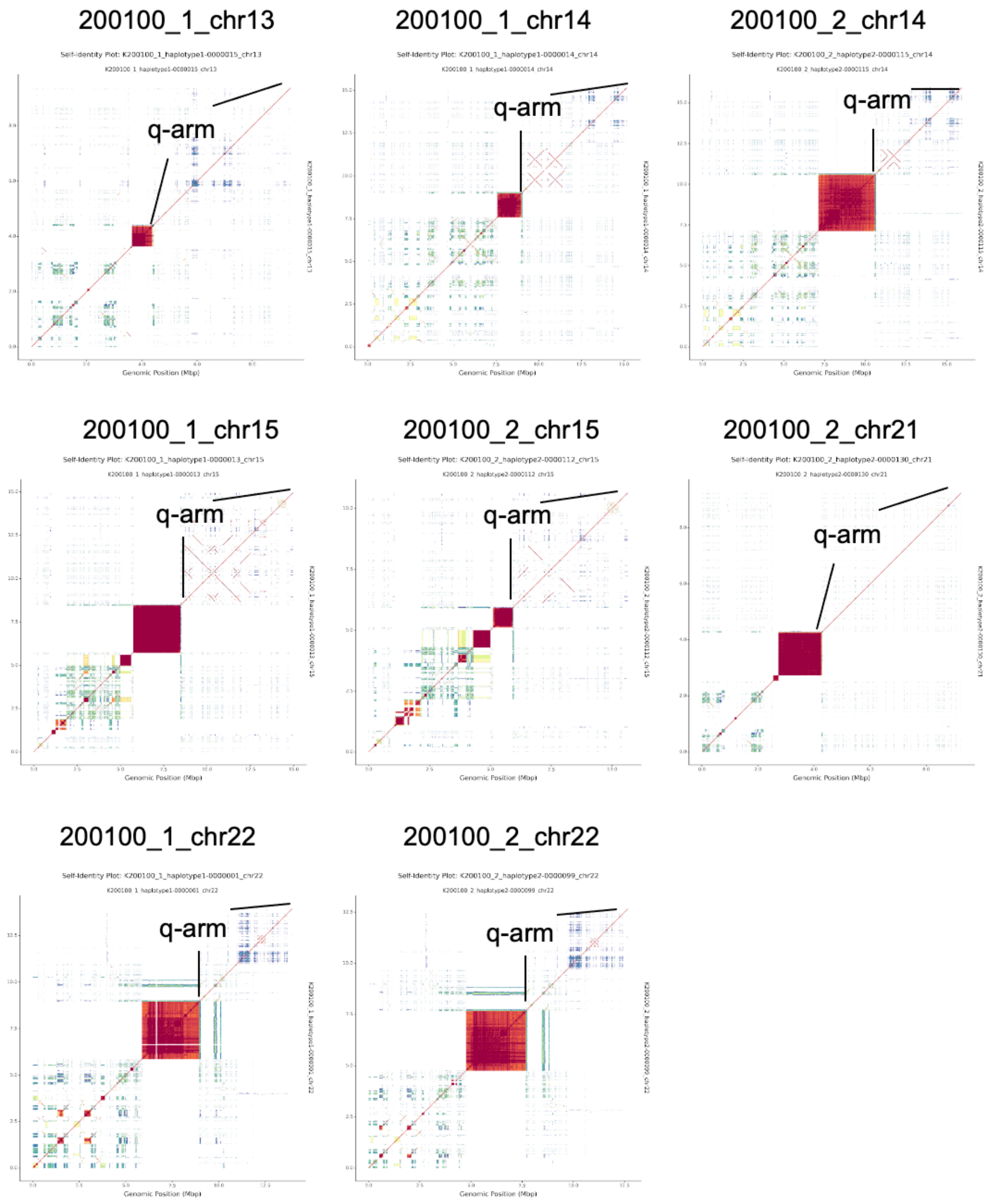


**Figure 1. ModDotPlot views of sample 200100 pq-scatigs.**

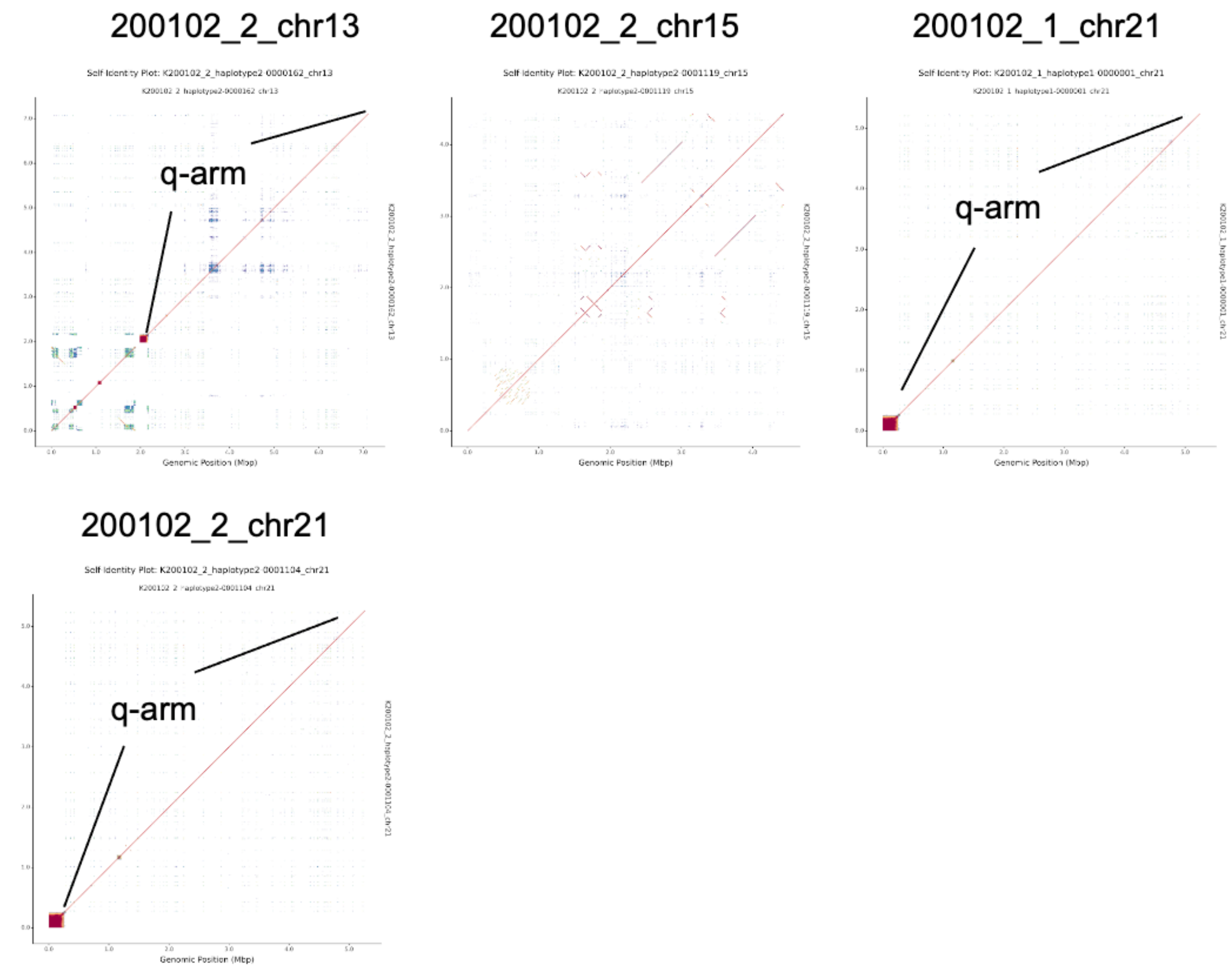


**Figure 2. ModDotPlot views of sample 200102 pq-scatigs**

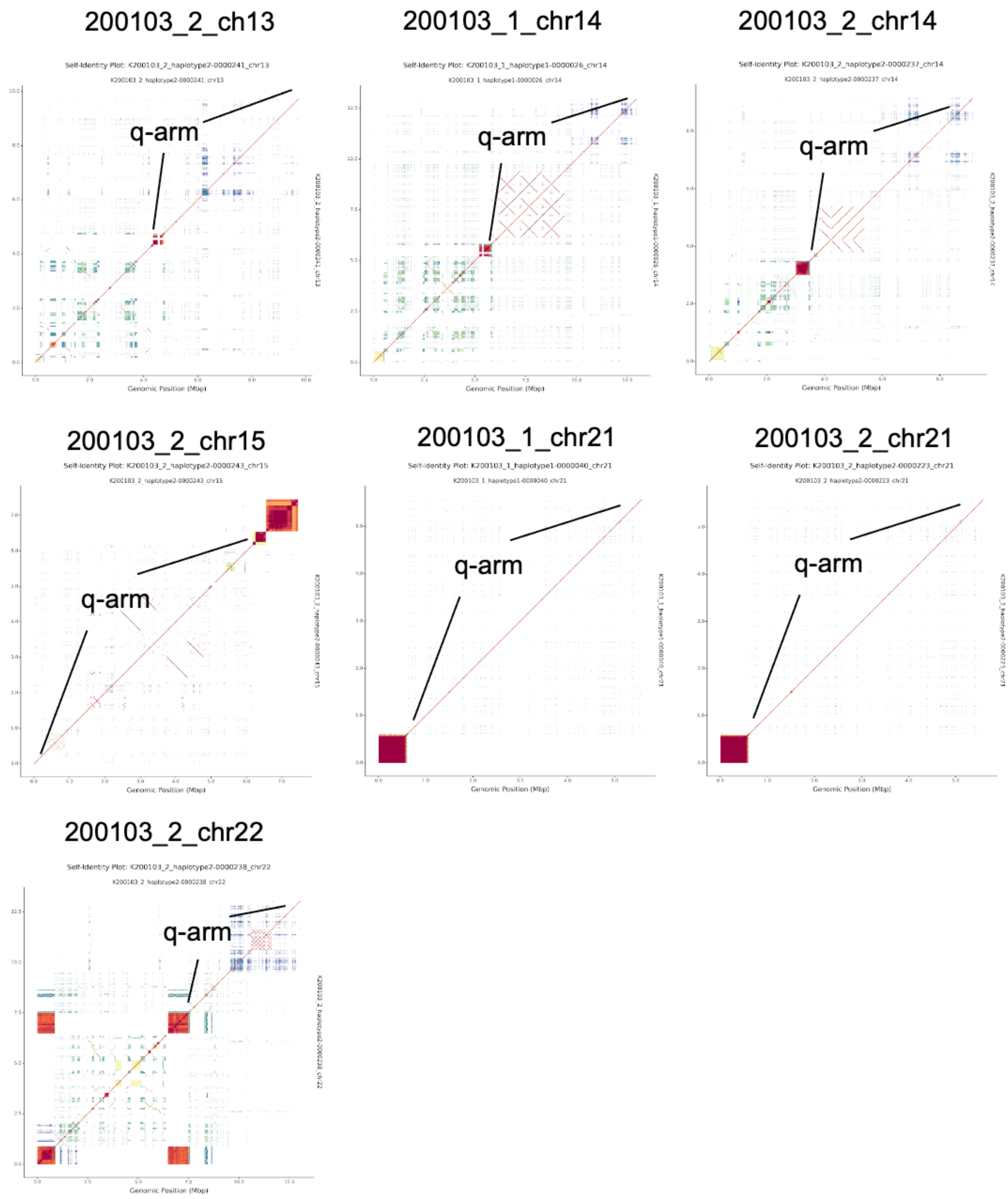


**Figure 3. ModDotPlot views of sample 200103 pq-scatigs.**

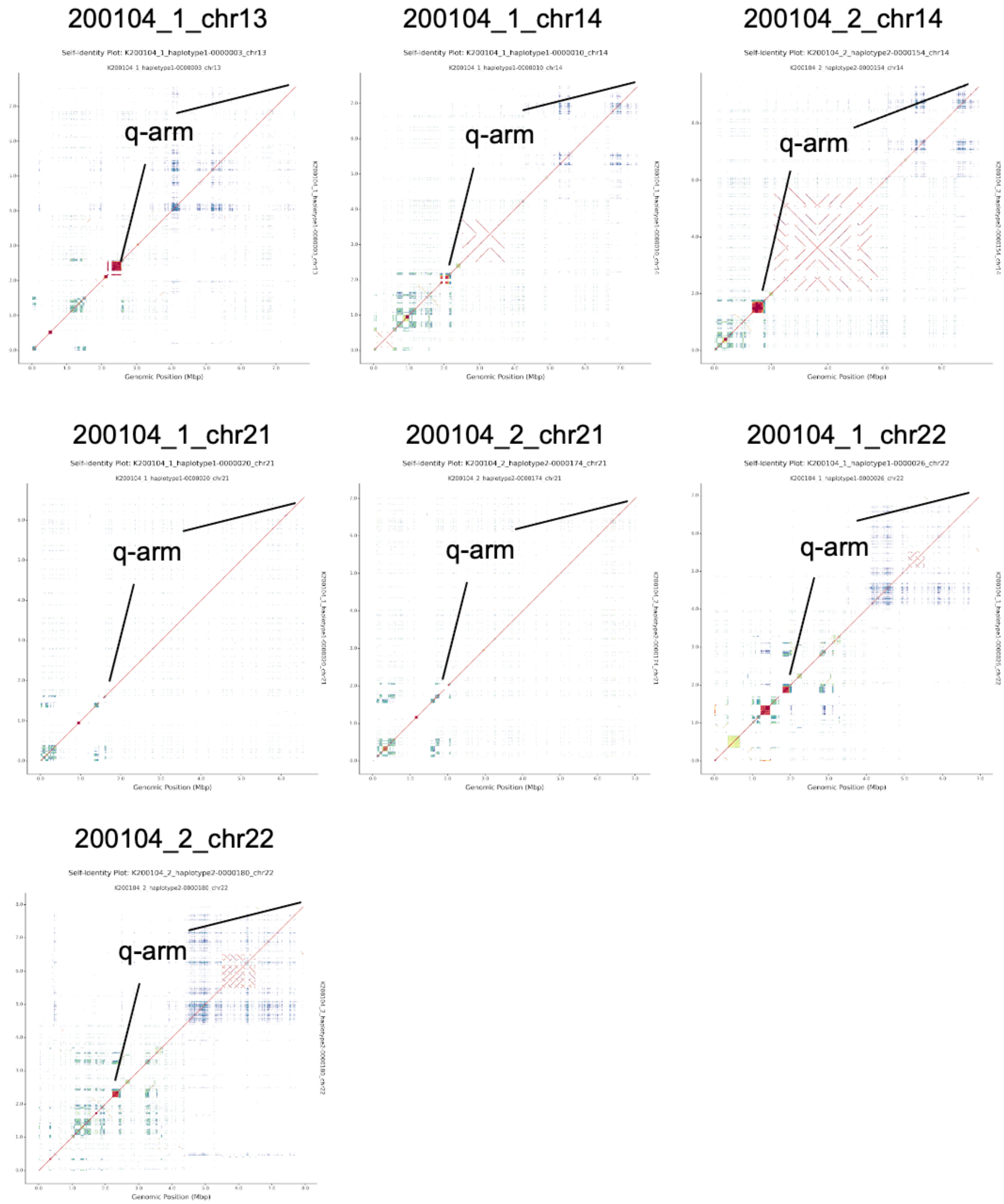


**Figure 4. ModDotPlot views of sample 200104 pq-scatigs.**

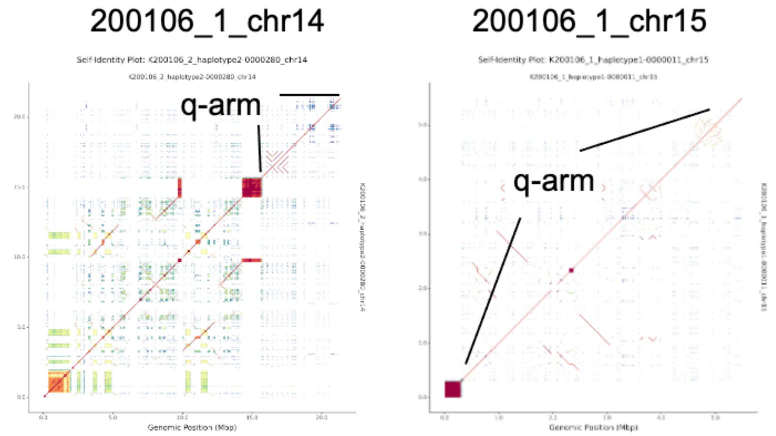


**Figure 5. ModDotPlot views of sample 200106 pq-scatigs.**

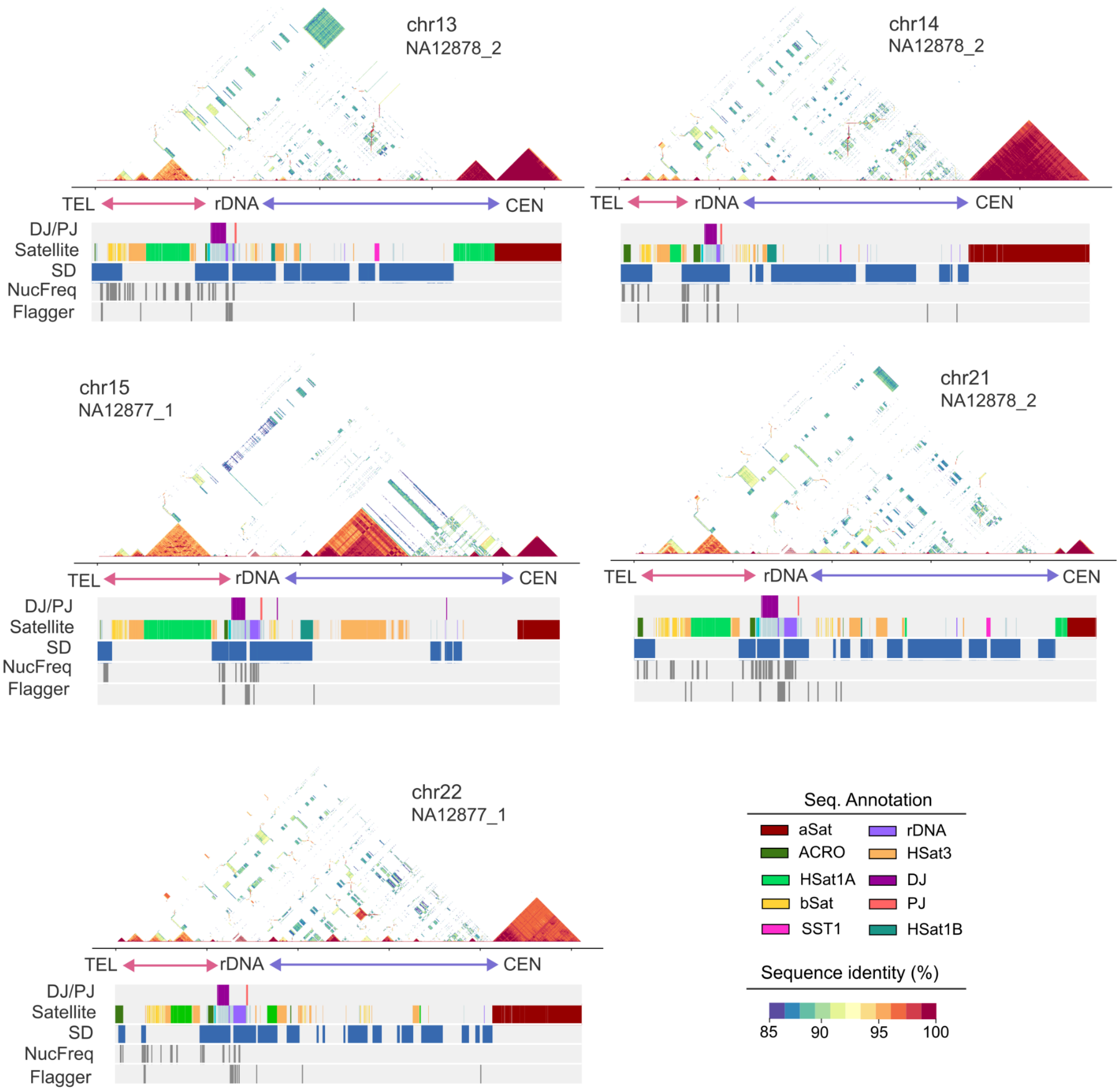


**Figure 6. Sequence organization of acrocentric short arms.** These five G2 haplotypes are the most frequently transmitted haplotypes to G4_fam1.

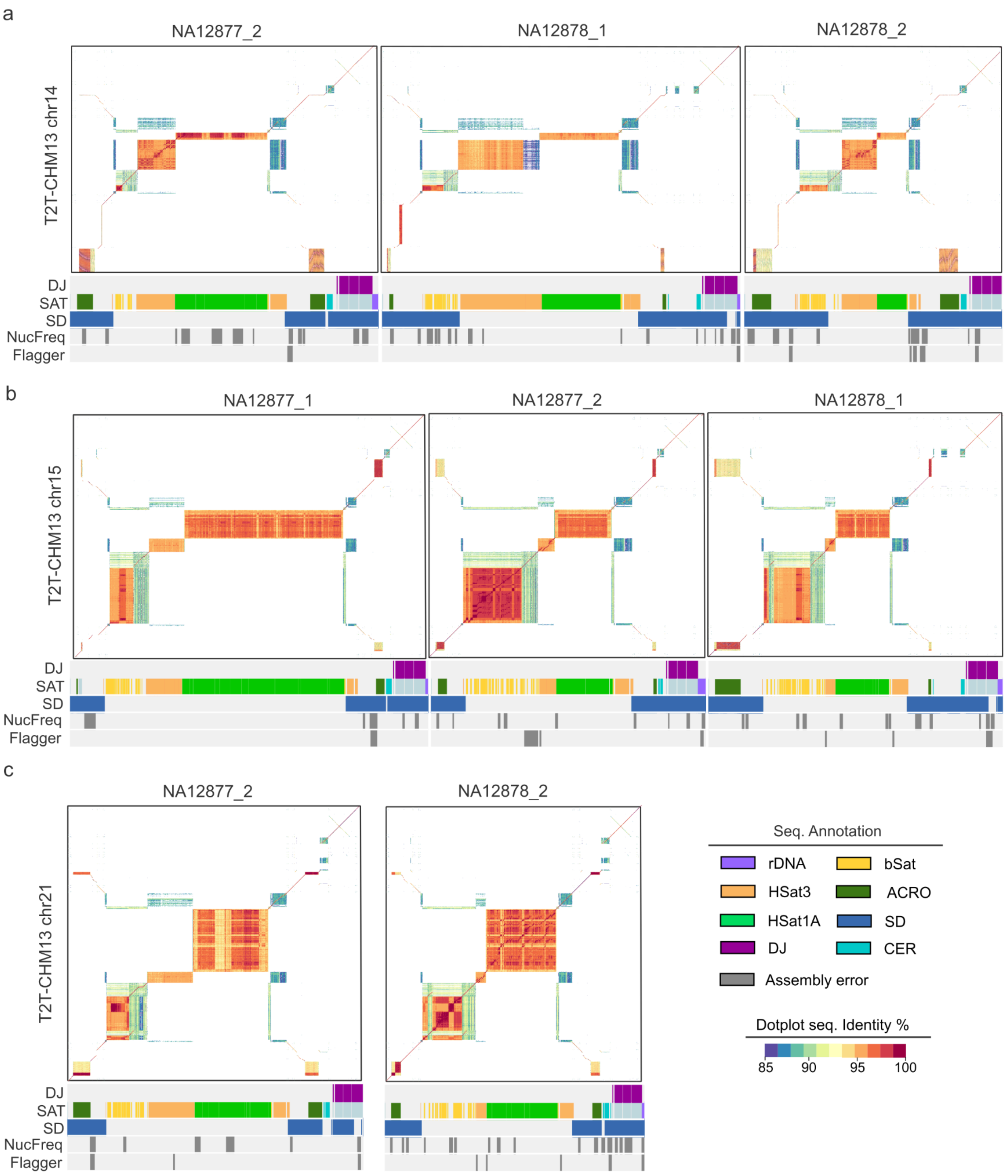


**Figure 7. Variation of distal sequences compared against T2T-CHM13 human reference genome.**

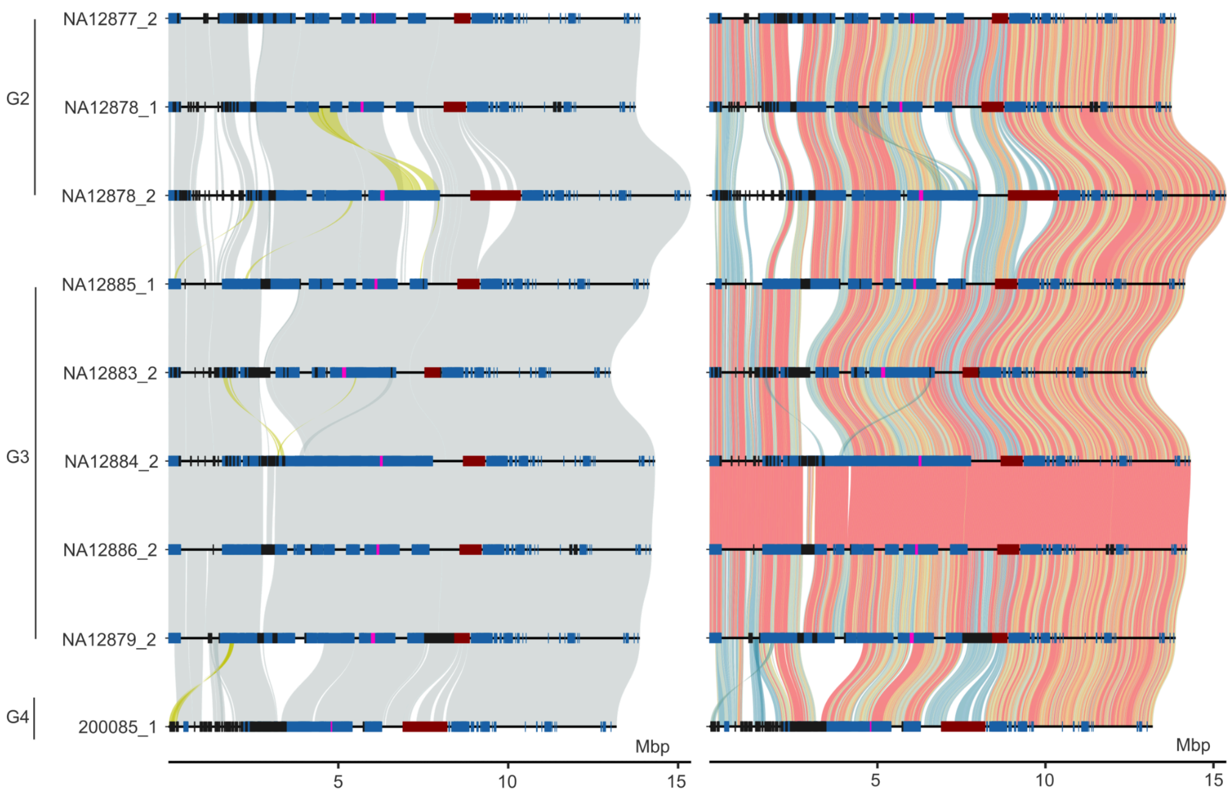


**Figure 8. The all-vs-all alignment of nine completely assembled chr13 pq-scatigs with distal (band p13) and proximal (band p11) sequence.** From top to bottom, the sequences are ordered by G2, G3, and G4. The left panel is colored by alignment orientation (yellow: reverse) and the right panel is colored by alignment identity. The red, blue, black, and pink rectangles on each haplotype represent aSat ($\alpha$-satellite), segmental duplication, assembly error, and SST1 array, respectively.

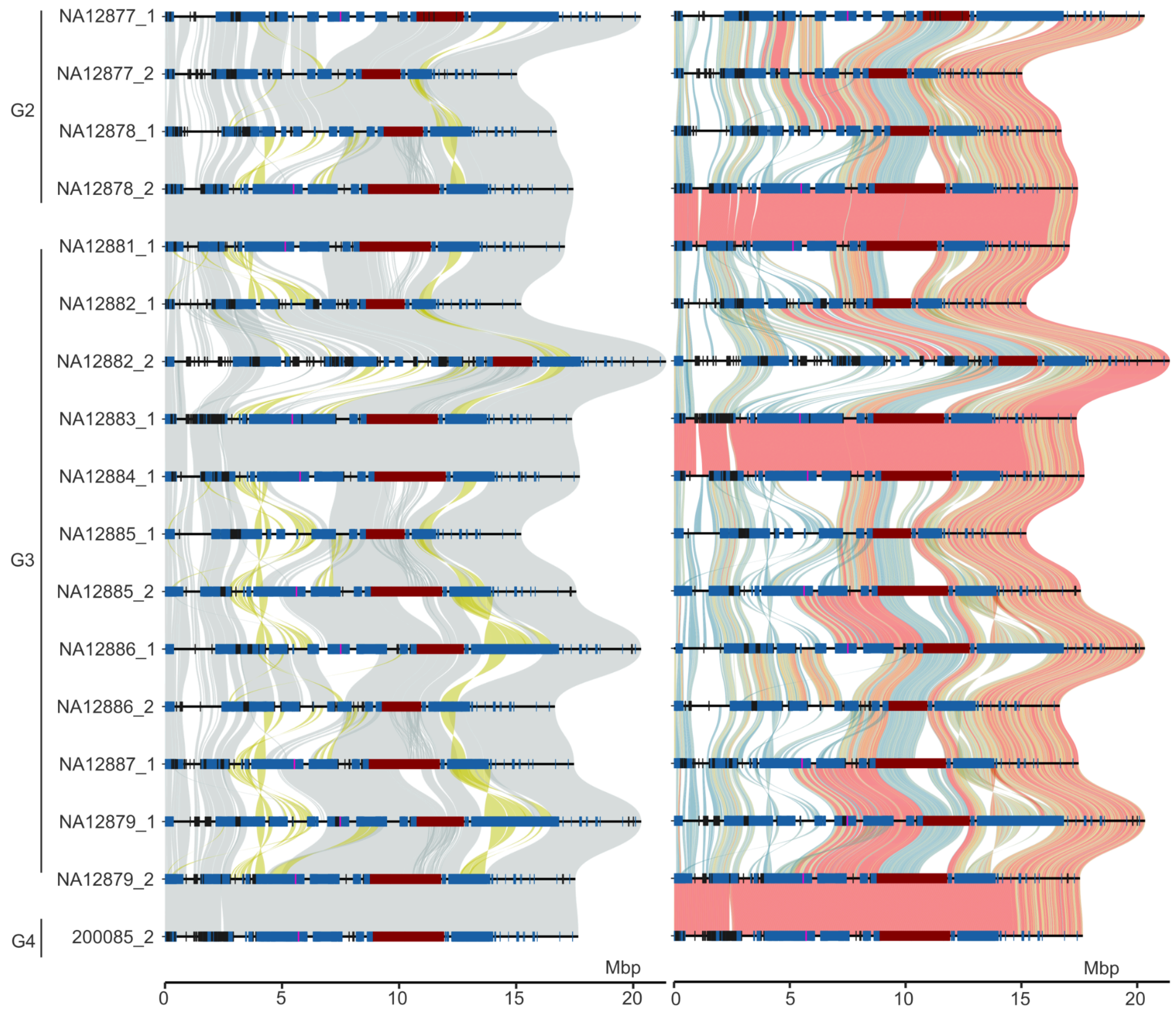


**Figure 9. The all-vs-all alignment of 17 completely assembled chr14 short arms pq-scatigs with distal (band p13) and proximal (band p11) sequence.** From top to bottom, the sequences are ordered by G2, G3, and G4. The left panel is colored by alignment orientation (yellow: reverse) and the right panel is colored by alignment identity. The red, blue, black, and pink rectangles on each haplotype represent aSat ($\alpha$-satellite), segmental duplication, assembly error, and SST1 array, respectively.

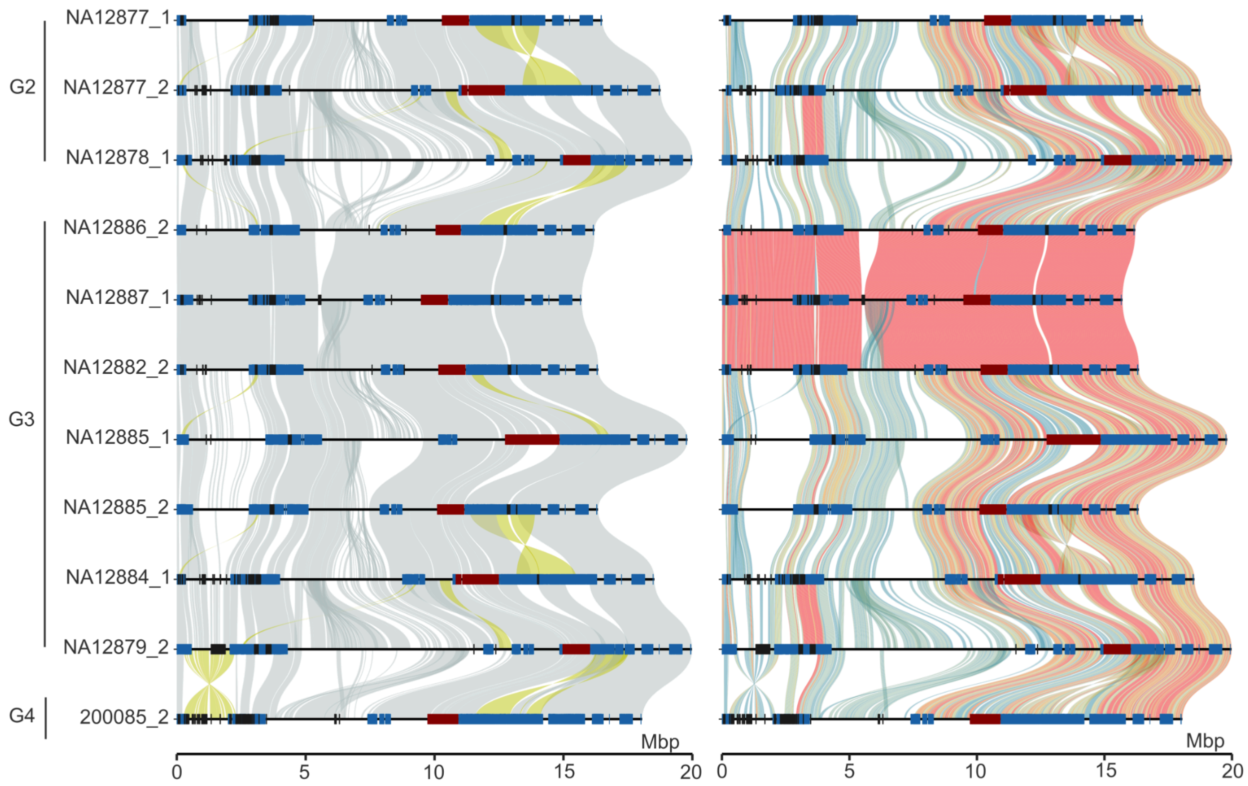


**Figure 10. The all-vs-all alignment of 11 completely assembled chr15 pq-scatigs with distal (band p13) and proximal (band p11) sequence.** From top to bottom, the sequences are ordered by G2, G3, and G4. The left panel is colored by alignment orientation (yellow: reverse) and the right panel is colored by alignment identity. The red, blue, and black rectangles on each haplotype represent aSat ($\alpha$-satellite), segmental duplication, and assembly error, respectively.

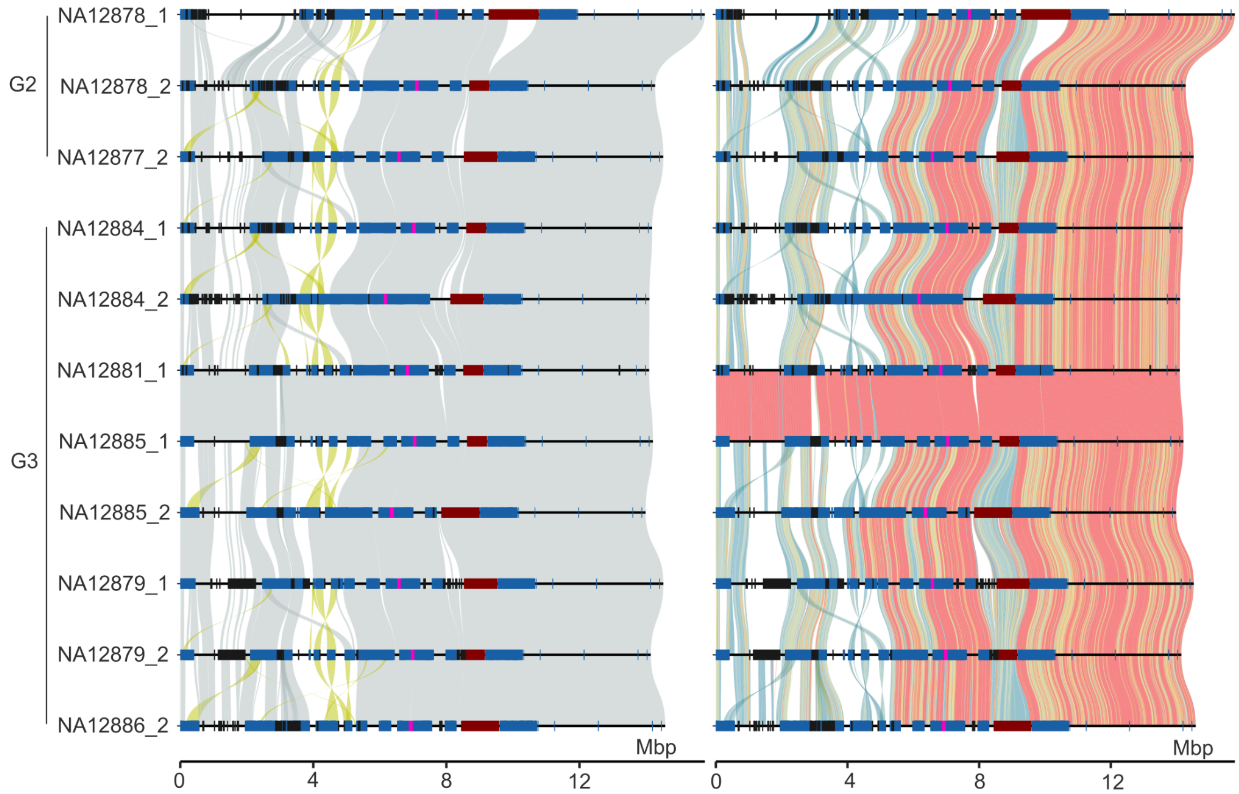


**Figure 11. The all-vs-all alignment of 11 completely assembled chr21 pq-scatigs with distal (band p13) and proximal (band p11) sequence.** From top to bottom, the sequences are ordered by G2, G3, and G4. The left panel is colored by alignment orientation (yellow: reverse) and the right panel is colored by alignment identity. The red, blue, black, and pink rectangles on each haplotype represent aSat ($\alpha$-satellite), segmental duplication, assembly error, and SST1 array, respectively.

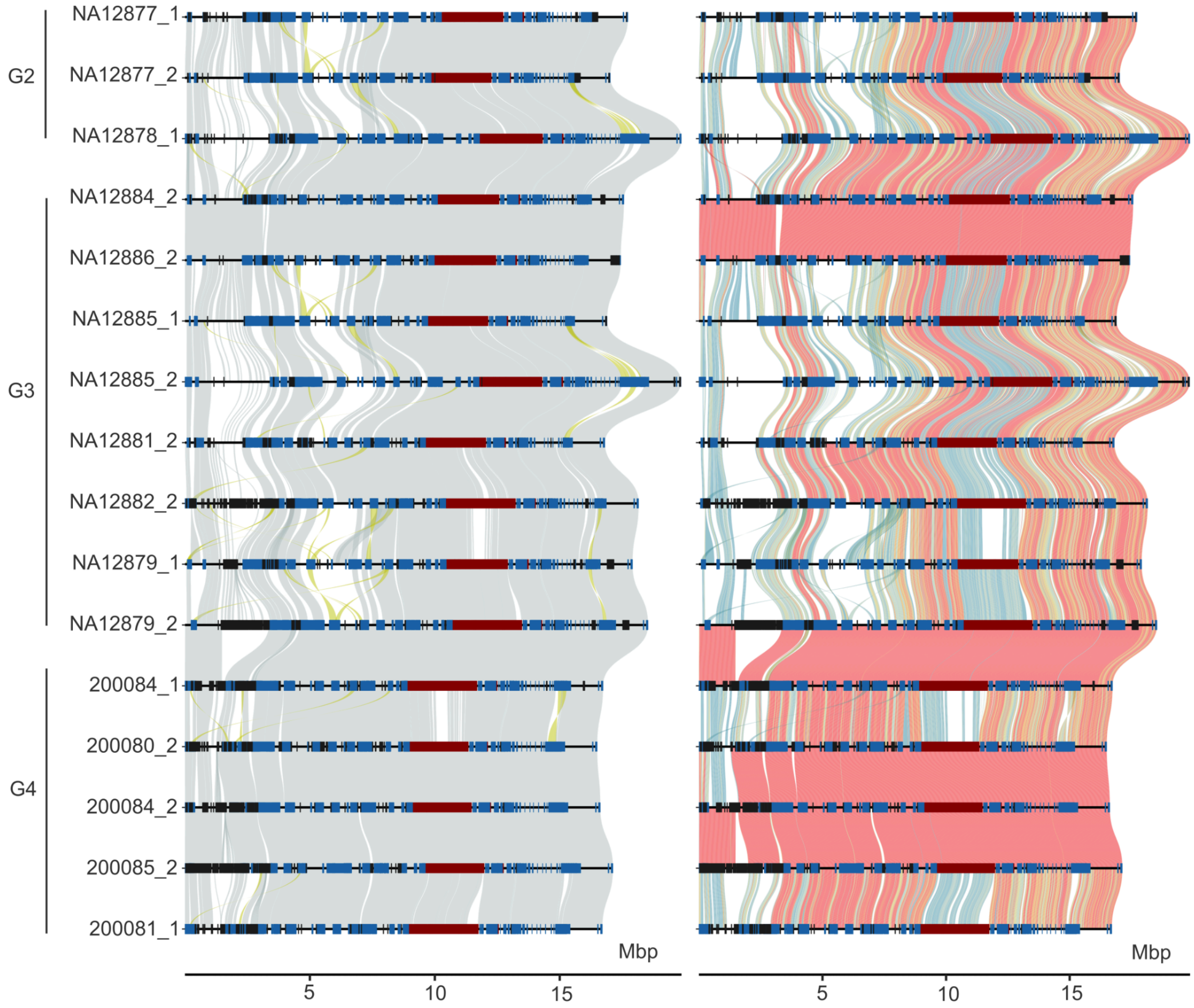


**Figure 12. The all-vs-all alignment of 16 completely assembled chr22 pq-scatigs with distal (band p13) and proximal (band p11) sequence.** From top to bottom, the sequences are ordered by G2, G3, and G4. The left panel is colored by alignment orientation (yellow: reverse) and the right panel is colored by alignment identity. The red, blue, and black rectangles on each haplotype represent aSat ($\alpha$-satellite), segmental duplication, and assembly error, respectively.

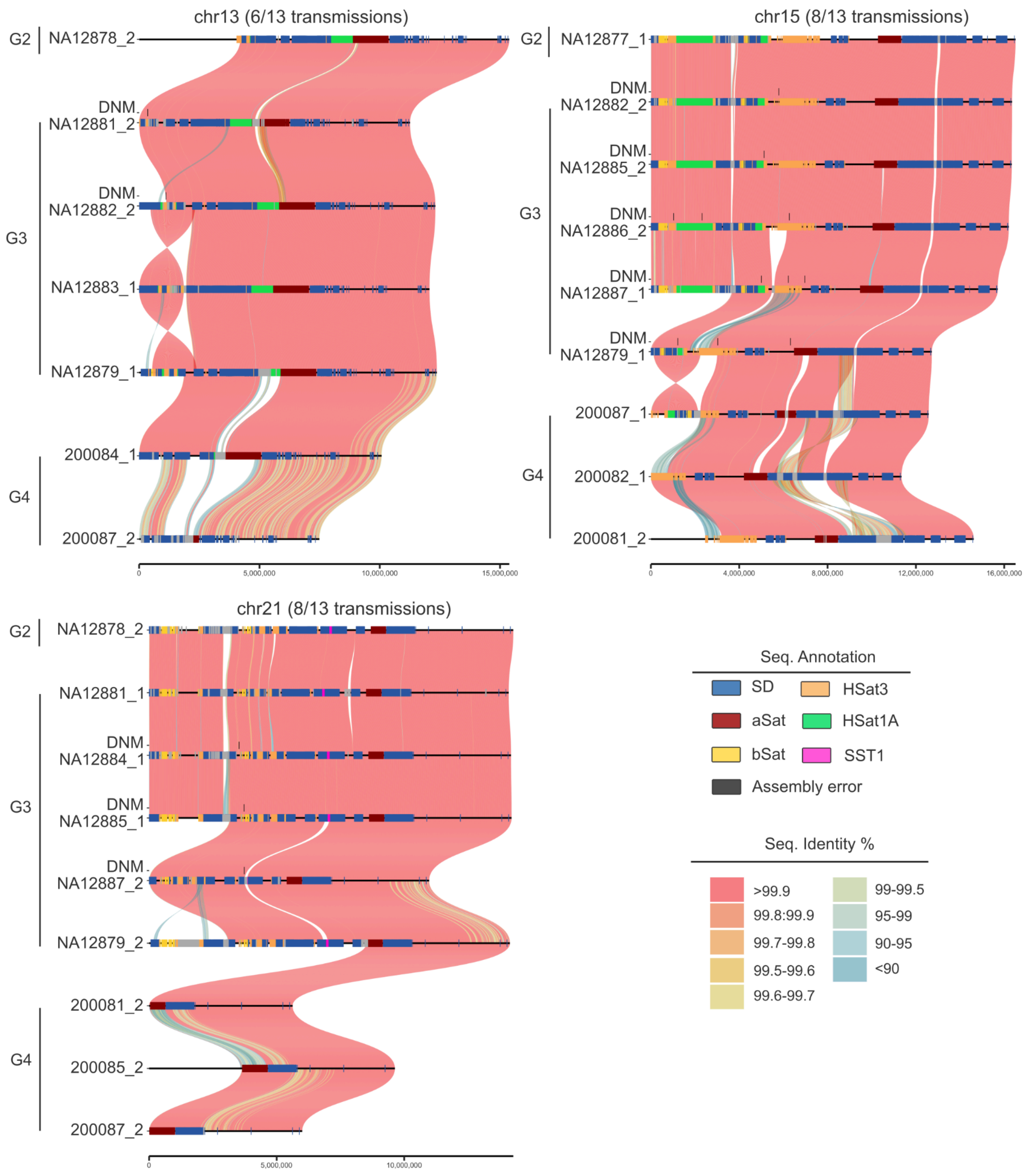


**Figure 13. Most frequently transmitted haplotypes of chr13, chr15, and chr21 from G2 to G4.** There are a total of 13 transmissions from G2 to G4 (eight in G3 and five in G4) for each haplotype in G2. Potential sites of *de novo* mutation (DNM) are indicated by black ticks along with SD annotation (blue bars). Note that the G3 DNMs are detected against the G2 reference and G4 DNMs are detected against the G3 parental haplotype.

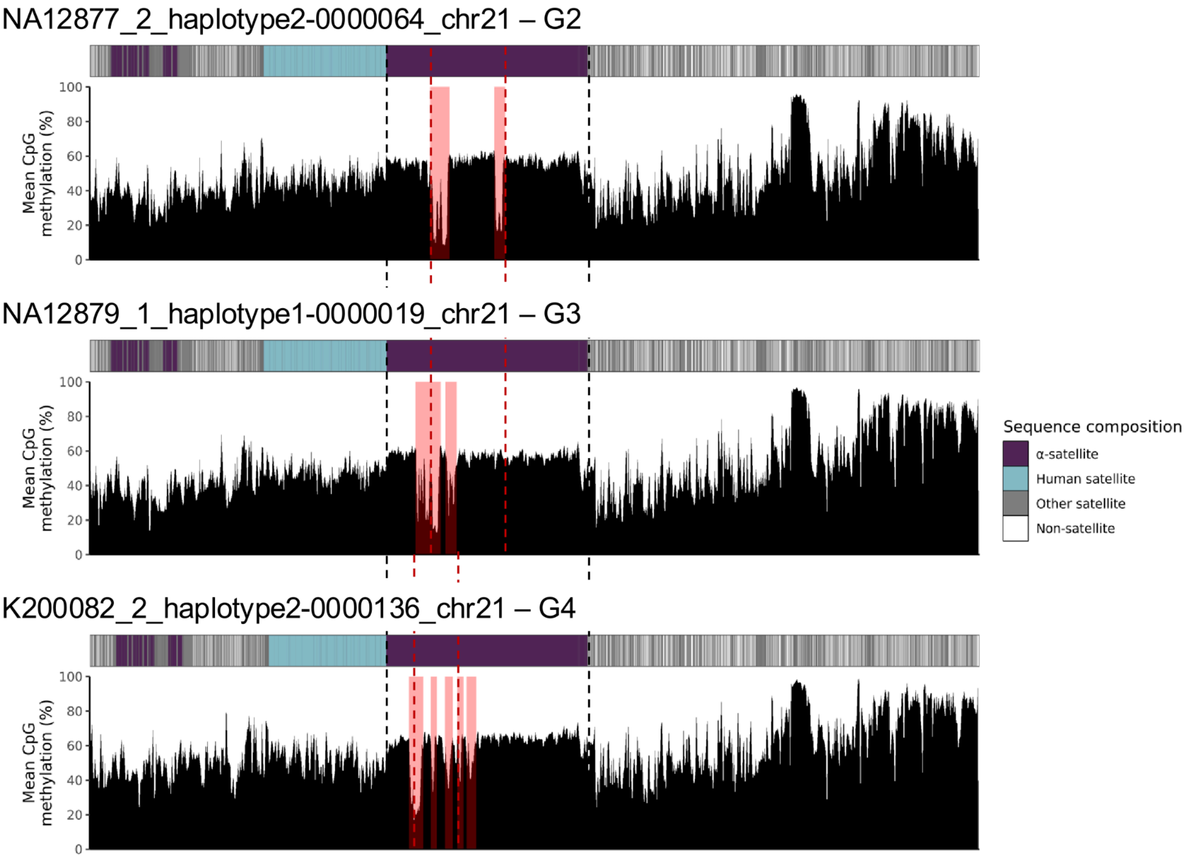


**Figure 14. Transmission of chr21 centromere from G2 to G4.** The pink rectangle indicates the centromere dip region (CDR) detected by CDR-Finder.

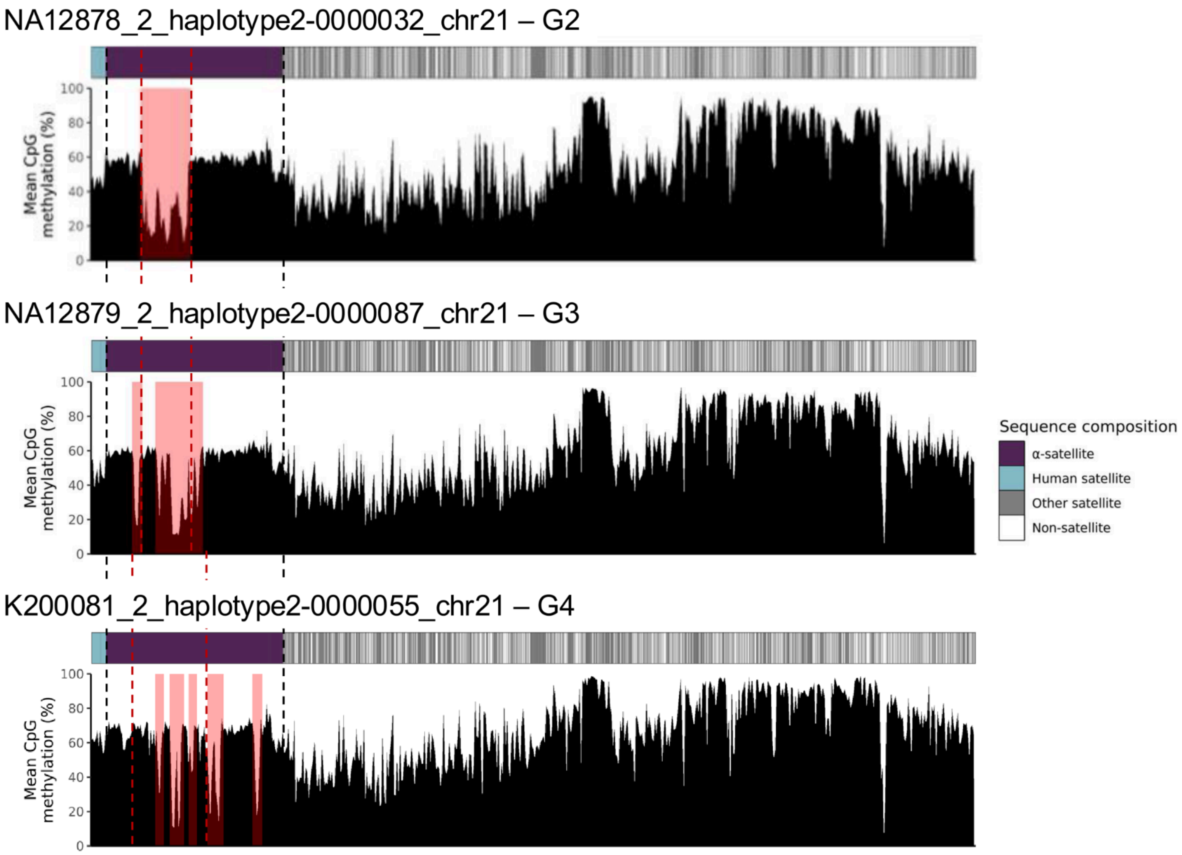


**Figure 15. Transmission of chr21 centromere from G2 to G4.** The pink rectangle indicates the CDR detected by CDR-Finder.

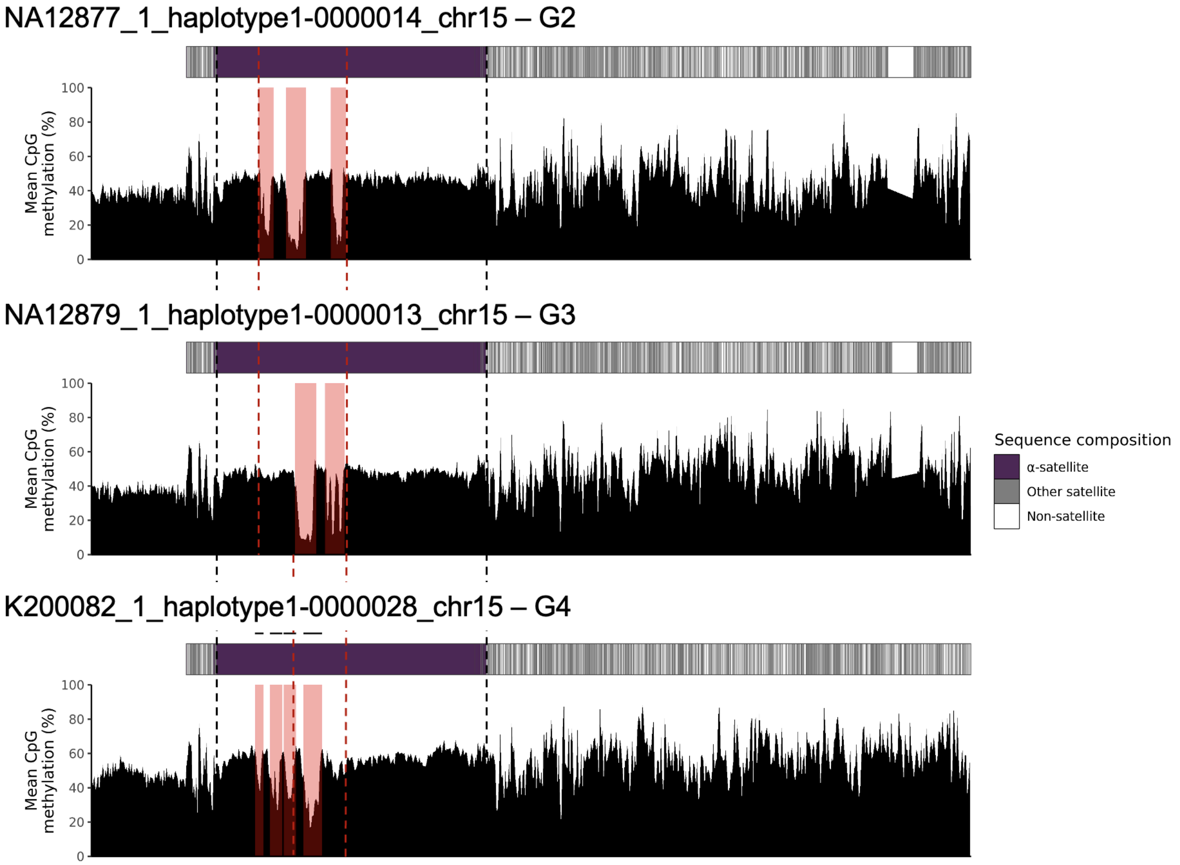


**Figure 16. Transmission of chr15 centromere from G2 to G4.** The pink rectangle indicates the CDR detected by CDR-Finder.

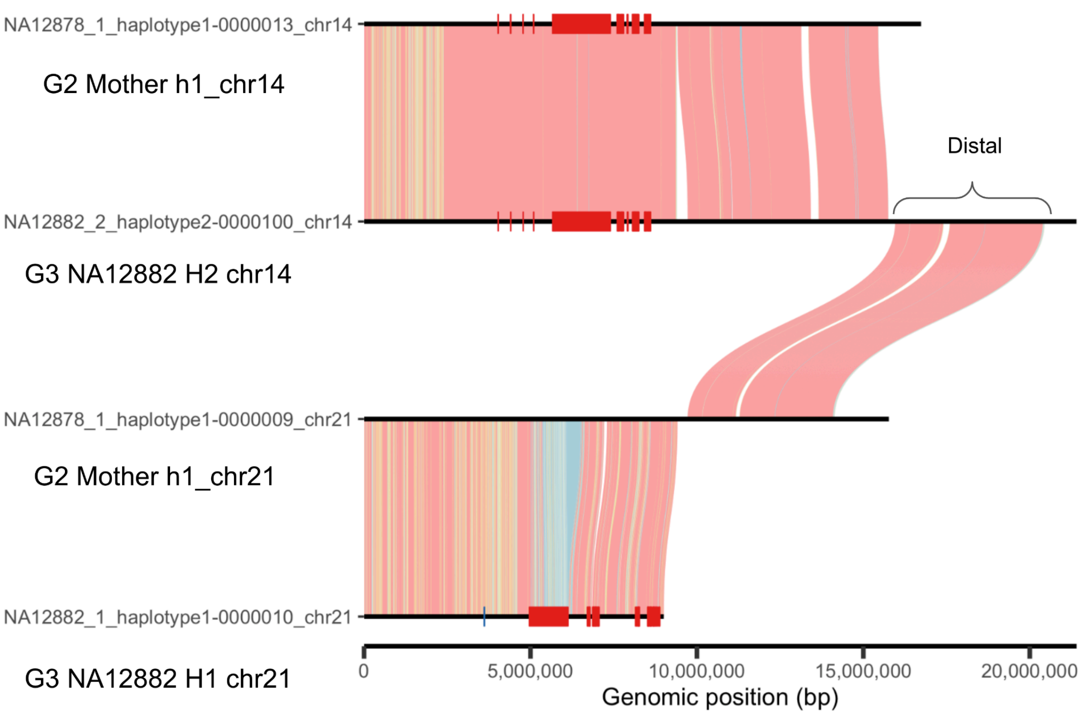


**Figure 17. One potentially incorrect recombination due to distal sequence scaffold error of child.** The distal sequence is incorrectly scaffolded to the child NA12882 chr14 haplotype. This caused an incorrect distal sequence alignment between NA12882 chr14 and NA12878 chr21.

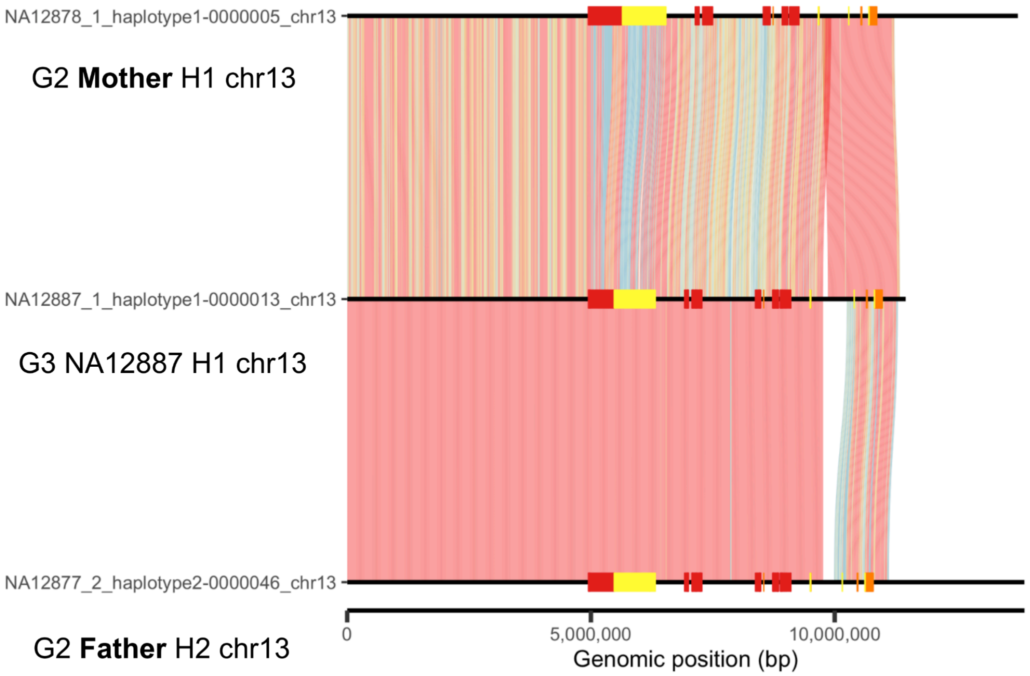


**Figure 18. One incorrect recombination due to haplotype switch error.**

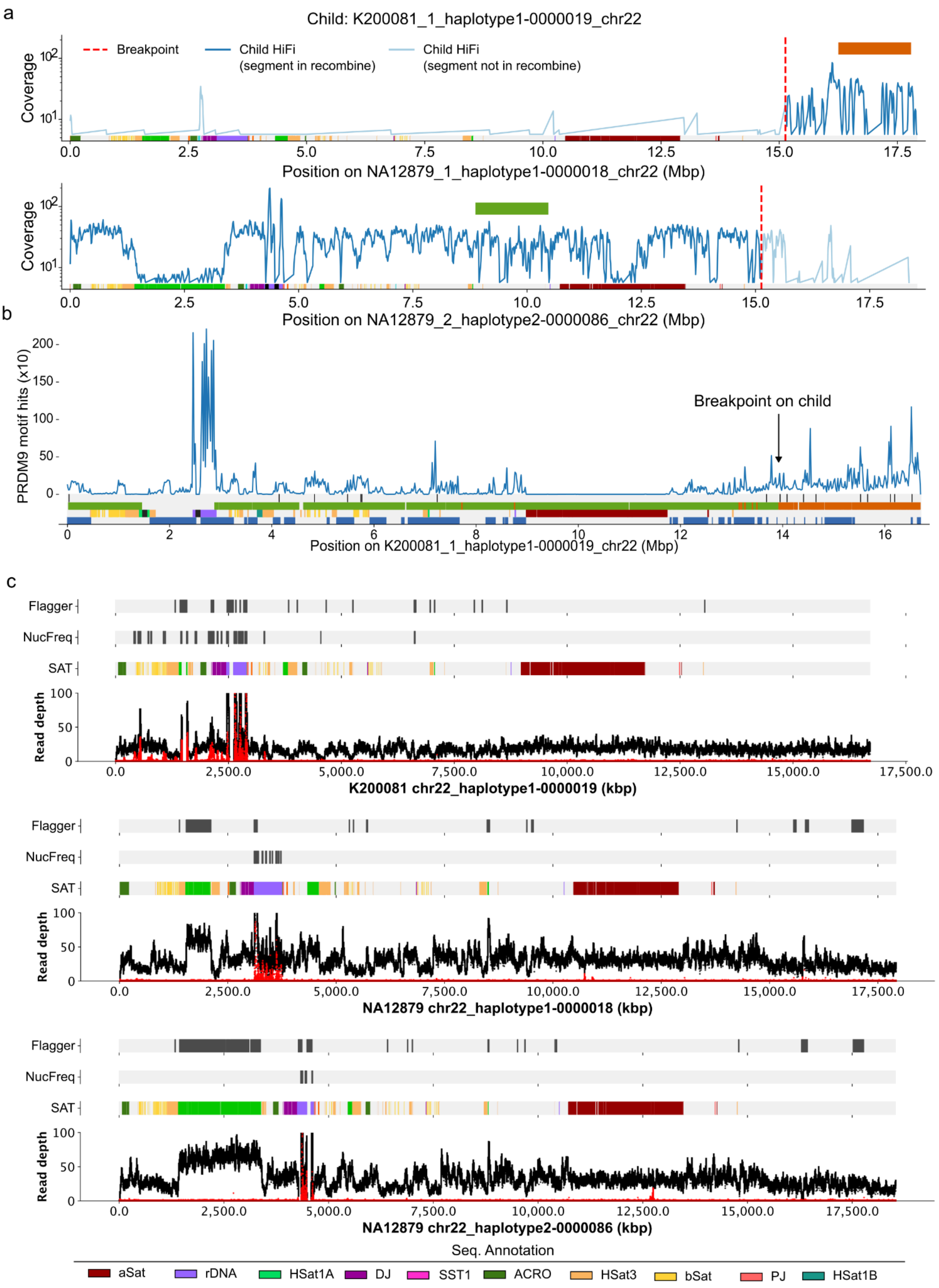

a
Child: K200081_1_haplotype1-0000019_chr22
Breakpoint
Child HiFi (segment in recombine)
Child HiFi (segment not in recombine)
Coverage
Position on NA12879_1_haplotype1-0000018_chr22 (Mbp)
Position on NA12879_2_haplotype2-0000086_chr22 (Mbp)
b
PRDM9 motif hits (x10)
Breakpoint on child
Position on K200081_1_haplotype1-0000019_chr22 (Mbp)
c
Flagger
NucFreq
SAT
Read depth
K200081 chr22_haplotype1-0000019 (kbp)
NA12879 chr22_haplotype1-0000018 (kbp)
NA12879 chr22_haplotype2-0000086 (kbp)
Seq. Annotation
aSat
rDNA
HSat1A
DJ
SST1
ACRO
HSat3
bSat
PJ
HSat1B

**Figure 19. The chr22 recombination on G4 sample 200081.** a) HiFi read coverage shows the alignment of the child's reads to parental haplotypes. b) The breakpoint characterization on the child recombinant. The tracks from top to bottom show PRDM9 motif hits, best PRDM9 motif hits, 10 kbp sliding aligner, satellite annotation, and segmental duplication. c) Each panel shows the quality of assemblies involved in the recombination. For each panel, from top to bottom the tracks show the Flagger- and NucFreq-identified assembly errors, satellite sequence (SAT), and NucFreq plot. (same legend for Figures 20-37)

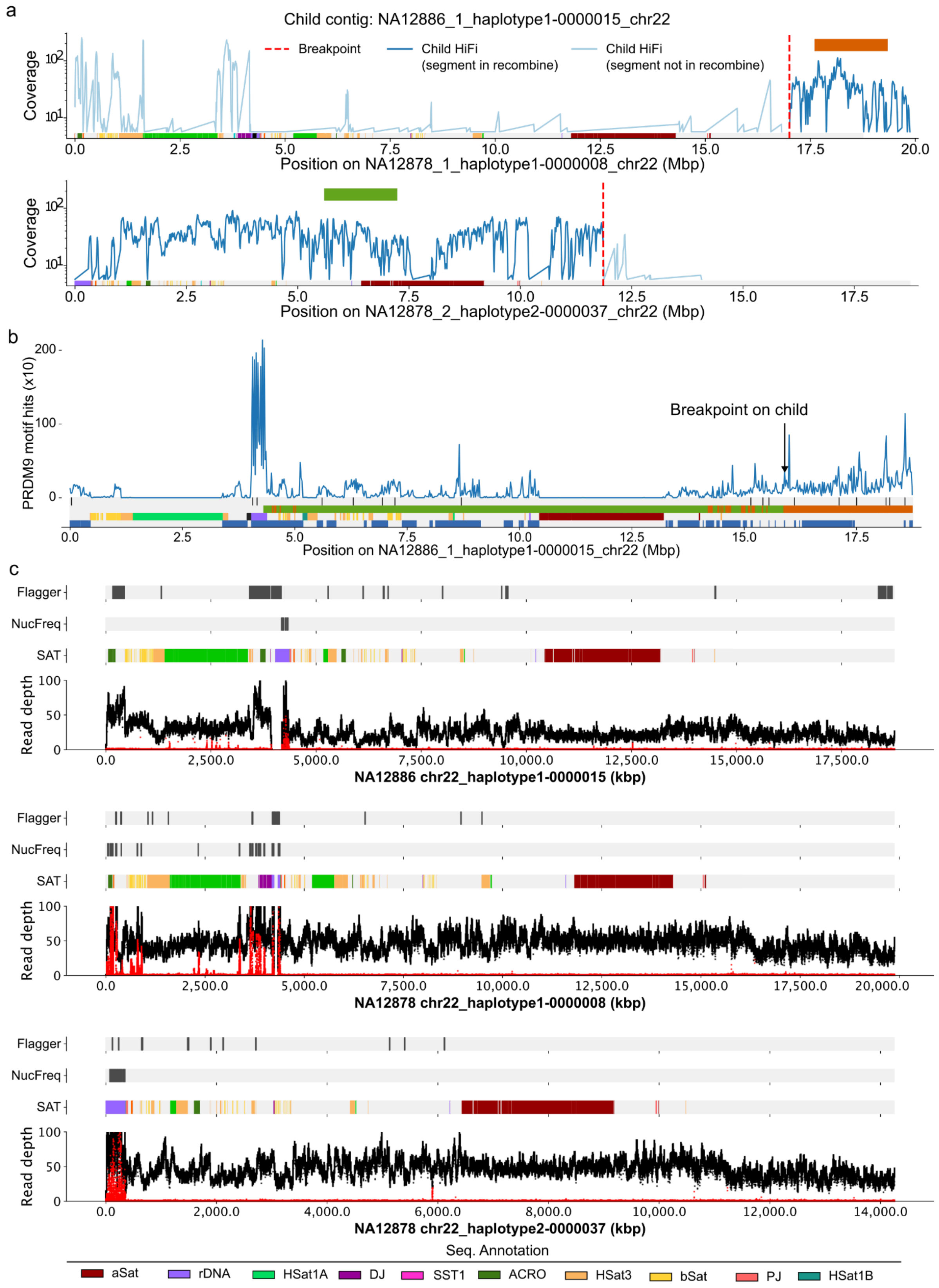


**Figure 20. The chr22 recombination on G3 sample NA12886.** (see Figure 19 legend).

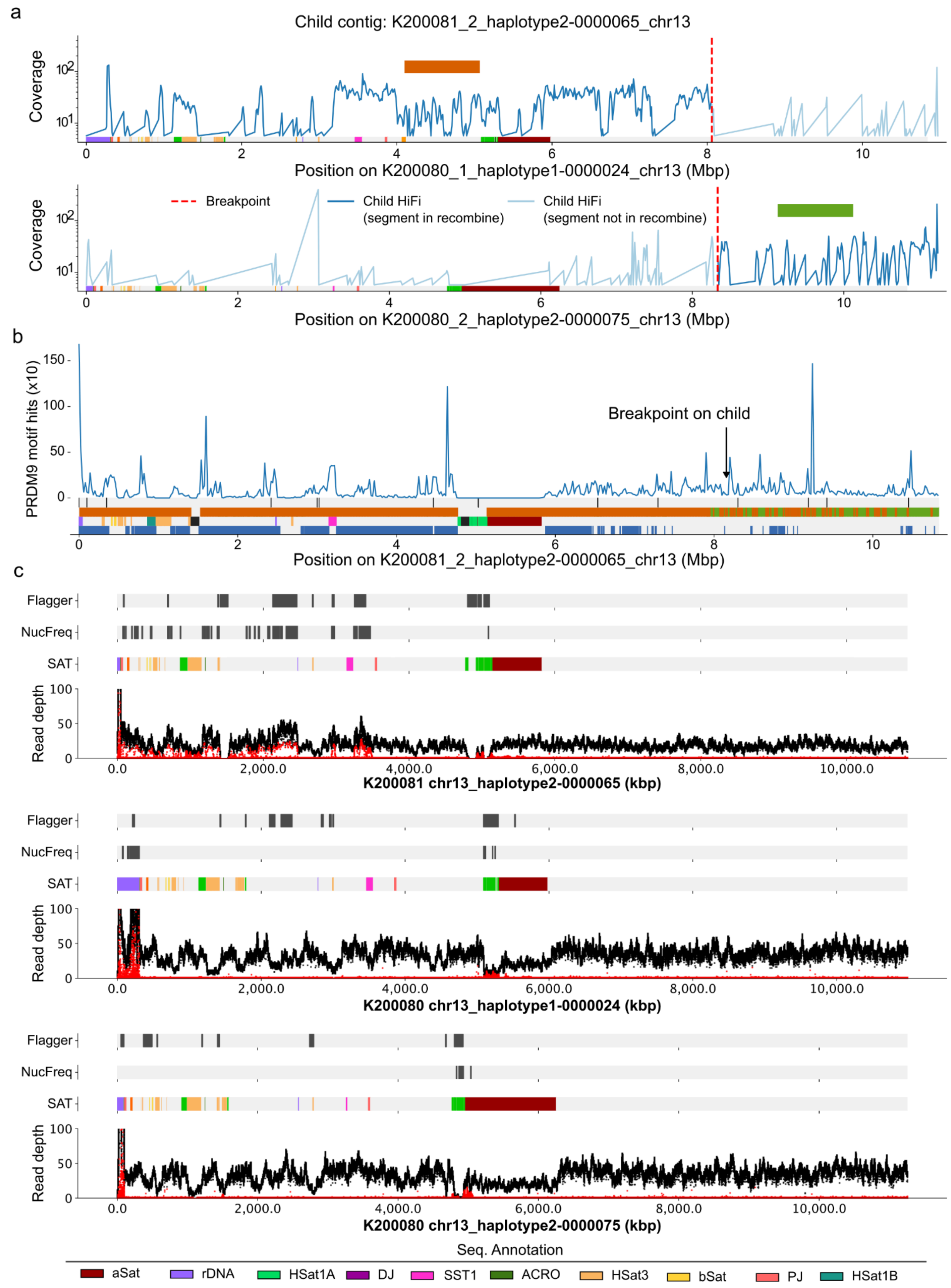


**Figure 21. The chr13 recombination on G4 sample 200081.** (see Figure 19 legend)

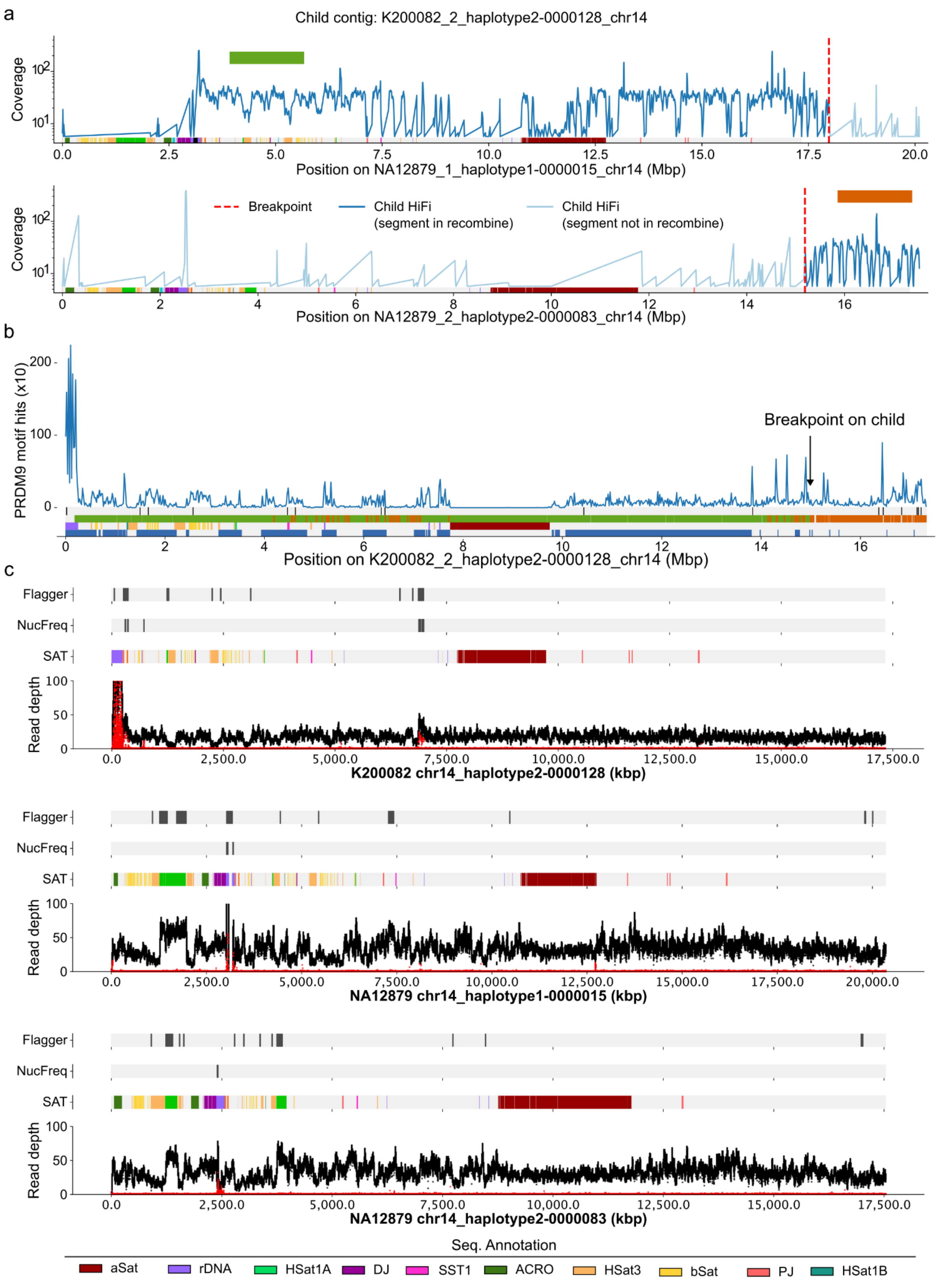


**Figure 22. The chr14 recombination on G4 sample 200082**. (see Figure 19 legend)

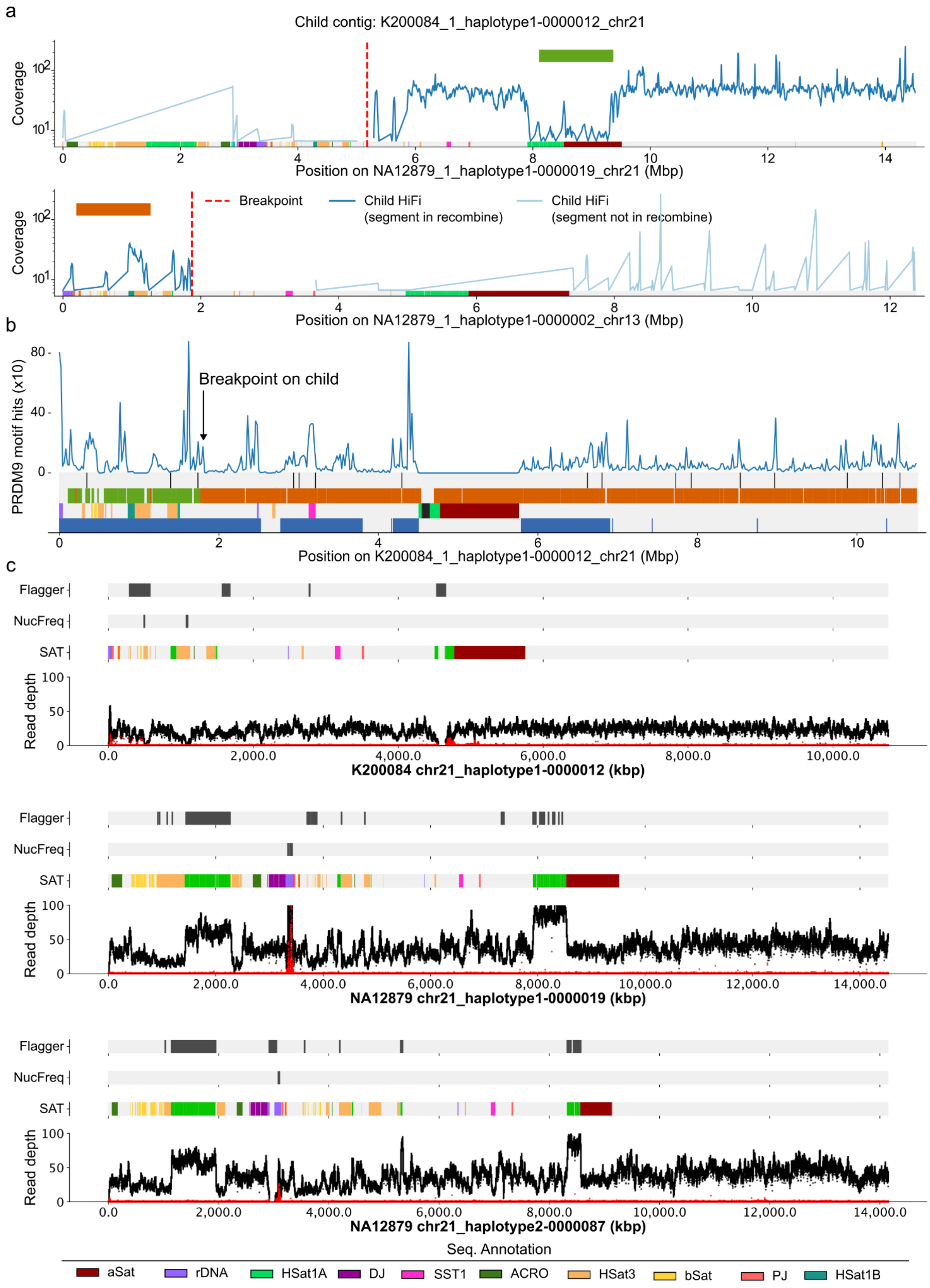


**Figure 23. The chr21 recombination on G4 sample 200084.** (see Figure 19 legend)

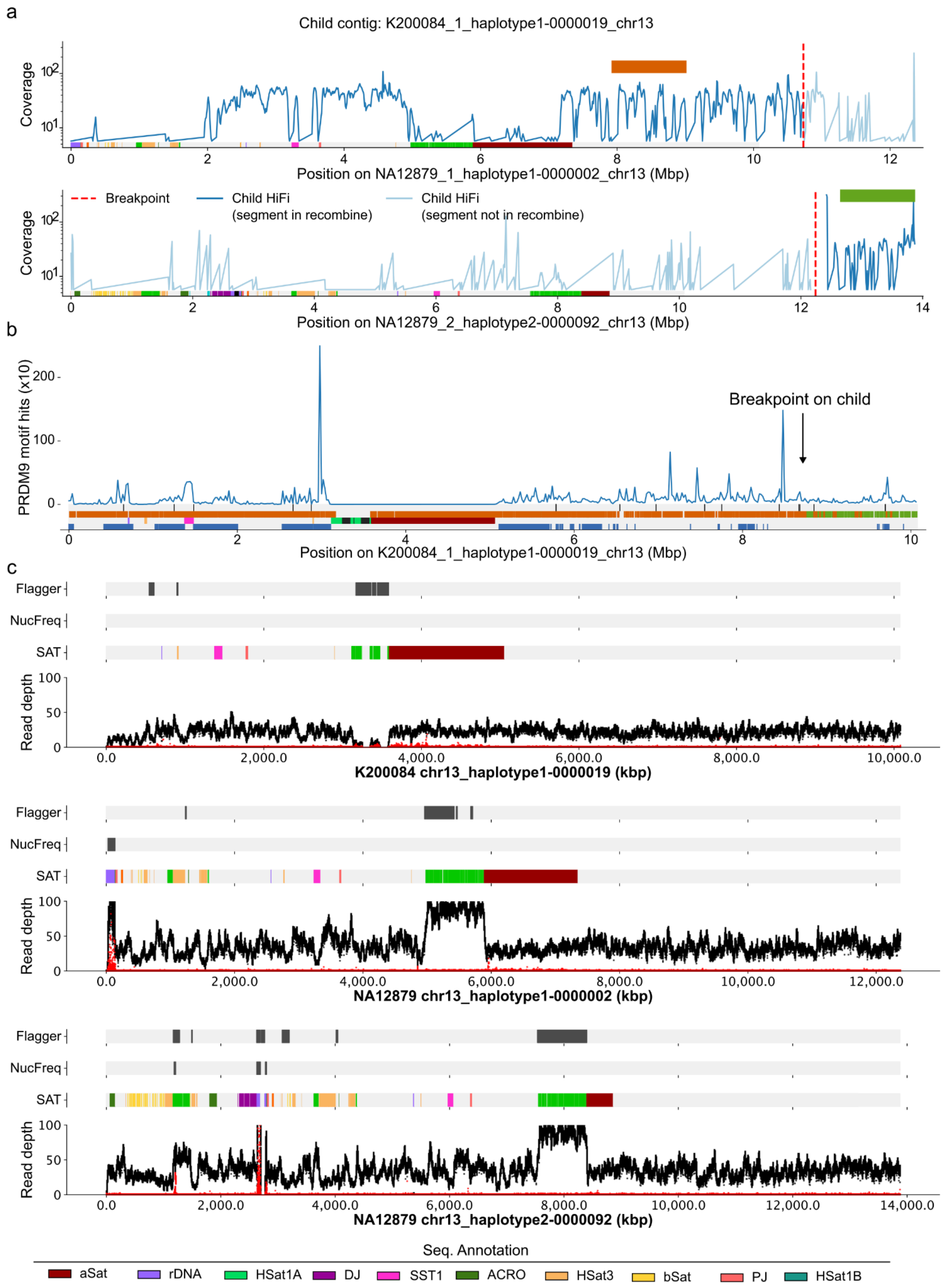


**Figure 24. The chr13 recombination on G4 sample 200084**. (see Figure 19 legend)

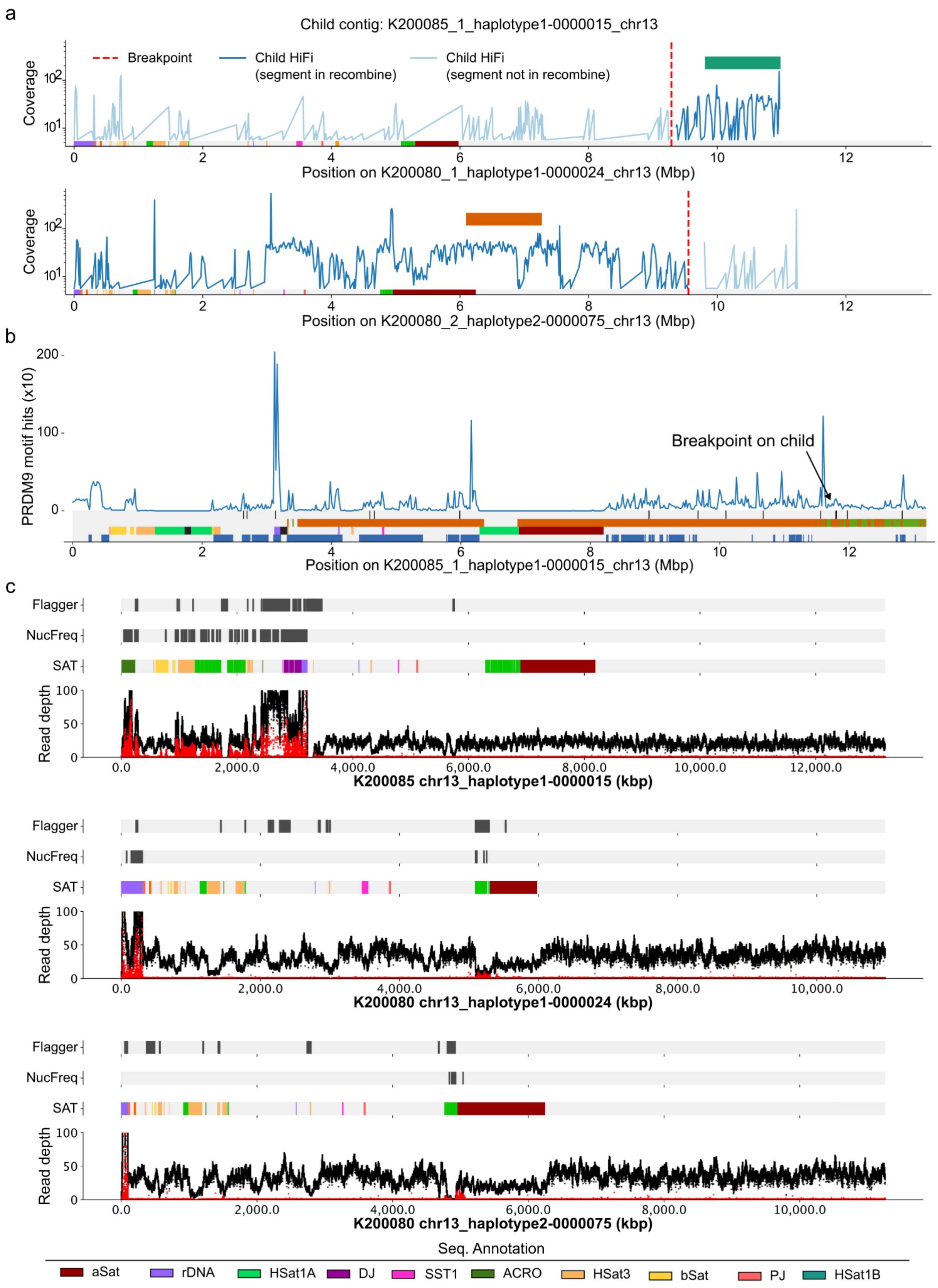


**Figure 25. The chr13 recombination on G4 sample 200085.** (see Figure 19 legend)

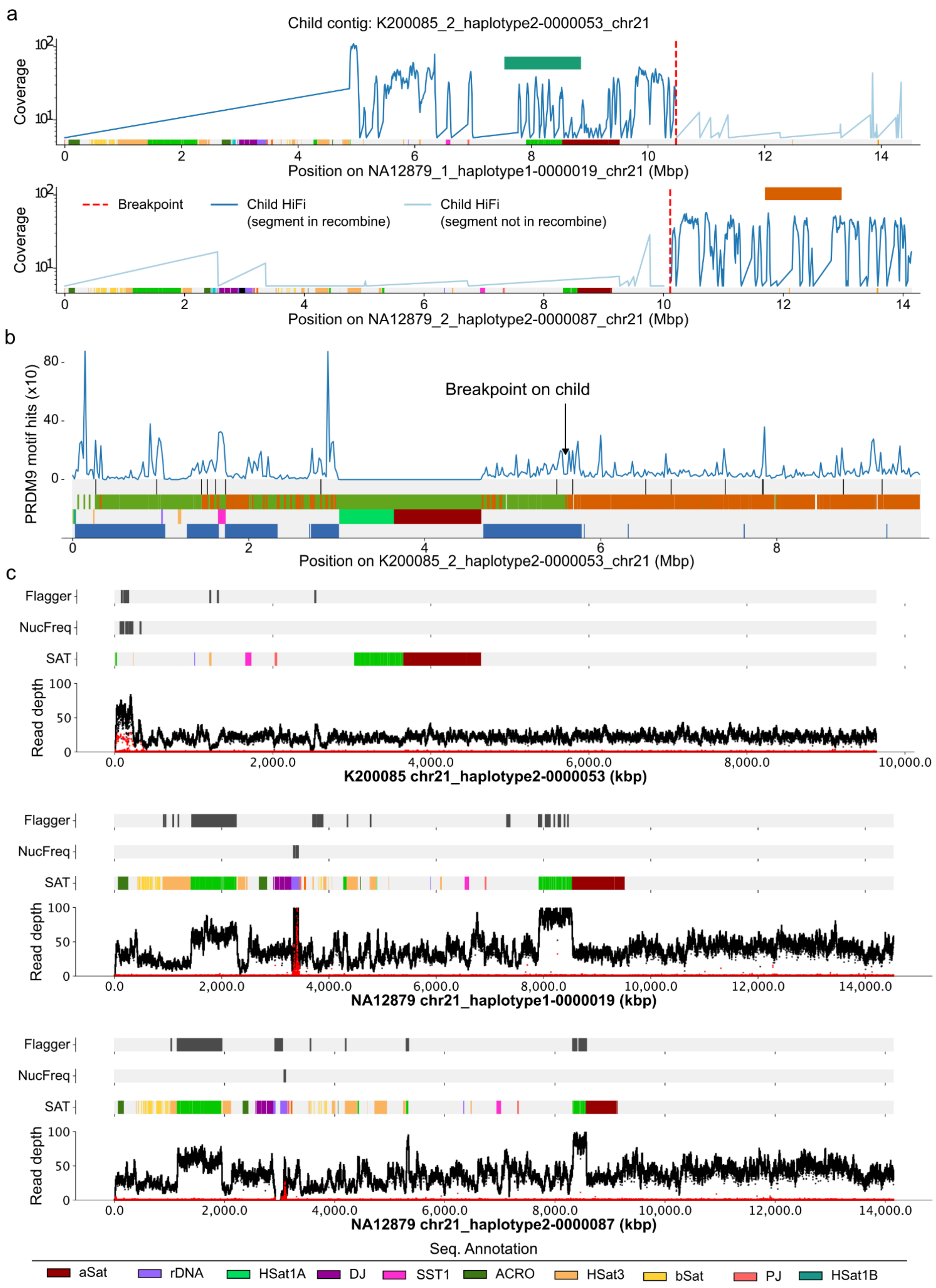


**Figure 26. The chr21 recombination on G4 sample 200085.** (see Figure 19 legend)

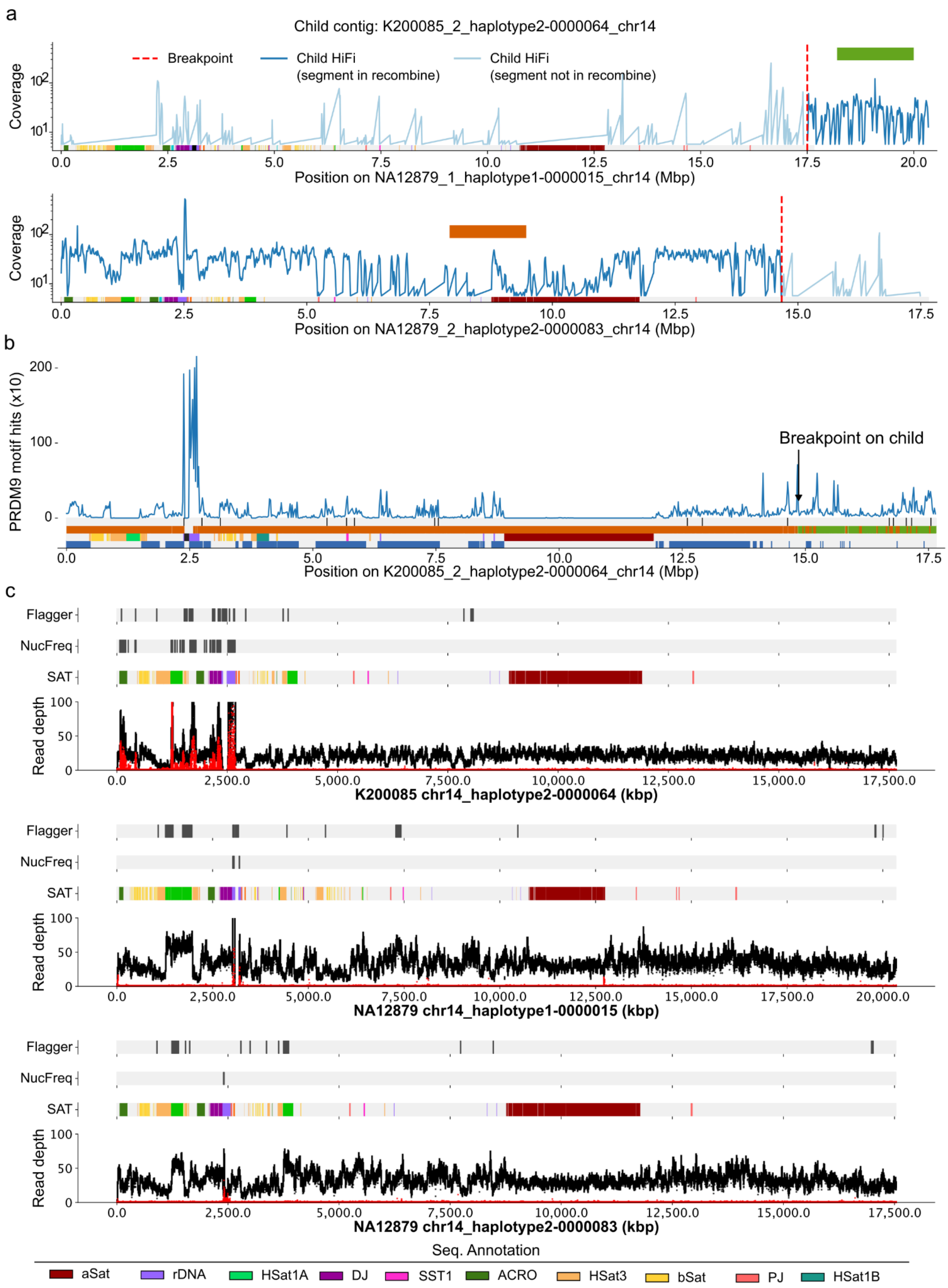


**Figure 27. The chr14 recombination on G4 sample 200085.** (see Figure 19 legend)

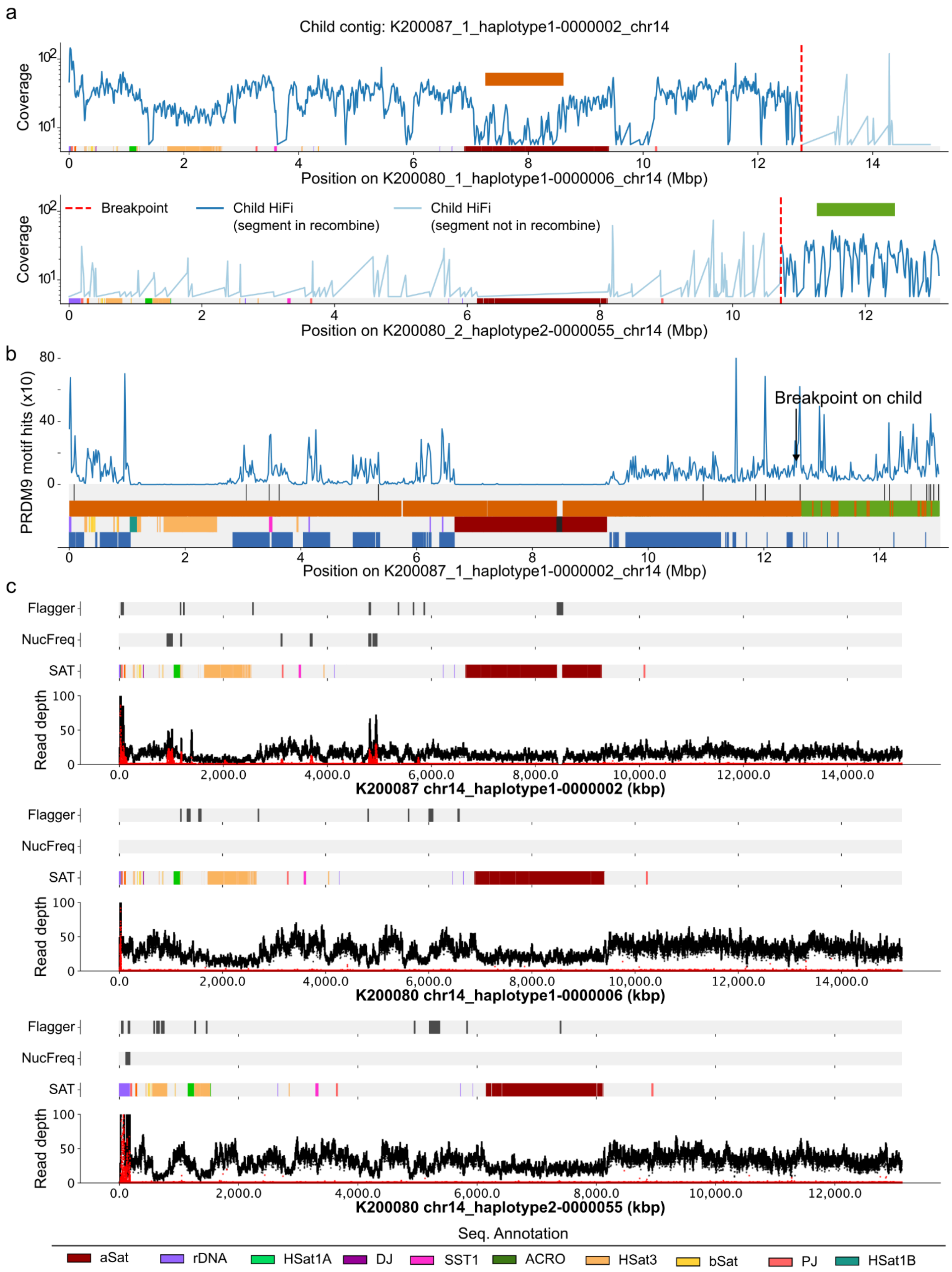


**Figure 28. The chr14 recombination on G4 sample 200087.** (see Figure 19 legend)

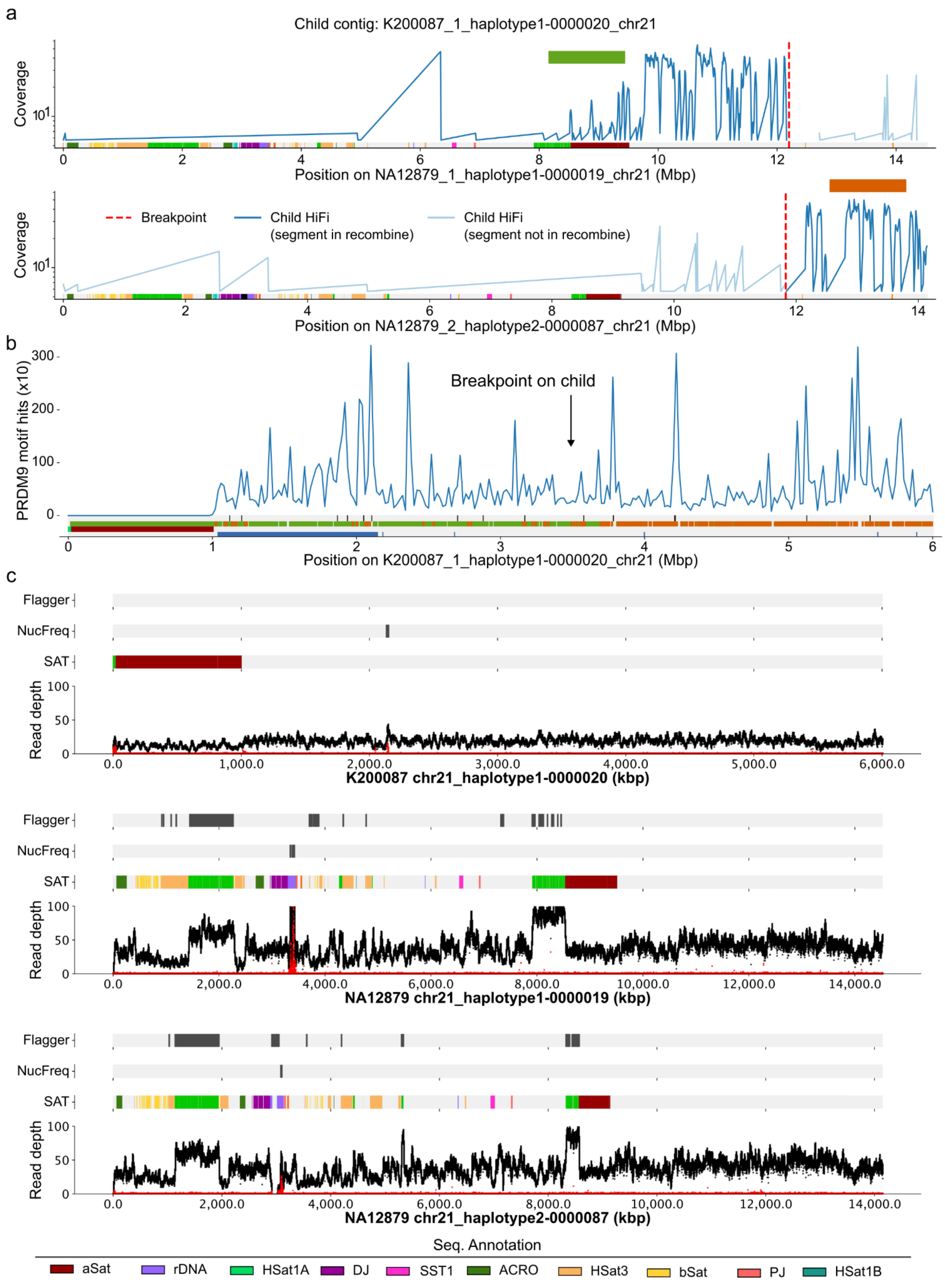


**Figure 29. The chr21 recombination on G4 sample 200087**. (see Figure 19 legend)

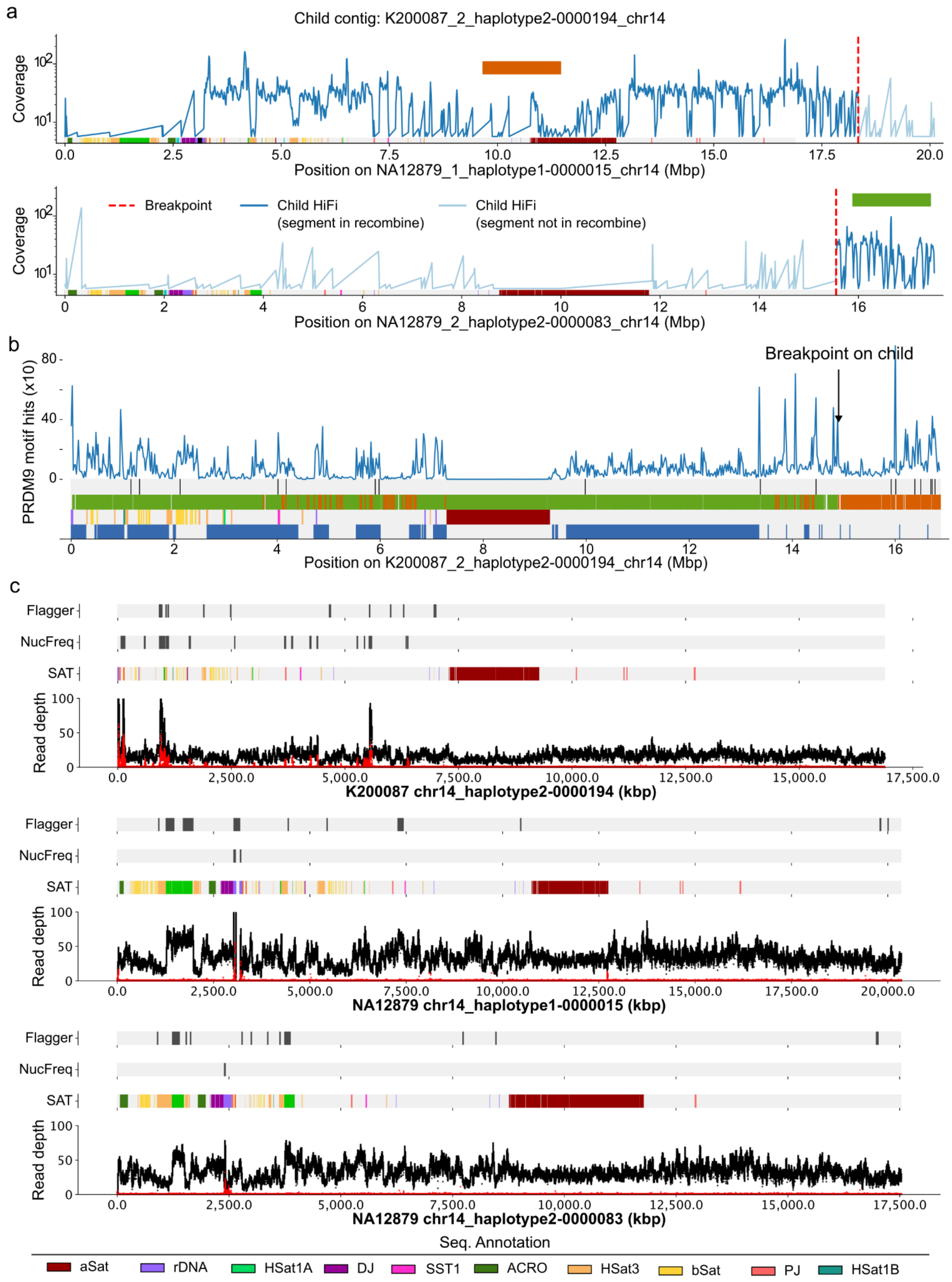


**Figure 30. The chr14 recombination on G4 sample 200087.** (see Figure 19 legend)

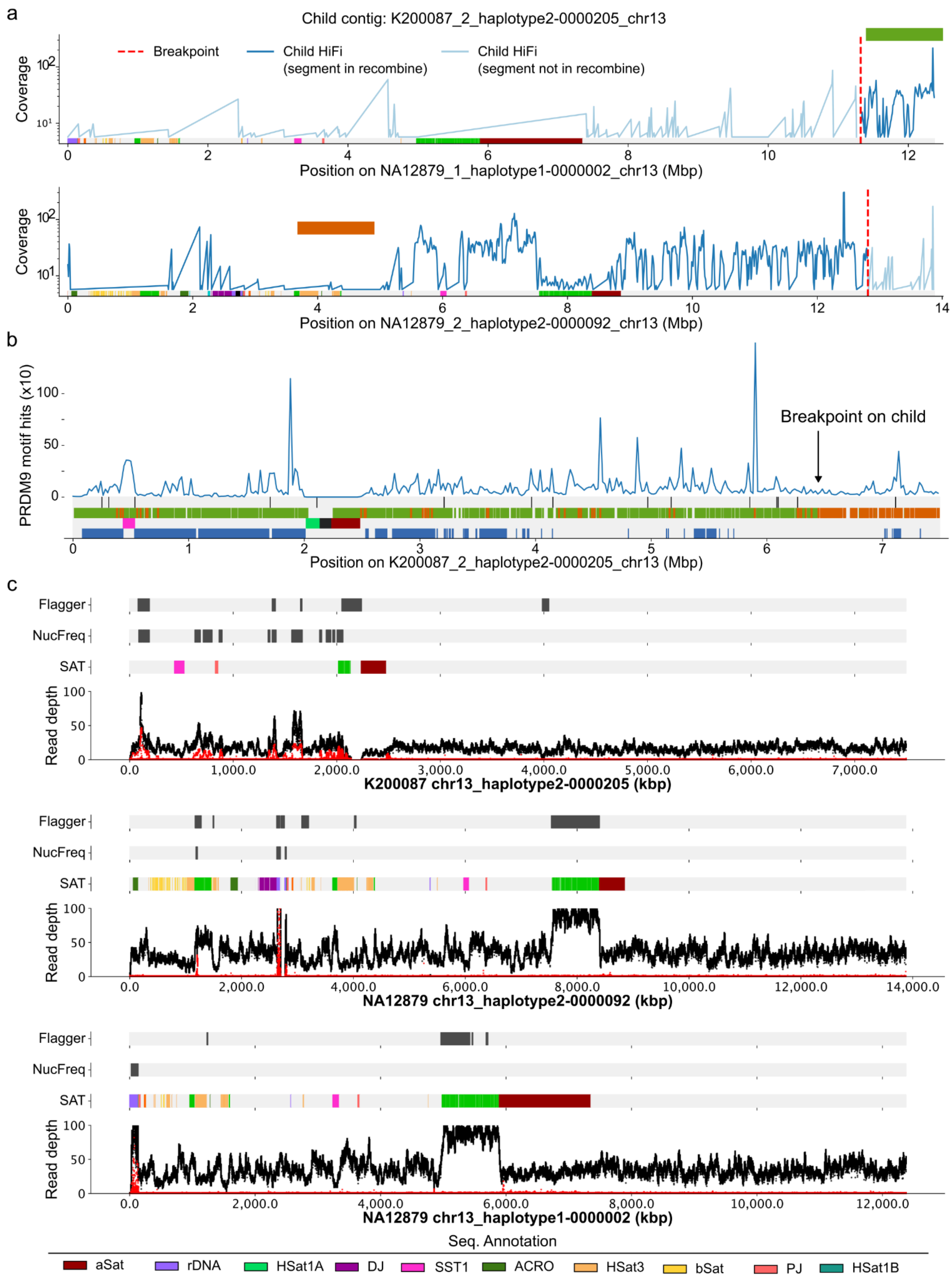


**Figure 31. The chr13 recombination on G4 sample 200087.** (see Figure 19 legend)

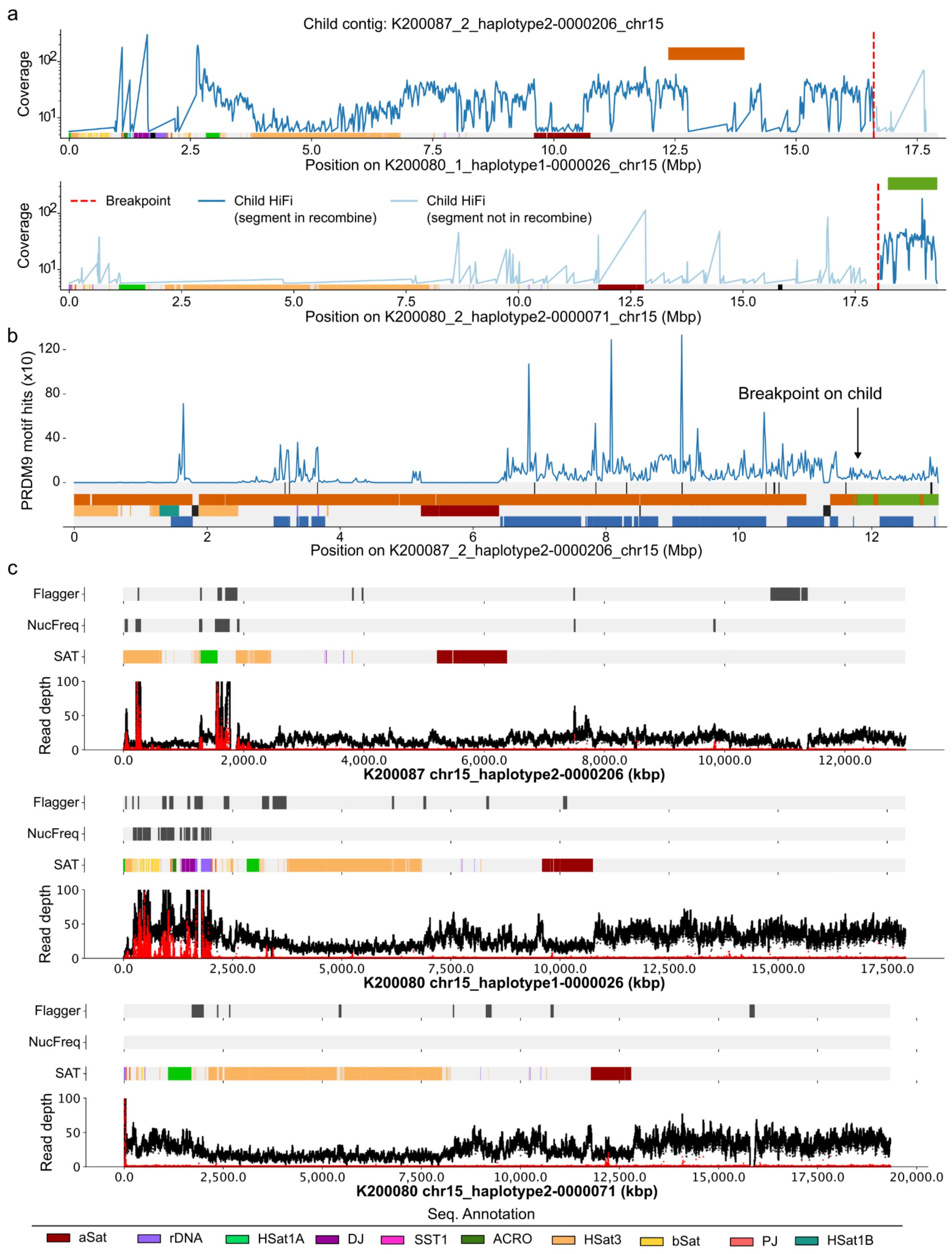


**Figure 32. The chr15 recombination on G4 sample 200087.** (see Figure 19 legend)

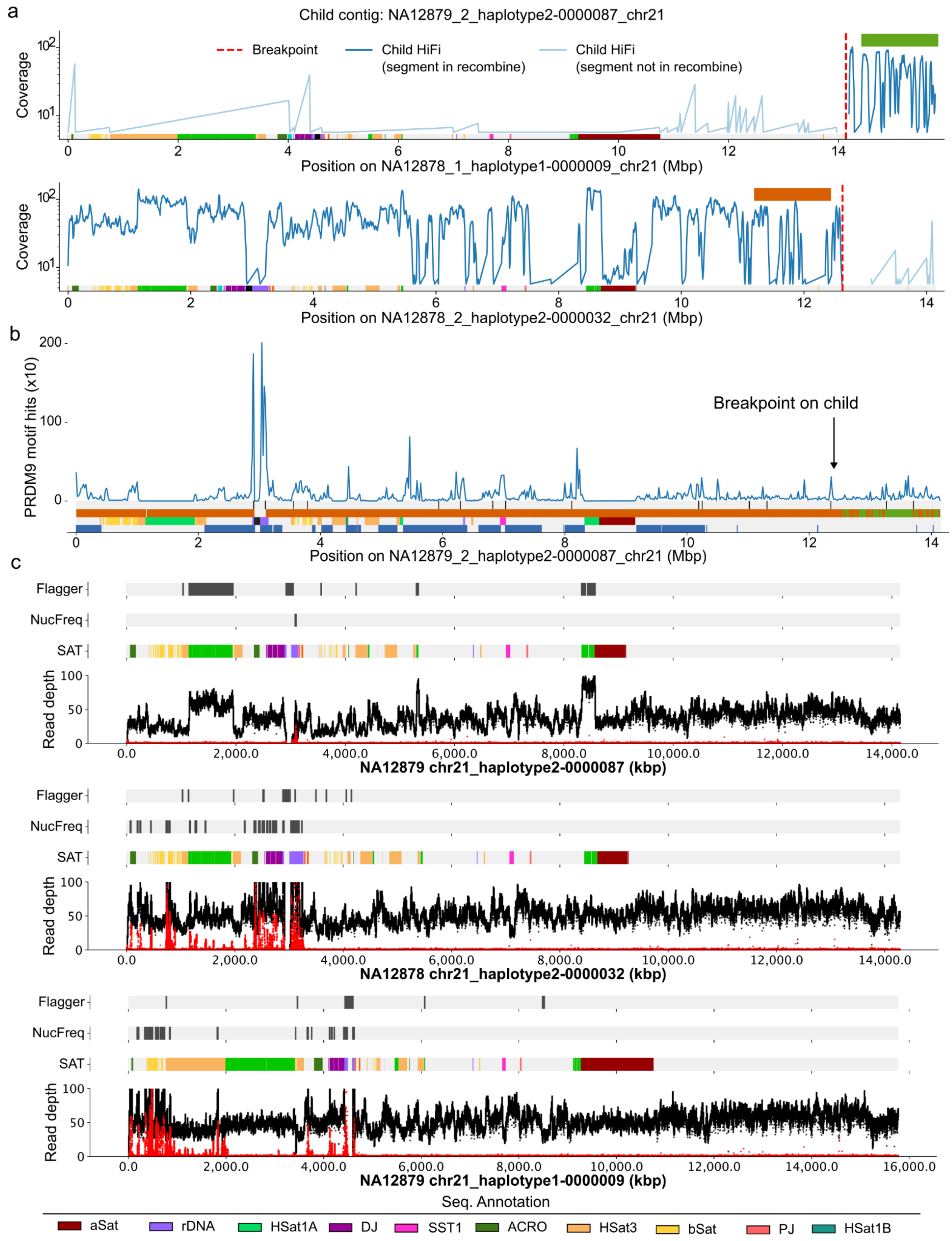


**Figure 33. The chr21 recombination on G3 sample NA12879.** (see Figure 19 legend)

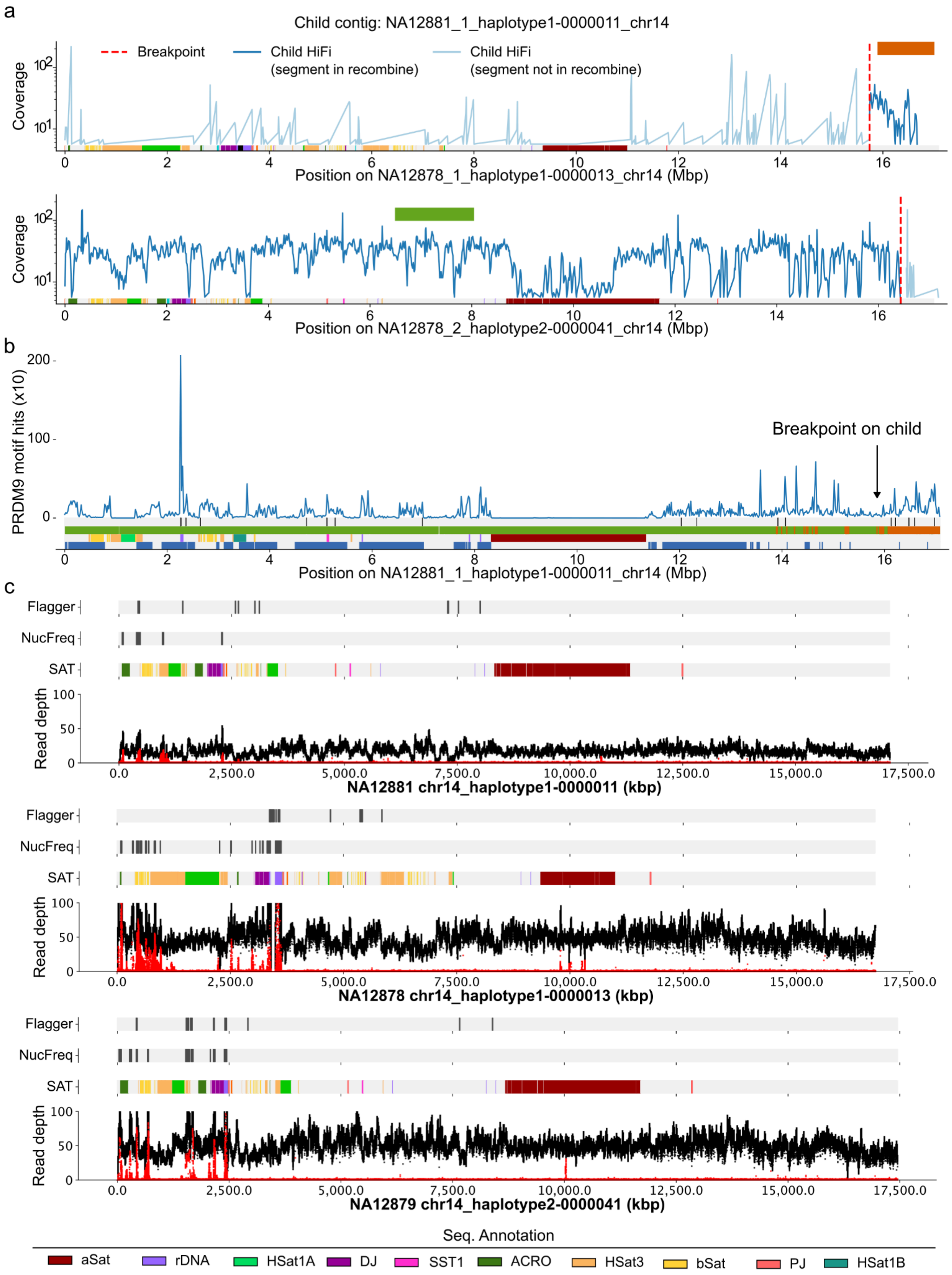


**Figure 34. The chr14 recombination on G3 sample NA12881.** (see Figure 19 legend)

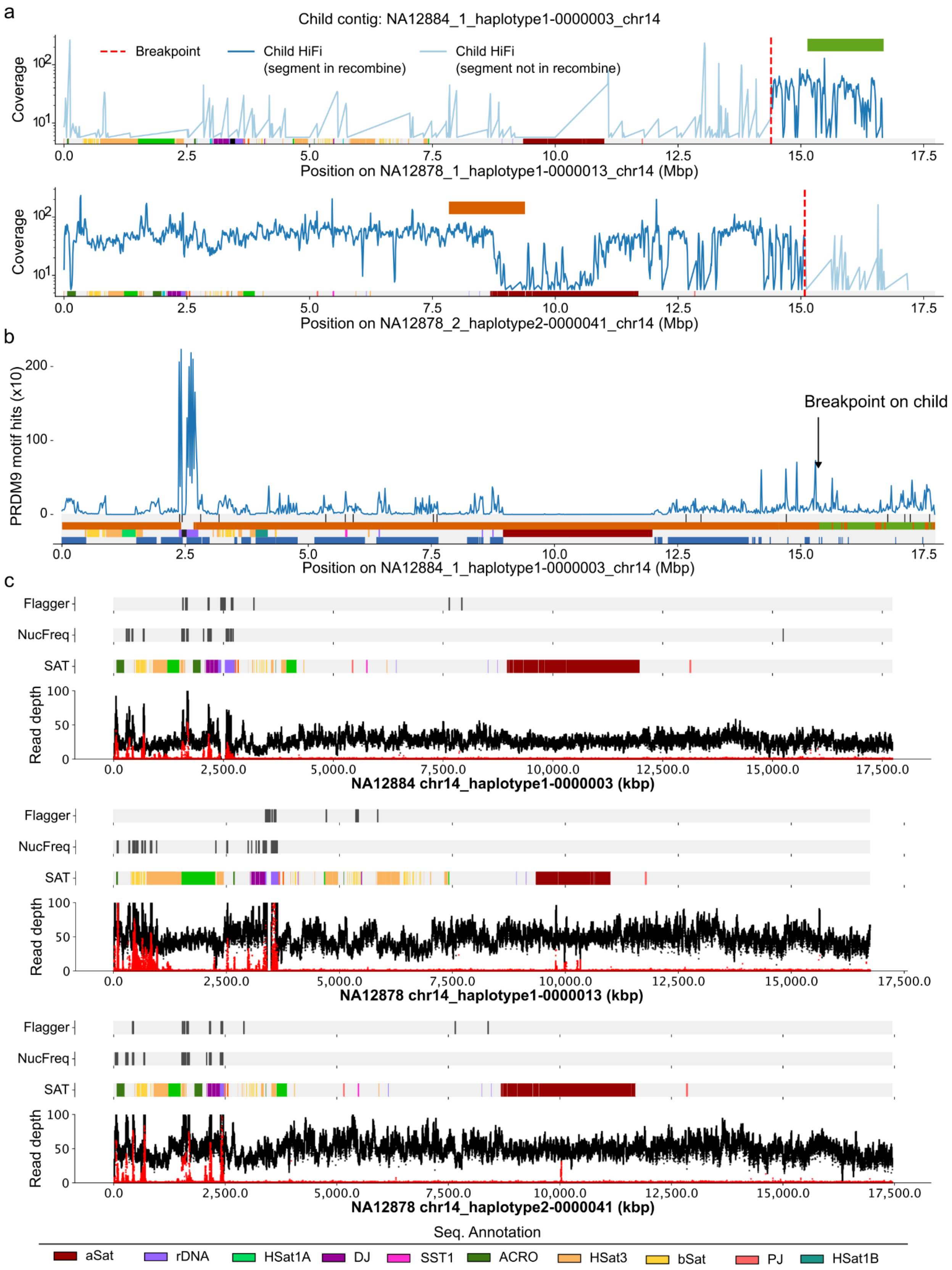


**Figure 35. The chr14 recombination on G3 sample NA12884.** (see Figure 19 legend)

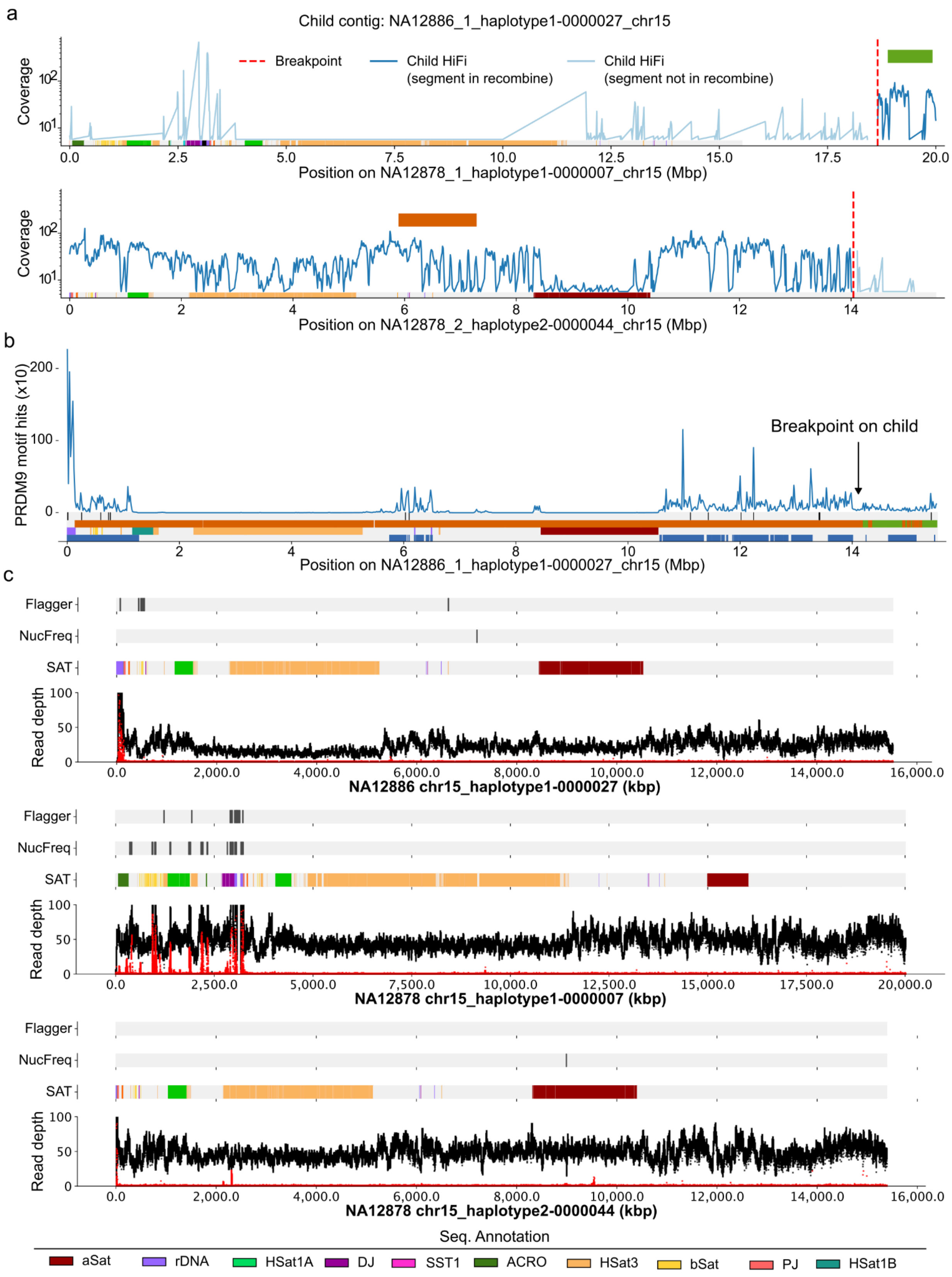


**Figure 36. The chr15 recombination on G3 sample NA12886**. (see Figure 19 legend)

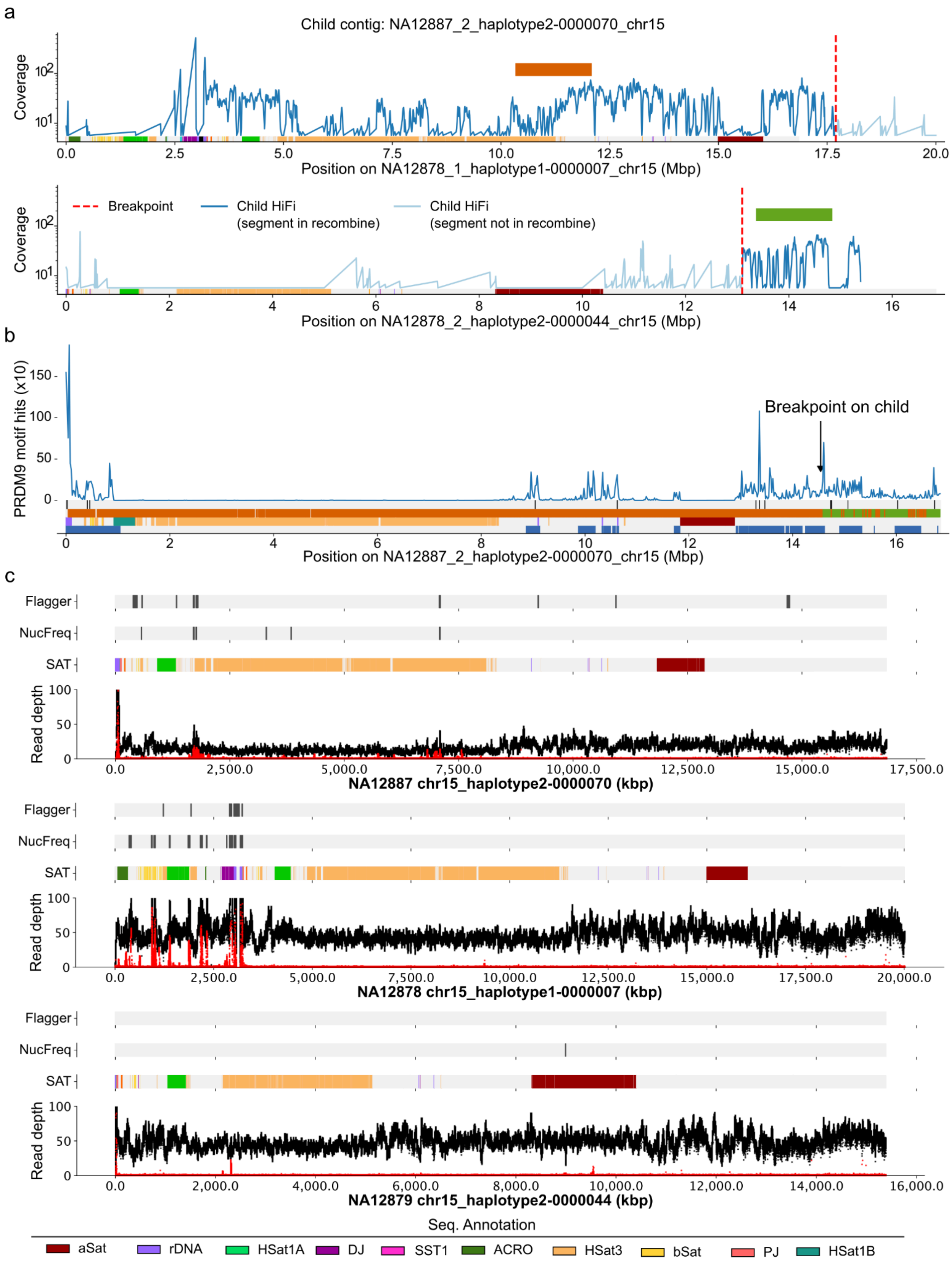


**Figure 37. The chr15 recombination on G3 sample NA12887.** (see Figure 19 legend)

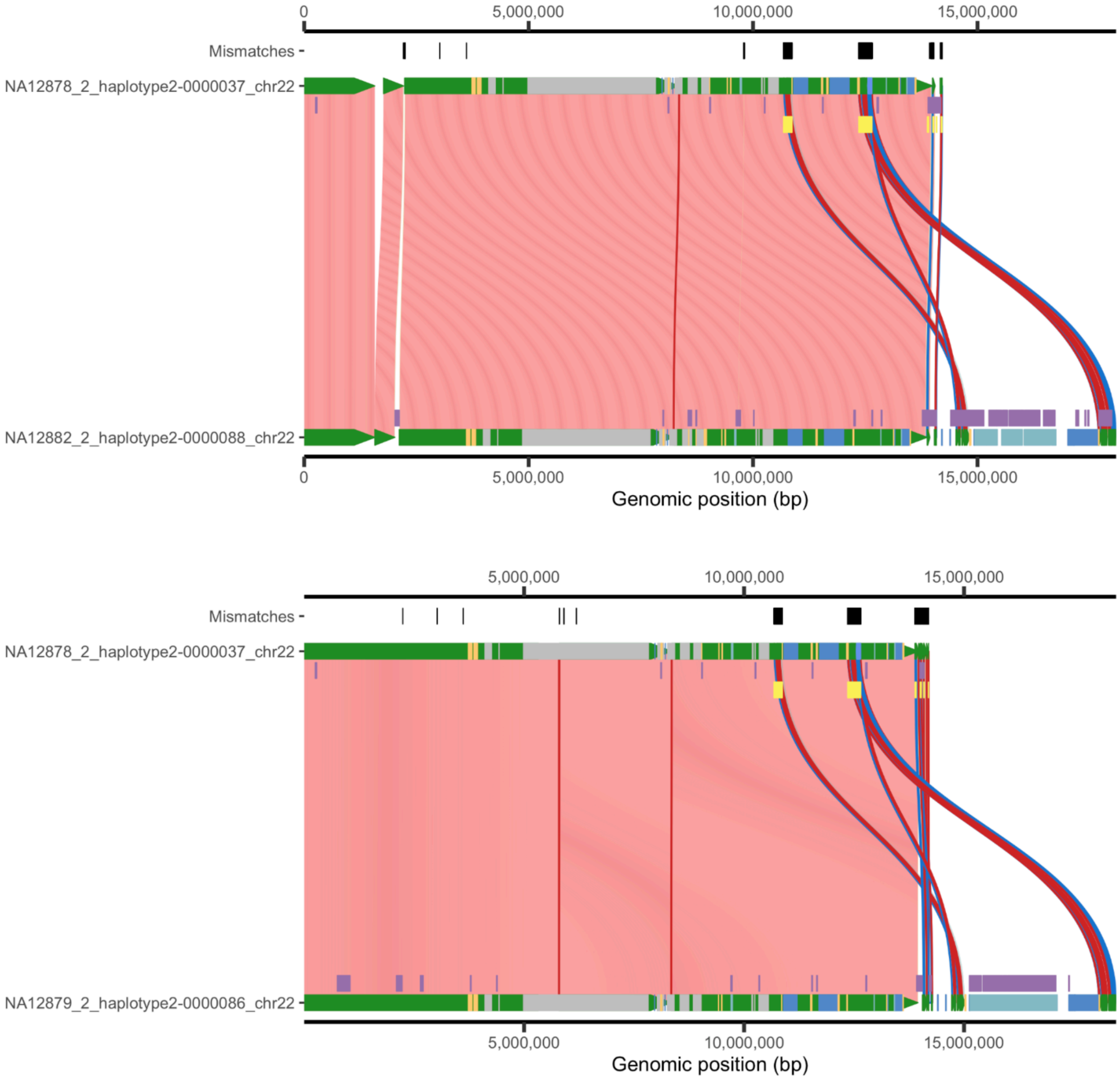


**Figure 38. Examples of misaligned distal sequences.** The misalignment of distal sequence from the child's haplotype (bottom) to parental haplotype (top) that missed the distal sequence in the assembly. The heatmap indicates the sequence identity (red > 99.9% identity). The blue and red lines in the figure between haplotypes represent insertion and deletion, respectively.

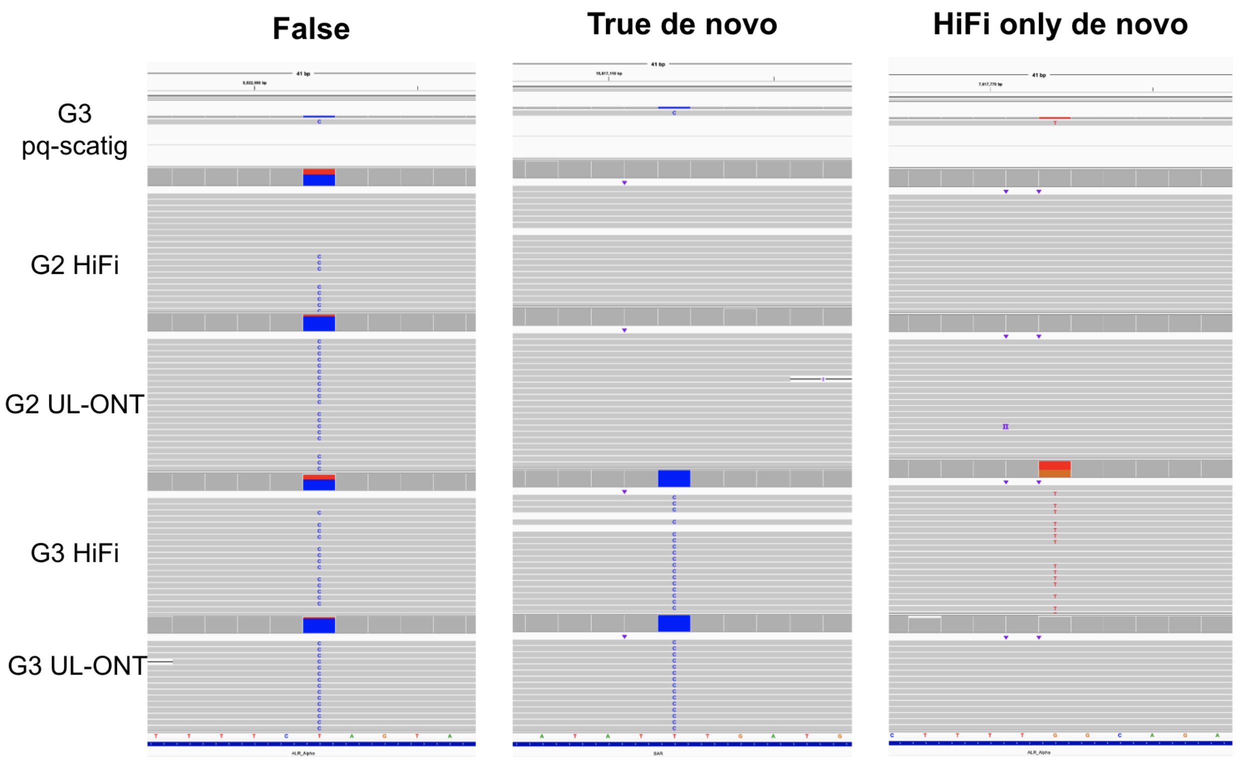


**Figure 39. Examples of validating *de novo* single-nucleotide variants (SNVs) detected from child to parent haplotype alignment.** The five tracks show the alignments to the reference parental haplotype. The variants are called from the G3 pq-scatig and further validated with HiFi and UL-ONT reads. The middle column is an example of the final true *de novo* SNV (variant is missed in parental reads and only shown in child reads). The HiFi-only *de novo* SNV indicates the variant is only supported by the child's HiFi reads.

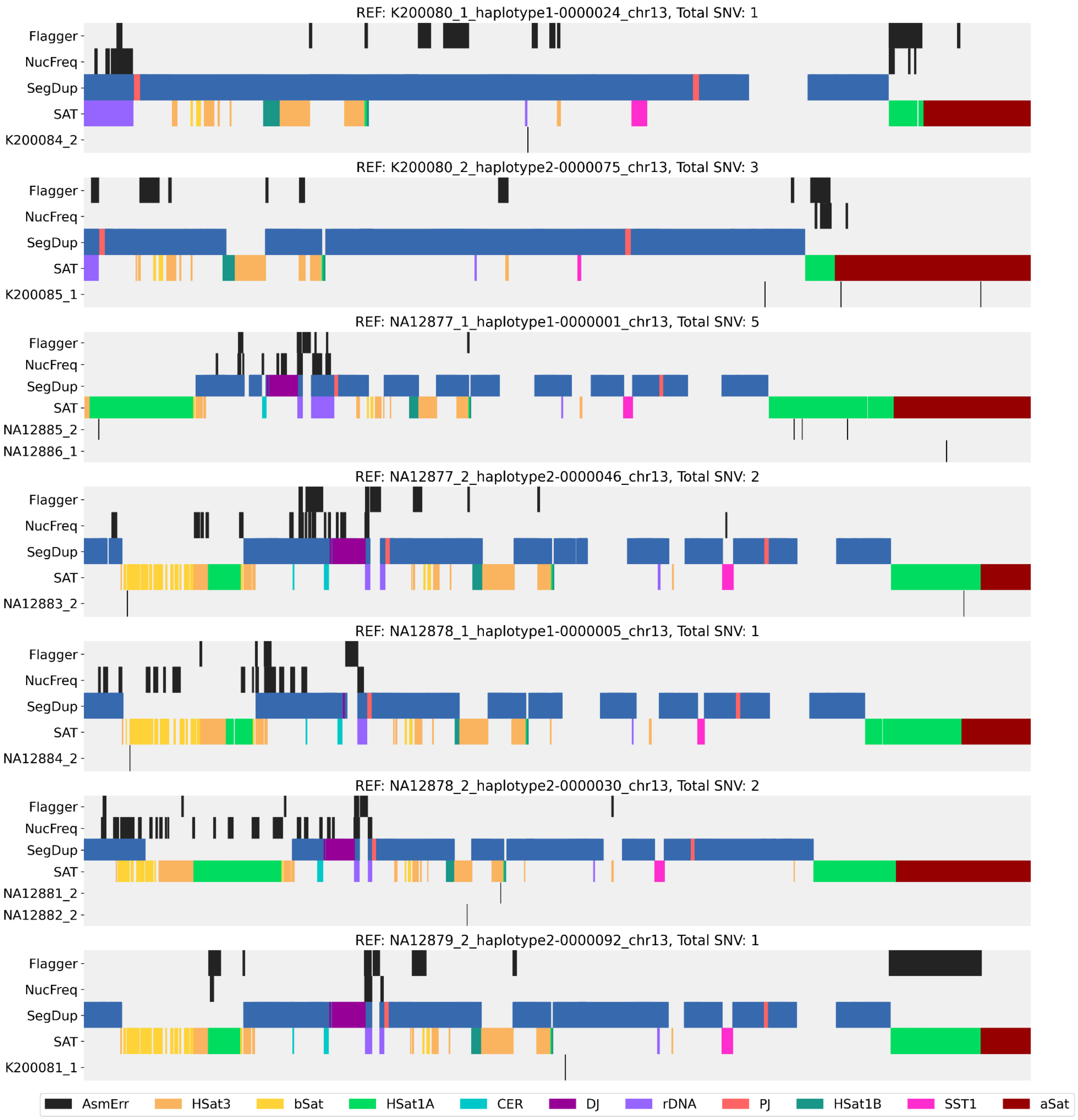


**Figure 40. Locations of chr13 *de novo* SNVs detected from each offspring against the reference parental haplotypes.** The tracks show the assembly error (Flagger and NucFreq), segmental duplication (SegDup), and satellite (SAT) annotations for the reference parental haplotype (REF). (same legend for Figures 41-44)

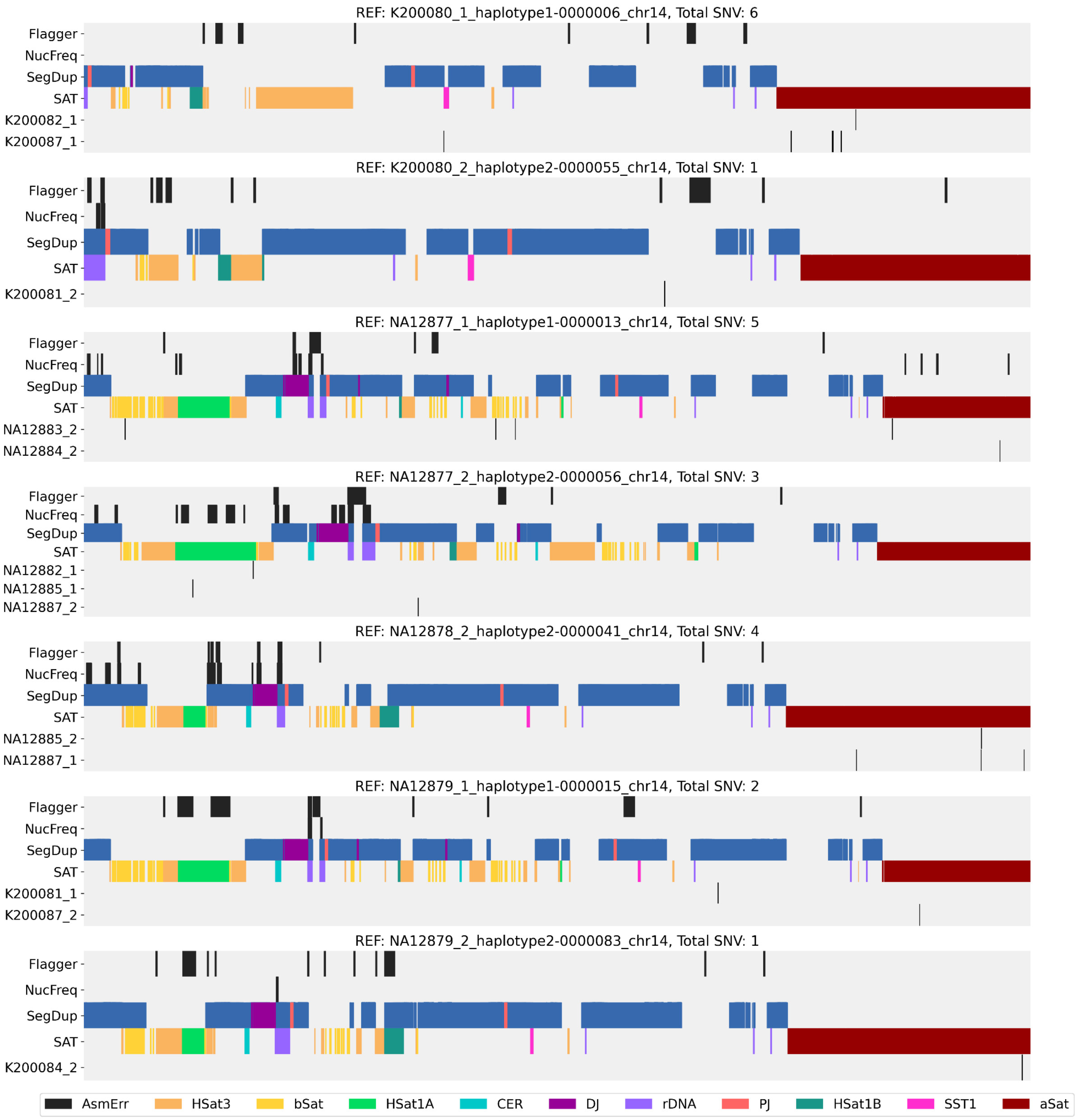


**Figure 41. Locations of chr14 *de novo* SNVs detected from each offspring against the reference parental haplotypes.** (see Figure 40 legend)

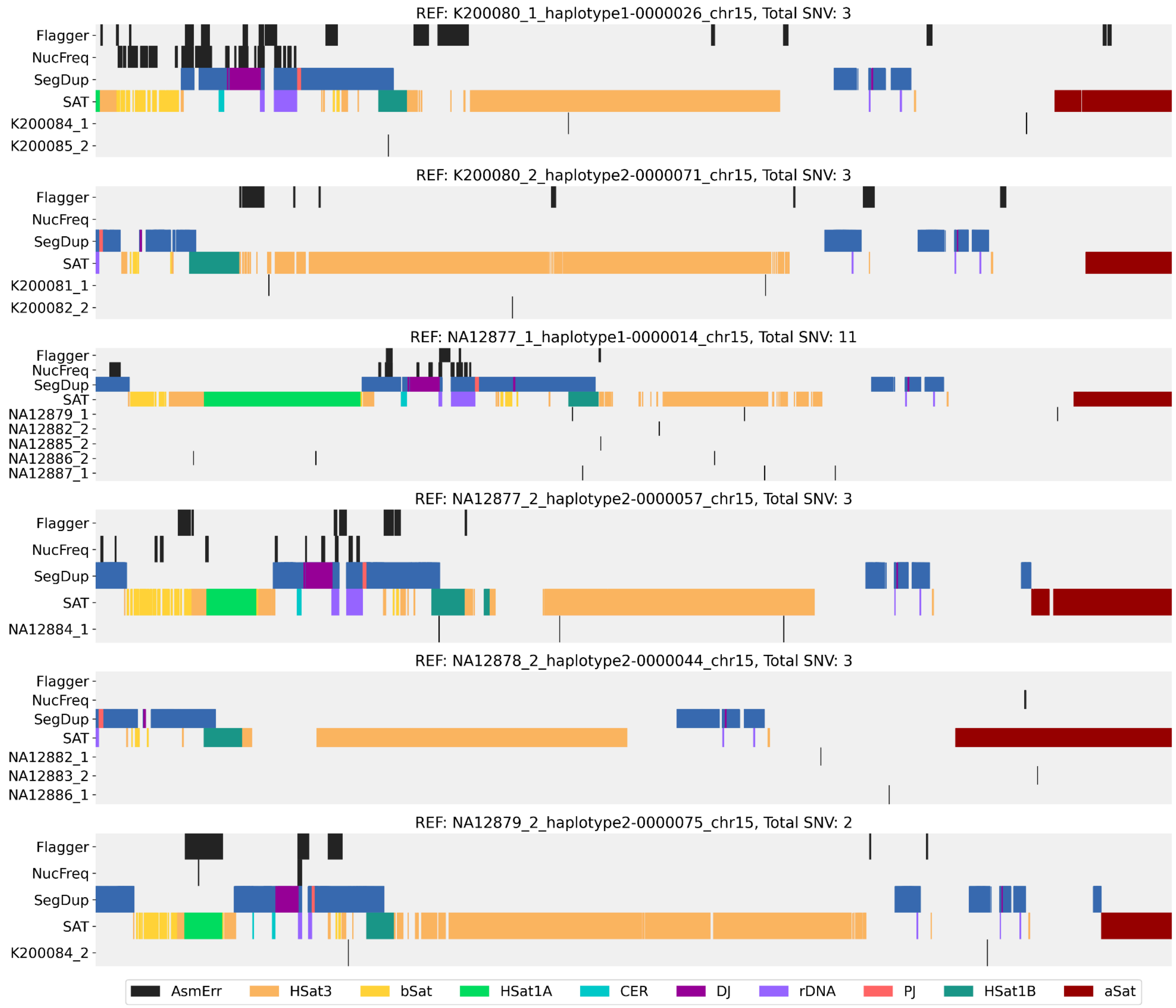


**Figure 42. Locations of chr15 *de novo* SNVs detected from each offspring against the reference parental haplotypes.** (see Figure 40 legend)

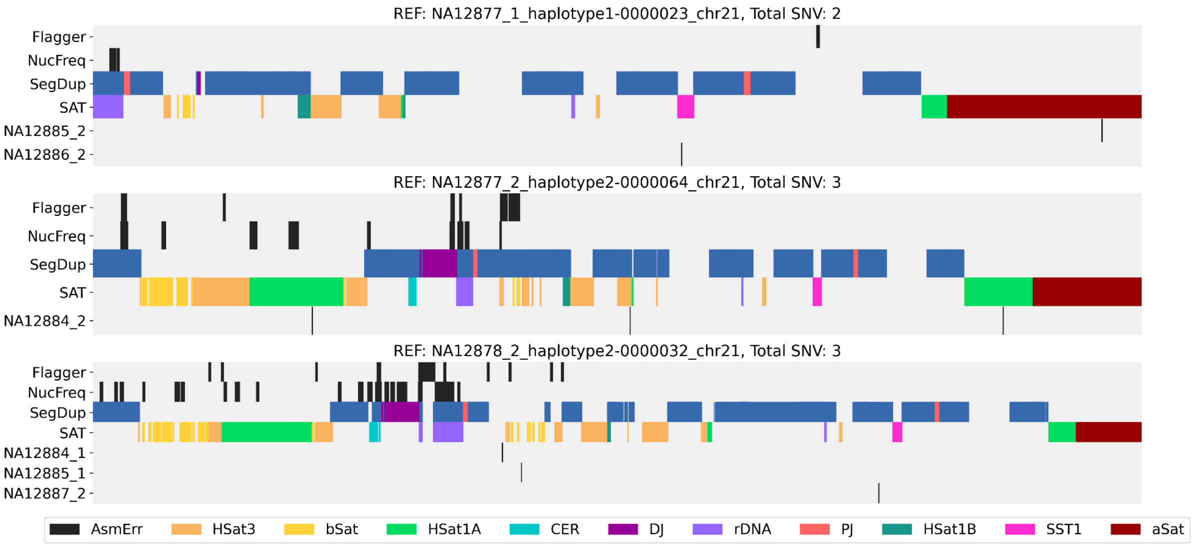


**Figure 43. Locations of chr21 *de novo* SNVs detected from each offspring against the reference parental haplotypes.** (see Figure 40 legend)

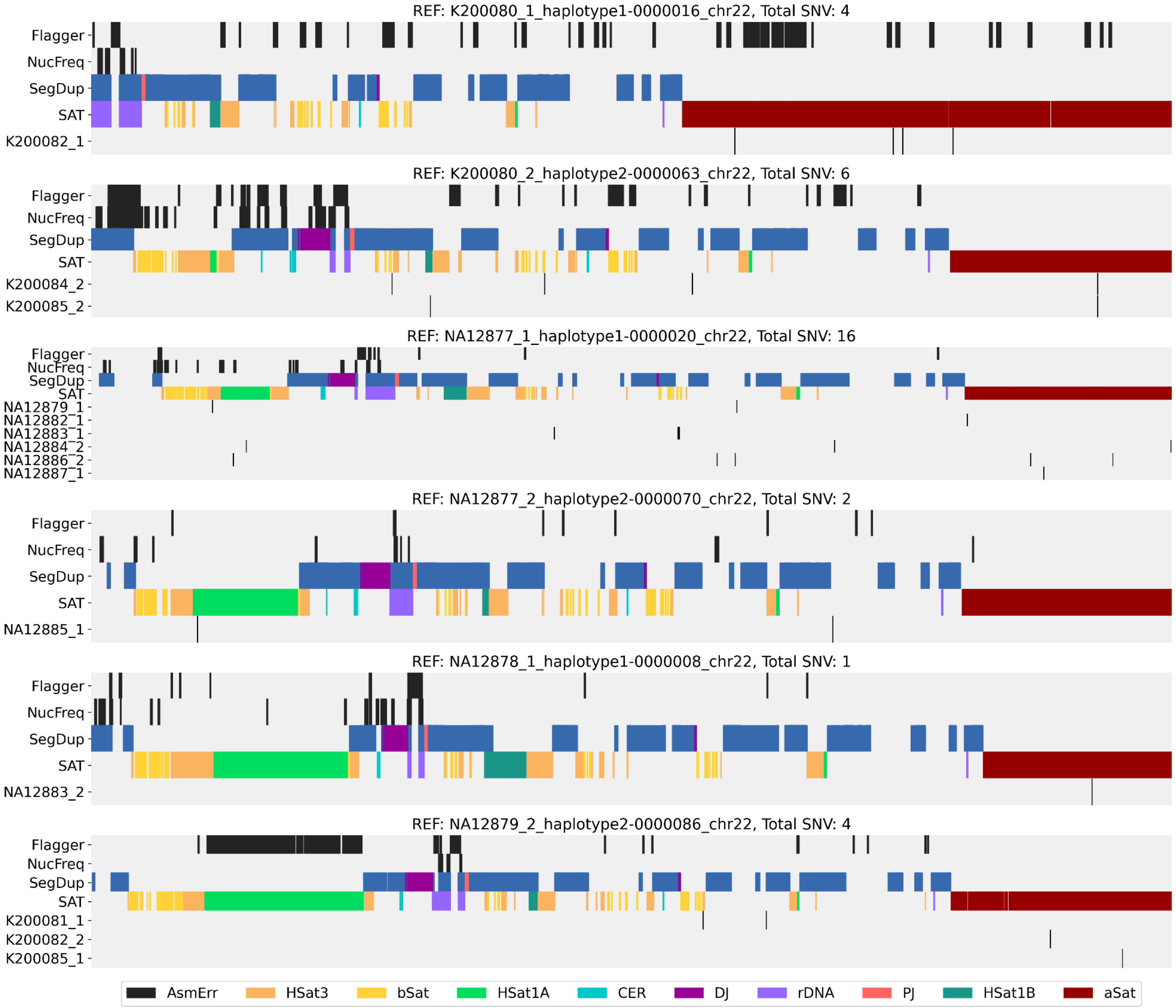


**Figure 44. Locations of chr22 *de novo* SNVs detected from each offspring against the reference parental haplotypes.** (see Figure 40 legend)

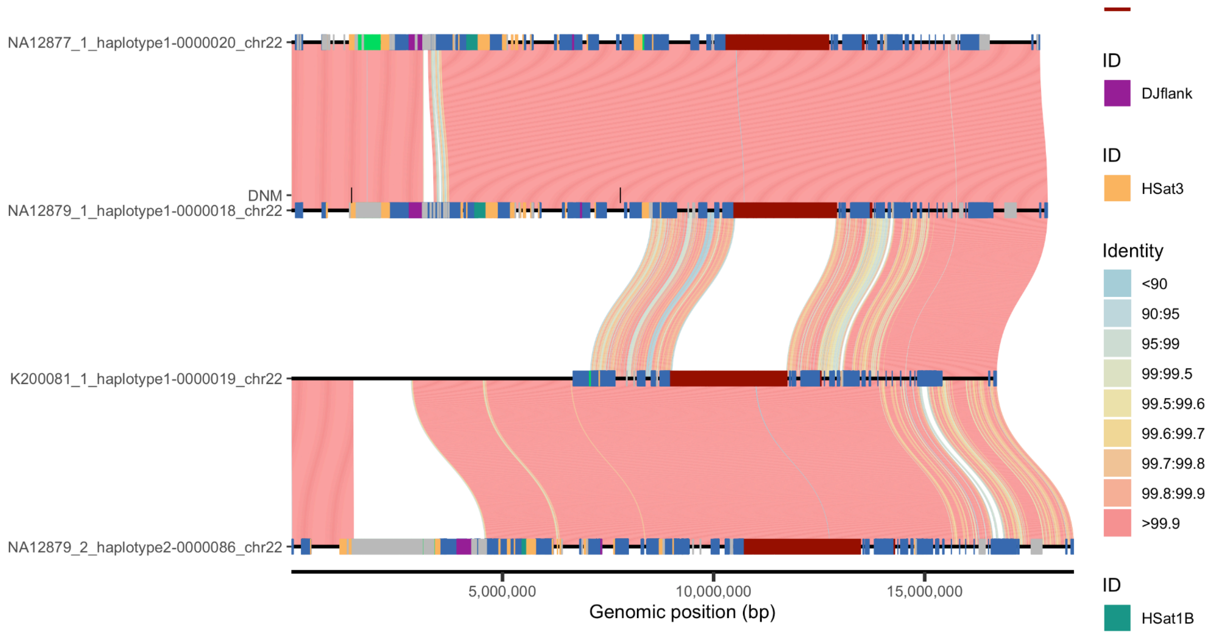


**Figure 45. Two *de novo* SNVs of NA12879 not inherited in G4 due to recombination.** The sequences (from top to bottom) represent:

- G2 father NA12877 (NA12877_1_haplotype1-0000020_chr22),
- G3 NA12879 (NA12879_1_haplotype1-0000018_chr22),
- G4 200081 (K200081_1_haplotype1-0000019_chr22), and
- G3 NA12879 (NA12879_2_haplotype2-0000086_chr22).

The recombination was found in G4 200081.

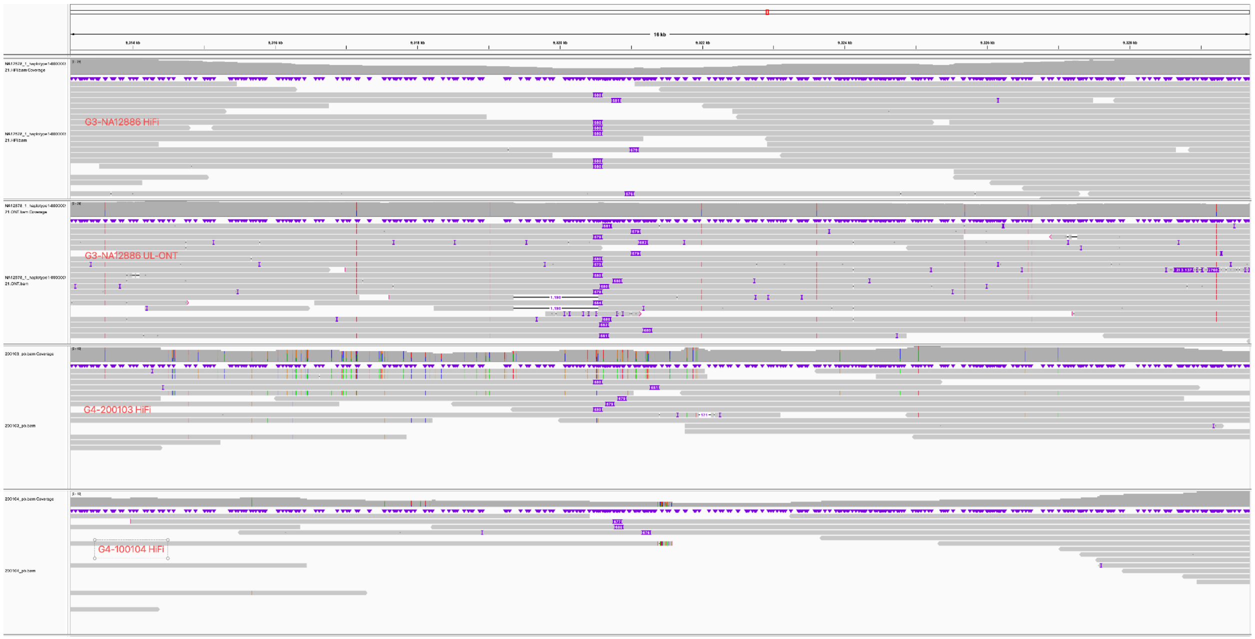


**Figure 46. Transmission of a *de novo* satellite insertion.** The tracks (from top to bottom) show the alignments from G3-NA12886 HiF, G3-NA12886 UL-ONT, G4-200103 HiFi, and G4-200104 HiFi. All reads are aligned to the reference haplotype G2-NA12878.

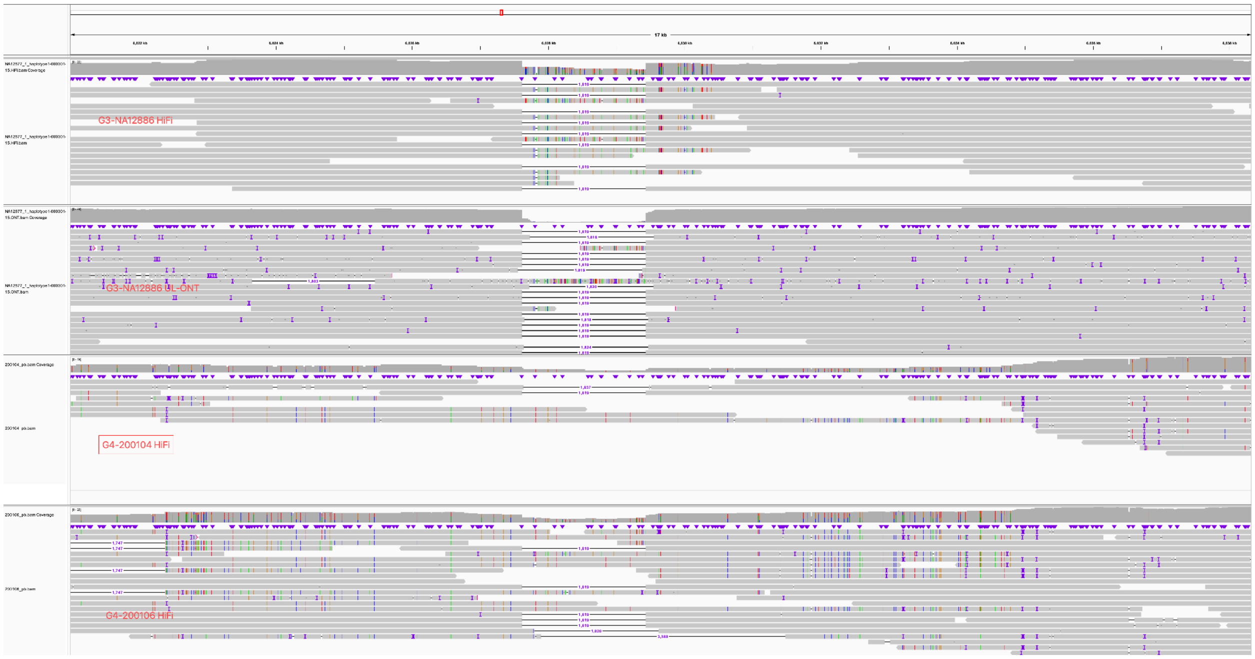


**Figure 47. Transmission of a *de novo* HSat3 deletion.** The tracks (from top to bottom) show the alignments from G3-NA12886 HiFi, G3-NA12886 UL-ONT, G4-200104 HiFi, and G4-200106 HiFi. All reads are aligned to the reference haplotype G2-NA12877.

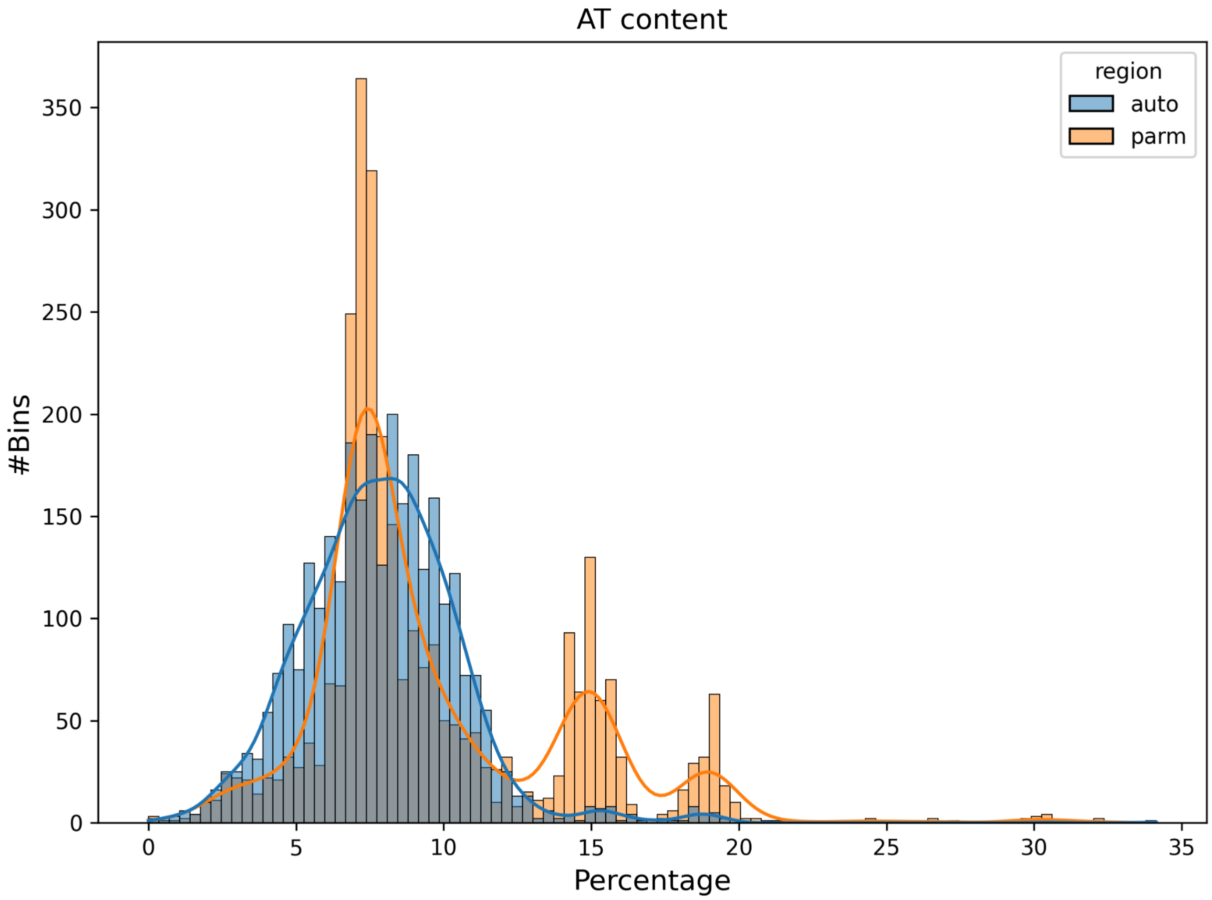


**Figure 48. Comparison of AT dimer percentage on acrocentric short arm and autosomal regions.** The percentage is calculated in 1 kbp bins randomly shuffled on p-arm (light orange) and autosomal regions (light blue).
