## Supplementary material for "Human acrocentric chromosome short arm *de novo* mutation and recombination": Data S2

This file contains all ModDotPlots, NucFreq plots and centromere validation plots for 156 pq-scatigs from 16 samples

NA12877

### chr13

#### NA12877_1_haplotype1-0000001_chr13

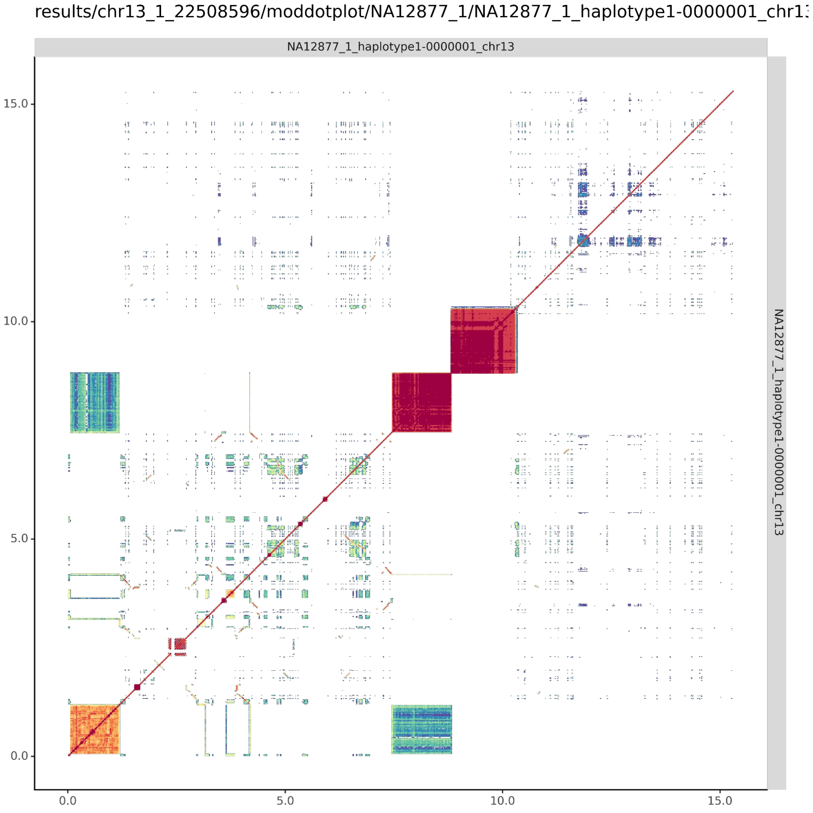


#### NA12877_2_haplotype2-0000046_chr13

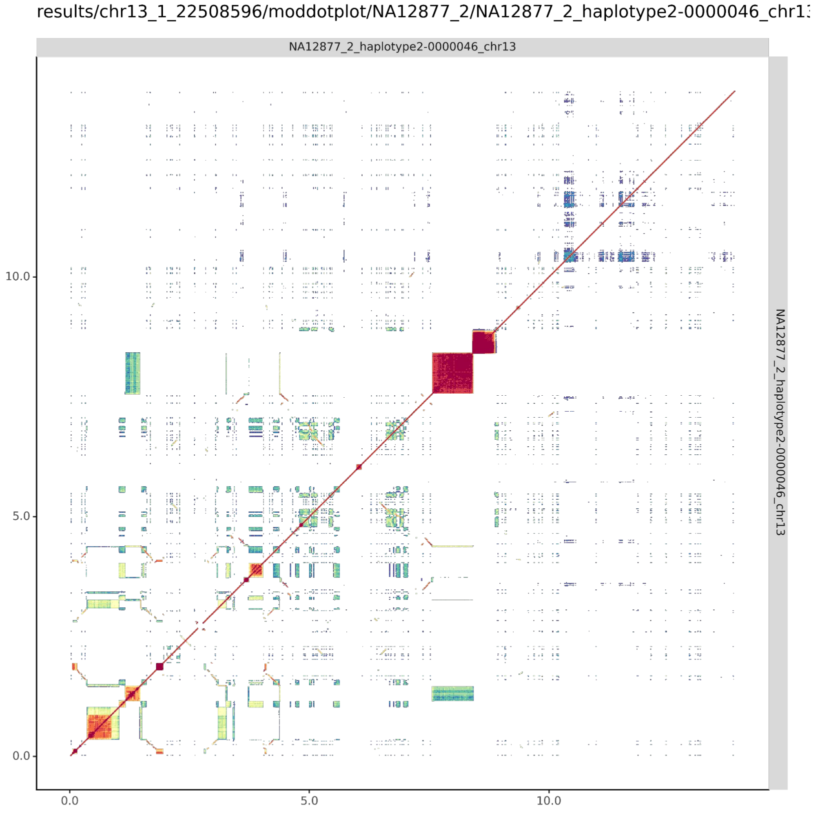

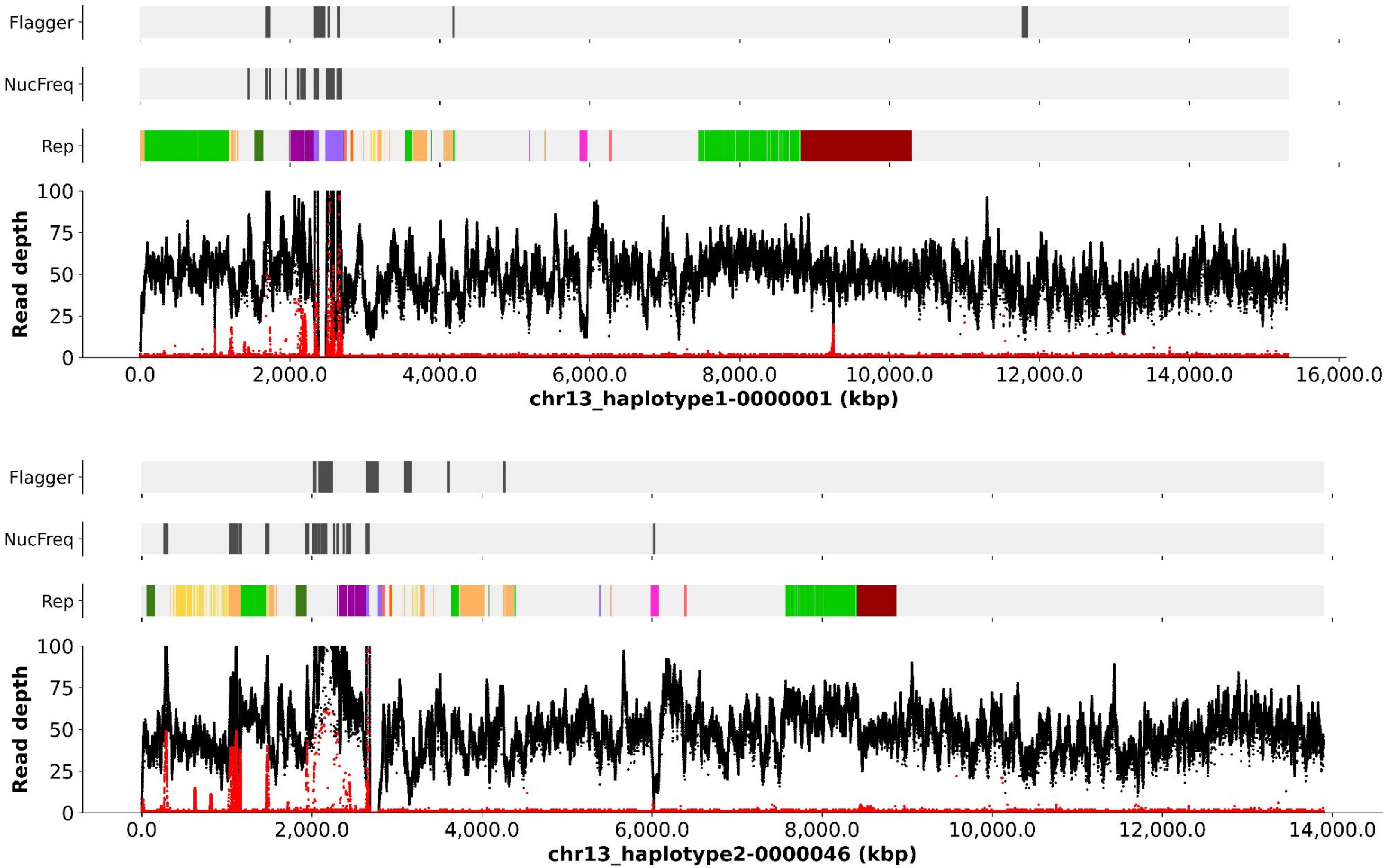
Flagger
NucFreq
Rep
Read depth
100
75
50
25
0
0.0
2,000.0
4,000.0
6,000.0
8,000.0
10,000.0
12,000.0
14,000.0
16,000.0
chr13_haplotype1-0000001 (kbp)
Flagger
NucFreq
Rep
Read depth
100
75
50
25
0
0.0
2,000.0
4,000.0
6,000.0
8,000.0
10,000.0
12,000.0
14,000.0
chr13_haplotype2-0000046 (kbp)

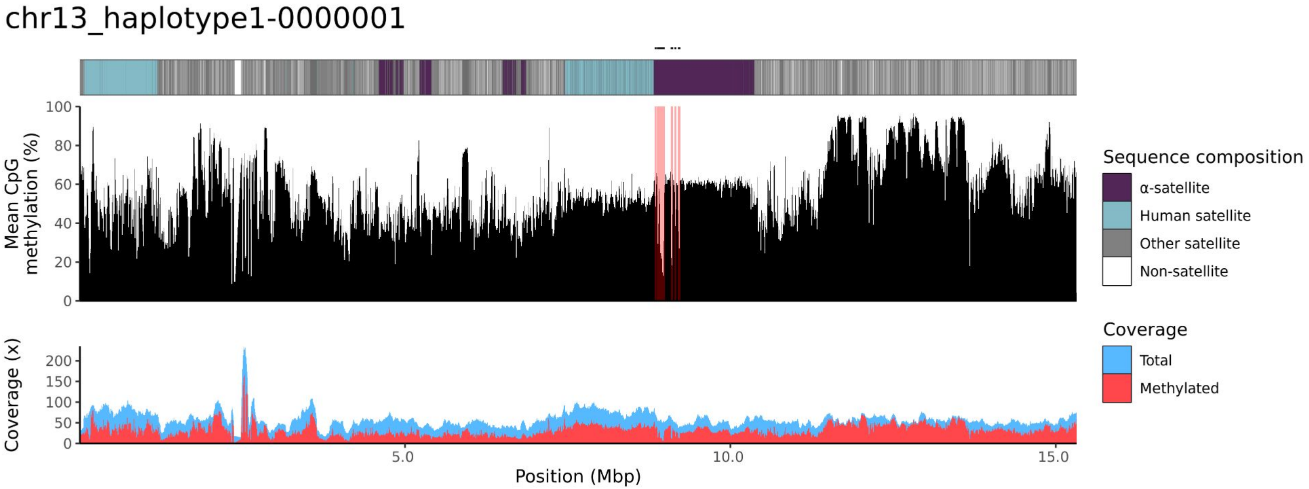
chr13_haplotype1-0000001
Mean CpG methylation (%)
100
80
60
40
20
0
Coverage (x)
200
150
100
50
0
5.0
10.0
15.0
Position (Mbp)
Sequence composition
α-satellite
Human satellite
Other satellite
Non-satellite
Coverage
Total
Methylated

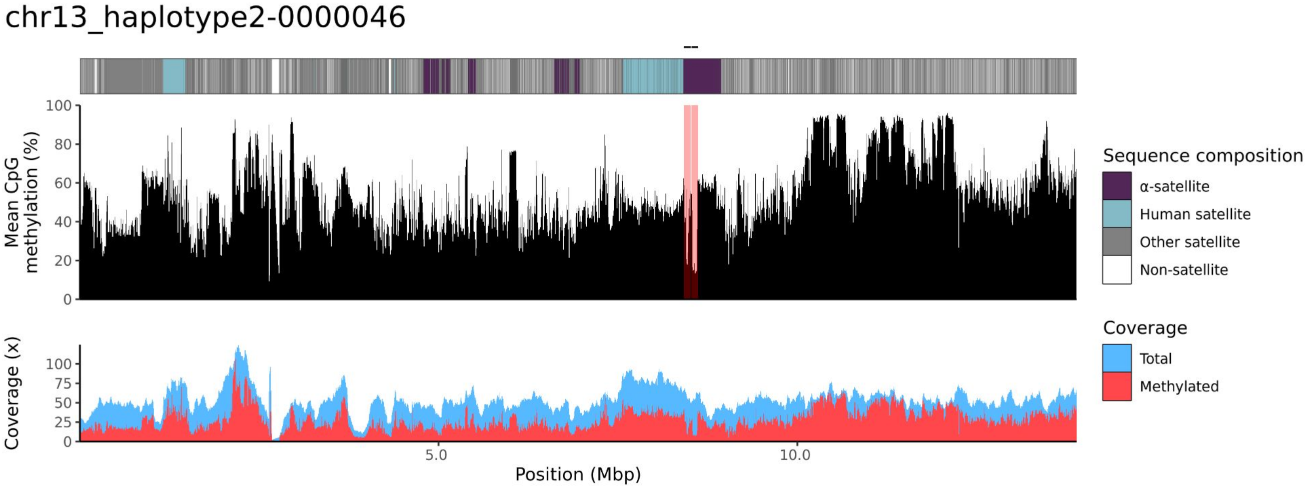
chr13_haplotype2-0000046
Mean CpG methylation (%)
100
80
60
40
20
0
Coverage (x)
100
75
50
25
0
5.0
10.0
Position (Mbp)
Sequence composition
α-satellite
Human satellite
Other satellite
Non-satellite
Coverage
Total
Methylated

### chr14

#### NA12877_1_haplotype1-0000013_chr14

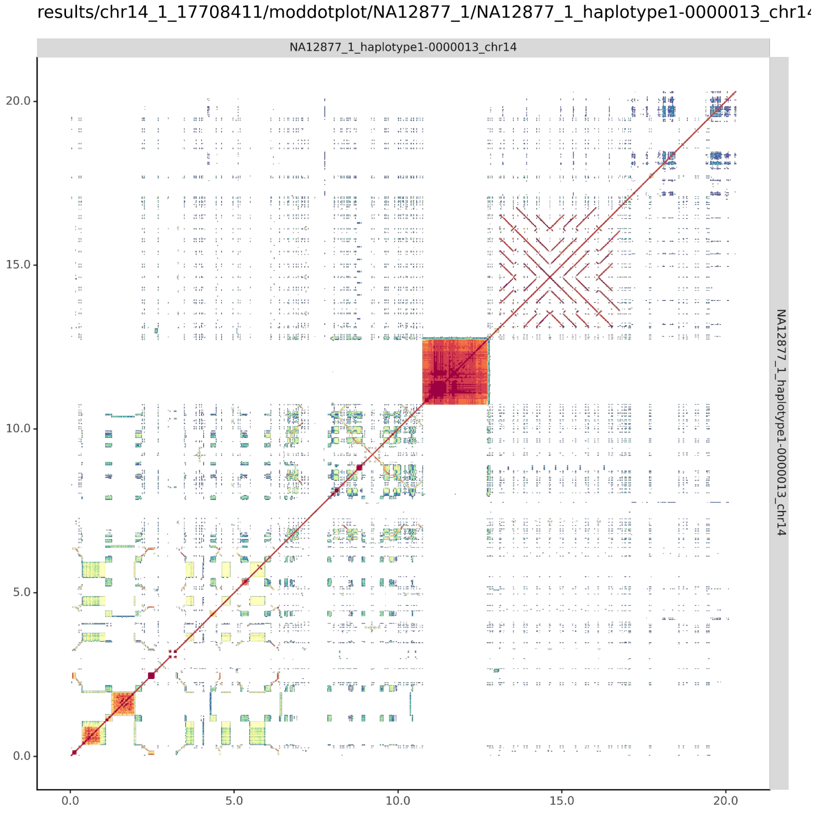


#### NA12877_2_haplotype2-0000056_chr14

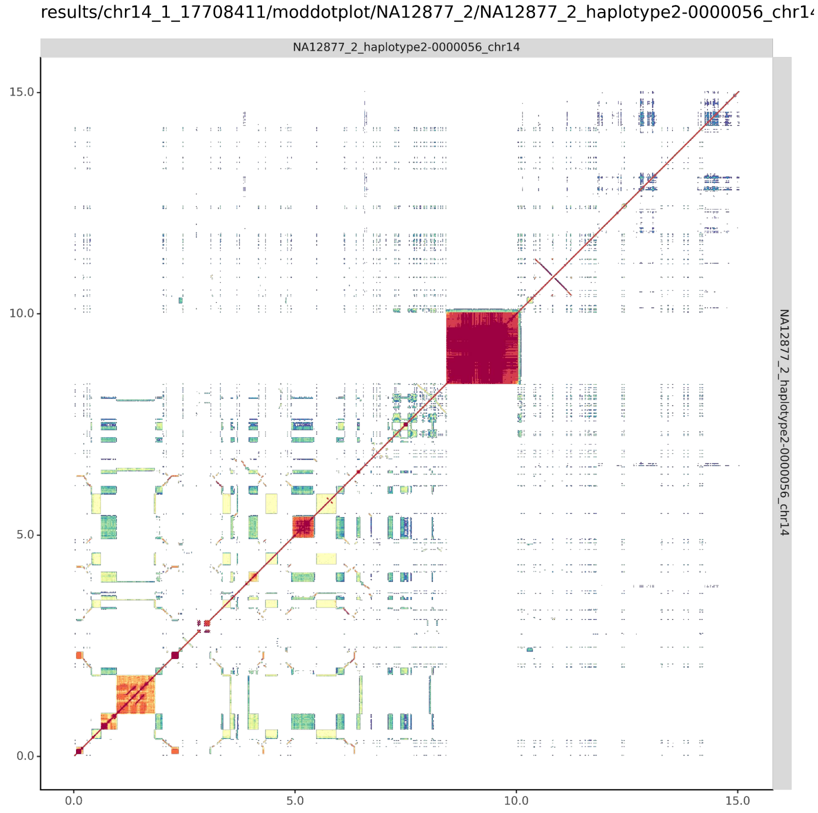

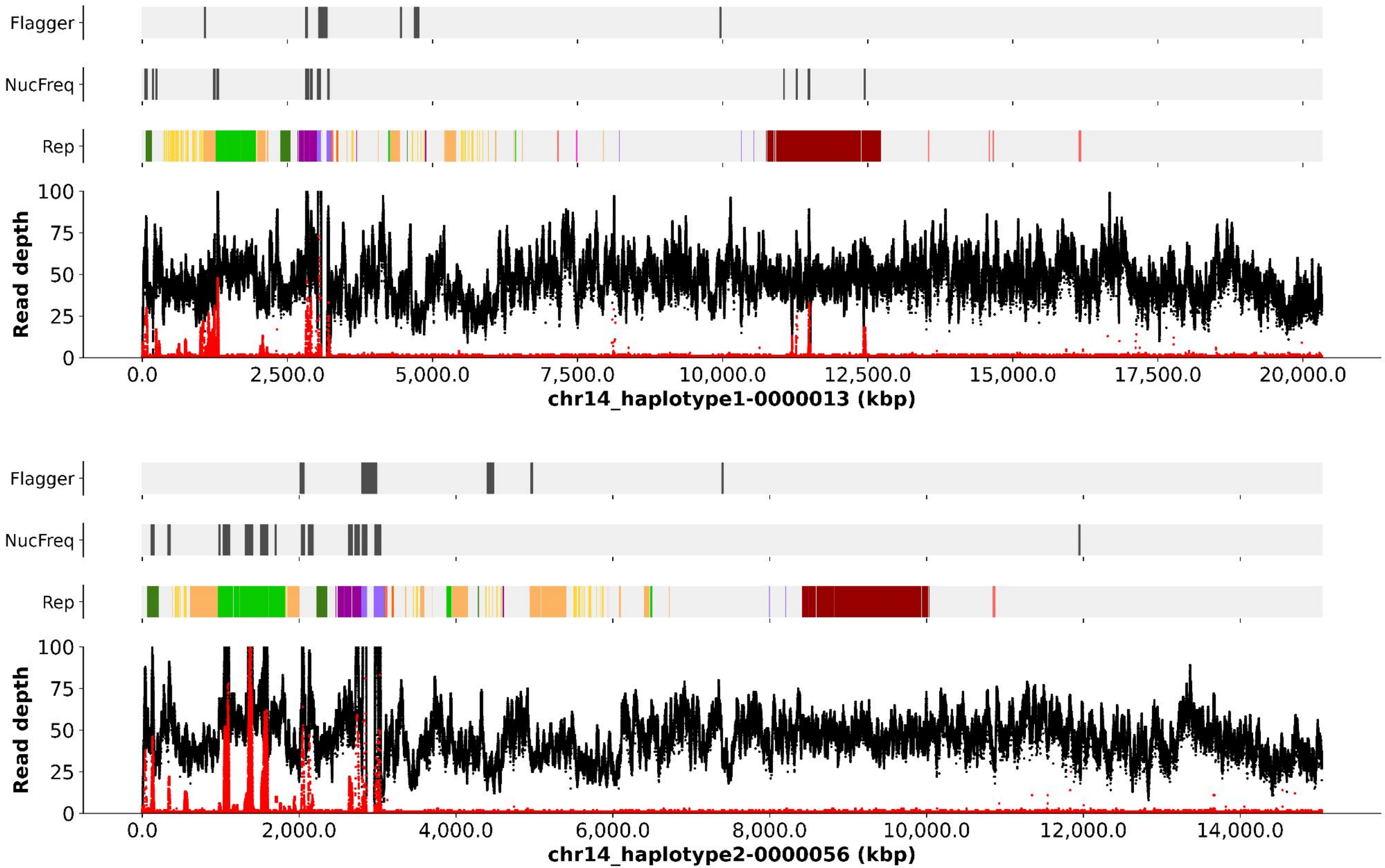
Flagger
NucFreq
Rep
Read depth
100
75
50
25
0
0.0
2,500.0
5,000.0
7,500.0
10,000.0
12,500.0
15,000.0
17,500.0
20,000.0
chr14_haplotype1-0000013 (kbp)
Flagger
NucFreq
Rep
Read depth
100
75
50
25
0
0.0
2,000.0
4,000.0
6,000.0
8,000.0
10,000.0
12,000.0
14,000.0
chr14_haplotype2-0000056 (kbp)

chr14_haplotype1-0000013

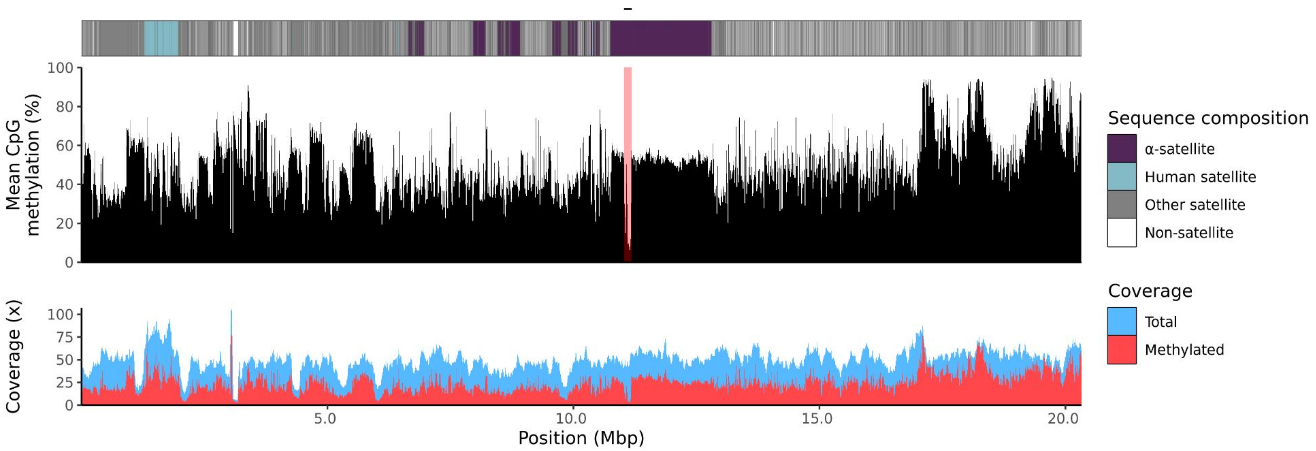


chr14_haplotype2-0000056

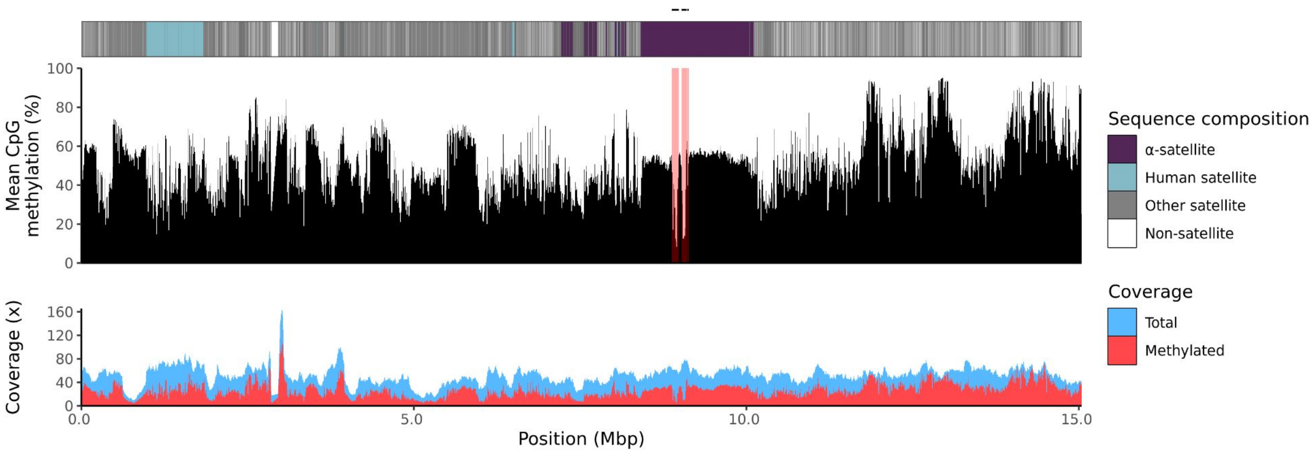

### chr15

#### NA12877_1_haplotype1-0000014_chr15

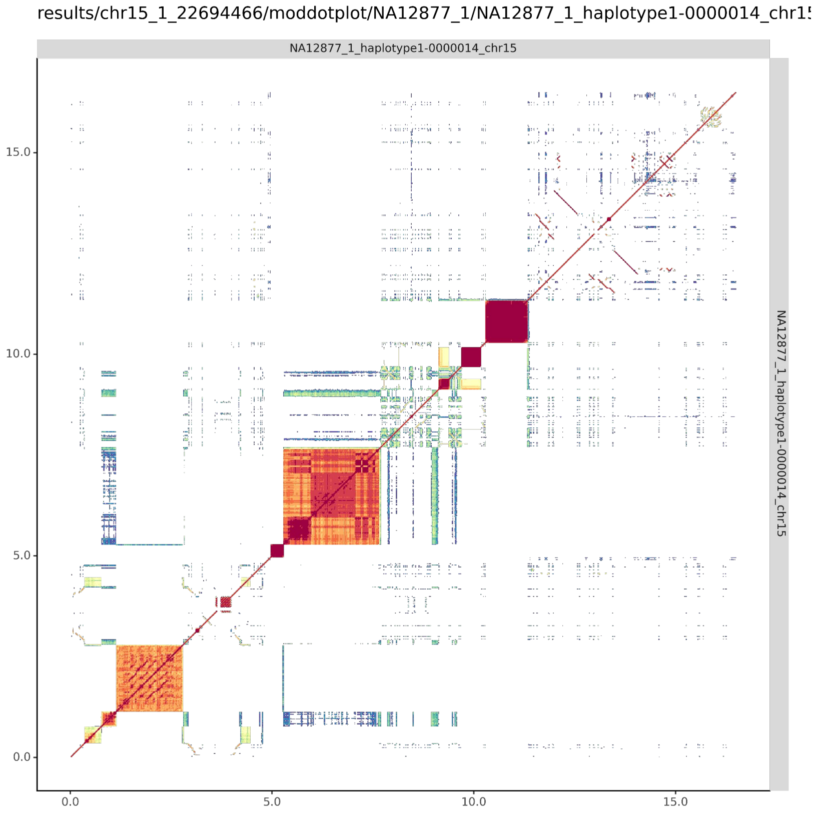


#### NA12877_2_haplotype2-0000057_chr15

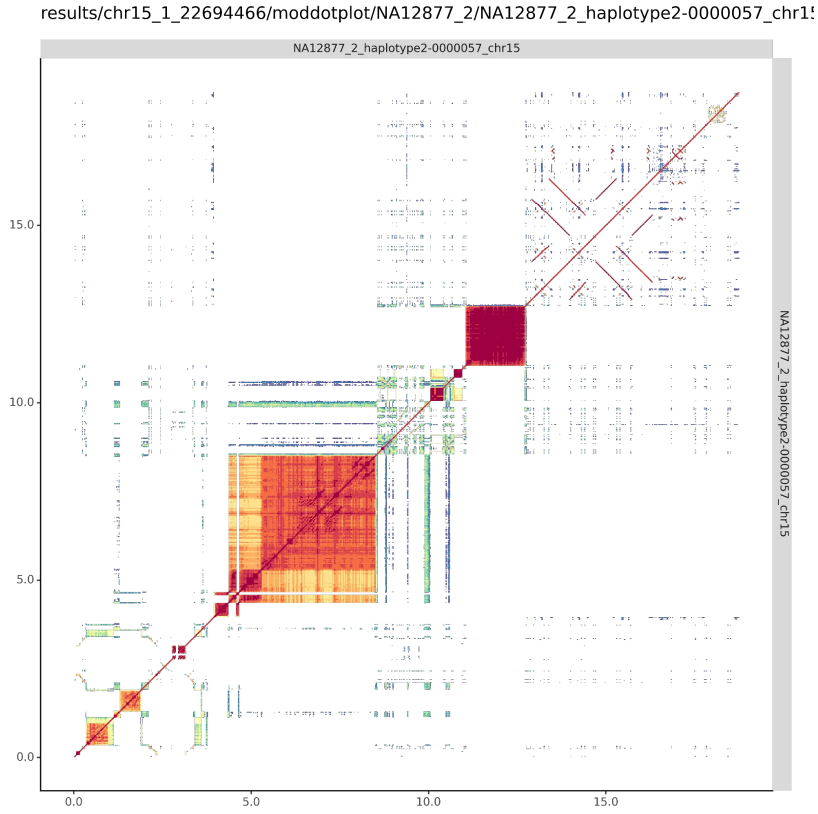

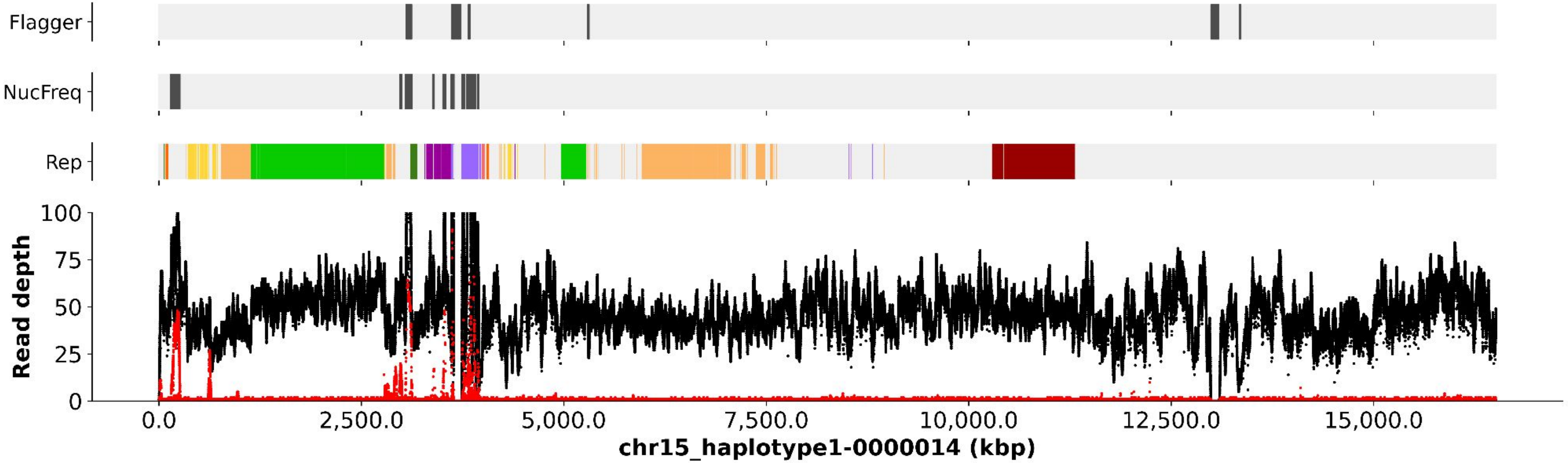
Flagger
NucFreq
Rep
Read depth
100
75
50
25
0
0.0
2,500.0
5,000.0
7,500.0
10,000.0
12,500.0
15,000.0
chr15_haplotype1-0000014 (kbp)

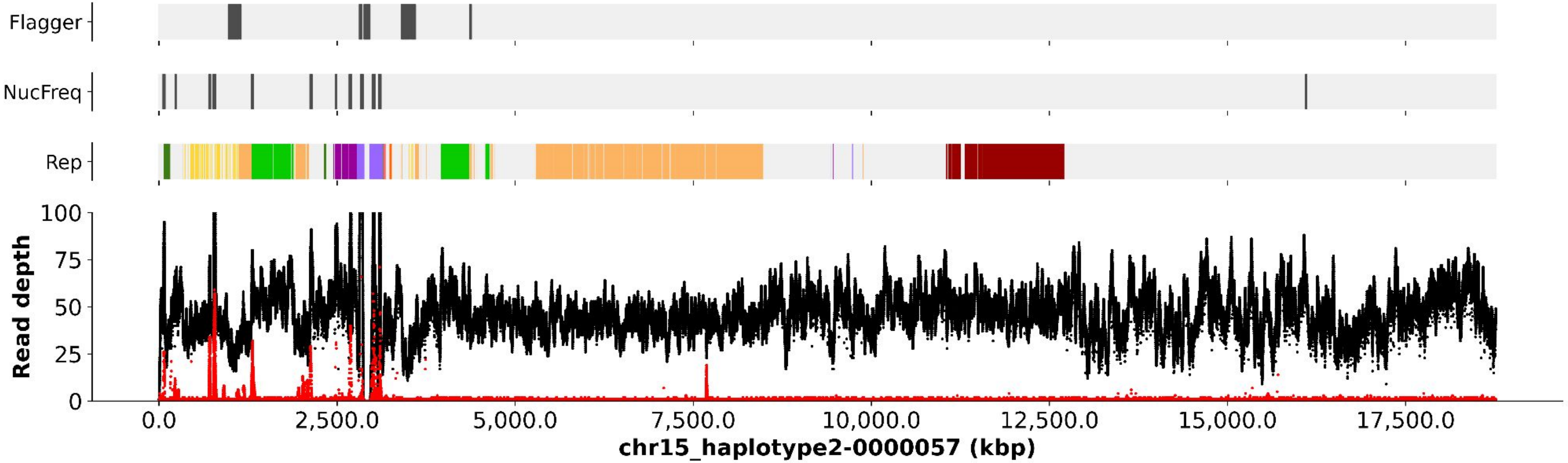
Flagger
NucFreq
Rep
Read depth
100
75
50
25
0
0.0
2,500.0
5,000.0
7,500.0
10,000.0
12,500.0
15,000.0
17,500.0
chr15_haplotype2-0000057 (kbp)

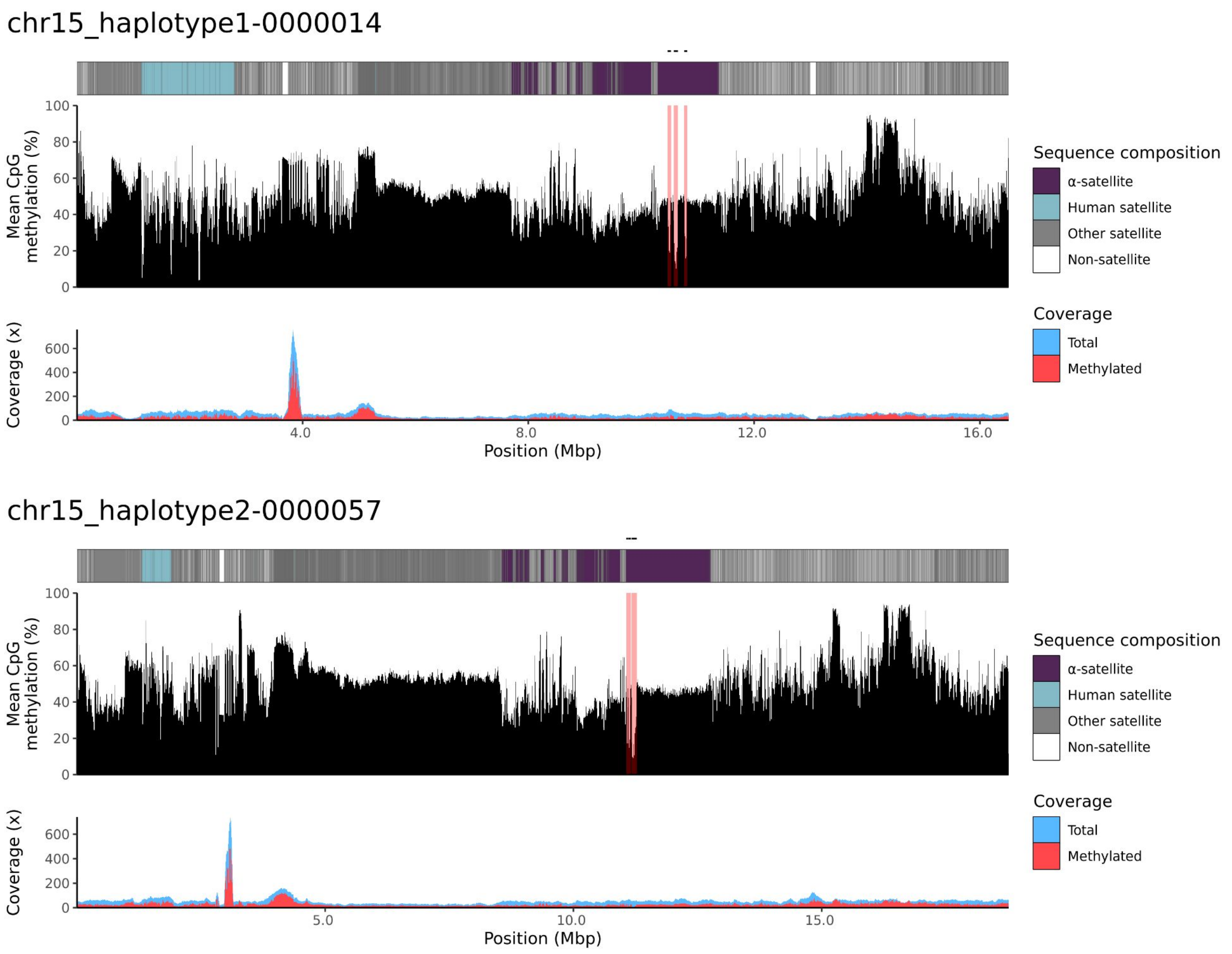
chr15_haplotype1-0000014
Mean CpG methylation (%)
100
80
60
40
20
0
Coverage (x)
600
400
200
0
4.0
8.0
12.0
16.0
Position (Mbp)
Sequence composition
α-satellite
Human satellite
Other satellite
Non-satellite
Coverage
Total
Methylated
chr15_haplotype2-0000057
Mean CpG methylation (%)
100
80
60
40
20
0
Coverage (x)
600
400
200
0
5.0
10.0
15.0
Position (Mbp)
Sequence composition
α-satellite
Human satellite
Other satellite
Non-satellite
Coverage
Total
Methylated

### chr21

#### NA12877_1_haplotype1-0000023_chr21

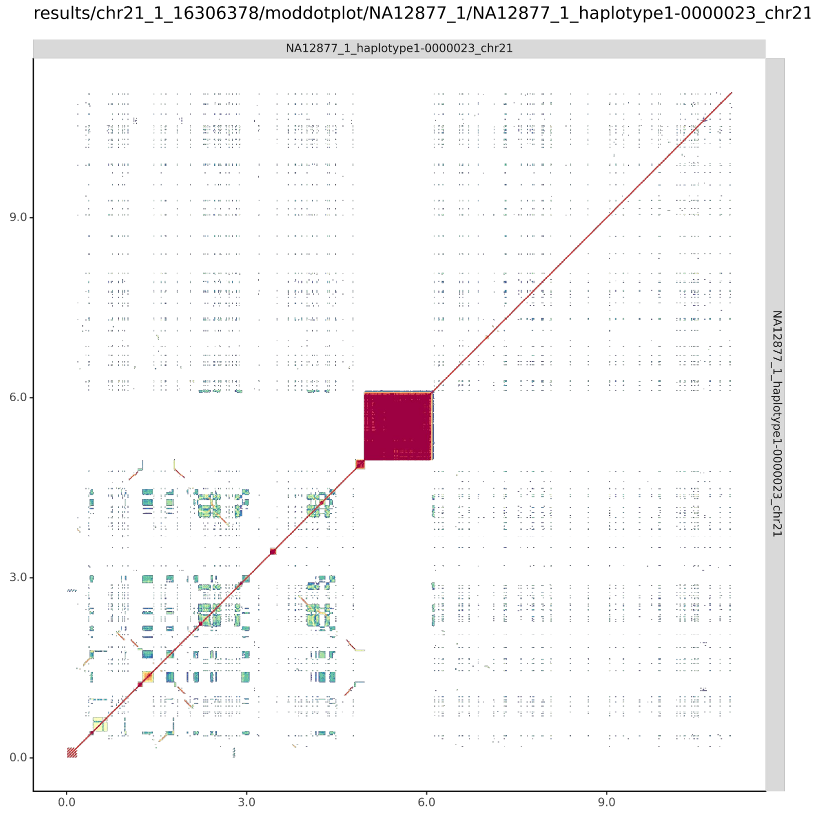


#### NA12877_2_haplotype2-0000064_chr21

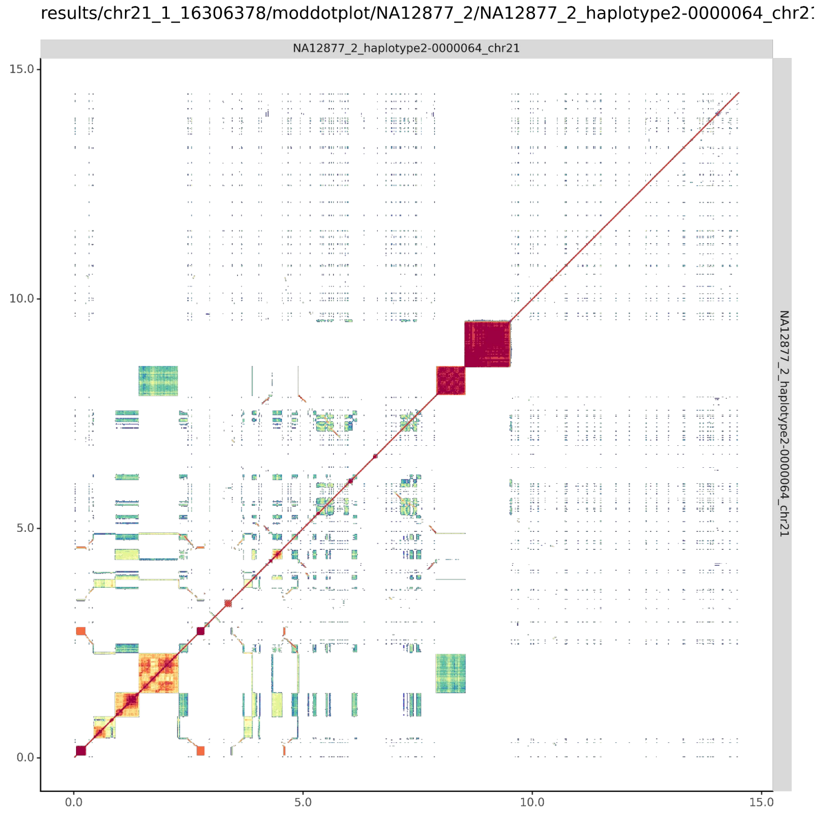

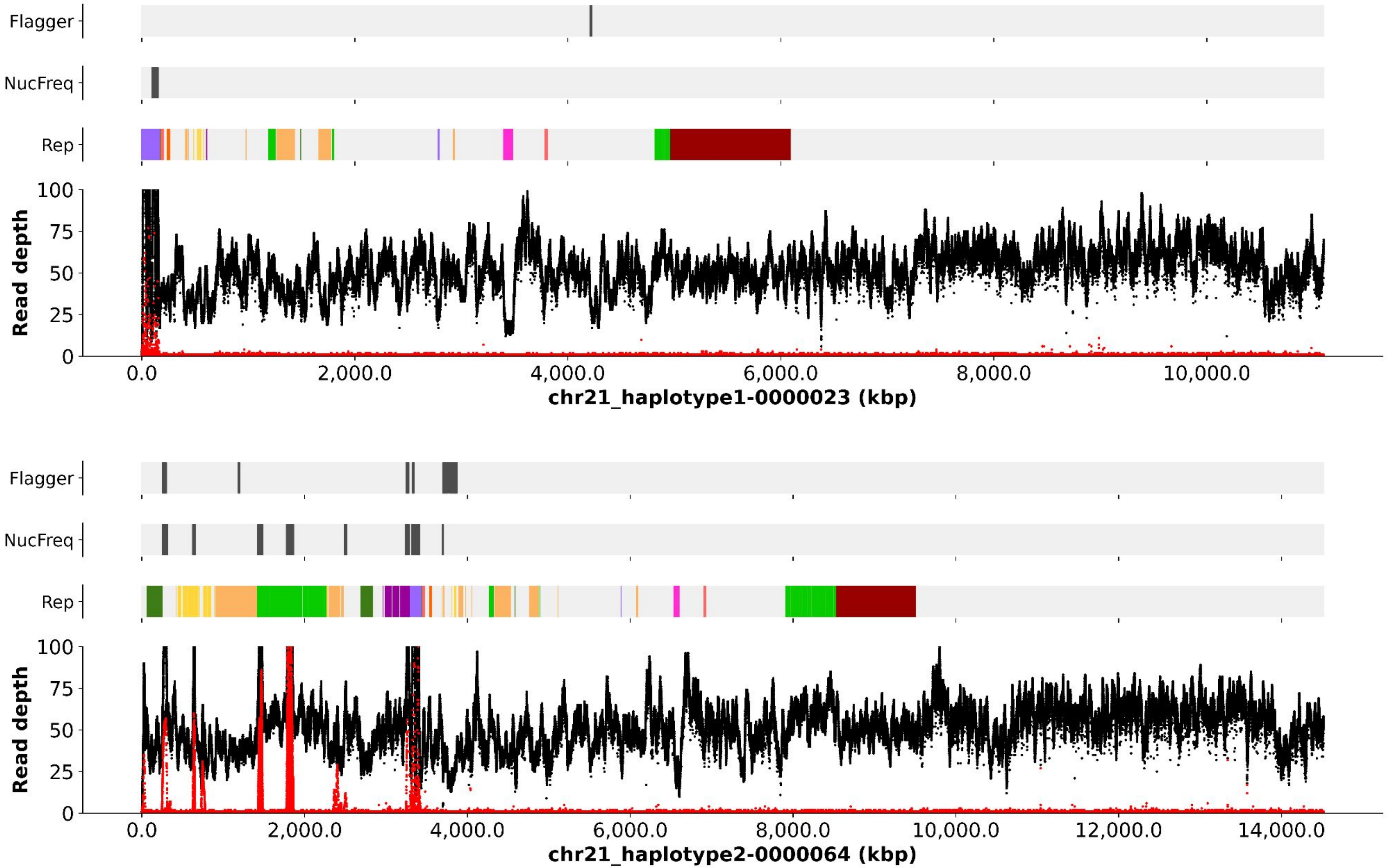
Flagger
NucFreq
Rep
Read depth
0
25
50
75
100
0.0
2,000.0
4,000.0
6,000.0
8,000.0
10,000.0
chr21_haplotype1-0000023 (kbp)
Flagger
NucFreq
Rep
Read depth
0
25
50
75
100
0.0
2,000.0
4,000.0
6,000.0
8,000.0
10,000.0
12,000.0
14,000.0
chr21_haplotype2-0000064 (kbp)

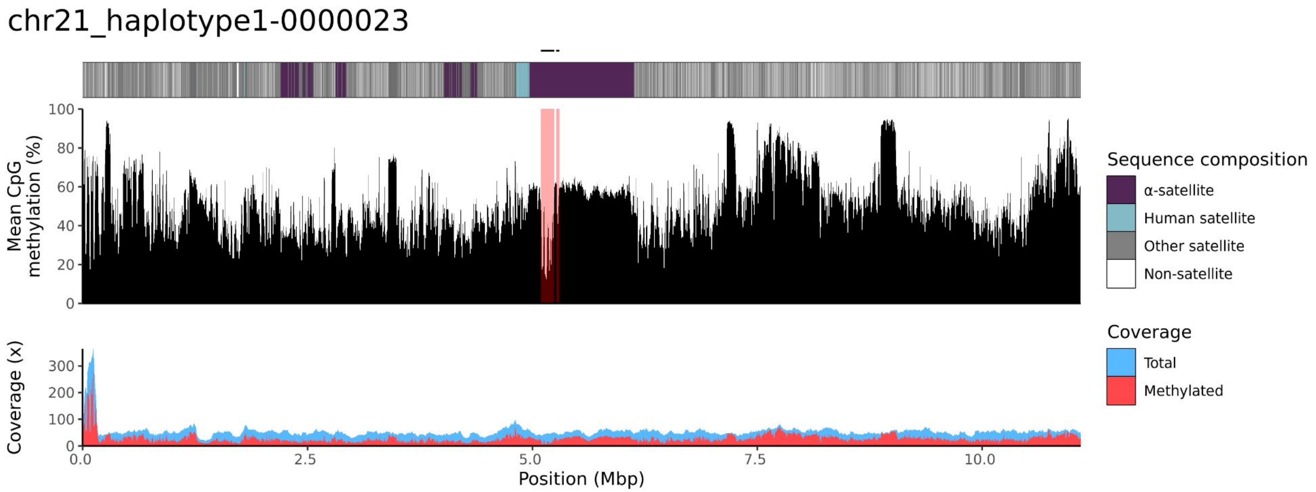
chr21_haplotype1-0000023
Mean CpG methylation (%)
100
80
60
40
20
0
Coverage (x)
300
200
100
0
0.0
2.5
5.0
7.5
10.0
Position (Mbp)
Sequence composition
α-satellite
Human satellite
Other satellite
Non-satellite
Coverage
Total
Methylated

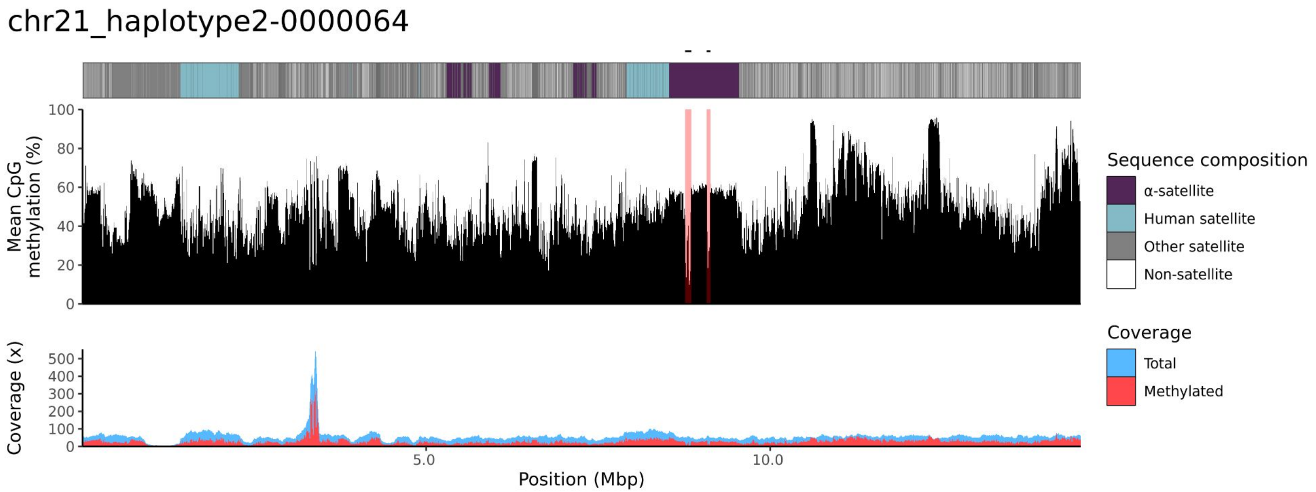
chr21_haplotype2-0000064
Mean CpG methylation (%)
100
80
60
40
20
0
Coverage (x)
500
400
300
200
100
0
5.0
10.0
Position (Mbp)
Sequence composition
α-satellite
Human satellite
Other satellite
Non-satellite
Coverage
Total
Methylated

### chr22

#### NA12877_1_haplotype1-0000020_chr22

results/chr22_1_20711065/moddotplot/NA12877_1/NA12877_1_haplotype1-0000020_chr2

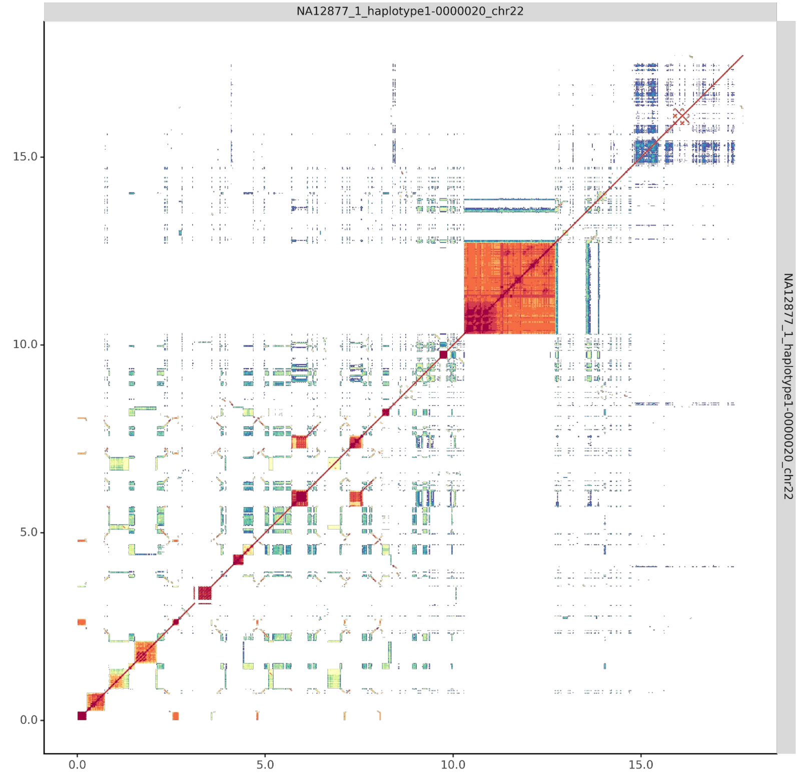


#### NA12877_2_haplotype2-0000070_chr22

results/chr22_1_20711065/moddotplot/NA12877_2/NA12877_2_haplotype2-0000070_chr2

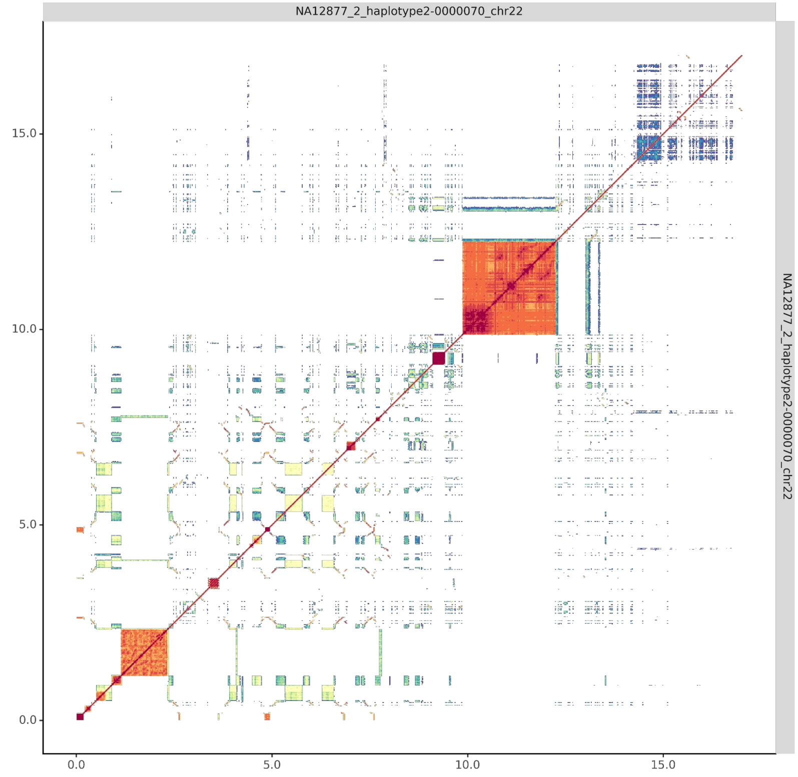

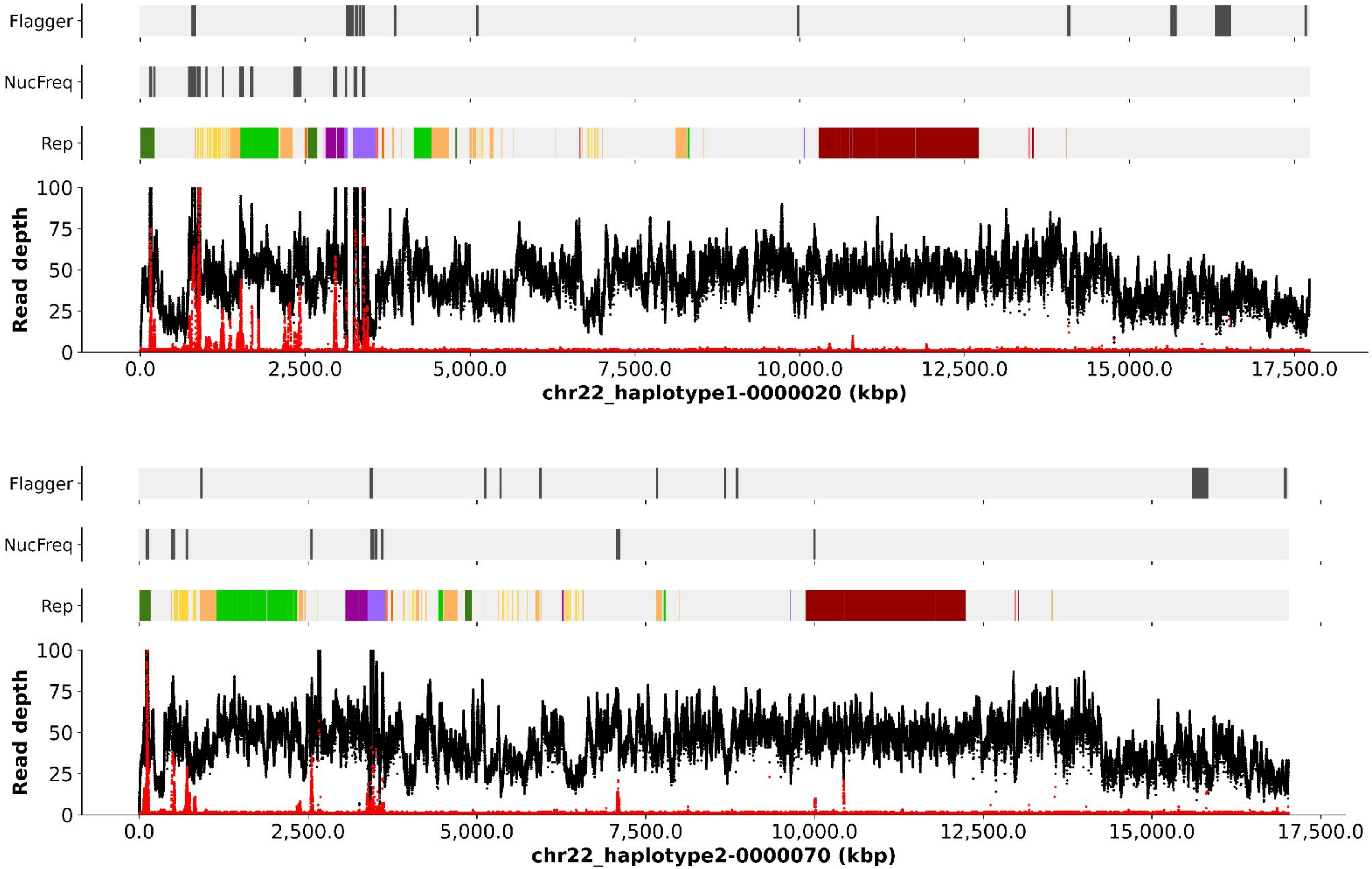

Flagger
NucFreq
Rep
Read depth
100
75
50
25
0
0.0
2,500.0
5,000.0
7,500.0
10,000.0
12,500.0
15,000.0
17,500.0
chr22_haplotype1-0000020 (kbp)
Flagger
NucFreq
Rep
Read depth
100
75
50
25
0
0.0
2,500.0
5,000.0
7,500.0
10,000.0
12,500.0
15,000.0
17,500.0
chr22_haplotype2-0000070 (kbp)

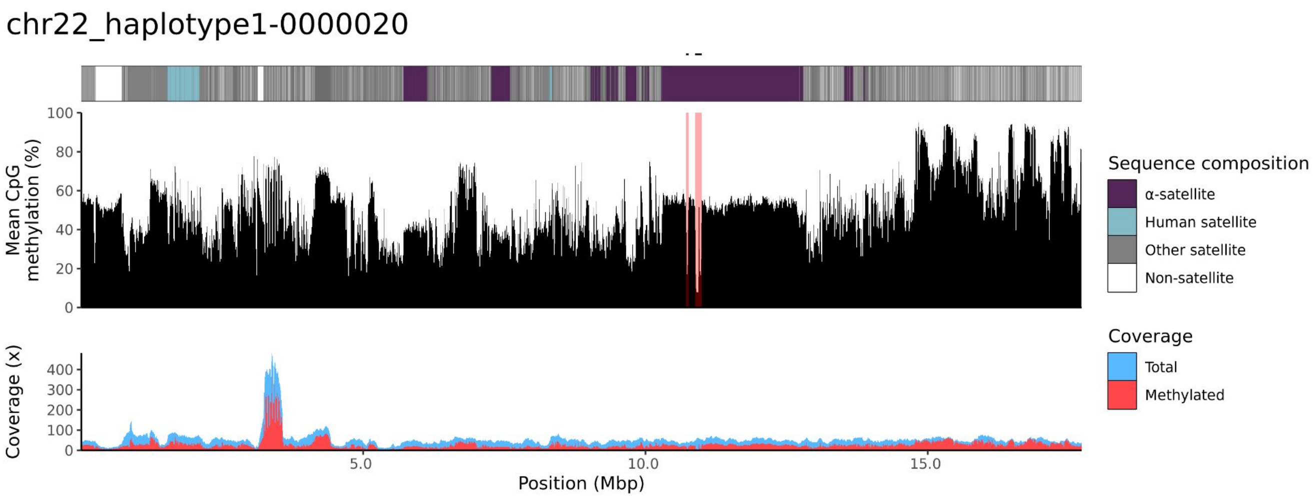
chr22_haplotype1-0000020
Mean CpG methylation (%)
100
80
60
40
20
0
Coverage (x)
400
300
200
100
0
5.0
10.0
15.0
Position (Mbp)
Sequence composition
α-satellite
Human satellite
Other satellite
Non-satellite
Coverage
Total
Methylated

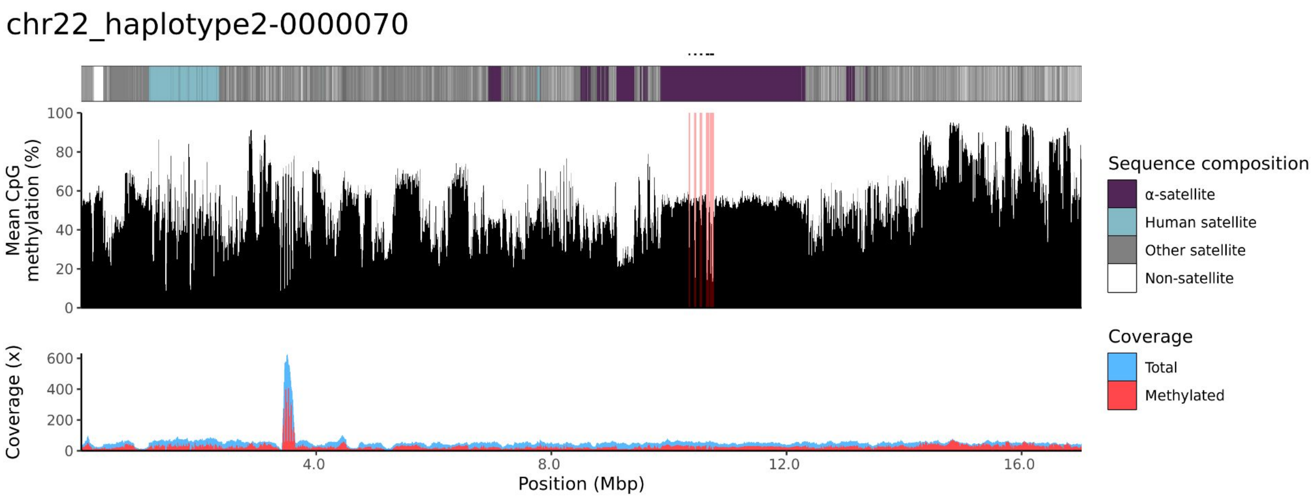
chr22_haplotype2-0000070
Mean CpG methylation (%)
100
80
60
40
20
0
Coverage (x)
600
400
200
0
4.0
8.0
12.0
16.0
Position (Mbp)
Sequence composition
α-satellite
Human satellite
Other satellite
Non-satellite
Coverage
Total
Methylated

NA12878

### chr13

#### NA12878_1_haplotype1-0000005_chr13

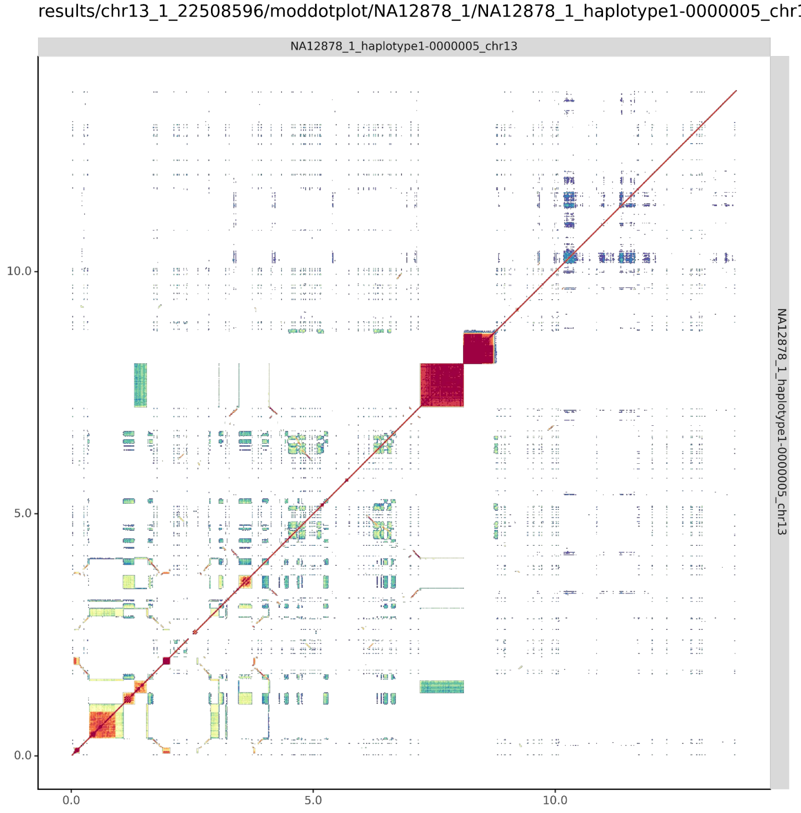


#### NA12878_2_haplotype2-0000030_chr13

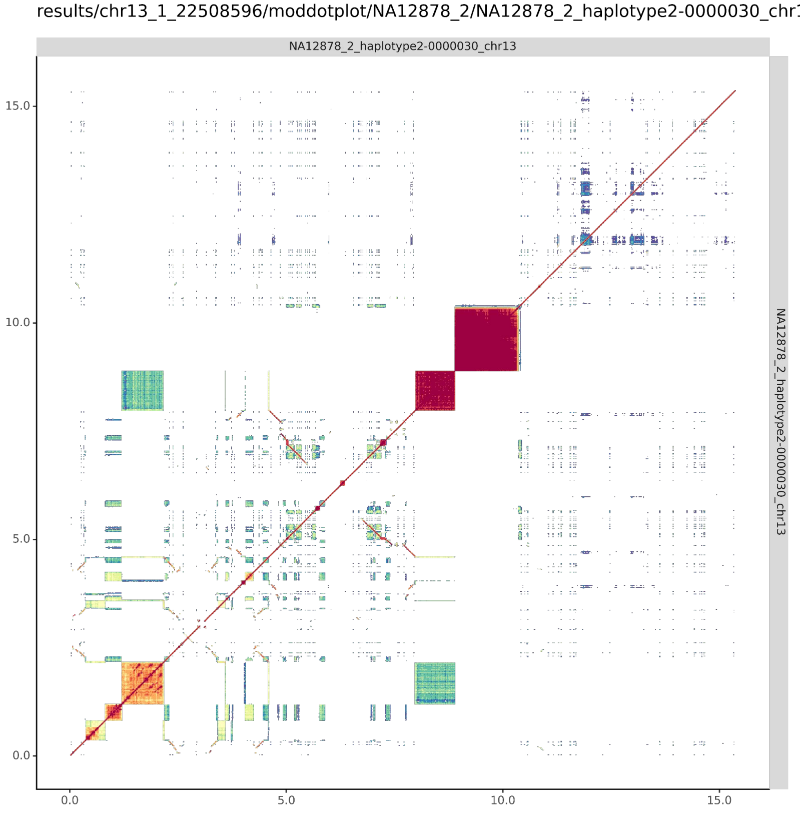

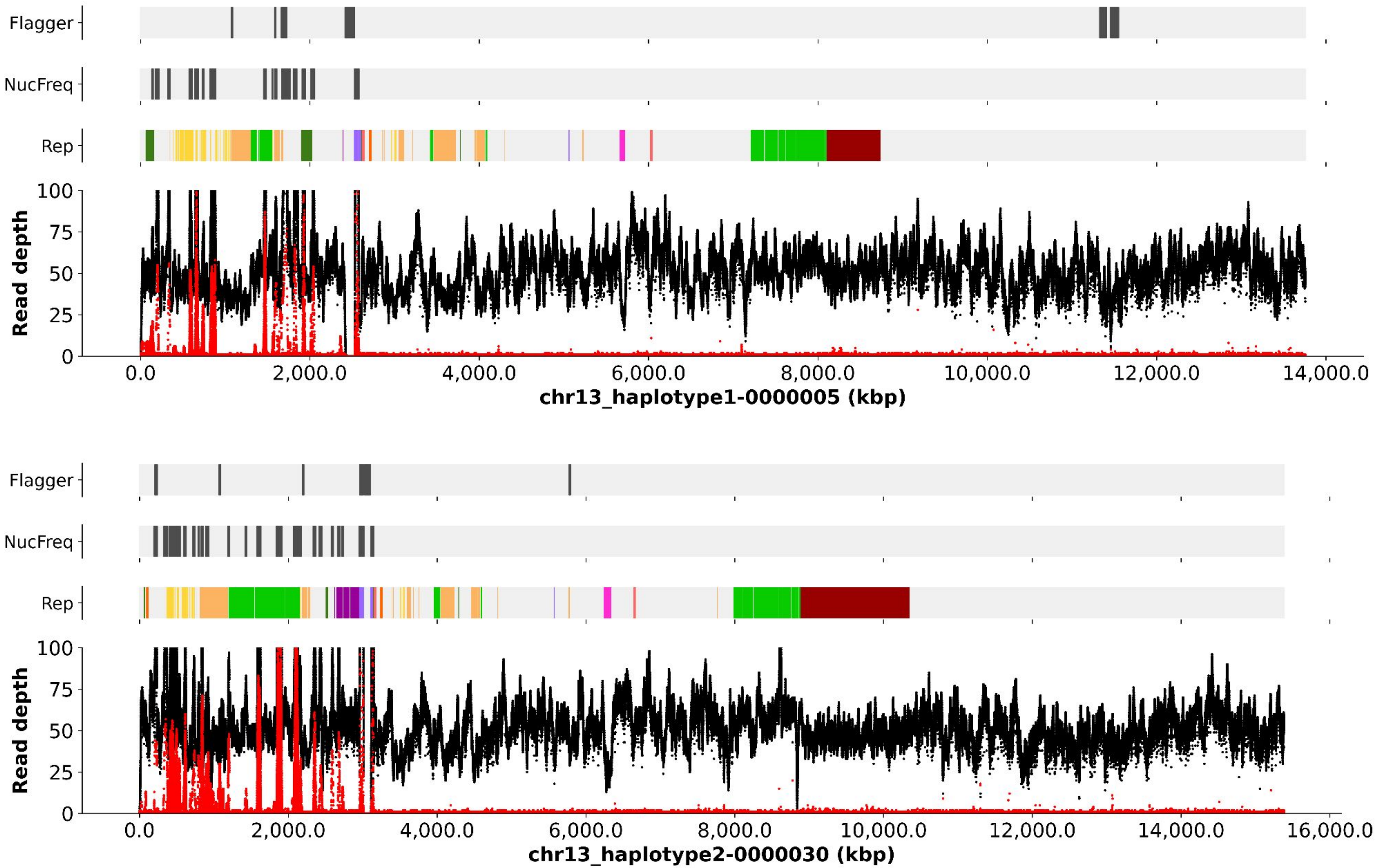
Flagger
NucFreq
Rep
Read depth
100
75
50
25
0
0.0
2,000.0
4,000.0
6,000.0
8,000.0
10,000.0
12,000.0
14,000.0
chr13_haplotype1-0000005 (kbp)
Flagger
NucFreq
Rep
Read depth
100
75
50
25
0
0.0
2,000.0
4,000.0
6,000.0
8,000.0
10,000.0
12,000.0
14,000.0
16,000.0
chr13_haplotype2-0000030 (kbp)

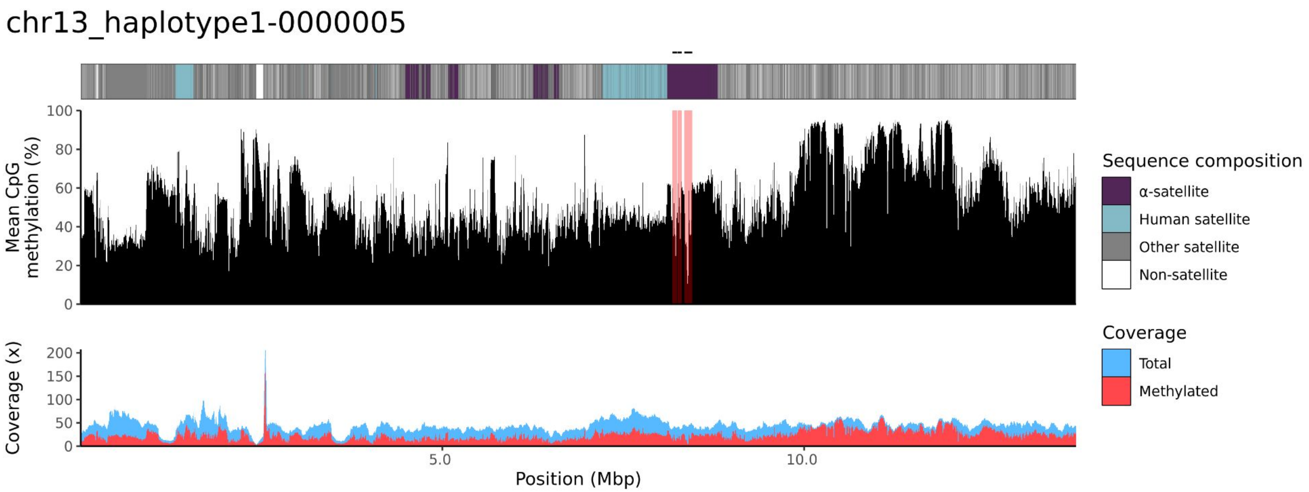
chr13_haplotype1-0000005
Mean CpG methylation (%)
100
80
60
40
20
0
Coverage (x)
200
150
100
50
0
5.0
10.0
Position (Mbp)
Sequence composition
α-satellite
Human satellite
Other satellite
Non-satellite
Coverage
Total
Methylated

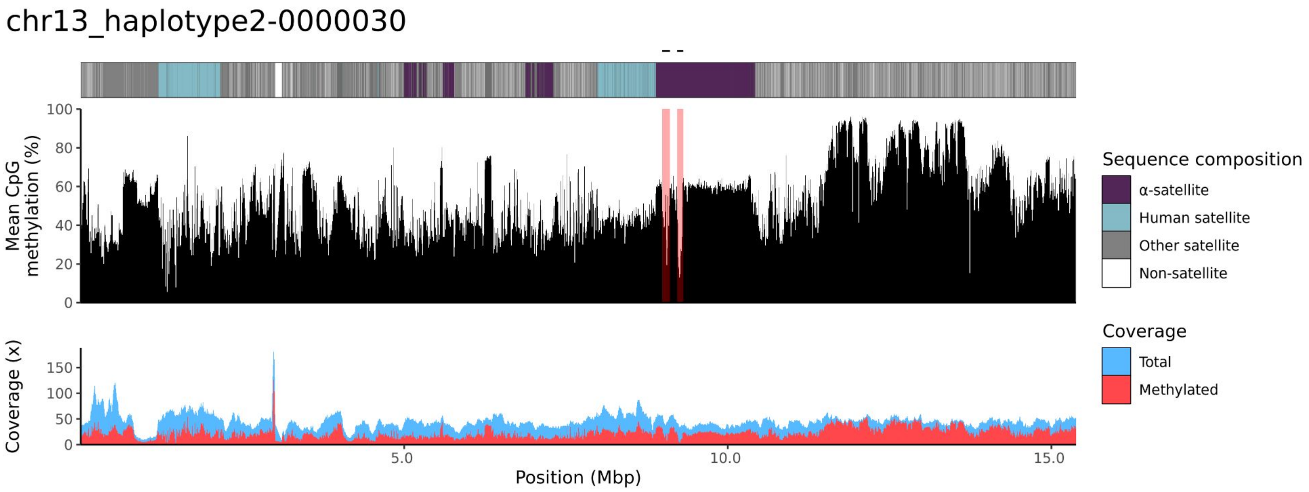
chr13_haplotype2-0000030
Mean CpG methylation (%)
100
80
60
40
20
0
Coverage (x)
150
100
50
0
5.0
10.0
15.0
Position (Mbp)
Sequence composition
α-satellite
Human satellite
Other satellite
Non-satellite
Coverage
Total
Methylated

### chr14

#### NA12878_1_haplotype1-0000013_chr14

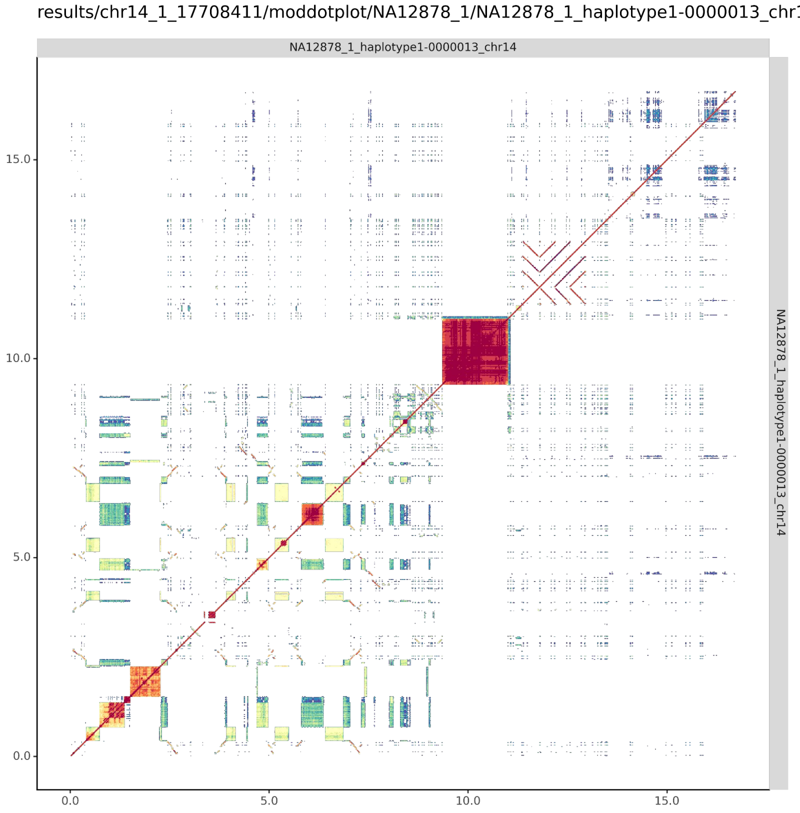


#### NA12878_2_haplotype2-0000041_chr14

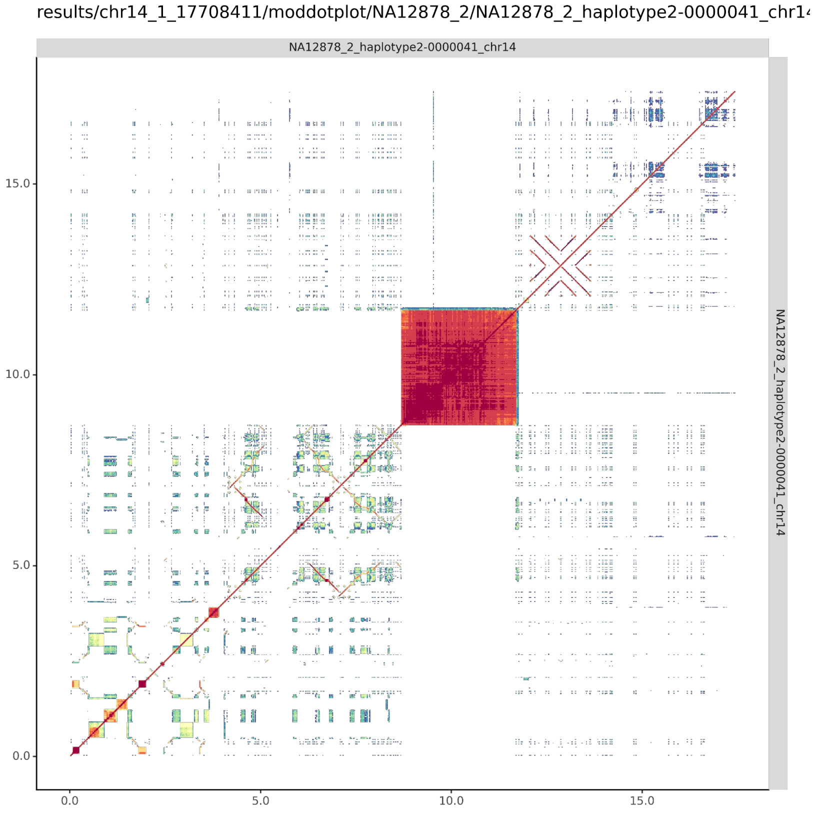

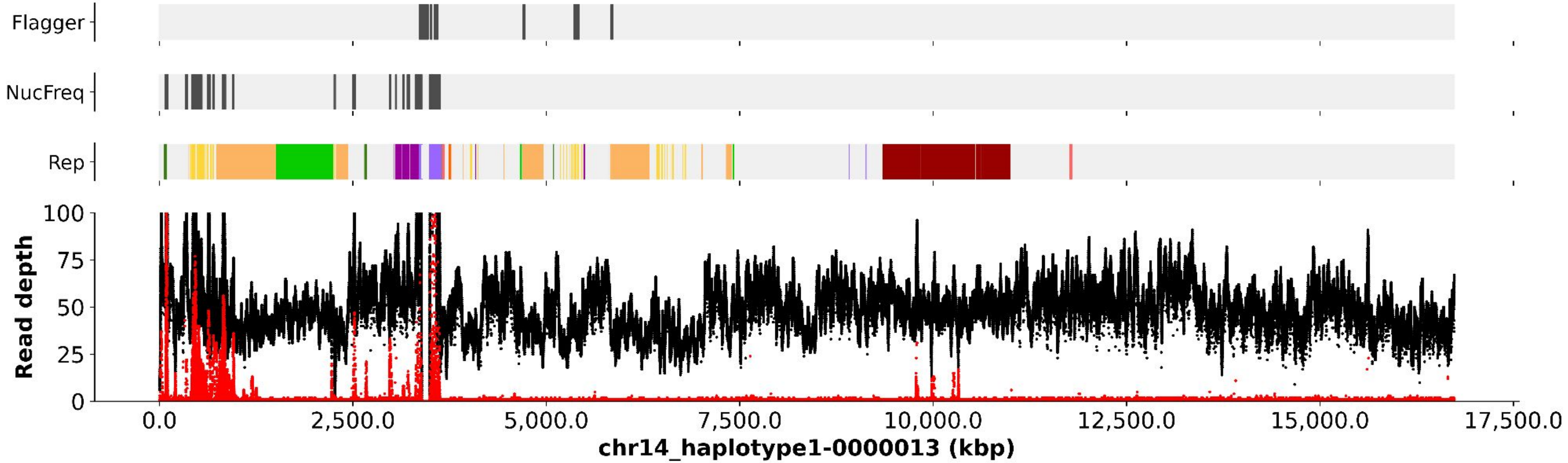

Flagger
NucFreq
Rep
Read depth
100
75
50
25
0
0.0
2,500.0
5,000.0
7,500.0
10,000.0
12,500.0
15,000.0
17,500.0
chr14_haplotype1-0000013 (kbp)


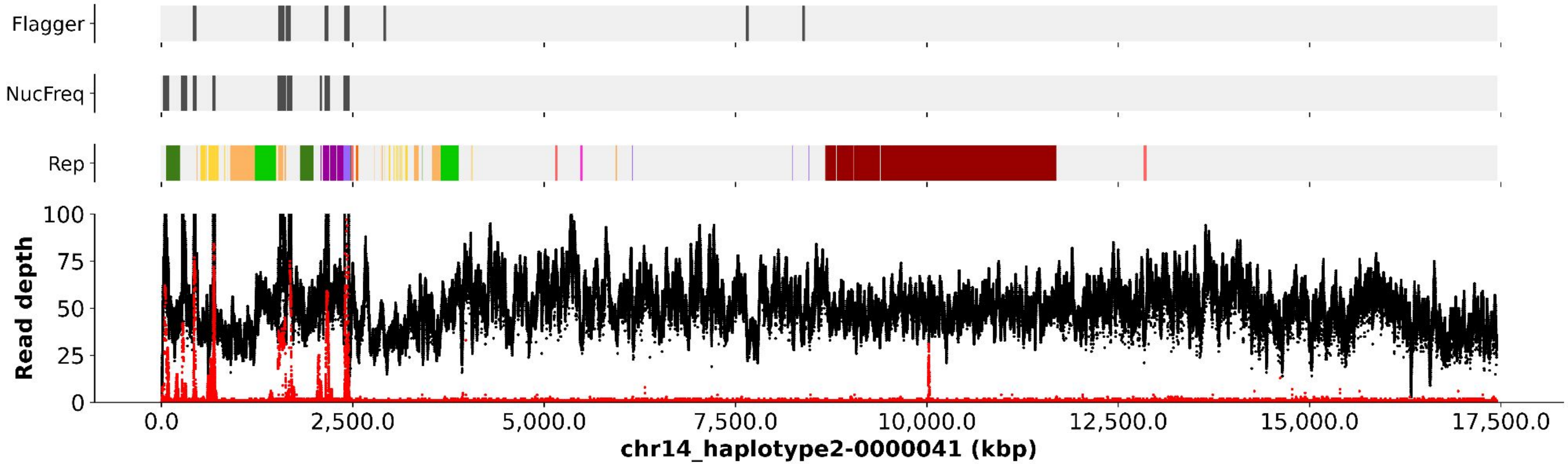

Flagger
NucFreq
Rep
Read depth
100
75
50
25
0
0.0
2,500.0
5,000.0
7,500.0
10,000.0
12,500.0
15,000.0
17,500.0
chr14_haplotype2-0000041 (kbp)

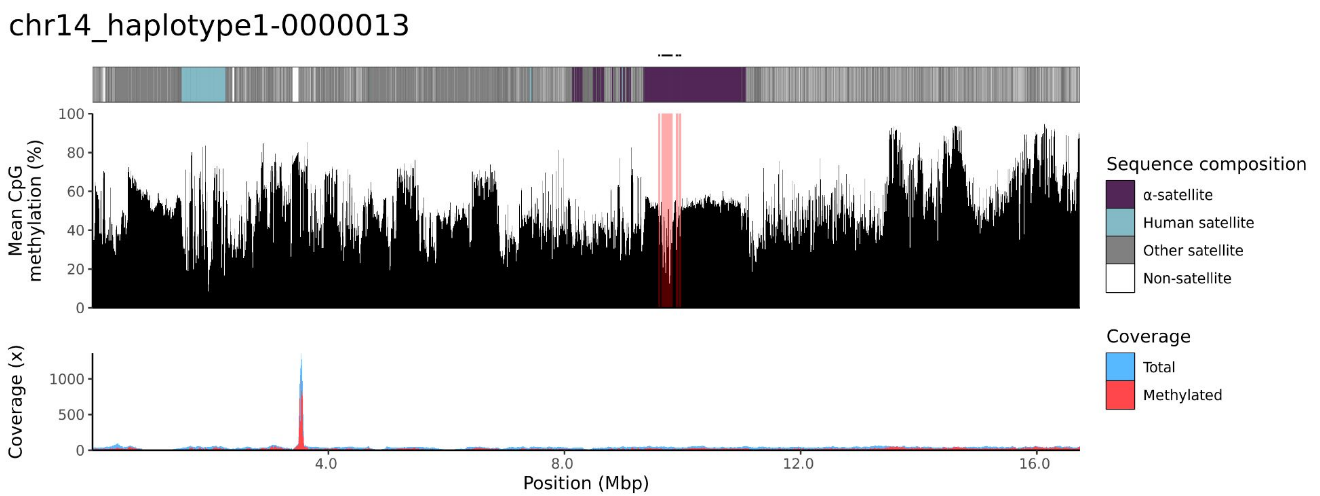
chr14_haplotype1-0000013
Mean CpG methylation (%)
100
80
60
40
20
0
Coverage (x)
1000
500
0
4.0
8.0
12.0
16.0
Position (Mbp)
Sequence composition
α-satellite
Human satellite
Other satellite
Non-satellite
Coverage
Total
Methylated

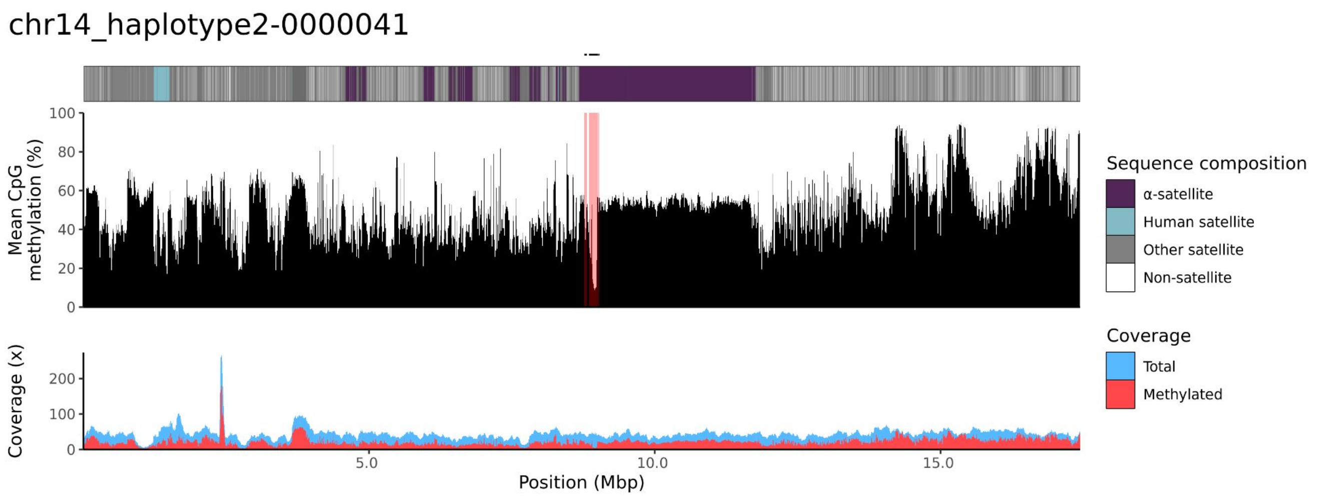
chr14_haplotype2-0000041
Mean CpG methylation (%)
100
80
60
40
20
0
Coverage (x)
200
100
0
5.0
10.0
15.0
Position (Mbp)
Sequence composition
α-satellite
Human satellite
Other satellite
Non-satellite
Coverage
Total
Methylated

### chr15

#### NA12878_1_haplotype1-0000007_chr15

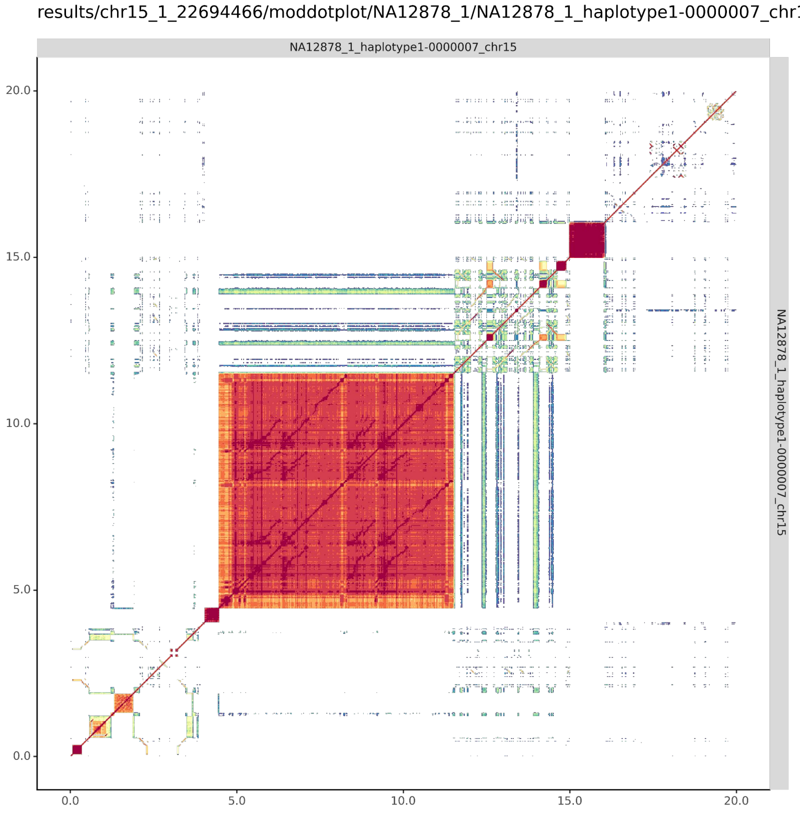


#### NA12878_2_haplotype2-0000044_chr15

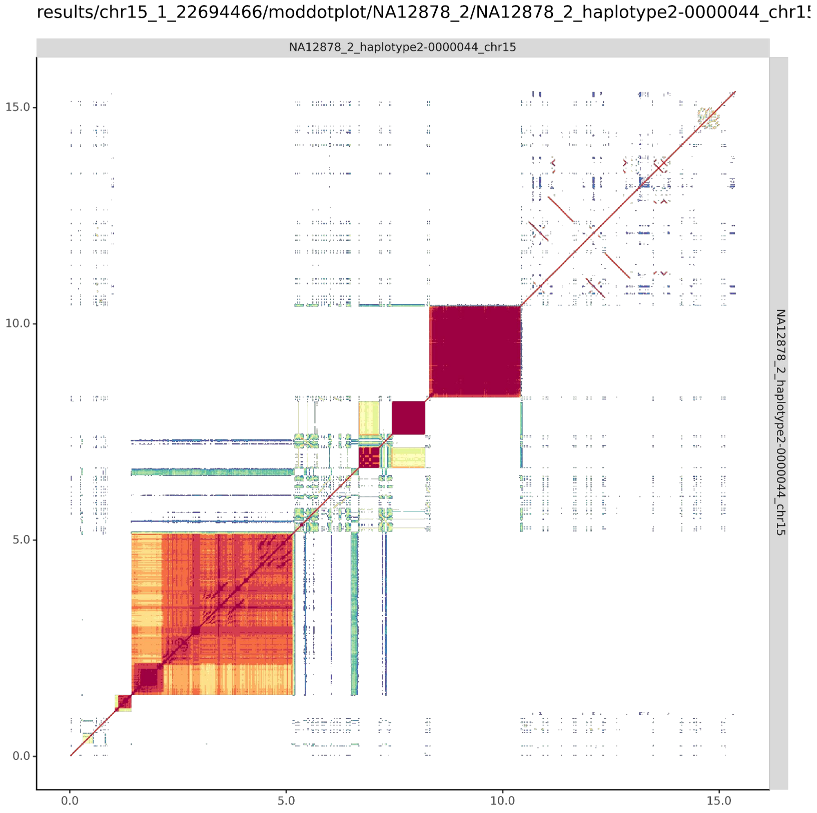

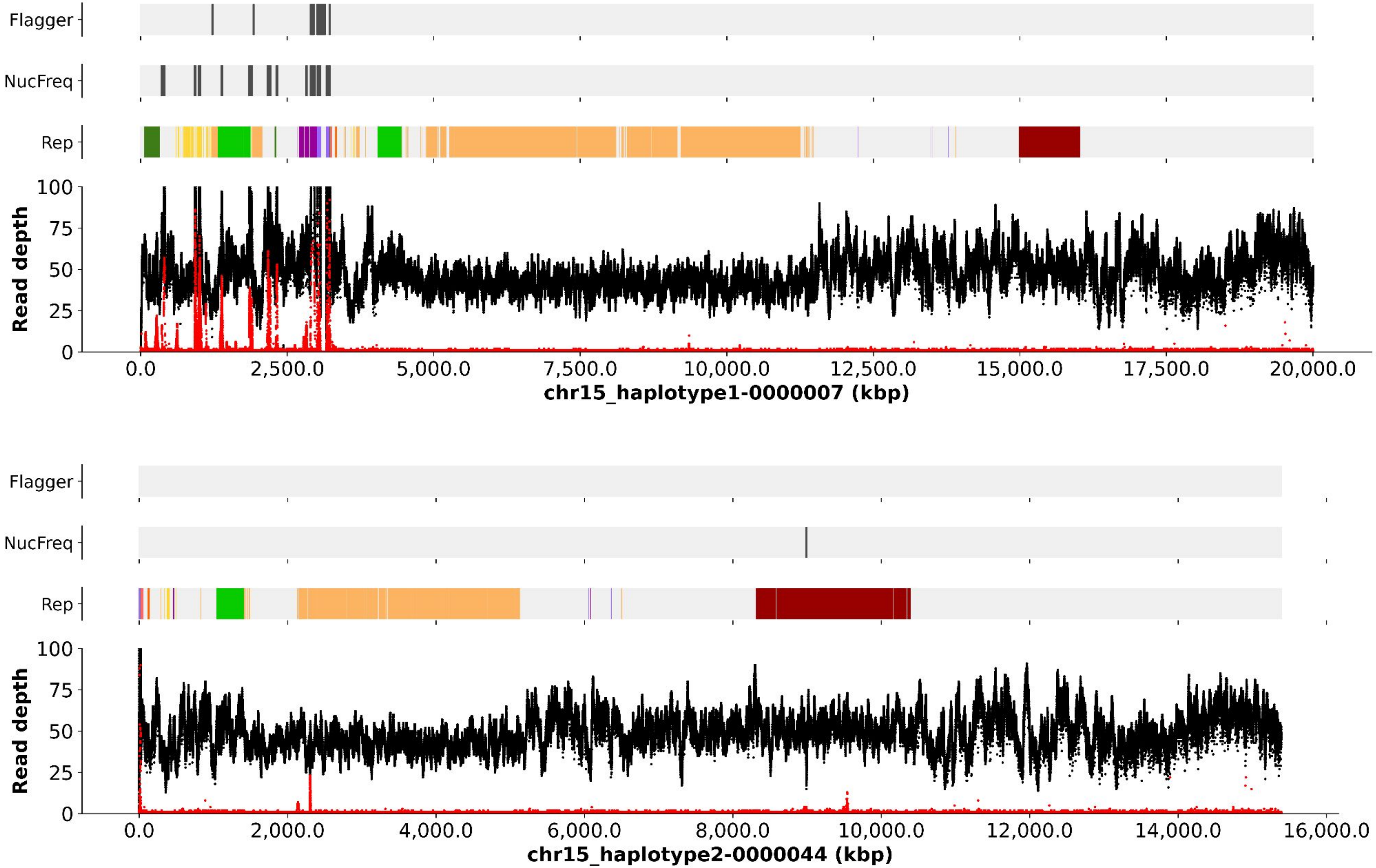
Flagger
NucFreq
Rep
Read depth
100
75
50
25
0
0.0
2,500.0
5,000.0
7,500.0
10,000.0
12,500.0
15,000.0
17,500.0
20,000.0
chr15_haplotype1-0000007 (kbp)
Flagger
NucFreq
Rep
Read depth
100
75
50
25
0
0.0
2,000.0
4,000.0
6,000.0
8,000.0
10,000.0
12,000.0
14,000.0
16,000.0
chr15_haplotype2-0000044 (kbp)

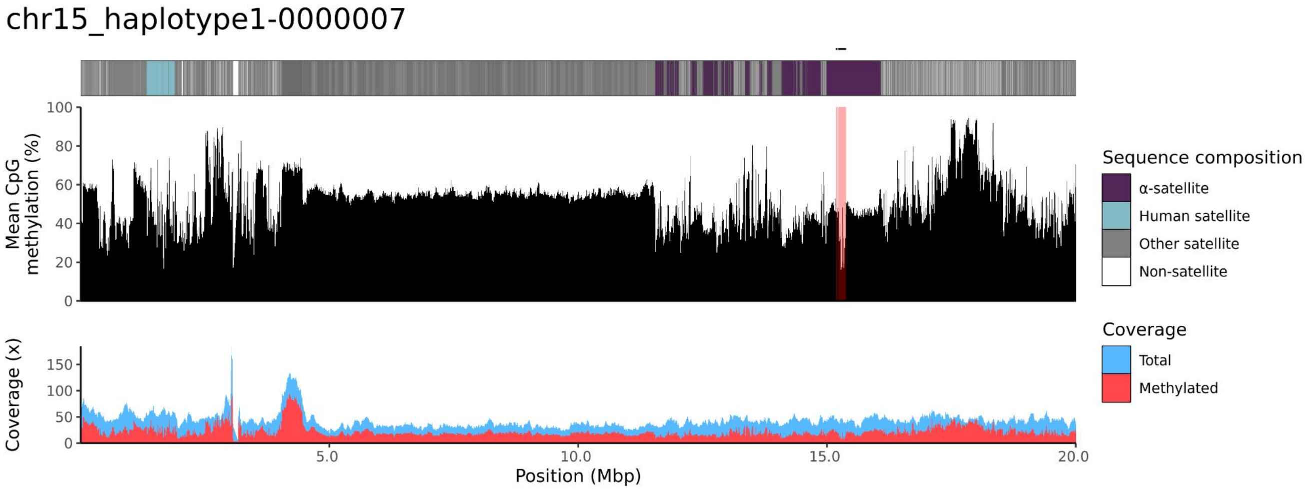
chr15_haplotype1-0000007
Mean CpG methylation (%)
0
20
40
60
80
100
Coverage (x)
0
50
100
150
5.0
10.0
15.0
20.0
Position (Mbp)
Sequence composition
α-satellite
Human satellite
Other satellite
Non-satellite
Coverage
Total
Methylated

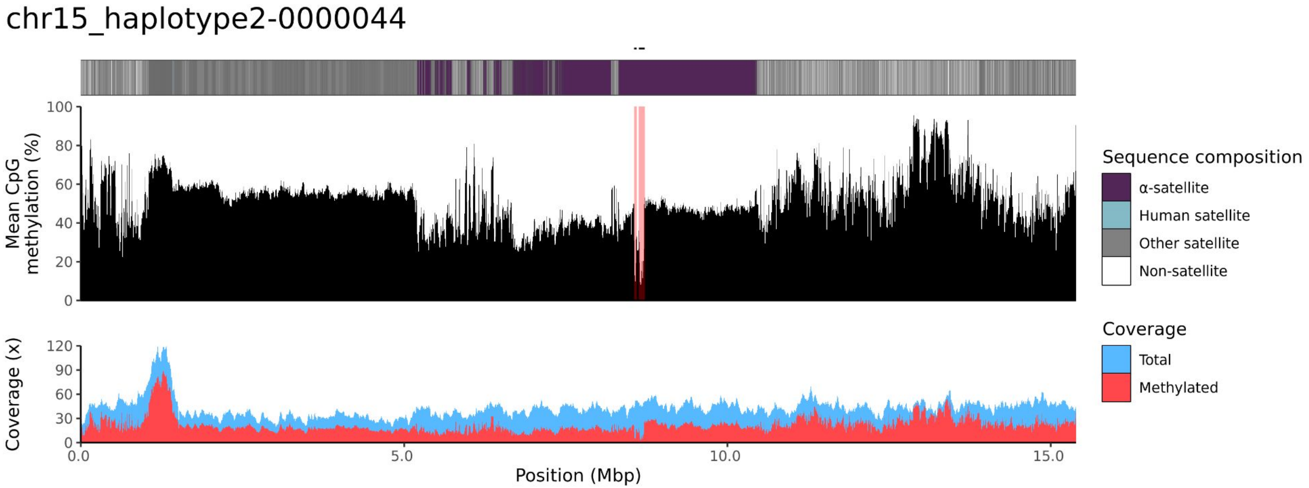
chr15_haplotype2-0000044
Mean CpG methylation (%)
0
20
40
60
80
100
Coverage (x)
0
30
60
90
120
0.0
5.0
10.0
15.0
Position (Mbp)
Sequence composition
α-satellite
Human satellite
Other satellite
Non-satellite
Coverage
Total
Methylated

### chr21

#### NA12878_1_haplotype1-0000009_chr21

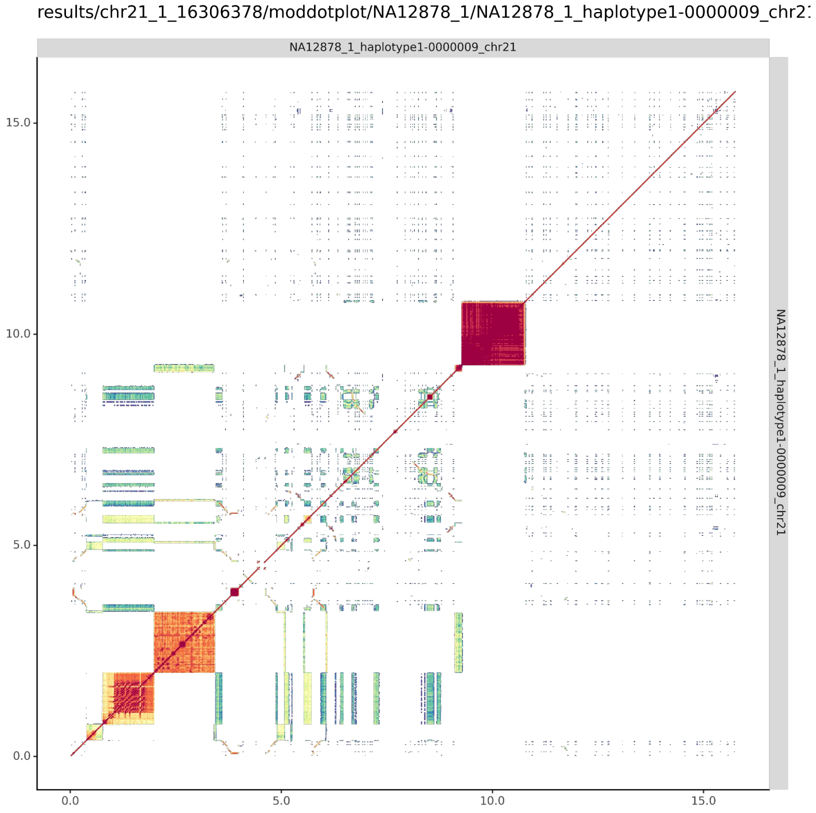


#### NA12878_2_haplotype2-0000032_chr21

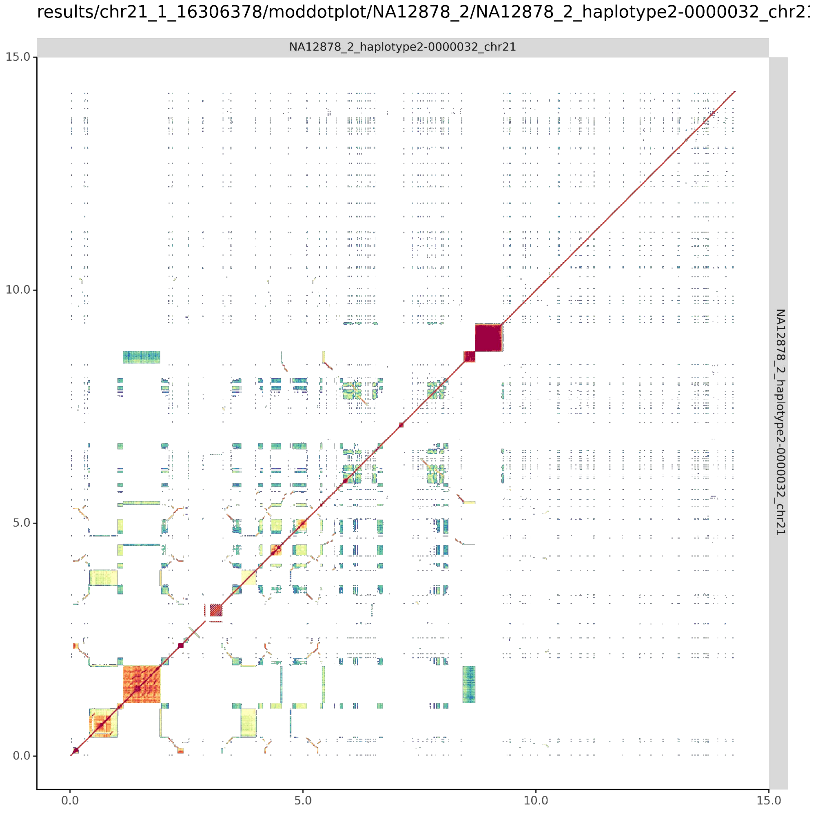

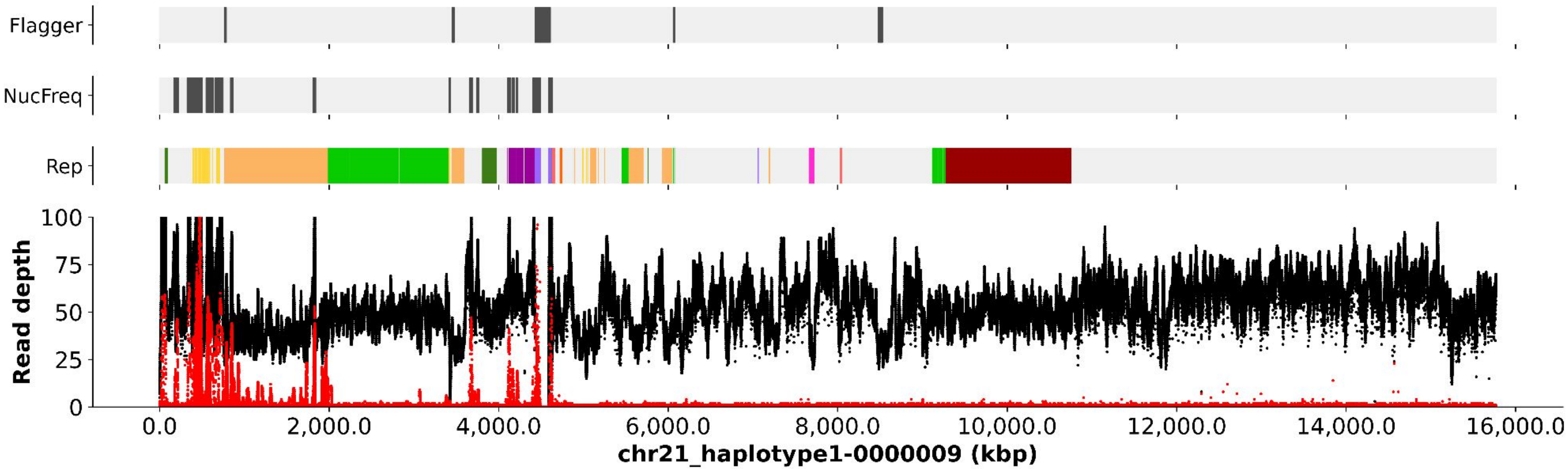
Flagger
NucFreq
Rep
Read depth
100
75
50
25
0
0.0
2,000.0
4,000.0
6,000.0
8,000.0
10,000.0
12,000.0
14,000.0
16,000.0
chr21_haplotype1-0000009 (kbp)

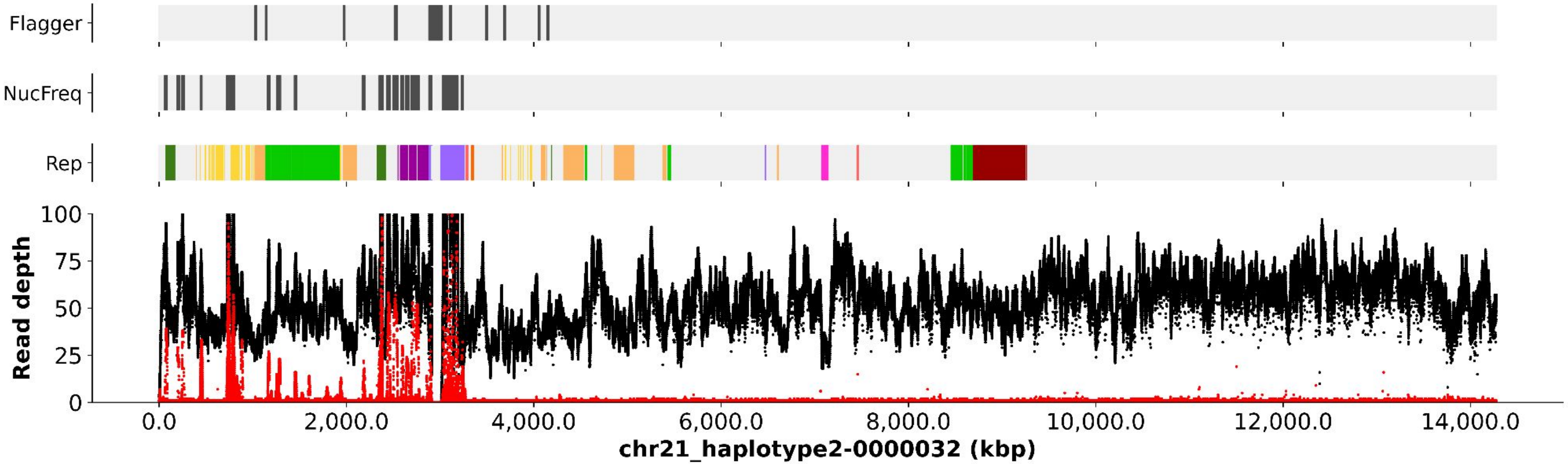
Flagger
NucFreq
Rep
Read depth
100
75
50
25
0
0.0
2,000.0
4,000.0
6,000.0
8,000.0
10,000.0
12,000.0
14,000.0
chr21_haplotype2-0000032 (kbp)

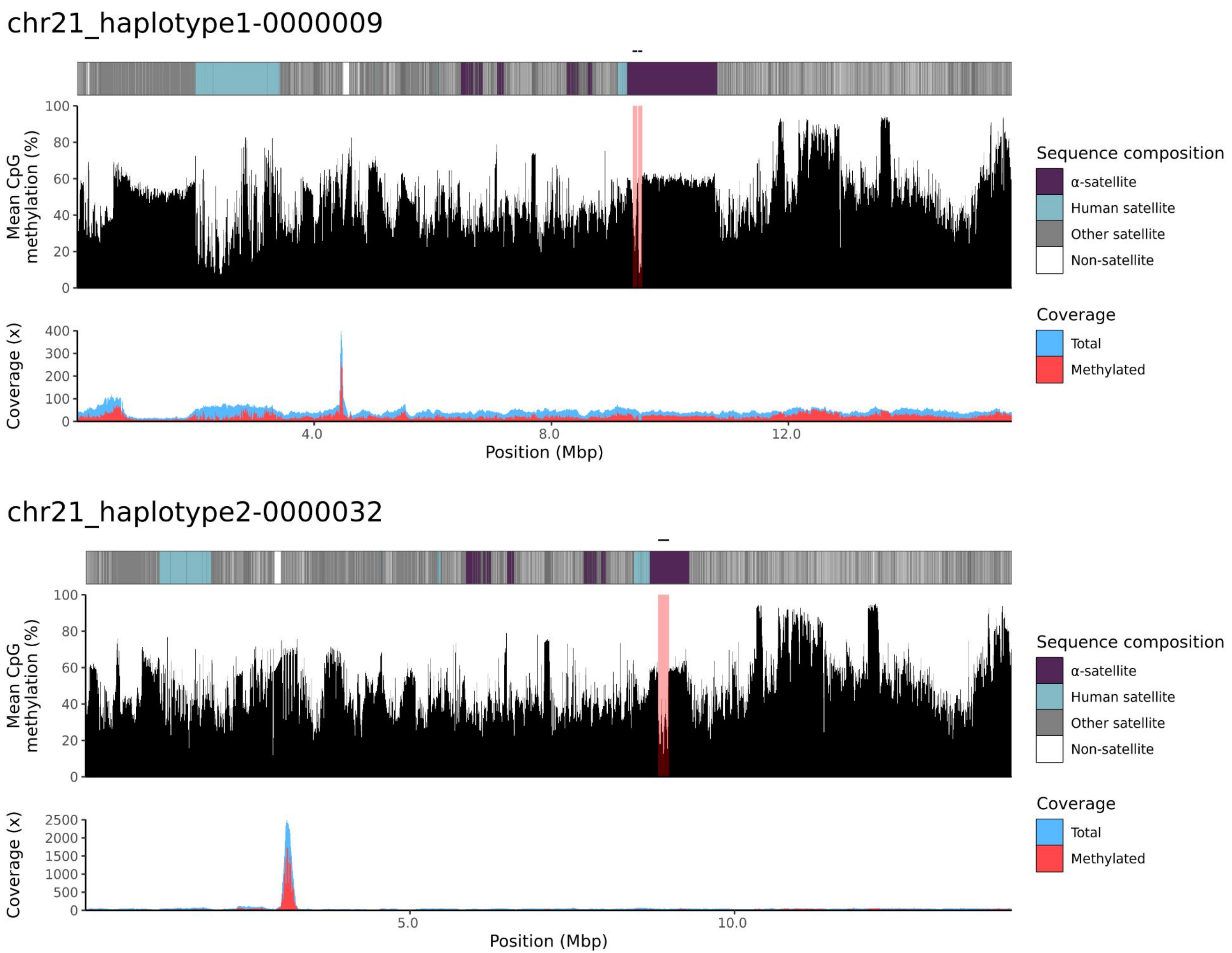

chr21_haplotype1-0000009
Mean CpG methylation (%)
Sequence composition
α-satellite
Human satellite
Other satellite
Non-satellite
Coverage (x)
Coverage
Total
Methylated
4.0
8.0
12.0
Position (Mbp)
chr21_haplotype2-0000032
Mean CpG methylation (%)
Sequence composition
α-satellite
Human satellite
Other satellite
Non-satellite
Coverage (x)
Coverage
Total
Methylated
5.0
10.0
Position (Mbp)

### chr22

#### NA12878_1_haplotype1-0000008_chr22

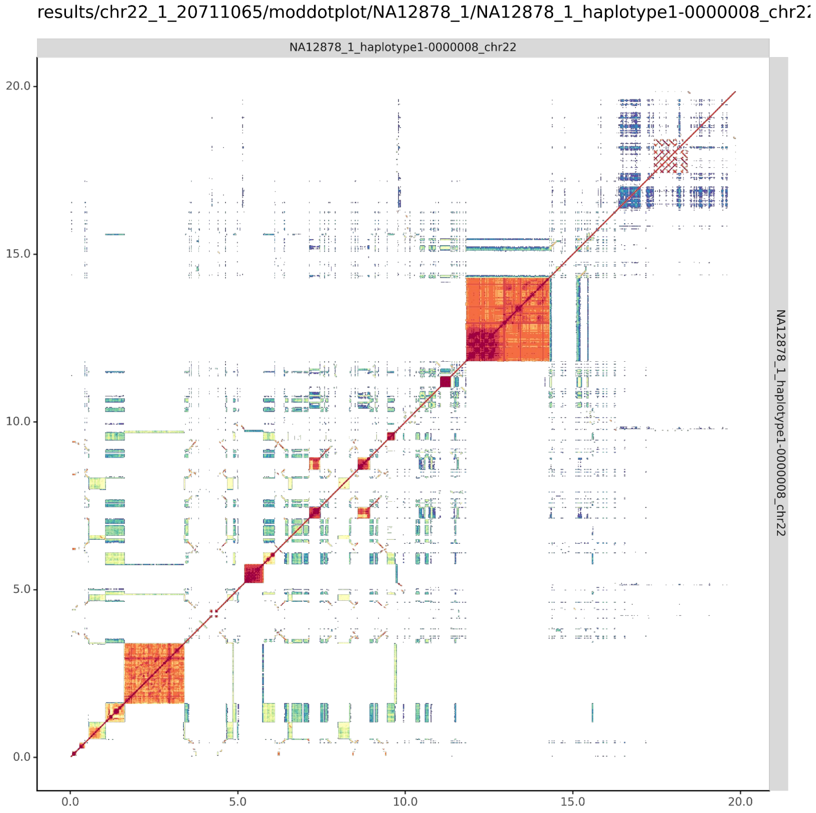


#### NA12878_2_haplotype2-0000037_chr22

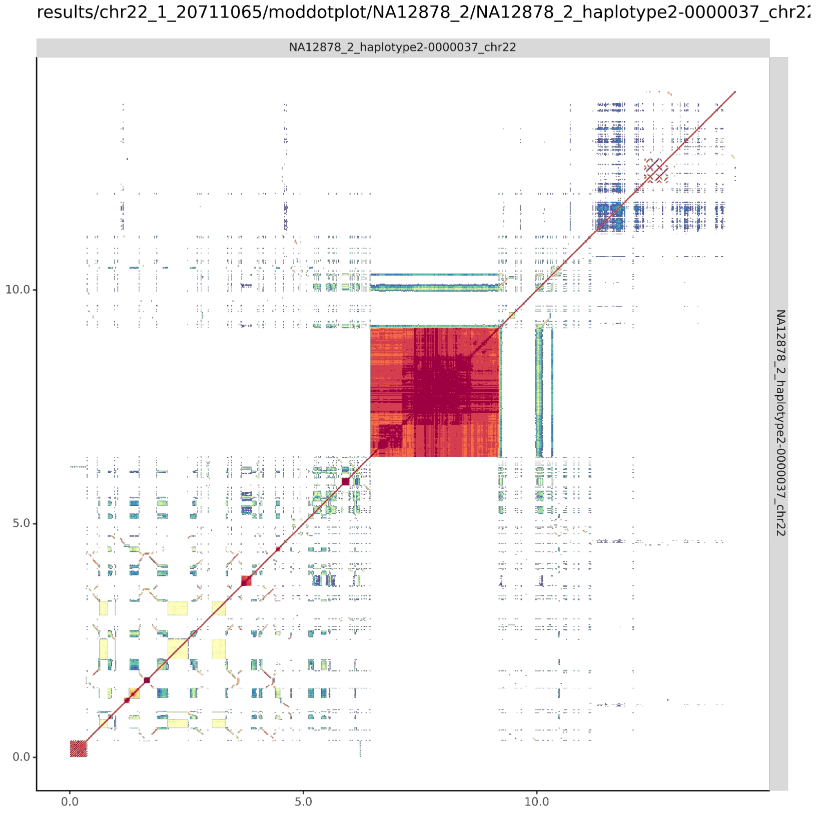

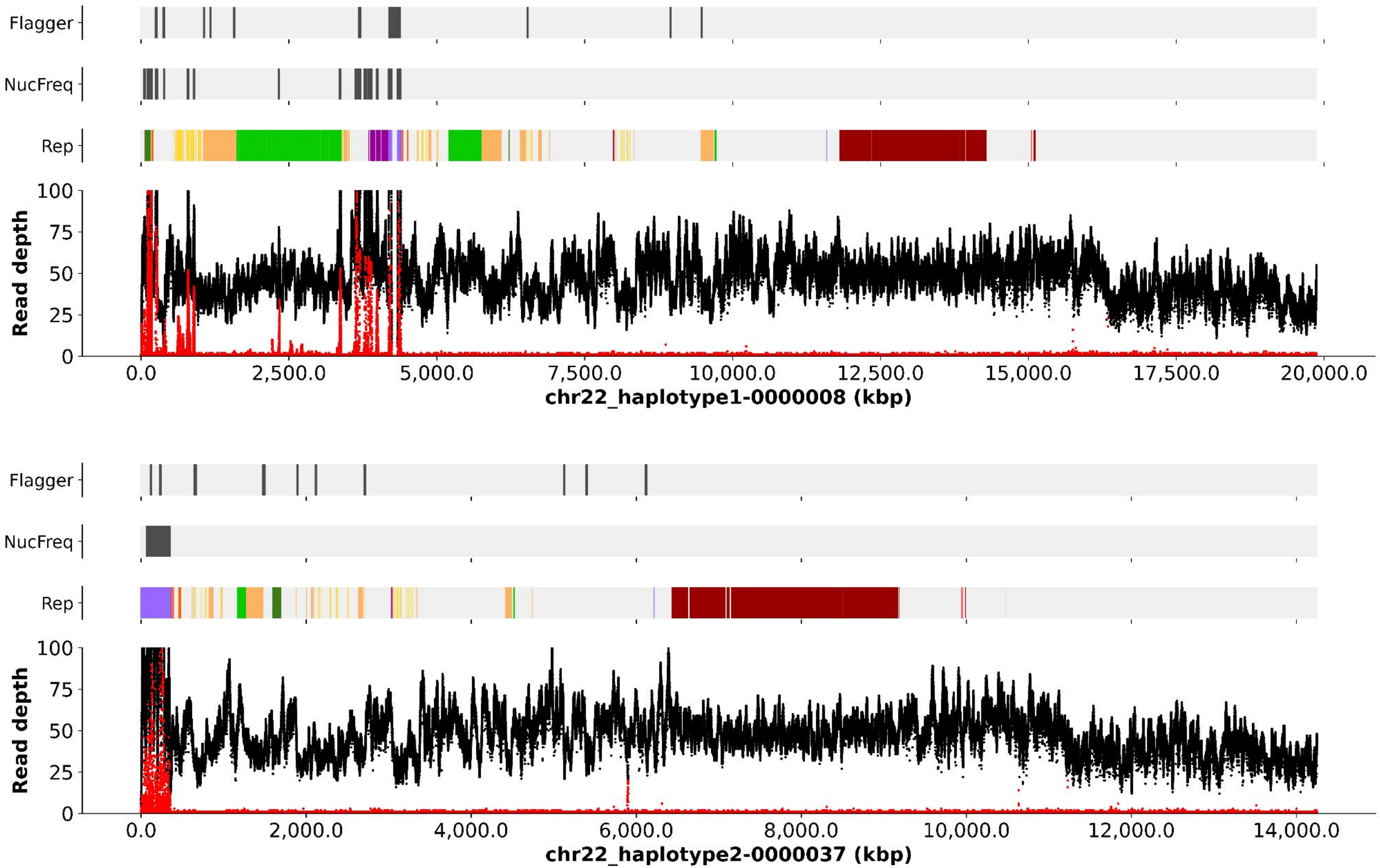

Flagger
NucFreq
Rep
Read depth
0
25
50
75
100
0.0
2,500.0
5,000.0
7,500.0
10,000.0
12,500.0
15,000.0
17,500.0
20,000.0
chr22_haplotype1-0000008 (kbp)
Flagger
NucFreq
Rep
Read depth
0
25
50
75
100
0.0
2,000.0
4,000.0
6,000.0
8,000.0
10,000.0
12,000.0
14,000.0
chr22_haplotype2-0000037 (kbp)

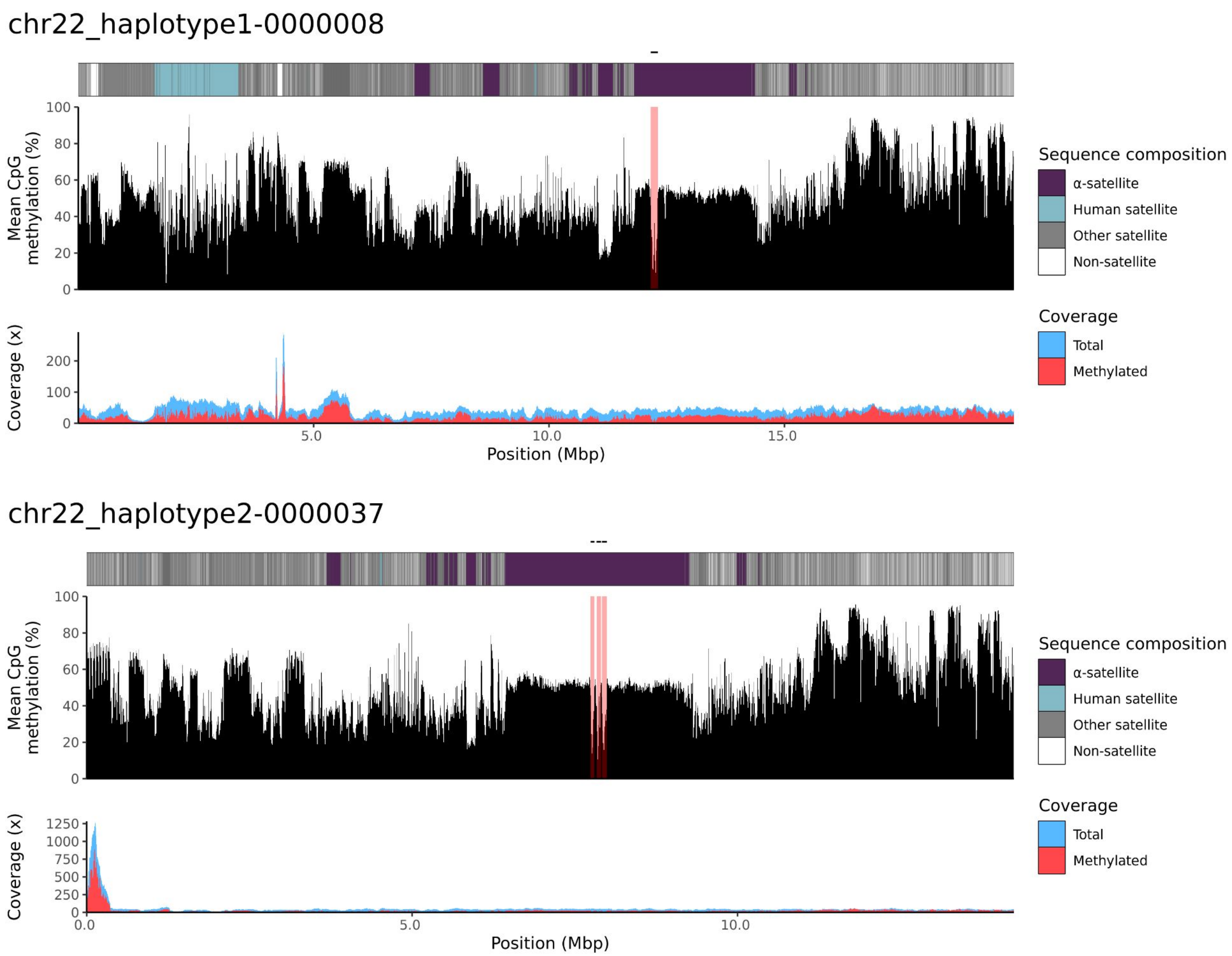
chr22_haplotype1-0000008
Mean CpG methylation (%)
100
80
60
40
20
0
Coverage (x)
200
100
0
5.0
10.0
15.0
Position (Mbp)
Sequence composition
α-satellite
Human satellite
Other satellite
Non-satellite
Coverage
Total
Methylated
chr22_haplotype2-0000037
Mean CpG methylation (%)
100
80
60
40
20
0
Coverage (x)
1250
1000
750
500
250
0
0.0
5.0
10.0
Position (Mbp)
Sequence composition
α-satellite
Human satellite
Other satellite
Non-satellite
Coverage
Total
Methylated

NA12879

### chr13

#### NA12879_1_haplotype1-0000002_chr13

results/chr13_1_22508596/moddotplot/NA12879_1/NA12879_1_haplotype1-0000002_chr1:

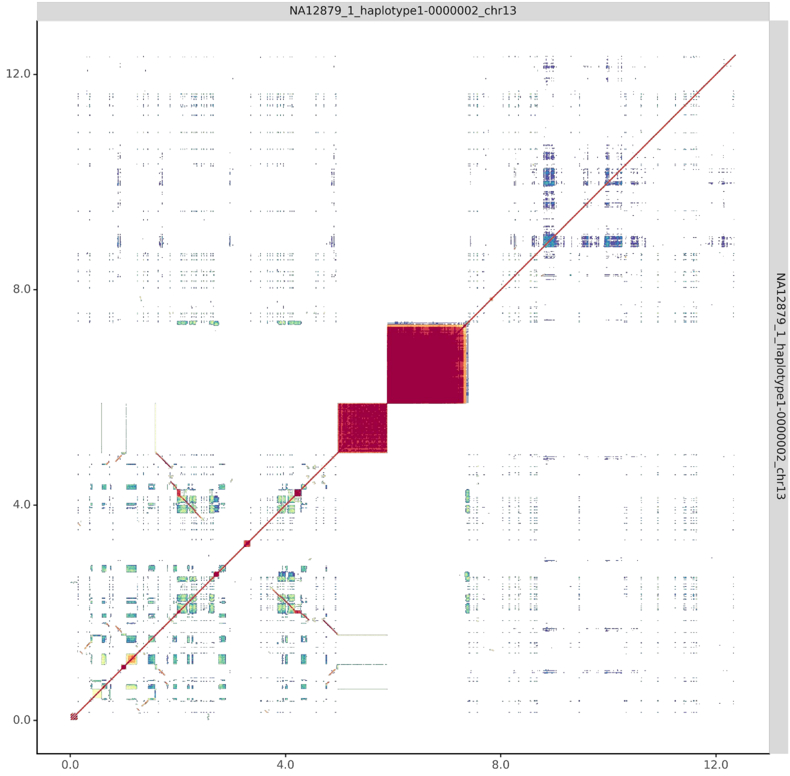


#### NA12879_2_haplotype2-0000092_chr13

results/chr13_1_22508596/moddotplot/NA12879_2/NA12879_2_haplotype2-0000092_chr1:

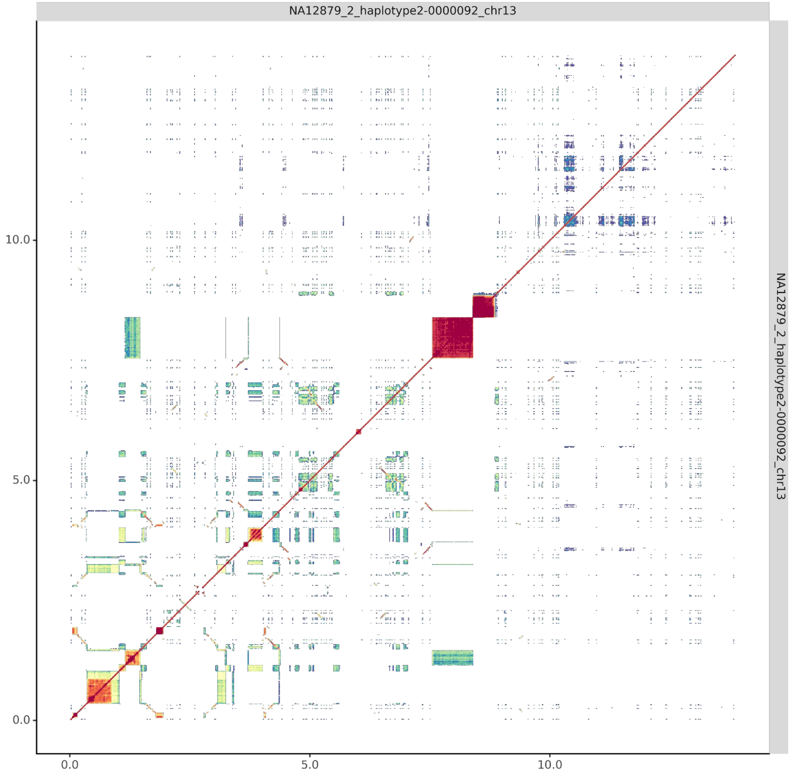

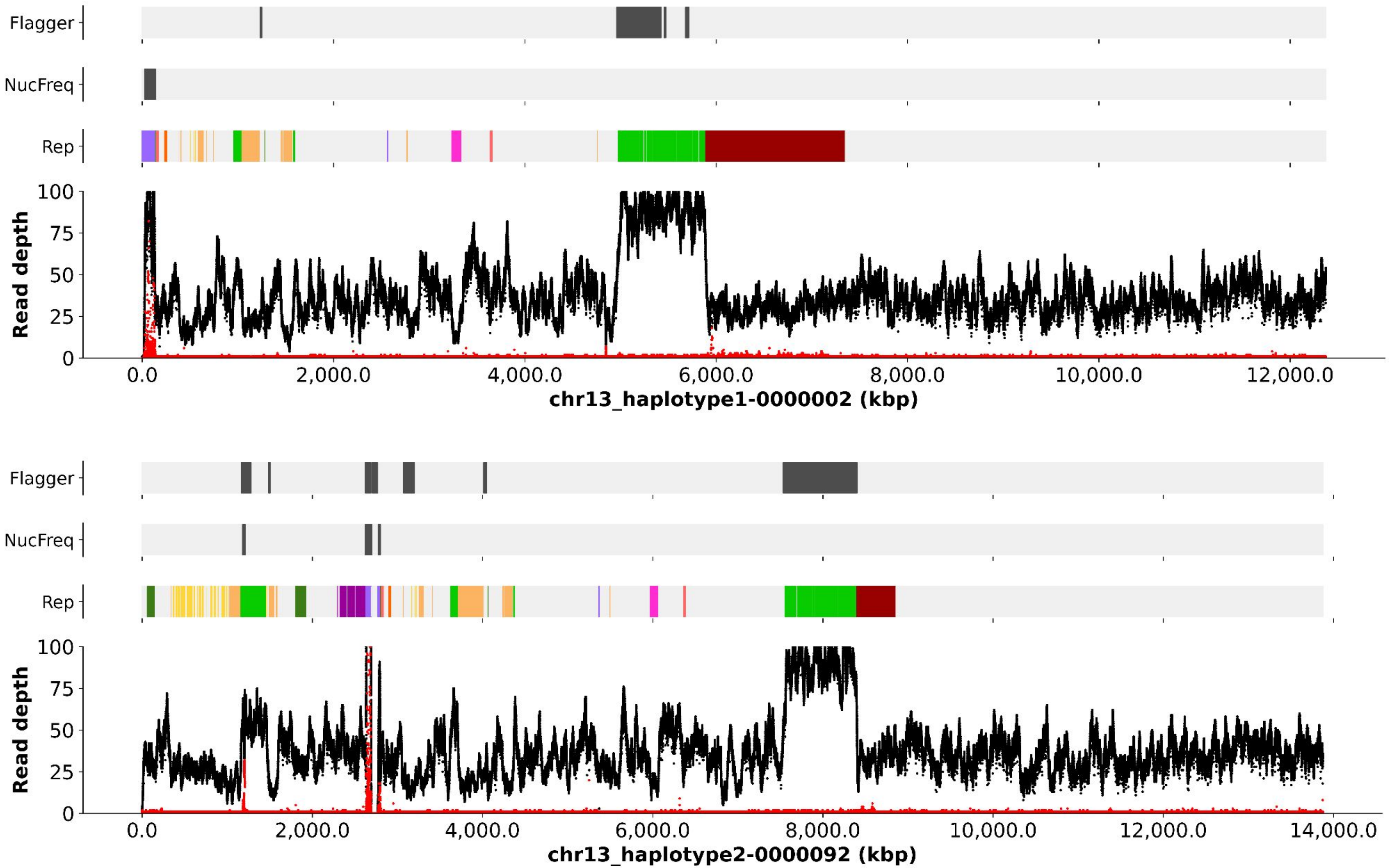

Flagger
NucFreq
Rep
Read depth
100
75
50
25
0
0.0
2,000.0
4,000.0
6,000.0
8,000.0
10,000.0
12,000.0
chr13_haplotype1-0000002 (kbp)
Flagger
NucFreq
Rep
Read depth
100
75
50
25
0
0.0
2,000.0
4,000.0
6,000.0
8,000.0
10,000.0
12,000.0
14,000.0
chr13_haplotype2-0000092 (kbp)

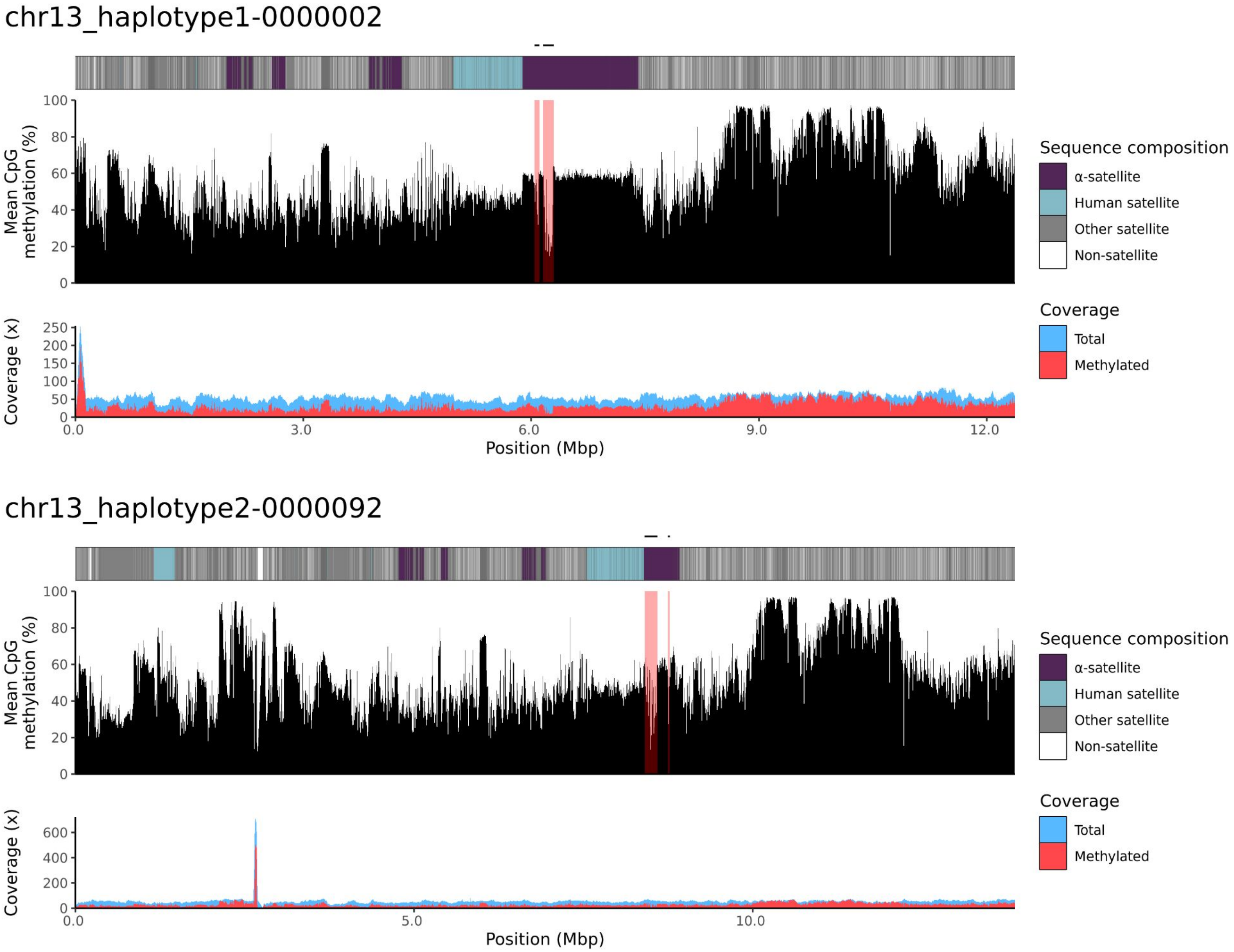
chr13_haplotype1-0000002
Mean CpG methylation (%)
100
80
60
40
20
0
Coverage (x)
250
200
150
100
50
0
0.0
3.0
6.0
9.0
12.0
Position (Mbp)
Sequence composition
α-satellite
Human satellite
Other satellite
Non-satellite
Coverage
Total
Methylated
chr13_haplotype2-0000092
Mean CpG methylation (%)
100
80
60
40
20
0
Coverage (x)
600
400
200
0
0.0
5.0
10.0
Position (Mbp)
Sequence composition
α-satellite
Human satellite
Other satellite
Non-satellite
Coverage
Total
Methylated

### chr14

#### NA12879_1_haplotype1-0000015_chr14

results/chr14_1_17708411/moddotplot/NA12879_1/NA12879_1_haplotype1-0000015_chr14

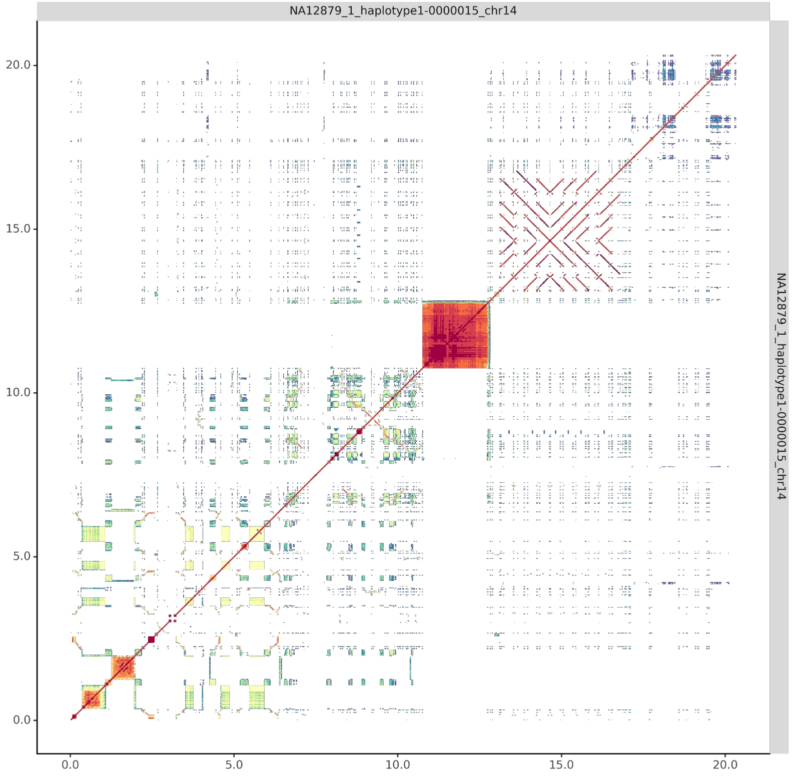


#### NA12879_2_haplotype2-0000083_chr14

results/chr14_1_17708411/moddotplot/NA12879_2/NA12879_2_haplotype2-0000083_chr14

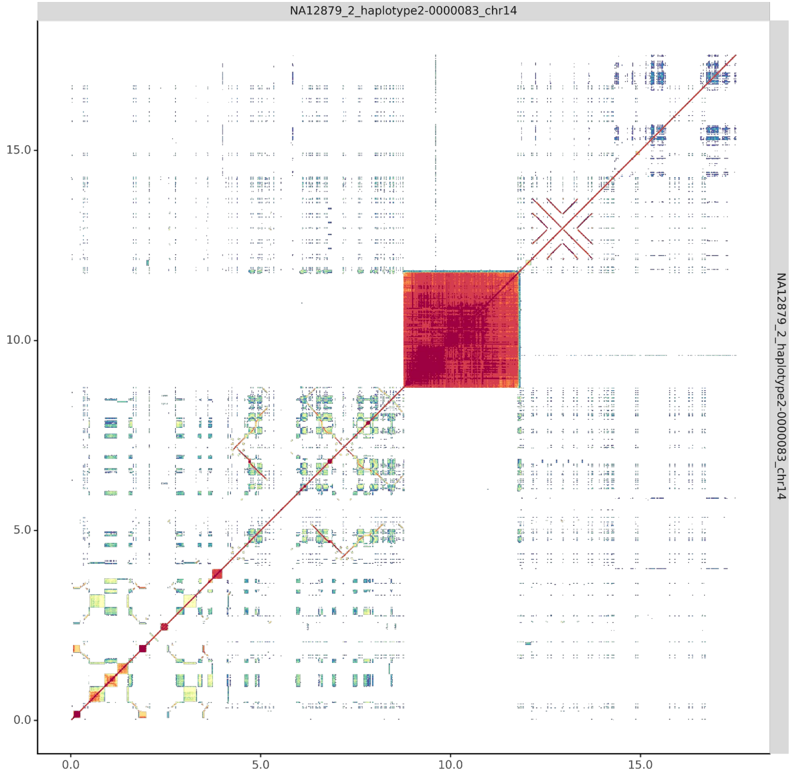

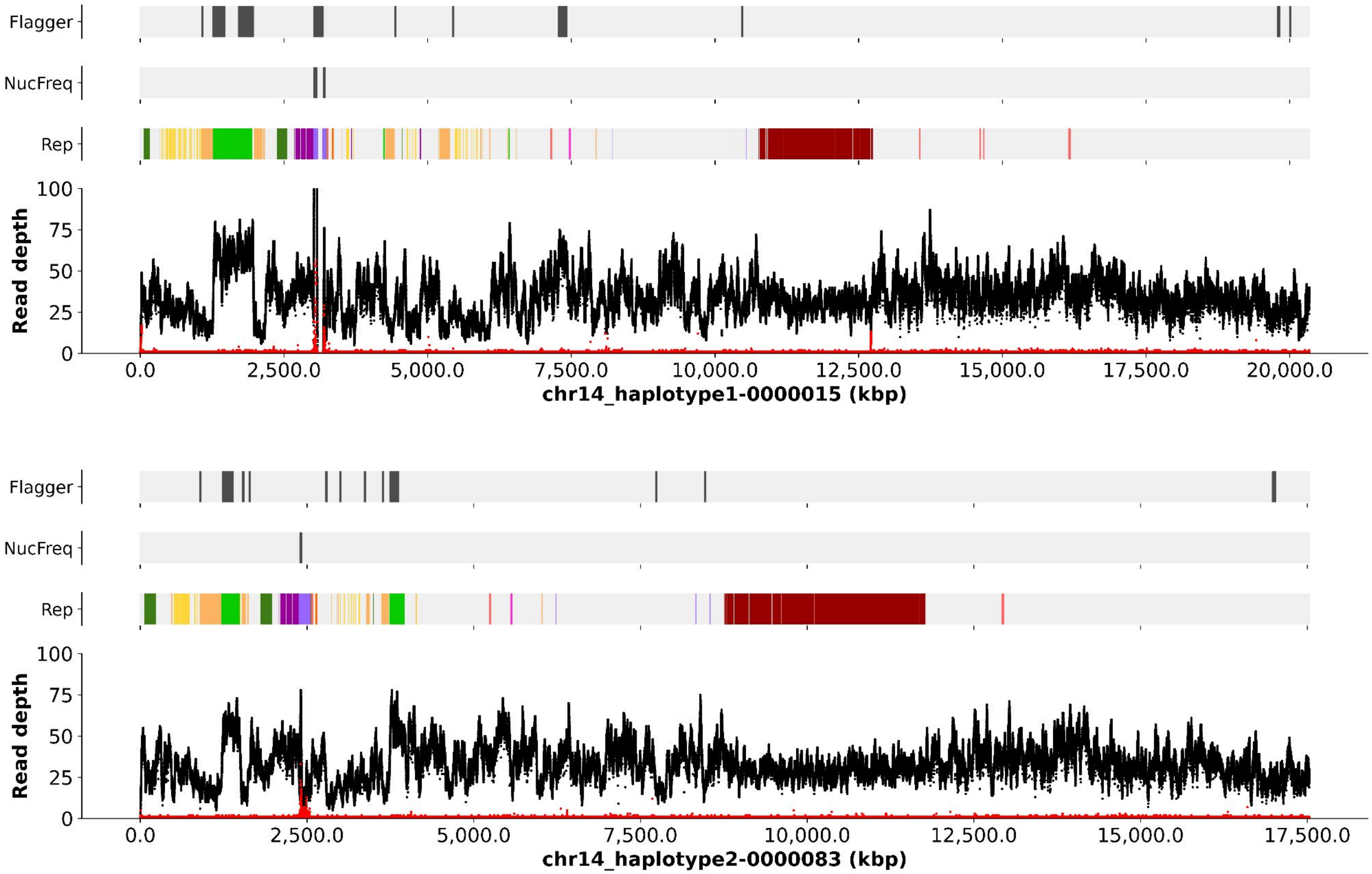
Flagger
NucFreq
Rep
Read depth
100
75
50
25
0
0.0
2,500.0
5,000.0
7,500.0
10,000.0
12,500.0
15,000.0
17,500.0
20,000.0
chr14_haplotype1-0000015 (kbp)
Flagger
NucFreq
Rep
Read depth
chr14_haplotype2-0000083 (kbp)

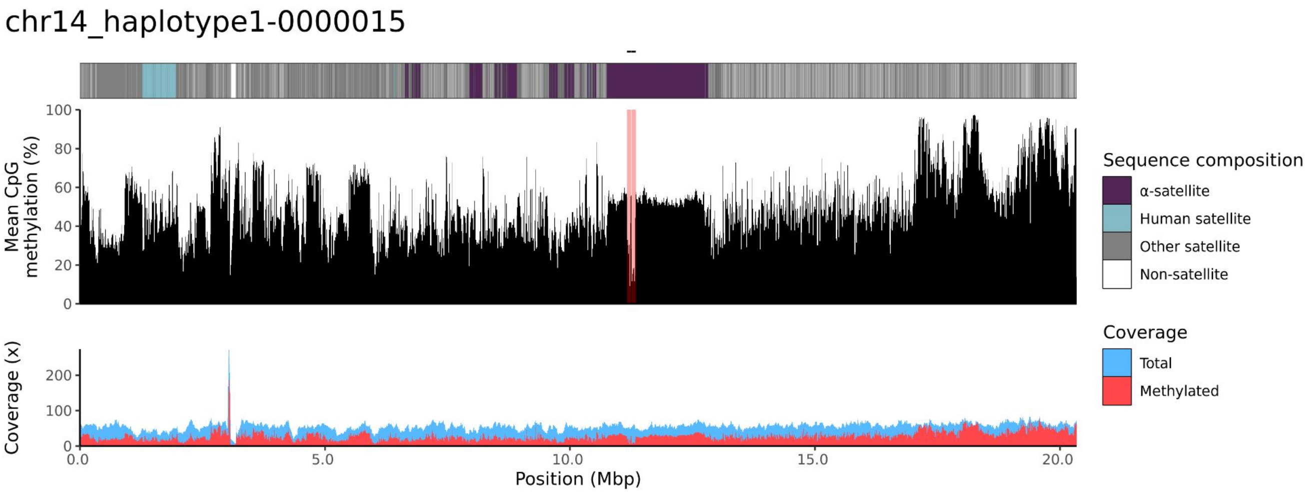
chr14_haplotype1-0000015
Mean CpG methylation (%)
Coverage (x)
Position (Mbp)
Sequence composition
α-satellite
Human satellite
Other satellite
Non-satellite
Coverage
Total
Methylated

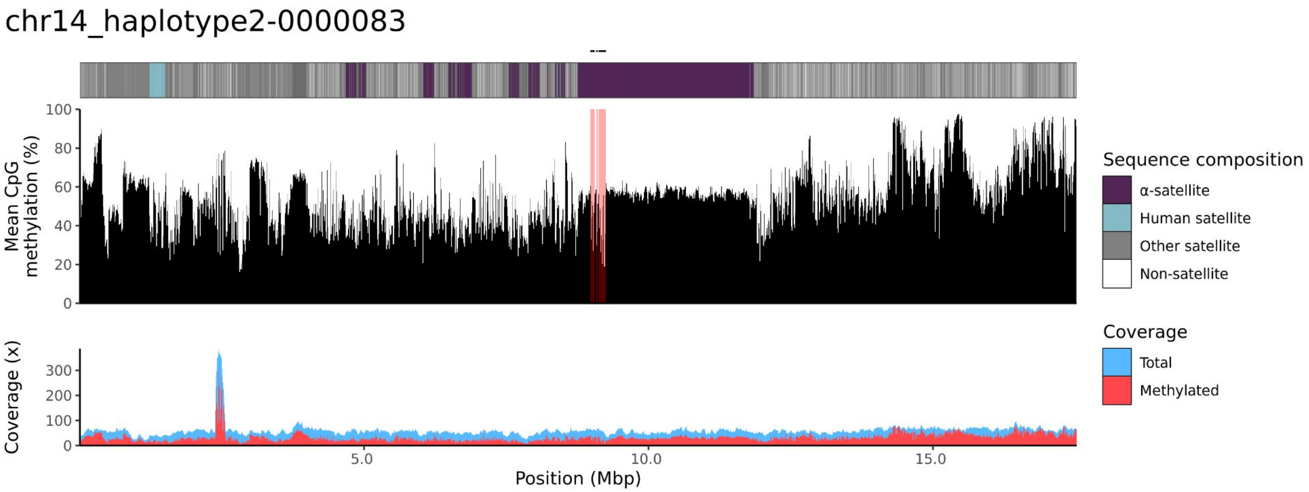
chr14_haplotype2-0000083
Mean CpG methylation (%)
Coverage (x)
Position (Mbp)
Sequence composition
α-satellite
Human satellite
Other satellite
Non-satellite
Coverage
Total
Methylated

### chr15

#### NA12879_1_haplotype1-0000013_chr15

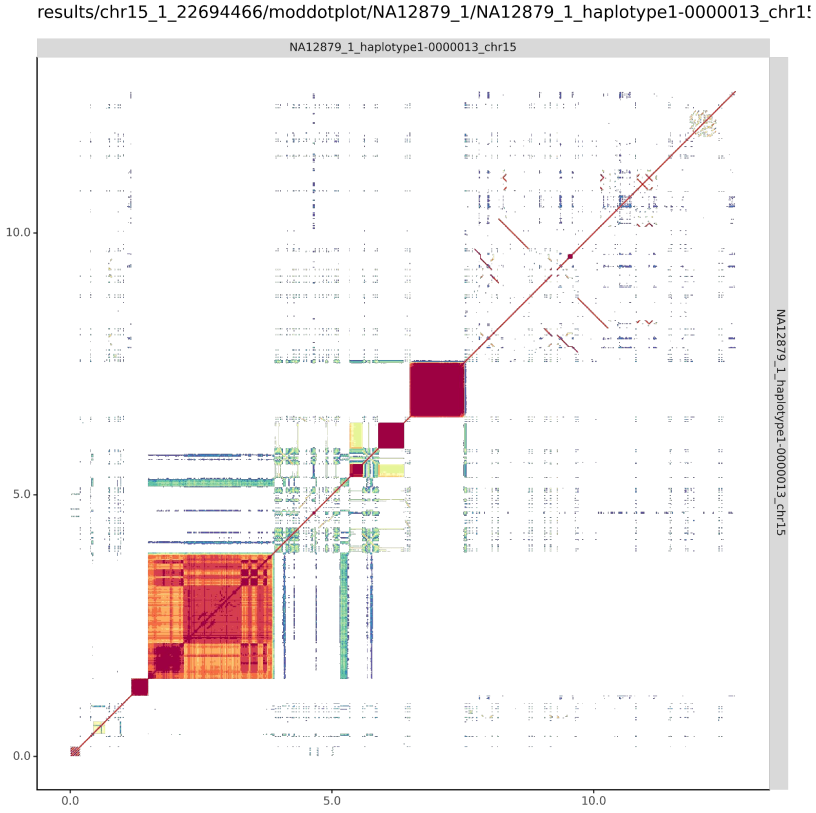


#### NA12879_2_haplotype2-0000075_chr15

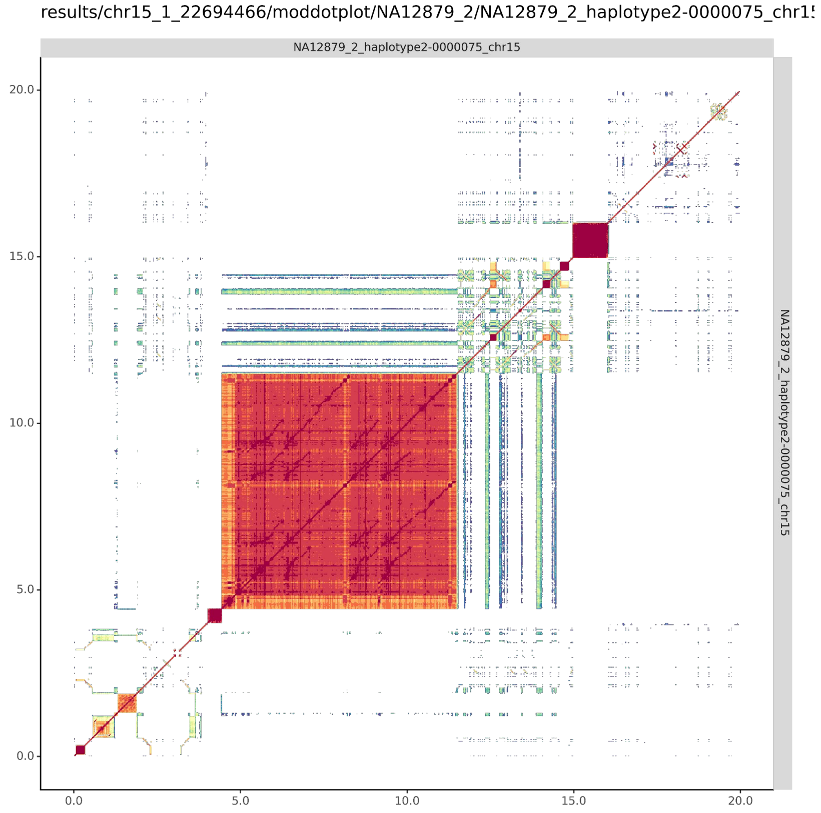

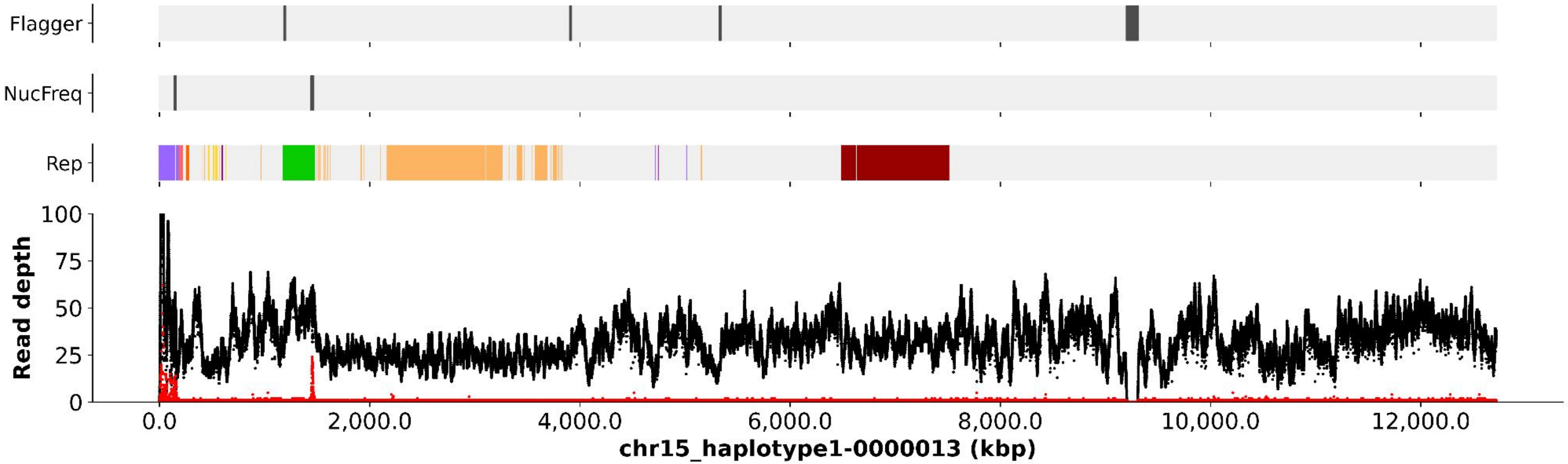
Flagger
NucFreq
Rep
Read depth
100
75
50
25
0
0.0
2,000.0
4,000.0
6,000.0
8,000.0
10,000.0
12,000.0
chr15_haplotype1-0000013 (kbp)

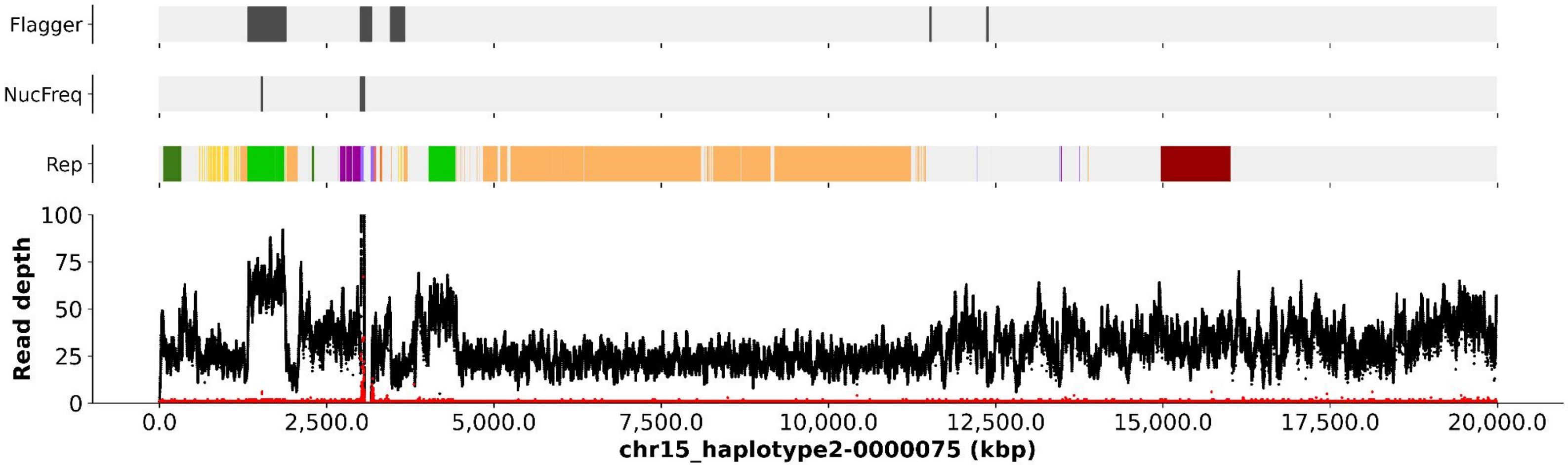
Flagger
NucFreq
Rep
Read depth
100
75
50
25
0
0.0
2,500.0
5,000.0
7,500.0
10,000.0
12,500.0
15,000.0
17,500.0
20,000.0
chr15_haplotype2-0000075 (kbp)

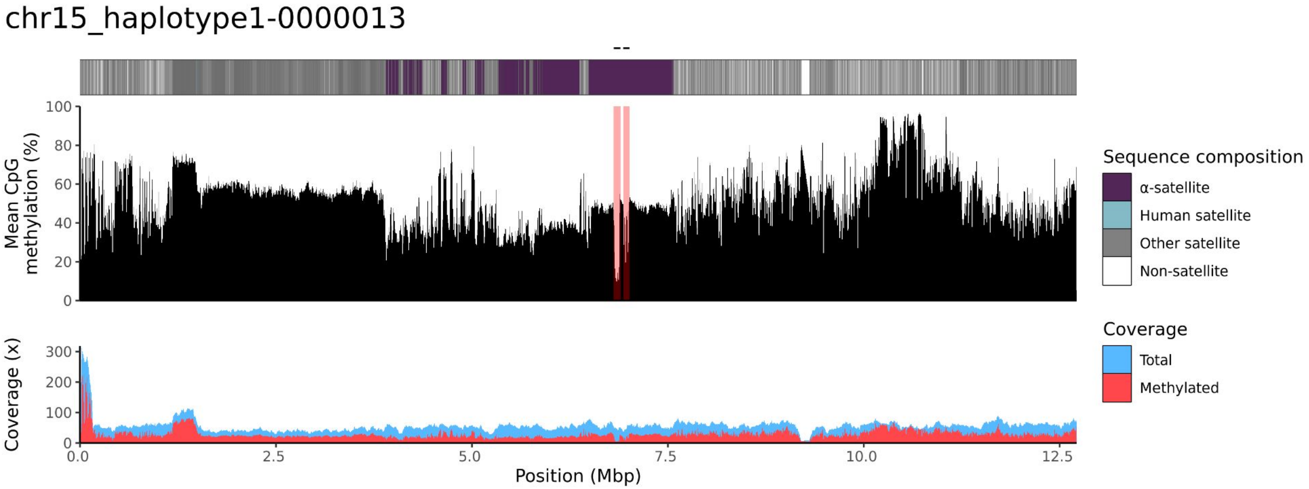
chr15_haplotype1-0000013
Mean CpG methylation (%)
100
80
60
40
20
0
Coverage (x)
300
200
100
0
0.0
2.5
5.0
7.5
10.0
12.5
Position (Mbp)
Sequence composition
α-satellite
Human satellite
Other satellite
Non-satellite
Coverage
Total
Methylated

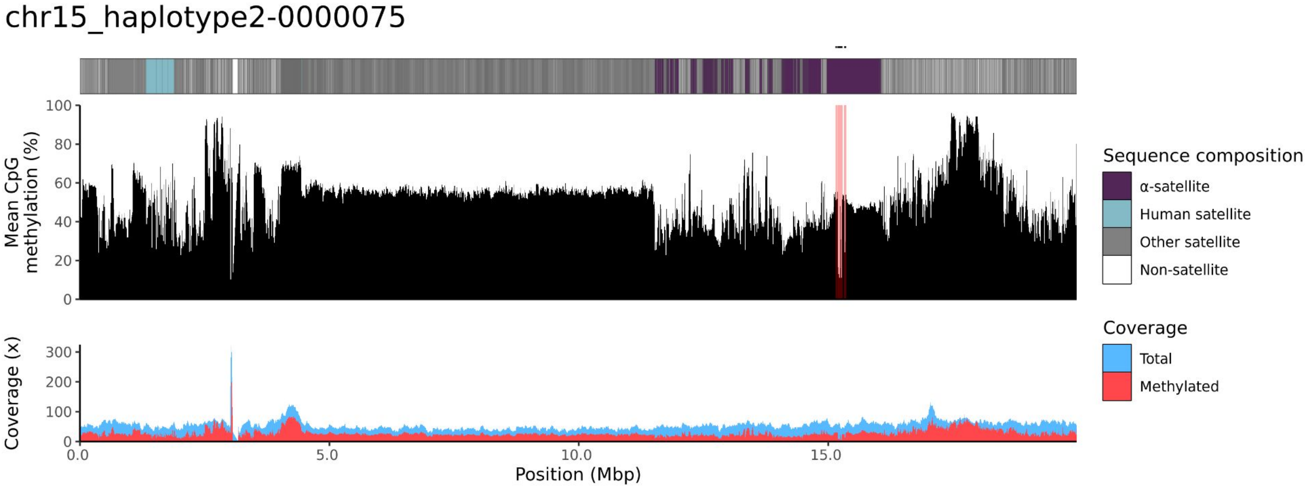
chr15_haplotype2-0000075
Mean CpG methylation (%)
100
80
60
40
20
0
Coverage (x)
300
200
100
0
0.0
5.0
10.0
15.0
Position (Mbp)
Sequence composition
α-satellite
Human satellite
Other satellite
Non-satellite
Coverage
Total
Methylated

### chr21

#### NA12879_1_haplotype1-0000019_chr21

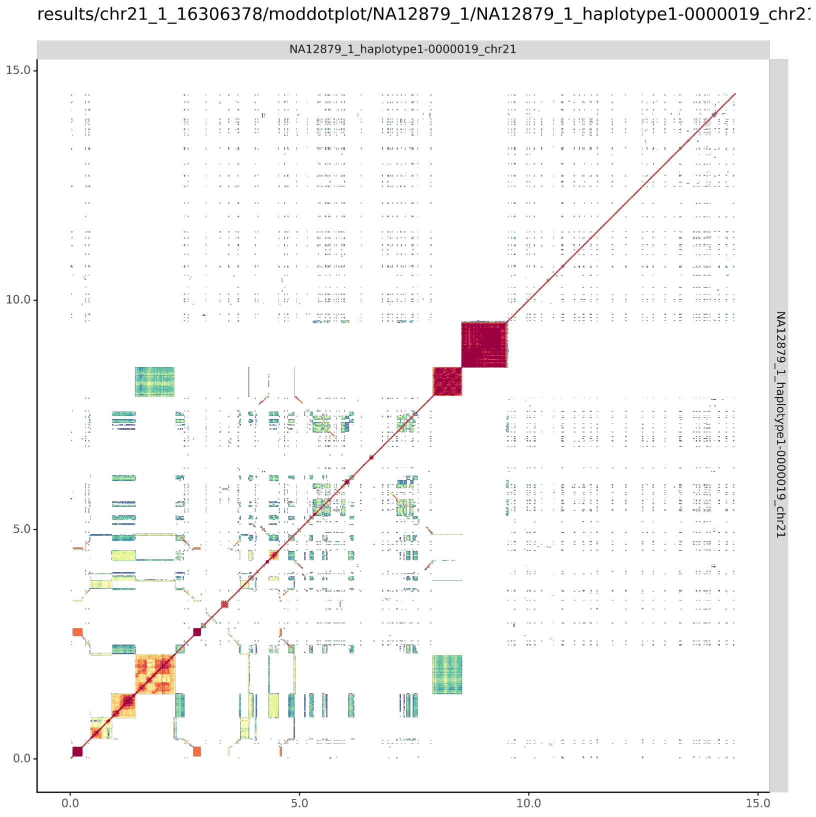


#### NA12879_2_haplotype2-0000087_chr21

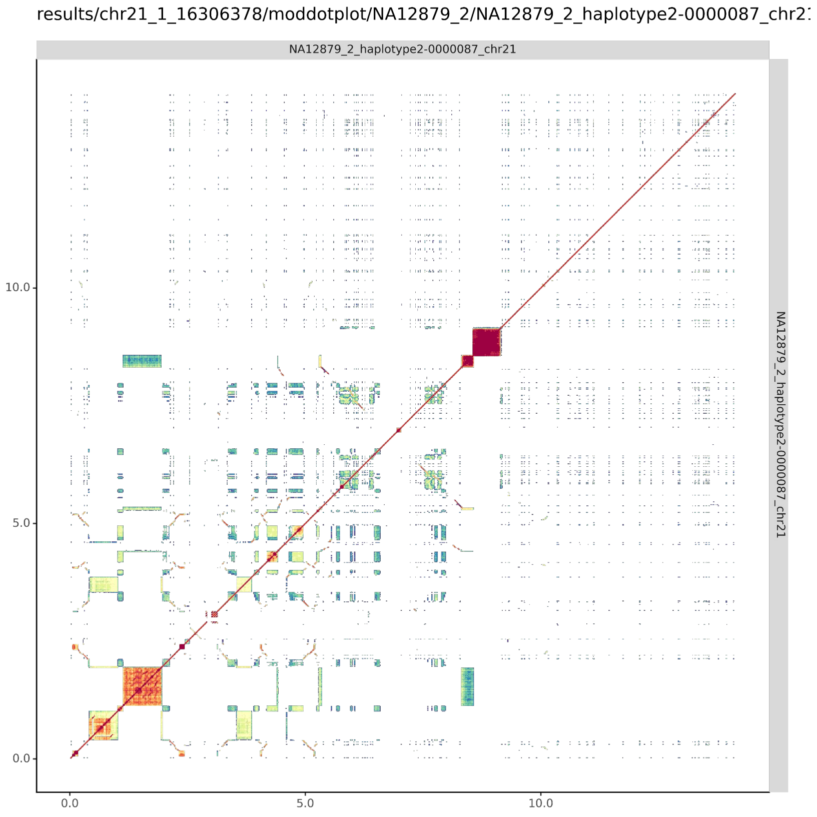

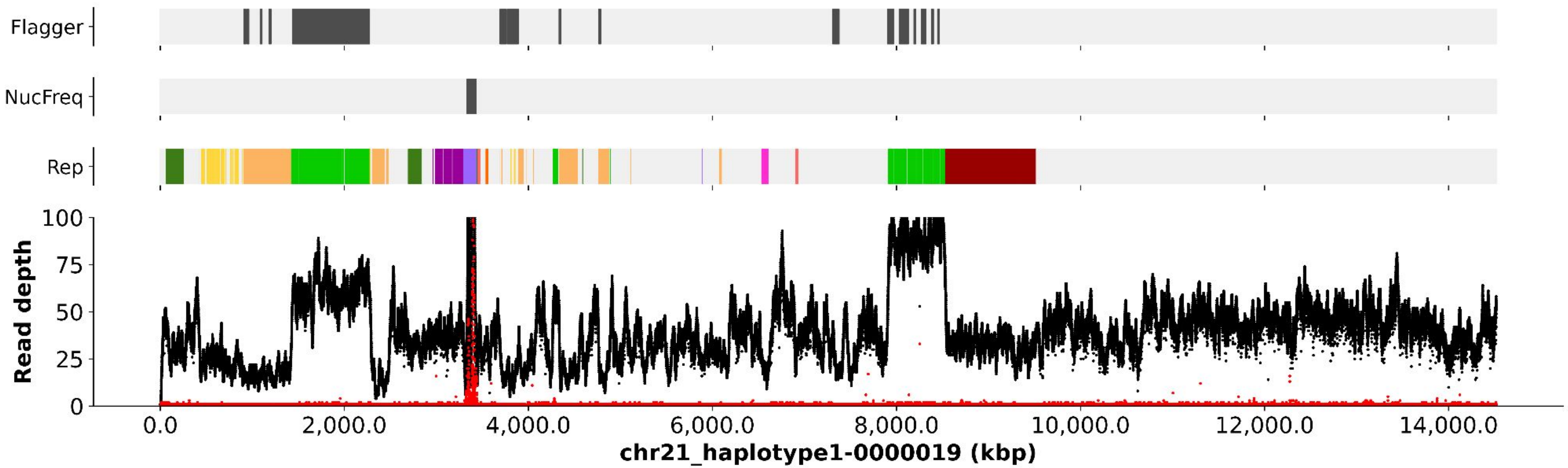

Flagger
NucFreq
Rep
Read depth
100
75
50
25
0
0.0
2,000.0
4,000.0
6,000.0
8,000.0
10,000.0
12,000.0
14,000.0
chr21_haplotype1-0000019 (kbp)


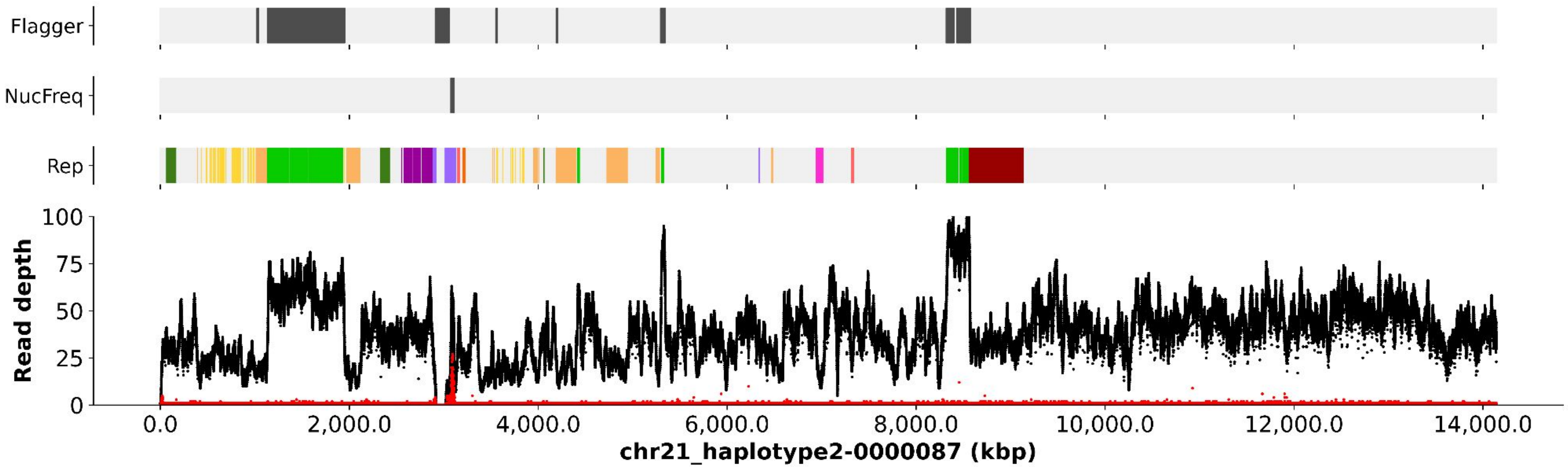

Flagger
NucFreq
Rep
Read depth
100
75
50
25
0
0.0
2,000.0
4,000.0
6,000.0
8,000.0
10,000.0
12,000.0
14,000.0
chr21_haplotype2-0000087 (kbp)

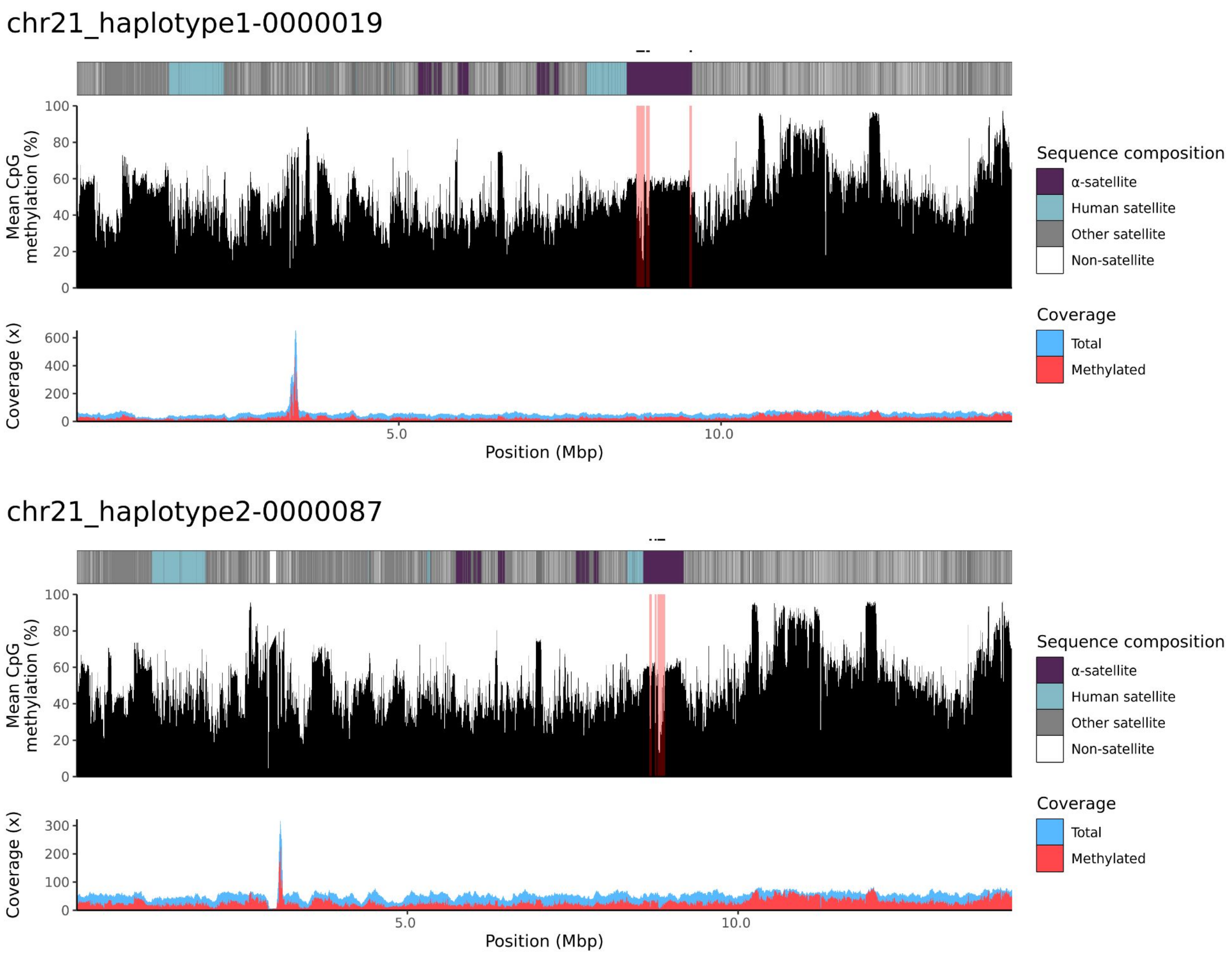

chr21_haplotype1-0000019
Mean CpG methylation (%)
100
80
60
40
20
0
Coverage (x)
600
400
200
0
5.0
10.0
Position (Mbp)
Sequence composition
α-satellite
Human satellite
Other satellite
Non-satellite
Coverage
Total
Methylated
chr21_haplotype2-0000087
Mean CpG methylation (%)
100
80
60
40
20
0
Coverage (x)
300
200
100
0
5.0
10.0
Position (Mbp)
Sequence composition
α-satellite
Human satellite
Other satellite
Non-satellite
Coverage
Total
Methylated

### chr22

#### NA12879_1_haplotype1-0000018_chr22

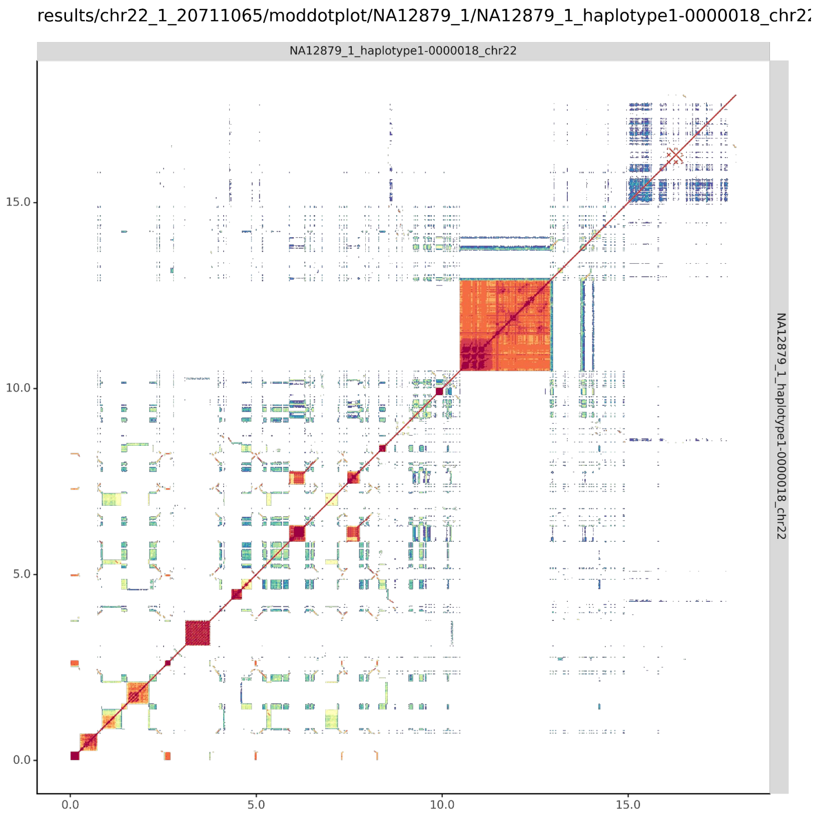


#### NA12879_2_haplotype2-0000086_chr22

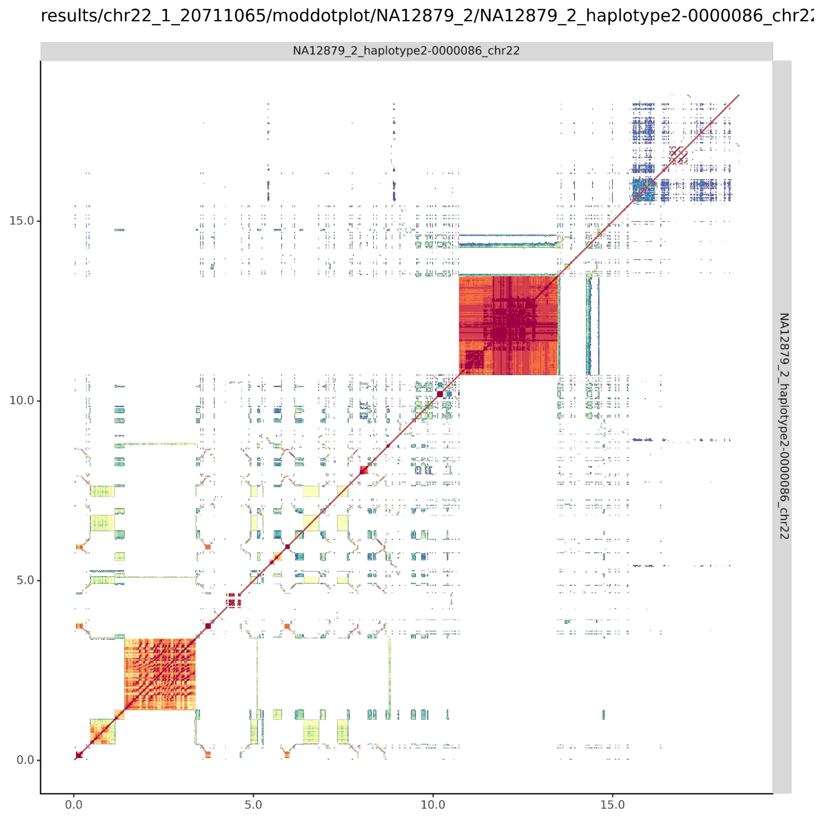

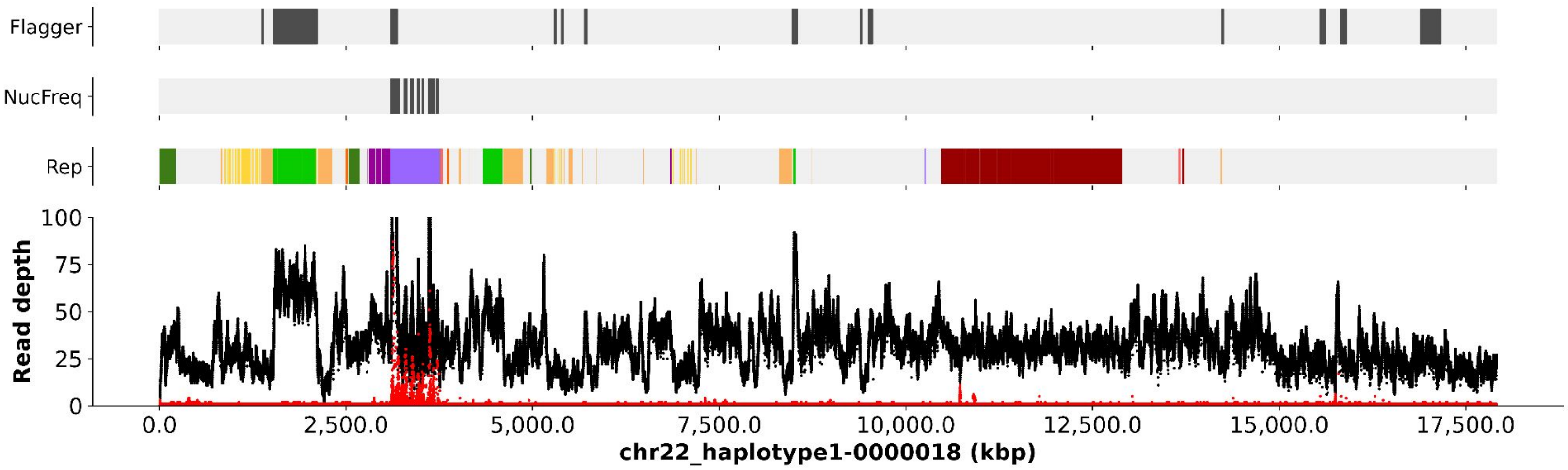
Flagger
NucFreq
Rep
Read depth
0
25
50
75
100
0.0
2,500.0
5,000.0
7,500.0
10,000.0
12,500.0
15,000.0
17,500.0
chr22_haplotype1-0000018 (kbp)

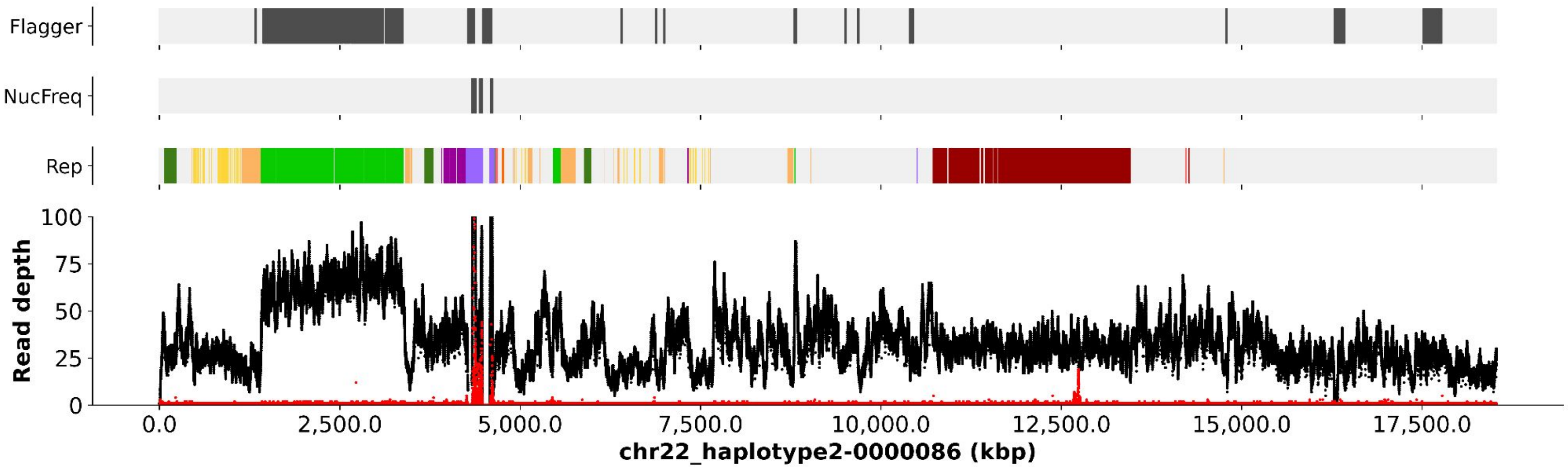
Flagger
NucFreq
Rep
Read depth
0
25
50
75
100
0.0
2,500.0
5,000.0
7,500.0
10,000.0
12,500.0
15,000.0
17,500.0
chr22_haplotype2-0000086 (kbp)

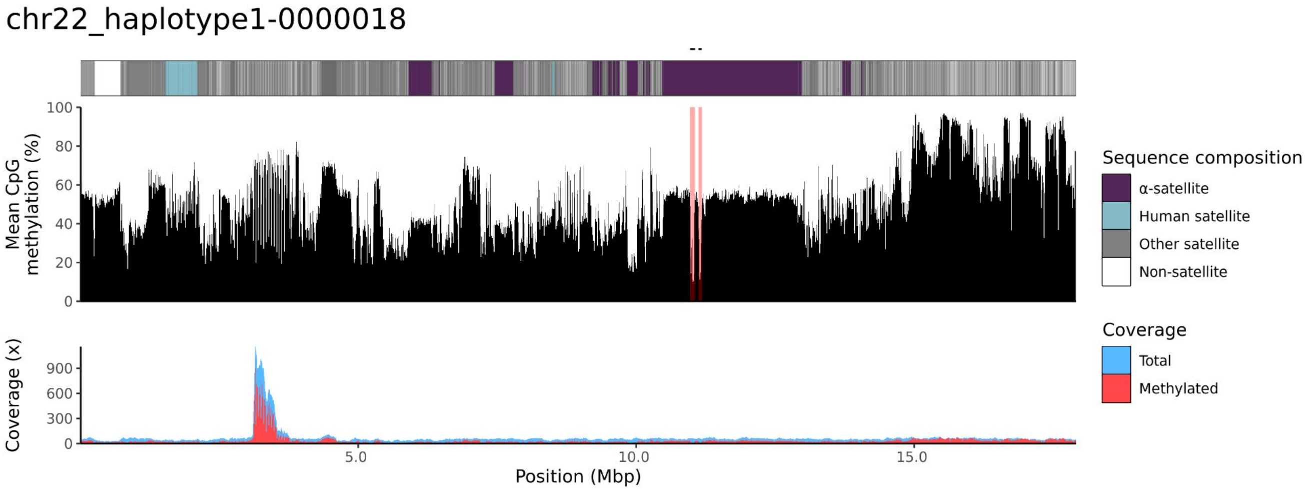
chr22_haplotype1-0000018
Mean CpG methylation (%)
100
80
60
40
20
0
Coverage (x)
900
600
300
0
5.0
10.0
15.0
Position (Mbp)
Sequence composition
α-satellite
Human satellite
Other satellite
Non-satellite
Coverage
Total
Methylated

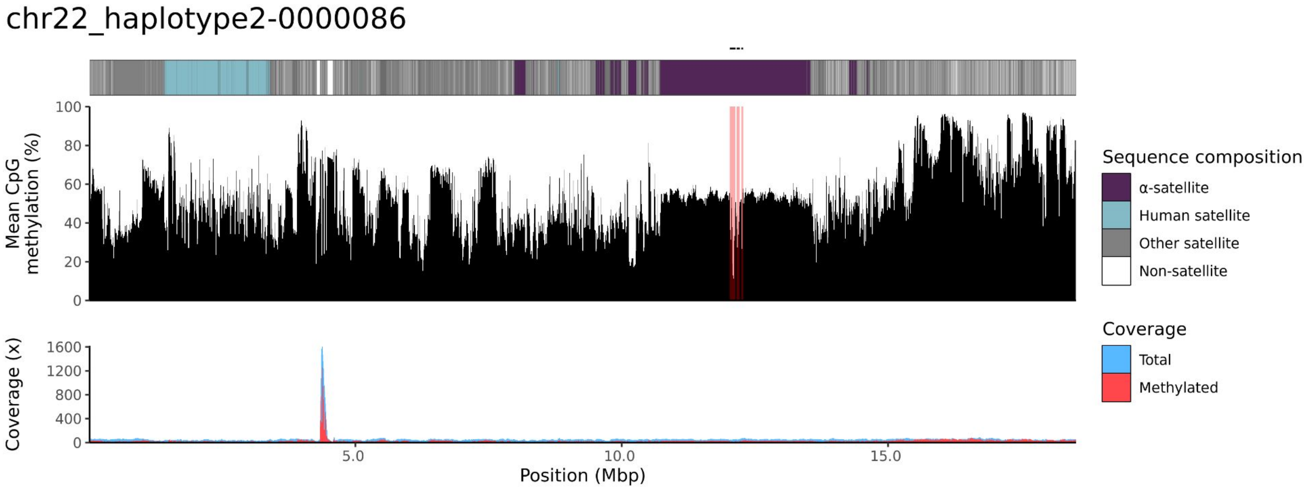
chr22_haplotype2-0000086
Mean CpG methylation (%)
100
80
60
40
20
0
Coverage (x)
1600
1200
800
400
0
5.0
10.0
15.0
Position (Mbp)
Sequence composition
α-satellite
Human satellite
Other satellite
Non-satellite
Coverage
Total
Methylated

NA12881

### chr13

#### NA12881_1_haplotype1-0000010_chr13

results/chr13_1_22508596/moddotplot/NA12881_1/NA12881_1_haplotype1-0000010_chr13

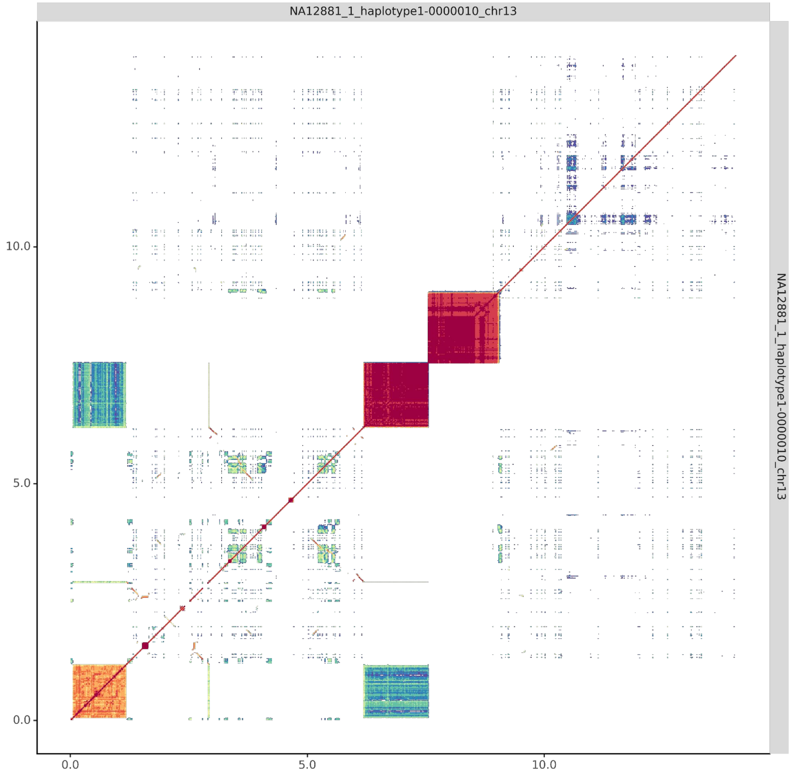


#### NA12881_2_haplotype2-0000061_chr13

results/chr13_1_22508596/moddotplot/NA12881_2/NA12881_2_haplotype2-0000061_chr13

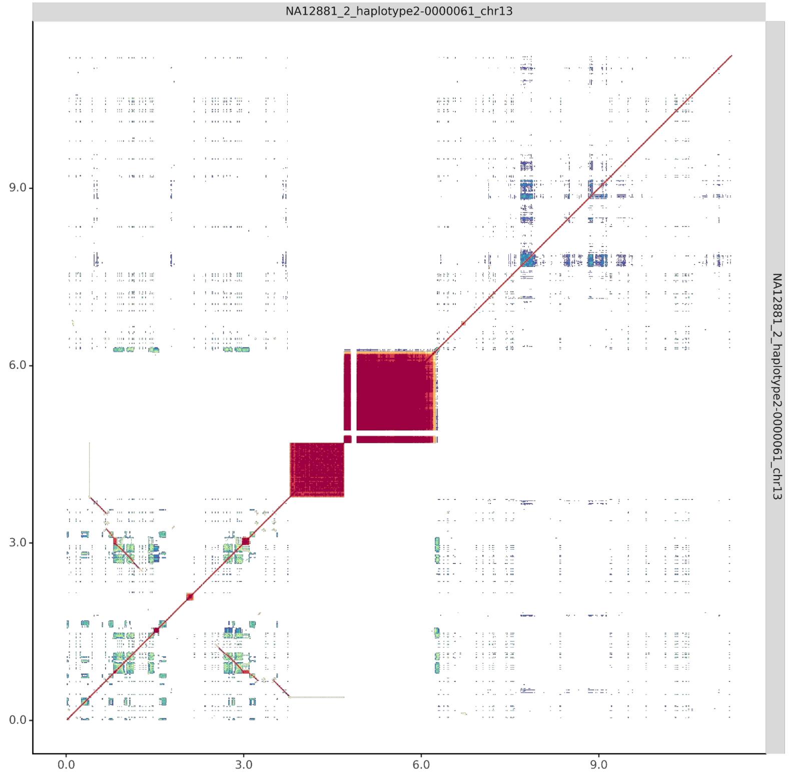

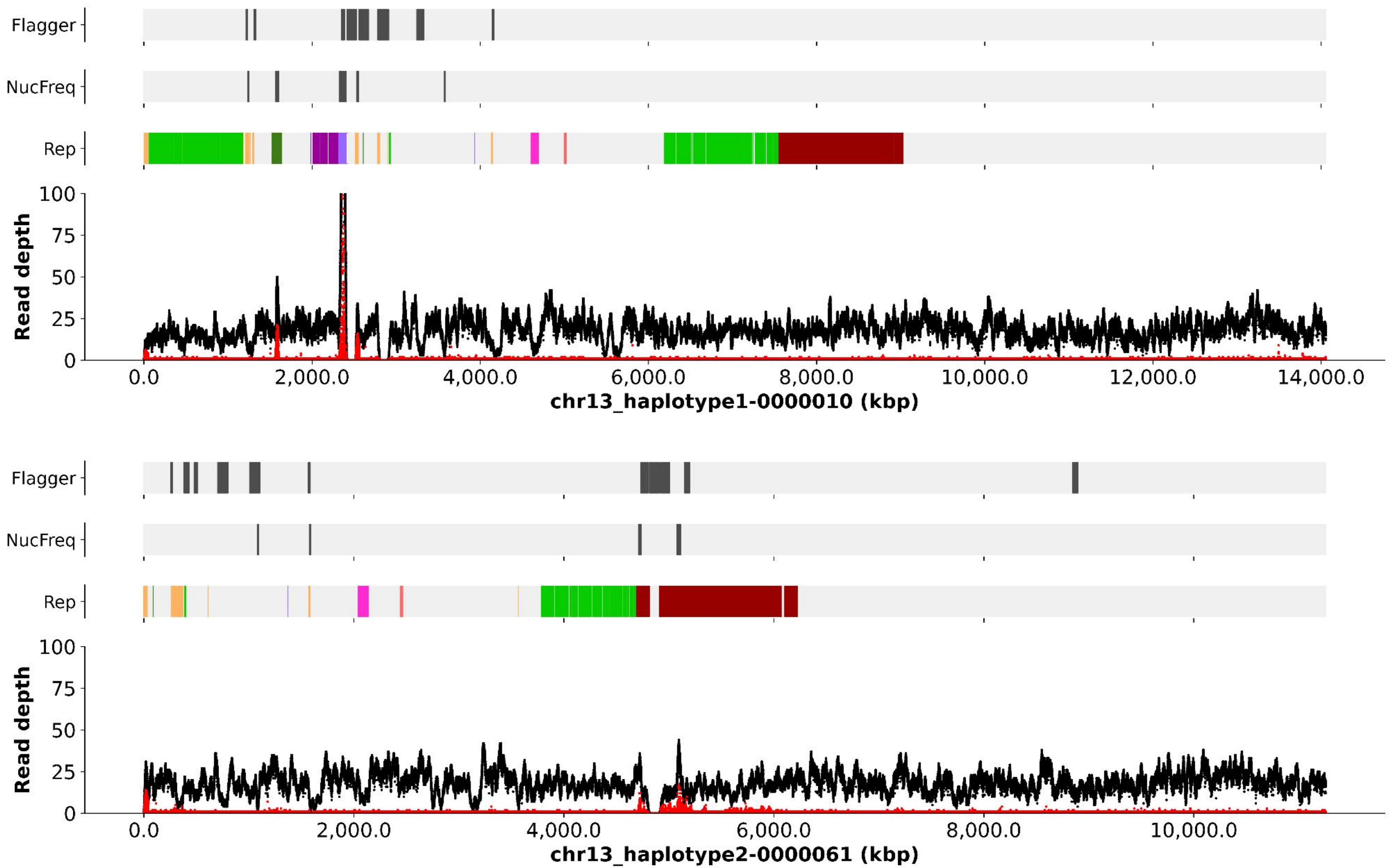
Flagger
NucFreq
Rep
Read depth
100
75
50
25
0
0.0
2,000.0
4,000.0
6,000.0
8,000.0
10,000.0
12,000.0
14,000.0
chr13_haplotype1-0000010 (kbp)
Flagger
NucFreq
Rep
Read depth
100
75
50
25
0
0.0
2,000.0
4,000.0
6,000.0
8,000.0
10,000.0
chr13_haplotype2-0000061 (kbp)

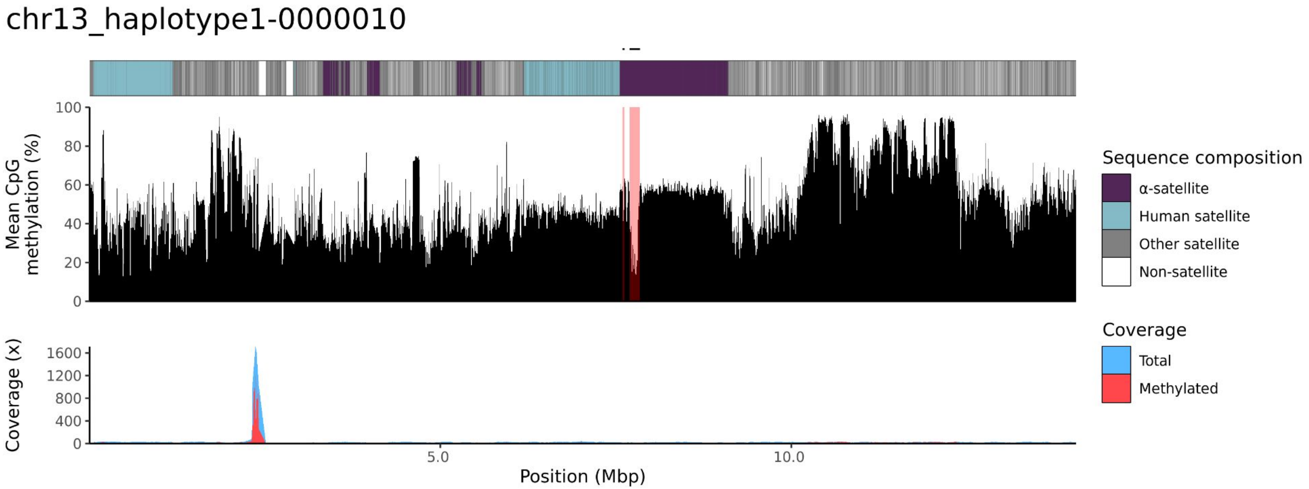
chr13_haplotype1-0000010
Mean CpG methylation (%)
0
20
40
60
80
100
Coverage (x)
0
400
800
1200
1600
5.0
10.0
Position (Mbp)
Sequence composition
α-satellite
Human satellite
Other satellite
Non-satellite
Coverage
Total
Methylated

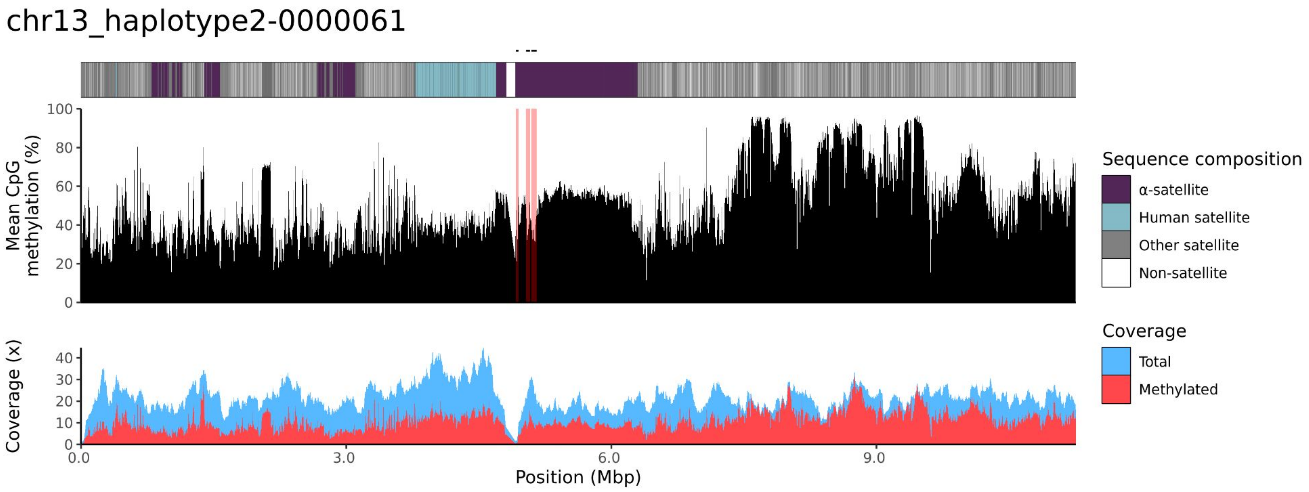
chr13_haplotype2-0000061
Mean CpG methylation (%)
0
20
40
60
80
100
Coverage (x)
0
10
20
30
40
0.0
3.0
6.0
9.0
Position (Mbp)
Sequence composition
α-satellite
Human satellite
Other satellite
Non-satellite
Coverage
Total
Methylated

### chr14

#### NA12881_1_haplotype1-0000011_chr14

results/chr14_1_17708411/moddotplot/NA12881_1/NA12881_1_haplotype1-0000011_chr14

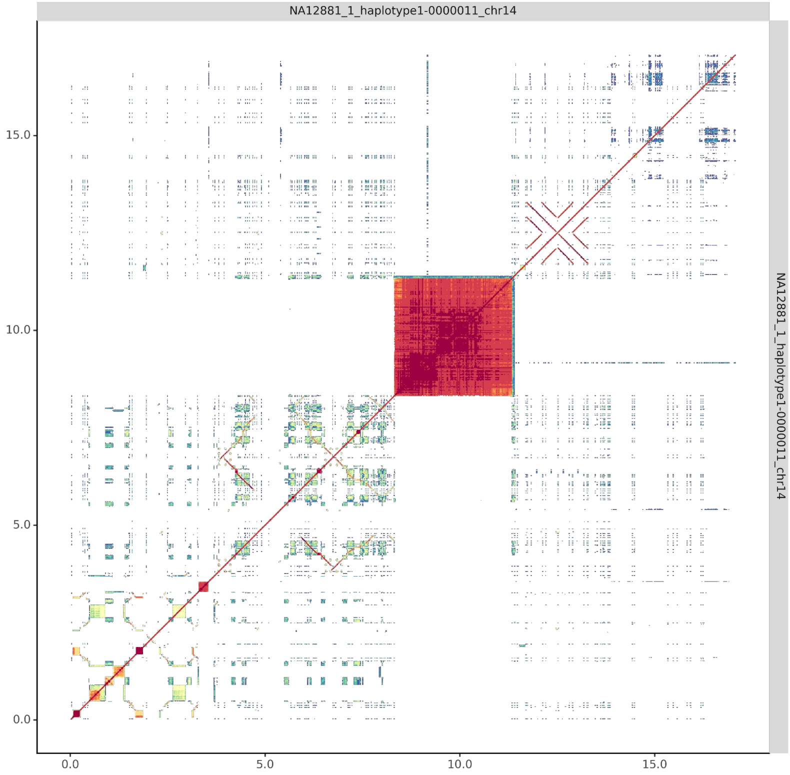


#### NA12881_2_haplotype2-0000062_chr14

results/chr14_1_17708411/moddotplot/NA12881_2/NA12881_2_haplotype2-0000062_chr14

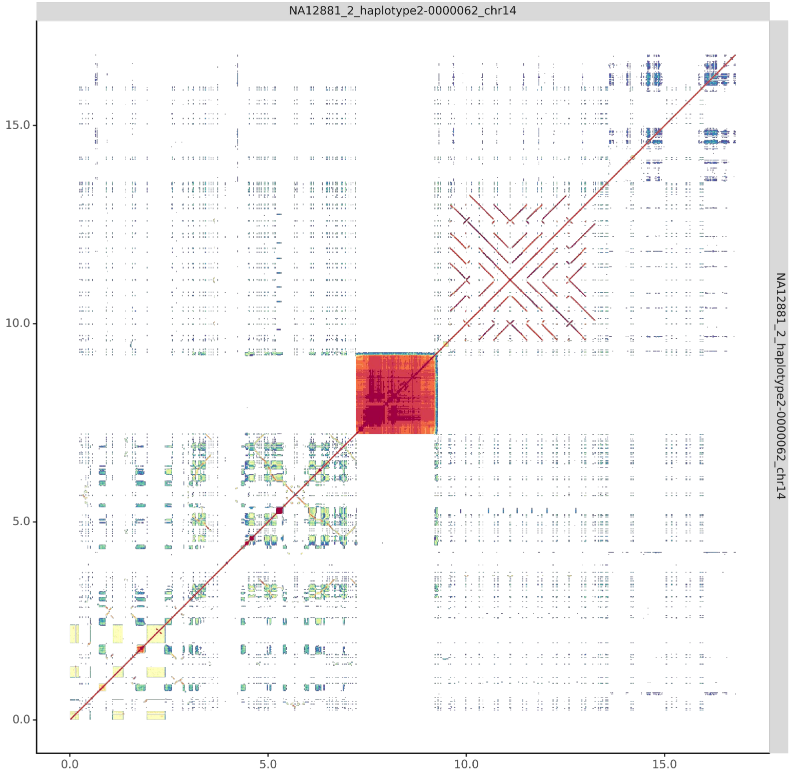

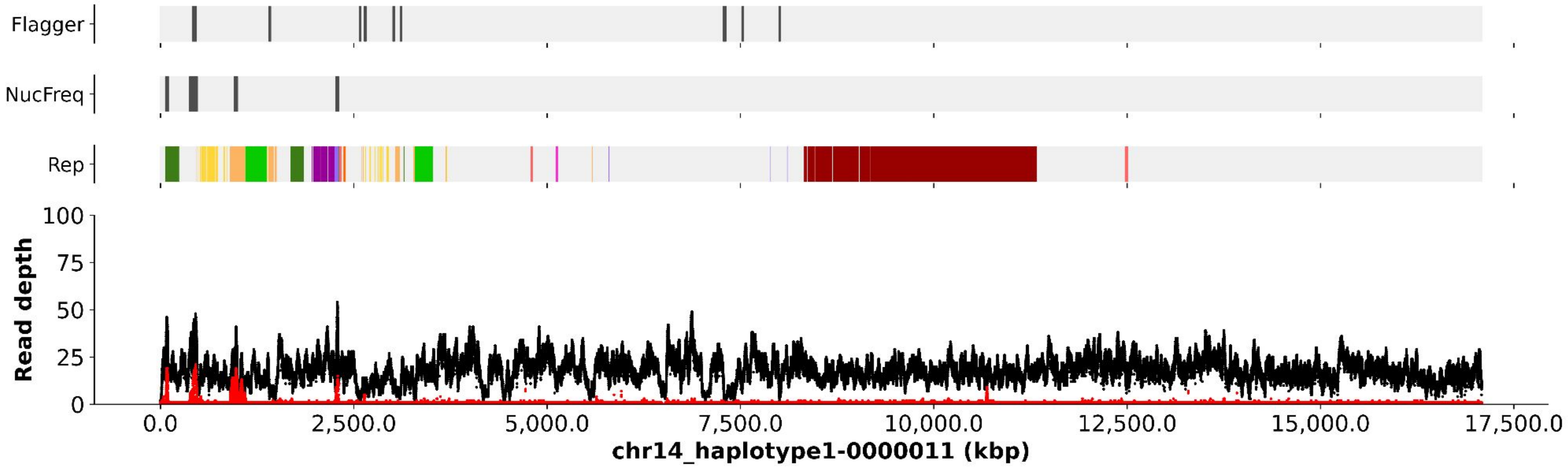

Flagger
NucFreq
Rep
100
75
50
25
0
Read depth
0.0
2,500.0
5,000.0
7,500.0
10,000.0
12,500.0
15,000.0
17,500.0
chr14_haplotype1-0000011 (kbp)


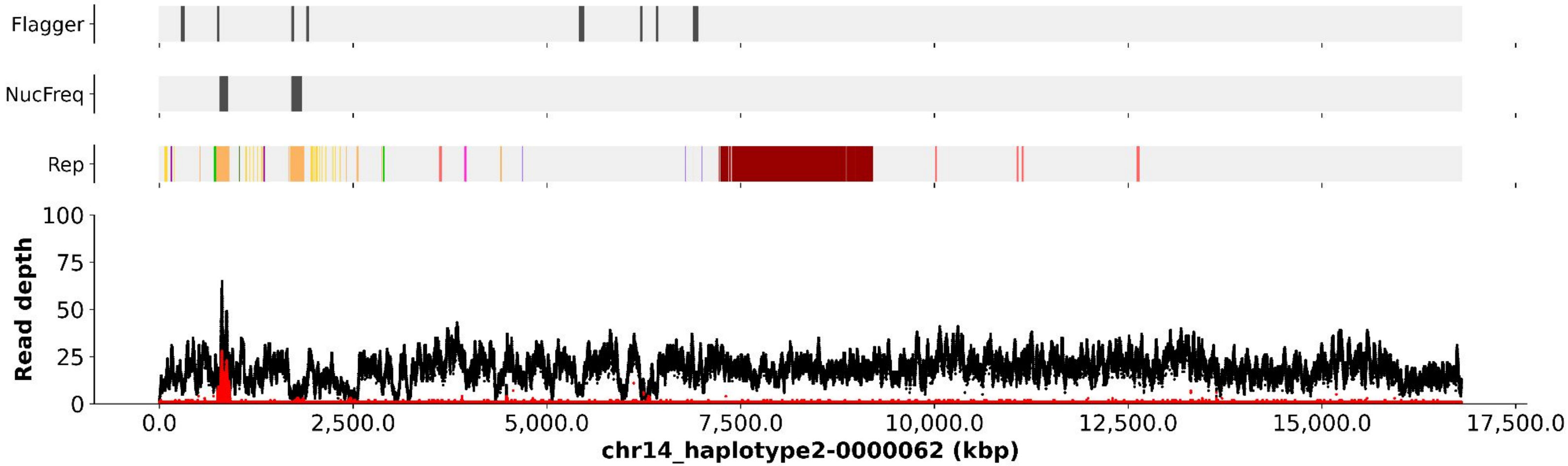

Flagger
NucFreq
Rep
100
75
50
25
0
Read depth
0.0
2,500.0
5,000.0
7,500.0
10,000.0
12,500.0
15,000.0
17,500.0
chr14_haplotype2-0000062 (kbp)

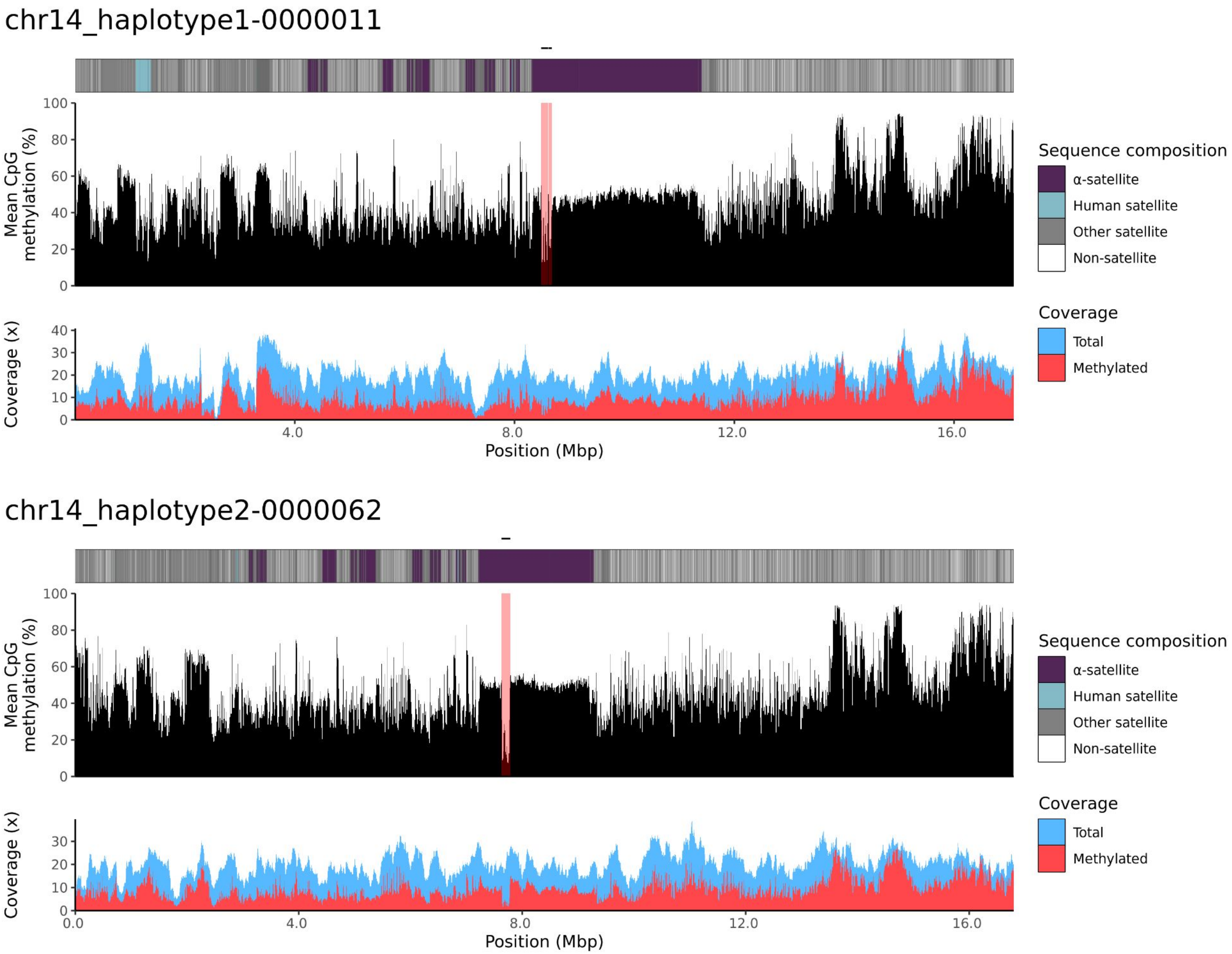
chr14_haplotype1-0000011
Mean CpG methylation (%)
Coverage (x)
Position (Mbp)
Sequence composition
α-satellite
Human satellite
Other satellite
Non-satellite
Coverage
Total
Methylated
chr14_haplotype2-0000062
Mean CpG methylation (%)
Coverage (x)
Position (Mbp)
Sequence composition
α-satellite
Human satellite
Other satellite
Non-satellite
Coverage
Total
Methylated

### chr15

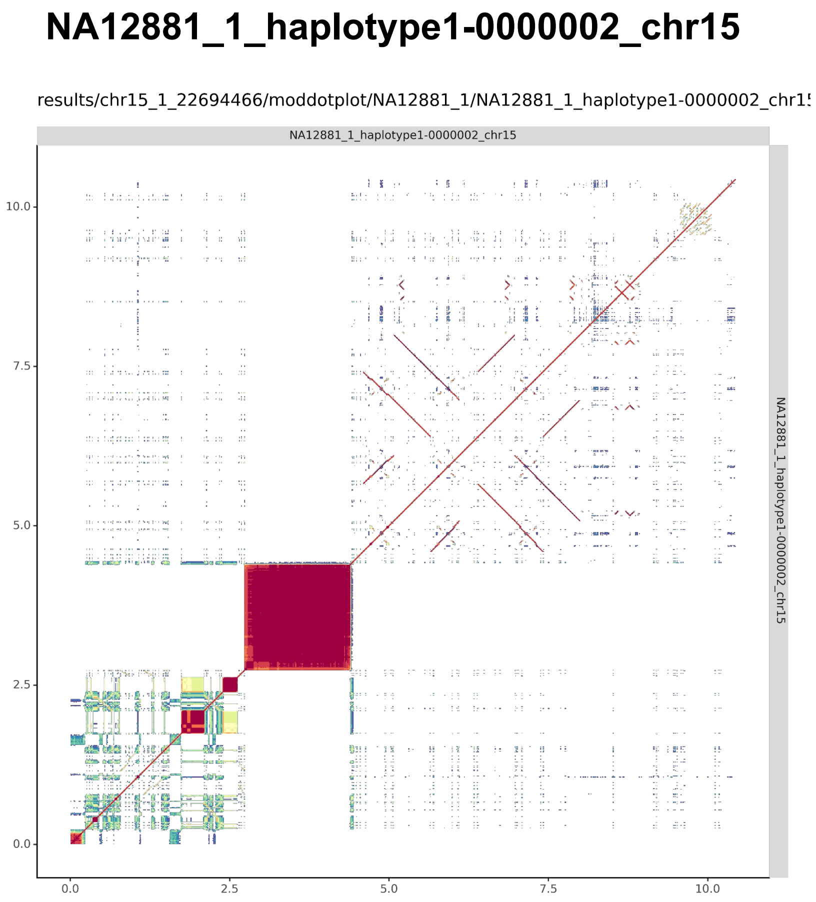
NA12881_1_haplotype1-0000002_chr15
results/chr15_1_22694466/moddotplot/NA12881_1/NA12881_1_haplotype1-0000002_chr1
NA12881_1_haplotype1-0000002_chr15
0.0
2.5
5.0
7.5
10.0
NA12881_1_haplotype1-0000002_chr15

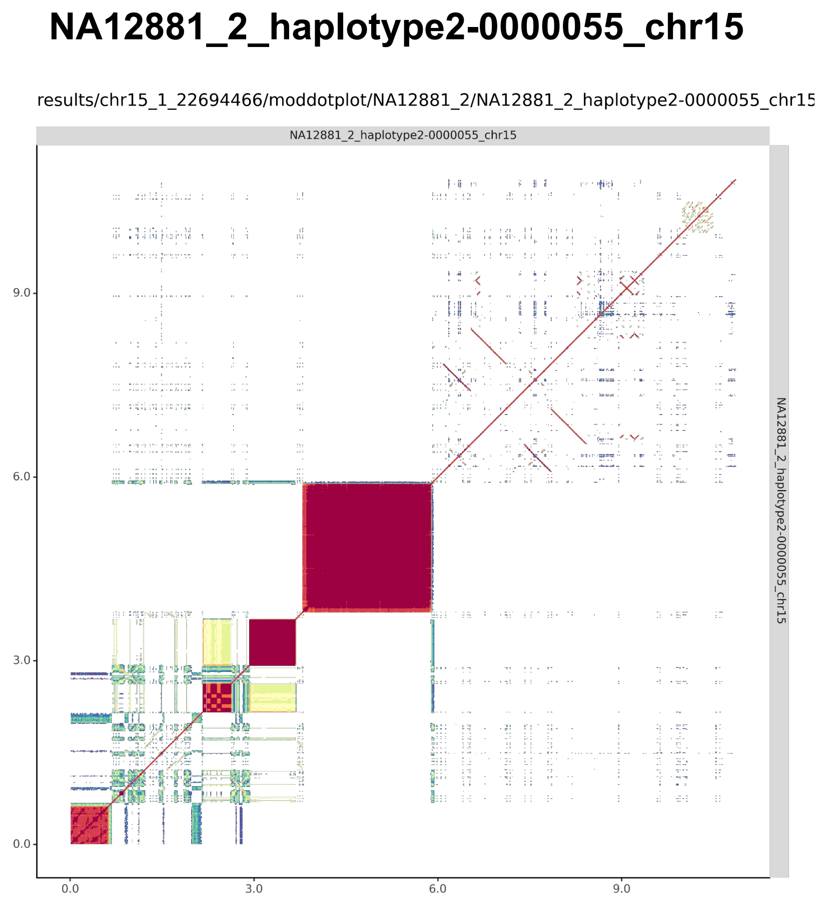
NA12881_2_haplotype2-0000055_chr15
results/chr15_1_22694466/moddotplot/NA12881_2/NA12881_2_haplotype2-0000055_chr15
NA12881_2_haplotype2-0000055_chr15
0.0
3.0
6.0
9.0
NA12881_2_haplotype2-0000055_chr15

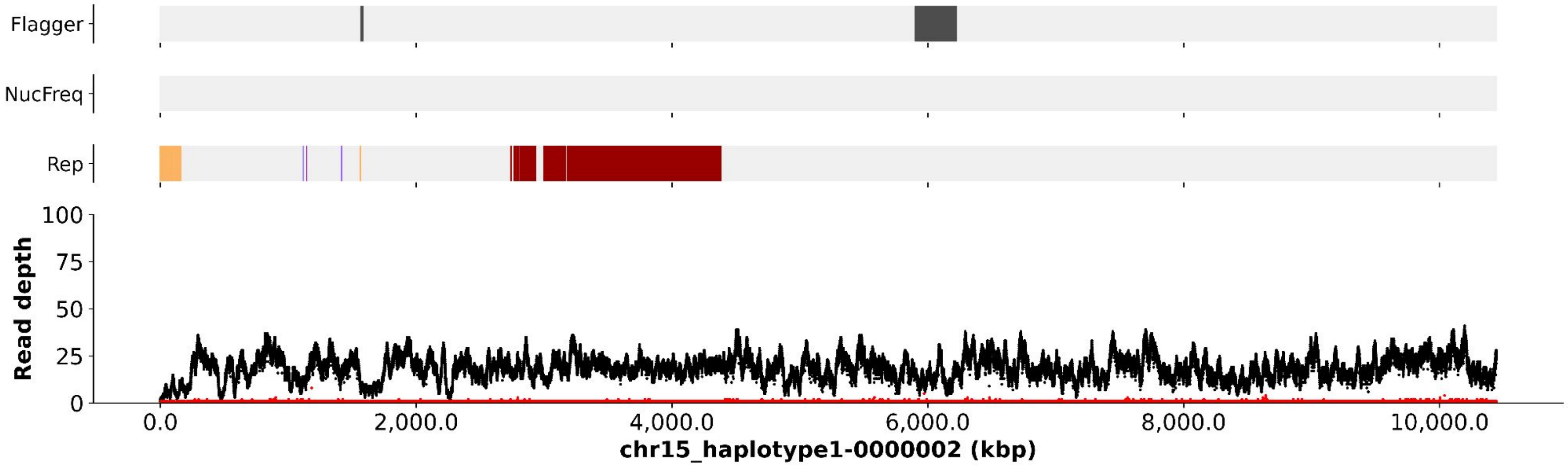

Flagger
NucFreq
Rep
Read depth
100
75
50
25
0
0.0
2,000.0
4,000.0
6,000.0
8,000.0
10,000.0
chr15_haplotype1-0000002 (kbp)


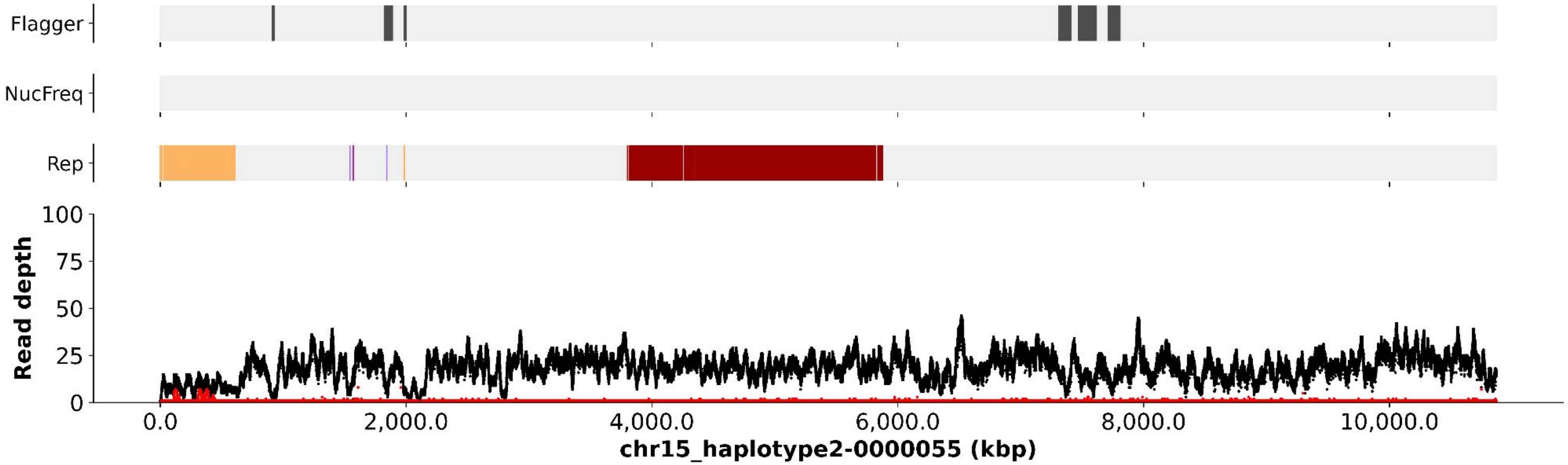

Flagger
NucFreq
Rep
Read depth
100
75
50
25
0
0.0
2,000.0
4,000.0
6,000.0
8,000.0
10,000.0
chr15_haplotype2-0000055 (kbp)

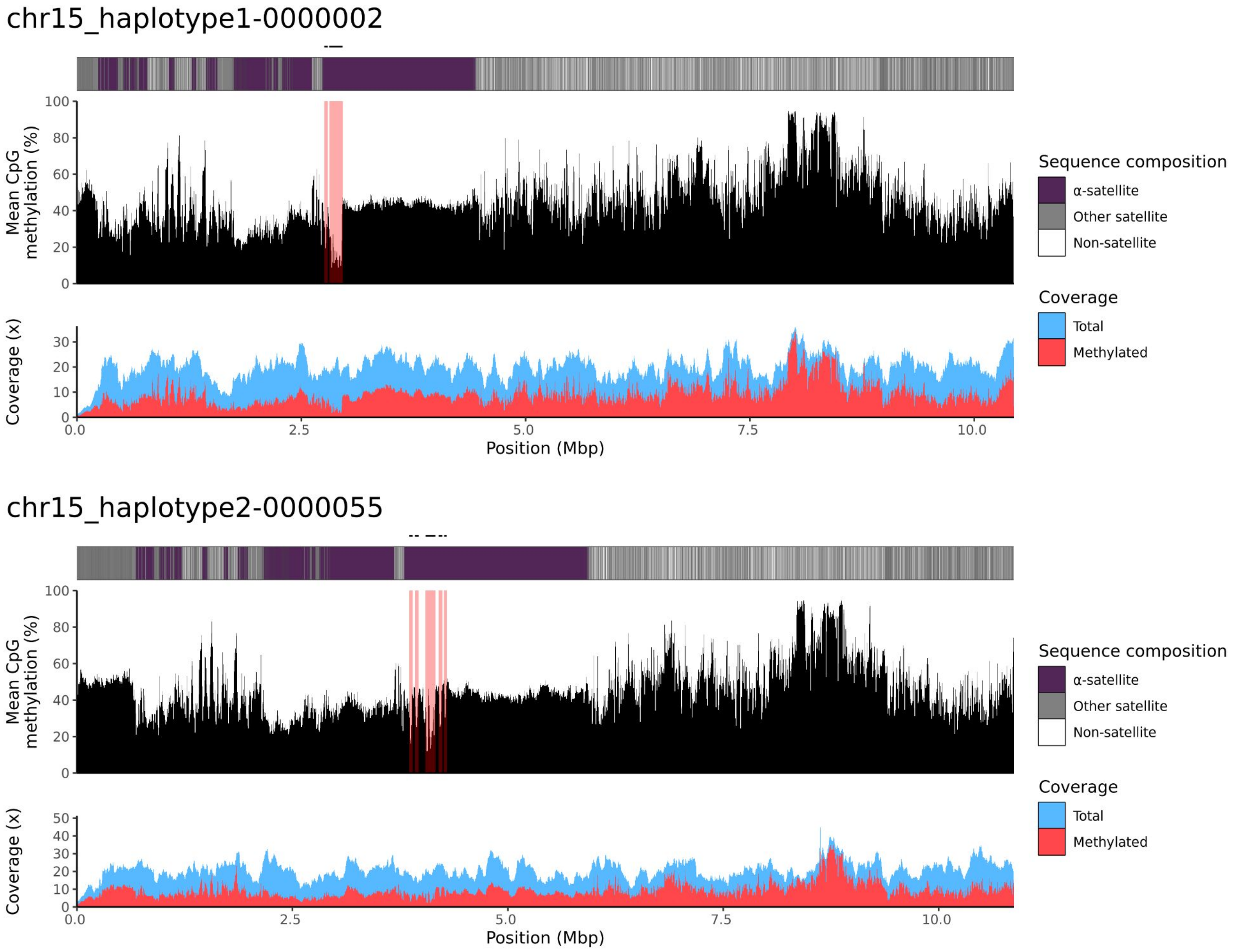
chr15_haplotype1-0000002
Mean CpG methylation (%)
Coverage (x)
Position (Mbp)
Sequence composition
α-satellite
Other satellite
Non-satellite
Coverage
Total
Methylated
chr15_haplotype2-0000055
Mean CpG methylation (%)
Coverage (x)
Position (Mbp)
Sequence composition
α-satellite
Other satellite
Non-satellite
Coverage
Total
Methylated

### chr21

#### NA12881_1_haplotype1-0000007_chr21

results/chr21_1_16306378/moddotplot/NA12881_1/NA12881_1_haplotype1-0000007_chr2

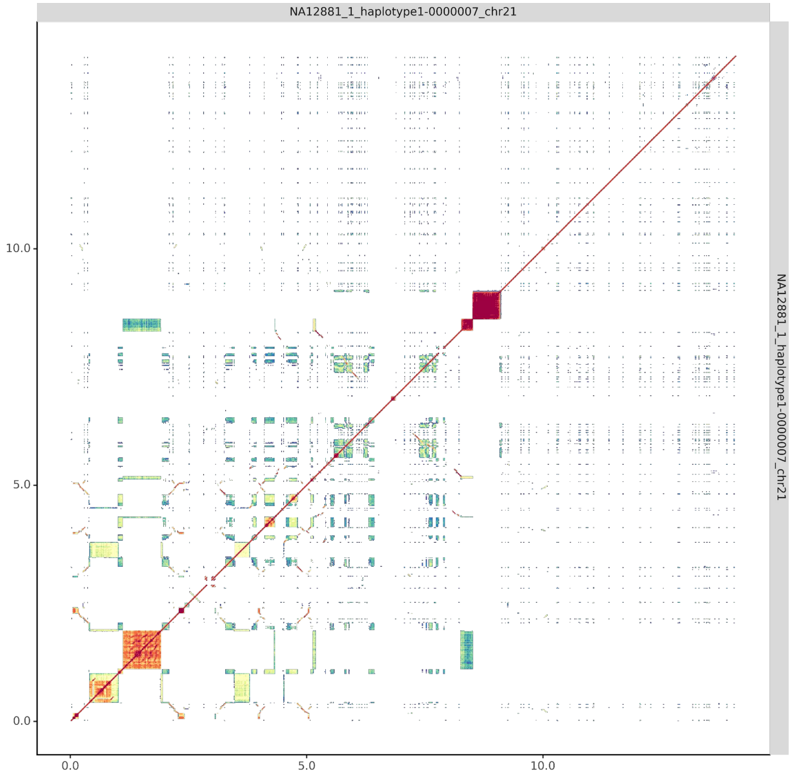


#### NA12881_2_haplotype2-0000058_chr21

results/chr21_1_16306378/moddotplot/NA12881_2/NA12881_2_haplotype2-0000058_chr21

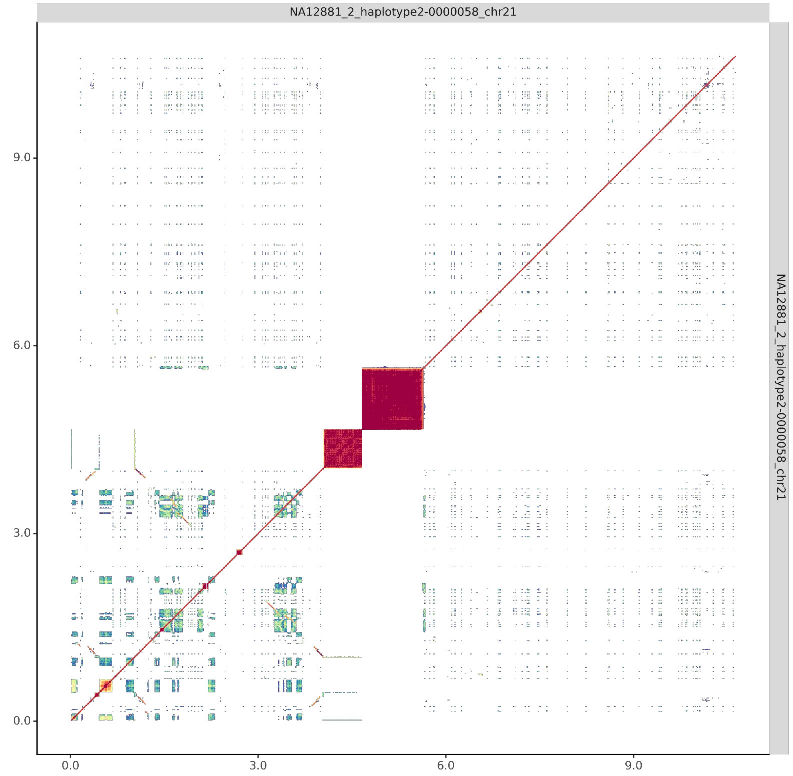

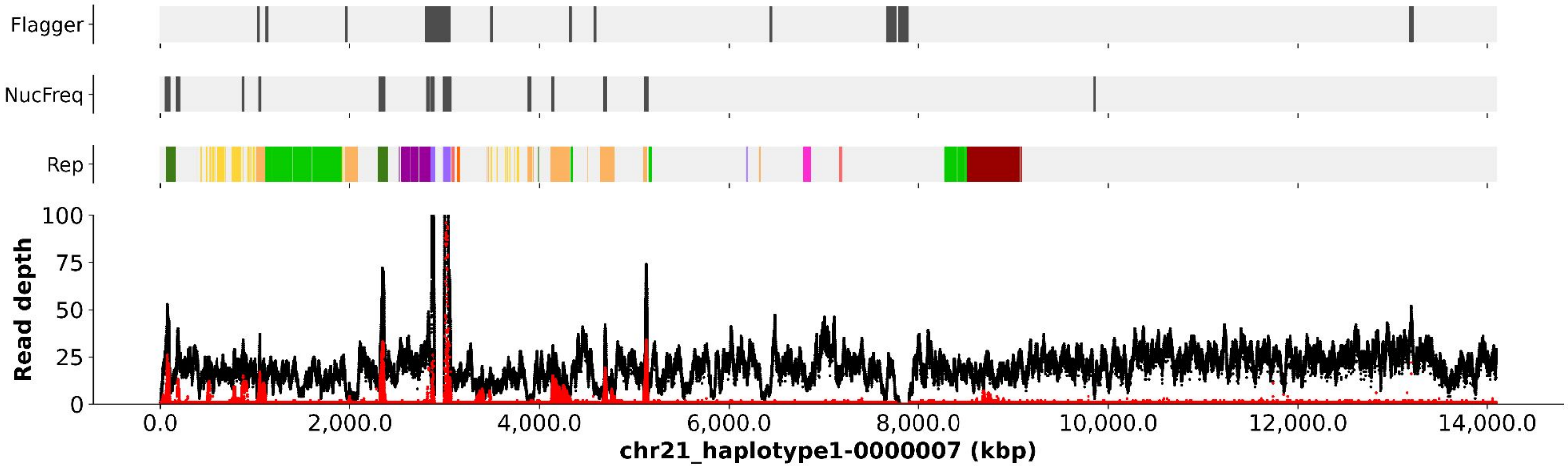
Flagger
NucFreq
Rep
Read depth
100
75
50
25
0
0.0
2,000.0
4,000.0
6,000.0
8,000.0
10,000.0
12,000.0
14,000.0
chr21_haplotype1-0000007 (kbp)

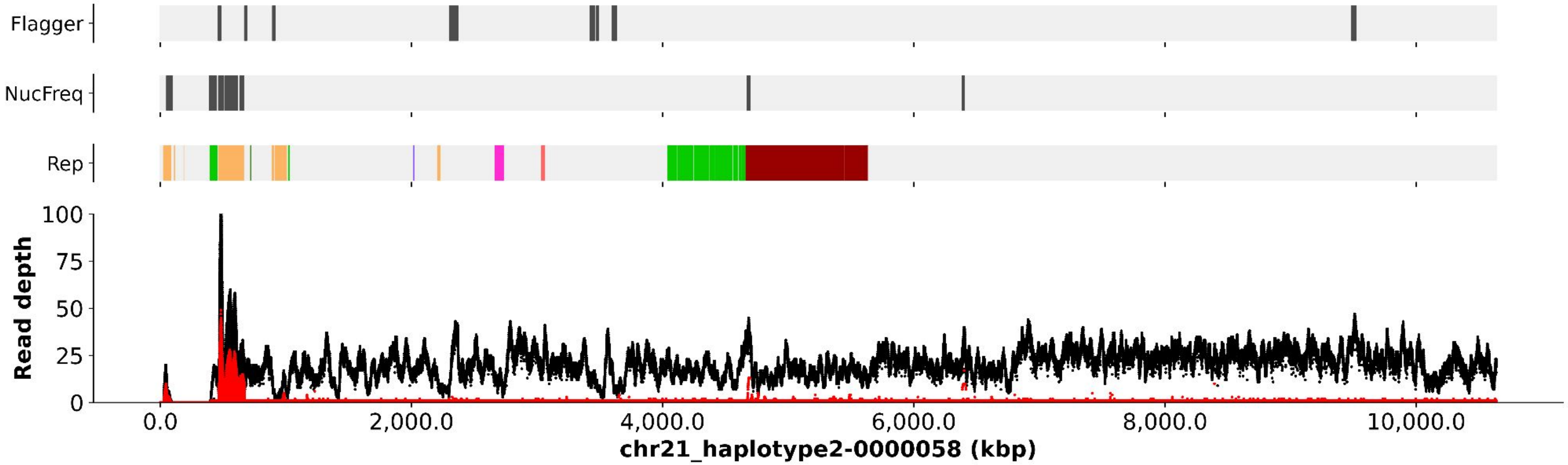
Flagger
NucFreq
Rep
Read depth
100
75
50
25
0
0.0
2,000.0
4,000.0
6,000.0
8,000.0
10,000.0
chr21_haplotype2-0000058 (kbp)

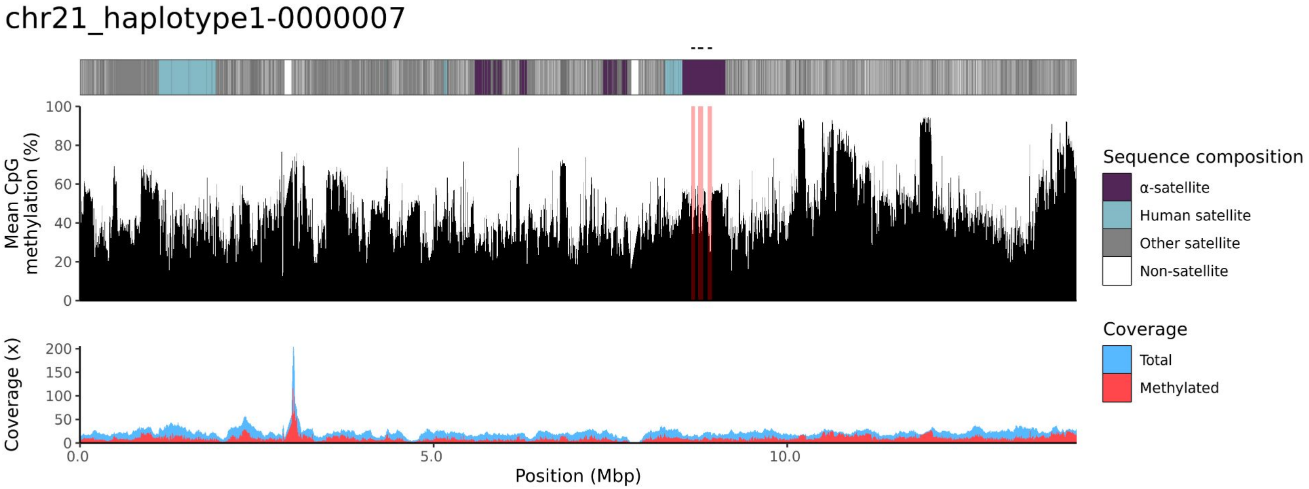
chr21_haplotype1-0000007
Mean CpG methylation (%)
100
80
60
40
20
0
Coverage (x)
200
150
100
50
0
0.0
5.0
10.0
Position (Mbp)
Sequence composition
α-satellite
Human satellite
Other satellite
Non-satellite
Coverage
Total
Methylated

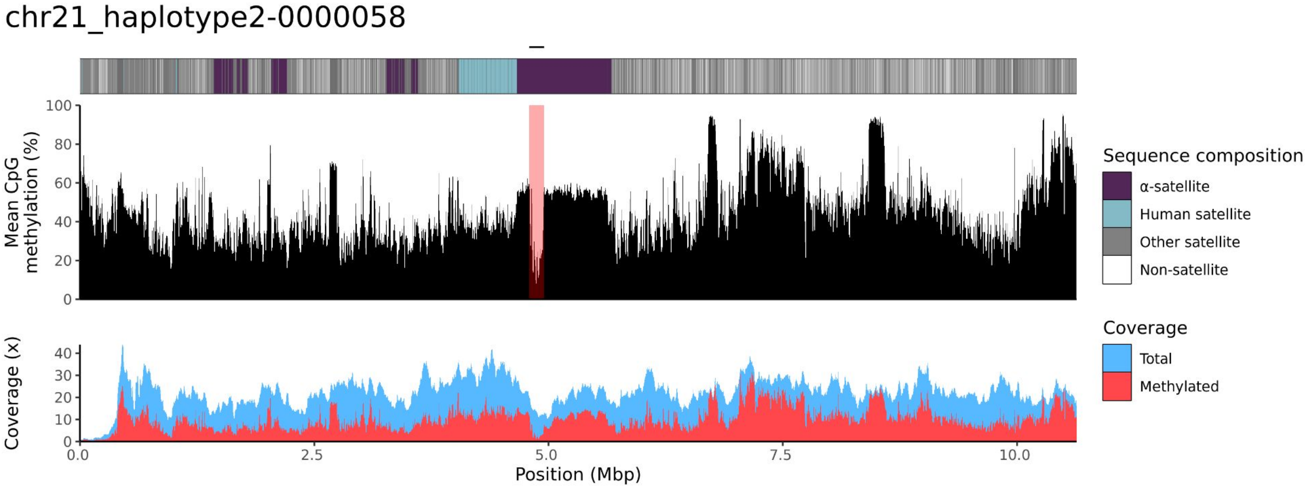
chr21_haplotype2-0000058
Mean CpG methylation (%)
100
80
60
40
20
0
Coverage (x)
40
30
20
10
0
0.0
2.5
5.0
7.5
10.0
Position (Mbp)
Sequence composition
α-satellite
Human satellite
Other satellite
Non-satellite
Coverage
Total
Methylated

### chr22

#### NA12881_1_haplotype1-0000024_chr22

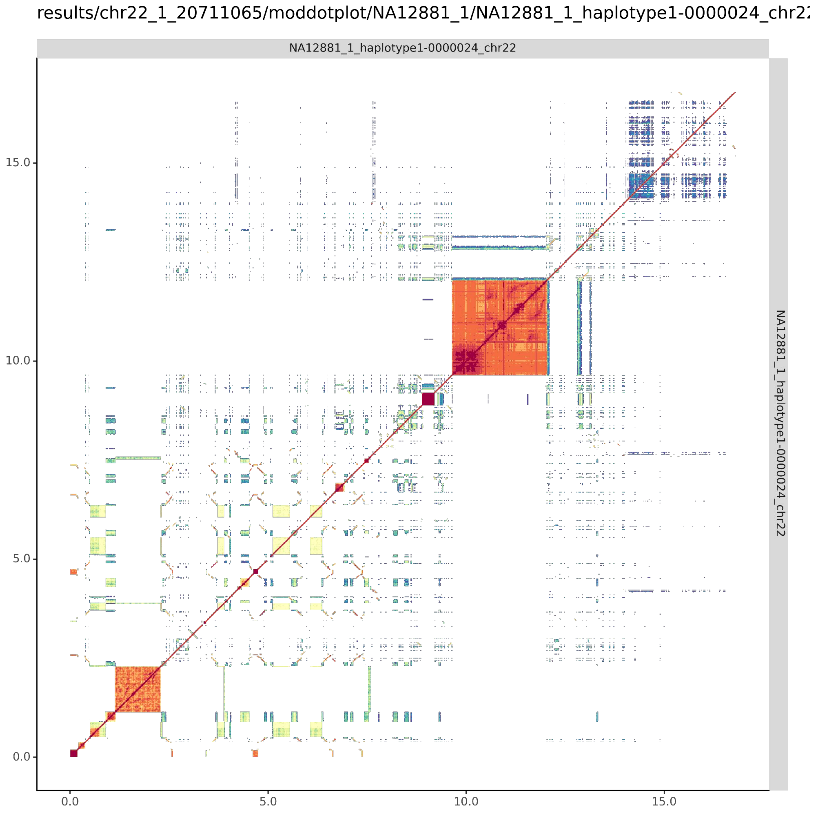


#### NA12881_2_haplotype2-0000077_chr22

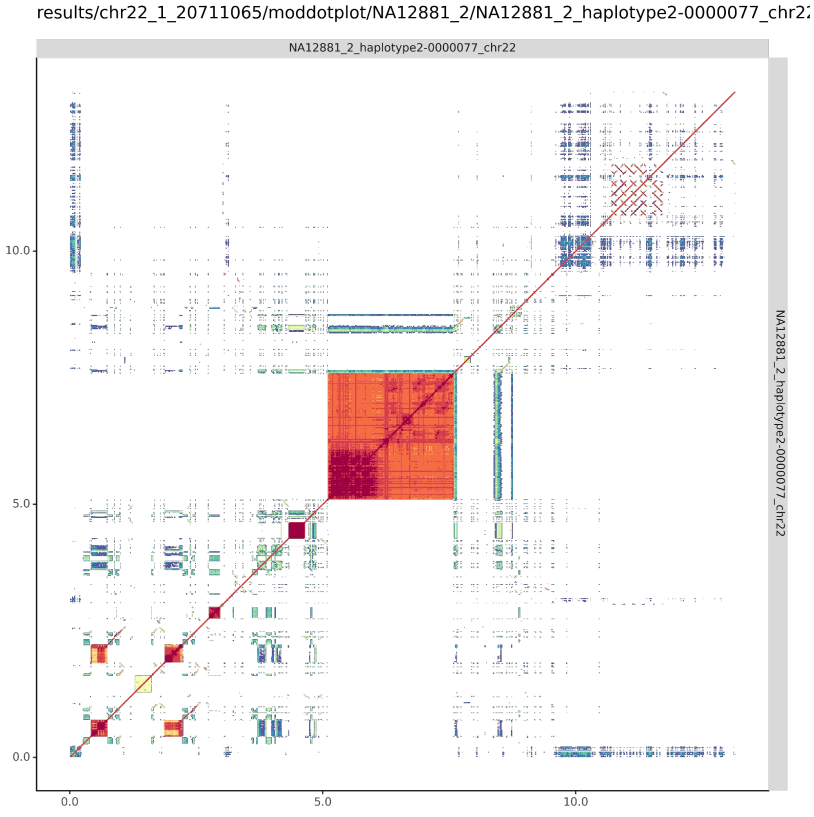

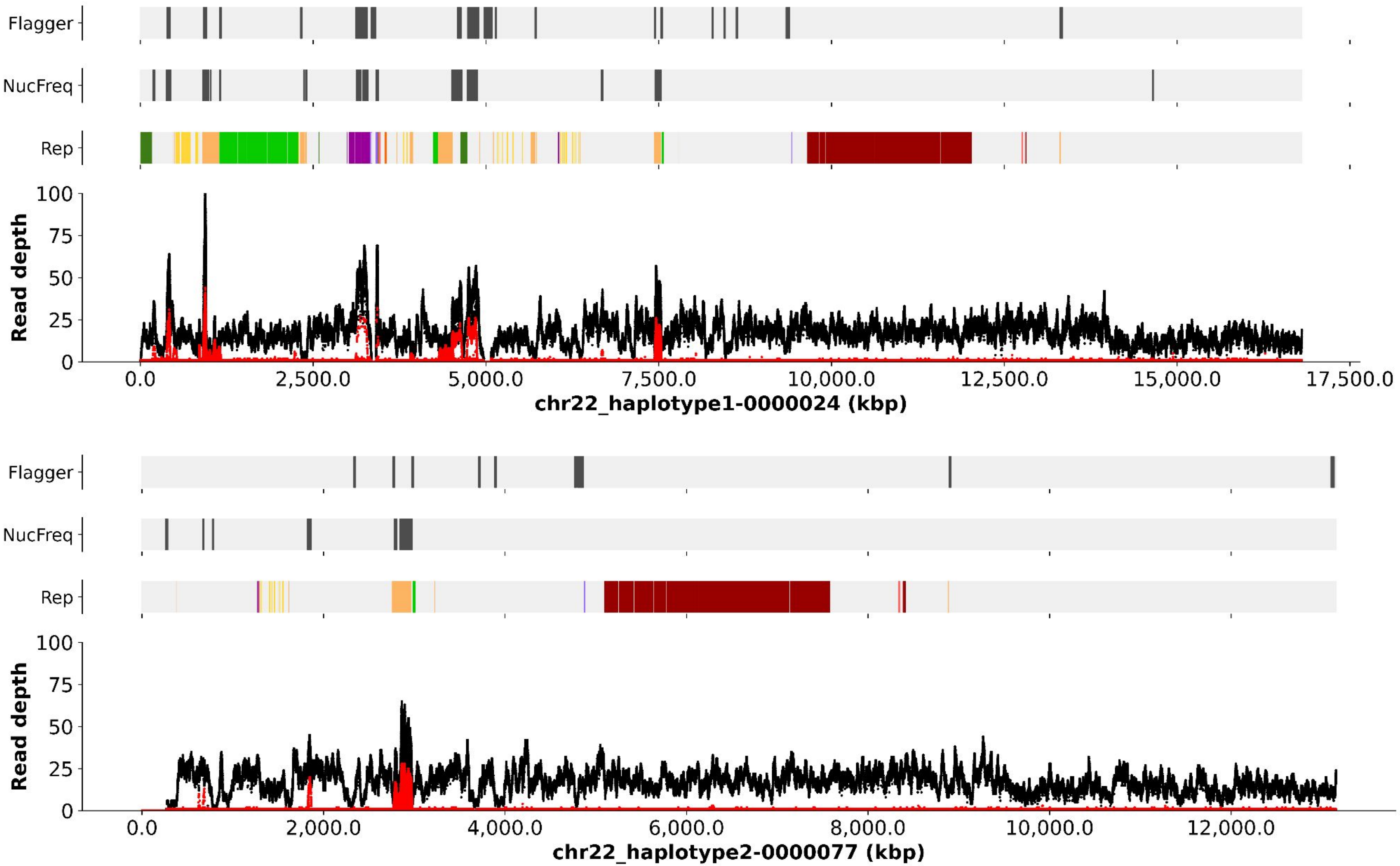

Flagger
NucFreq
Rep
Read depth
0
25
50
75
100
0.0
2,500.0
5,000.0
7,500.0
10,000.0
12,500.0
15,000.0
17,500.0
chr22_haplotype1-0000024 (kbp)
Flagger
NucFreq
Rep
Read depth
0
25
50
75
100
0.0
2,000.0
4,000.0
6,000.0
8,000.0
10,000.0
12,000.0
chr22_haplotype2-0000077 (kbp)

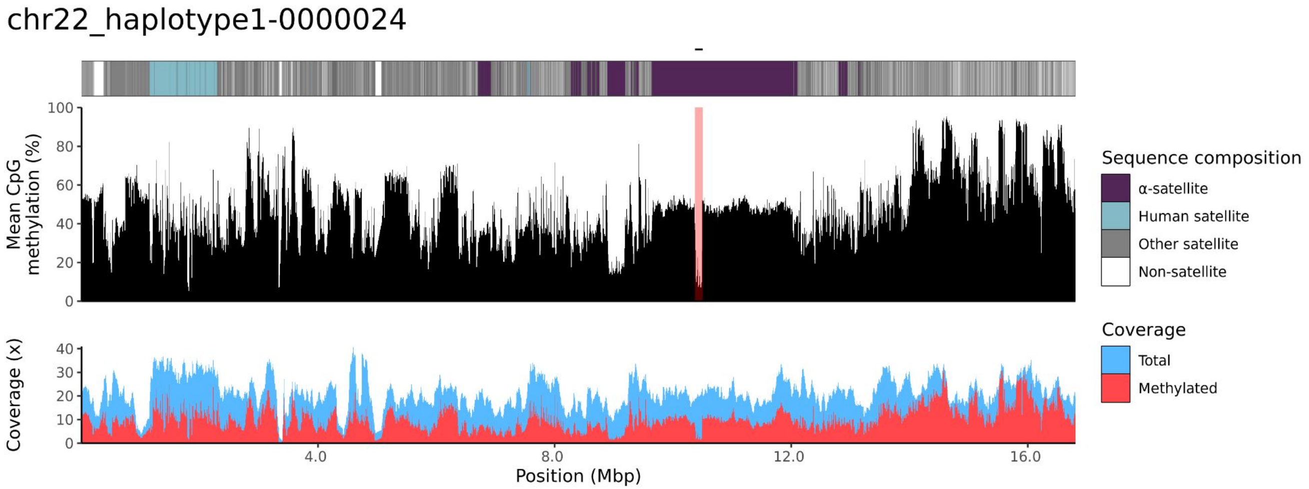
chr22_haplotype1-0000024
Mean CpG methylation (%)
Coverage (x)
Position (Mbp)
Sequence composition
α-satellite
Human satellite
Other satellite
Non-satellite
Coverage
Total
Methylated

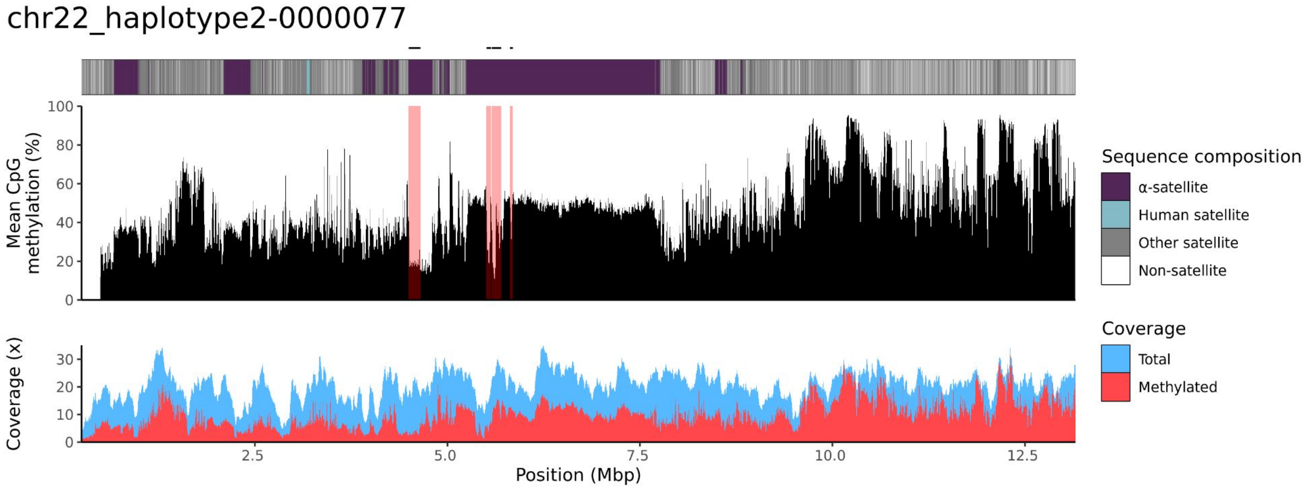
chr22_haplotype2-0000077
Mean CpG methylation (%)
Coverage (x)
Position (Mbp)
Sequence composition
α-satellite
Human satellite
Other satellite
Non-satellite
Coverage
Total
Methylated

NA12882

### chr13

#### NA12882_1_haplotype1-0000028_chr13

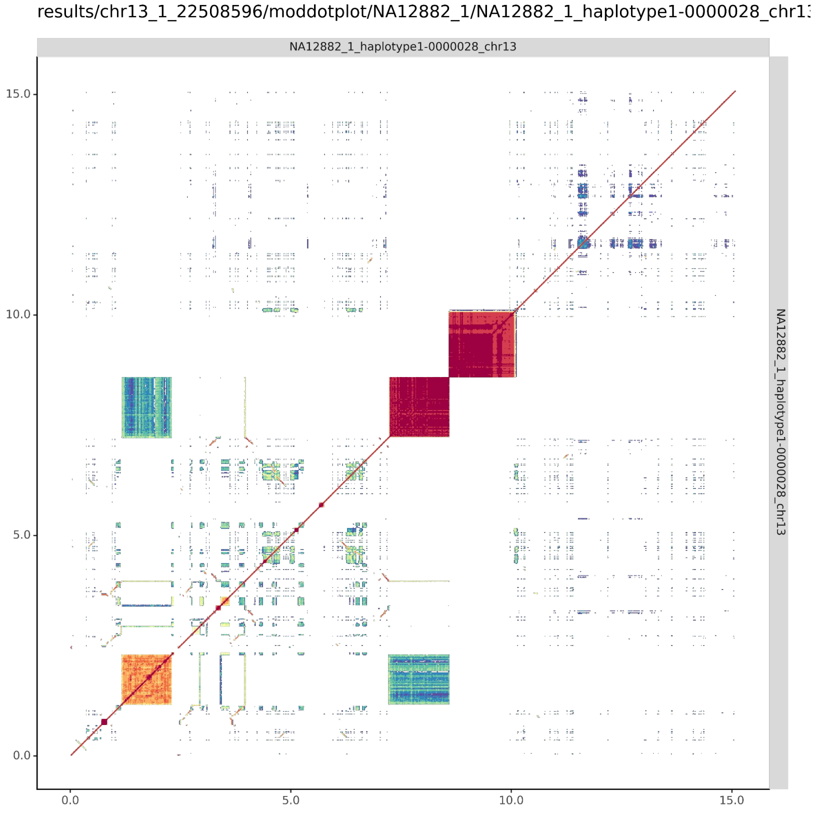


#### NA12882_2_haplotype2-0000122_chr13

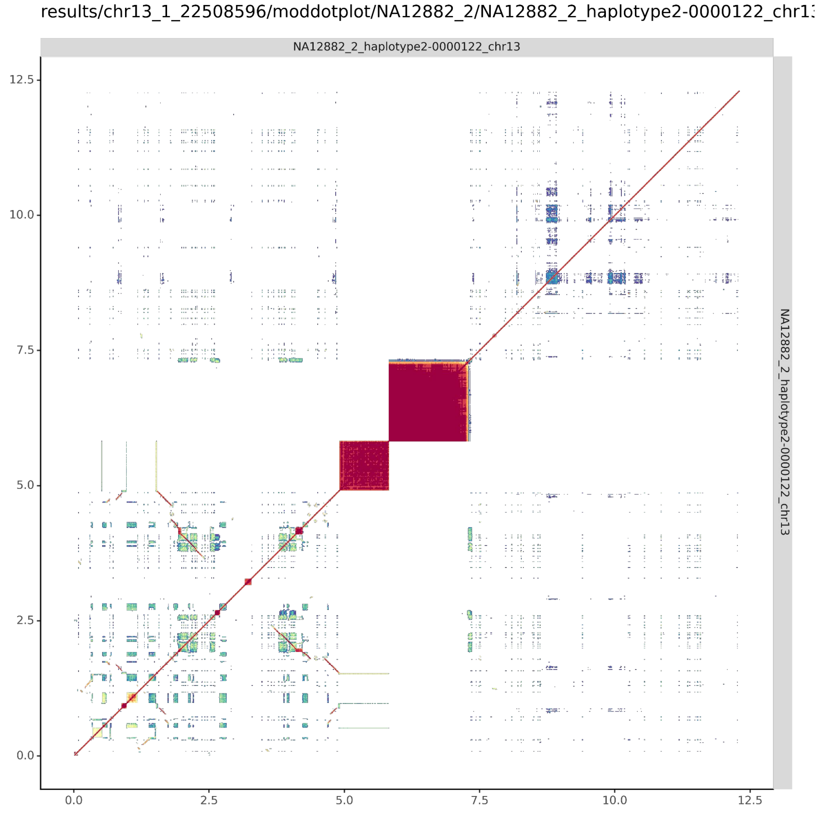

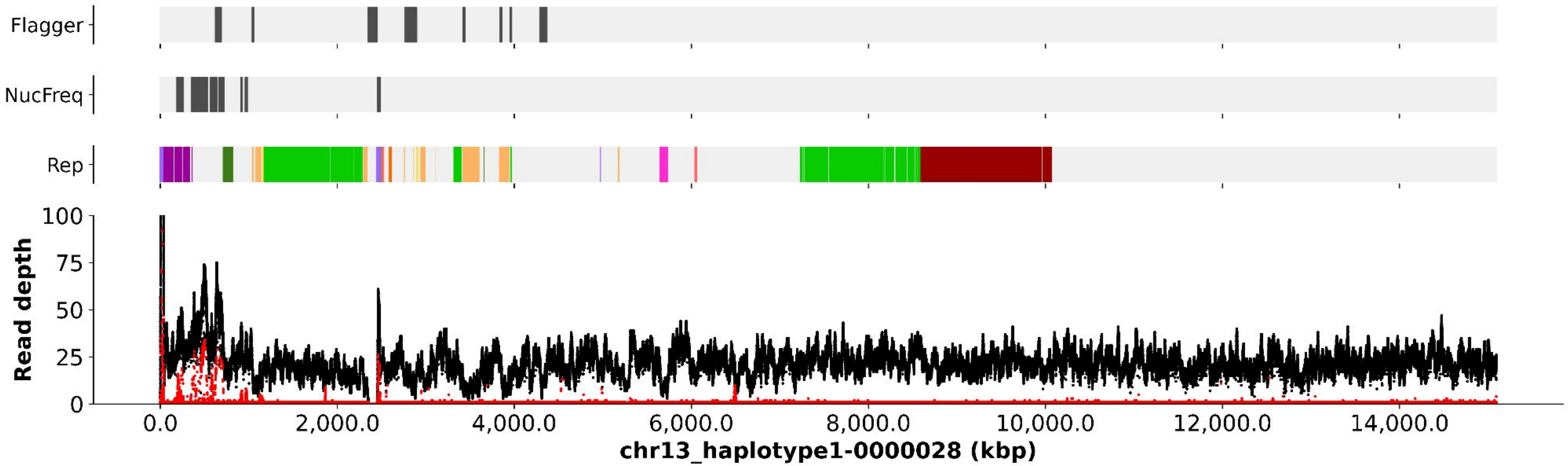
Flagger
NucFreq
Rep
Read depth
100
75
50
25
0
0.0
2,000.0
4,000.0
6,000.0
8,000.0
10,000.0
12,000.0
14,000.0
chr13_haplotype1-0000028 (kbp)

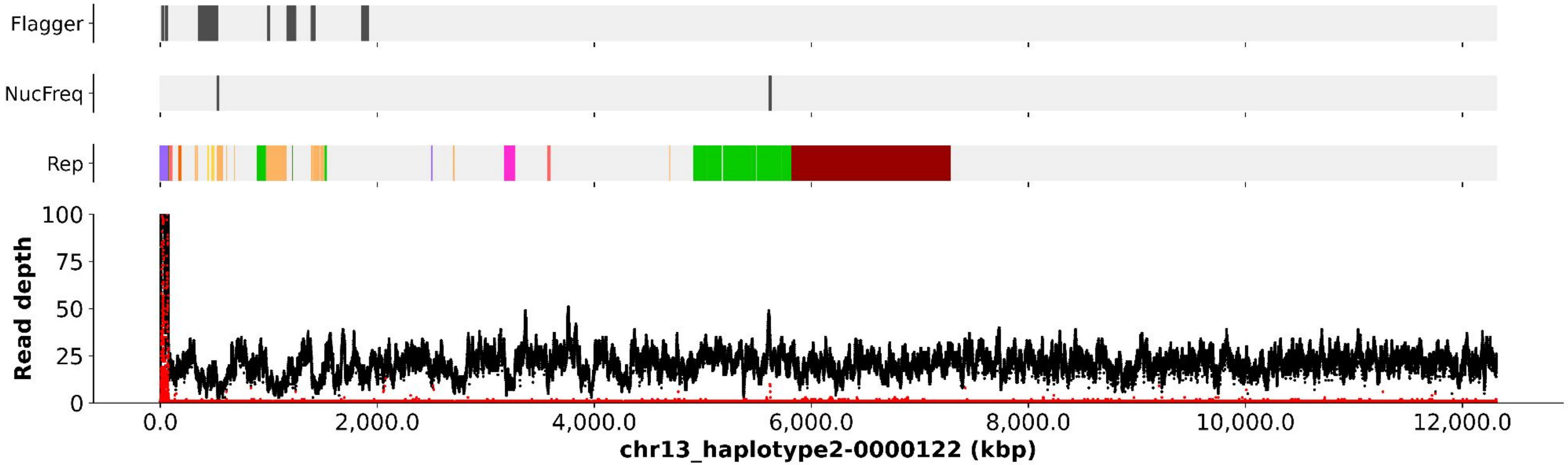
Flagger
NucFreq
Rep
Read depth
100
75
50
25
0
0.0
2,000.0
4,000.0
6,000.0
8,000.0
10,000.0
12,000.0
chr13_haplotype2-0000122 (kbp)

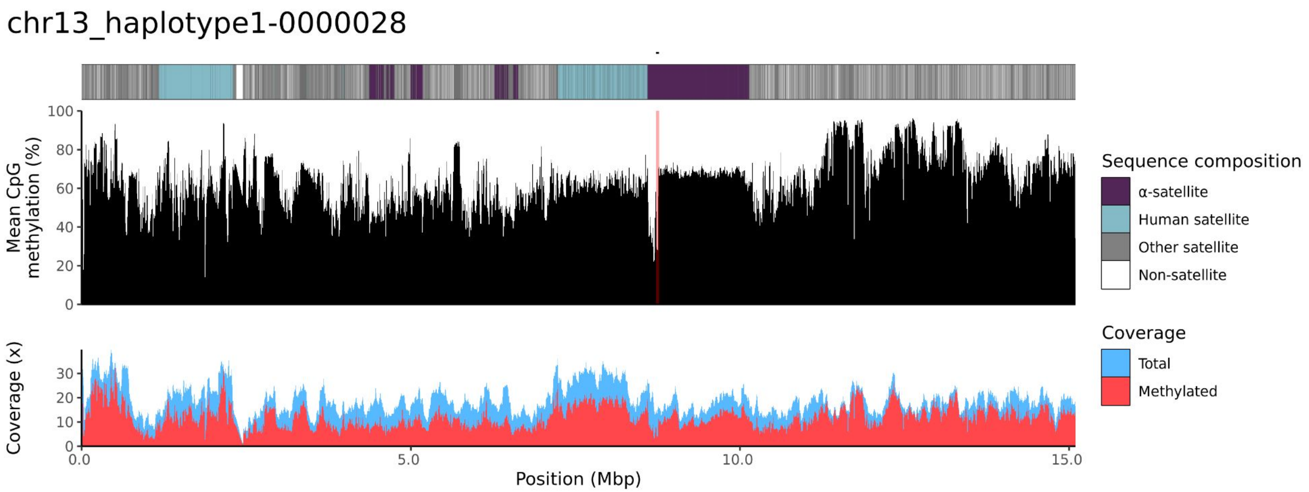

chr13_haplotype1-0000028
Mean CpG methylation (%)
Coverage (x)
Position (Mbp)
Sequence composition
α-satellite
Human satellite
Other satellite
Non-satellite
Coverage
Total
Methylated


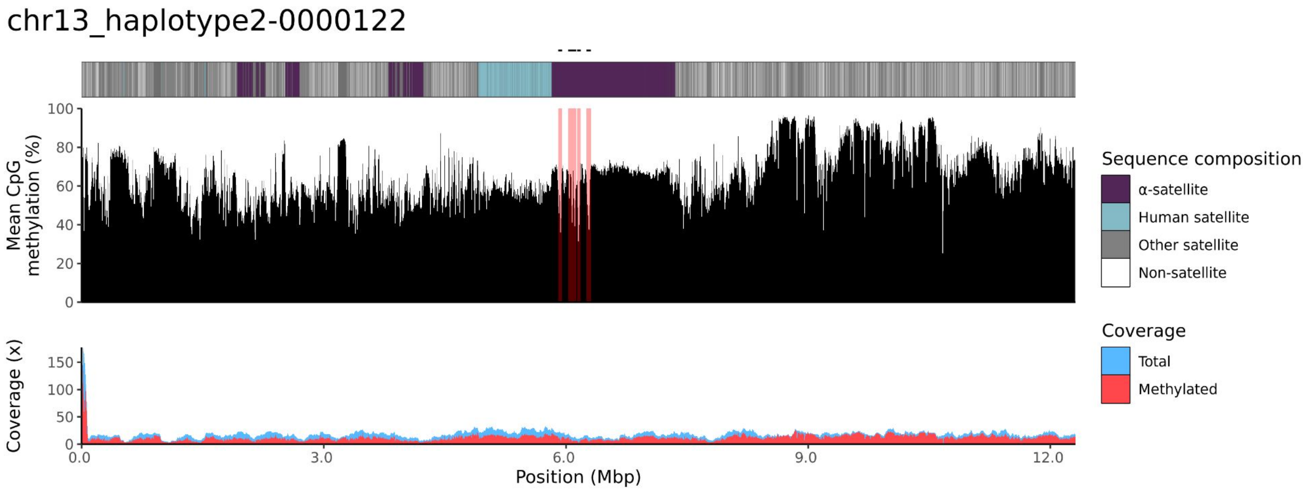

chr13_haplotype2-0000122
Mean CpG methylation (%)
Coverage (x)
Position (Mbp)
Sequence composition
α-satellite
Human satellite
Other satellite
Non-satellite
Coverage
Total
Methylated

### chr14

#### NA12882_1_haplotype1-0000016_chr14

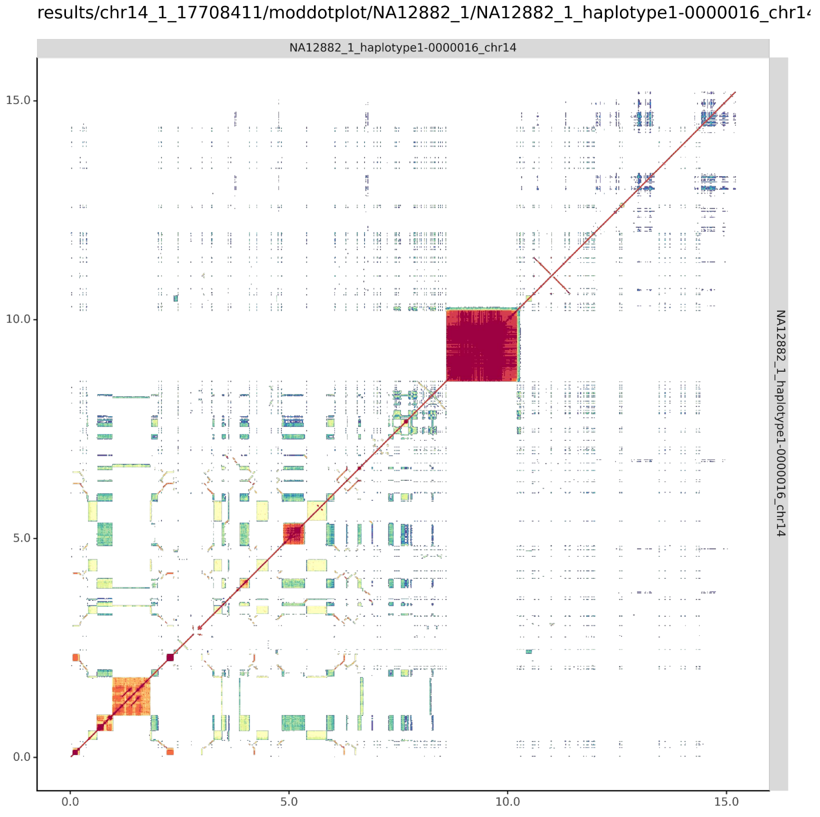


#### NA12882_2_haplotype2-0000112_chr14

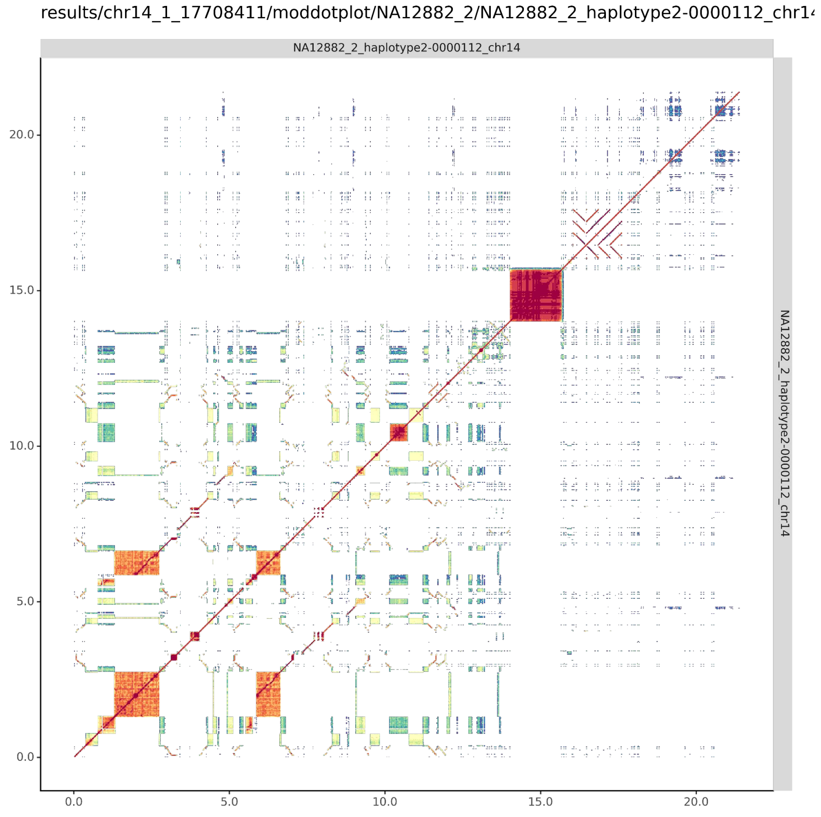

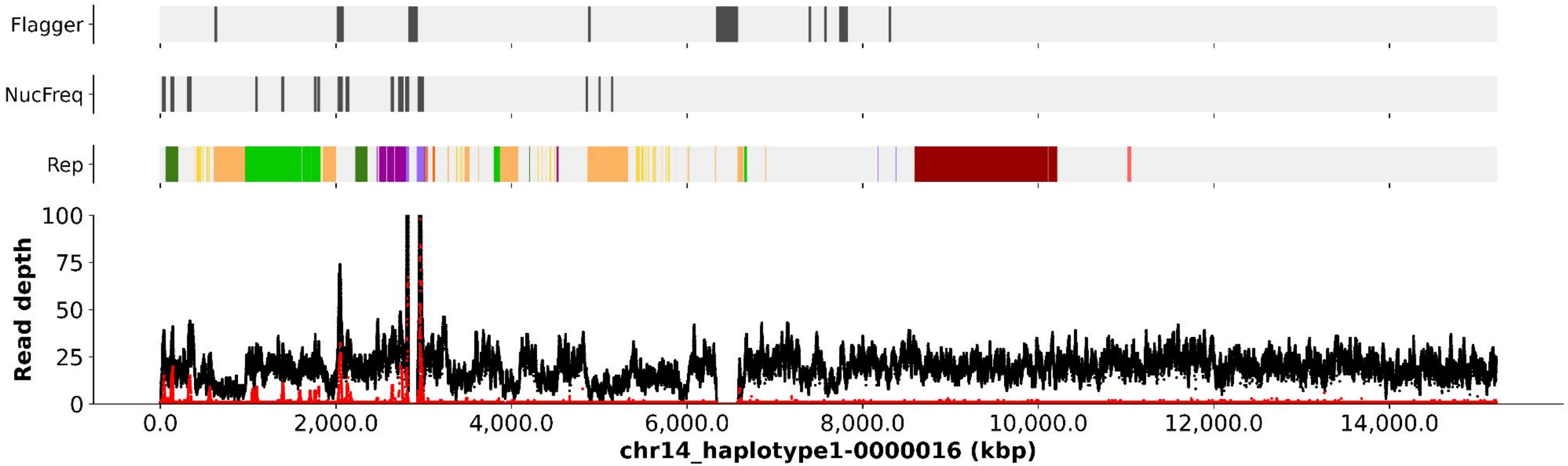

Flagger
NucFreq
Rep
Read depth
100
75
50
25
0
0.0
2,000.0
4,000.0
6,000.0
8,000.0
10,000.0
12,000.0
14,000.0
chr14_haplotype1-0000016 (kbp)


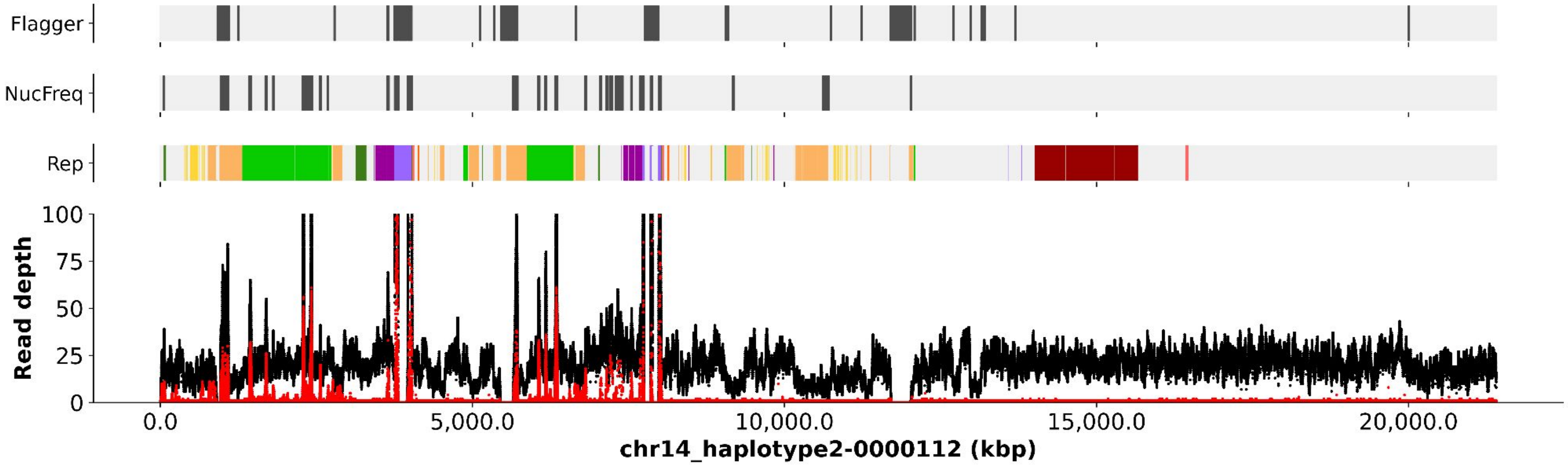

Flagger
NucFreq
Rep
Read depth
100
75
50
25
0
0.0
5,000.0
10,000.0
15,000.0
20,000.0
chr14_haplotype2-0000112 (kbp)

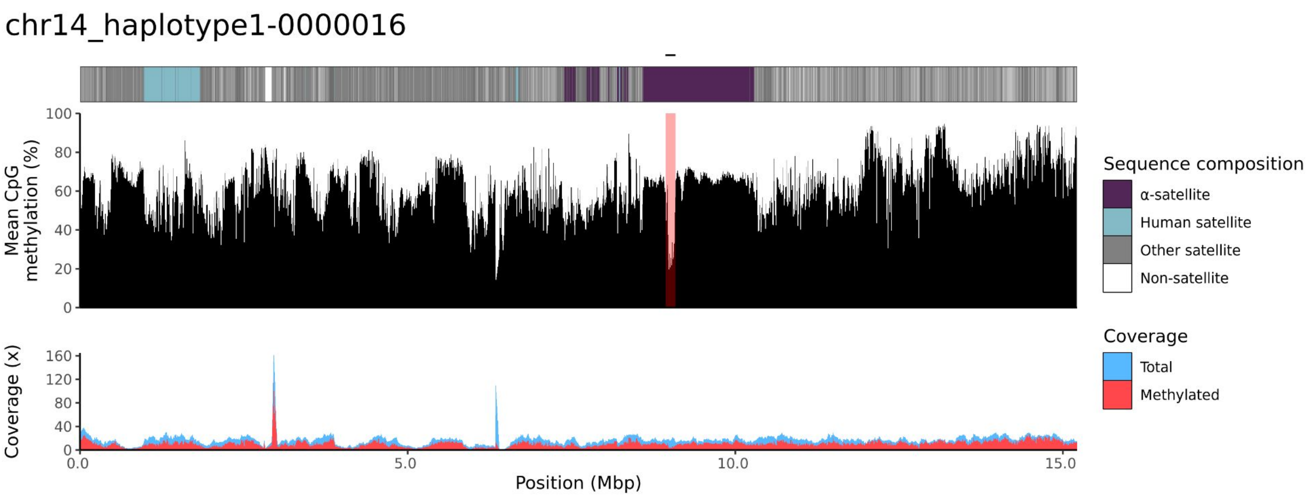
chr14_haplotype1-0000016
Mean CpG methylation (%)
Coverage (x)
Position (Mbp)
Sequence composition
α-satellite
Human satellite
Other satellite
Non-satellite
Coverage
Total
Methylated

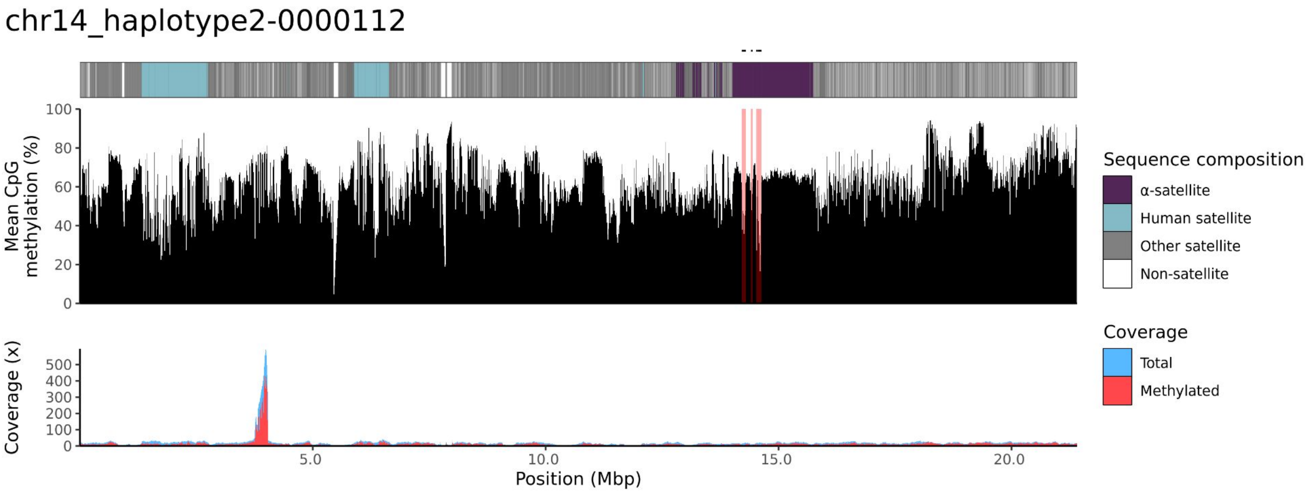
chr14_haplotype2-0000112
Mean CpG methylation (%)
Coverage (x)
Position (Mbp)
Sequence composition
α-satellite
Human satellite
Other satellite
Non-satellite
Coverage
Total
Methylated

### chr15

#### NA12882_1_haplotype1-0000019_chr15

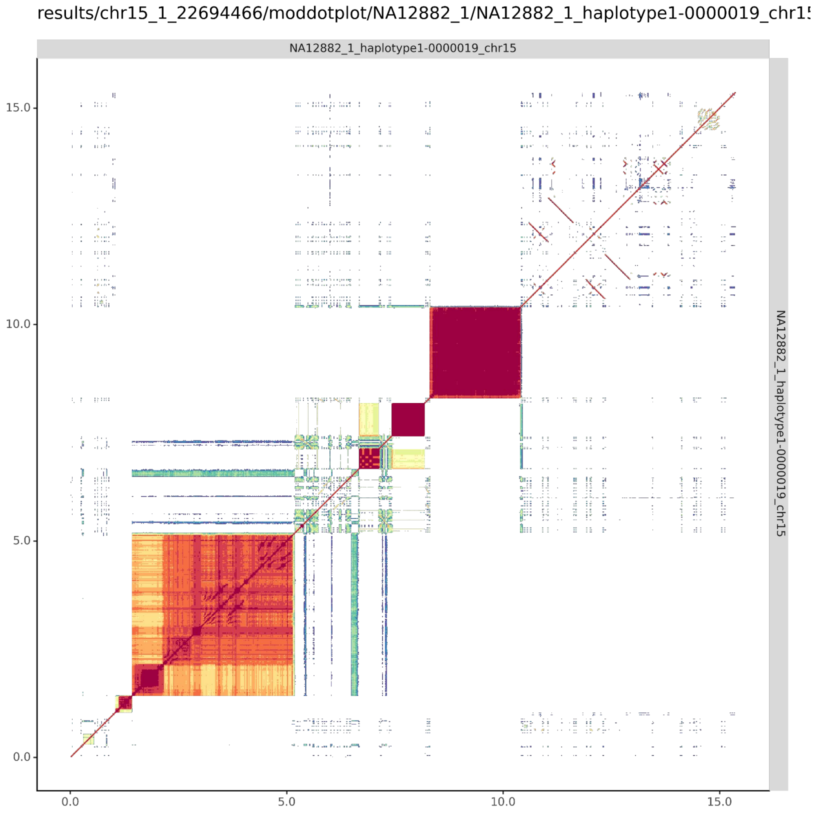


#### NA12882_2_haplotype2-0000116_chr15

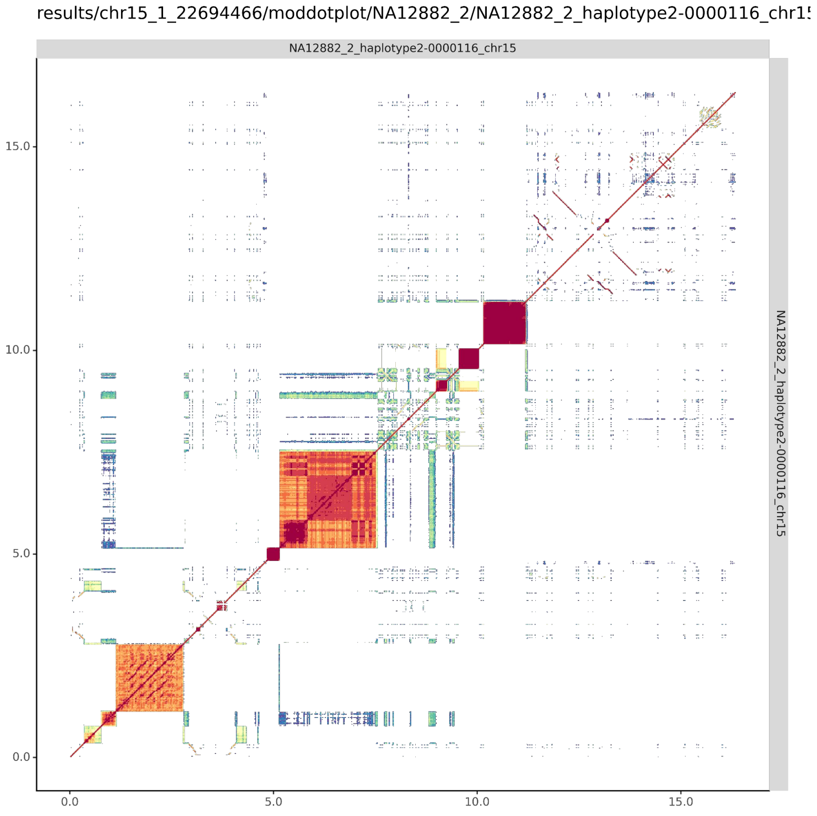

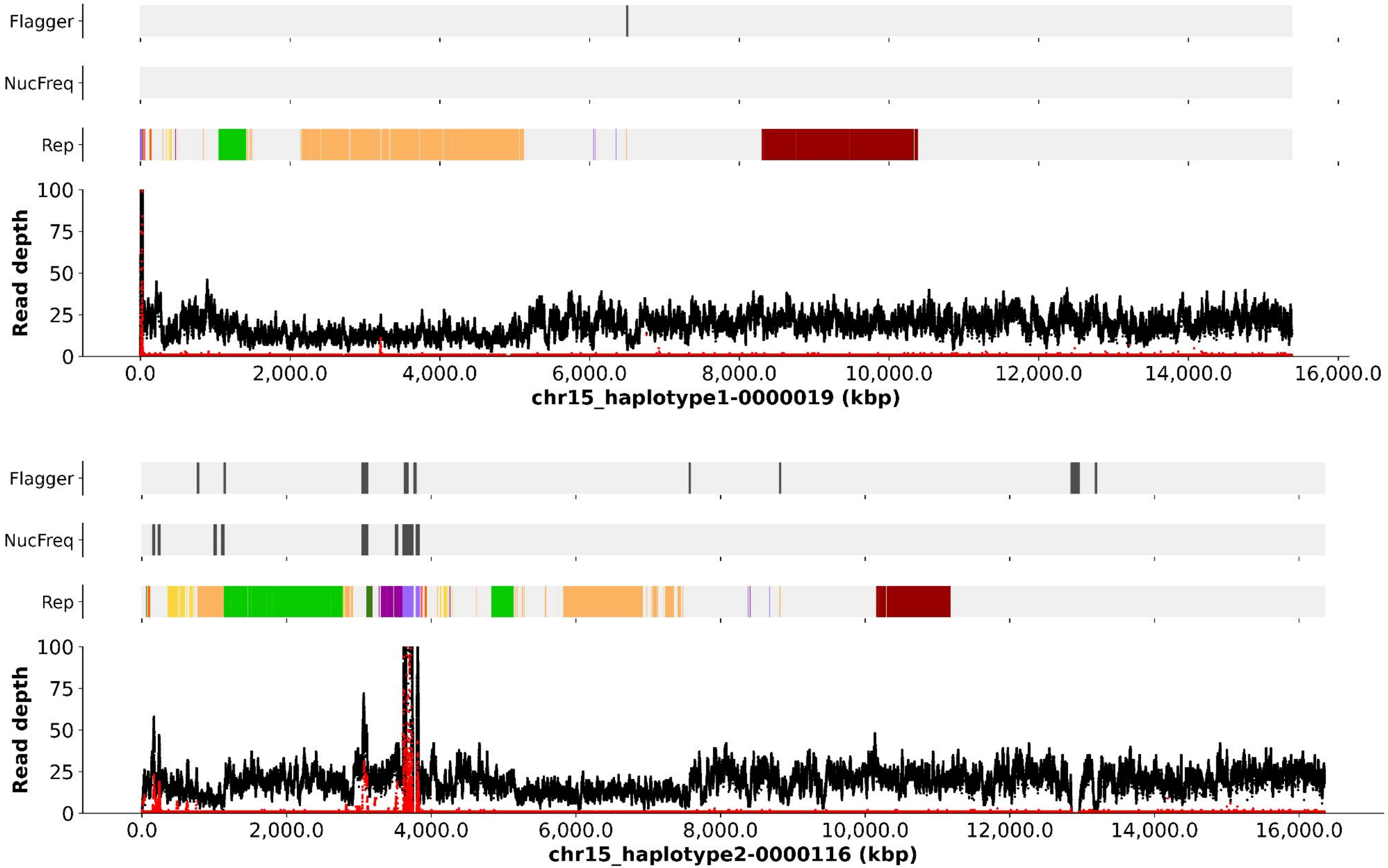
Flagger
NucFreq
Rep
Read depth
100
75
50
25
0
0.0
2,000.0
4,000.0
6,000.0
8,000.0
10,000.0
12,000.0
14,000.0
16,000.0
chr15_haplotype1-0000019 (kbp)
Flagger
NucFreq
Rep
Read depth
100
75
50
25
0
0.0
2,000.0
4,000.0
6,000.0
8,000.0
10,000.0
12,000.0
14,000.0
16,000.0
chr15_haplotype2-0000116 (kbp)

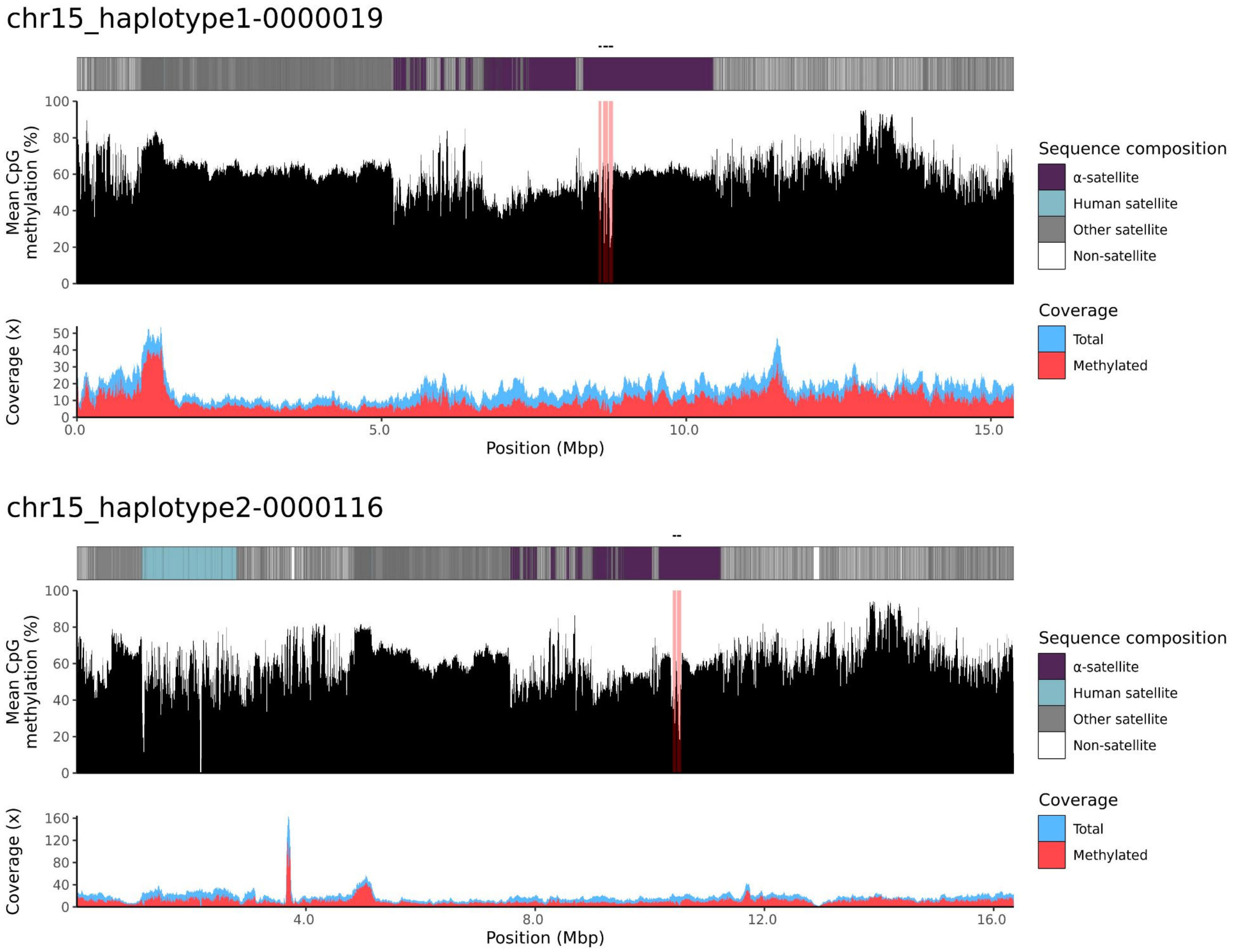

chr15_haplotype1-0000019
100
80
60
40
20
0
Mean CpG methylation (%)
50
40
30
20
10
0
Coverage (x)
0.0
5.0
10.0
15.0
Position (Mbp)
Sequence composition
α-satellite
Human satellite
Other satellite
Non-satellite
Coverage
Total
Methylated
chr15_haplotype2-0000116
100
80
60
40
20
0
Mean CpG methylation (%)
160
120
80
40
0
Coverage (x)
4.0
8.0
12.0
16.0
Position (Mbp)
Sequence composition
α-satellite
Human satellite
Other satellite
Non-satellite
Coverage
Total
Methylated

### chr21

#### NA12882_1_haplotype1-0000010_chr21

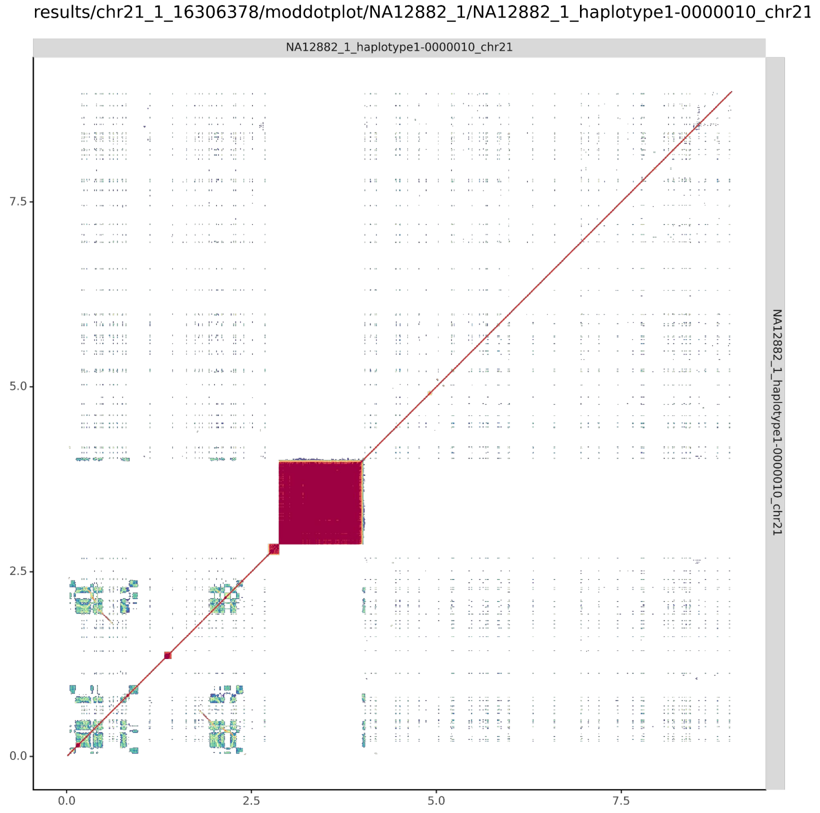


#### NA12882_2_haplotype2-0000107_chr21

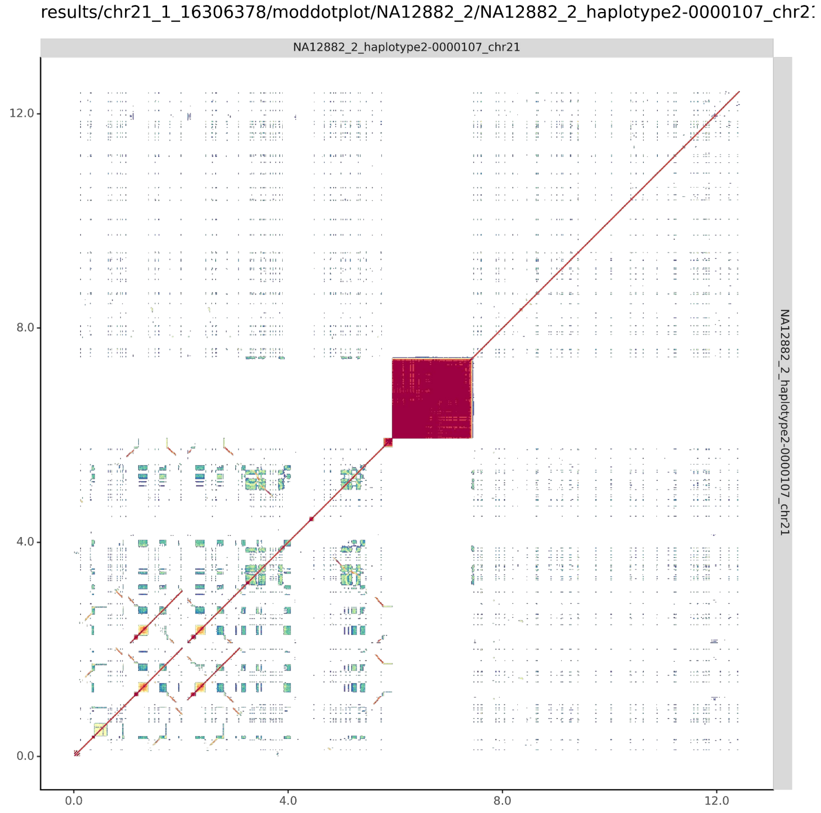

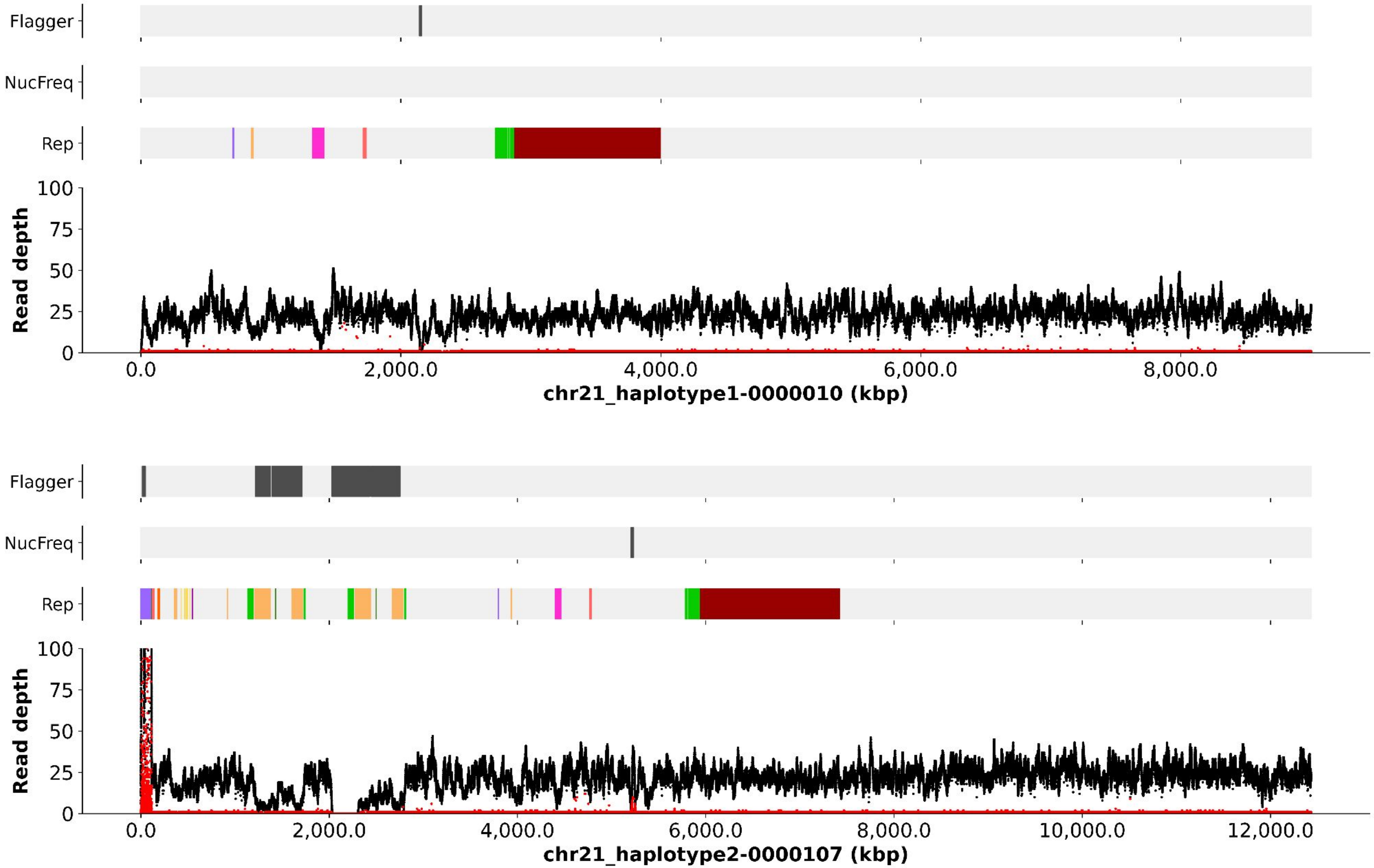

Flagger
NucFreq
Rep
Read depth
100
75
50
25
0
0.0
2,000.0
4,000.0
6,000.0
8,000.0
chr21_haplotype1-0000010 (kbp)
Flagger
NucFreq
Rep
Read depth
100
75
50
25
0
0.0
2,000.0
4,000.0
6,000.0
8,000.0
10,000.0
12,000.0
chr21_haplotype2-0000107 (kbp)

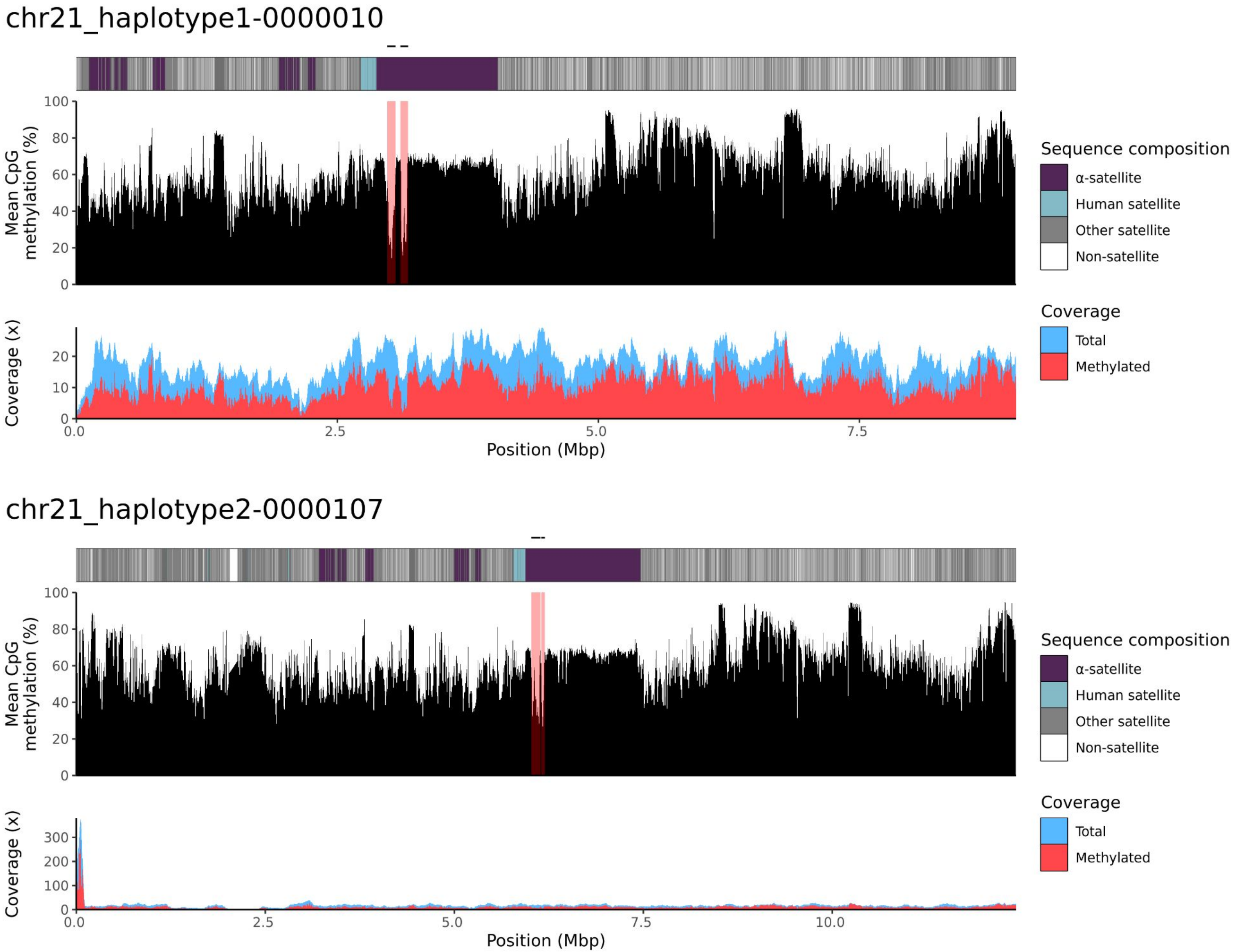

chr21_haplotype1-0000010
Mean CpG methylation (%)
Coverage (x)
Position (Mbp)
Sequence composition
α-satellite
Human satellite
Other satellite
Non-satellite
Coverage
Total
Methylated
chr21_haplotype2-0000107
Mean CpG methylation (%)
Coverage (x)
Position (Mbp)
Sequence composition
α-satellite
Human satellite
Other satellite
Non-satellite
Coverage
Total
Methylated

### chr22

#### NA12882_1_haplotype1-0000004_chr22

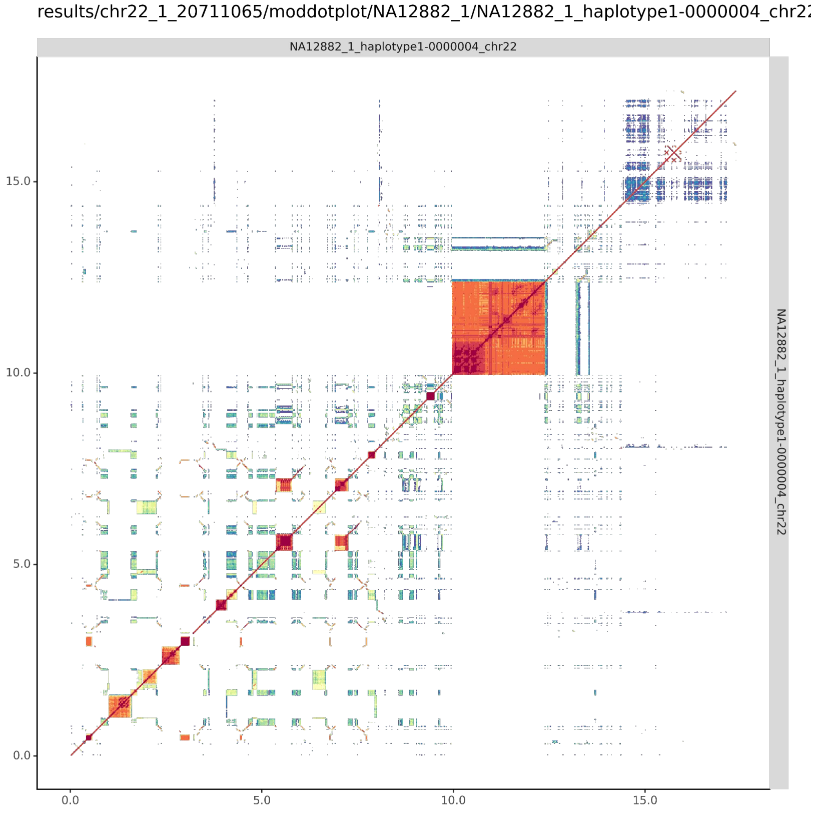


#### NA12882_2_haplotype2-0000099_chr22

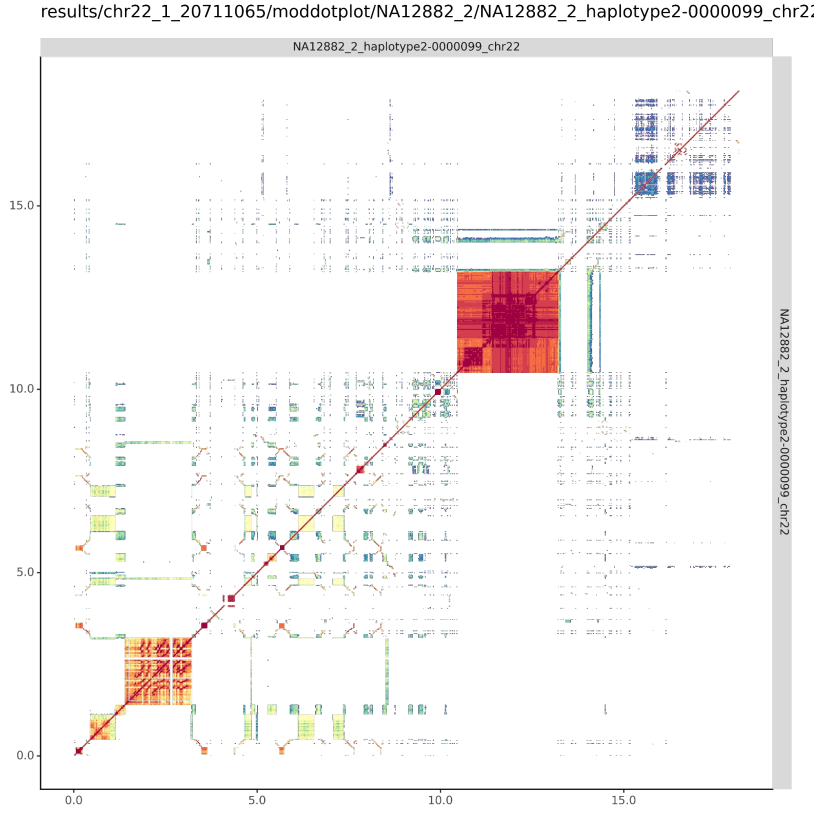

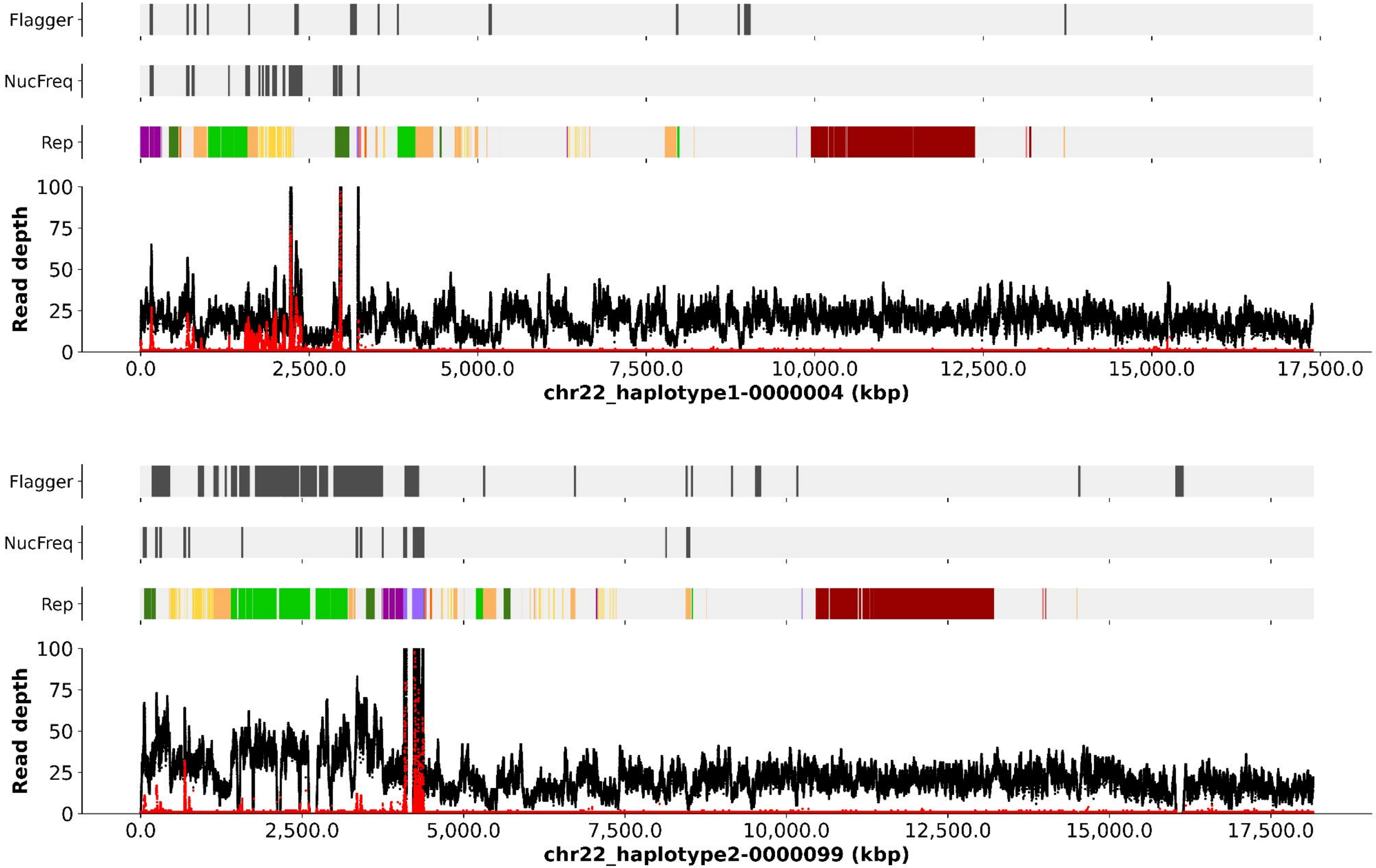
Flagger
NucFreq
Rep
Read depth
chr22_haplotype1-0000004 (kbp)
Flagger
NucFreq
Rep
Read depth
chr22_haplotype2-0000099 (kbp)

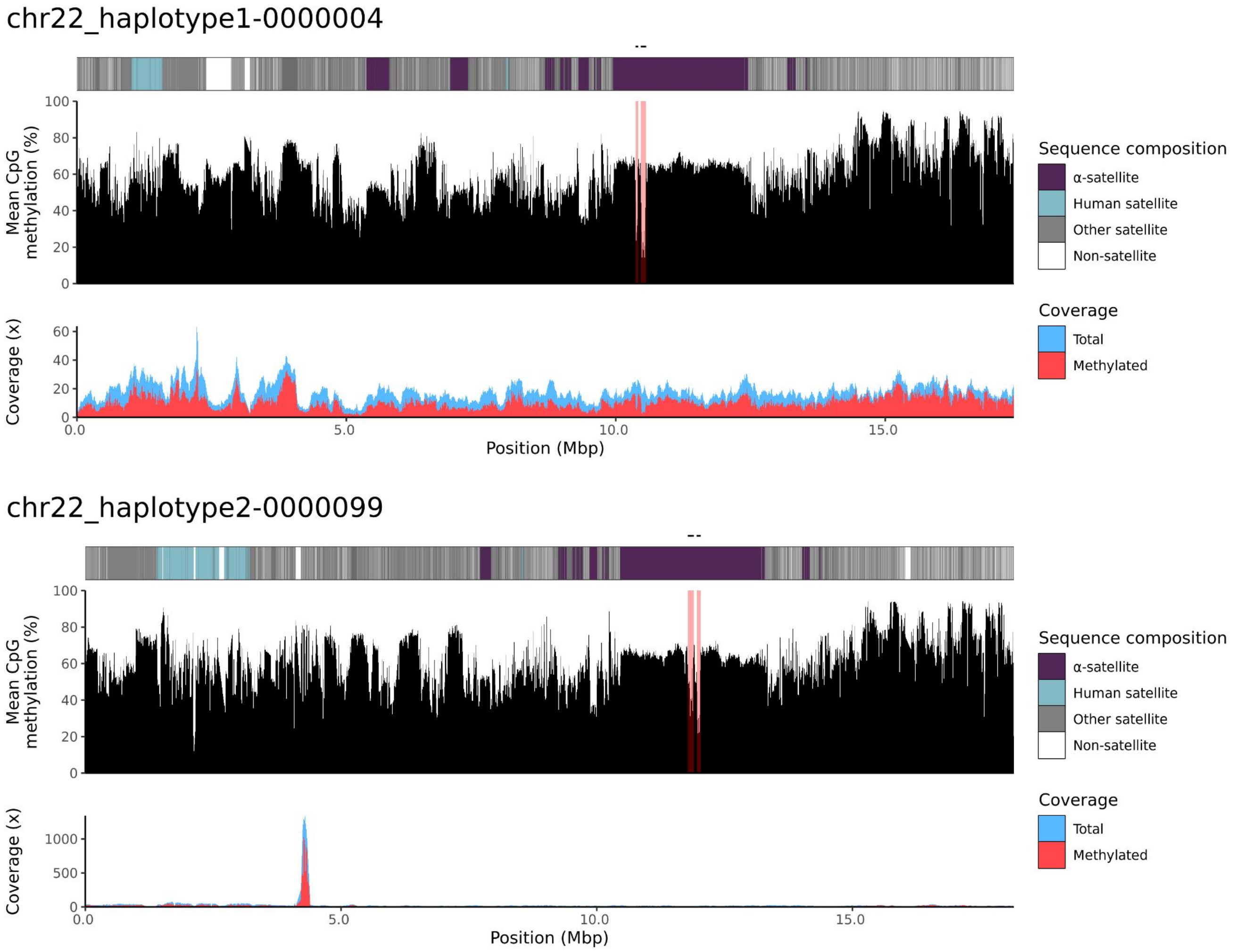
chr22_haplotype1-0000004
Mean CpG methylation (%)
Coverage (x)
Position (Mbp)
Sequence composition
α-satellite
Human satellite
Other satellite
Non-satellite
Coverage
Total
Methylated
chr22_haplotype2-0000099
Mean CpG methylation (%)
Coverage (x)
Position (Mbp)
Sequence composition
α-satellite
Human satellite
Other satellite
Non-satellite
Coverage
Total
Methylated

NA12883

### chr13

#### NA12883_1_haplotype1-0000004_chr13

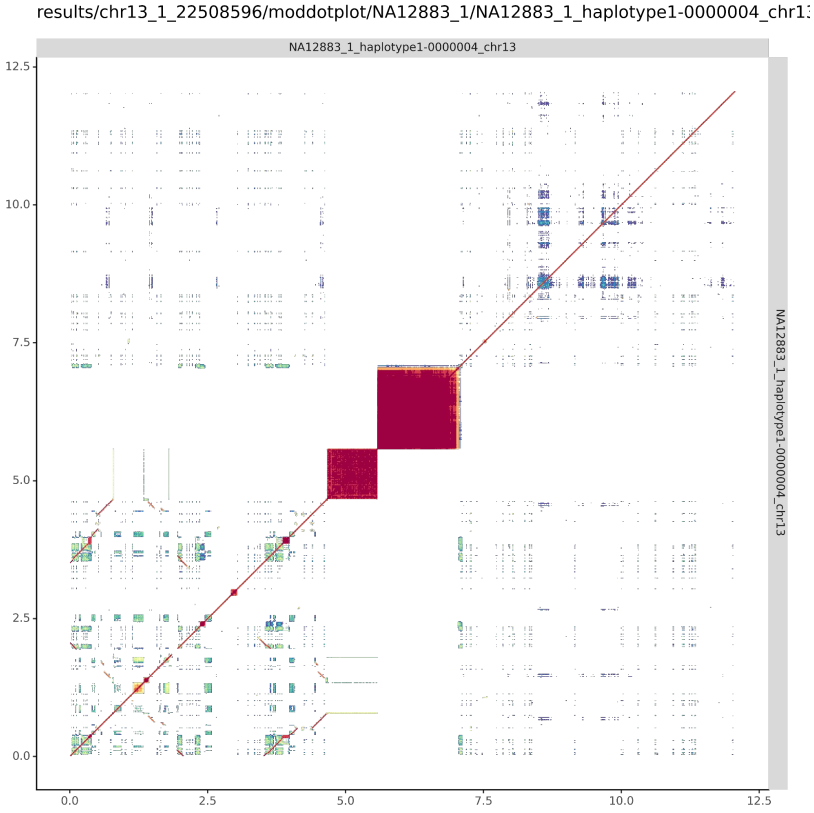


#### NA12883_2_haplotype2-0000078_chr13

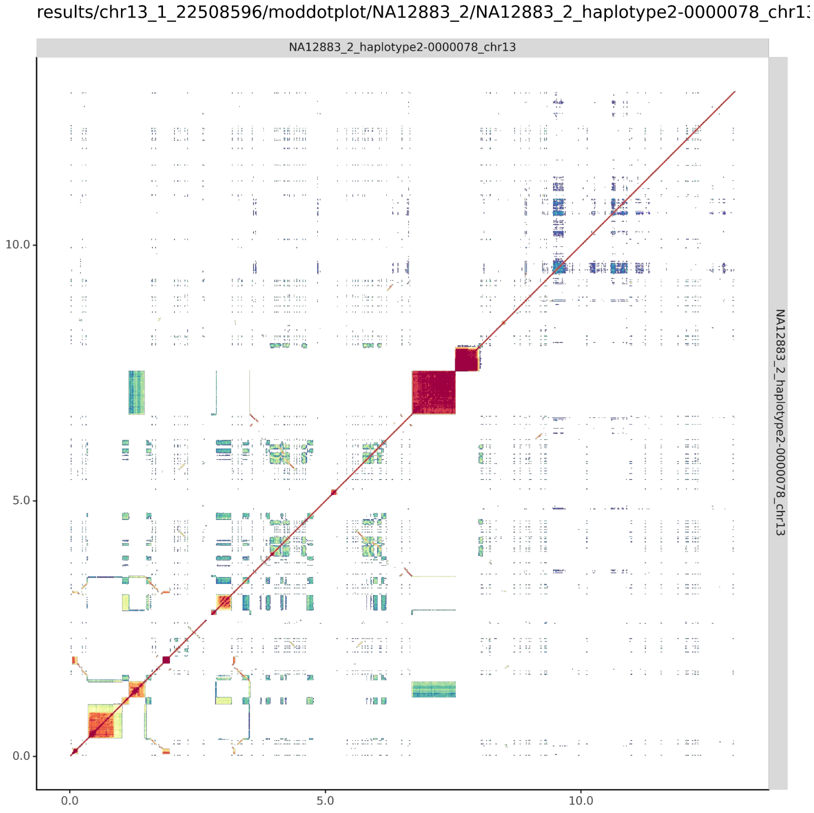

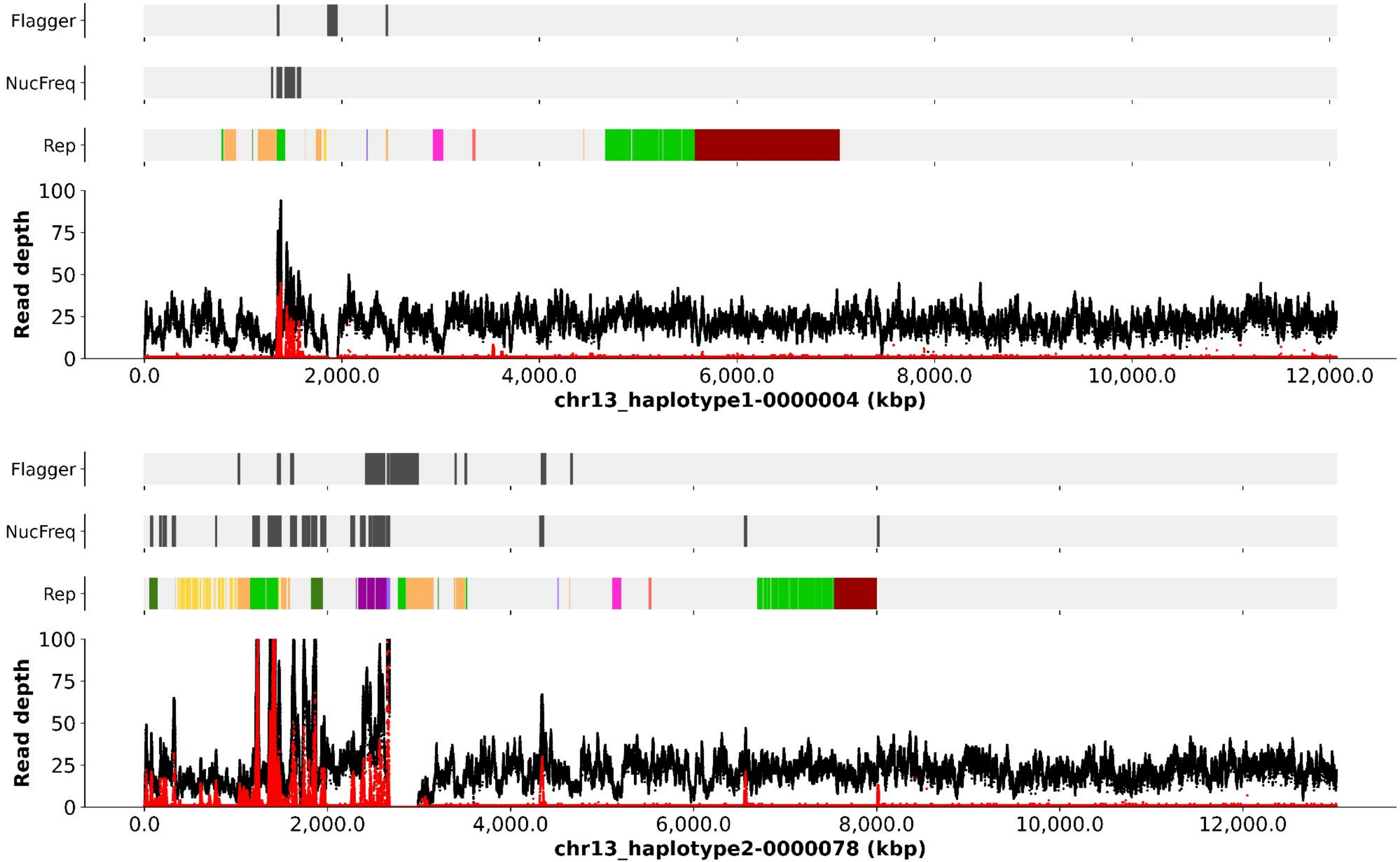
Flagger
NucFreq
Rep
Read depth
100
75
50
25
0
0.0
2,000.0
4,000.0
6,000.0
8,000.0
10,000.0
12,000.0
chr13_haplotype1-0000004 (kbp)
Flagger
NucFreq
Rep
Read depth
100
75
50
25
0
0.0
2,000.0
4,000.0
6,000.0
8,000.0
10,000.0
12,000.0
chr13_haplotype2-0000078 (kbp)

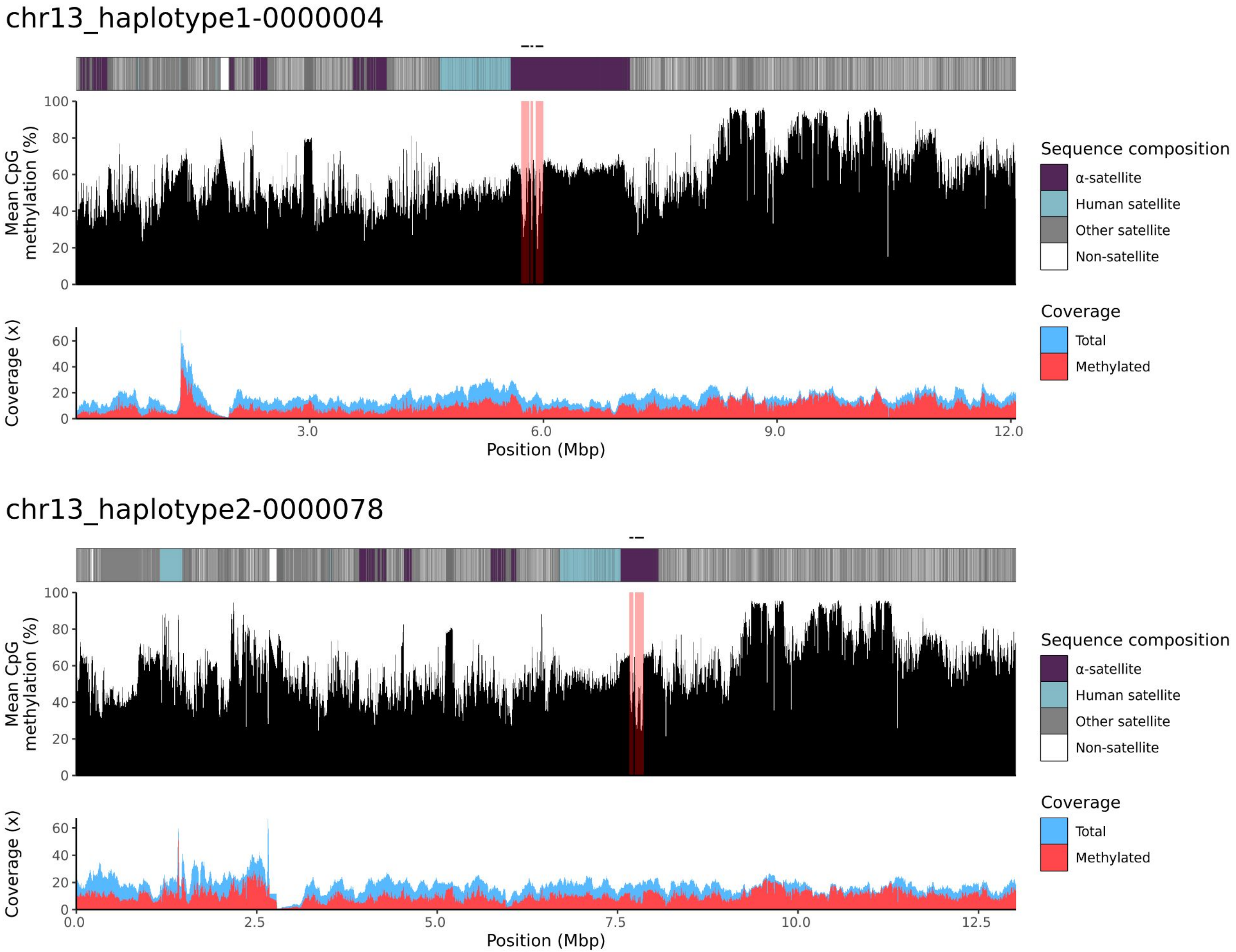
chr13_haplotype1-0000004
Mean CpG methylation (%)
Coverage (x)
Position (Mbp)
Sequence composition
α-satellite
Human satellite
Other satellite
Non-satellite
Coverage
Total
Methylated
chr13_haplotype2-0000078
Mean CpG methylation (%)
Coverage (x)
Position (Mbp)
Sequence composition
α-satellite
Human satellite
Other satellite
Non-satellite
Coverage
Total
Methylated

### chr14

#### NA12883_1_haplotype1-0000025_chr14

results/chr14_1_17708411/moddotplot/NA12883_1/NA12883_1_haplotype1-0000025_chr14

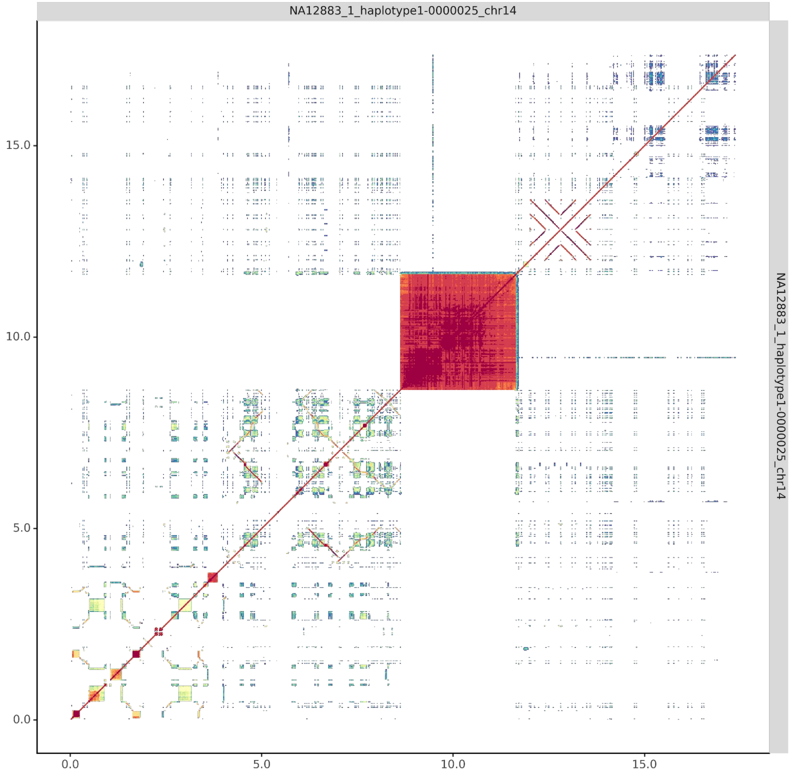


#### NA12883_2_haplotype2-0000084_chr14

results/chr14_1_17708411/moddotplot/NA12883_2/NA12883_2_haplotype2-0000084_chr14

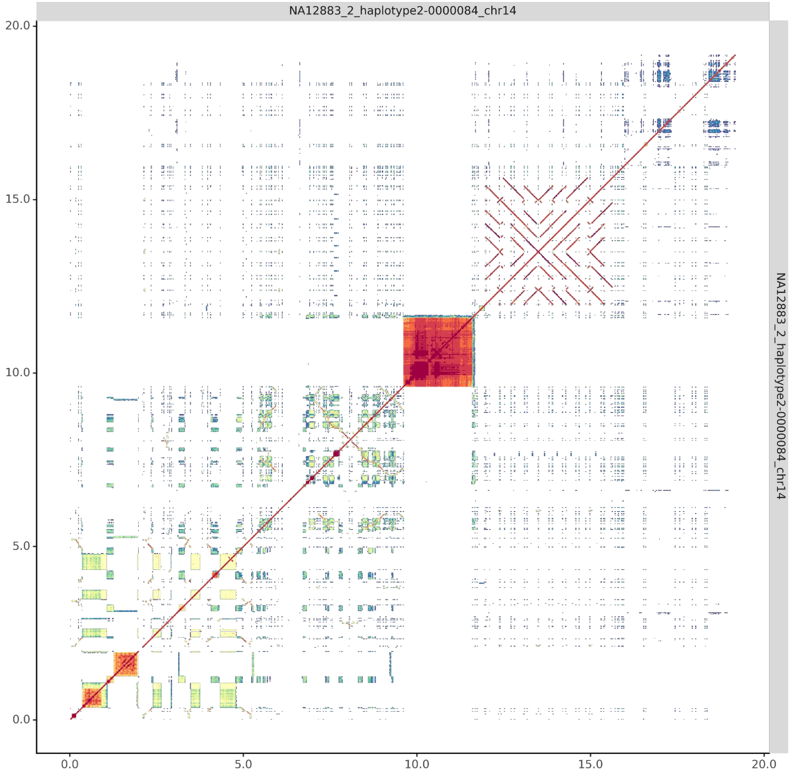

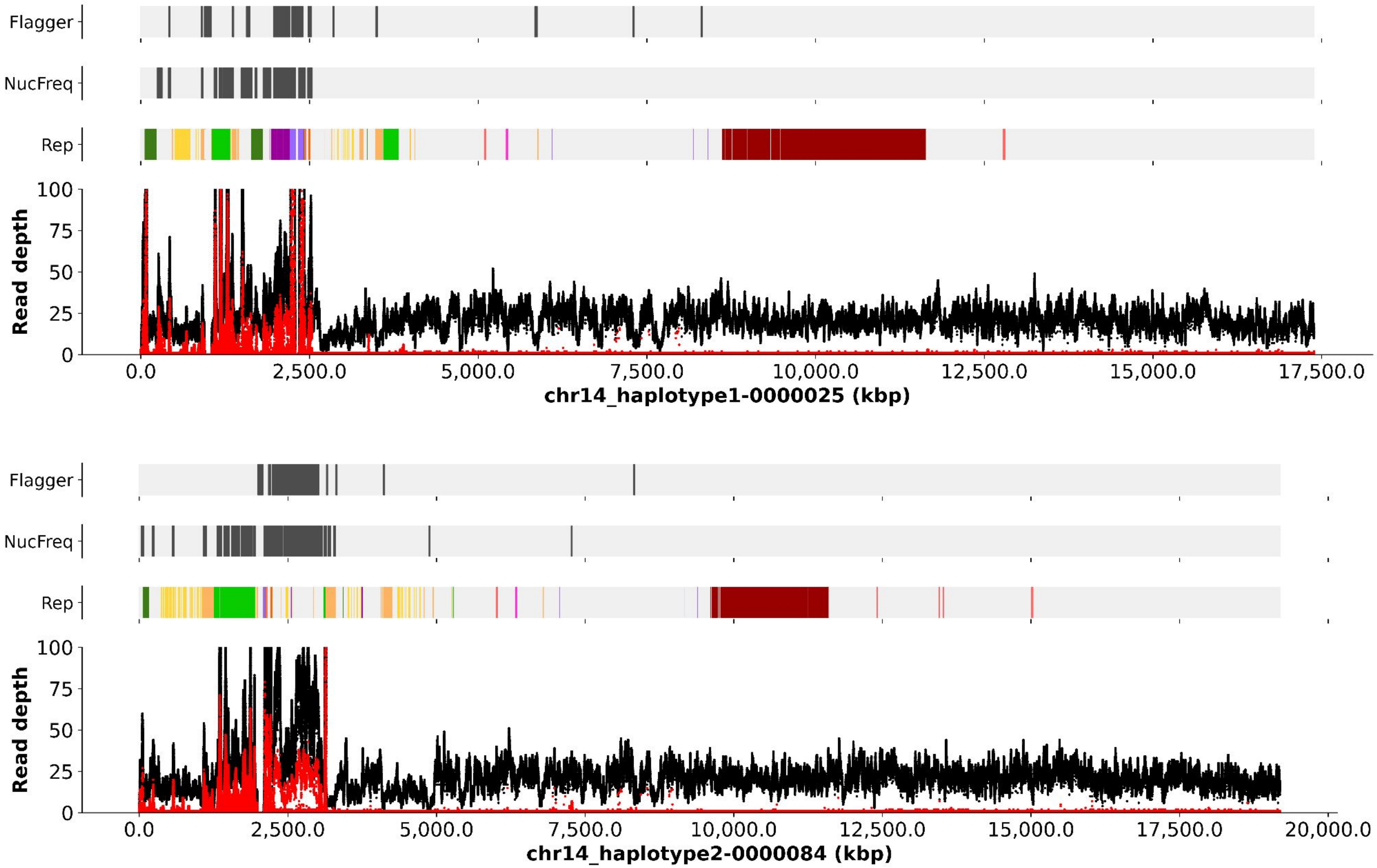

Flagger
NucFreq
Rep
Read depth
100
75
50
25
0
0.0
2,500.0
5,000.0
7,500.0
10,000.0
12,500.0
15,000.0
17,500.0
chr14_haplotype1-0000025 (kbp)
Flagger
NucFreq
Rep
Read depth
100
75
50
25
0
0.0
2,500.0
5,000.0
7,500.0
10,000.0
12,500.0
15,000.0
17,500.0
20,000.0
chr14_haplotype2-0000084 (kbp)

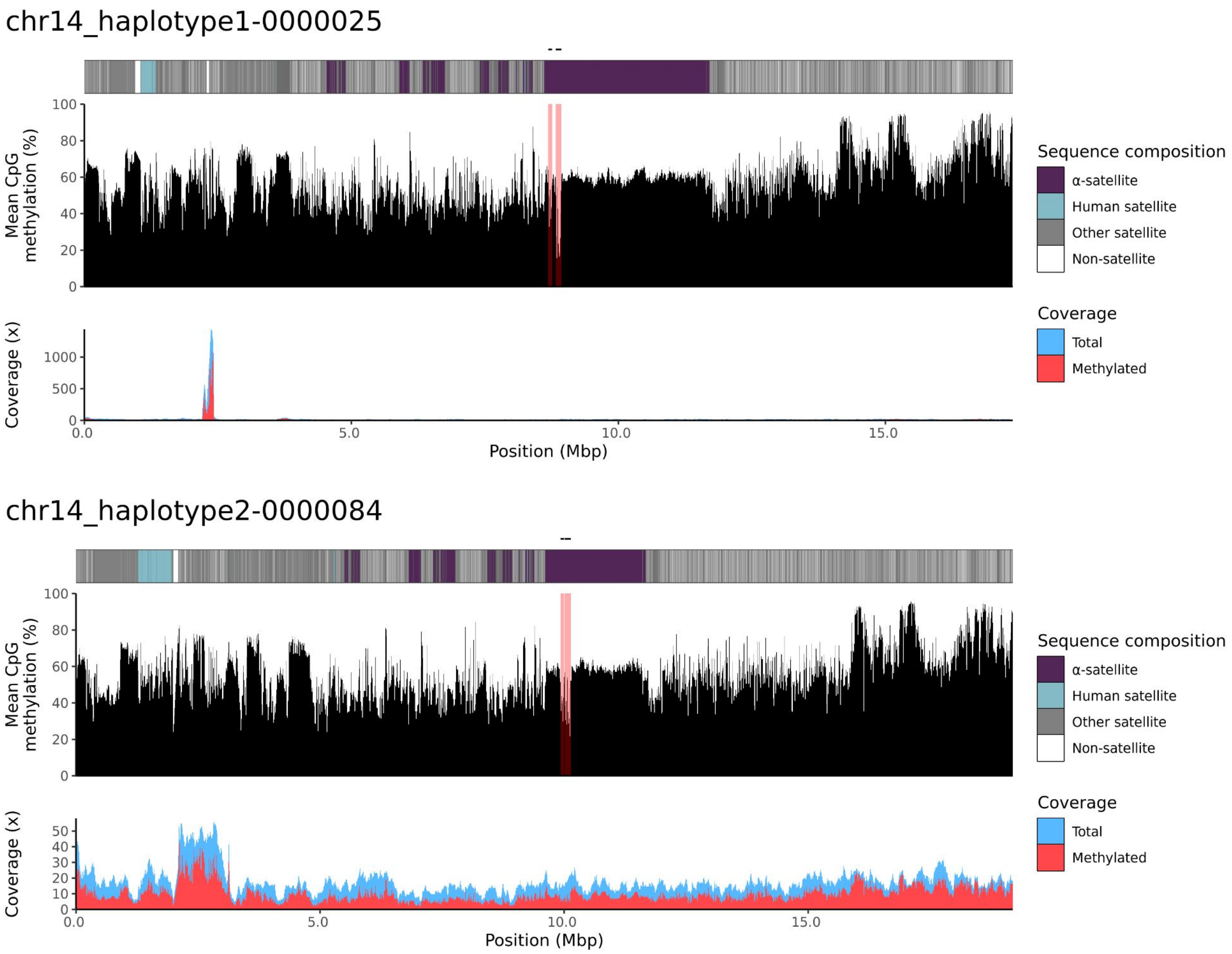
chr14_haplotype1-0000025
Mean CpG methylation (%)
100
80
60
40
20
0
Coverage (x)
1000
500
0
0.0
5.0
10.0
15.0
Position (Mbp)
Sequence composition
α-satellite
Human satellite
Other satellite
Non-satellite
Coverage
Total
Methylated
chr14_haplotype2-0000084
Mean CpG methylation (%)
100
80
60
40
20
0
Coverage (x)
50
40
30
20
10
0
0.0
5.0
10.0
15.0
Position (Mbp)
Sequence composition
α-satellite
Human satellite
Other satellite
Non-satellite
Coverage
Total
Methylated

### chr15

#### NA12883_2_haplotype2-0000087_chr15

results/chr15_1_22694466/moddotplot/NA12883_2/NA12883_2_haplotype2-0000087_chr15

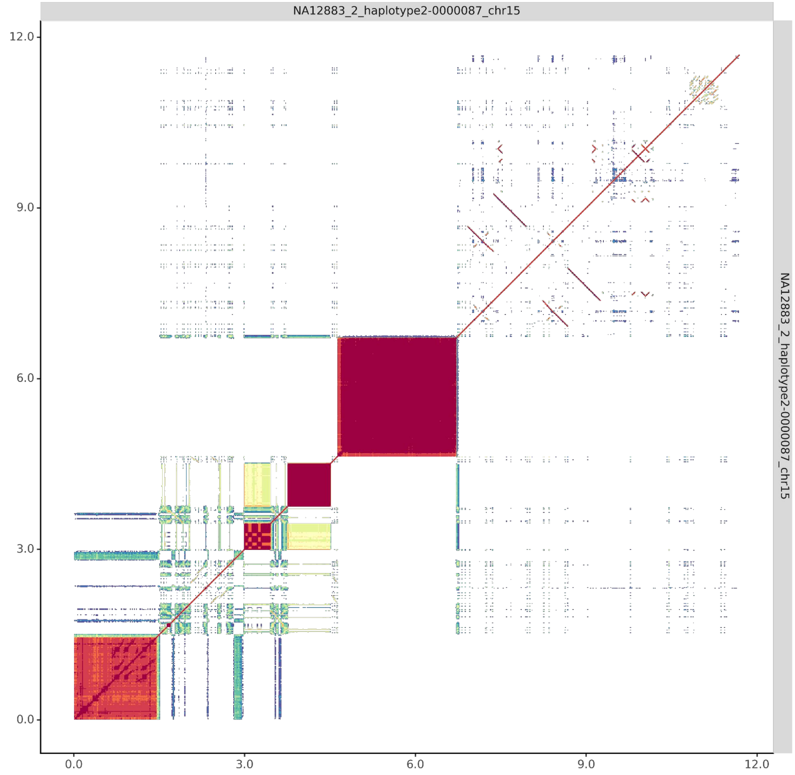

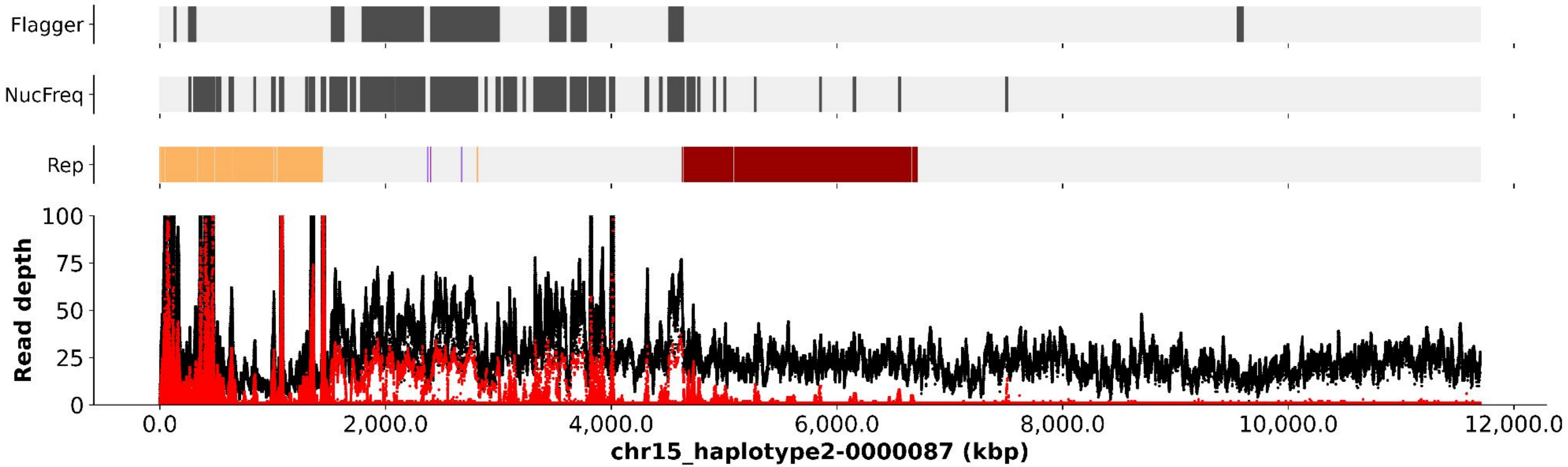
Flagger
NucFreq
Rep
Read depth
100
75
50
25
0
0.0
2,000.0
4,000.0
6,000.0
8,000.0
10,000.0
12,000.0
chr15_haplotype2-0000087 (kbp)

chr15_haplotype2-0000087

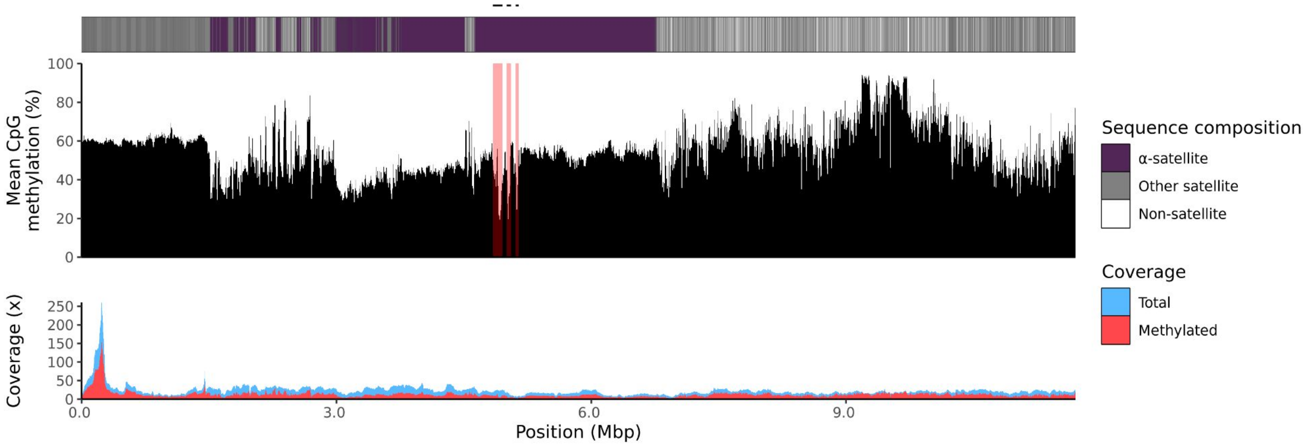

### chr21

#### NA12883_1_haplotype1-0000019_chr21

results/chr21_1_16306378/moddotplot/NA12883_1/NA12883_1_haplotype1-0000019_chr21

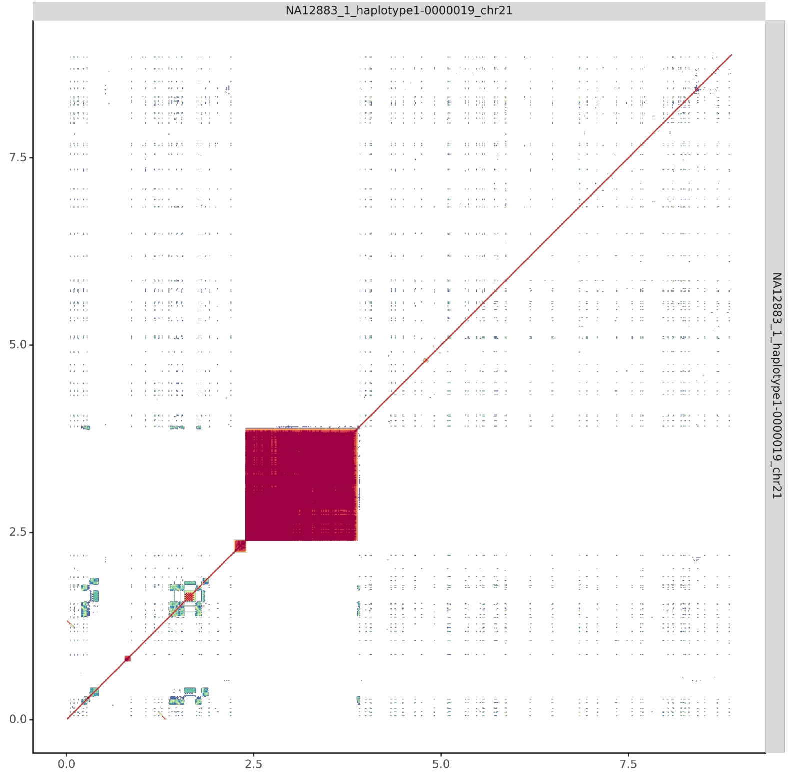


#### NA12883_2_haplotype2-0000080_chr21

results/chr21_1_16306378/moddotplot/NA12883_2/NA12883_2_haplotype2-0000080_chr21

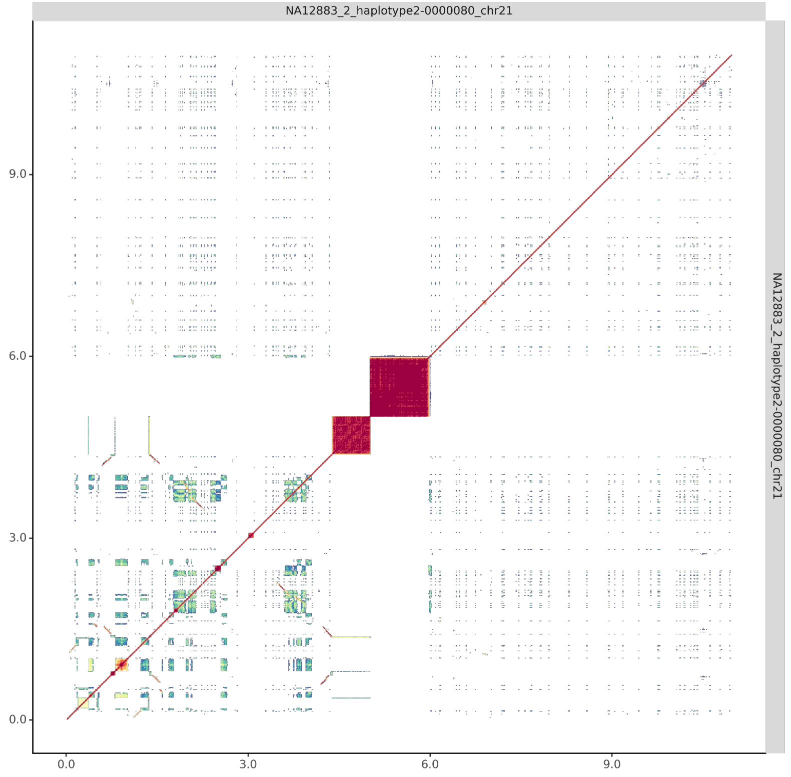

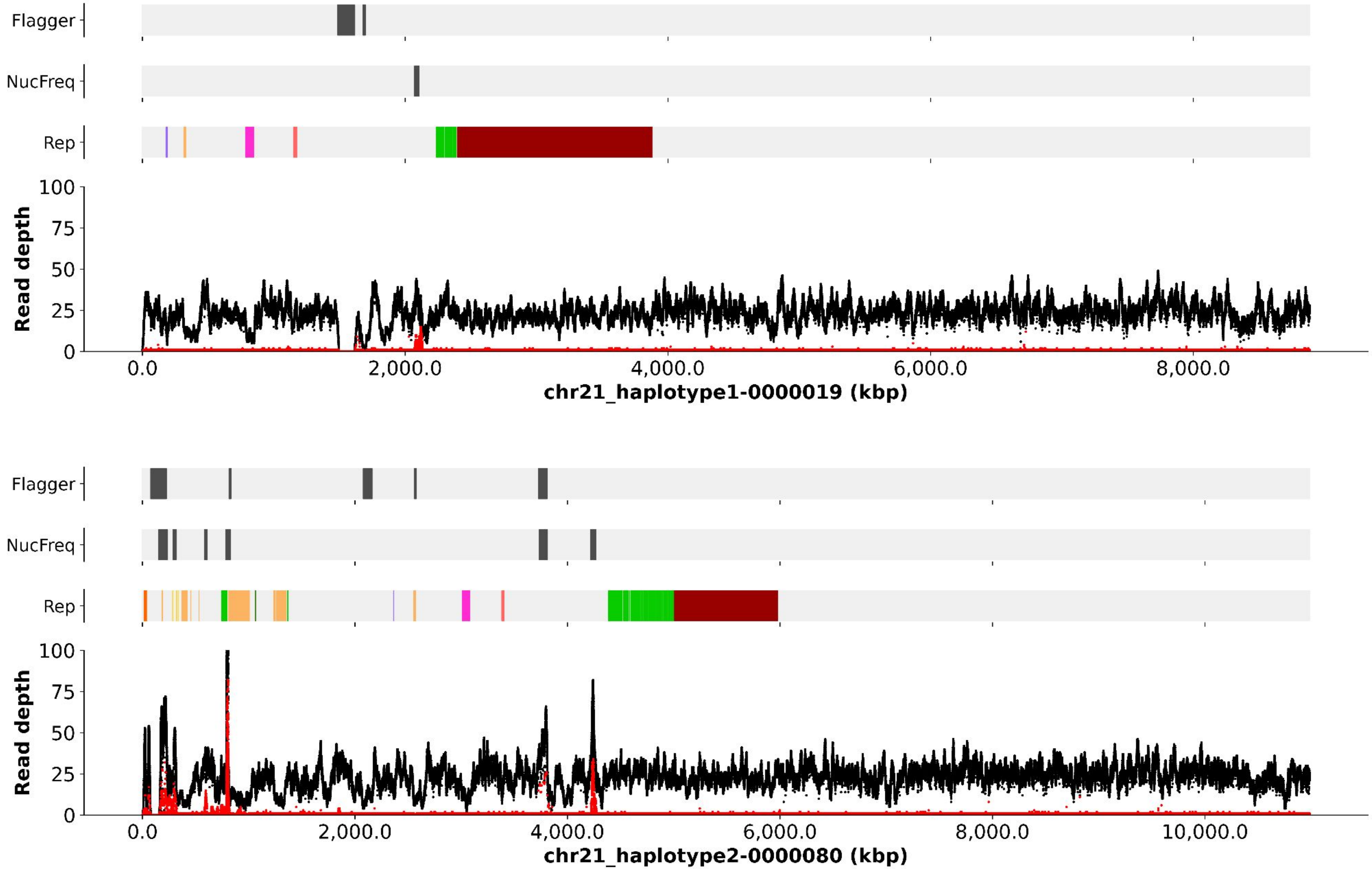
Flagger
NucFreq
Rep
Read depth
100
75
50
25
0
0.0
2,000.0
4,000.0
6,000.0
8,000.0
chr21_haplotype1-0000019 (kbp)
Flagger
NucFreq
Rep
Read depth
100
75
50
25
0
0.0
2,000.0
4,000.0
6,000.0
8,000.0
10,000.0
chr21_haplotype2-0000080 (kbp)

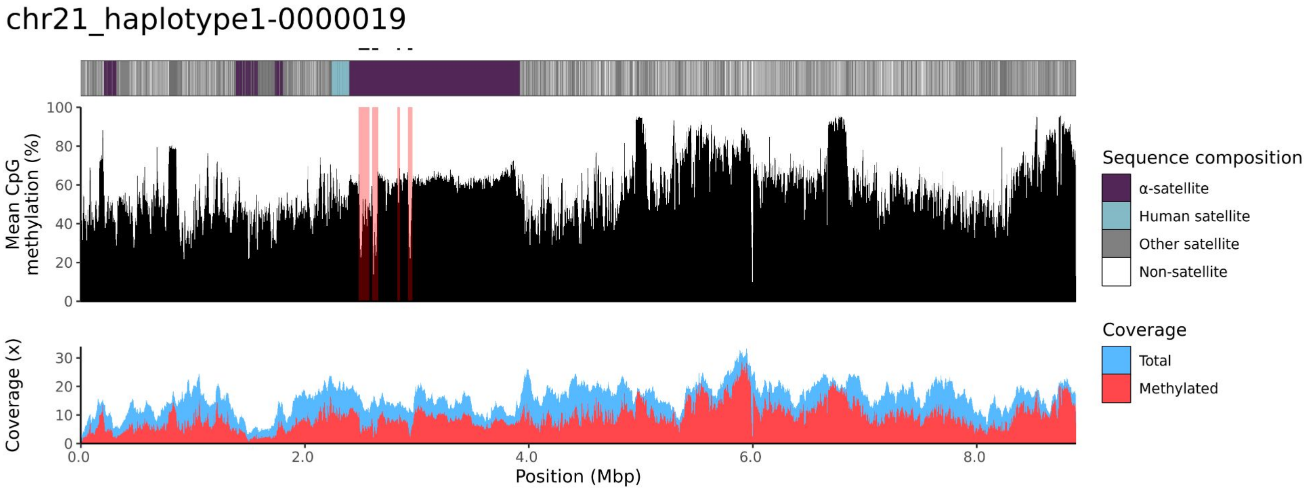
chr21_haplotype1-0000019
Mean CpG methylation (%)
Coverage (x)
Position (Mbp)
Sequence composition
α-satellite
Human satellite
Other satellite
Non-satellite
Coverage
Total
Methylated

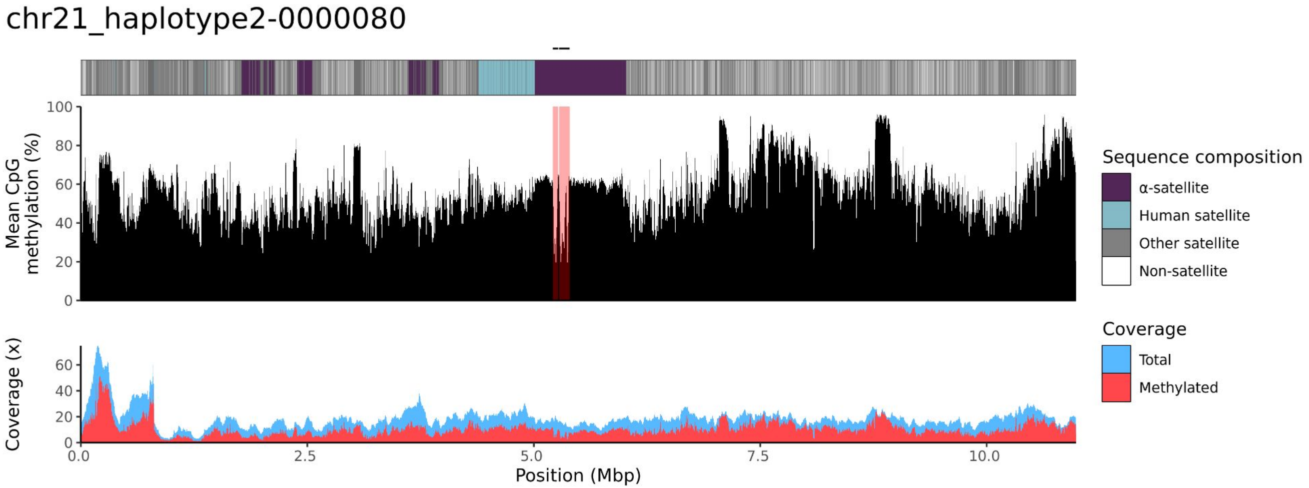
chr21_haplotype2-0000080
Mean CpG methylation (%)
Coverage (x)
Position (Mbp)
Sequence composition
α-satellite
Human satellite
Other satellite
Non-satellite
Coverage
Total
Methylated

### chr22

#### NA12883_1_haplotype1-0000024_chr22

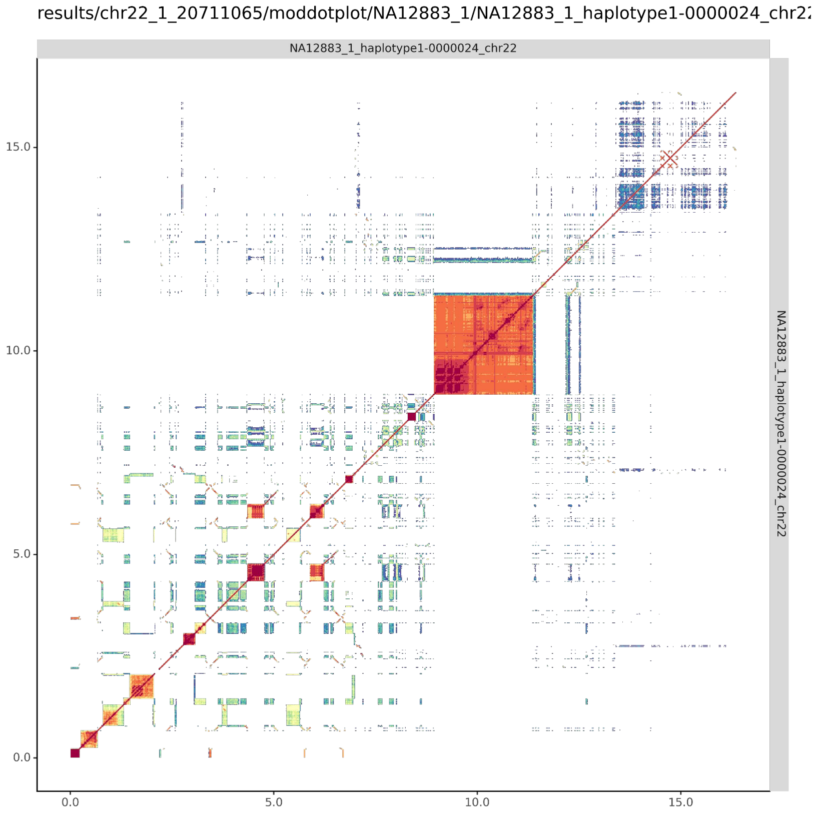


#### NA12883_2_haplotype2-0000081_chr22

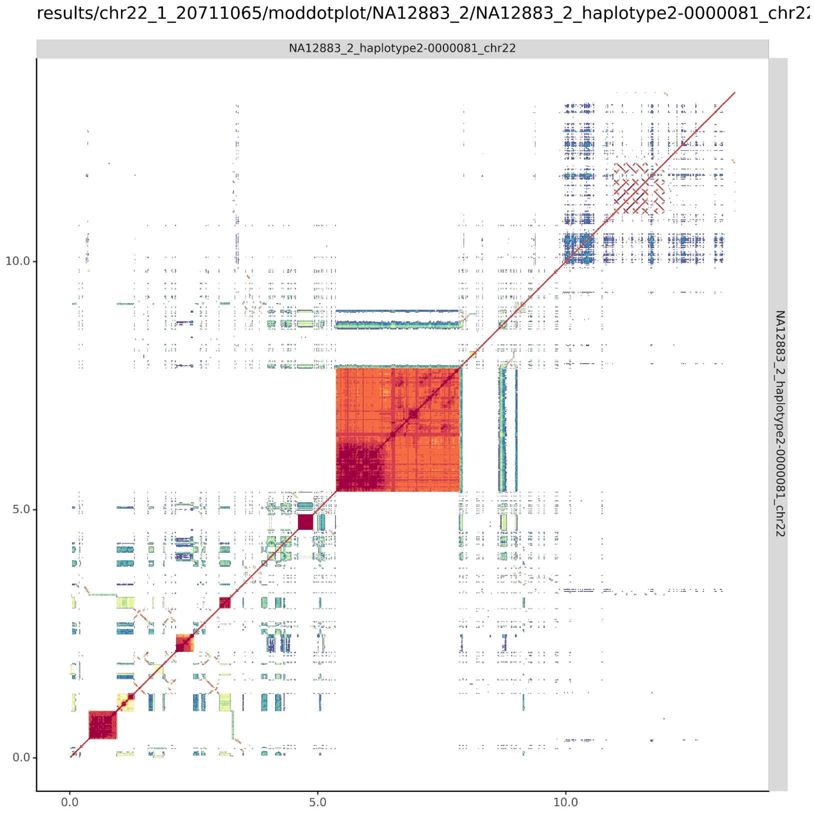

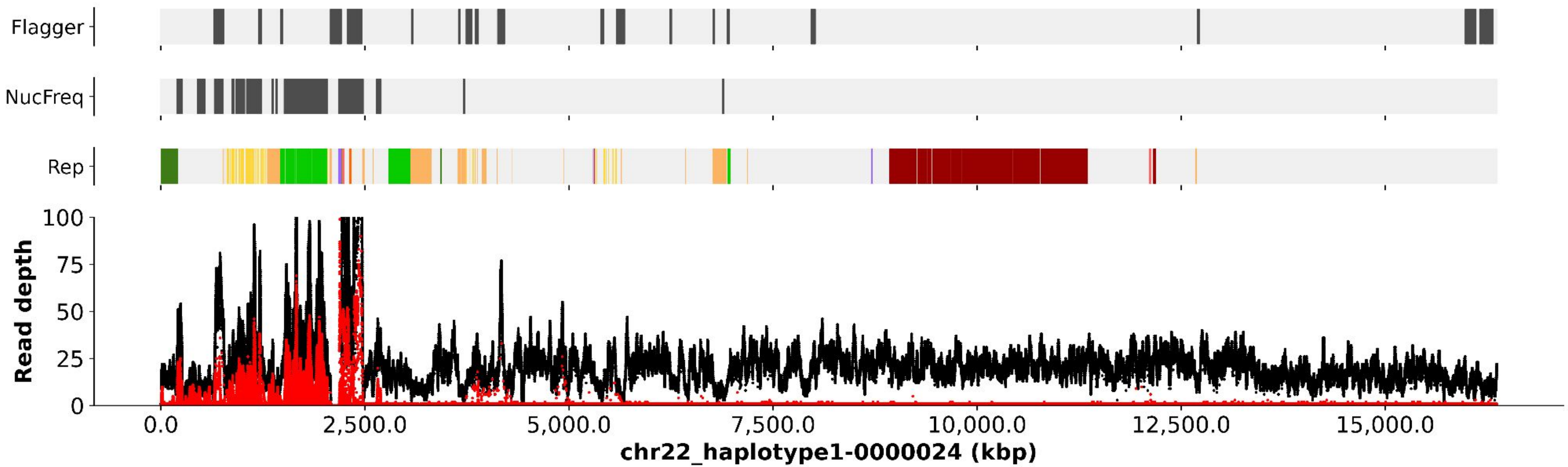
Flagger
NucFreq
Rep
Read depth
0
25
50
75
100
0.0
2,500.0
5,000.0
7,500.0
10,000.0
12,500.0
15,000.0
chr22_haplotype1-0000024 (kbp)

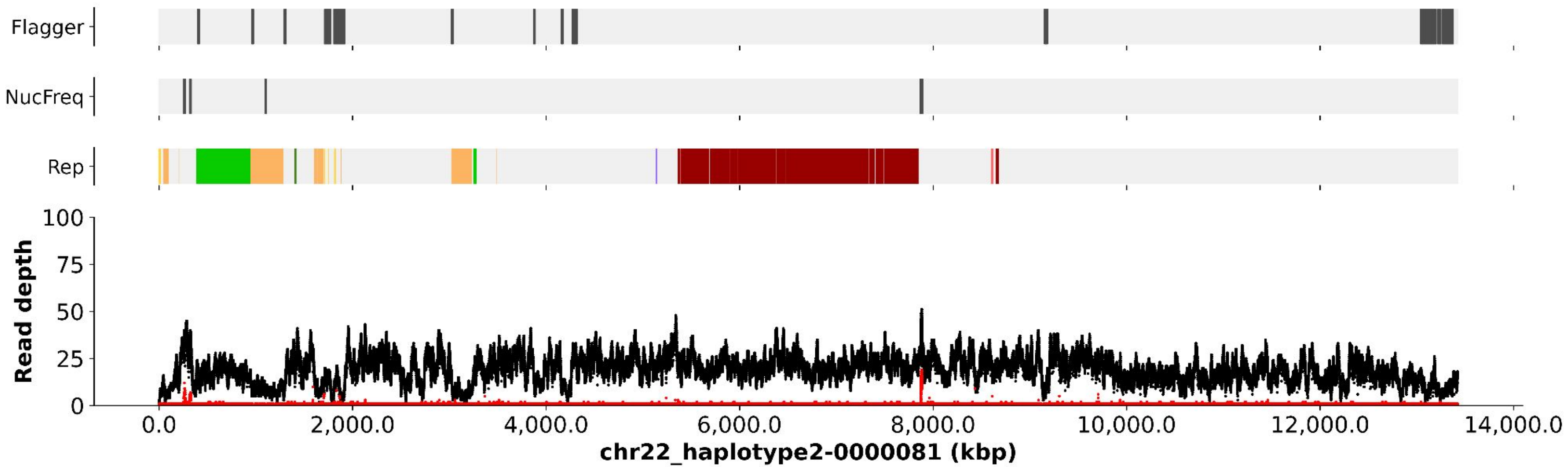
Flagger
NucFreq
Rep
Read depth
0
25
50
75
100
0.0
2,000.0
4,000.0
6,000.0
8,000.0
10,000.0
12,000.0
14,000.0
chr22_haplotype2-0000081 (kbp)

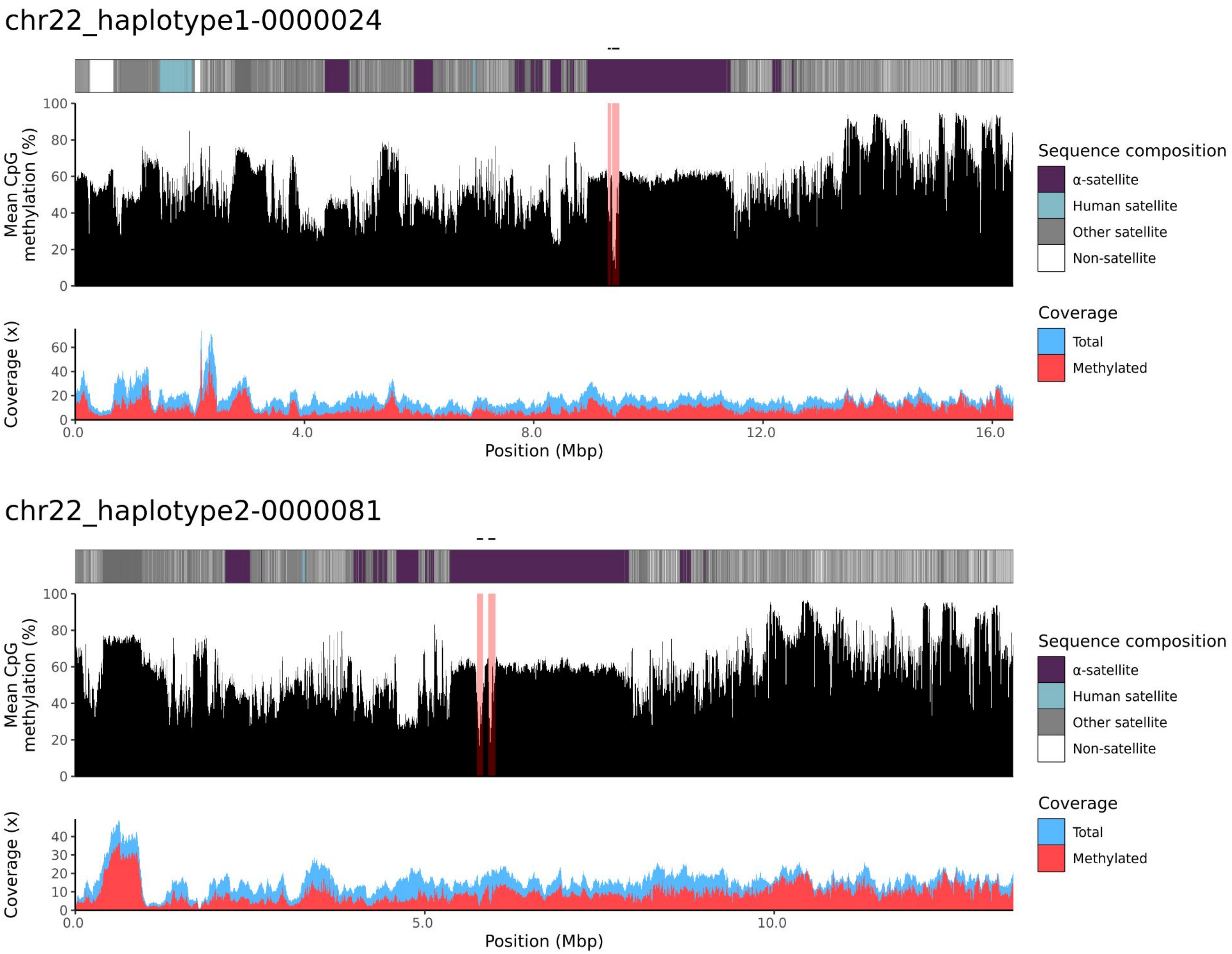
chr22_haplotype1-0000024
Mean CpG methylation (%)
Coverage (x)
Position (Mbp)
Sequence composition
α-satellite
Human satellite
Other satellite
Non-satellite
Coverage
Total
Methylated
chr22_haplotype2-0000081
Mean CpG methylation (%)
Coverage (x)
Position (Mbp)
Sequence composition
α-satellite
Human satellite
Other satellite
Non-satellite
Coverage
Total
Methylated

NA12884

### chr13

#### NA12884_1_haplotype1-0000016_chr13

results/chr13_1_22508596/moddotplot/NA12884_1/NA12884_1_haplotype1-0000016_chr13

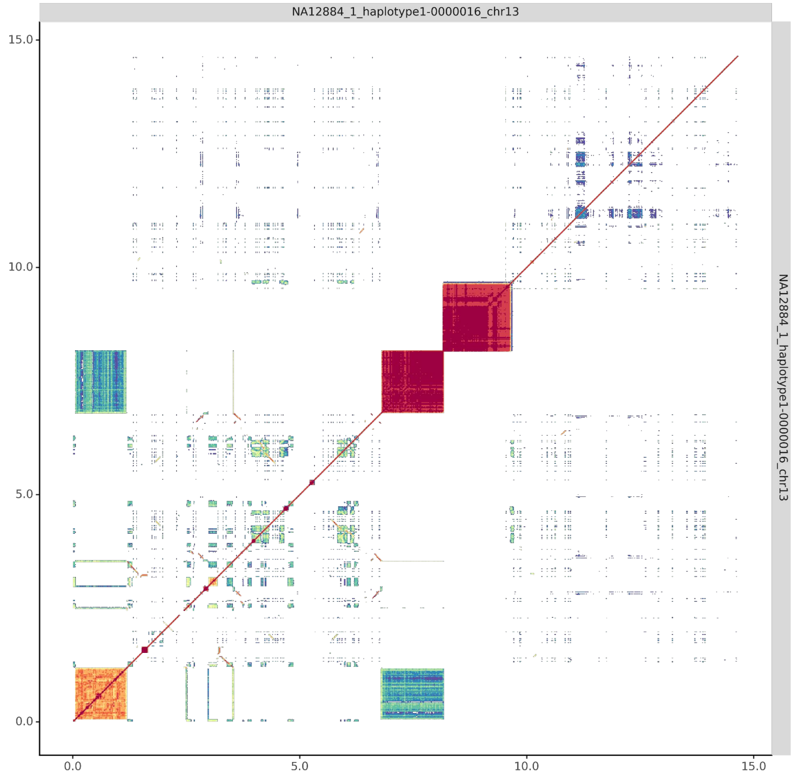


#### NA12884_2_haplotype2-0000068_chr13

results/chr13_1_22508596/moddotplot/NA12884_2/NA12884_2_haplotype2-0000068_chr13

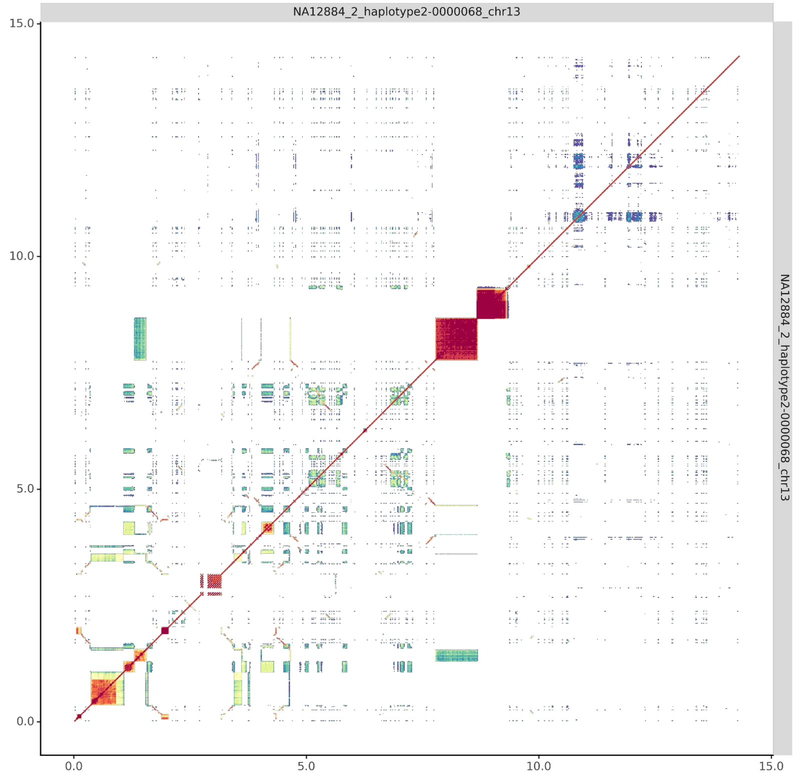

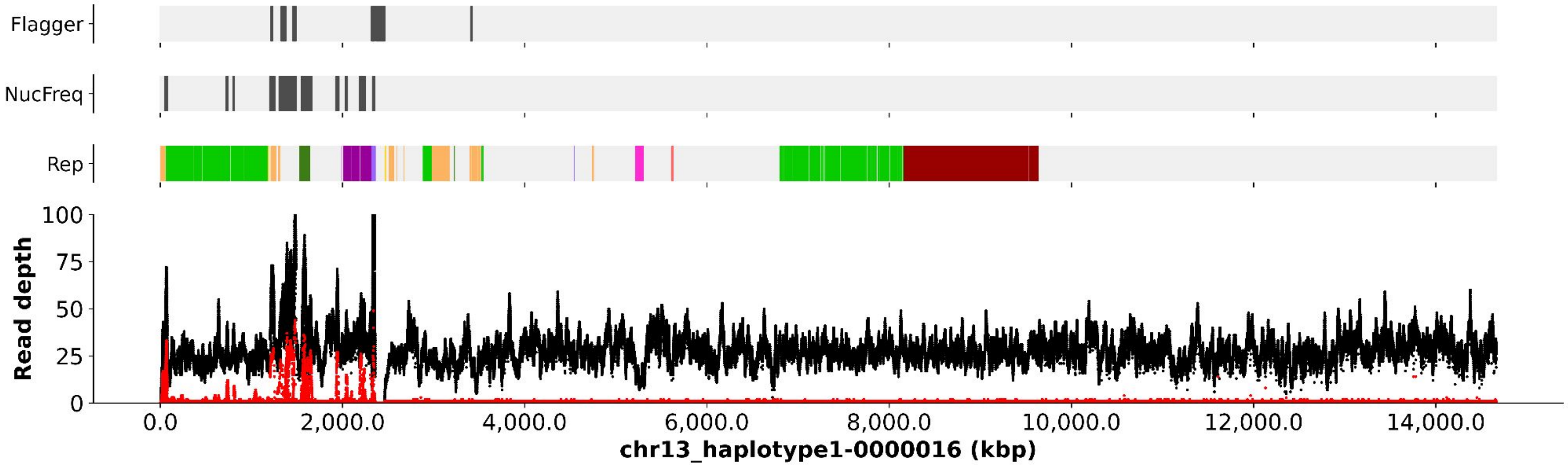

Flagger
NucFreq
Rep
Read depth
0
25
50
75
100
0.0
2,000.0
4,000.0
6,000.0
8,000.0
10,000.0
12,000.0
14,000.0
chr13_haplotype1-0000016 (kbp)


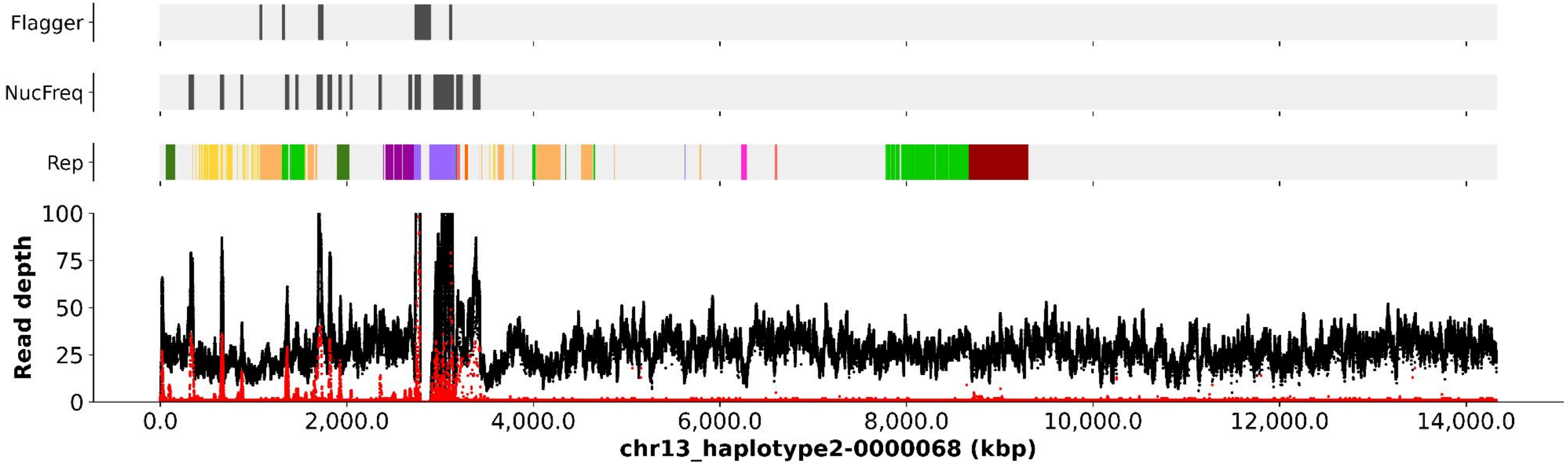

Flagger
NucFreq
Rep
Read depth
0
25
50
75
100
0.0
2,000.0
4,000.0
6,000.0
8,000.0
10,000.0
12,000.0
14,000.0
chr13_haplotype2-0000068 (kbp)

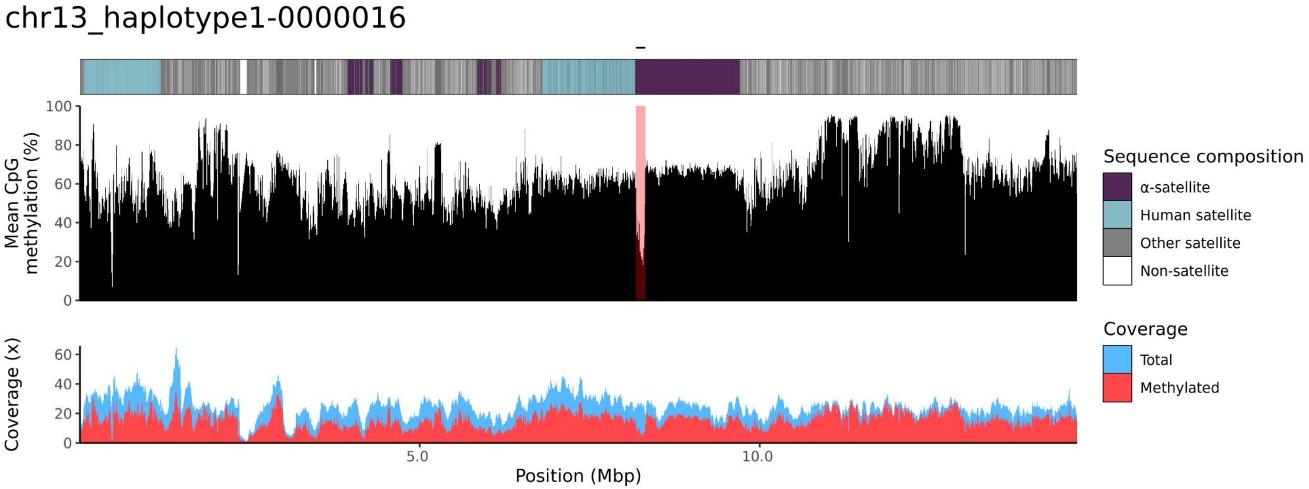
chr13_haplotype1-0000016
Mean CpG methylation (%)
0
20
40
60
80
100
Coverage (x)
0
20
40
60
5.0
10.0
Position (Mbp)
Sequence composition
α-satellite
Human satellite
Other satellite
Non-satellite
Coverage
Total
Methylated

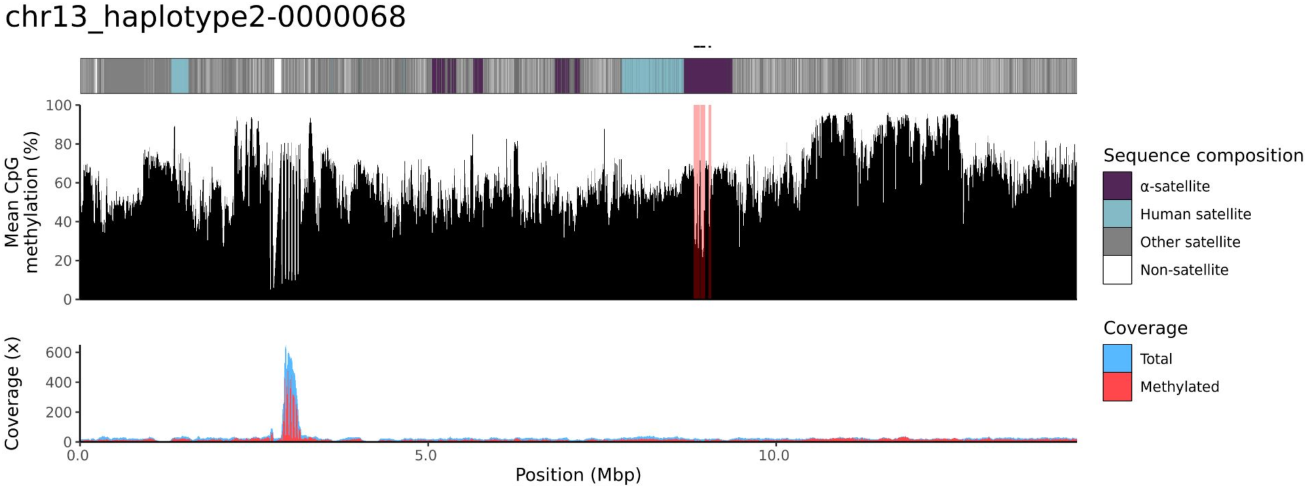
chr13_haplotype2-0000068
Mean CpG methylation (%)
0
20
40
60
80
100
Coverage (x)
0
200
400
600
0.0
5.0
10.0
Position (Mbp)
Sequence composition
α-satellite
Human satellite
Other satellite
Non-satellite
Coverage
Total
Methylated

### chr14

#### NA12884_1_haplotype1-0000003_chr14

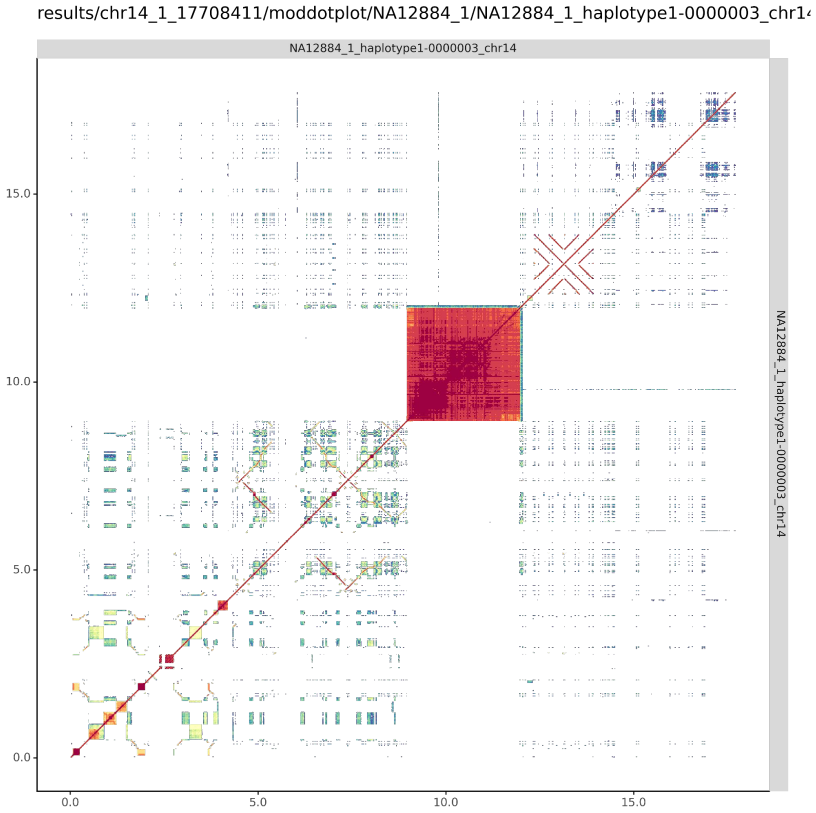


#### NA12884_2_haplotype2-0000058_chr14

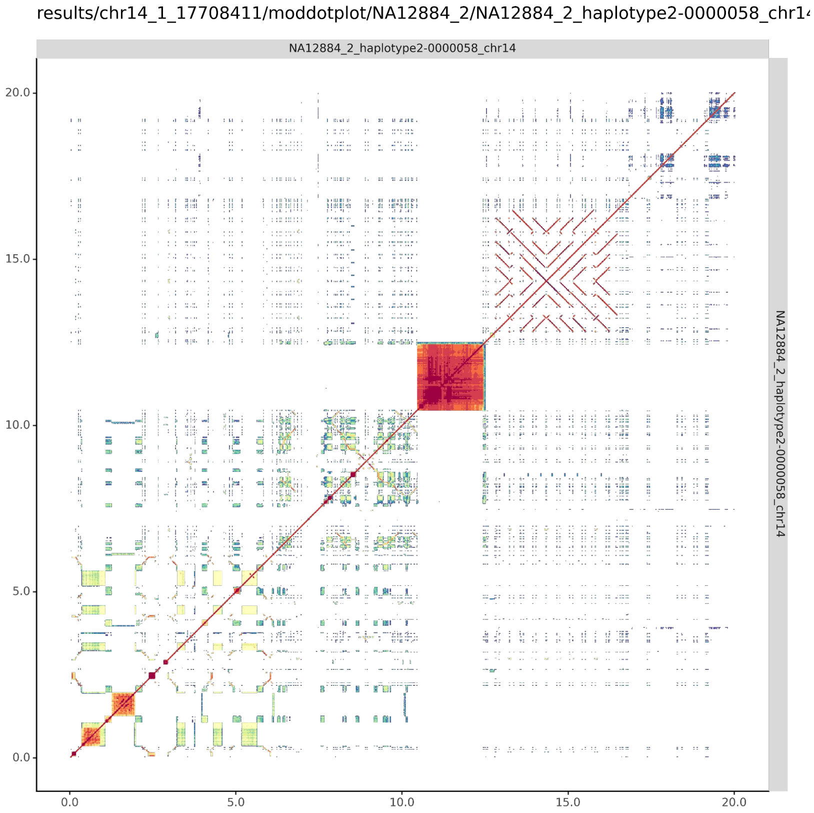

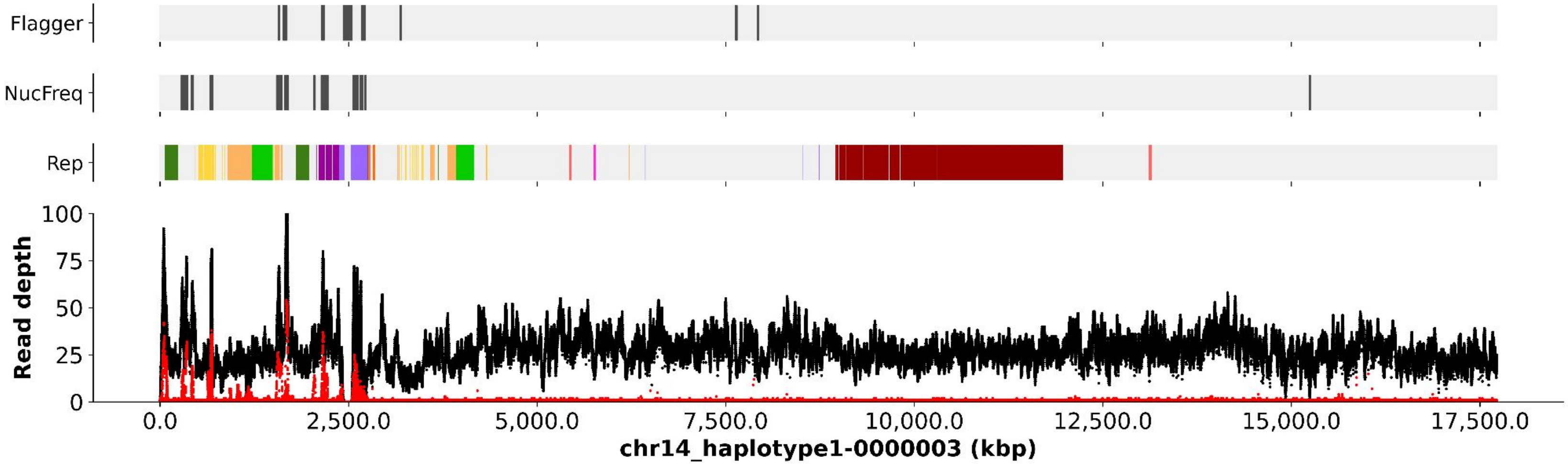
Flagger
NucFreq
Rep
Read depth
100
75
50
25
0
0.0
2,500.0
5,000.0
7,500.0
10,000.0
12,500.0
15,000.0
17,500.0
chr14_haplotype1-0000003 (kbp)

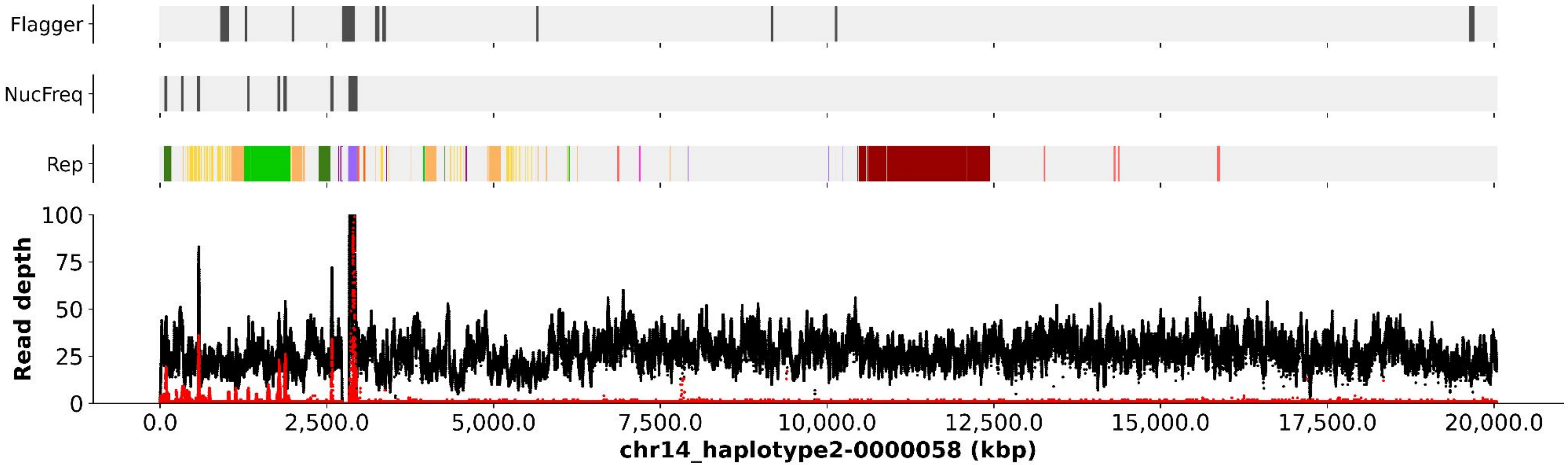
Flagger
NucFreq
Rep
Read depth
100
75
50
25
0
0.0
2,500.0
5,000.0
7,500.0
10,000.0
12,500.0
15,000.0
17,500.0
20,000.0
chr14_haplotype2-0000058 (kbp)

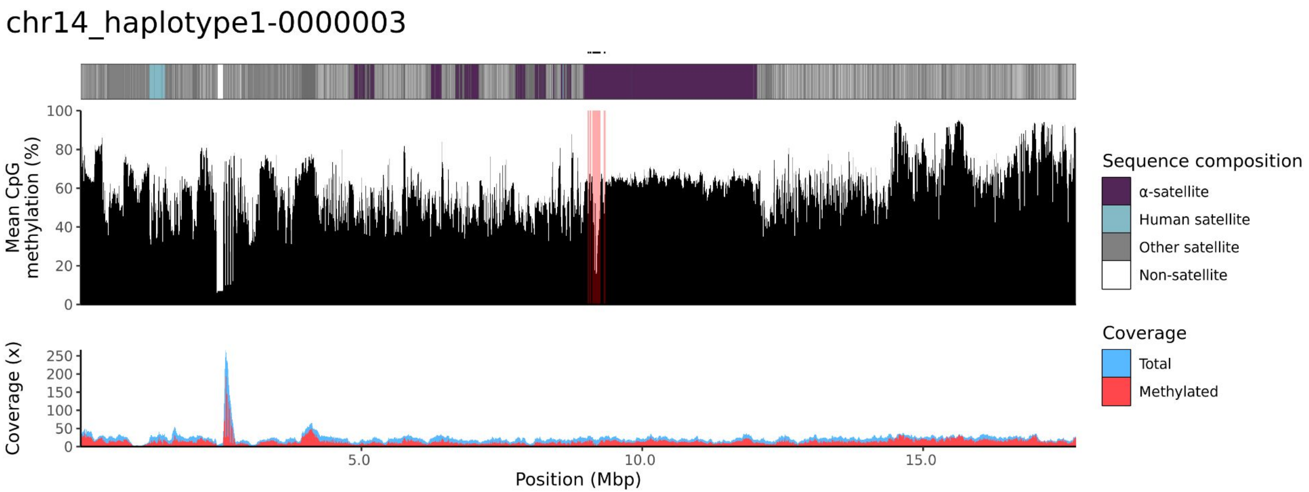
chr14_haplotype1-0000003
Mean CpG methylation (%)
100
80
60
40
20
0
Coverage (x)
250
200
150
100
50
0
5.0
10.0
15.0
Position (Mbp)
Sequence composition
α-satellite
Human satellite
Other satellite
Non-satellite
Coverage
Total
Methylated

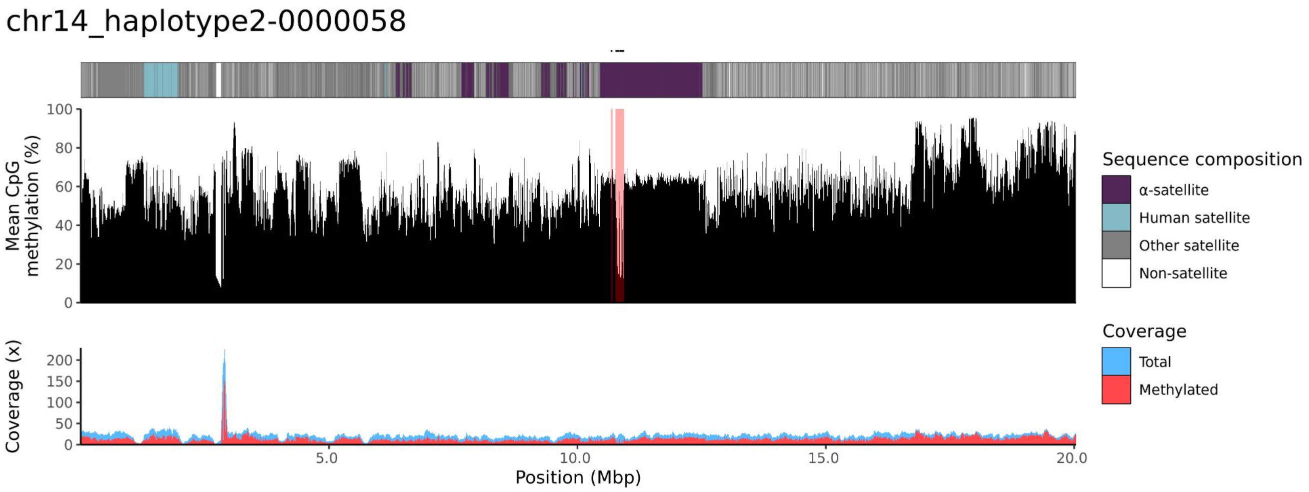
chr14_haplotype2-0000058
Mean CpG methylation (%)
100
80
60
40
20
0
Coverage (x)
200
150
100
50
0
5.0
10.0
15.0
20.0
Position (Mbp)
Sequence composition
α-satellite
Human satellite
Other satellite
Non-satellite
Coverage
Total
Methylated

### chr15

#### NA12884_1_haplotype1-0000015_chr15

results/chr15_1_22694466/moddotplot/NA12884_1/NA12884_1_haplotype1-0000015_chr1!

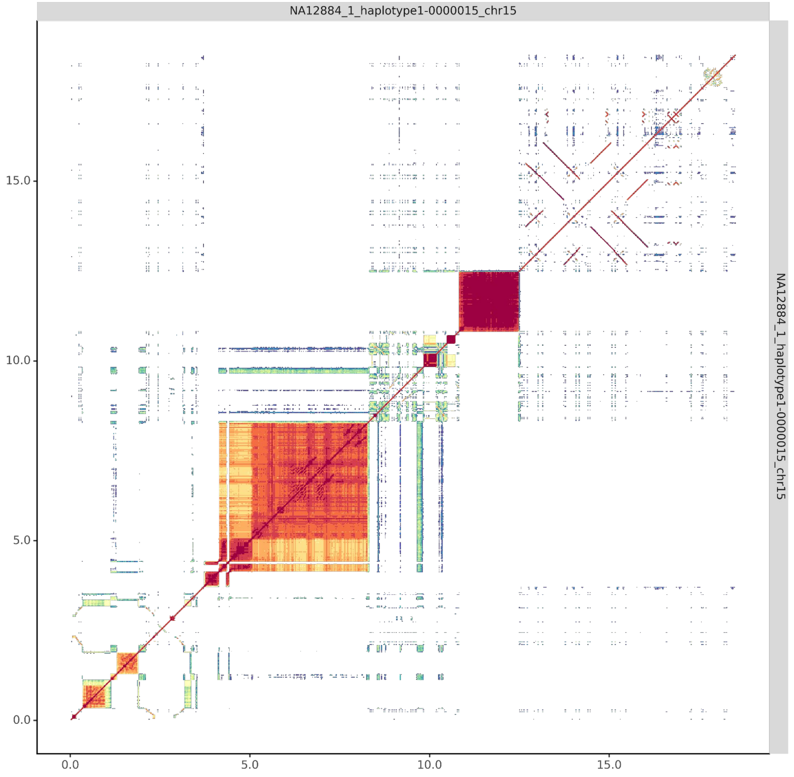


#### NA12884_2_haplotype2-0000067_chr15

results/chr15_1_22694466/moddotplot/NA12884_2/NA12884_2_haplotype2-0000067_chr1!

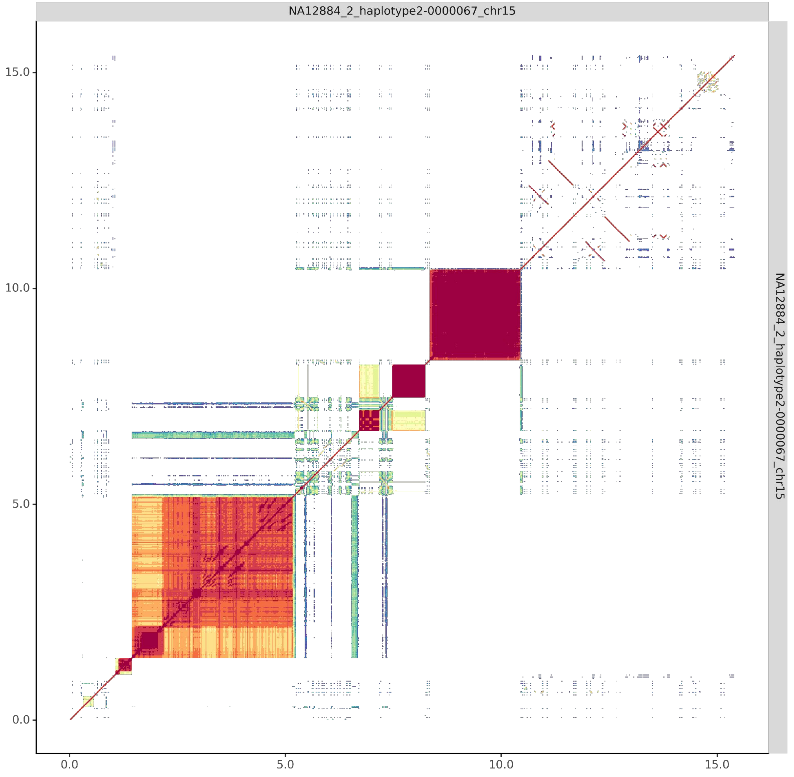

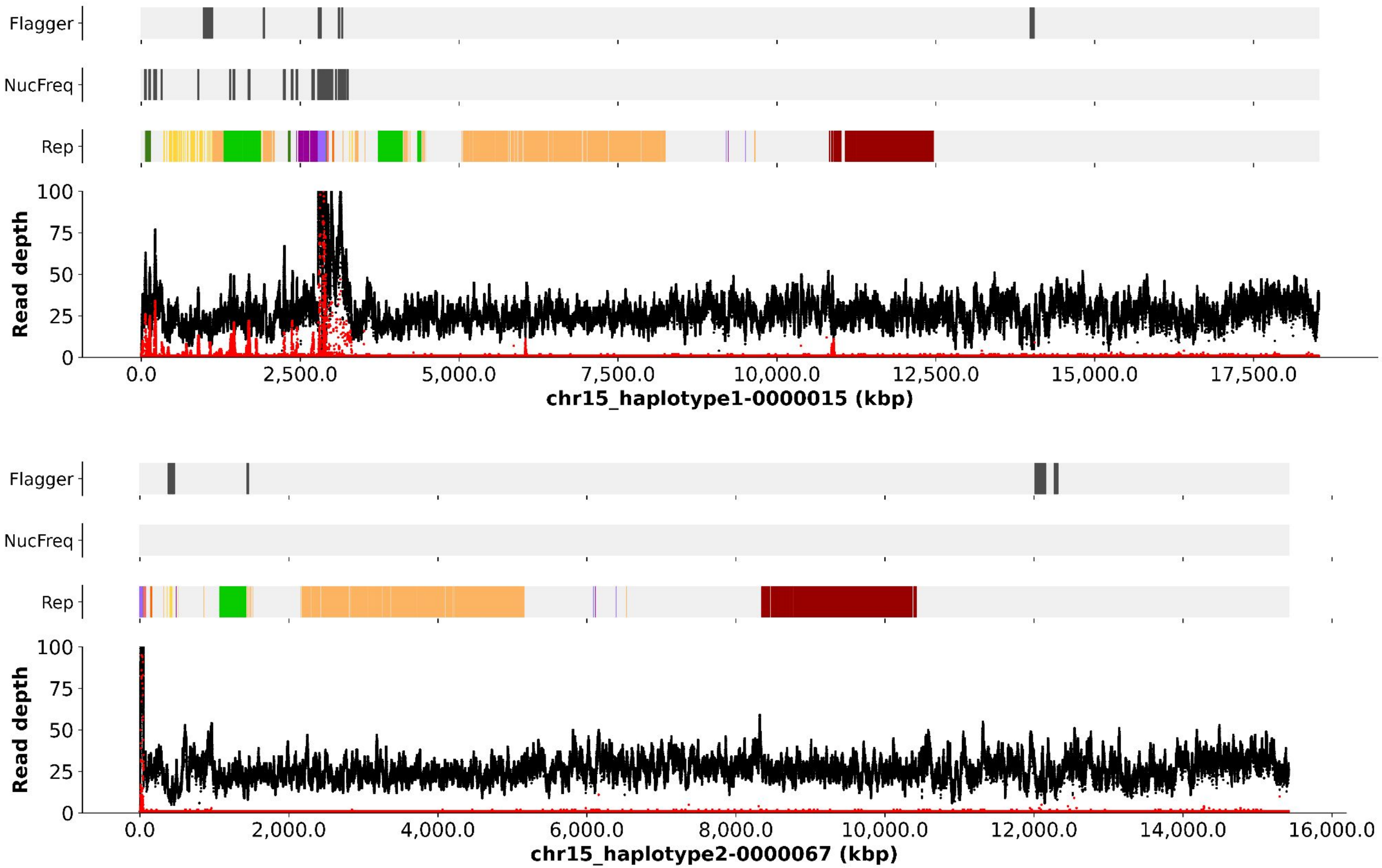

Flagger
NucFreq
Rep
Read depth
chr15_haplotype1-0000015 (kbp)
Flagger
NucFreq
Rep
Read depth
chr15_haplotype2-0000067 (kbp)

chr15_haplotype1-0000015

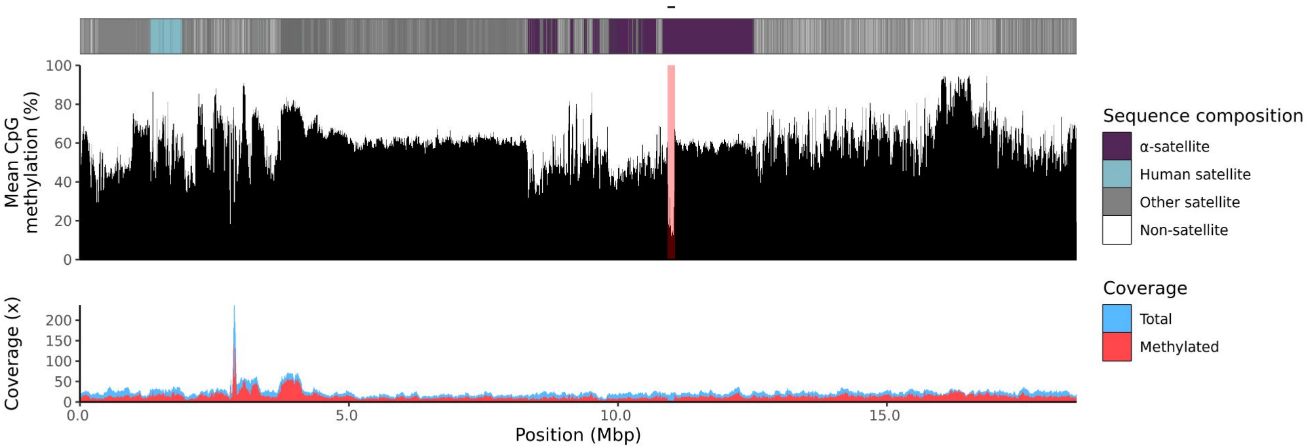


chr15_haplotype2-0000067

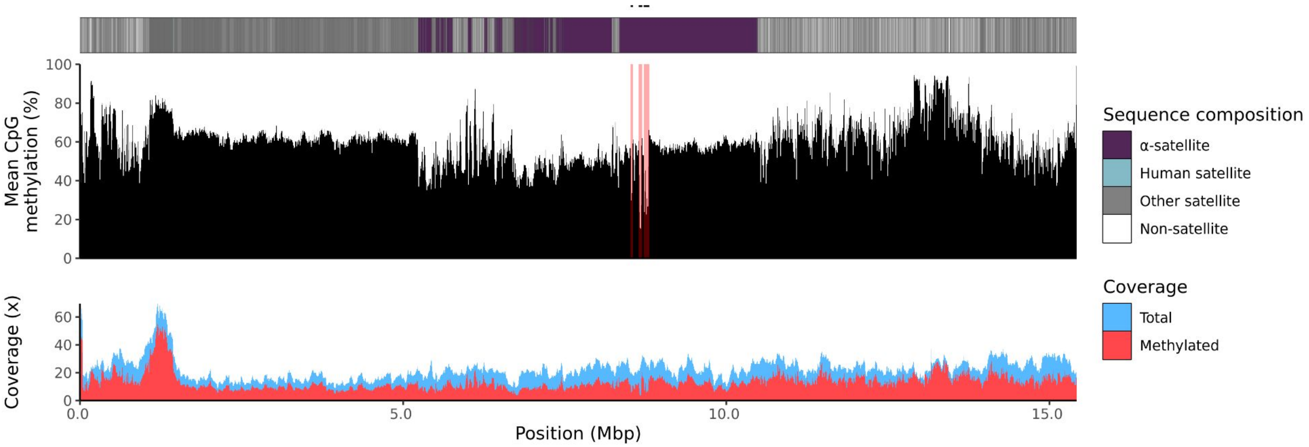

### chr21

#### NA12884_1_haplotype1-0000010_chr21

results/chr21_1_16306378/moddotplot/NA12884_1/NA12884_1_haplotype1-0000010_chr2[illegible]

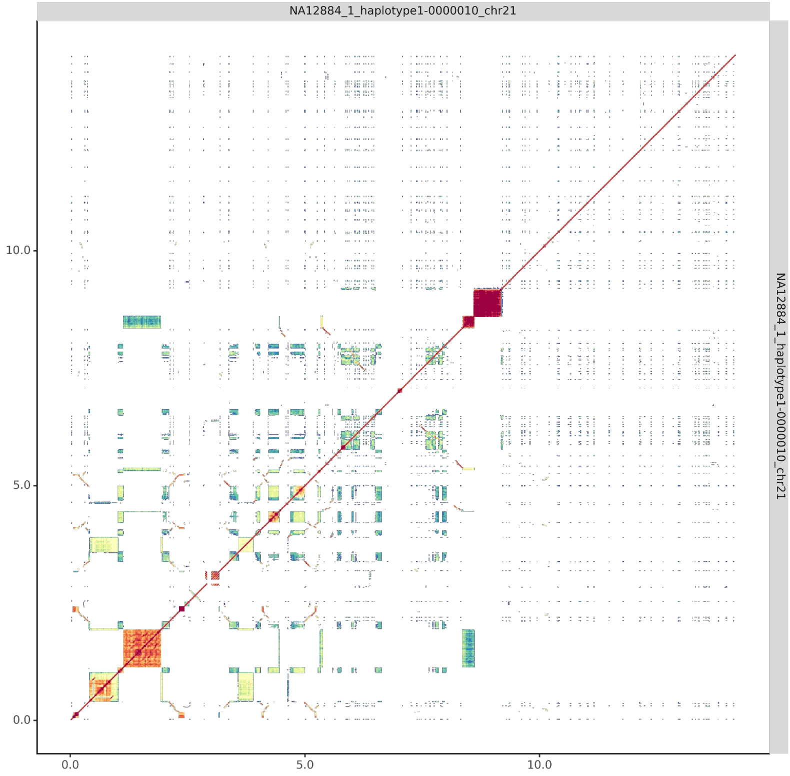


#### NA12884_2_haplotype2-0000075_chr21

results/chr21_1_16306378/moddotplot/NA12884_2/NA12884_2_haplotype2-0000075_chr2[illegible]

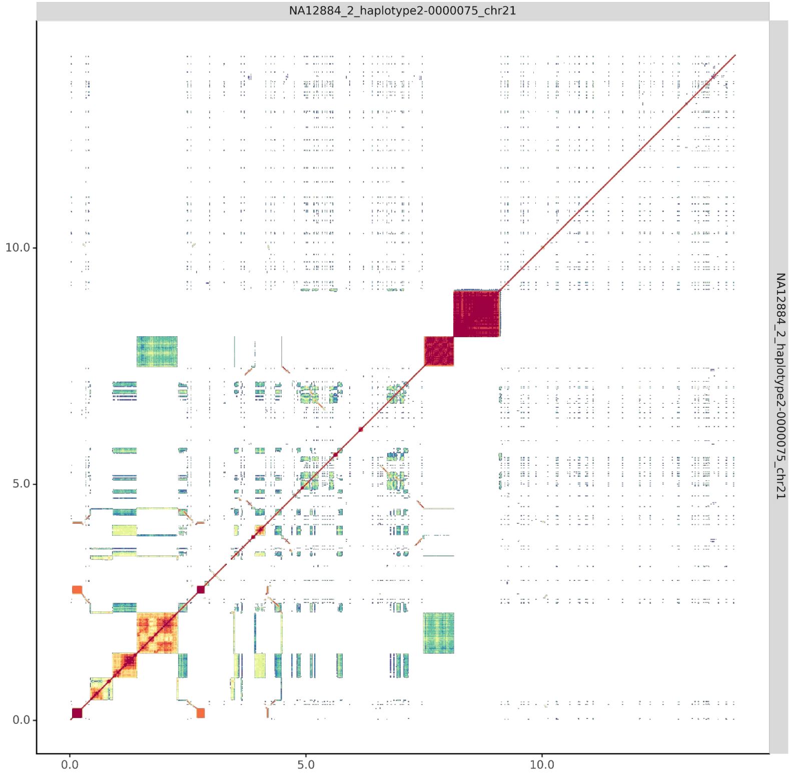

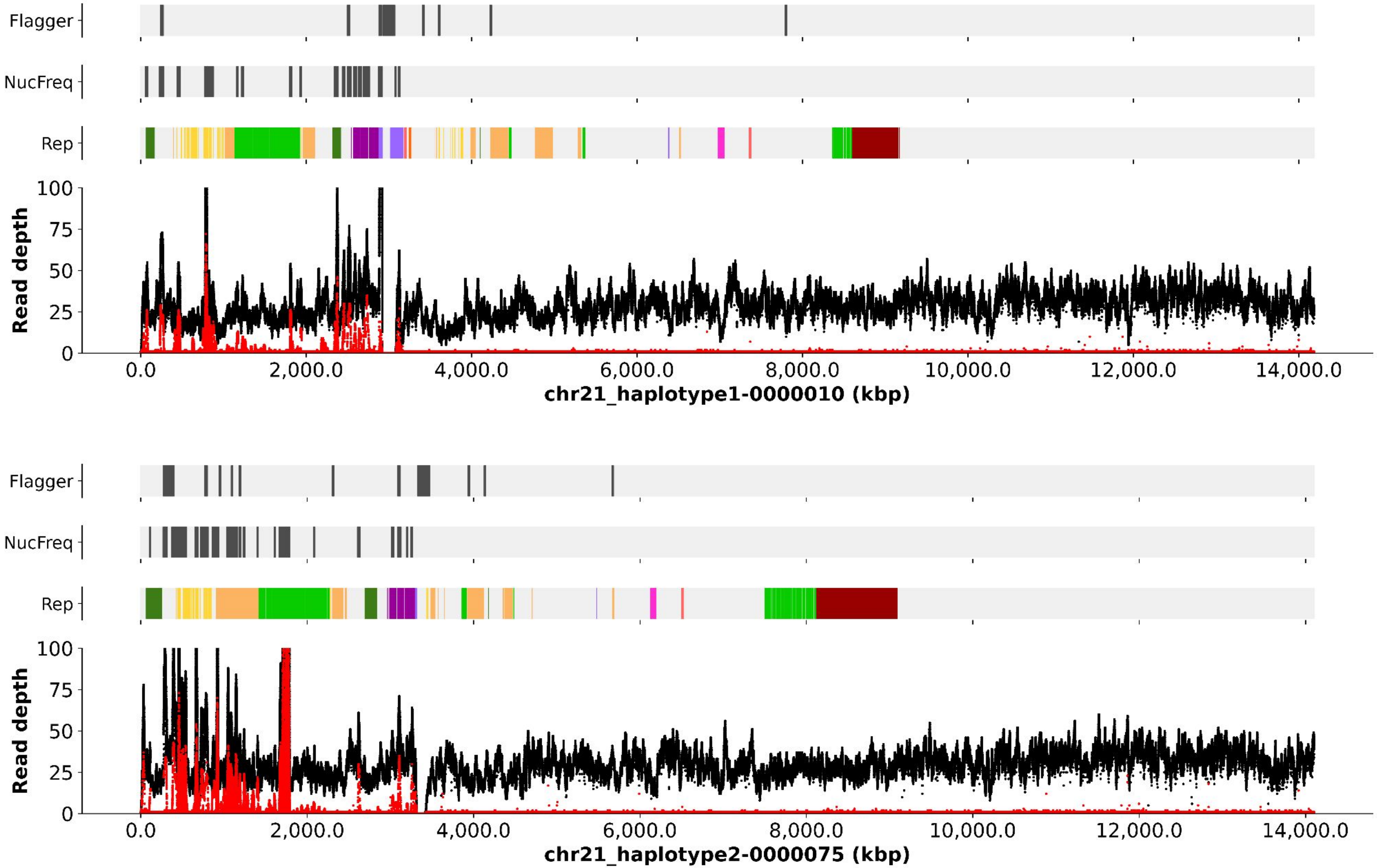

Flagger
NucFreq
Rep
Read depth
100
75
50
25
0
0.0
2,000.0
4,000.0
6,000.0
8,000.0
10,000.0
12,000.0
14,000.0
chr21_haplotype1-0000010 (kbp)
Flagger
NucFreq
Rep
Read depth
100
75
50
25
0
0.0
2,000.0
4,000.0
6,000.0
8,000.0
10,000.0
12,000.0
14,000.0
chr21_haplotype2-0000075 (kbp)

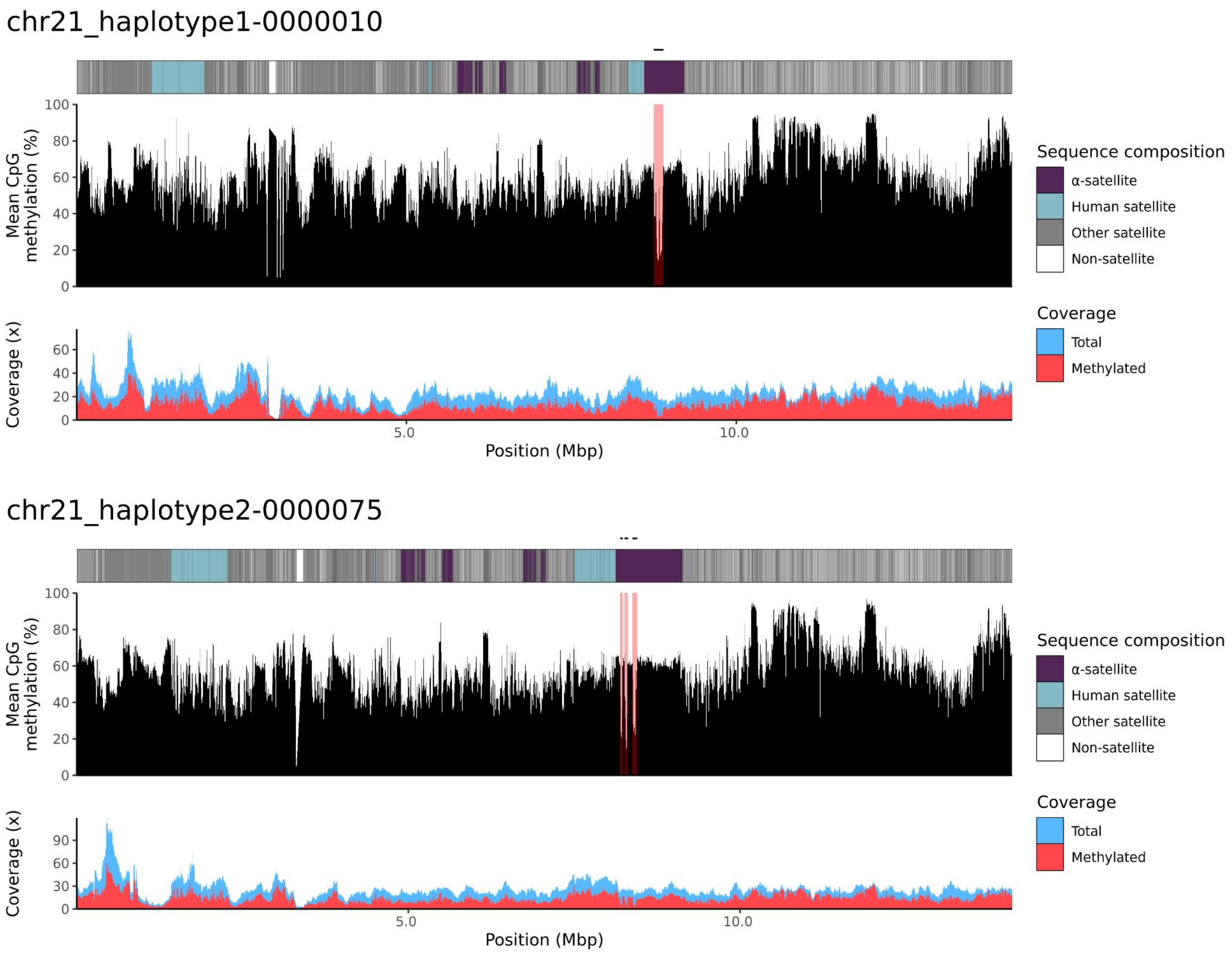

chr21_haplotype1-0000010
Mean CpG methylation (%)
100
80
60
40
20
0
Coverage (x)
60
40
20
0
5.0
10.0
Position (Mbp)
Sequence composition
α-satellite
Human satellite
Other satellite
Non-satellite
Coverage
Total
Methylated
chr21_haplotype2-0000075
Mean CpG methylation (%)
100
80
60
40
20
0
Coverage (x)
90
60
30
0
5.0
10.0
Position (Mbp)
Sequence composition
α-satellite
Human satellite
Other satellite
Non-satellite
Coverage
Total
Methylated

### chr22

#### NA12884_1_haplotype1-0000009_chr22

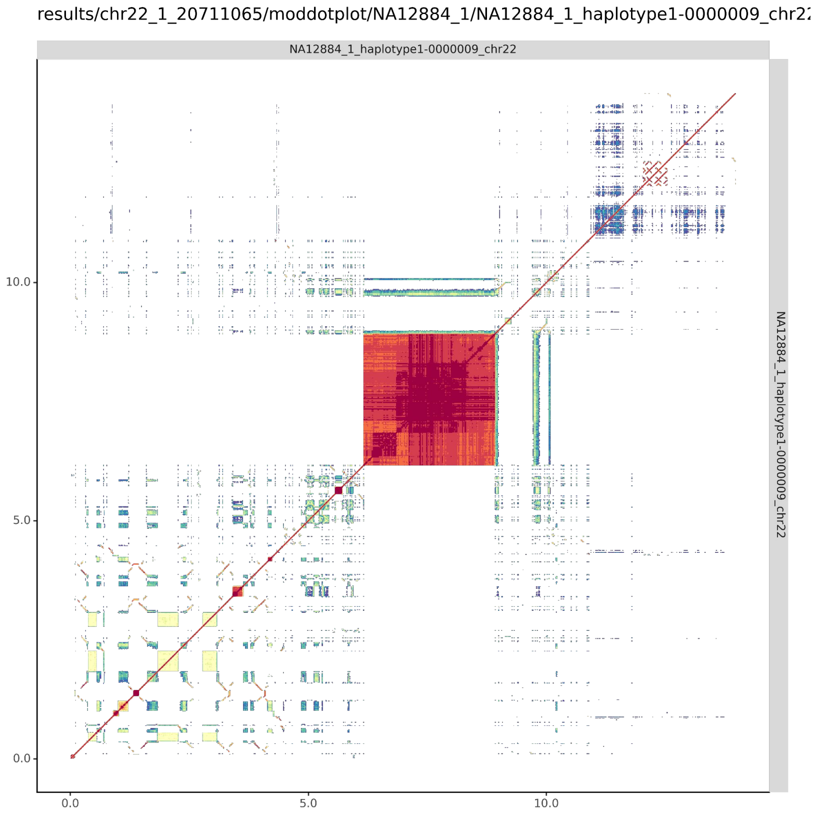


#### NA12884_2_haplotype2-0000071_chr22

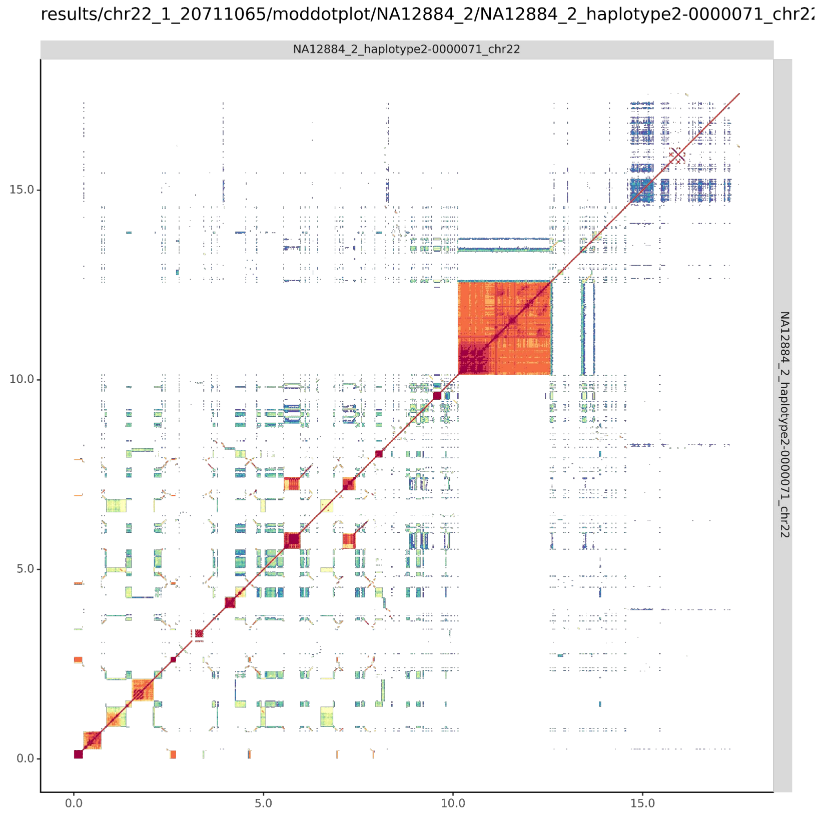

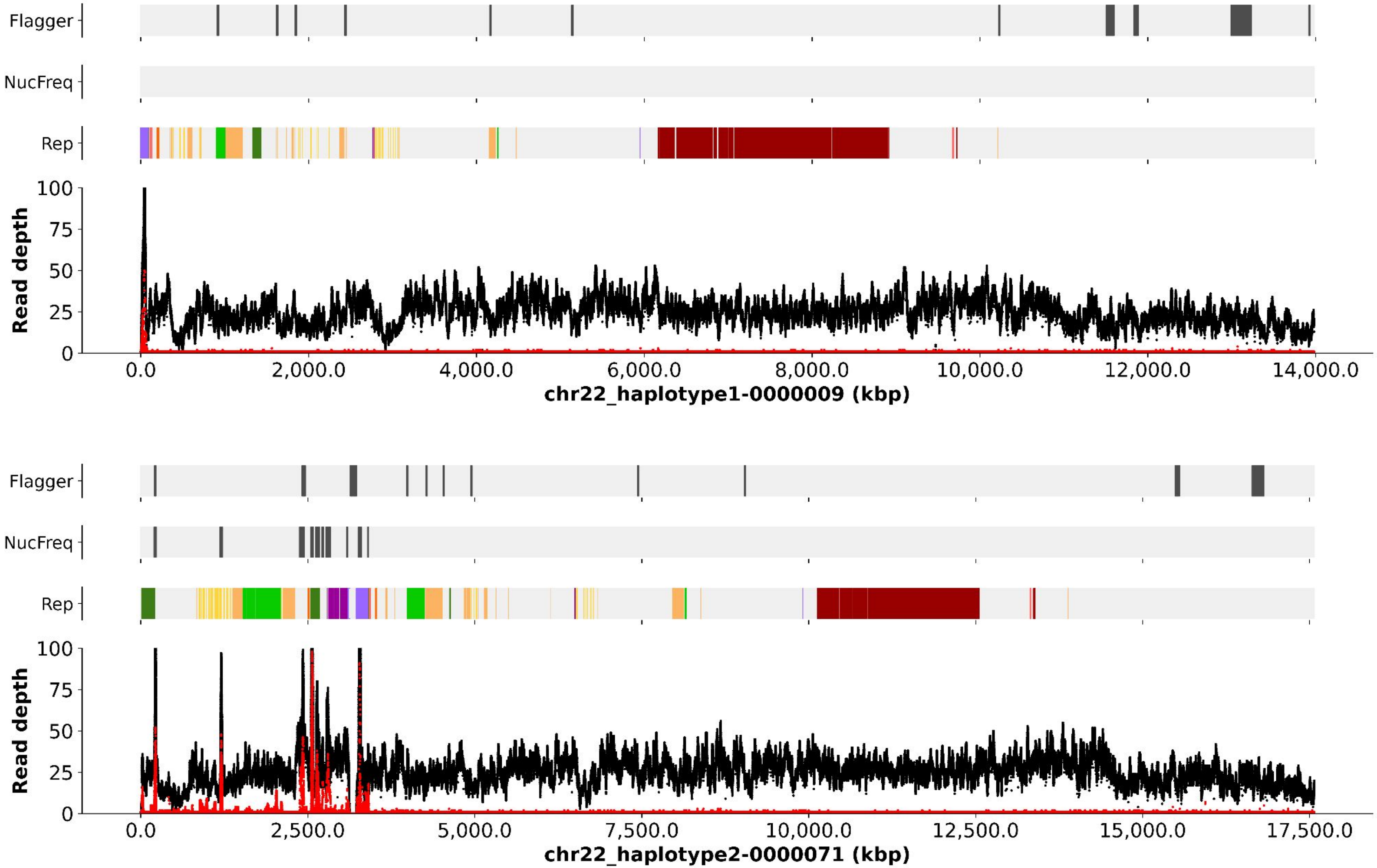
Flagger
NucFreq
Rep
Read depth
100
75
50
25
0
0.0
2,000.0
4,000.0
6,000.0
8,000.0
10,000.0
12,000.0
14,000.0
chr22_haplotype1-0000009 (kbp)
Flagger
NucFreq
Rep
Read depth
100
75
50
25
0
0.0
2,500.0
5,000.0
7,500.0
10,000.0
12,500.0
15,000.0
17,500.0
chr22_haplotype2-0000071 (kbp)

chr22_haplotype1-0000009

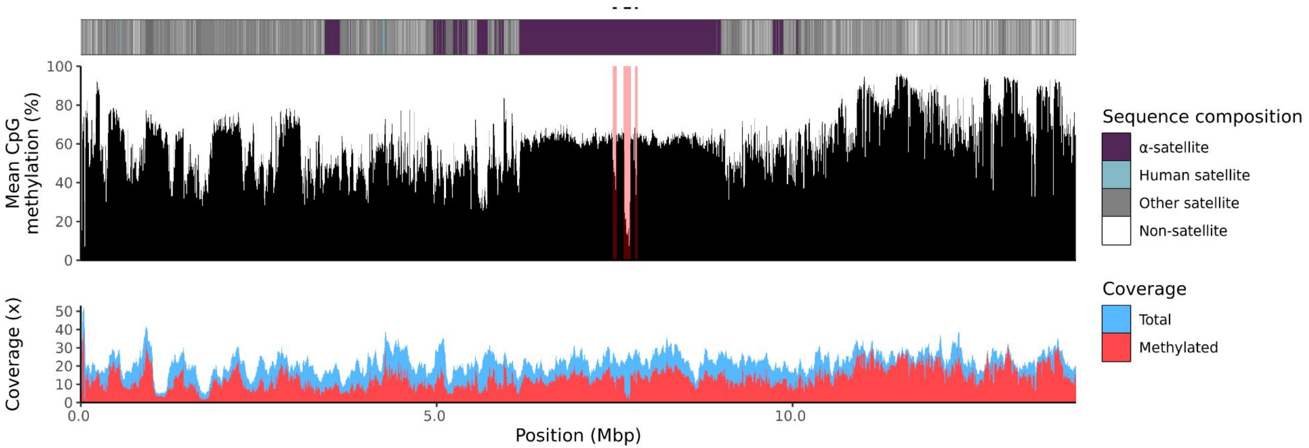


chr22_haplotype2-0000071

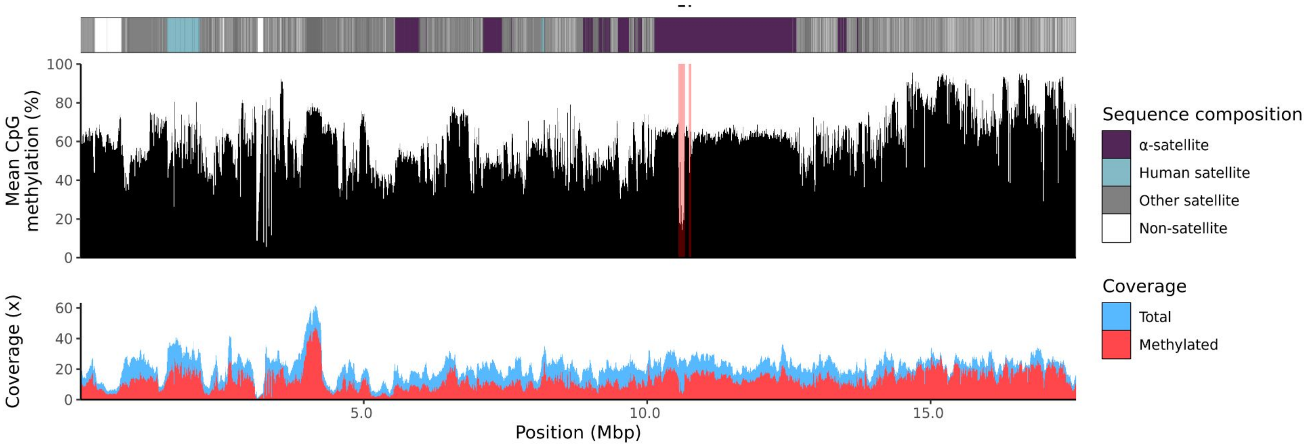

NA12885

### chr13

#### NA12885_1_haplotype1-0000011_chr13

results/chr13_1_22508596/moddotplot/NA12885_1/NA12885_1_haplotype1-0000011_chr1

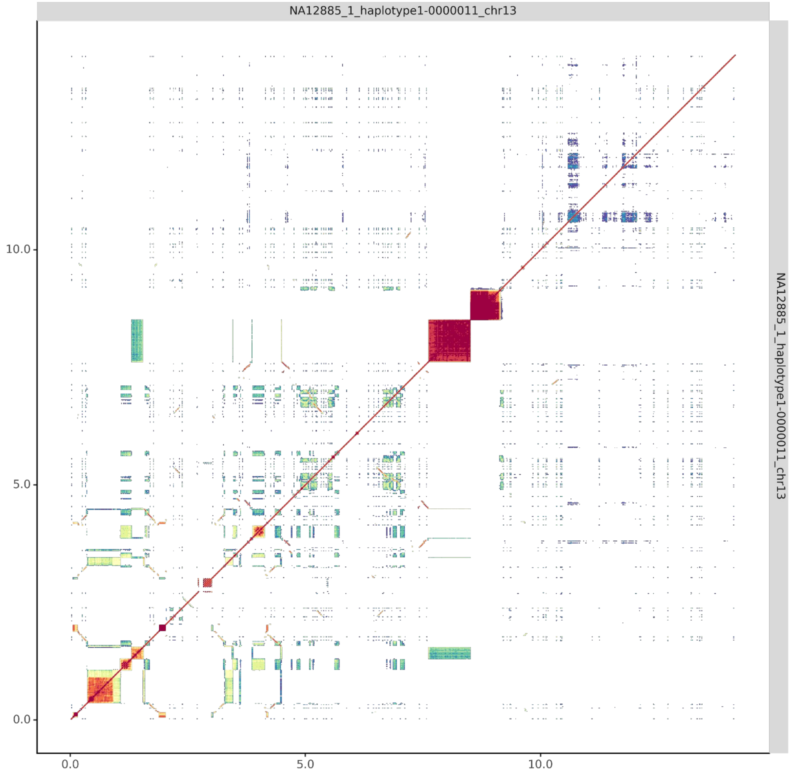


#### NA12885_2_haplotype2-0000069_chr13

results/chr13_1_22508596/moddotplot/NA12885_2/NA12885_2_haplotype2-0000069_chr1

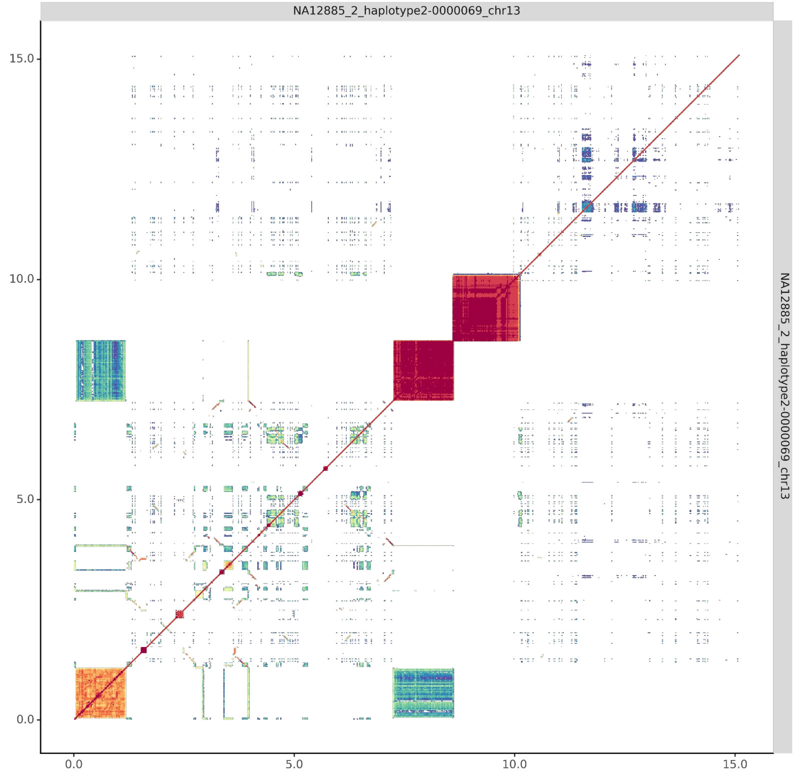

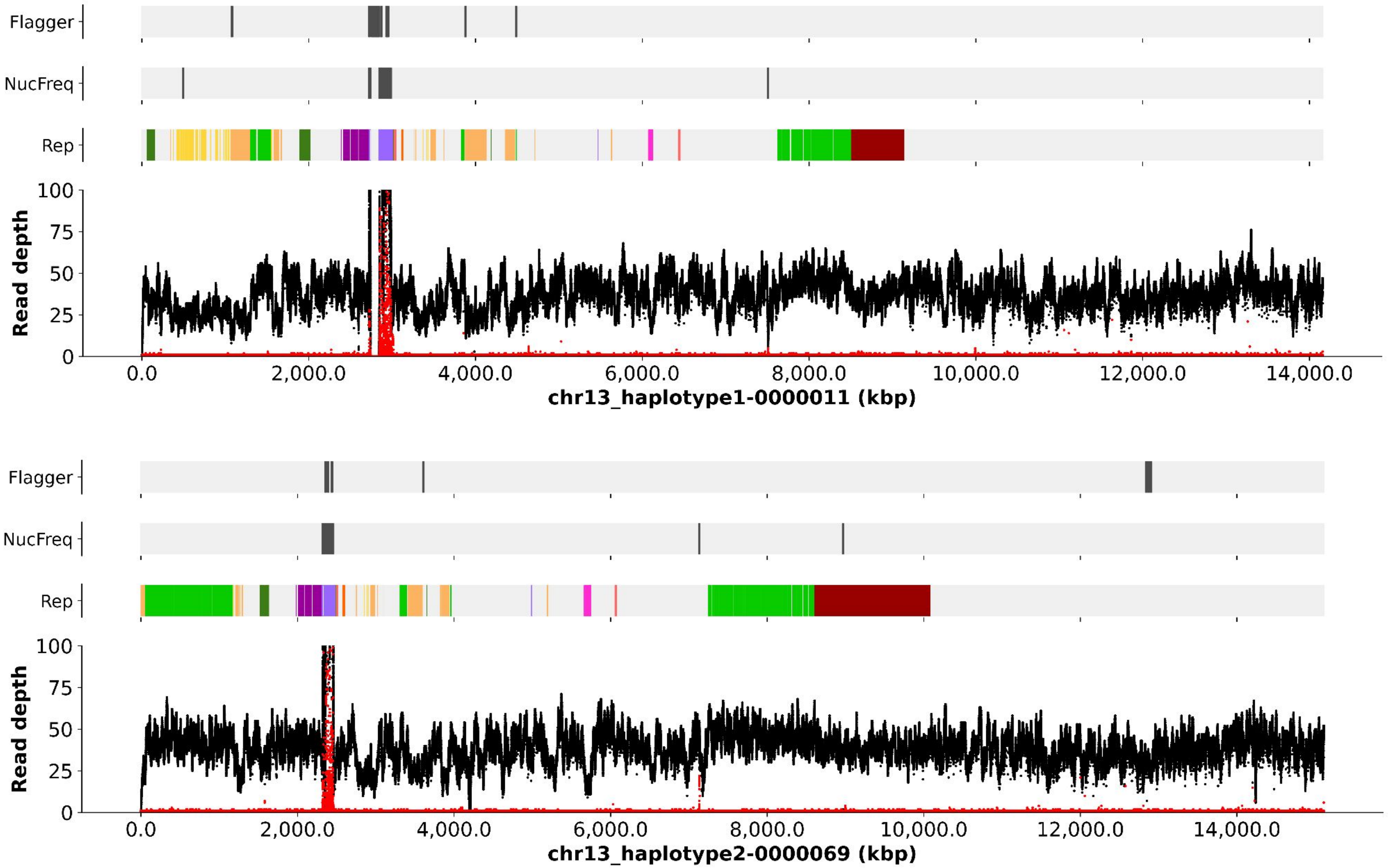
Flagger
NucFreq
Rep
Read depth
100
75
50
25
0
0.0
2,000.0
4,000.0
6,000.0
8,000.0
10,000.0
12,000.0
14,000.0
chr13_haplotype1-0000011 (kbp)
Flagger
NucFreq
Rep
Read depth
100
75
50
25
0
0.0
2,000.0
4,000.0
6,000.0
8,000.0
10,000.0
12,000.0
14,000.0
chr13_haplotype2-0000069 (kbp)

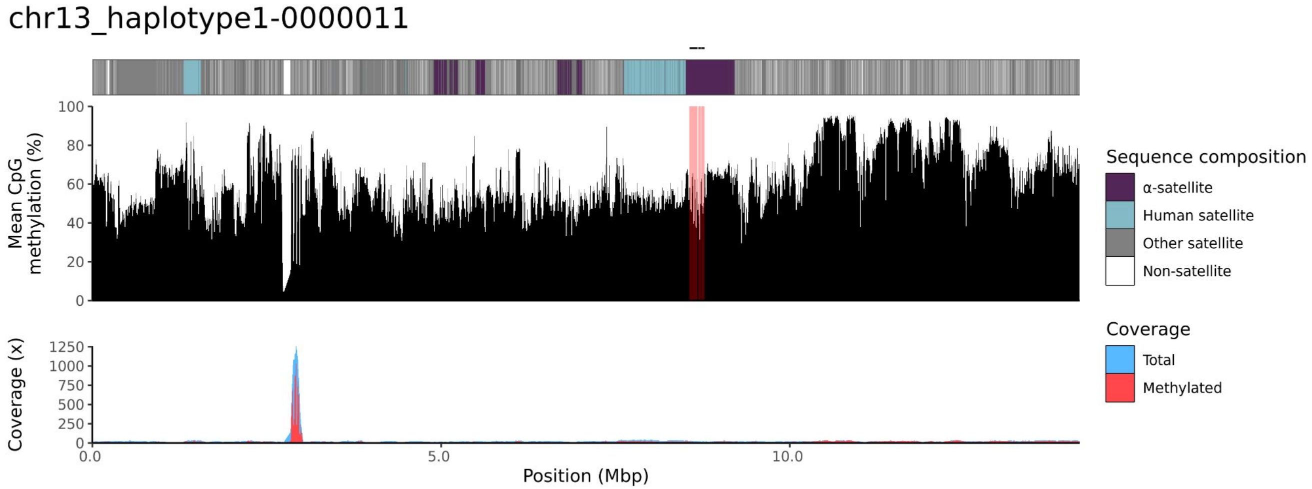
chr13_haplotype1-0000011
Mean CpG methylation (%)
100
80
60
40
20
0
Coverage (x)
1250
1000
750
500
250
0
0.0
5.0
10.0
Position (Mbp)
Sequence composition
α-satellite
Human satellite
Other satellite
Non-satellite
Coverage
Total
Methylated

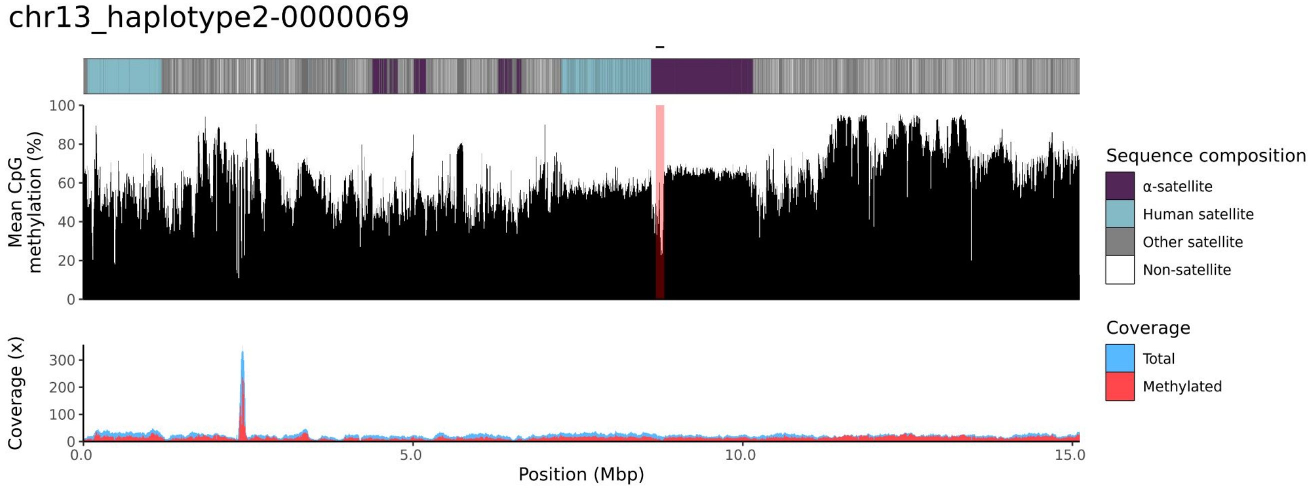
chr13_haplotype2-0000069
Mean CpG methylation (%)
100
80
60
40
20
0
Coverage (x)
300
200
100
0
0.0
5.0
10.0
15.0
Position (Mbp)
Sequence composition
α-satellite
Human satellite
Other satellite
Non-satellite
Coverage
Total
Methylated

### chr14

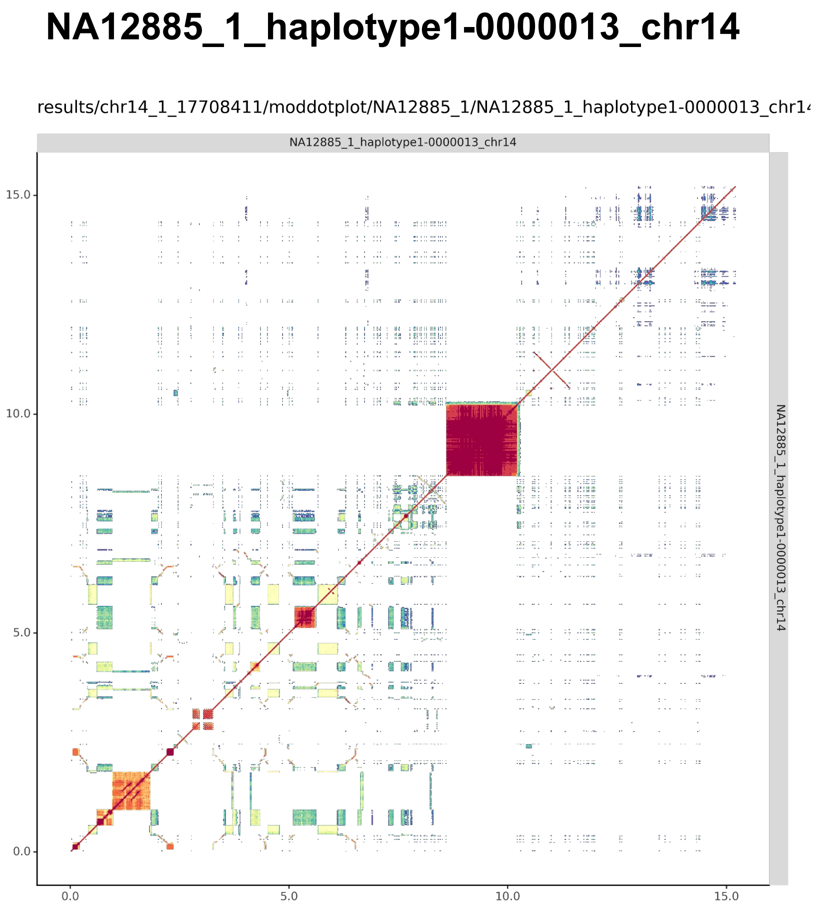
NA12885_1_haplotype1-0000013_chr14
results/chr14_1_17708411/moddotplot/NA12885_1/NA12885_1_haplotype1-0000013_chr1
NA12885_1_haplotype1-0000013_chr14
15.0
10.0
5.0
0.0
0.0
5.0
10.0
15.0
NA12885_1_haplotype1-0000013_chr14

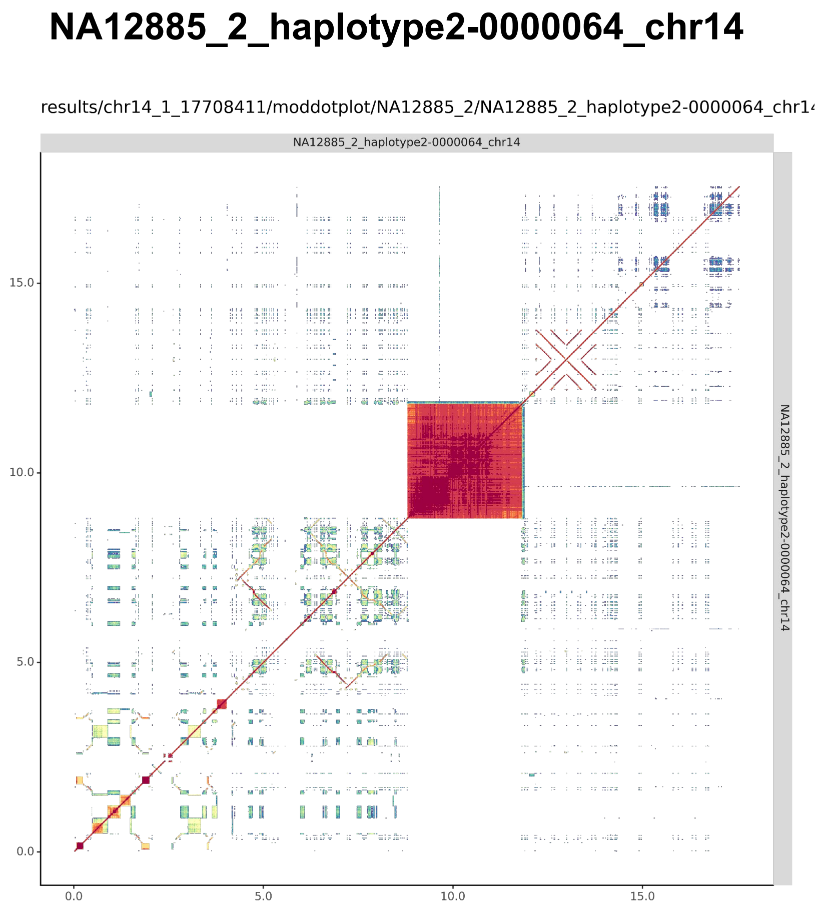
NA12885_2_haplotype2-0000064_chr14
results/chr14_1_17708411/moddotplot/NA12885_2/NA12885_2_haplotype2-0000064_chr1
NA12885_2_haplotype2-0000064_chr14
15.0
10.0
5.0
0.0
0.0
5.0
10.0
15.0
NA12885_2_haplotype2-0000064_chr14

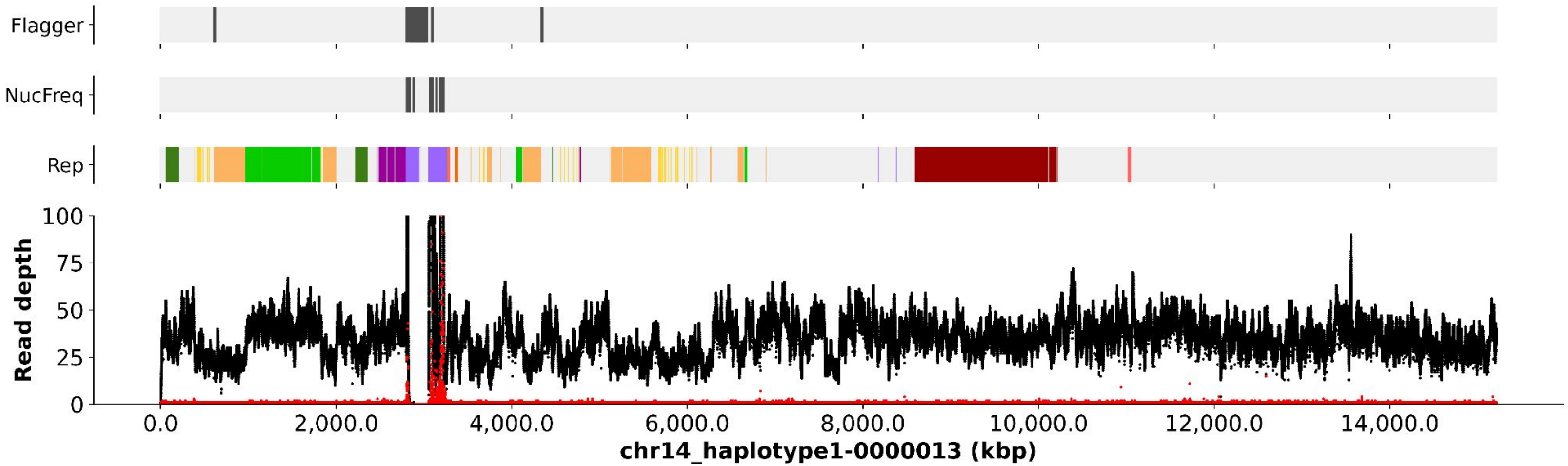
Flagger
NucFreq
Rep
Read depth
100
75
50
25
0
0.0
2,000.0
4,000.0
6,000.0
8,000.0
10,000.0
12,000.0
14,000.0
chr14_haplotype1-0000013 (kbp)

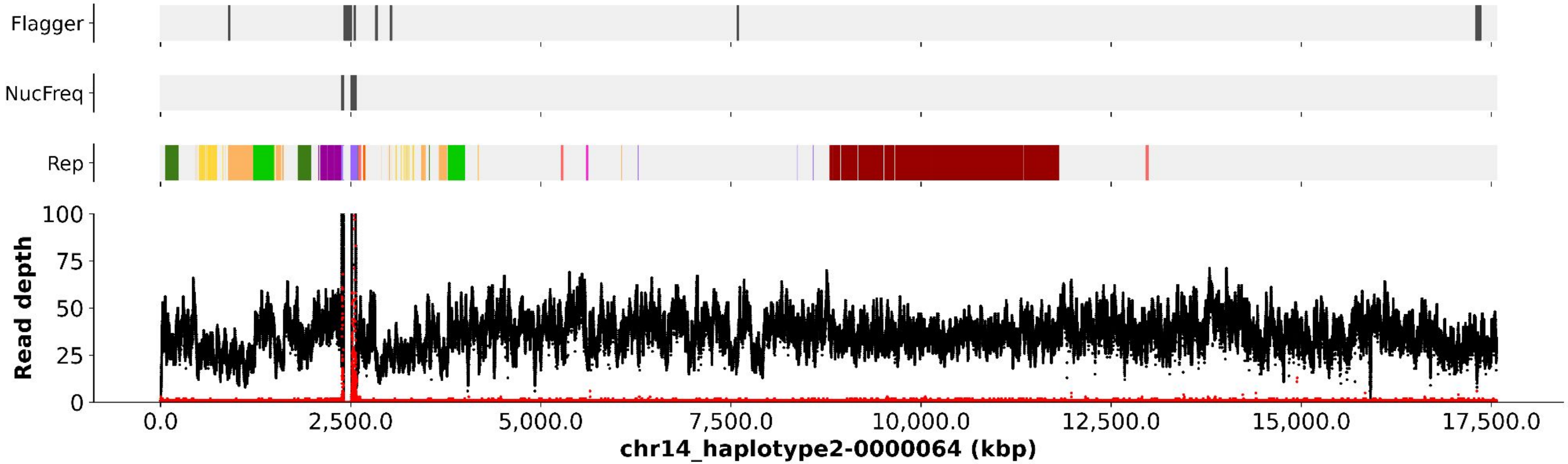
Flagger
NucFreq
Rep
Read depth
100
75
50
25
0
0.0
2,500.0
5,000.0
7,500.0
10,000.0
12,500.0
15,000.0
17,500.0
chr14_haplotype2-0000064 (kbp)

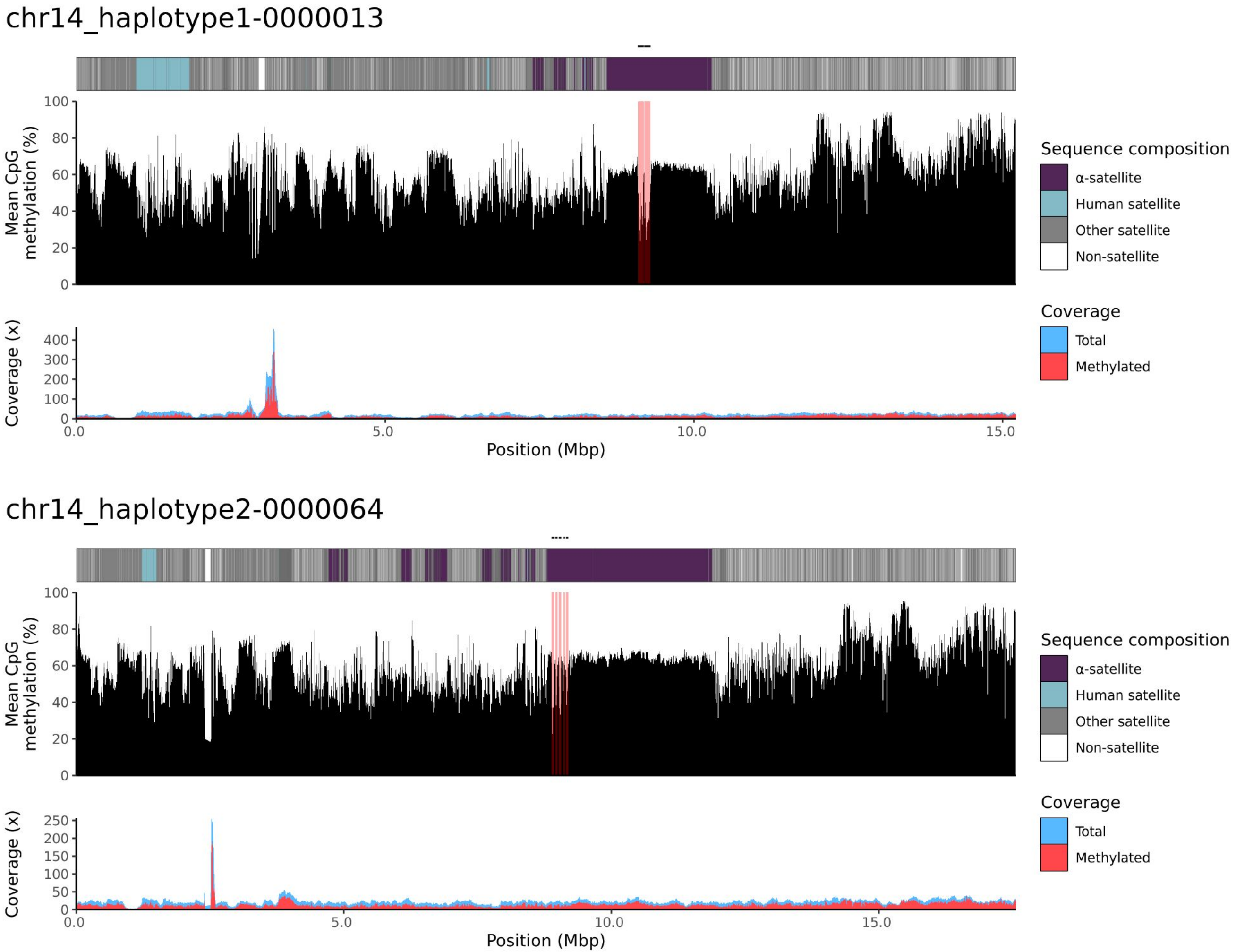

chr14_haplotype1-0000013
Mean CpG methylation (%)
Coverage (x)
Position (Mbp)
Sequence composition
α-satellite
Human satellite
Other satellite
Non-satellite
Coverage
Total
Methylated
chr14_haplotype2-0000064
Mean CpG methylation (%)
Coverage (x)
Position (Mbp)
Sequence composition
α-satellite
Human satellite
Other satellite
Non-satellite
Coverage
Total
Methylated

### chr15

#### NA12885_1_haplotype1-0000016_chr15

results/chr15_1_22694466/moddotplot/NA12885_1/NA12885_1_haplotype1-0000016_chr1!

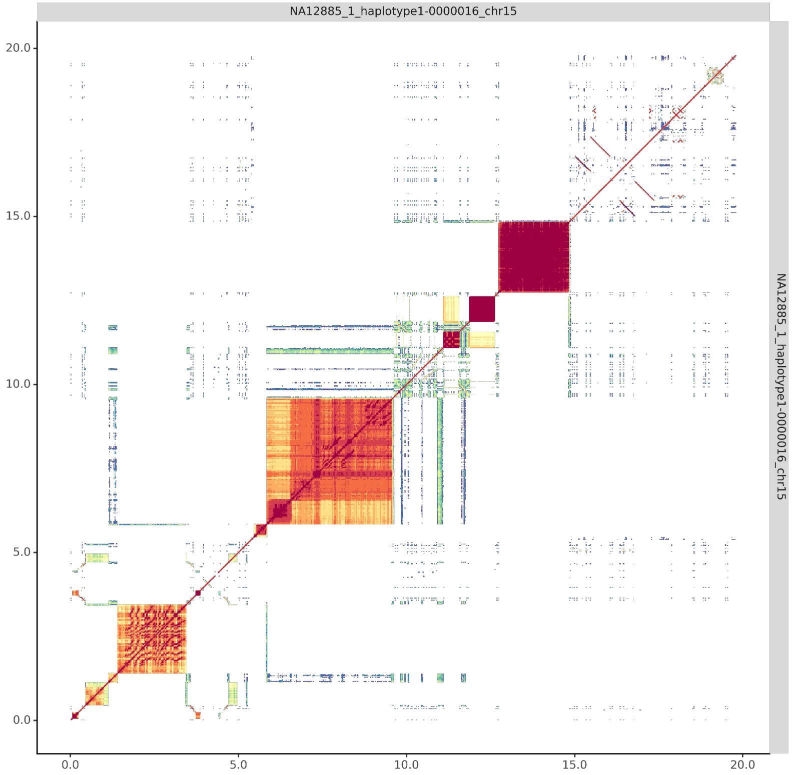


#### NA12885_2_haplotype2-0000072_chr15

results/chr15_1_22694466/moddotplot/NA12885_2/NA12885_2_haplotype2-0000072_chr1!

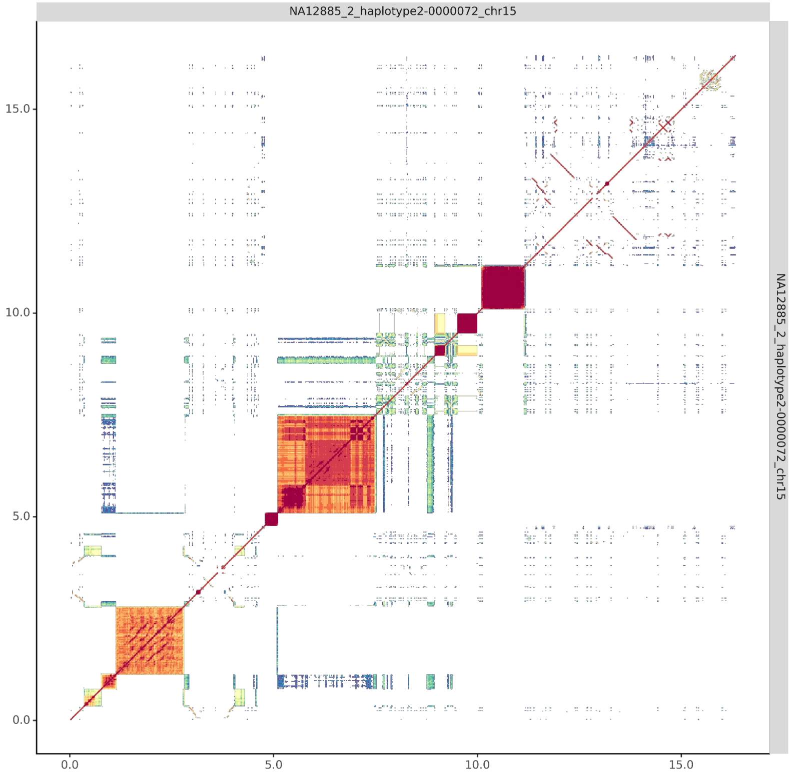

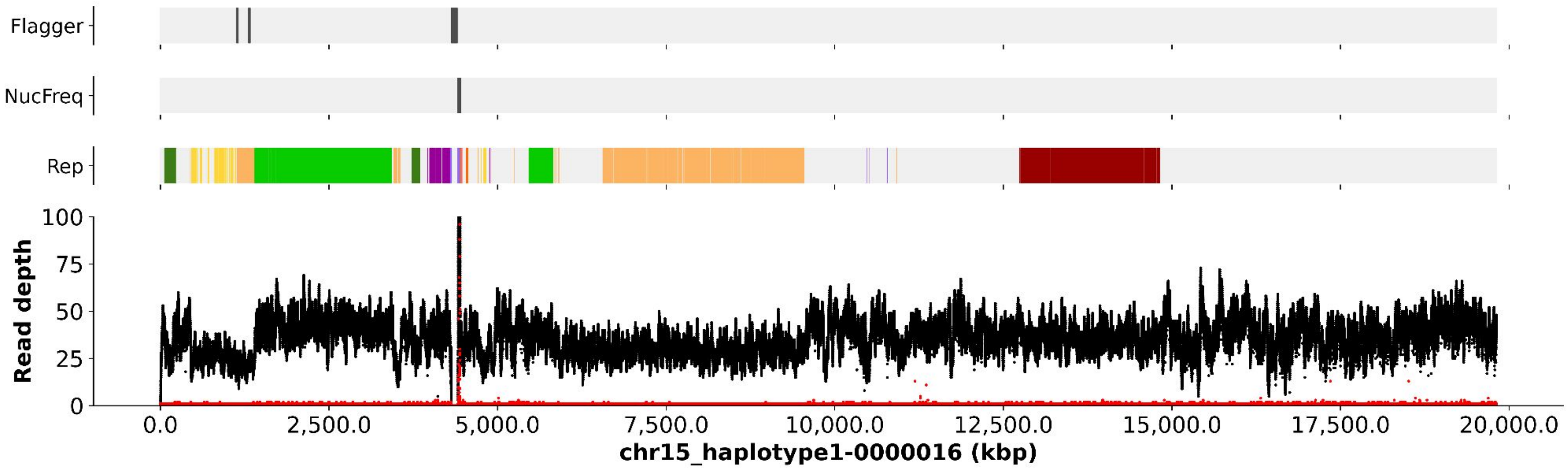
Flagger
NucFreq
Rep
Read depth
100
75
50
25
0
0.0
2,500.0
5,000.0
7,500.0
10,000.0
12,500.0
15,000.0
17,500.0
20,000.0
chr15_haplotype1-0000016 (kbp)

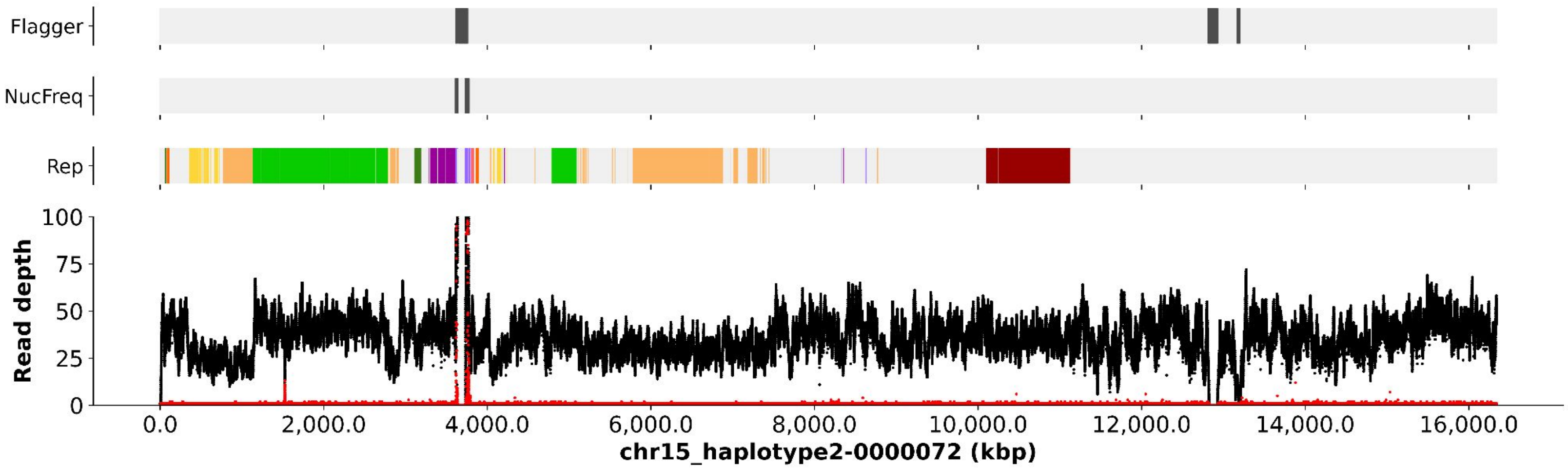
Flagger
NucFreq
Rep
Read depth
100
75
50
25
0
0.0
2,000.0
4,000.0
6,000.0
8,000.0
10,000.0
12,000.0
14,000.0
16,000.0
chr15_haplotype2-0000072 (kbp)

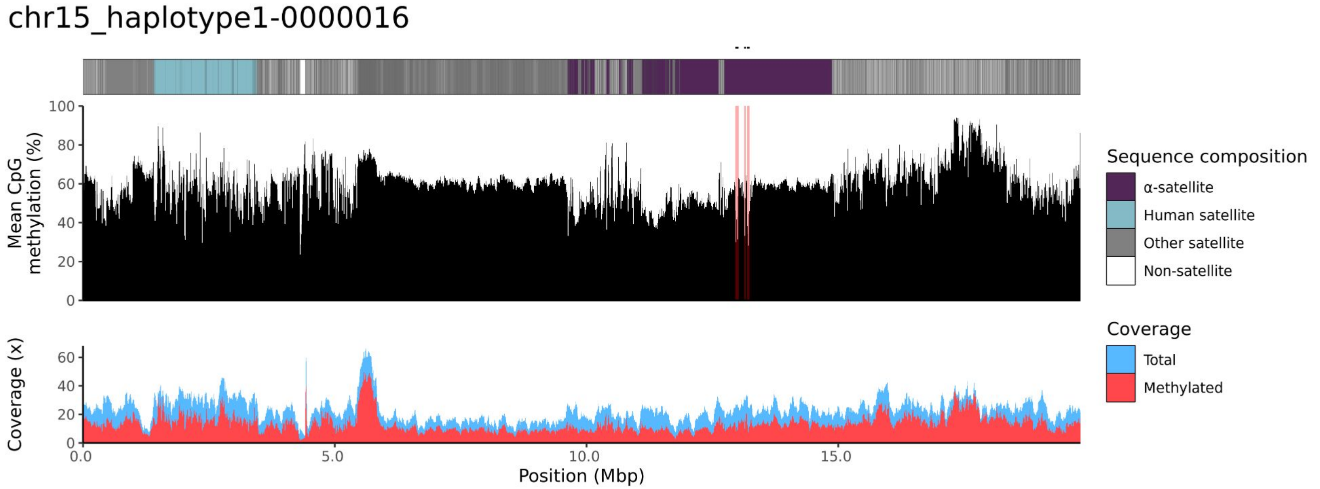
chr15_haplotype1-0000016
Mean CpG methylation (%)
Coverage (x)
Position (Mbp)
0.0
5.0
10.0
15.0
Sequence composition
α-satellite
Human satellite
Other satellite
Non-satellite
Coverage
Total
Methylated

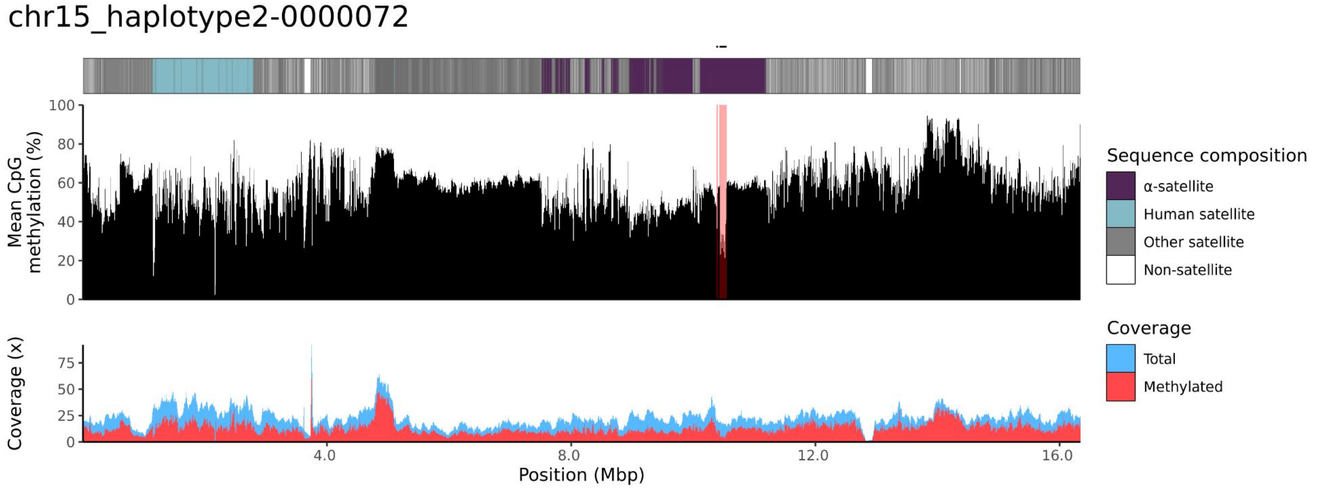
chr15_haplotype2-0000072
Mean CpG methylation (%)
Coverage (x)
Position (Mbp)
4.0
8.0
12.0
16.0
Sequence composition
α-satellite
Human satellite
Other satellite
Non-satellite
Coverage
Total
Methylated

### chr21

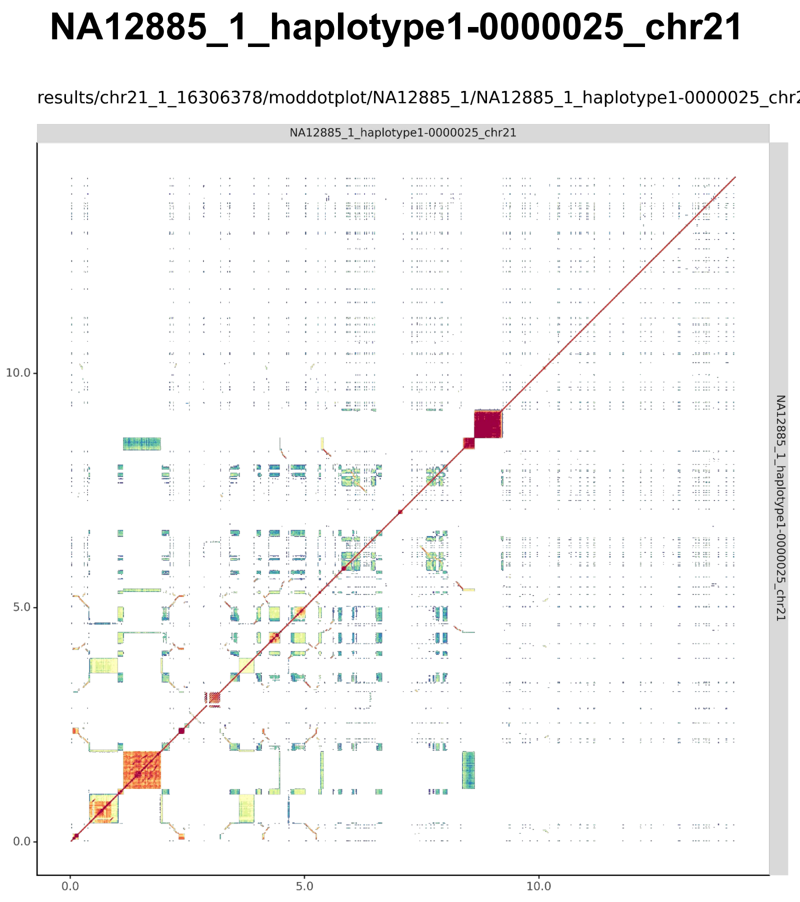
NA12885_1_haplotype1-0000025_chr21
results/chr21_1_16306378/moddotplot/NA12885_1/NA12885_1_haplotype1-0000025_chr2
NA12885_1_haplotype1-0000025_chr21
10.0
5.0
0.0
0.0
5.0
10.0
NA12885_1_haplotype1-0000025_chr21

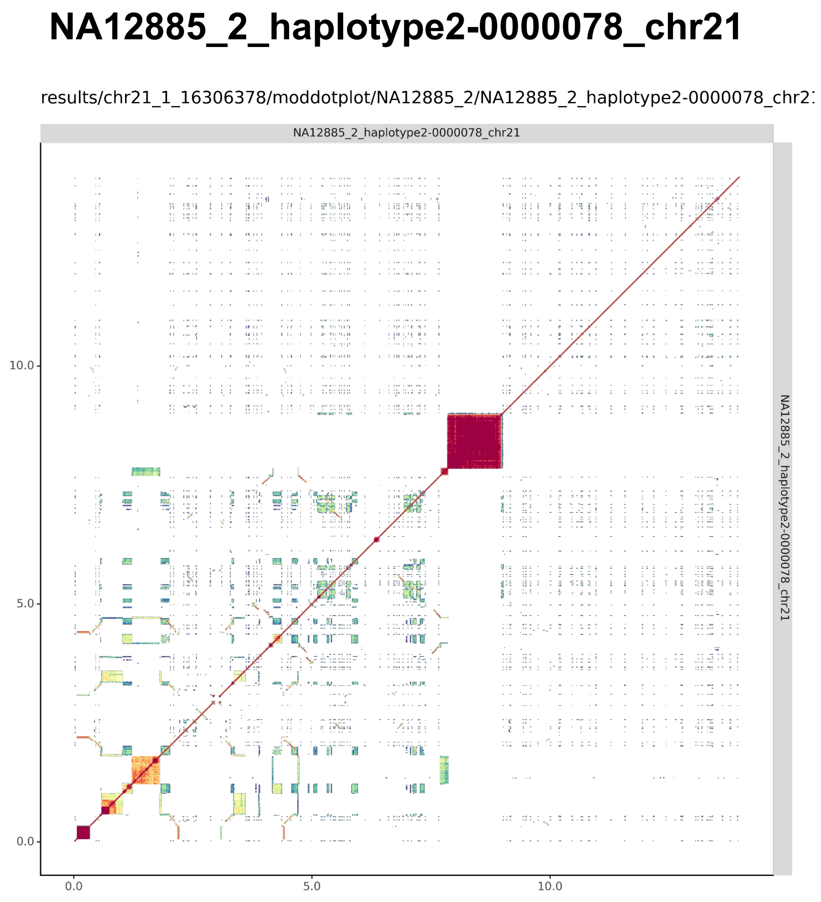
NA12885_2_haplotype2-0000078_chr21
results/chr21_1_16306378/moddotplot/NA12885_2/NA12885_2_haplotype2-0000078_chr2
NA12885_2_haplotype2-0000078_chr21
10.0
5.0
0.0
0.0
5.0
10.0
NA12885_2_haplotype2-0000078_chr21

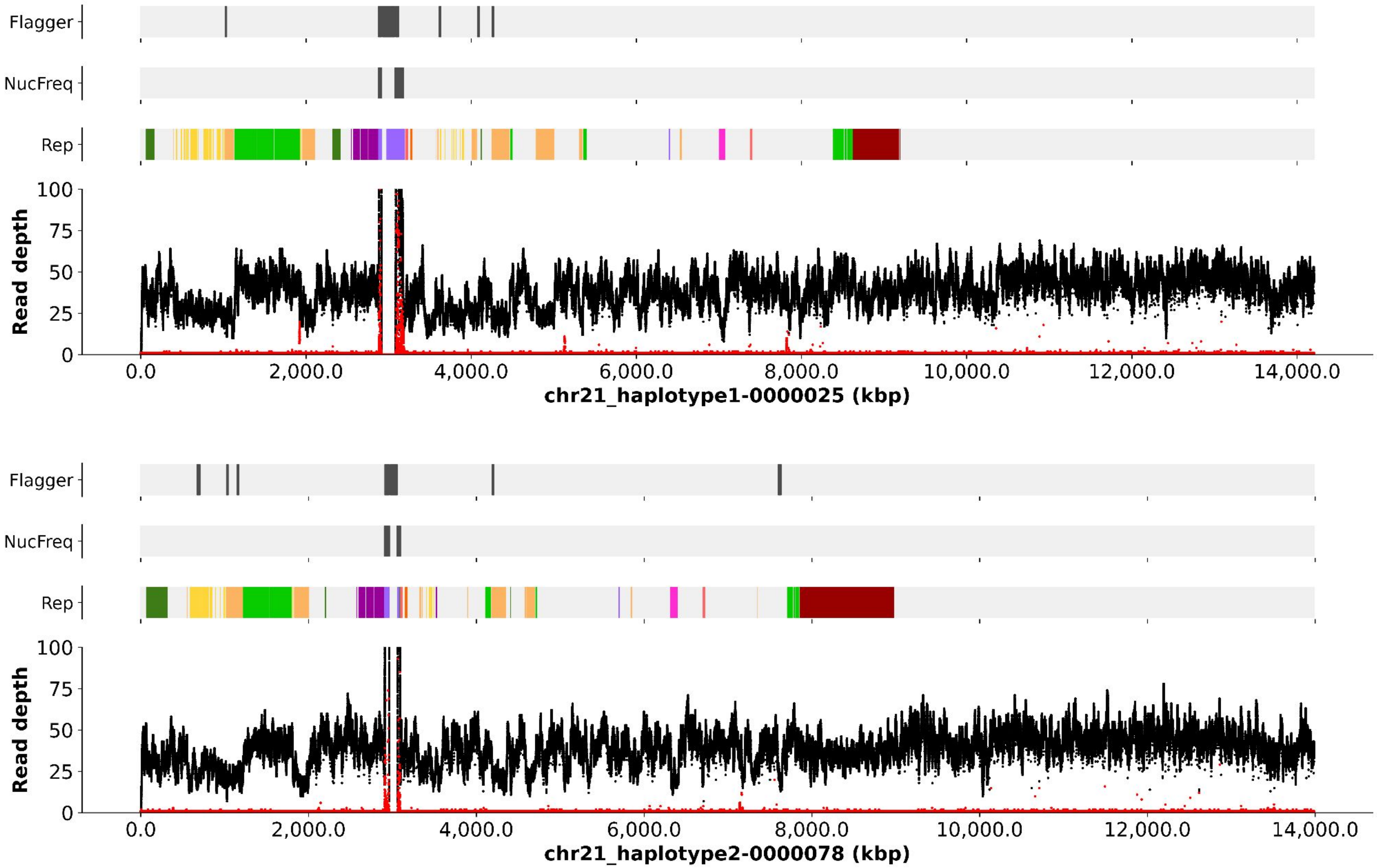

Flagger
NucFreq
Rep
Read depth
100
75
50
25
0
0.0
2,000.0
4,000.0
6,000.0
8,000.0
10,000.0
12,000.0
14,000.0
chr21_haplotype1-0000025 (kbp)
Flagger
NucFreq
Rep
Read depth
100
75
50
25
0
0.0
2,000.0
4,000.0
6,000.0
8,000.0
10,000.0
12,000.0
14,000.0
chr21_haplotype2-0000078 (kbp)

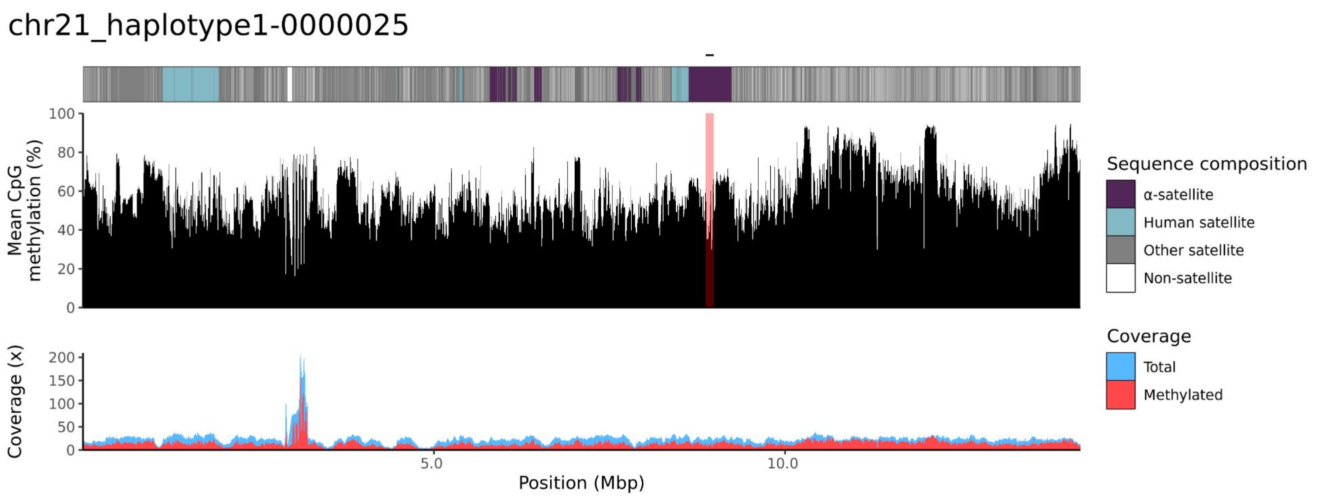
chr21_haplotype1-0000025
Mean CpG methylation (%)
100
80
60
40
20
0
Coverage (x)
200
150
100
50
0
5.0
10.0
Position (Mbp)
Sequence composition
α-satellite
Human satellite
Other satellite
Non-satellite
Coverage
Total
Methylated

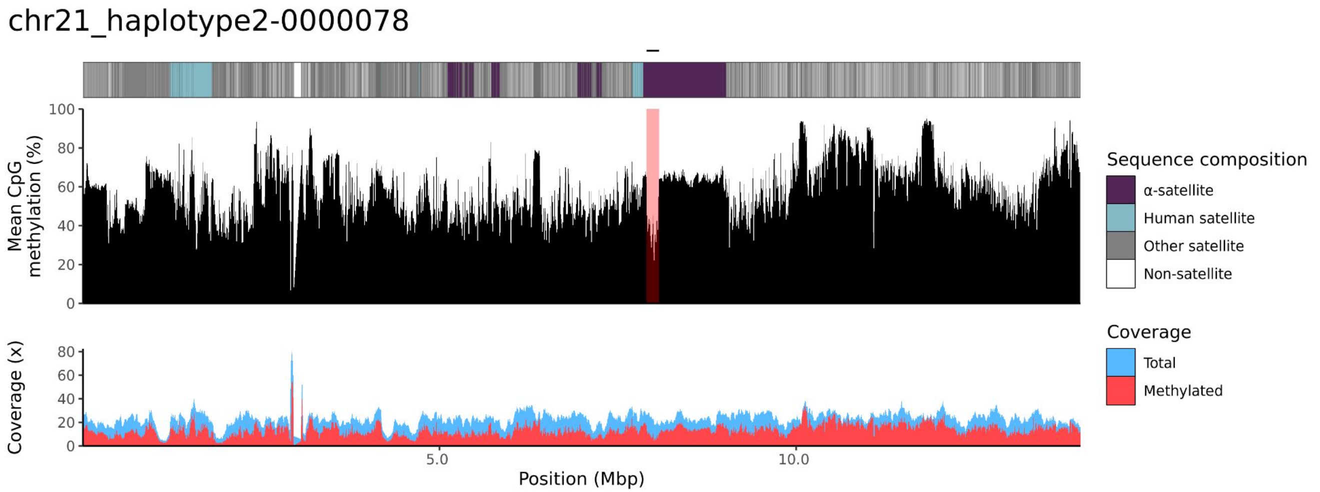
chr21_haplotype2-0000078
Mean CpG methylation (%)
100
80
60
40
20
0
Coverage (x)
80
60
40
20
0
5.0
10.0
Position (Mbp)
Sequence composition
α-satellite
Human satellite
Other satellite
Non-satellite
Coverage
Total
Methylated

### chr22

#### NA12885_1_haplotype1-0000009_chr22

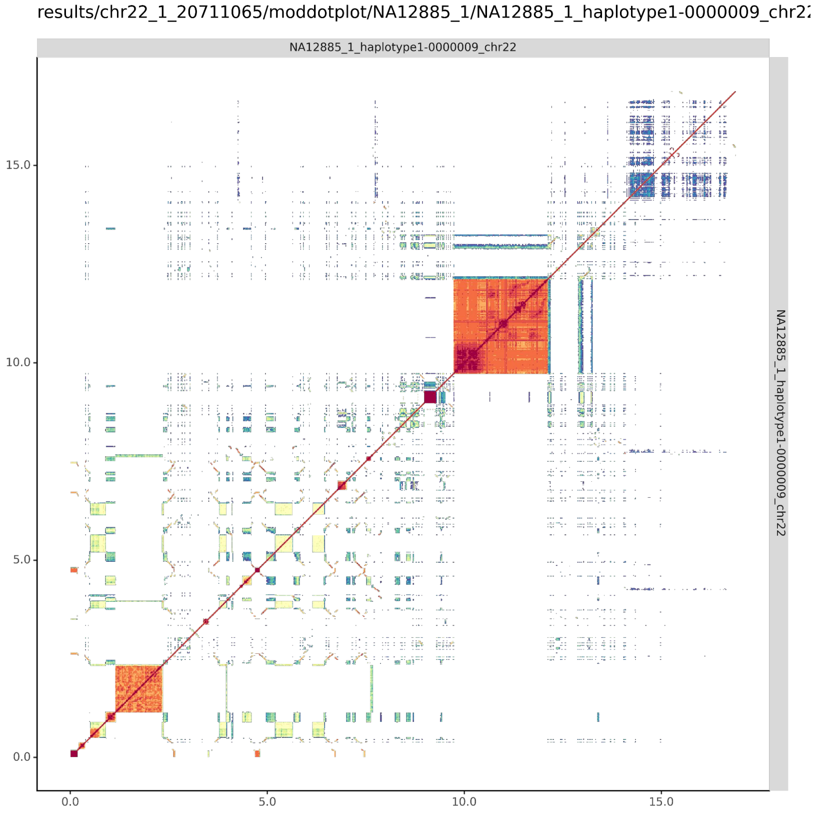


#### NA12885_2_haplotype2-0000067_chr22

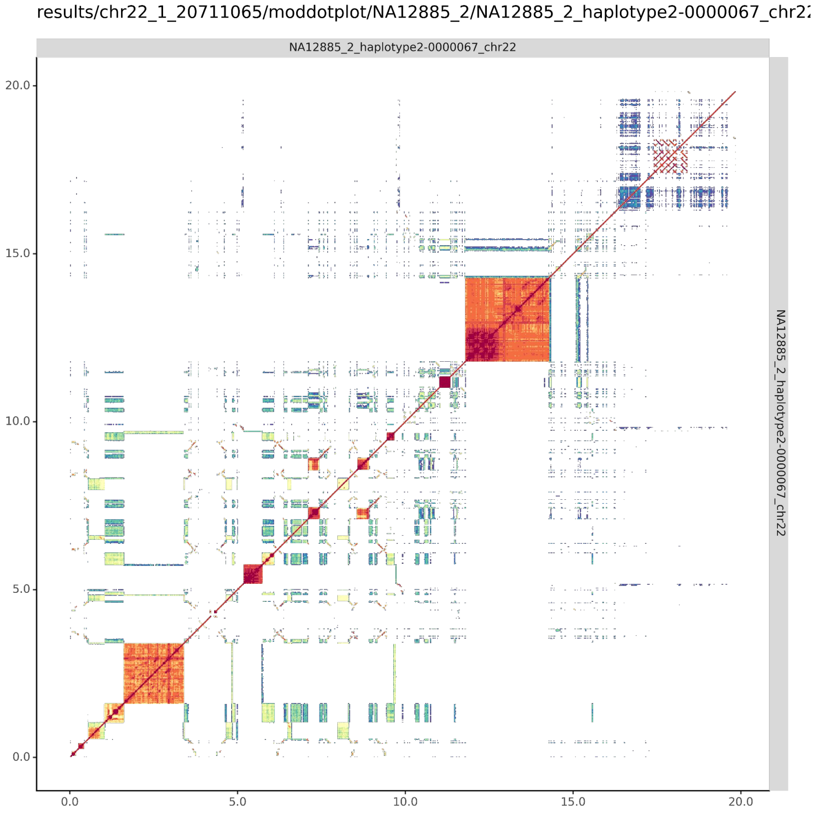

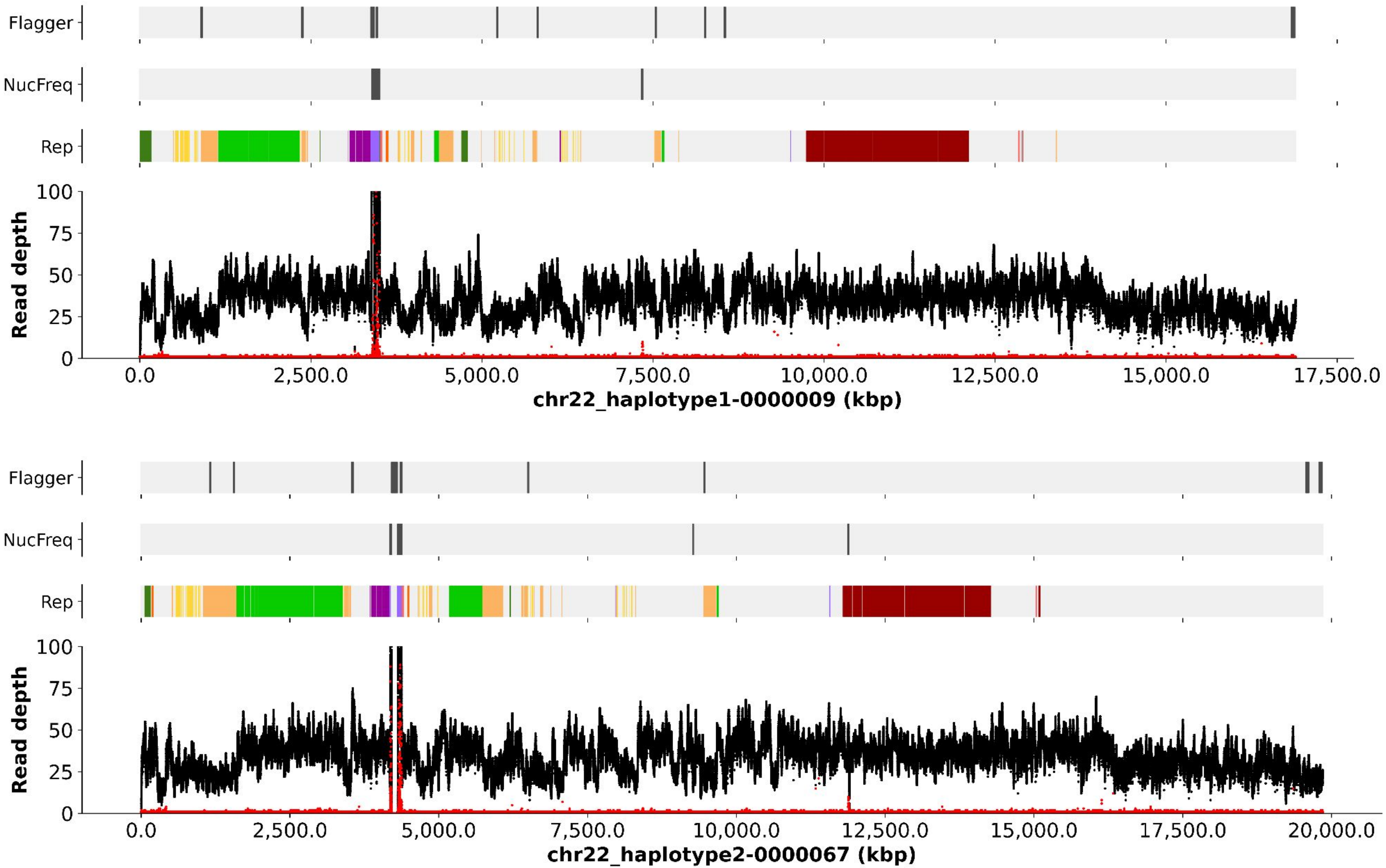
Flagger
NucFreq
Rep
Read depth
100
75
50
25
0
0.0
2,500.0
5,000.0
7,500.0
10,000.0
12,500.0
15,000.0
17,500.0
chr22_haplotype1-0000009 (kbp)
Flagger
NucFreq
Rep
Read depth
100
75
50
25
0
0.0
2,500.0
5,000.0
7,500.0
10,000.0
12,500.0
15,000.0
17,500.0
20,000.0
chr22_haplotype2-0000067 (kbp)

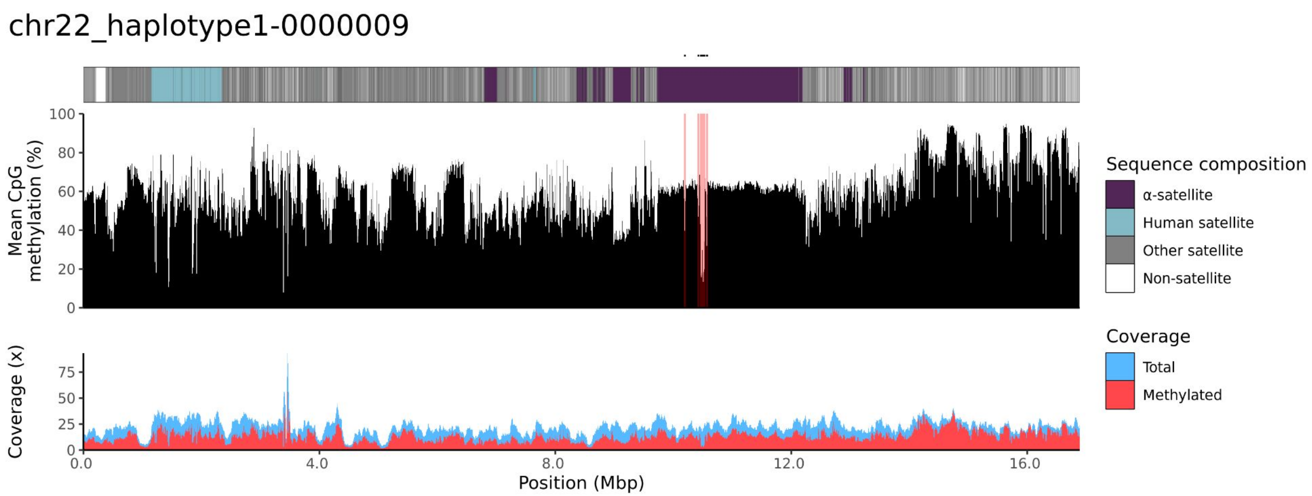
chr22_haplotype1-0000009
Mean CpG methylation (%)
Coverage (x)
Position (Mbp)
Sequence composition
α-satellite
Human satellite
Other satellite
Non-satellite
Coverage
Total
Methylated

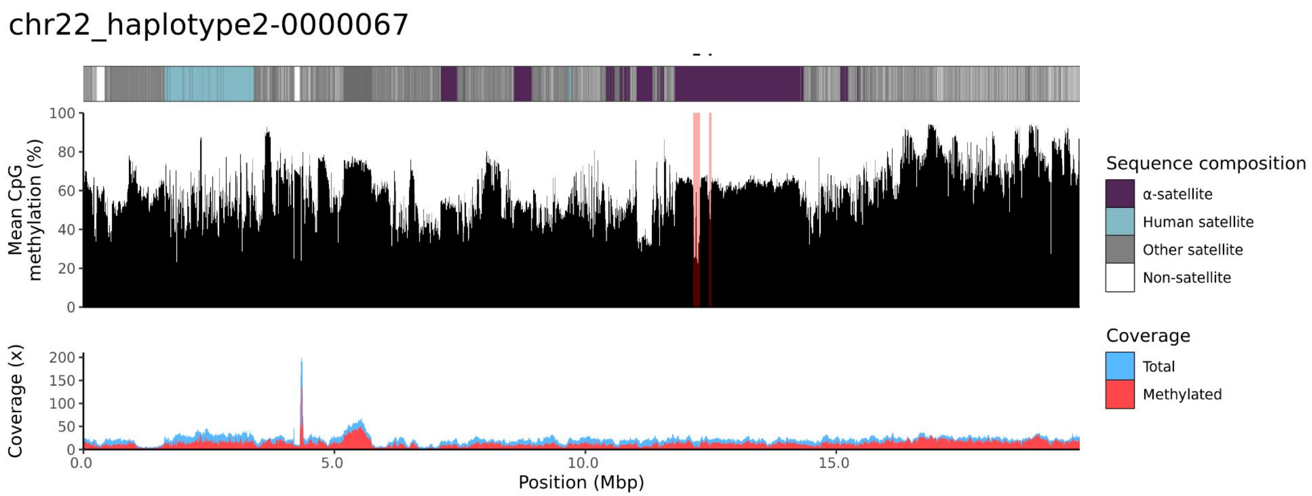
chr22_haplotype2-0000067
Mean CpG methylation (%)
Coverage (x)
Position (Mbp)
Sequence composition
α-satellite
Human satellite
Other satellite
Non-satellite
Coverage
Total
Methylated

NA12886

### chr13

#### NA12886_1_haplotype1-0000017_chr13

results/chr13_1_22508596/moddotplot/NA12886_1/NA12886_1_haplotype1-0000017_chr1:

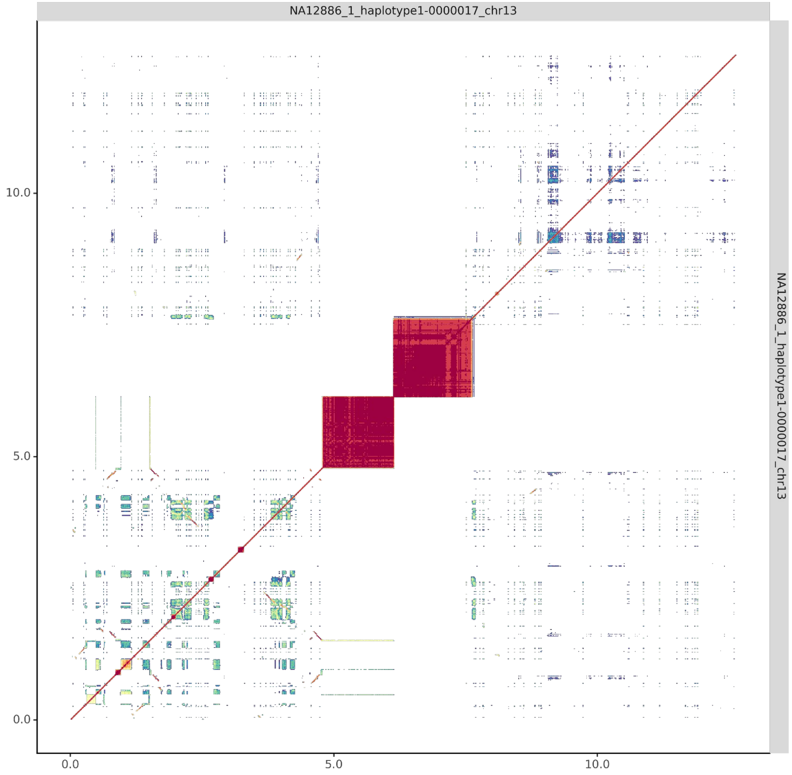


#### NA12886_2_haplotype2-0000086_chr13

results/chr13_1_22508596/moddotplot/NA12886_2/NA12886_2_haplotype2-0000086_chr1:

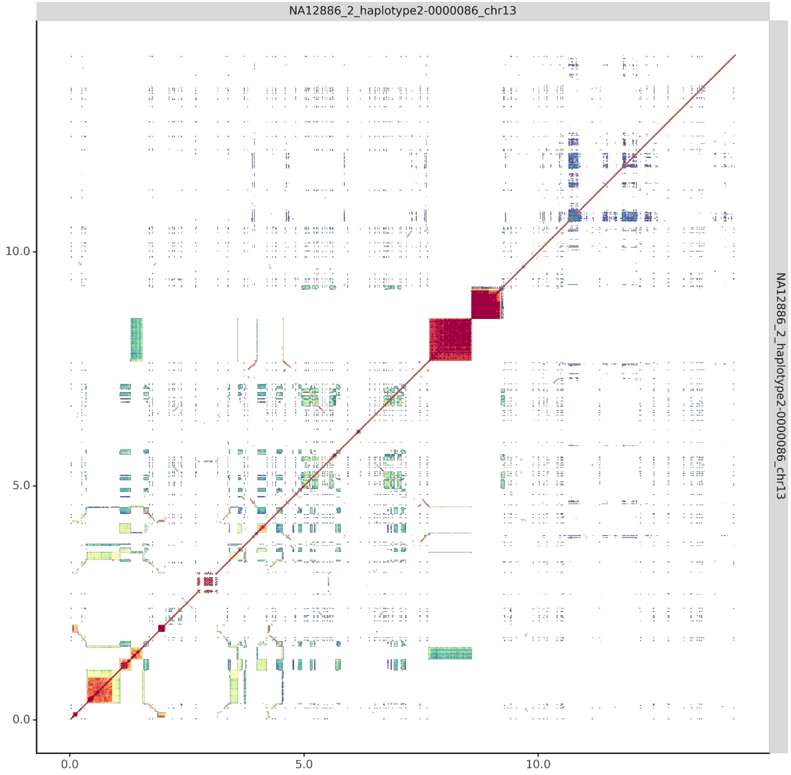

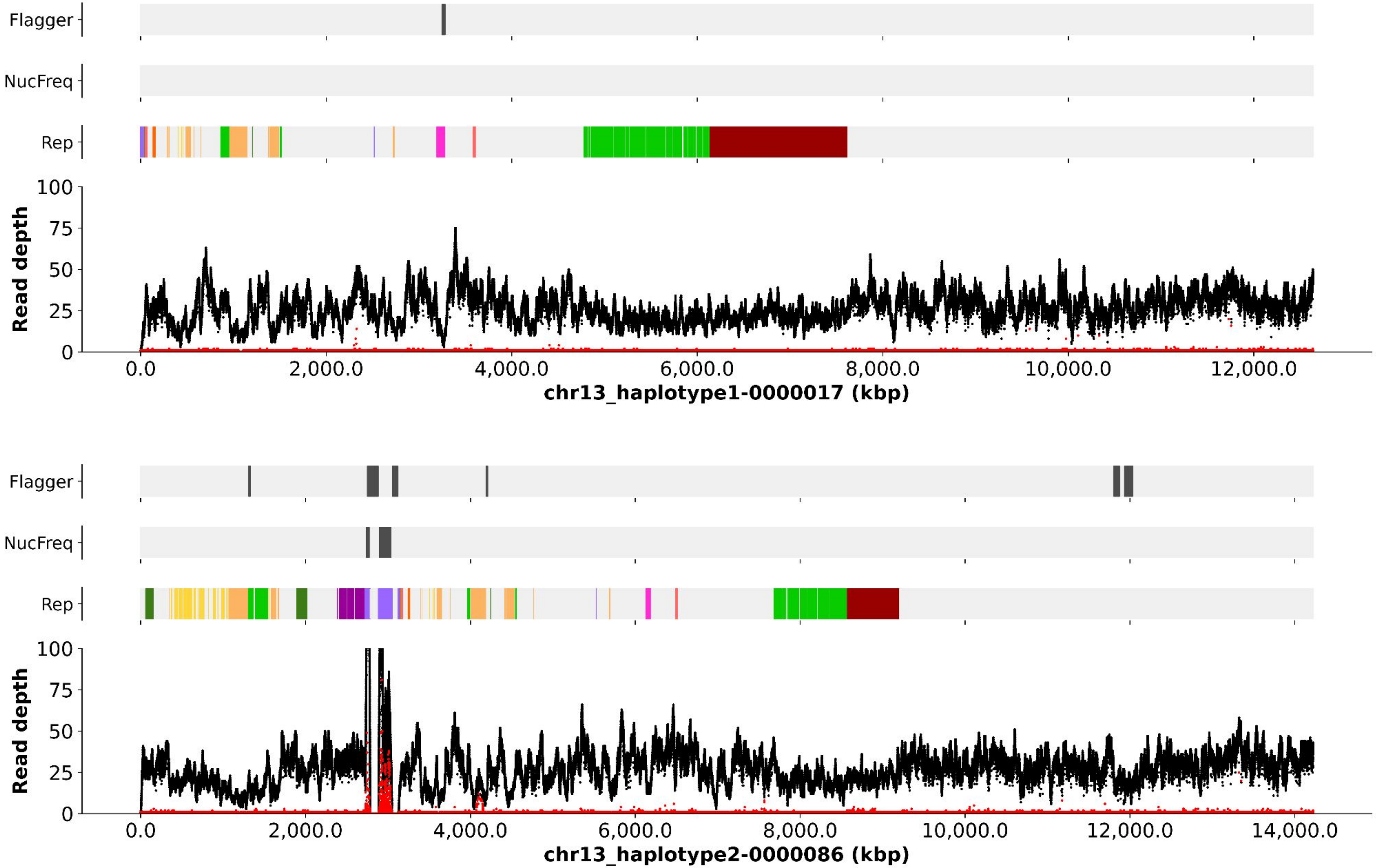

Flagger
NucFreq
Rep
Read depth
100
75
50
25
0
0.0
2,000.0
4,000.0
6,000.0
8,000.0
10,000.0
12,000.0
chr13_haplotype1-0000017 (kbp)
Flagger
NucFreq
Rep
Read depth
100
75
50
25
0
0.0
2,000.0
4,000.0
6,000.0
8,000.0
10,000.0
12,000.0
14,000.0
chr13_haplotype2-0000086 (kbp)

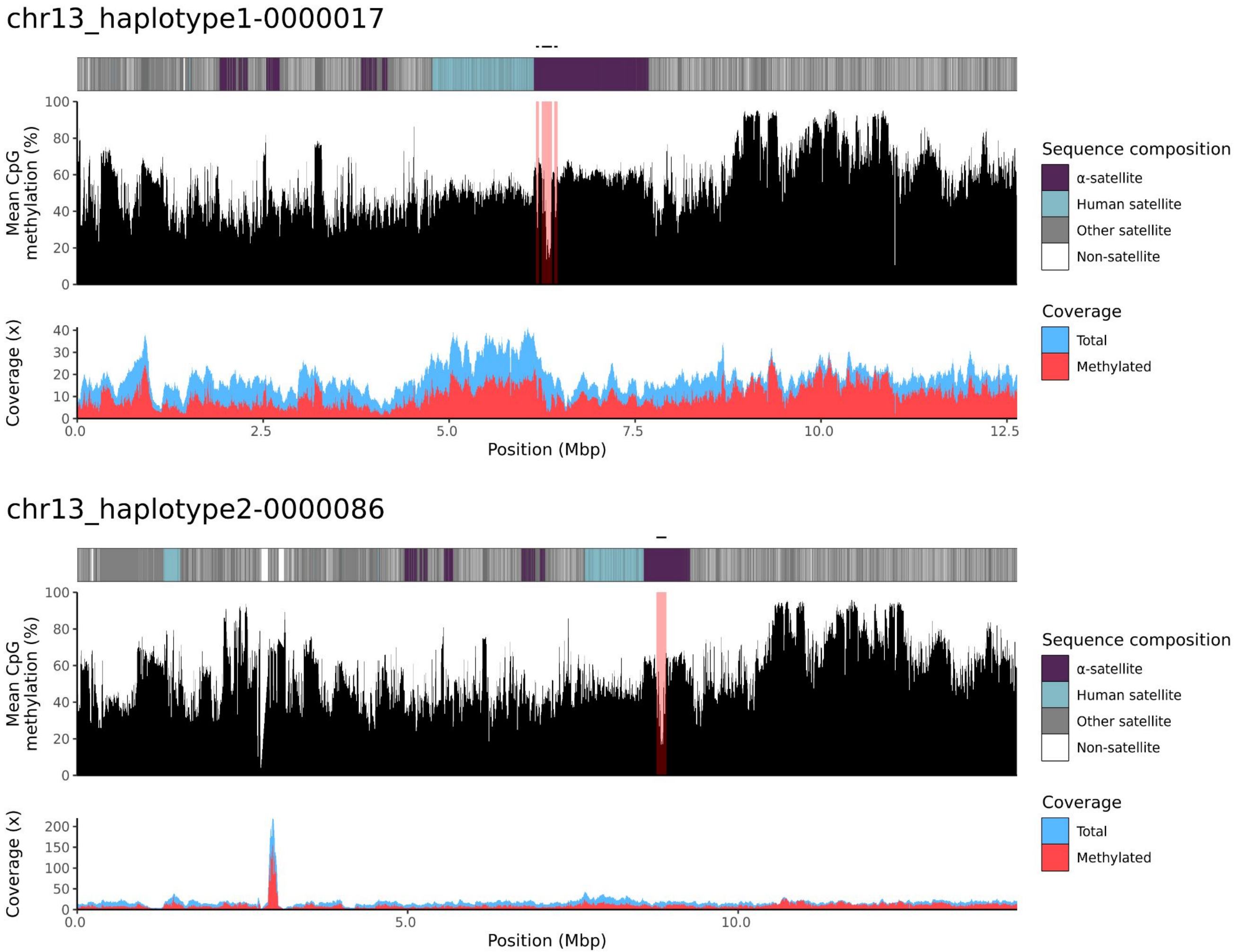
chr13_haplotype1-0000017
Mean CpG methylation (%)
100
80
60
40
20
0
Coverage (x)
40
30
20
10
0
0.0
2.5
5.0
7.5
10.0
12.5
Position (Mbp)
Sequence composition
α-satellite
Human satellite
Other satellite
Non-satellite
Coverage
Total
Methylated
chr13_haplotype2-0000086
Mean CpG methylation (%)
100
80
60
40
20
0
Coverage (x)
200
150
100
50
0
0.0
5.0
10.0
Position (Mbp)
Sequence composition
α-satellite
Human satellite
Other satellite
Non-satellite
Coverage
Total
Methylated

### chr14

#### NA12886_1_haplotype1-0000009_chr14

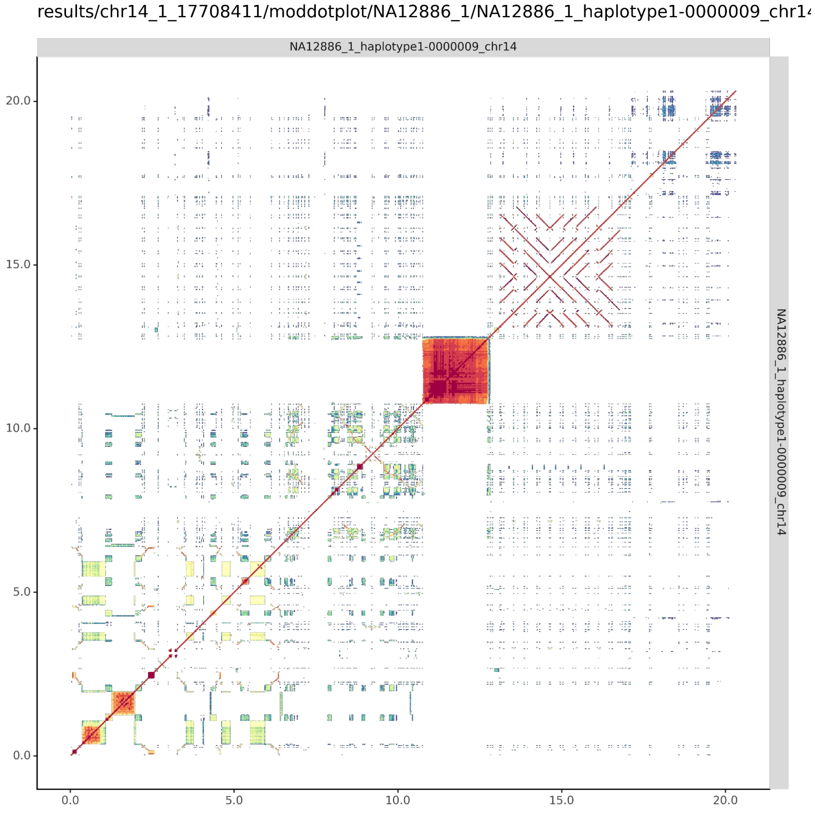


#### NA12886_2_haplotype2-0000085_chr14

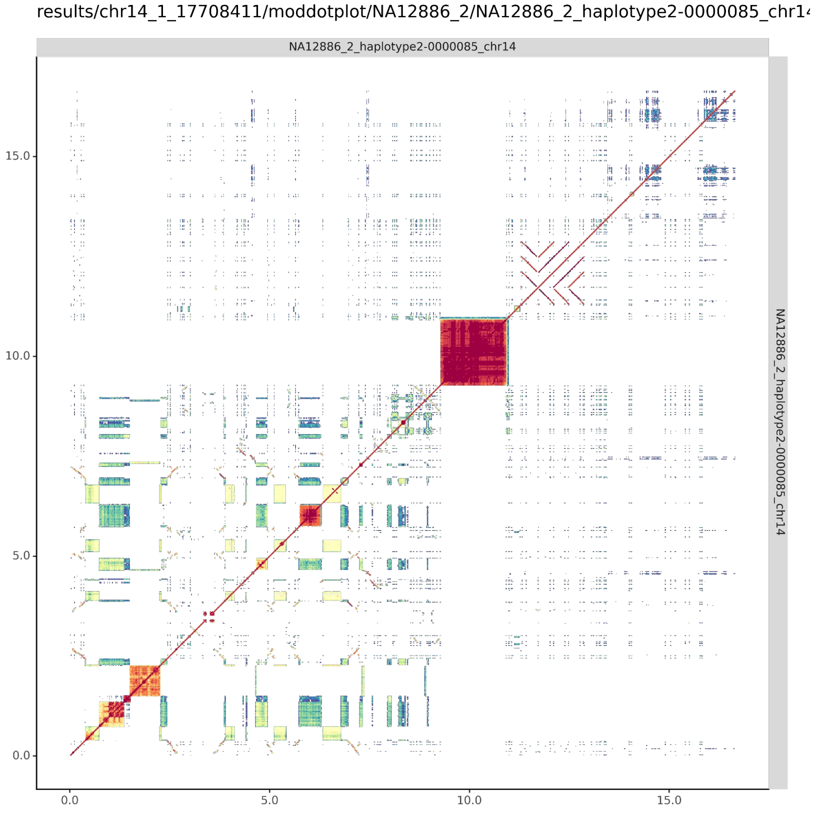

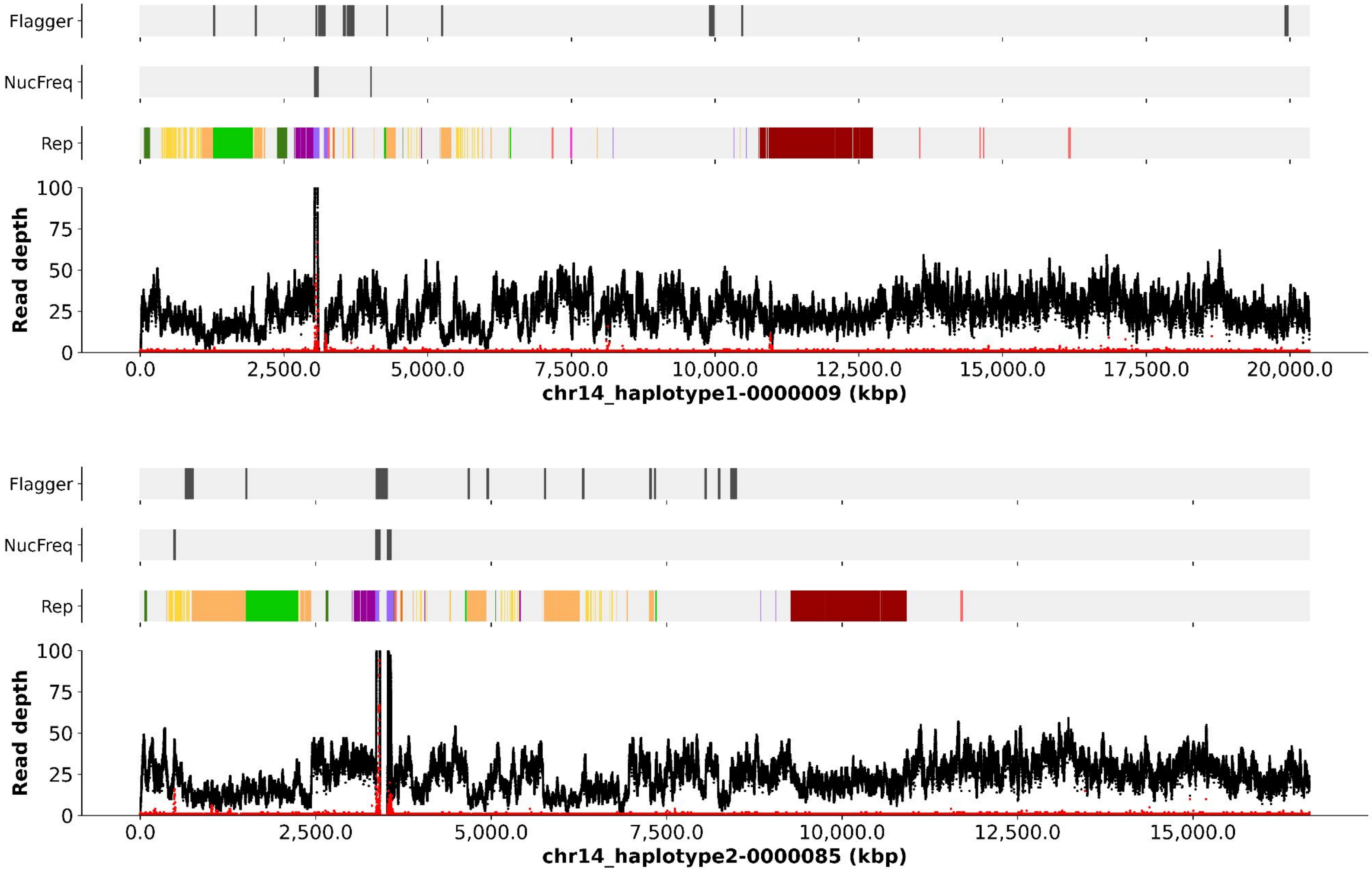
Flagger
NucFreq
Rep
Read depth
100
75
50
25
0
0.0
2,500.0
5,000.0
7,500.0
10,000.0
12,500.0
15,000.0
17,500.0
20,000.0
chr14_haplotype1-0000009 (kbp)
Flagger
NucFreq
Rep
Read depth
100
75
50
25
0
0.0
2,500.0
5,000.0
7,500.0
10,000.0
12,500.0
15,000.0
chr14_haplotype2-0000085 (kbp)

#### chr14_haplotype1-0000009

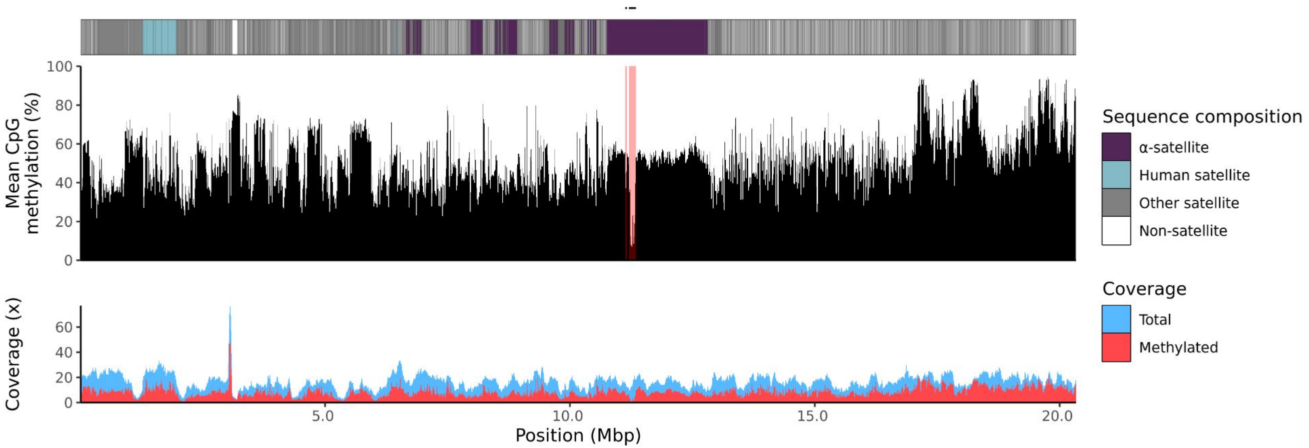


#### chr14_haplotype2-0000085

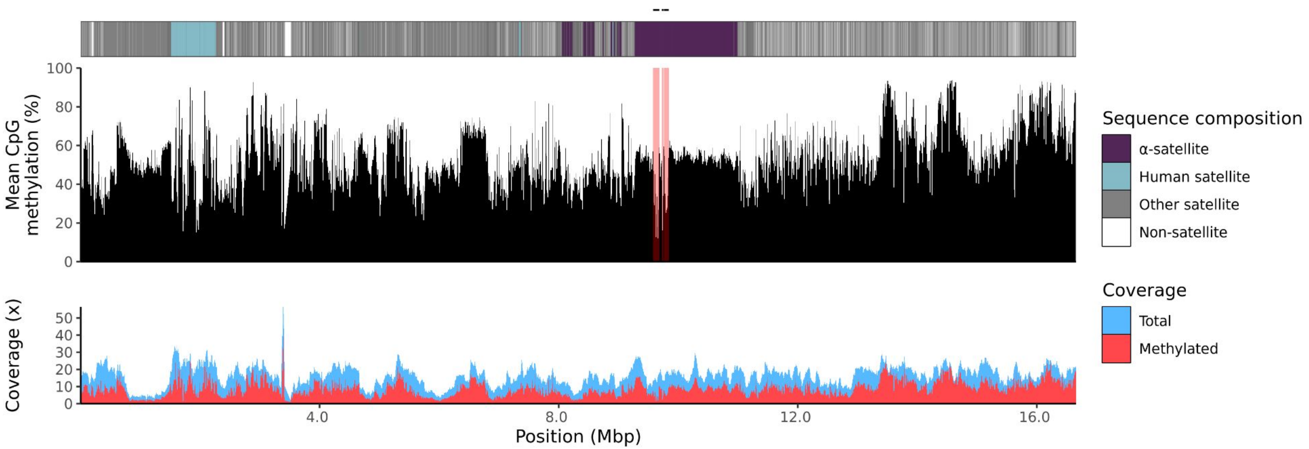

### chr15

#### NA12886_1_haplotype1-0000027_chr15

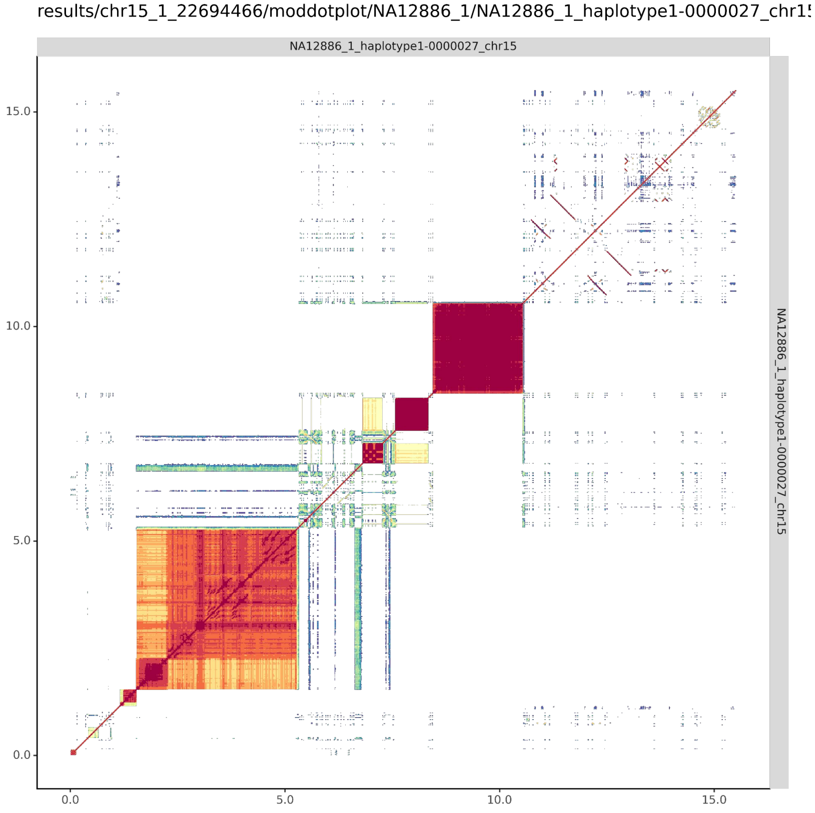


#### NA12886_2_haplotype2-0000095_chr15

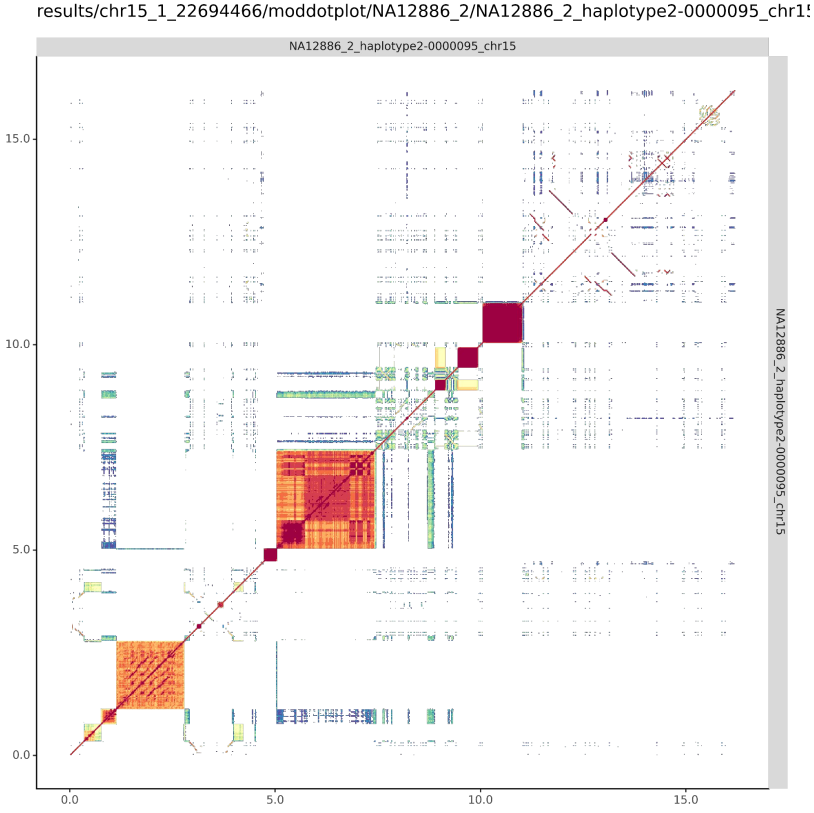

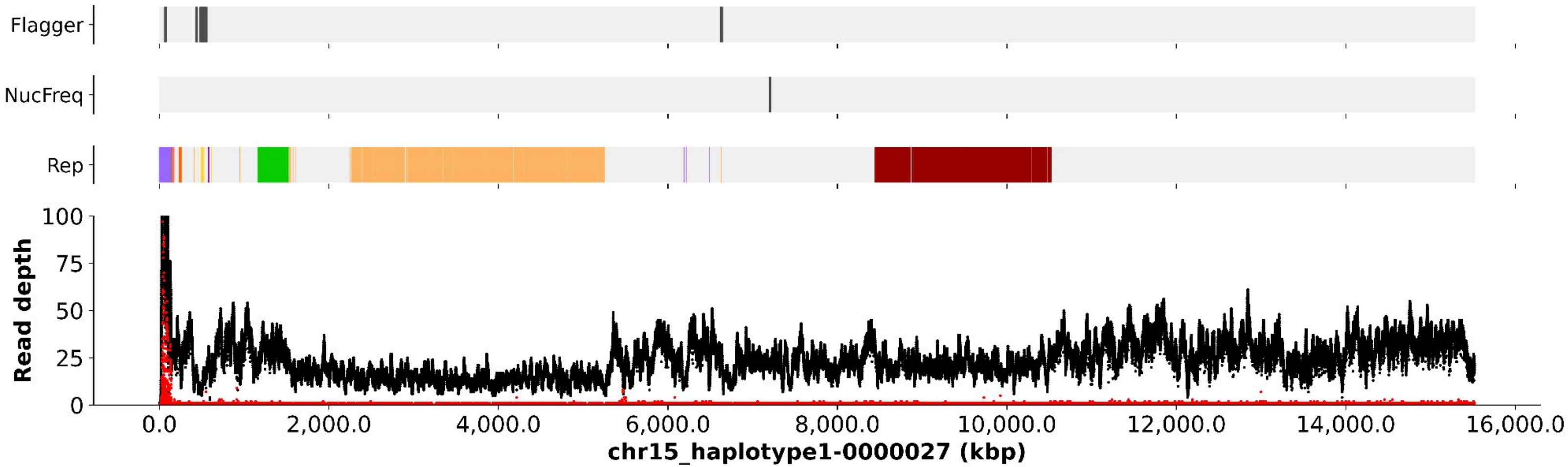
Flagger
NucFreq
Rep
Read depth
100
75
50
25
0
0.0
2,000.0
4,000.0
6,000.0
8,000.0
10,000.0
12,000.0
14,000.0
16,000.0
chr15_haplotype1-0000027 (kbp)

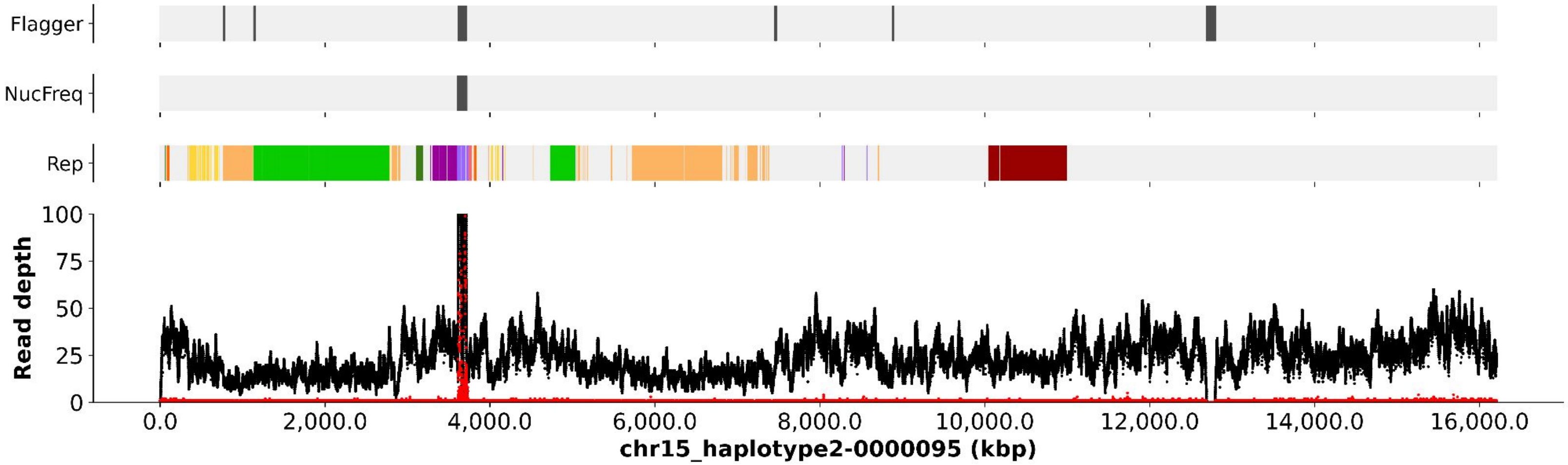
Flagger
NucFreq
Rep
Read depth
100
75
50
25
0
0.0
2,000.0
4,000.0
6,000.0
8,000.0
10,000.0
12,000.0
14,000.0
16,000.0
chr15_haplotype2-0000095 (kbp)

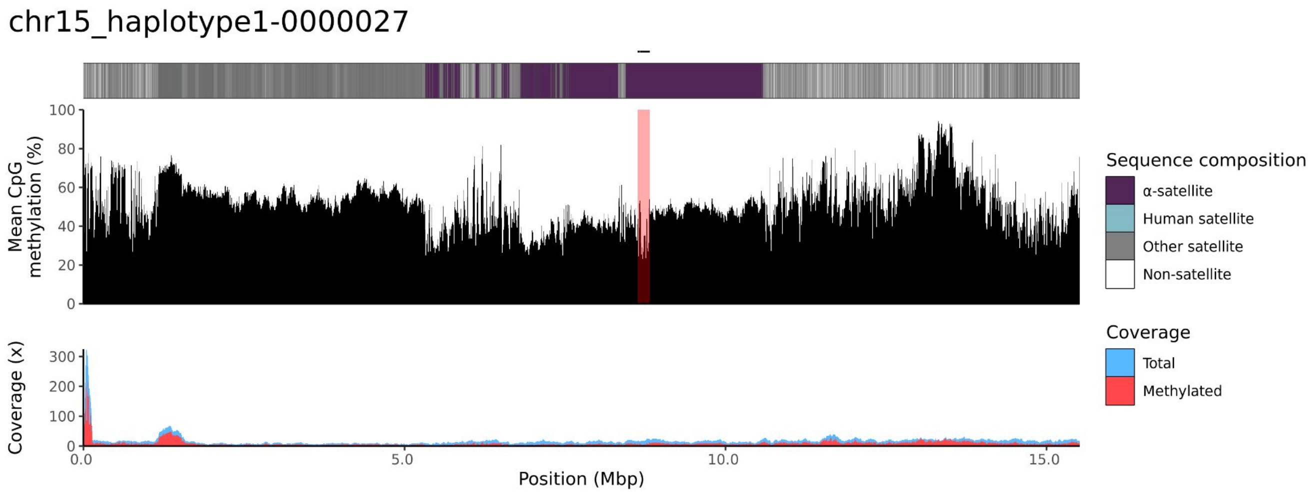
chr15_haplotype1-0000027
Mean CpG methylation (%)
100
80
60
40
20
0
Coverage (x)
300
200
100
0
0.0
5.0
10.0
15.0
Position (Mbp)
Sequence composition
α-satellite
Human satellite
Other satellite
Non-satellite
Coverage
Total
Methylated

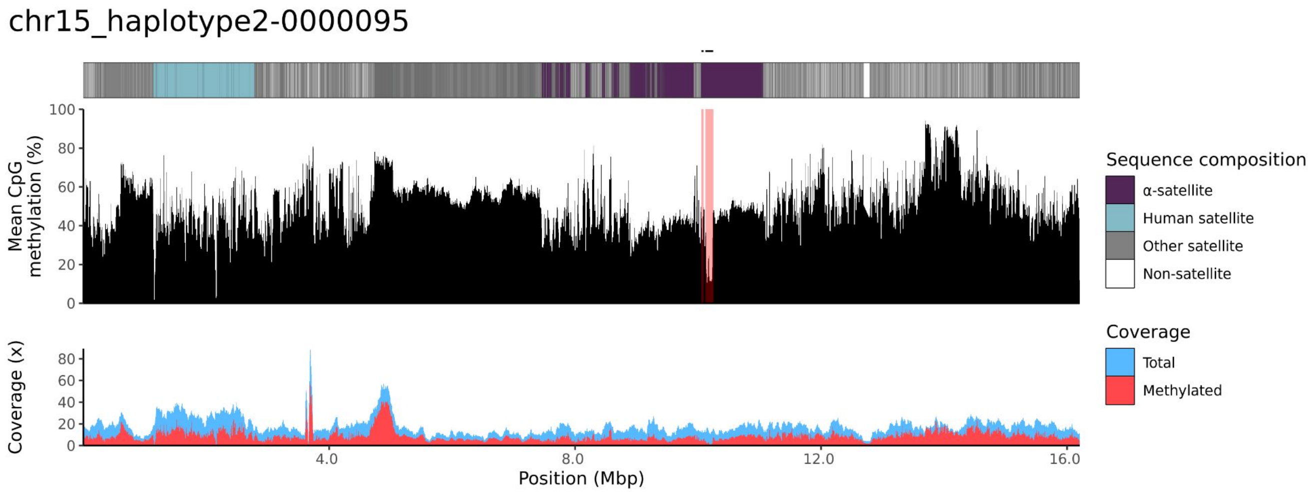
chr15_haplotype2-0000095
Mean CpG methylation (%)
100
80
60
40
20
0
Coverage (x)
80
60
40
20
0
4.0
8.0
12.0
16.0
Position (Mbp)
Sequence composition
α-satellite
Human satellite
Other satellite
Non-satellite
Coverage
Total
Methylated

### chr21

#### NA12886_1_haplotype1-0000005_chr21

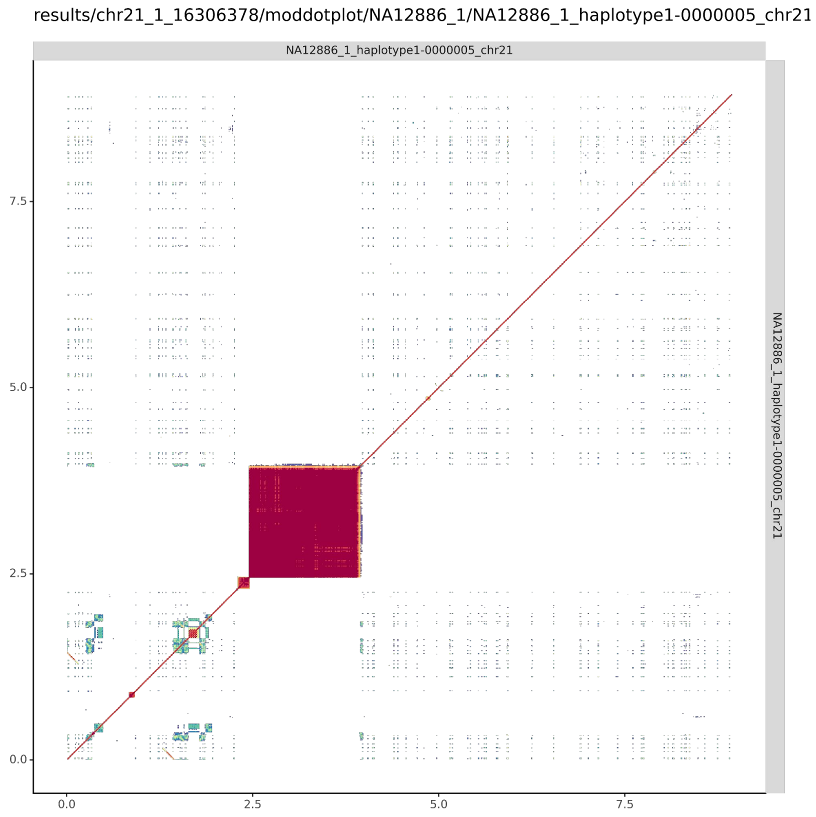


#### NA12886_2_haplotype2-0000078_chr21

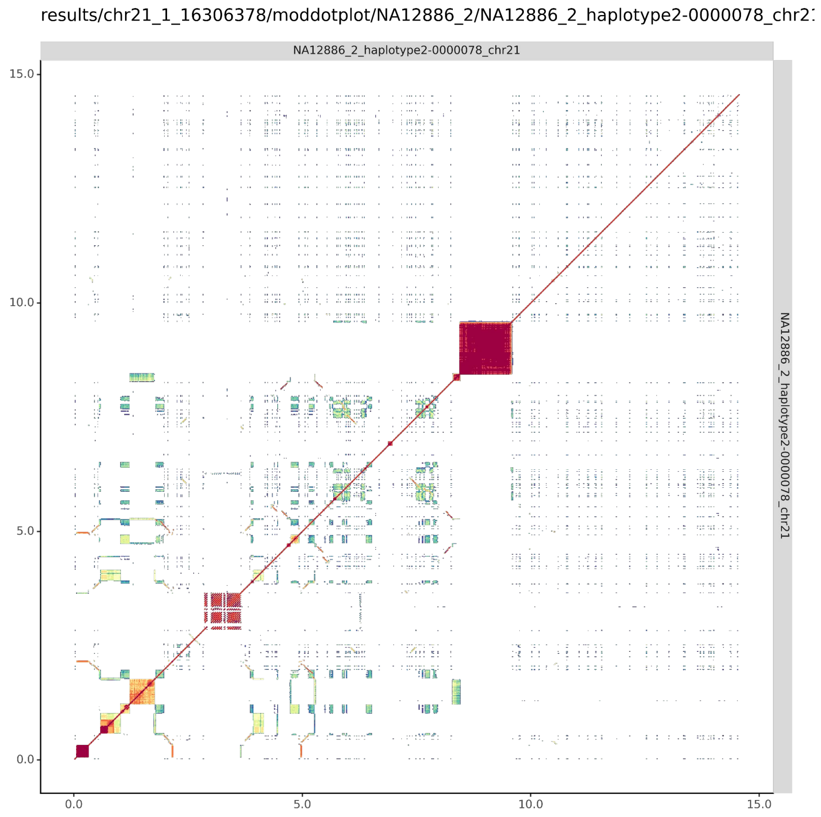

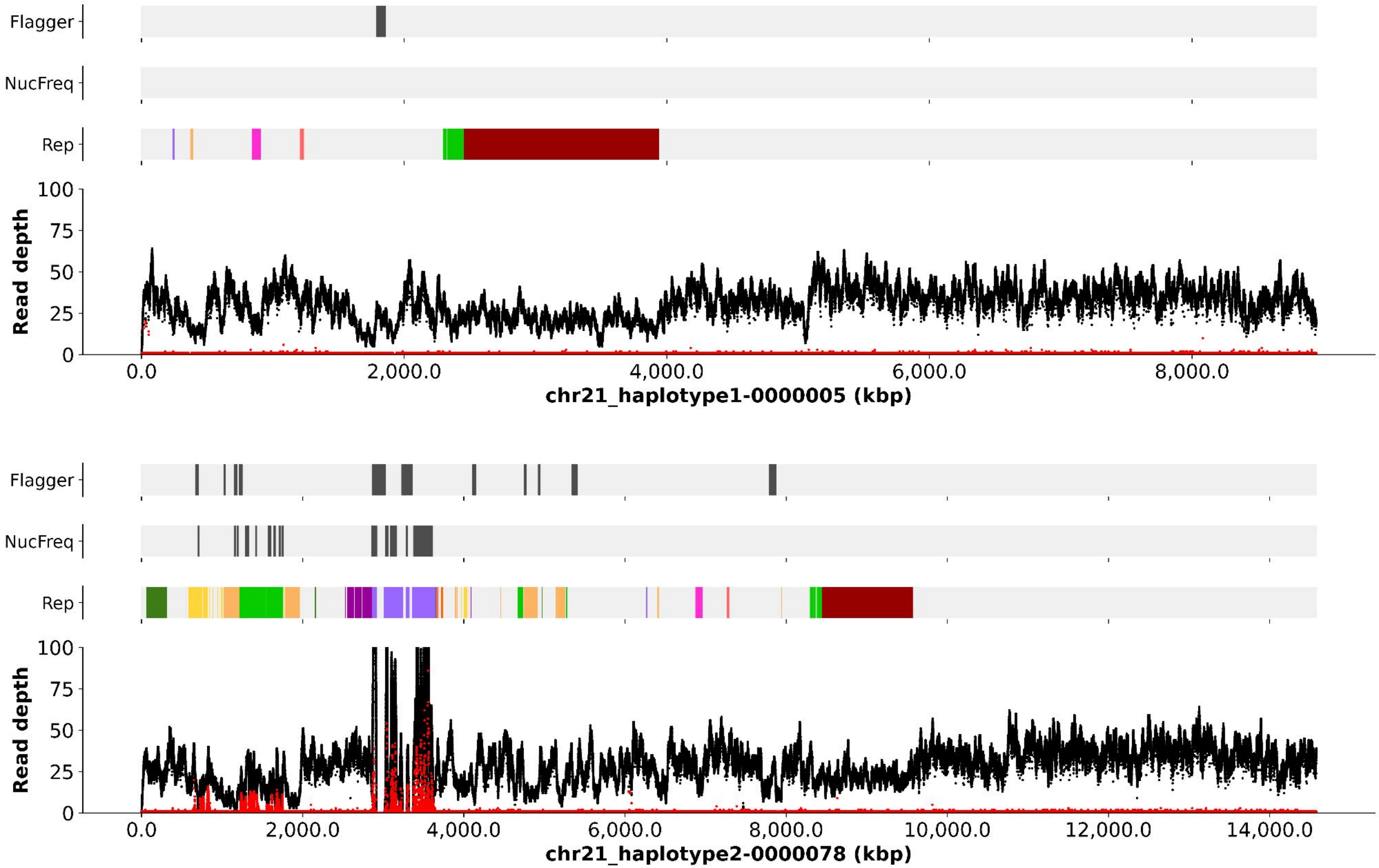
Flagger
NucFreq
Rep
Read depth
100
75
50
25
0
0.0
2,000.0
4,000.0
6,000.0
8,000.0
chr21_haplotype1-0000005 (kbp)
Flagger
NucFreq
Rep
Read depth
100
75
50
25
0
0.0
2,000.0
4,000.0
6,000.0
8,000.0
10,000.0
12,000.0
14,000.0
chr21_haplotype2-0000078 (kbp)

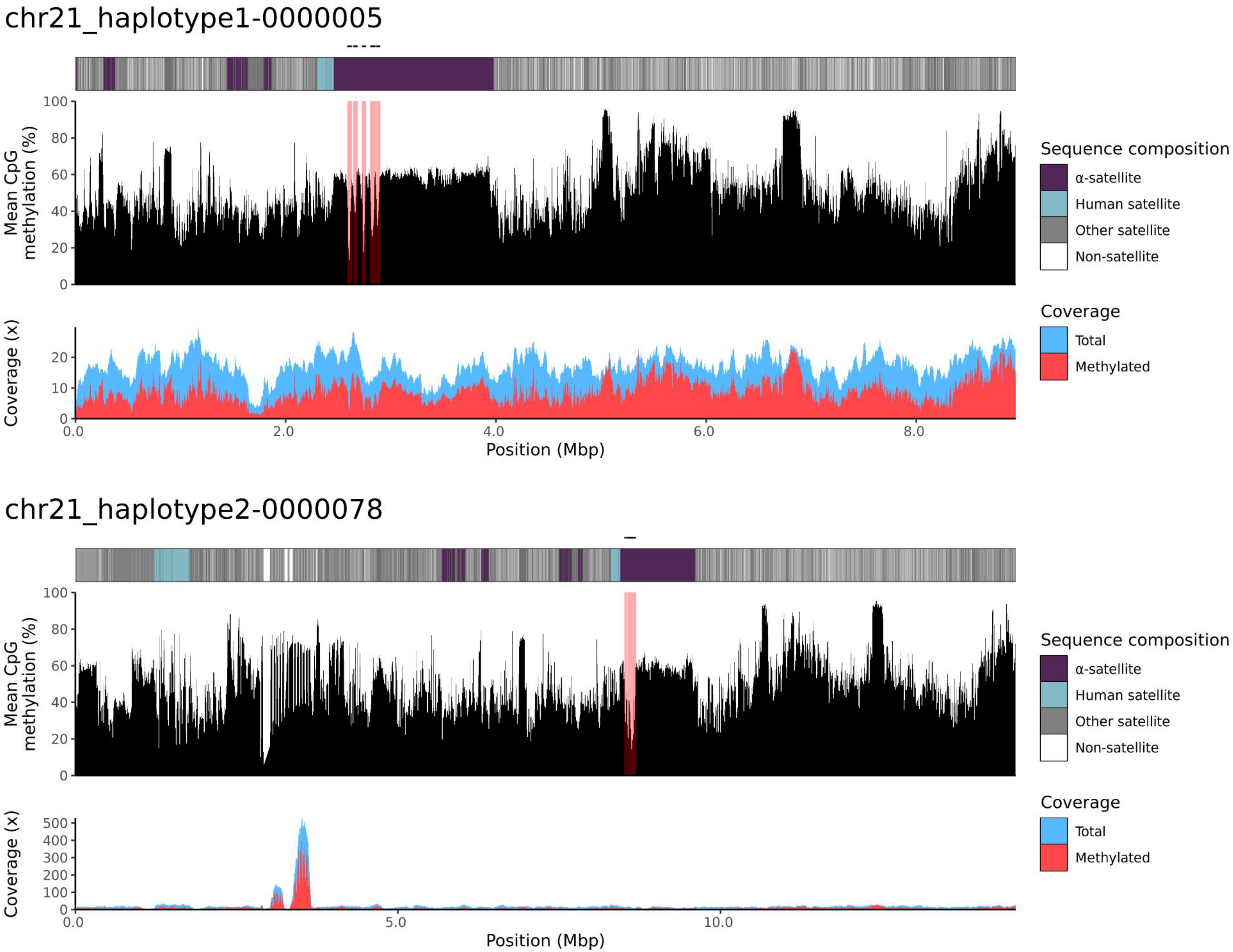
chr21_haplotype1-0000005
Mean CpG methylation (%)
100
80
60
40
20
0
Coverage (x)
20
10
0
0.0
2.0
4.0
6.0
8.0
Position (Mbp)
Sequence composition
α-satellite
Human satellite
Other satellite
Non-satellite
Coverage
Total
Methylated
chr21_haplotype2-0000078
Mean CpG methylation (%)
100
80
60
40
20
0
Coverage (x)
500
400
300
200
100
0
0.0
5.0
10.0
Position (Mbp)
Sequence composition
α-satellite
Human satellite
Other satellite
Non-satellite
Coverage
Total
Methylated

### chr22

#### NA12886_1_haplotype1-0000015_chr22

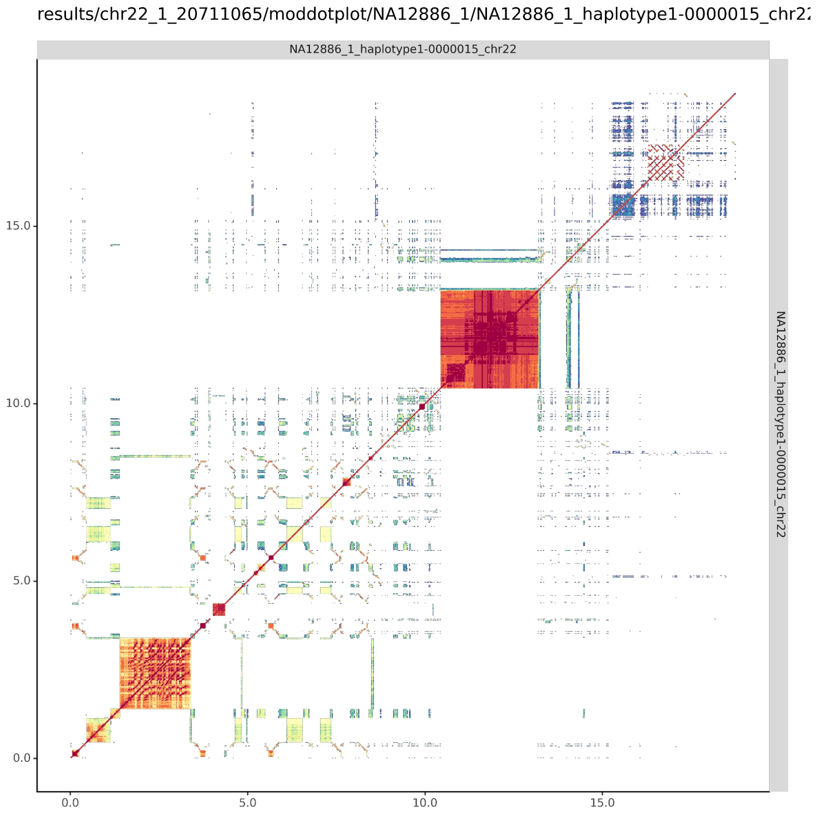


#### NA12886_2_haplotype2-0000091_chr22

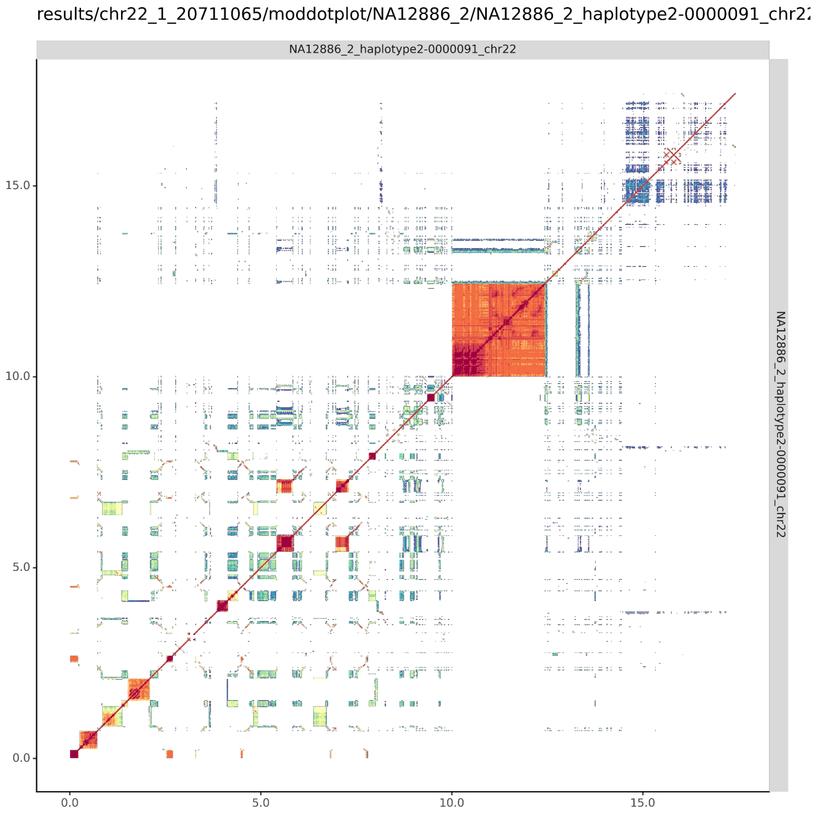

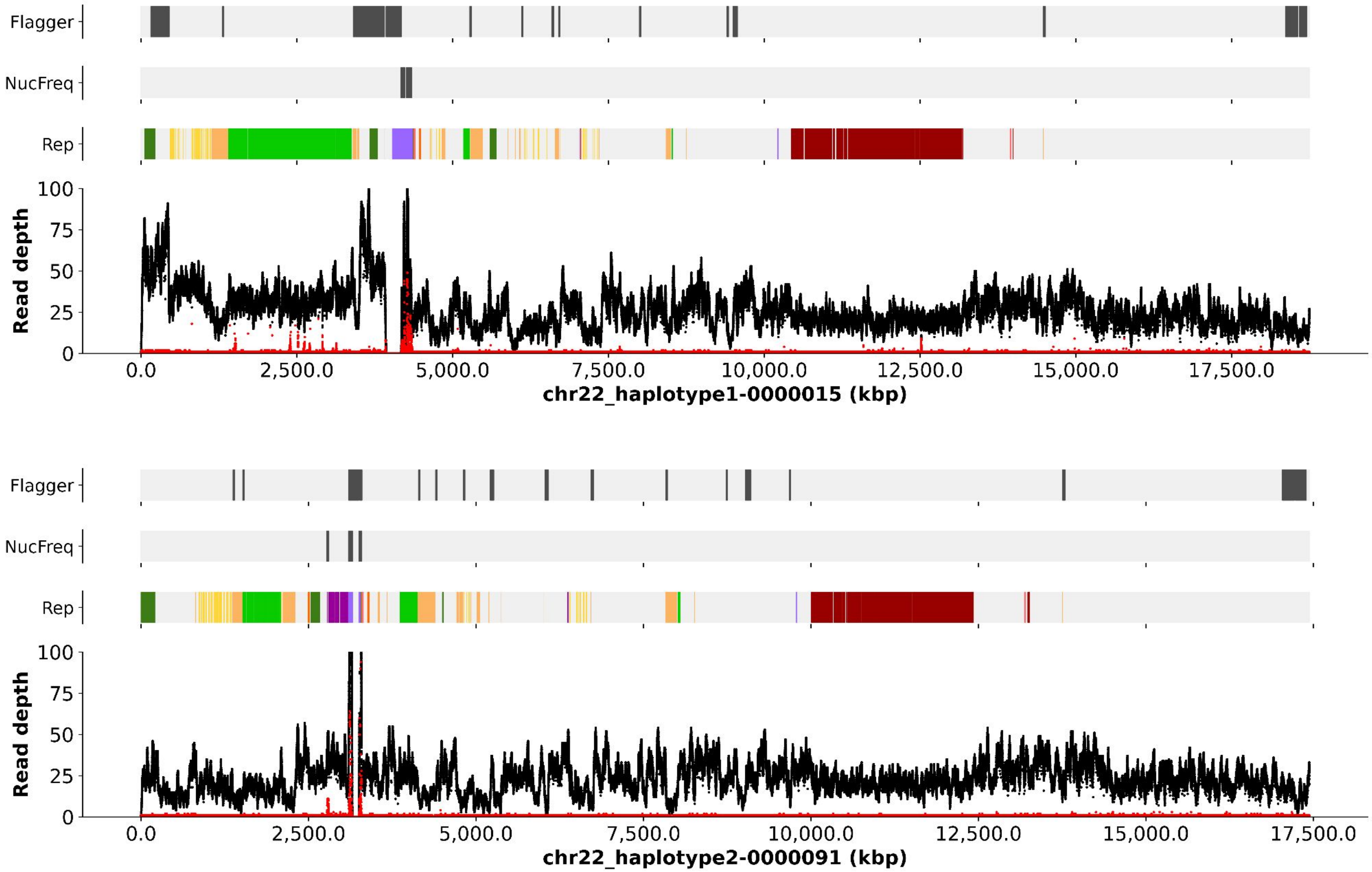
Flagger
NucFreq
Rep
Read depth
100
75
50
25
0
0.0
2,500.0
5,000.0
7,500.0
10,000.0
12,500.0
15,000.0
17,500.0
chr22_haplotype1-0000015 (kbp)
Flagger
NucFreq
Rep
Read depth
100
75
50
25
0
0.0
2,500.0
5,000.0
7,500.0
10,000.0
12,500.0
15,000.0
17,500.0
chr22_haplotype2-0000091 (kbp)

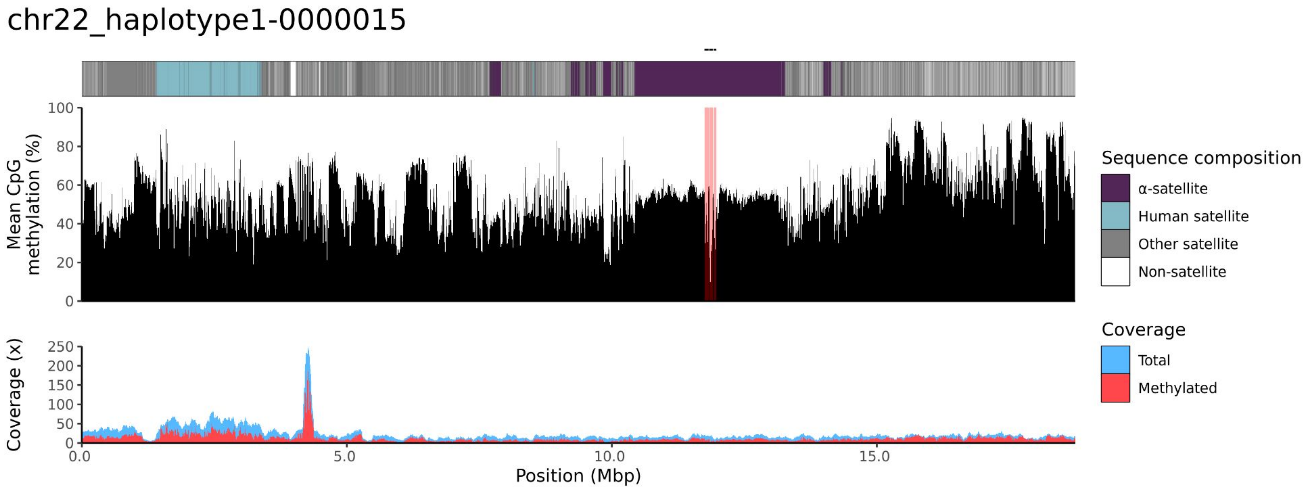
chr22_haplotype1-0000015
Mean CpG methylation (%)
100
80
60
40
20
0
Coverage (x)
250
200
150
100
50
0
0.0
5.0
10.0
15.0
Position (Mbp)
Sequence composition
α-satellite
Human satellite
Other satellite
Non-satellite
Coverage
Total
Methylated

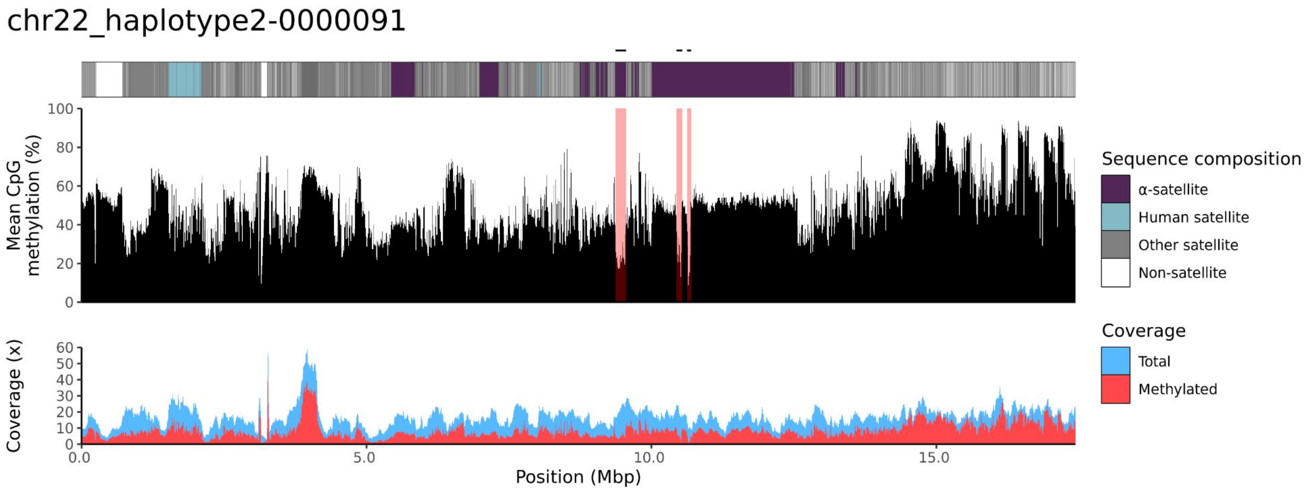
chr22_haplotype2-0000091
Mean CpG methylation (%)
100
80
60
40
20
0
Coverage (x)
60
50
40
30
20
10
0
0.0
5.0
10.0
15.0
Position (Mbp)
Sequence composition
α-satellite
Human satellite
Other satellite
Non-satellite
Coverage
Total
Methylated

NA12887

### chr13

#### NA12887_1_haplotype1-0000013_chr13

results/chr13_1_22508596/moddotplot/NA12887_1/NA12887_1_haplotype1-0000013_chr1:

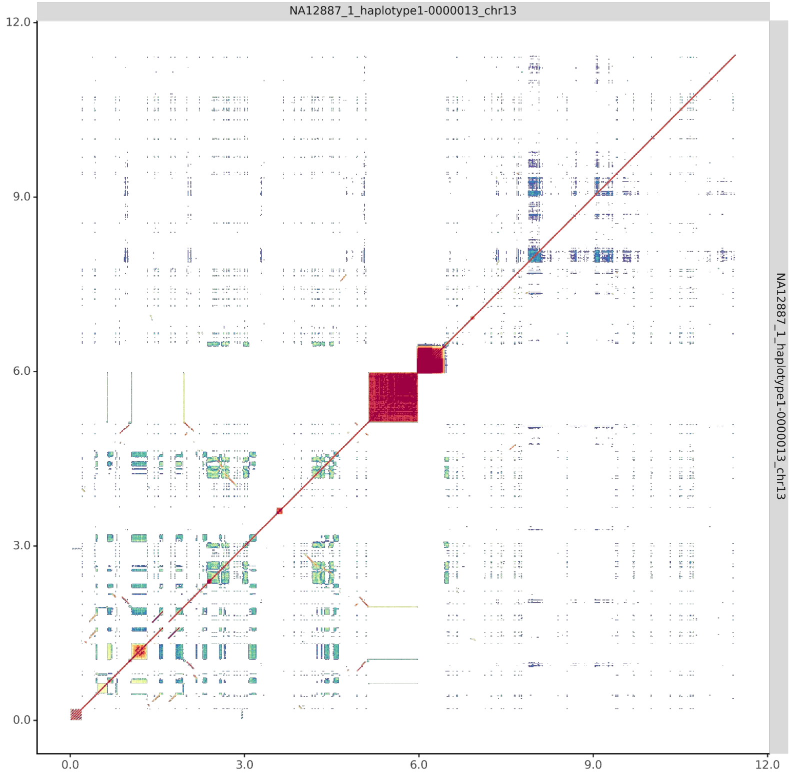


#### NA12887_2_haplotype2-0000076_chr13

results/chr13_1_22508596/moddotplot/NA12887_2/NA12887_2_haplotype2-0000076_chr1:

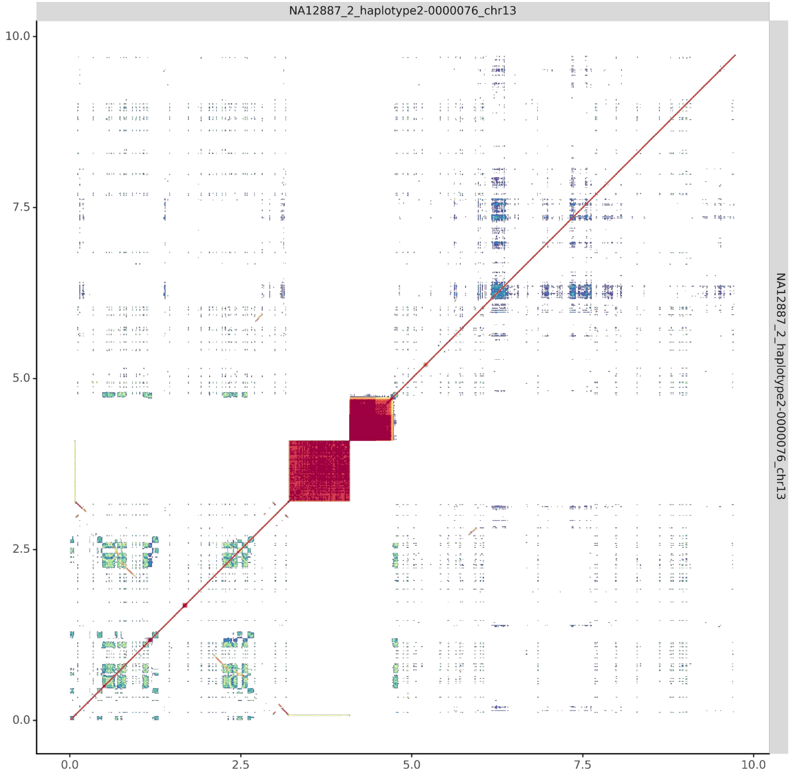

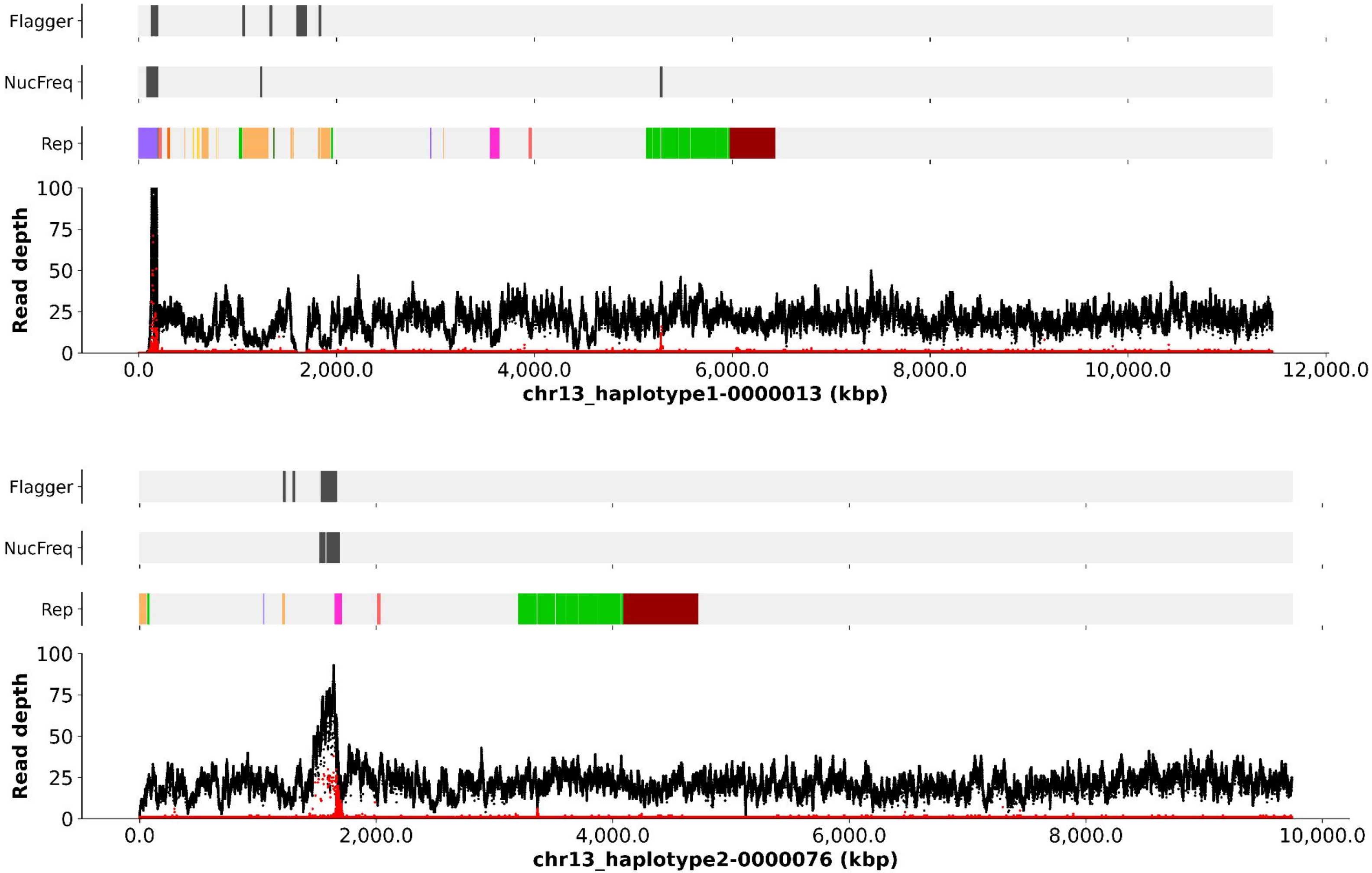
Flagger
NucFreq
Rep
Read depth
100
75
50
25
0
0.0
2,000.0
4,000.0
6,000.0
8,000.0
10,000.0
12,000.0
chr13_haplotype1-0000013 (kbp)
Flagger
NucFreq
Rep
Read depth
100
75
50
25
0
0.0
2,000.0
4,000.0
6,000.0
8,000.0
10,000.0
chr13_haplotype2-0000076 (kbp)

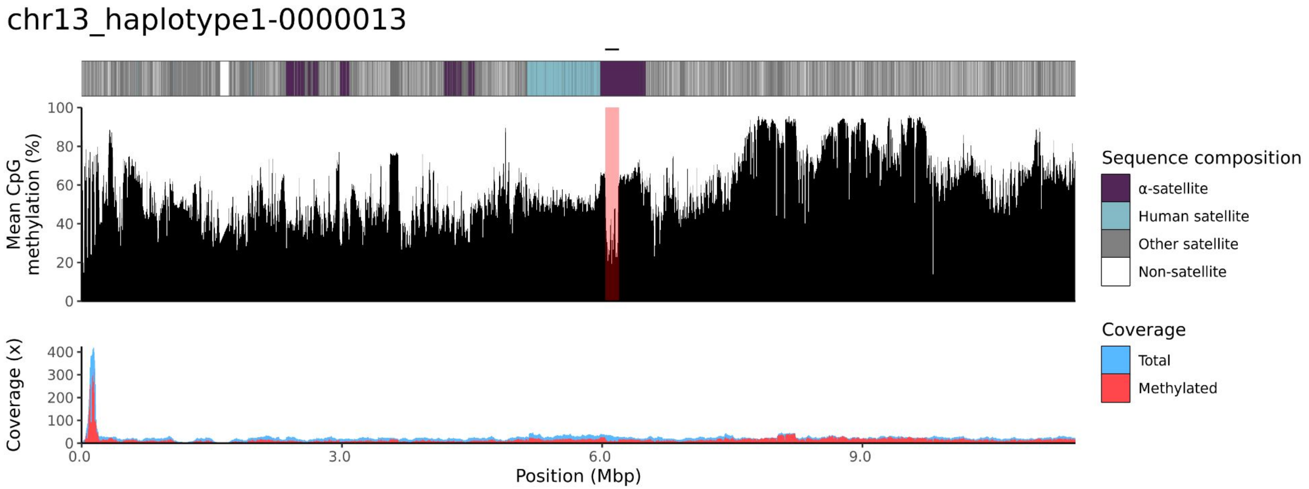
chr13_haplotype1-0000013
Mean CpG methylation (%)
100
80
60
40
20
0
Coverage (x)
400
300
200
100
0
0.0
3.0
6.0
9.0
Position (Mbp)
Sequence composition
α-satellite
Human satellite
Other satellite
Non-satellite
Coverage
Total
Methylated

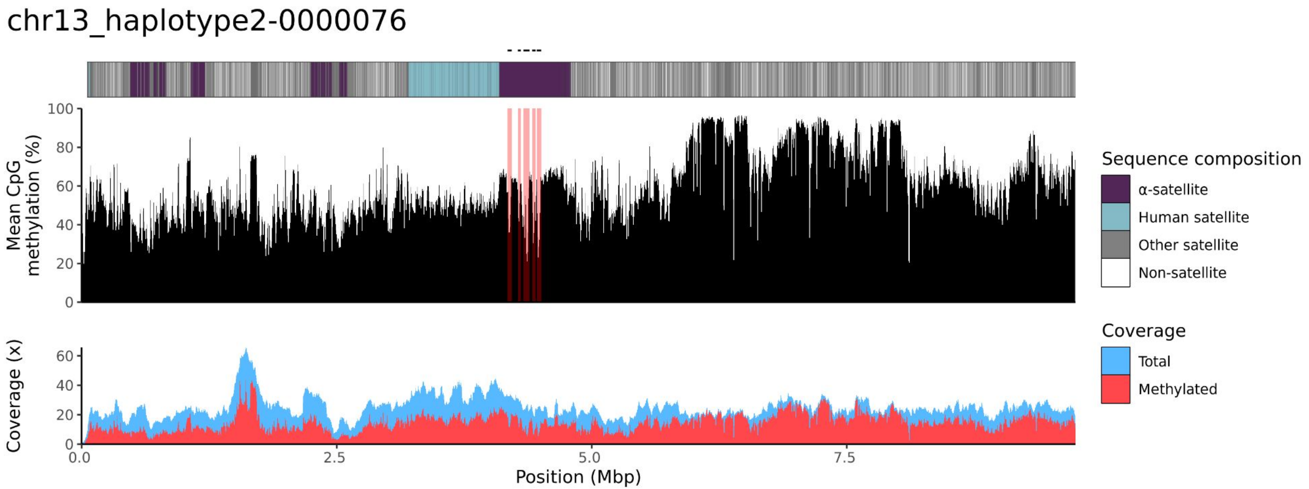
chr13_haplotype2-0000076
Mean CpG methylation (%)
100
80
60
40
20
0
Coverage (x)
60
40
20
0
0.0
2.5
5.0
7.5
Position (Mbp)
Sequence composition
α-satellite
Human satellite
Other satellite
Non-satellite
Coverage
Total
Methylated

### chr14

#### NA12887_1_haplotype1-0000019_chr14

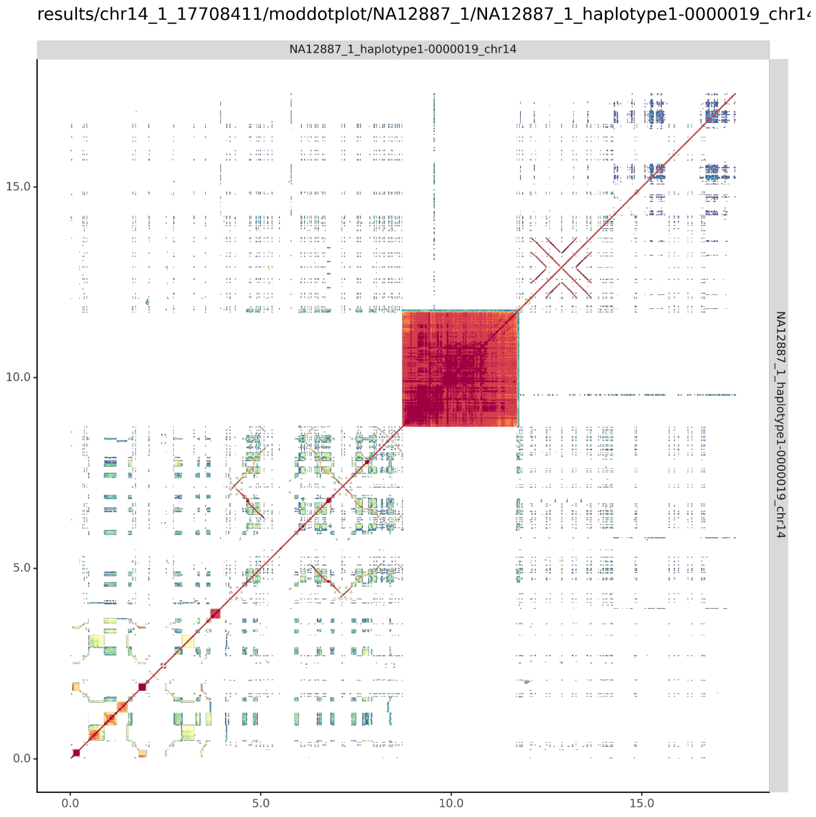


#### NA12887_2_haplotype2-0000093_chr14

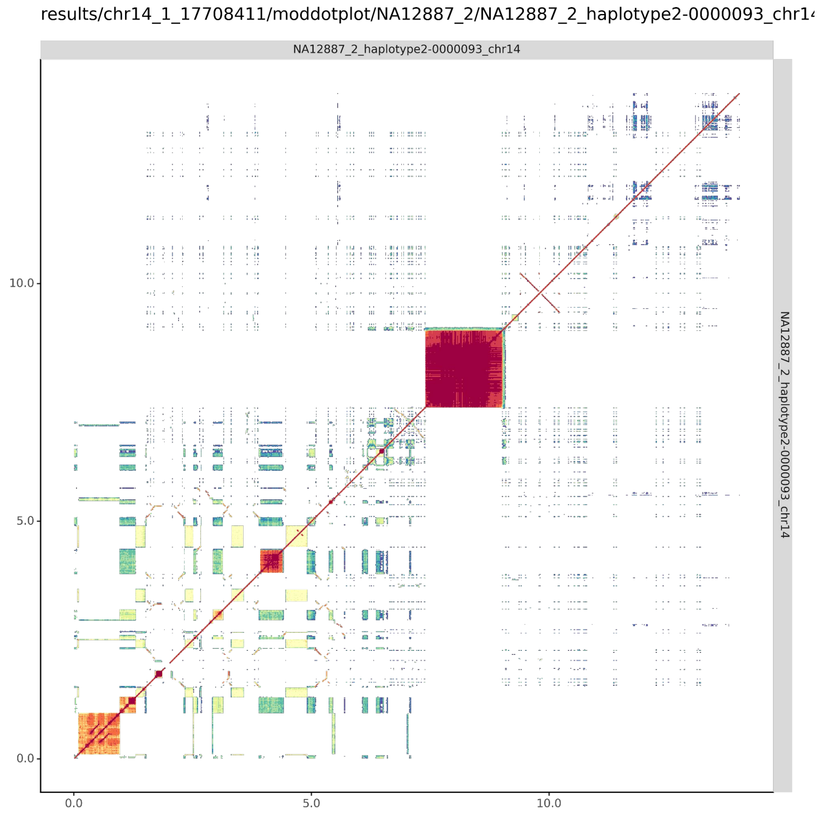

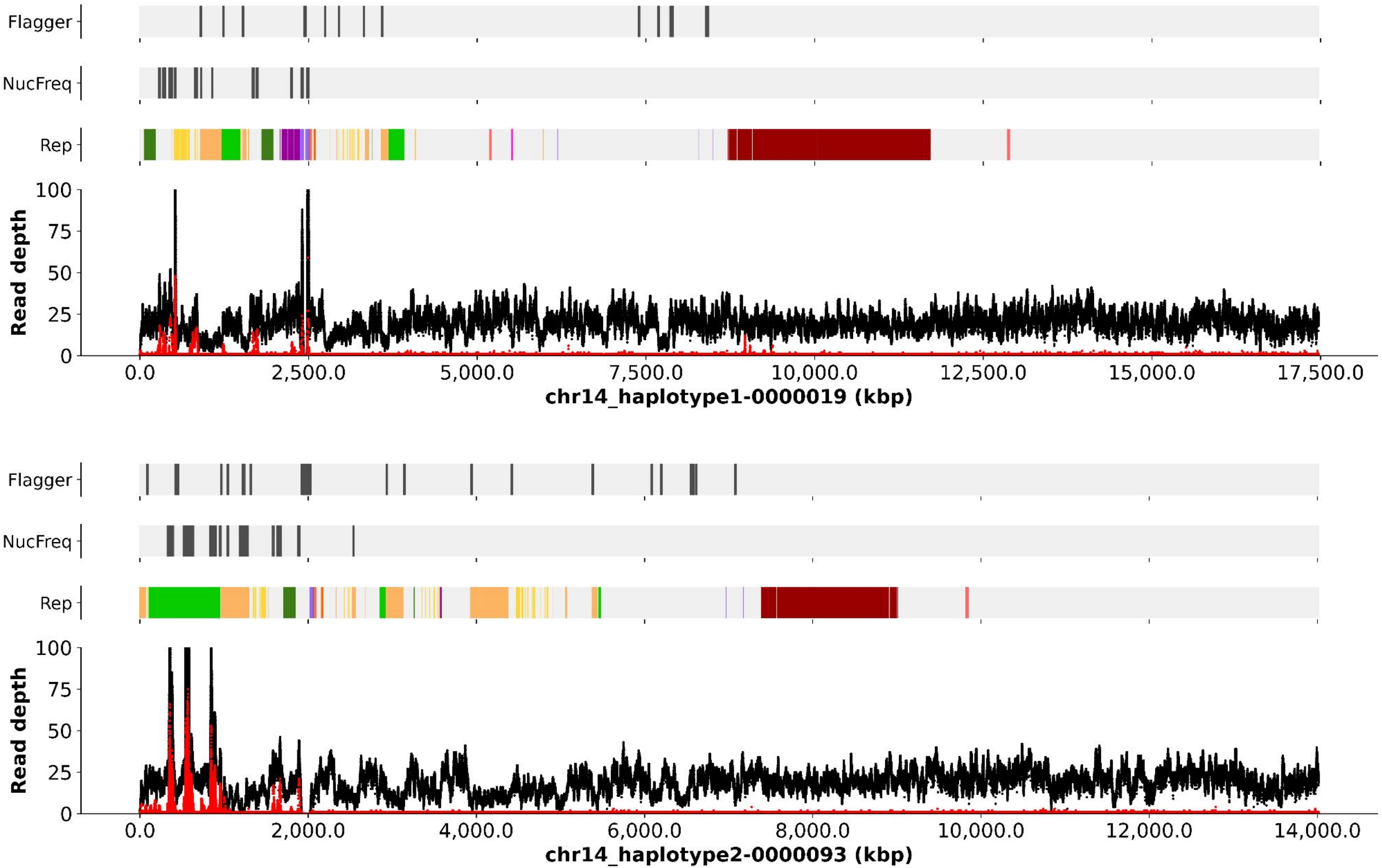

Flagger
NucFreq
Rep
Read depth
100
75
50
25
0
0.0
2,500.0
5,000.0
7,500.0
10,000.0
12,500.0
15,000.0
17,500.0
chr14_haplotype1-0000019 (kbp)
Flagger
NucFreq
Rep
Read depth
100
75
50
25
0
0.0
2,000.0
4,000.0
6,000.0
8,000.0
10,000.0
12,000.0
14,000.0
chr14_haplotype2-0000093 (kbp)

chr14_haplotype1-0000019

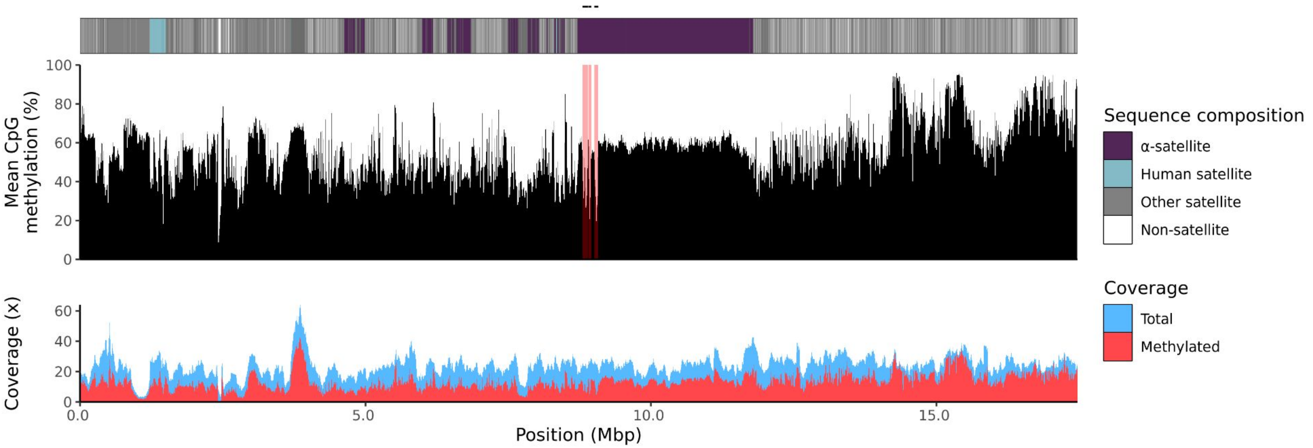


chr14_haplotype2-0000093

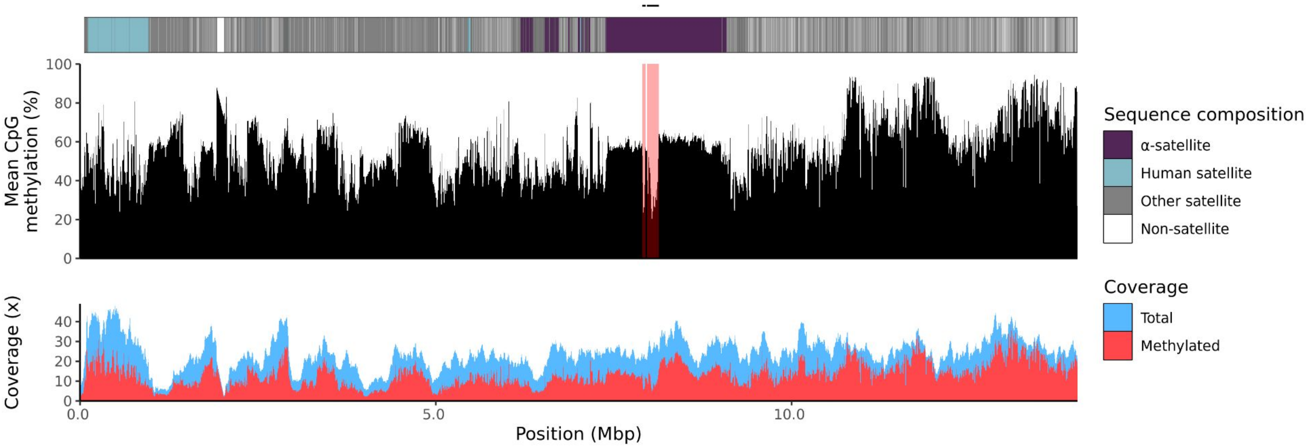

### chr15

#### NA12887_1_haplotype1-0000012_chr15

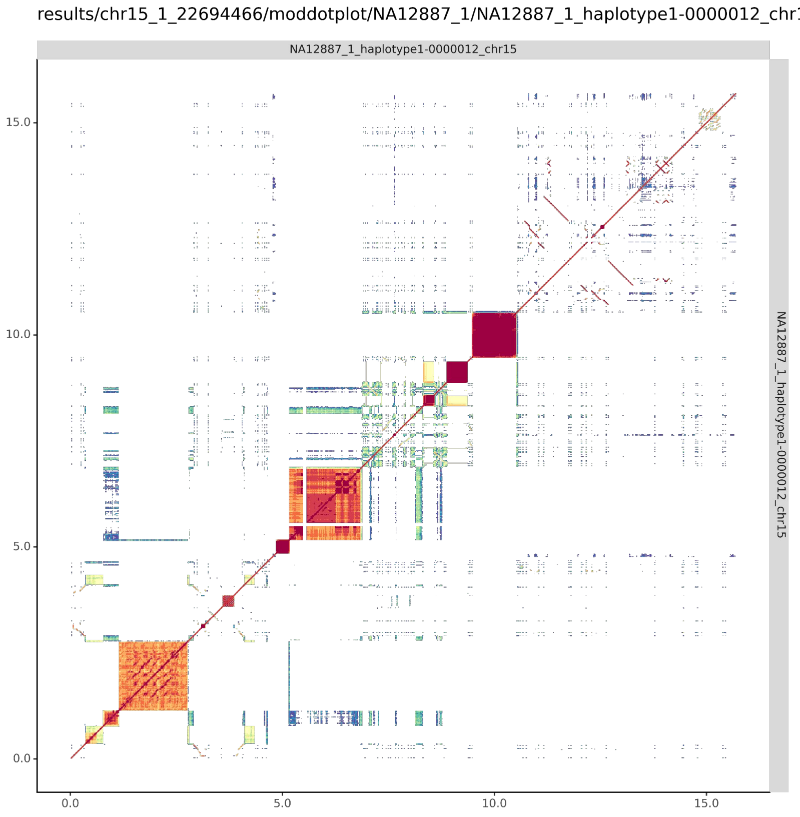


#### NA12887_2_haplotype2-0000070_chr15

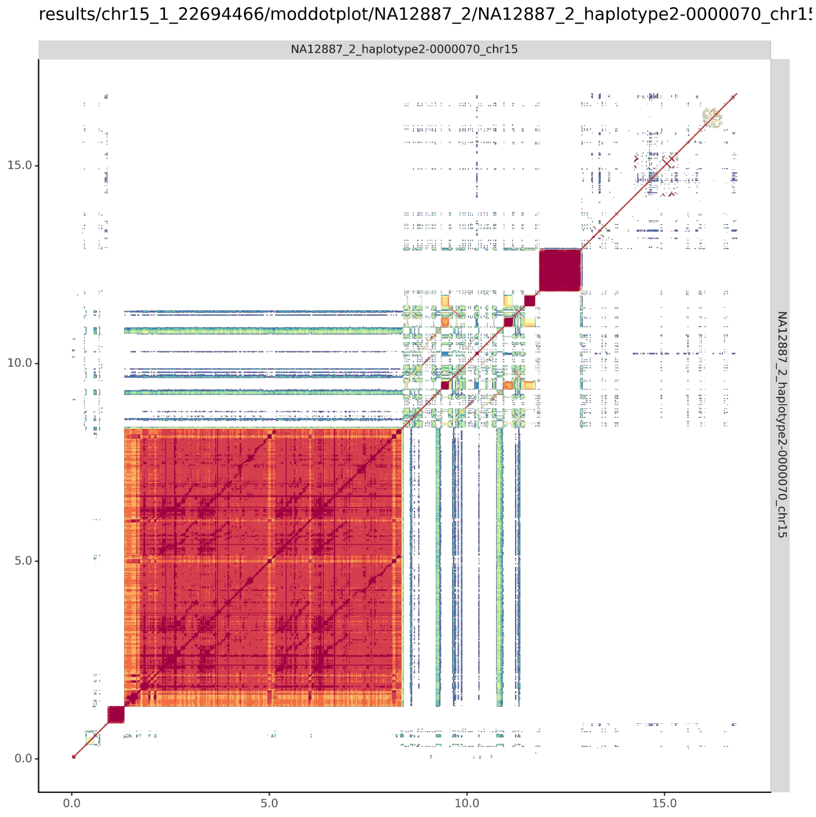

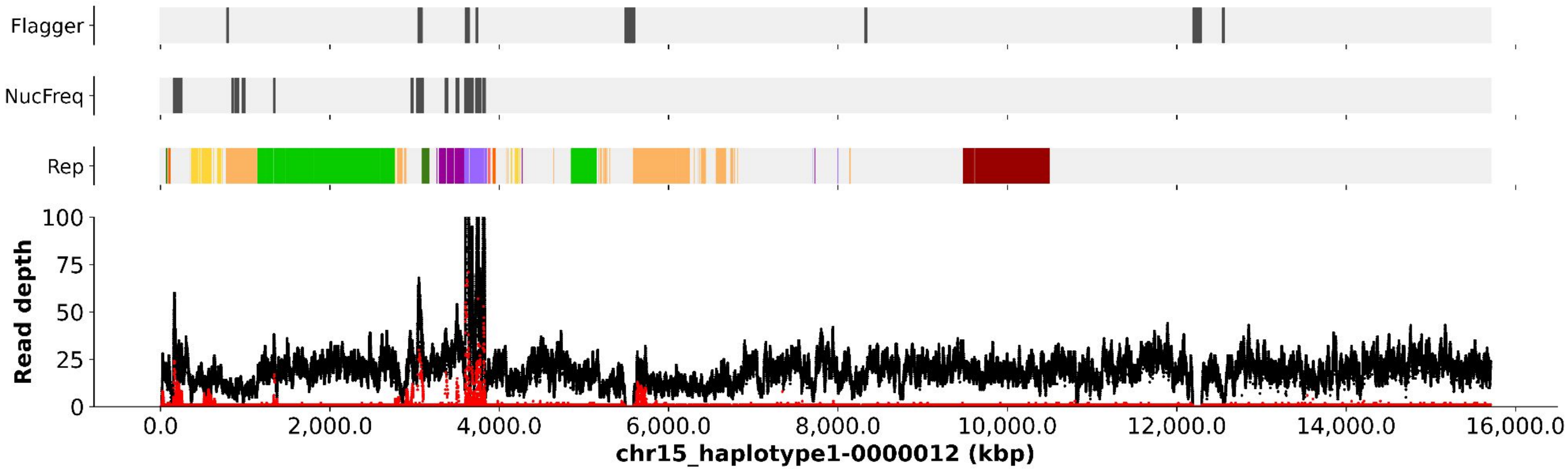
Flagger
NucFreq
Rep
Read depth
100
75
50
25
0
0.0
2,000.0
4,000.0
6,000.0
8,000.0
10,000.0
12,000.0
14,000.0
16,000.0
chr15_haplotype1-0000012 (kbp)

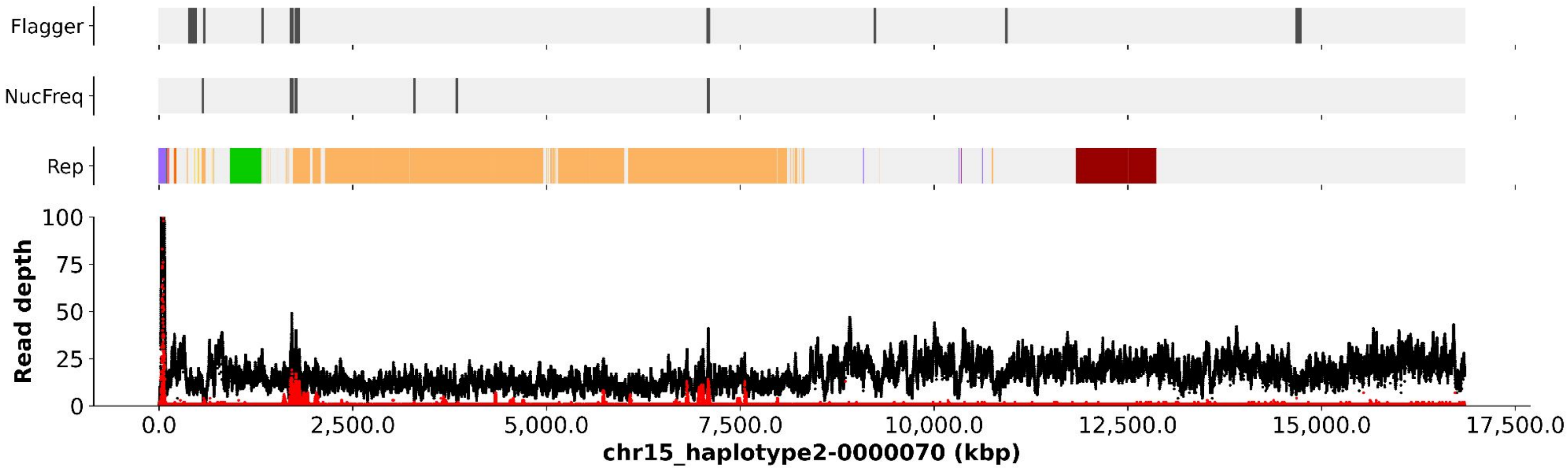
Flagger
NucFreq
Rep
Read depth
100
75
50
25
0
0.0
2,500.0
5,000.0
7,500.0
10,000.0
12,500.0
15,000.0
17,500.0
chr15_haplotype2-0000070 (kbp)

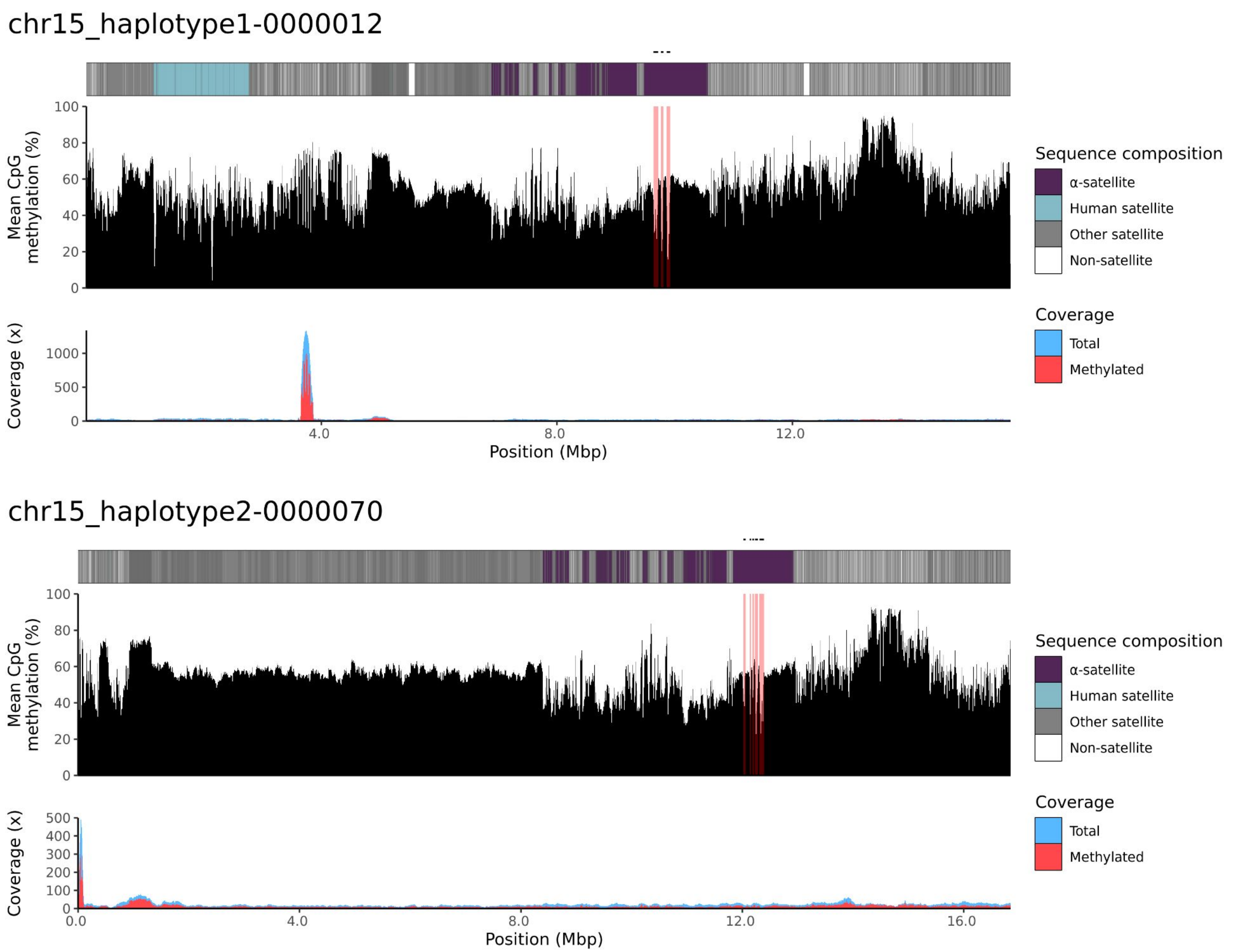
chr15_haplotype1-0000012
Mean CpG methylation (%)
100
80
60
40
20
0
Coverage (x)
1000
500
0
4.0
8.0
12.0
Position (Mbp)
Sequence composition
α-satellite
Human satellite
Other satellite
Non-satellite
Coverage
Total
Methylated
chr15_haplotype2-0000070
Mean CpG methylation (%)
100
80
60
40
20
0
Coverage (x)
500
400
300
200
100
0
0.0
4.0
8.0
12.0
16.0
Position (Mbp)
Sequence composition
α-satellite
Human satellite
Other satellite
Non-satellite
Coverage
Total
Methylated

### chr21

#### NA12887_1_haplotype1-0000006_chr21

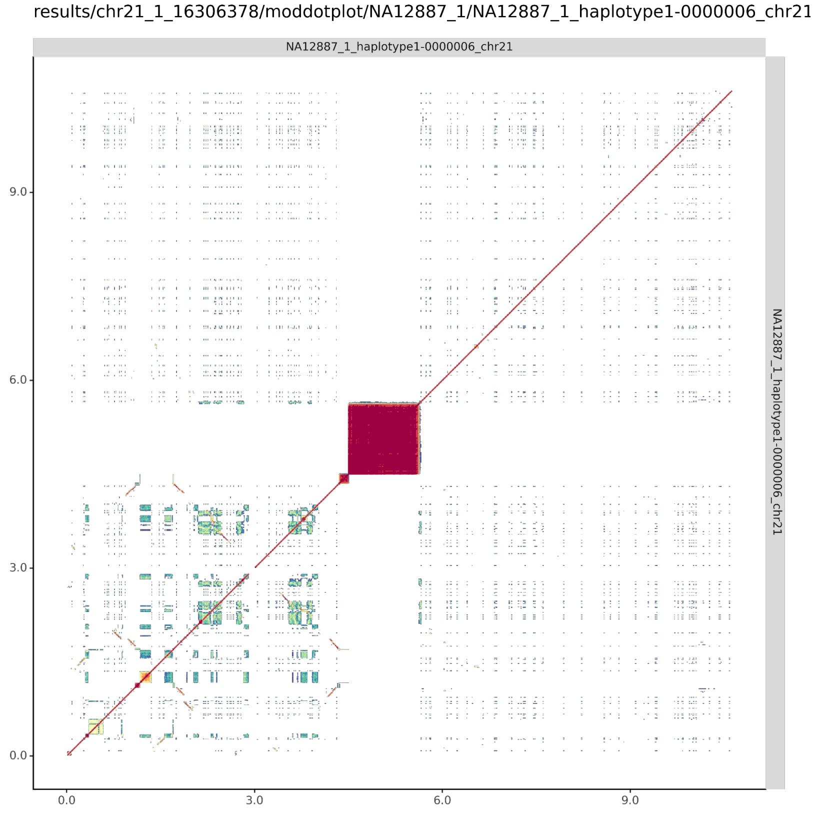


#### NA12887_2_haplotype2-0000065_chr21

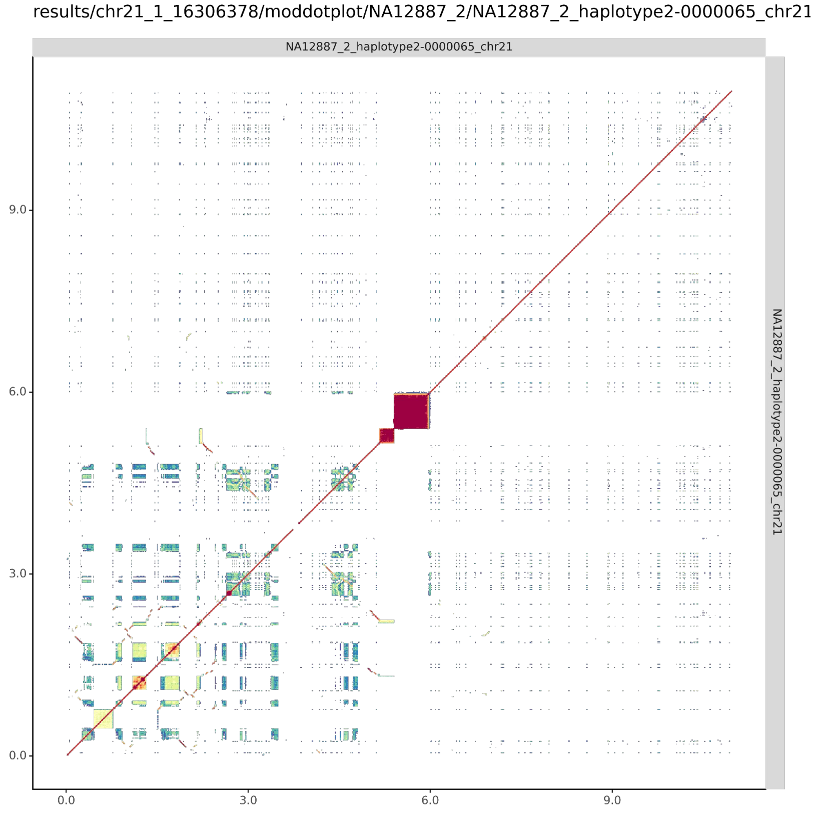

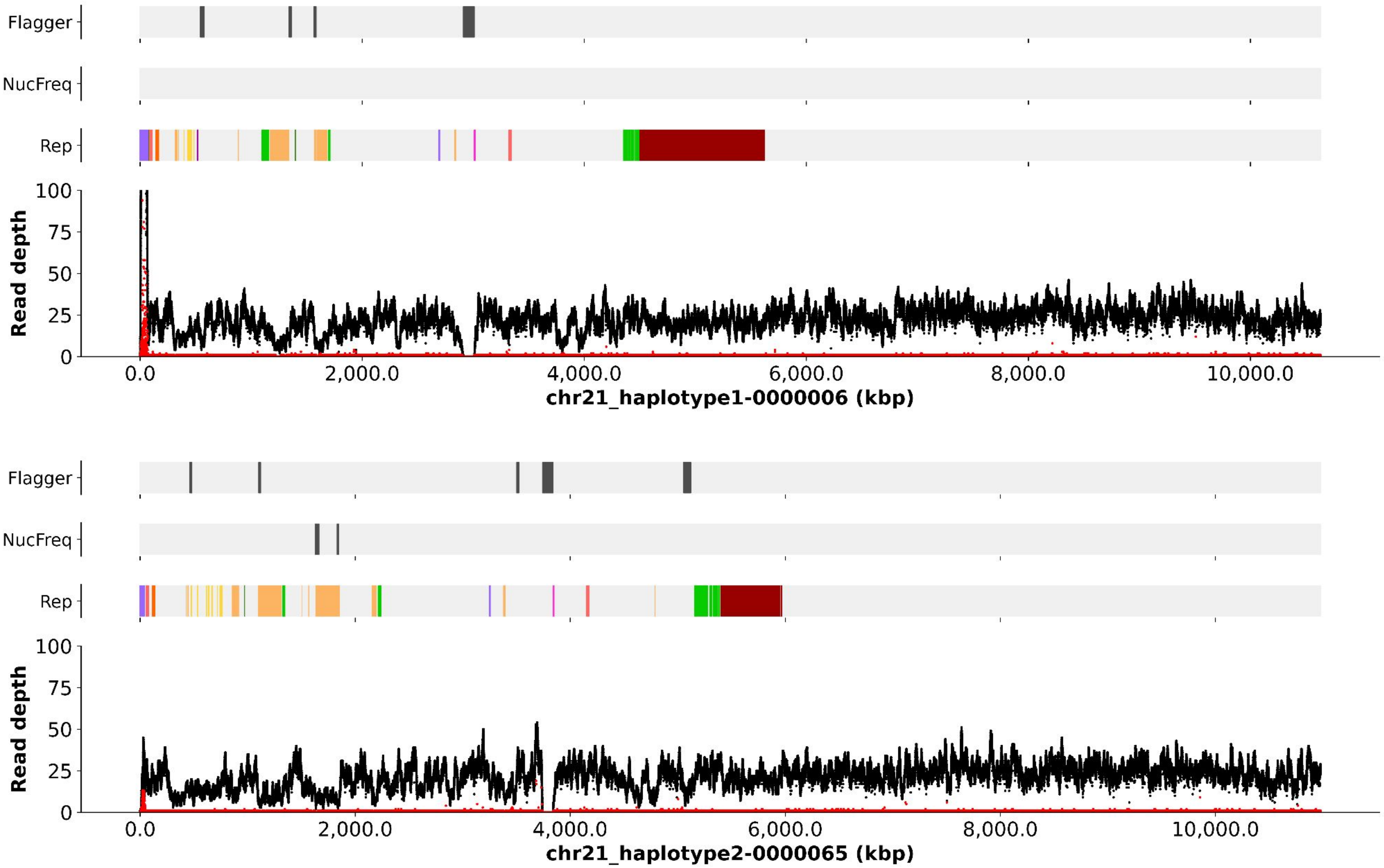
Flagger
NucFreq
Rep
Read depth
100
75
50
25
0
0.0
2,000.0
4,000.0
6,000.0
8,000.0
10,000.0
chr21_haplotype1-0000006 (kbp)
Flagger
NucFreq
Rep
Read depth
100
75
50
25
0
0.0
2,000.0
4,000.0
6,000.0
8,000.0
10,000.0
chr21_haplotype2-0000065 (kbp)

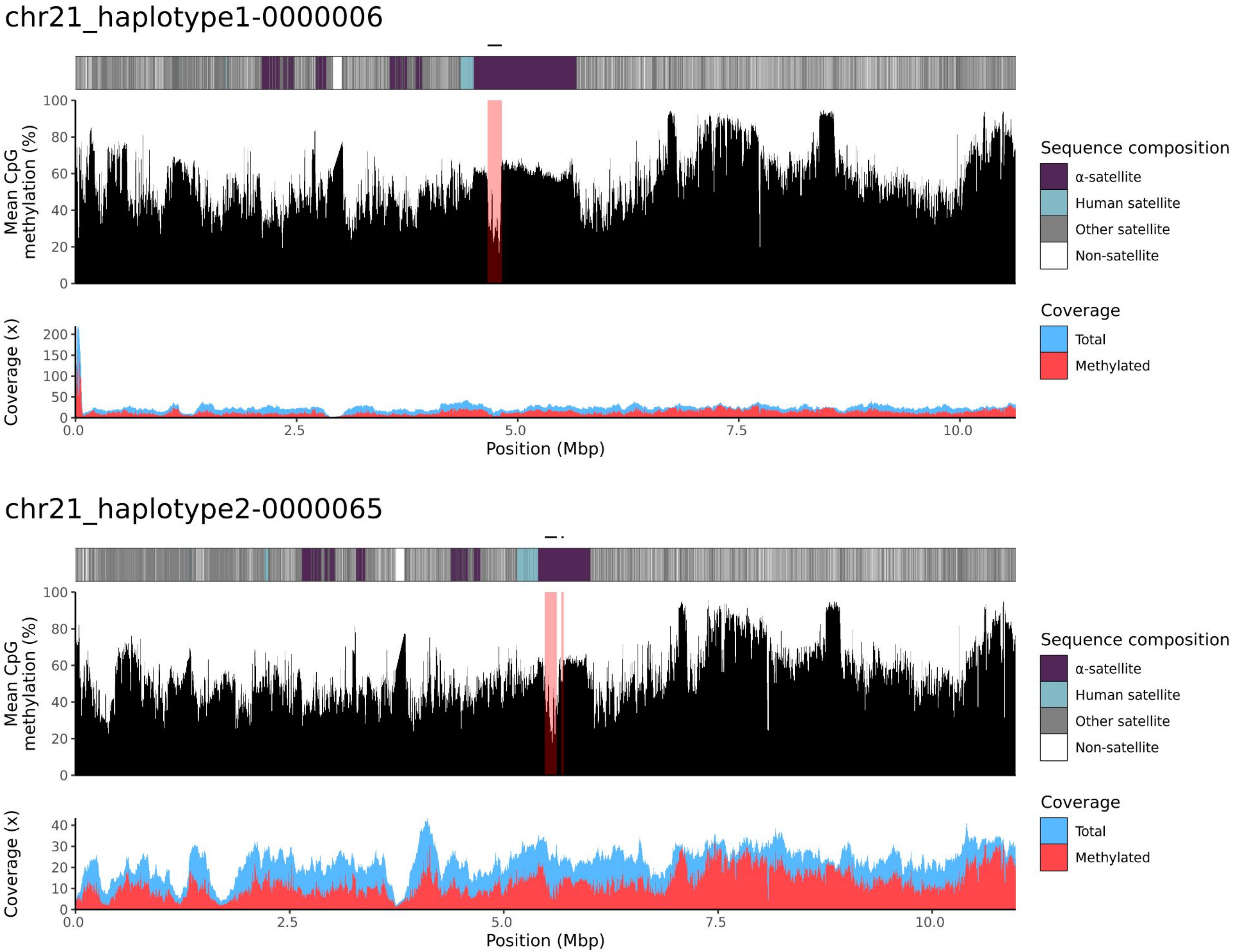
chr21_haplotype1-0000006
Mean CpG methylation (%)
Coverage (x)
Position (Mbp)
Sequence composition
α-satellite
Human satellite
Other satellite
Non-satellite
Coverage
Total
Methylated
chr21_haplotype2-0000065
Mean CpG methylation (%)
Coverage (x)
Position (Mbp)
Sequence composition
α-satellite
Human satellite
Other satellite
Non-satellite
Coverage
Total
Methylated

### chr22

#### NA12887_1_haplotype1-0000021_chr22

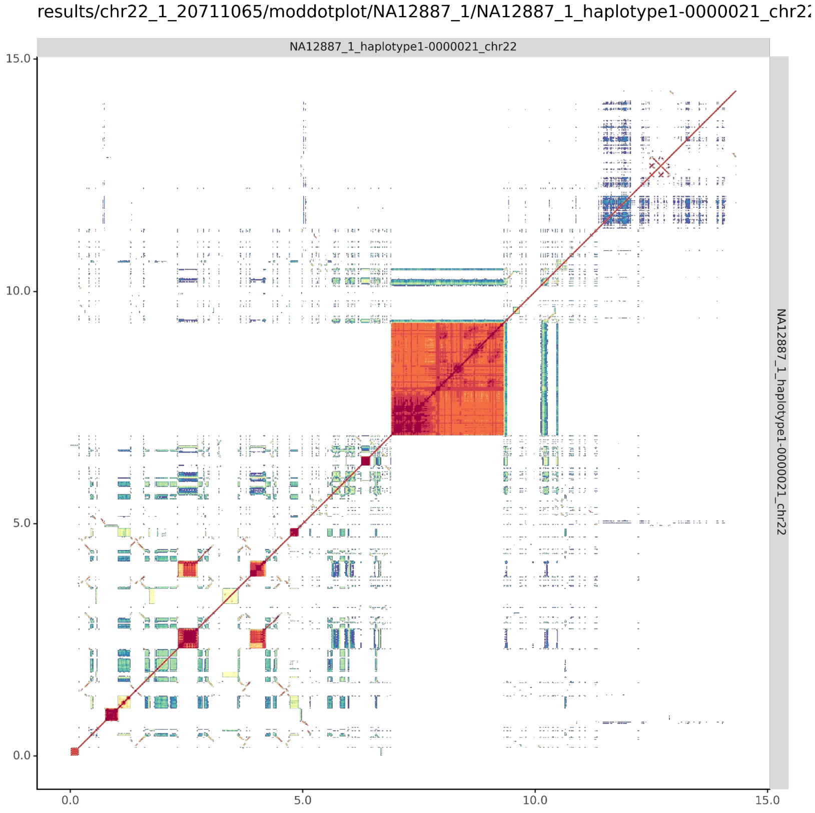


#### NA12887_2_haplotype2-0000080_chr22

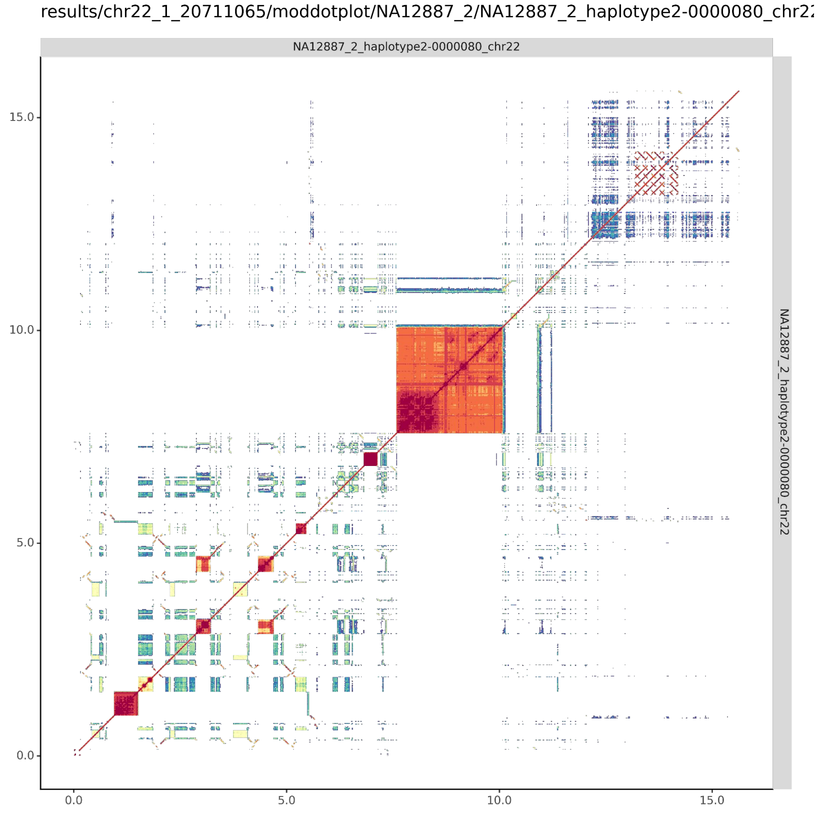

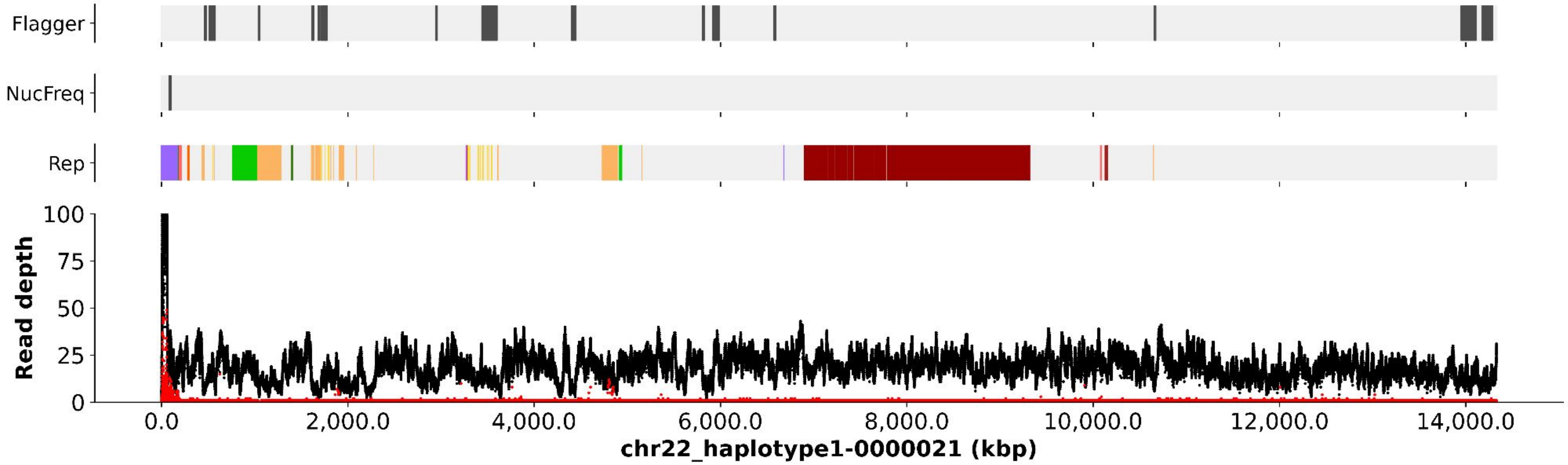
Flagger
NucFreq
Rep
Read depth
100
75
50
25
0
0.0
2,000.0
4,000.0
6,000.0
8,000.0
10,000.0
12,000.0
14,000.0
chr22_haplotype1-0000021 (kbp)

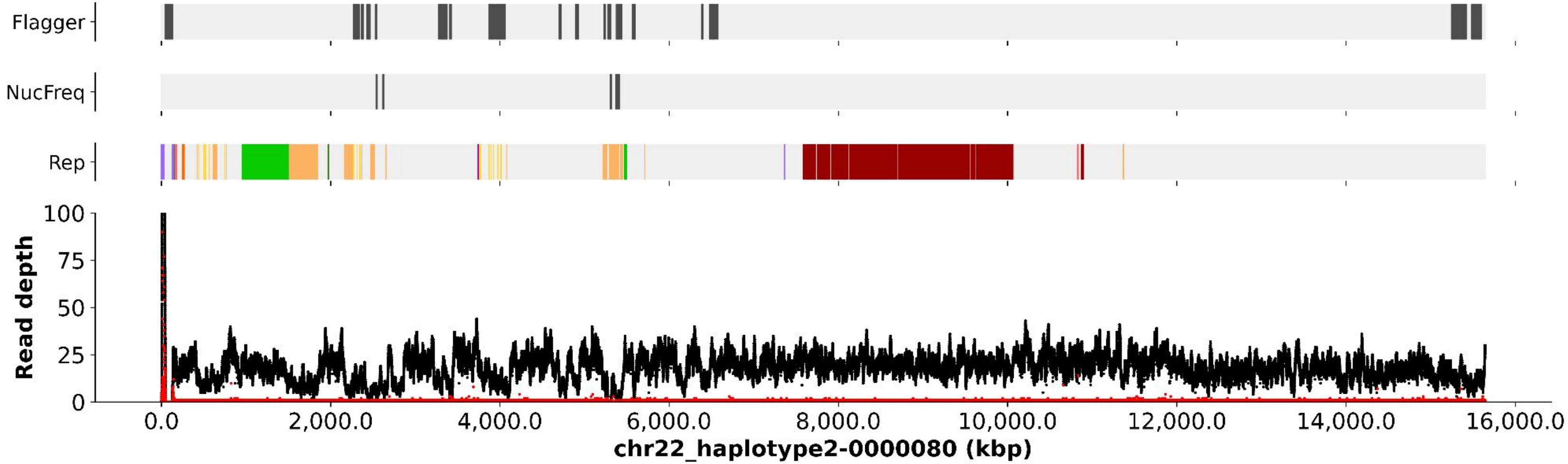
Flagger
NucFreq
Rep
Read depth
100
75
50
25
0
0.0
2,000.0
4,000.0
6,000.0
8,000.0
10,000.0
12,000.0
14,000.0
16,000.0
chr22_haplotype2-0000080 (kbp)

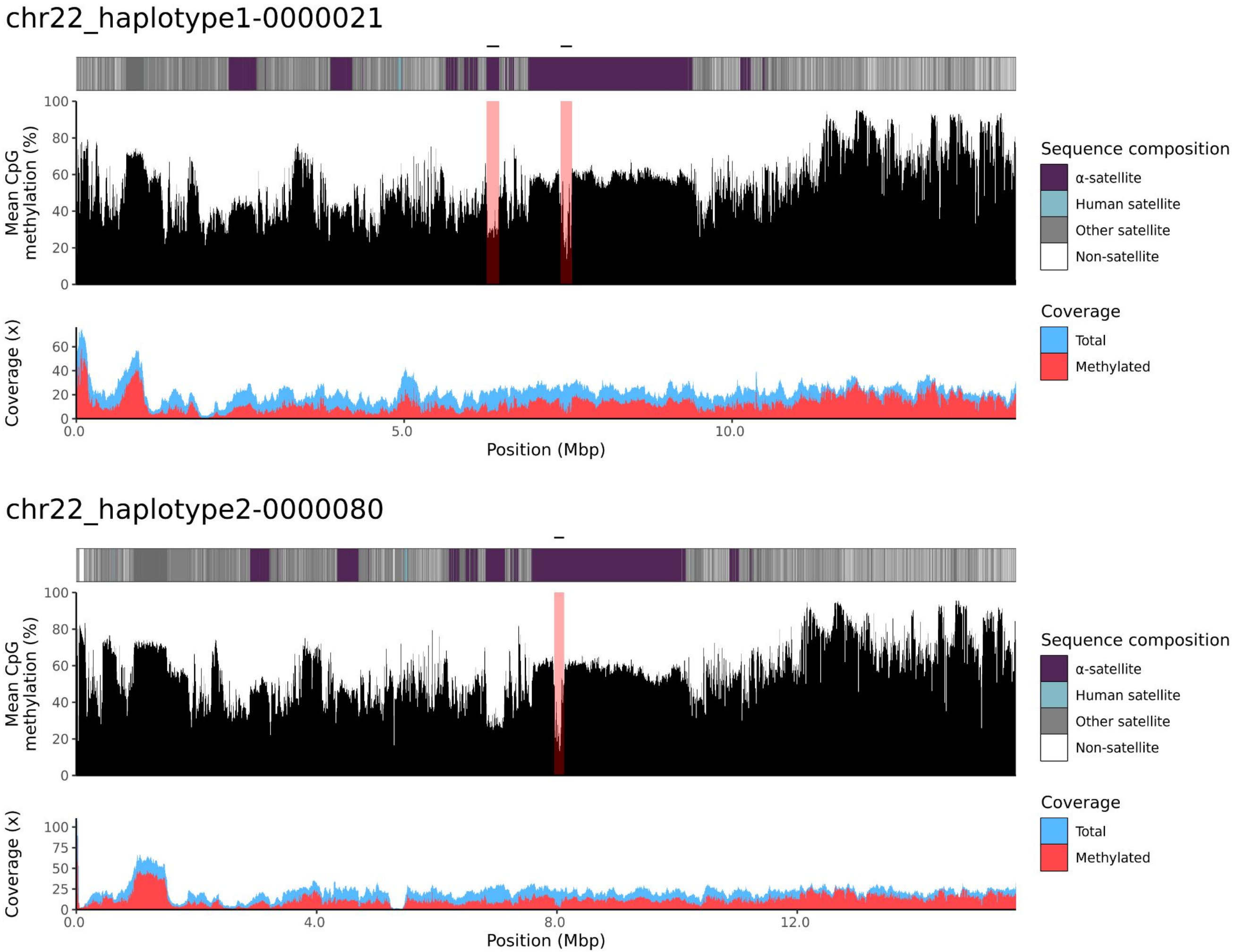
chr22_haplotype1-0000021
Mean CpG methylation (%)
Coverage (x)
Position (Mbp)
Sequence composition
α-satellite
Human satellite
Other satellite
Non-satellite
Coverage
Total
Methylated
chr22_haplotype2-0000080
Mean CpG methylation (%)
Coverage (x)
Position (Mbp)
Sequence composition
α-satellite
Human satellite
Other satellite
Non-satellite
Coverage
Total
Methylated

200080

### chr13

#### K200080_1_haplotype1-0000024_chr13

results/chr13_1_22508596/moddotplot/K200080_1/K200080_1_haplotype1-0000024_chr13

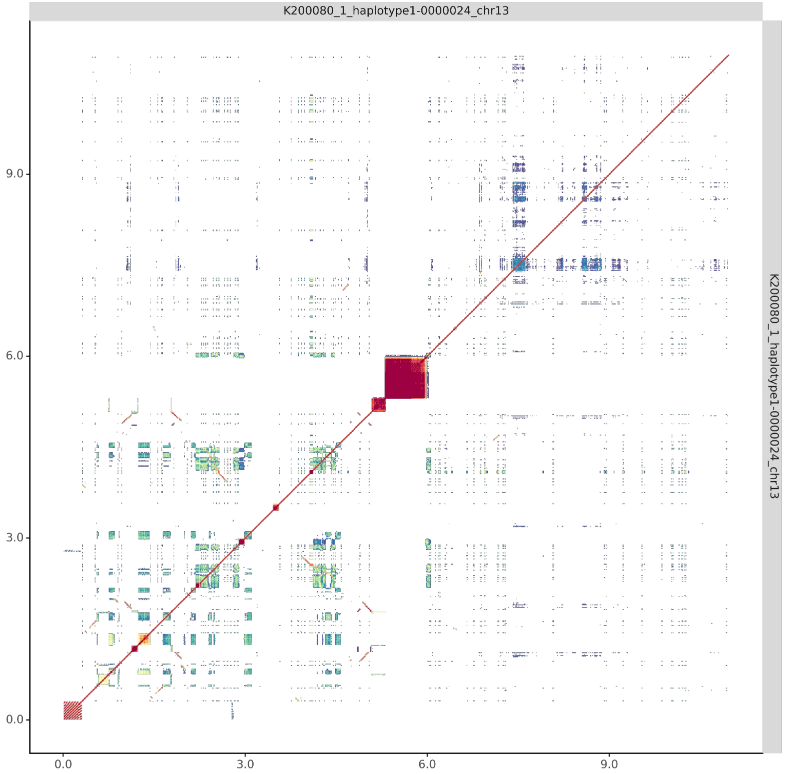


#### K200080_2_haplotype2-0000075_chr13

results/chr13_1_22508596/moddotplot/K200080_2/K200080_2_haplotype2-0000075_chr13

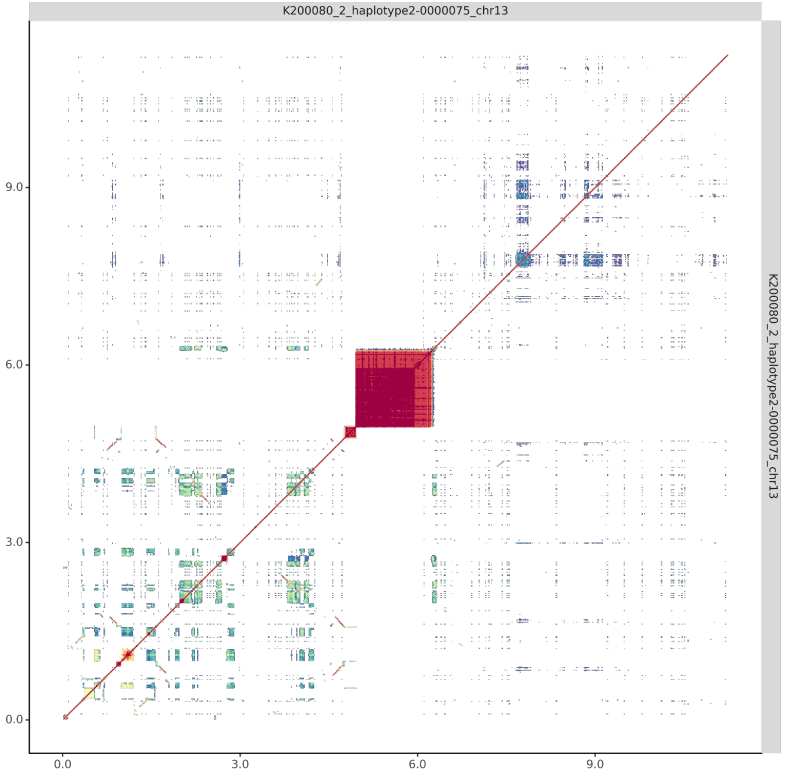

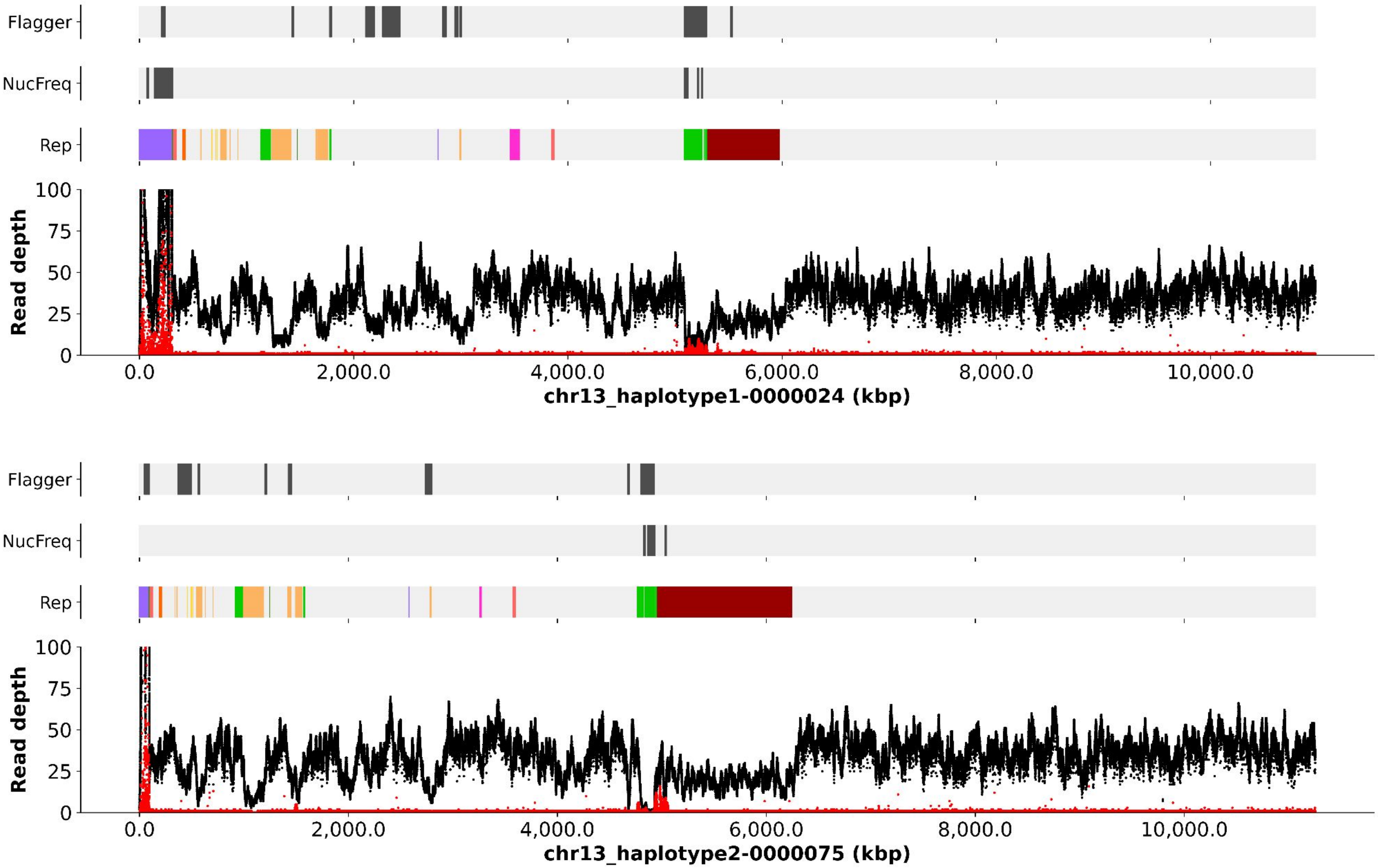
Flagger
NucFreq
Rep
Read depth
100
75
50
25
0
0.0
2,000.0
4,000.0
6,000.0
8,000.0
10,000.0
chr13_haplotype1-0000024 (kbp)
Flagger
NucFreq
Rep
Read depth
100
75
50
25
0
0.0
2,000.0
4,000.0
6,000.0
8,000.0
10,000.0
chr13_haplotype2-0000075 (kbp)

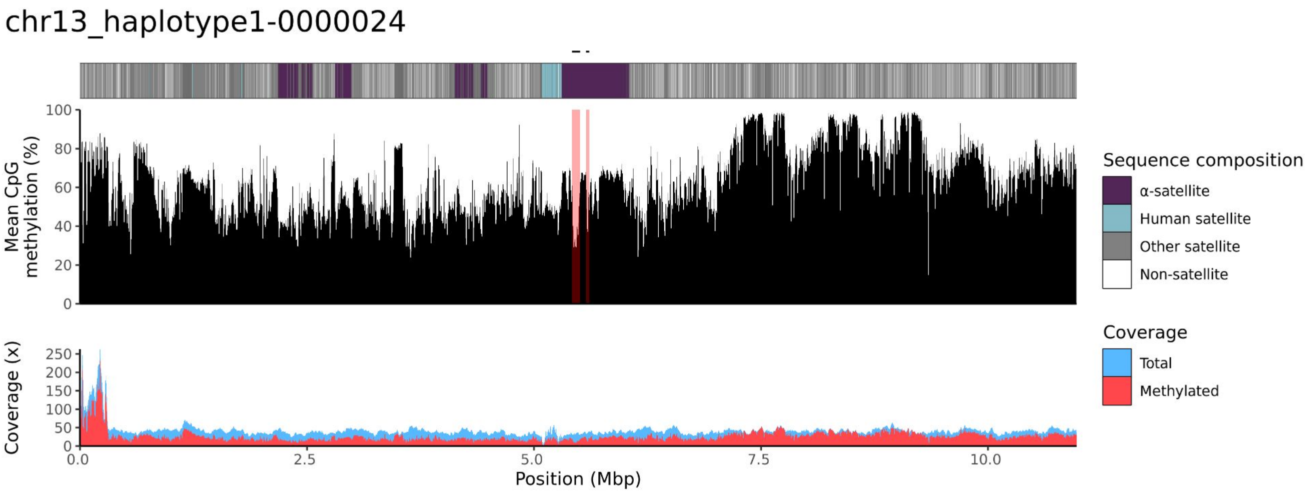
chr13_haplotype1-0000024
Mean CpG methylation (%)
100
80
60
40
20
0
Sequence composition
α-satellite
Human satellite
Other satellite
Non-satellite
Coverage (x)
250
200
150
100
50
0
0.0
2.5
5.0
7.5
10.0
Position (Mbp)
Coverage
Total
Methylated

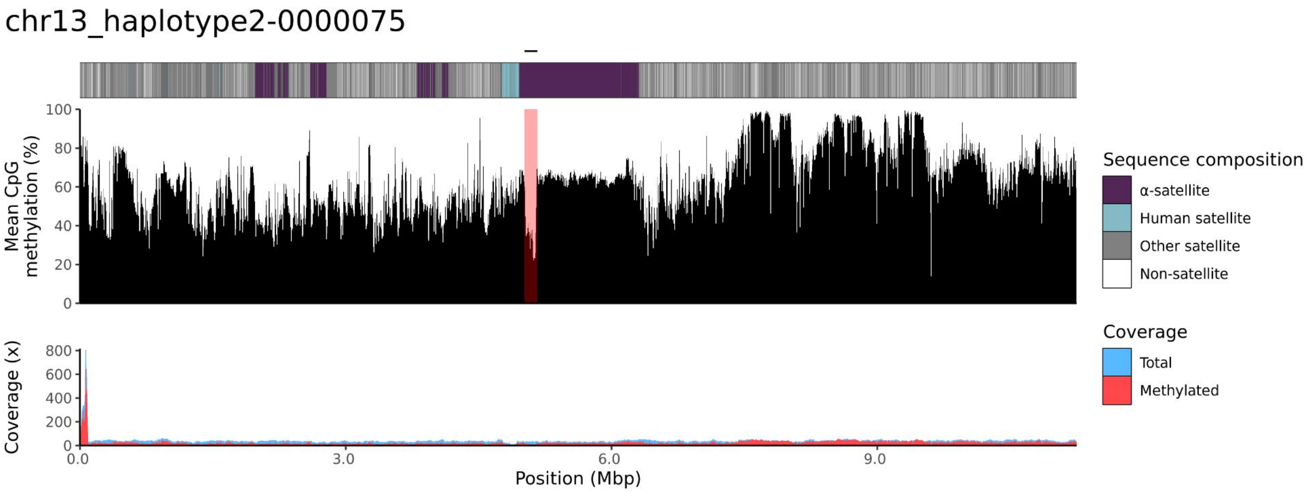
chr13_haplotype2-0000075
Mean CpG methylation (%)
100
80
60
40
20
0
Sequence composition
α-satellite
Human satellite
Other satellite
Non-satellite
Coverage (x)
800
600
400
200
0
0.0
3.0
6.0
9.0
Position (Mbp)
Coverage
Total
Methylated

### chr14

#### K200080_1_haplotype1-0000006_chr14

results/chr14_1_17708411/moddotplot/K200080_1/K200080_1_haplotype1-0000006_chr14

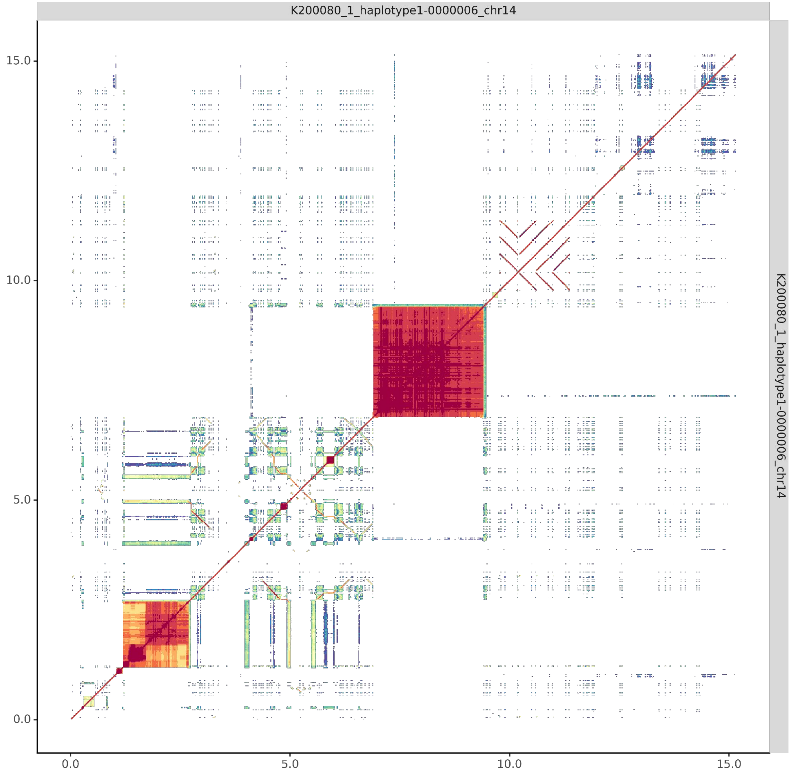


#### K200080_2_haplotype2-0000055_chr14

results/chr14_1_17708411/moddotplot/K200080_2/K200080_2_haplotype2-0000055_chr14

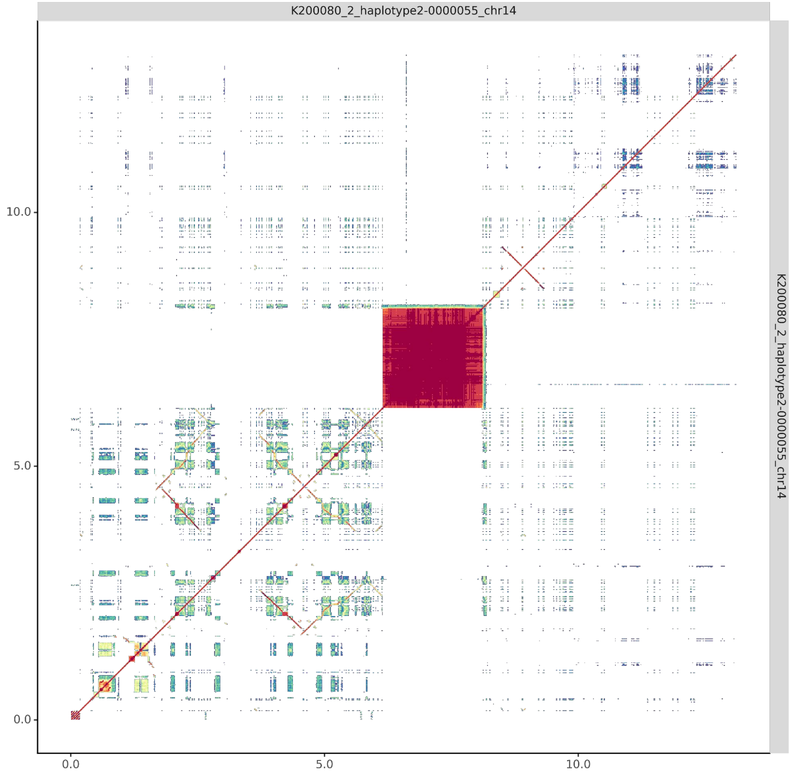

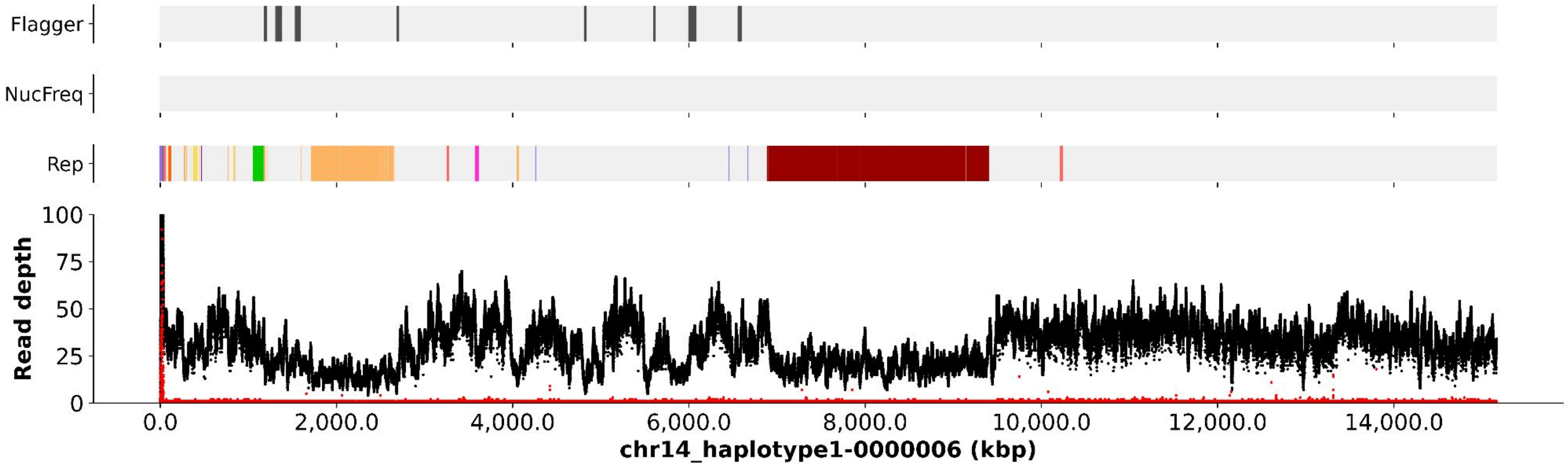
Flagger
NucFreq
Rep
Read depth
100
75
50
25
0
0.0
2,000.0
4,000.0
6,000.0
8,000.0
10,000.0
12,000.0
14,000.0
chr14_haplotype1-0000006 (kbp)

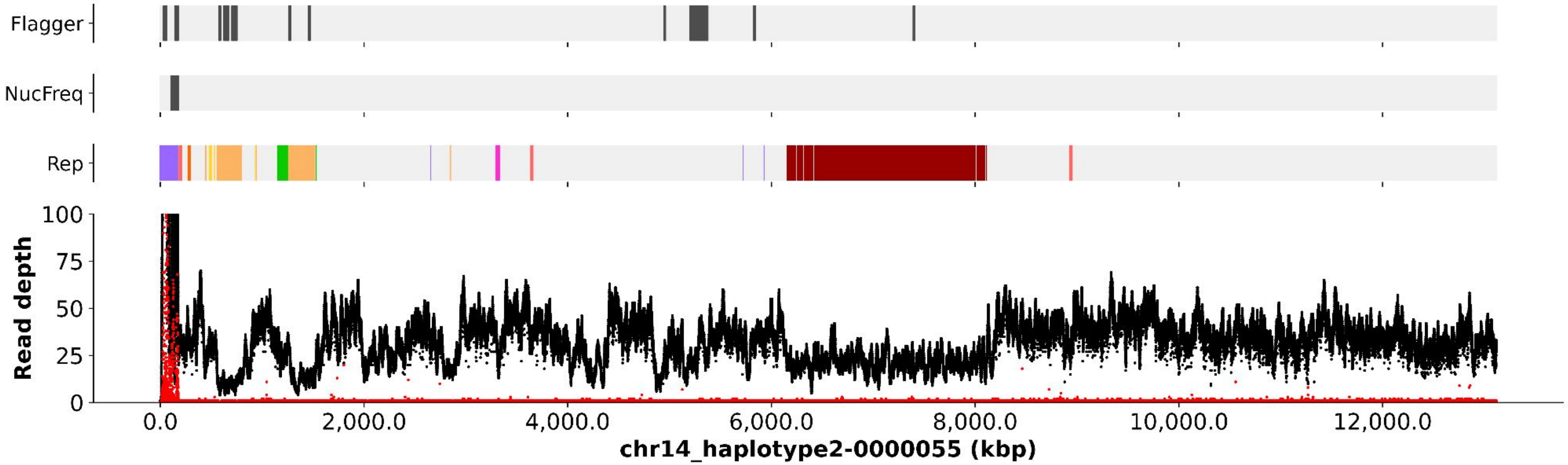
Flagger
NucFreq
Rep
Read depth
100
75
50
25
0
0.0
2,000.0
4,000.0
6,000.0
8,000.0
10,000.0
12,000.0
chr14_haplotype2-0000055 (kbp)

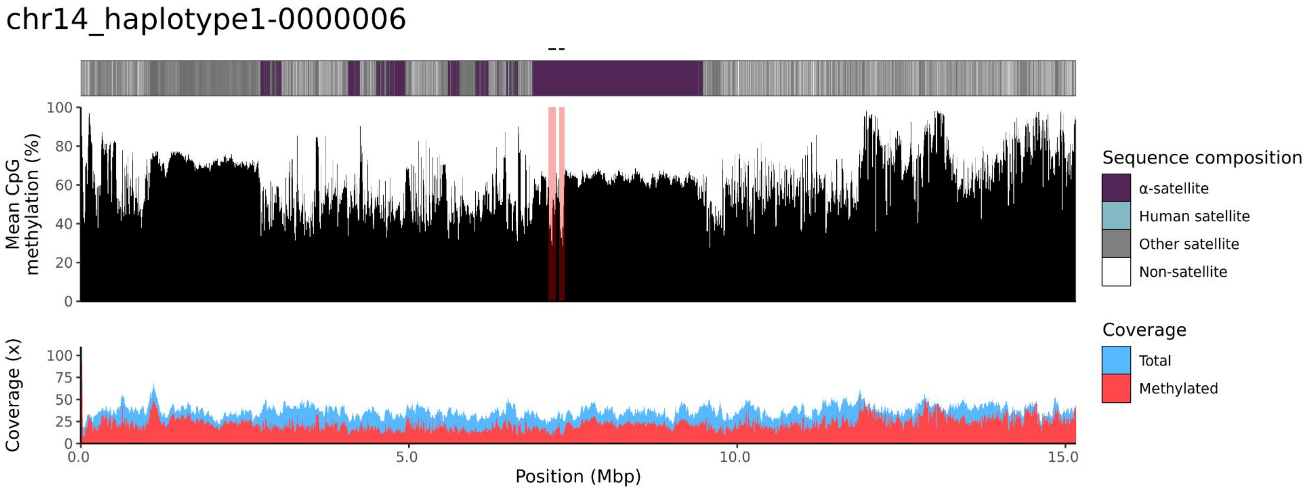
chr14_haplotype1-0000006
Mean CpG methylation (%)
100
80
60
40
20
0
Coverage (x)
100
75
50
25
0
0.0
5.0
10.0
15.0
Position (Mbp)
Sequence composition
α-satellite
Human satellite
Other satellite
Non-satellite
Coverage
Total
Methylated

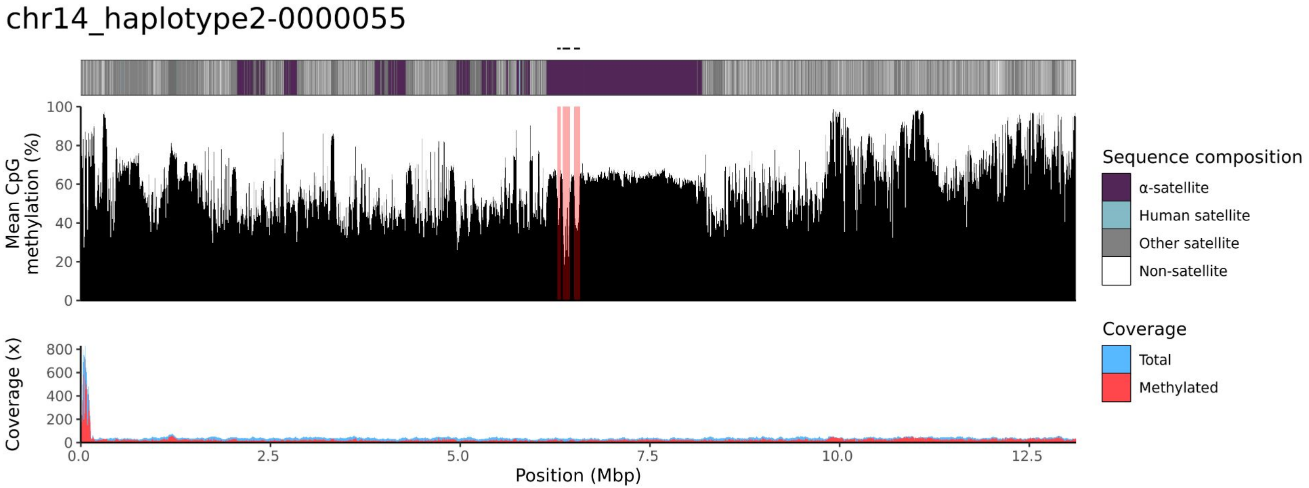
chr14_haplotype2-0000055
Mean CpG methylation (%)
100
80
60
40
20
0
Coverage (x)
800
600
400
200
0
0.0
2.5
5.0
7.5
10.0
12.5
Position (Mbp)
Sequence composition
α-satellite
Human satellite
Other satellite
Non-satellite
Coverage
Total
Methylated

### chr15

#### K200080_1_haplotype1-0000026_chr15

#### K200080_2_haplotype2-0000071_chr15

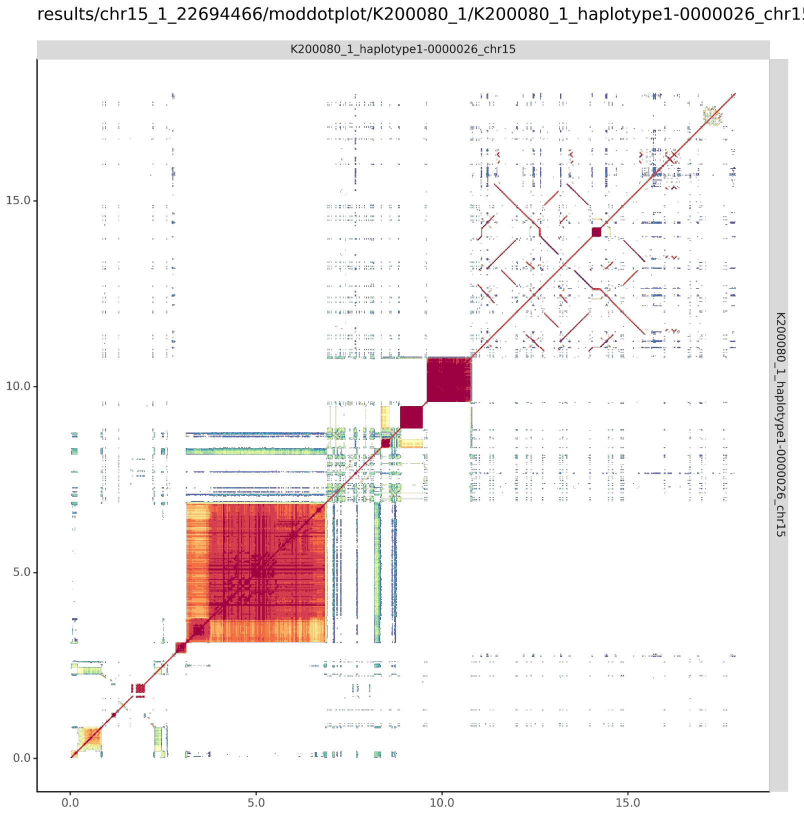


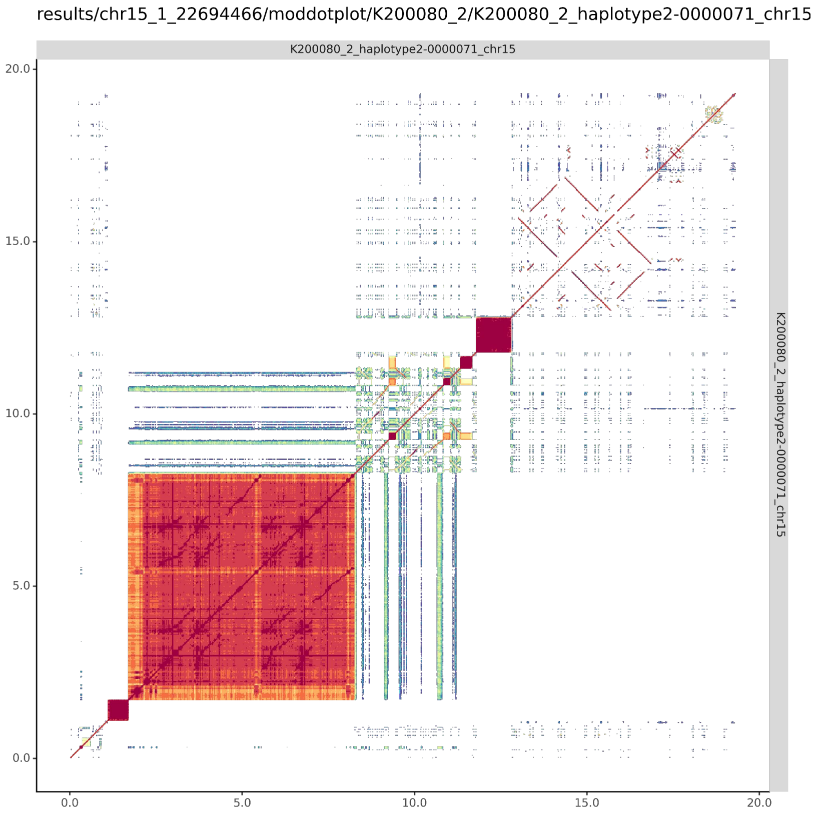

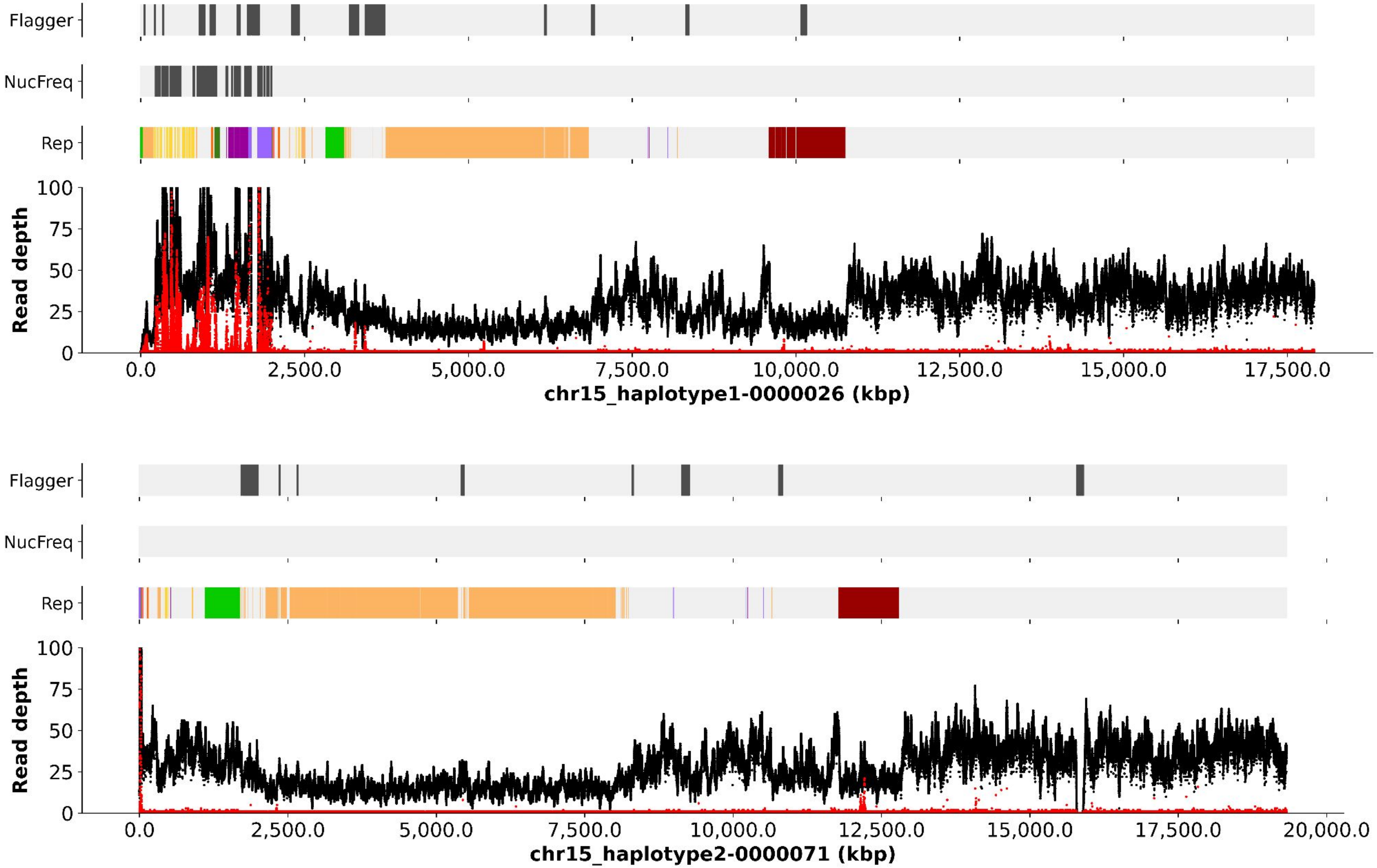
Flagger
NucFreq
Rep
Read depth
100
75
50
25
0
0.0
2,500.0
5,000.0
7,500.0
10,000.0
12,500.0
15,000.0
17,500.0
chr15_haplotype1-0000026 (kbp)
Flagger
NucFreq
Rep
Read depth
100
75
50
25
0
0.0
2,500.0
5,000.0
7,500.0
10,000.0
12,500.0
15,000.0
17,500.0
20,000.0
chr15_haplotype2-0000071 (kbp)

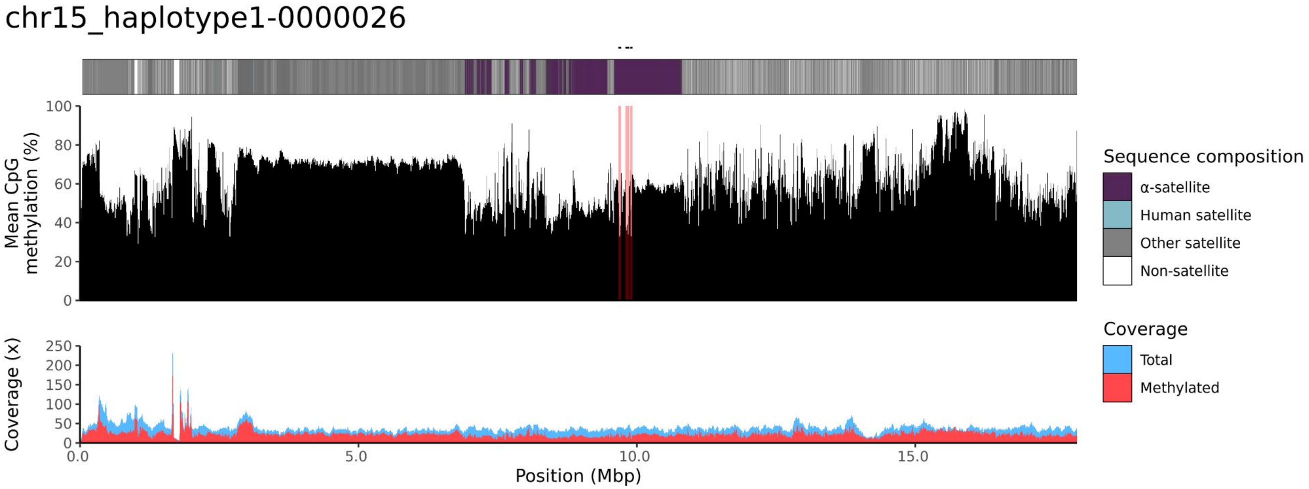
chr15_haplotype1-0000026
Mean CpG methylation (%)
100
80
60
40
20
0
Coverage (x)
250
200
150
100
50
0
0.0
5.0
10.0
15.0
Position (Mbp)
Sequence composition
α-satellite
Human satellite
Other satellite
Non-satellite
Coverage
Total
Methylated

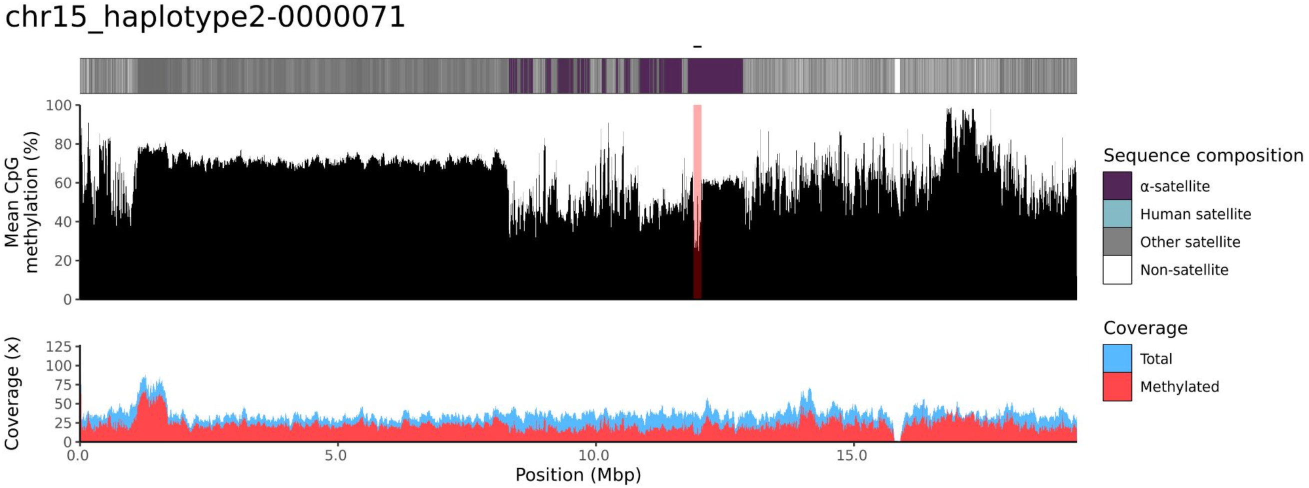
chr15_haplotype2-0000071
Mean CpG methylation (%)
100
80
60
40
20
0
Coverage (x)
125
100
75
50
25
0
0.0
5.0
10.0
15.0
Position (Mbp)
Sequence composition
α-satellite
Human satellite
Other satellite
Non-satellite
Coverage
Total
Methylated

### chr21

#### K200080_1_haplotype1-0000030_chr21

results/chr21_1_16306378/moddotplot/K200080_1/K200080_1_haplotype1-0000030_chr21

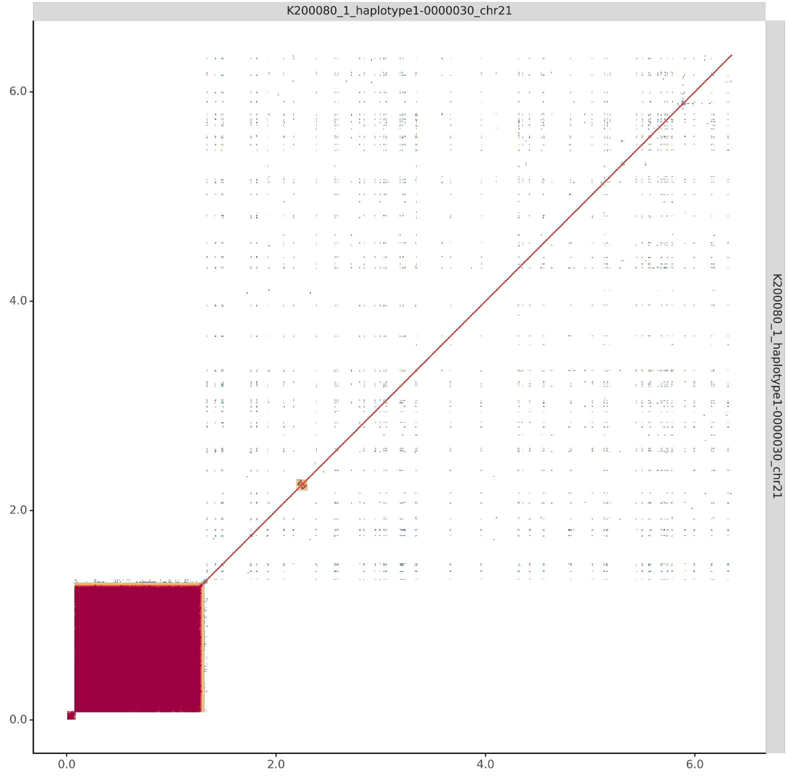


#### K200080_2_haplotype2-0000081_chr21

results/chr21_1_16306378/moddotplot/K200080_2/K200080_2_haplotype2-0000081_chr21

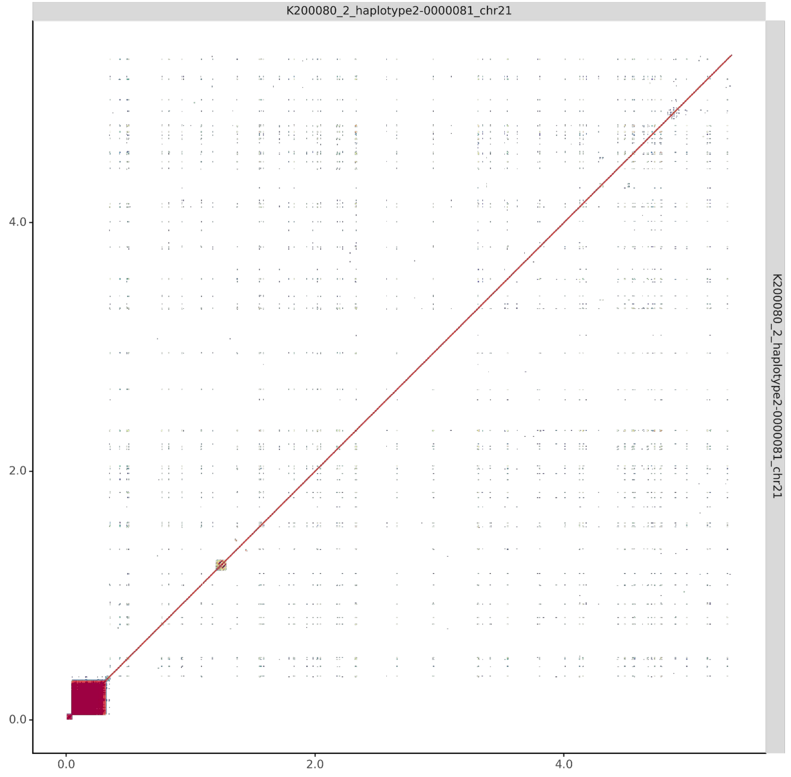

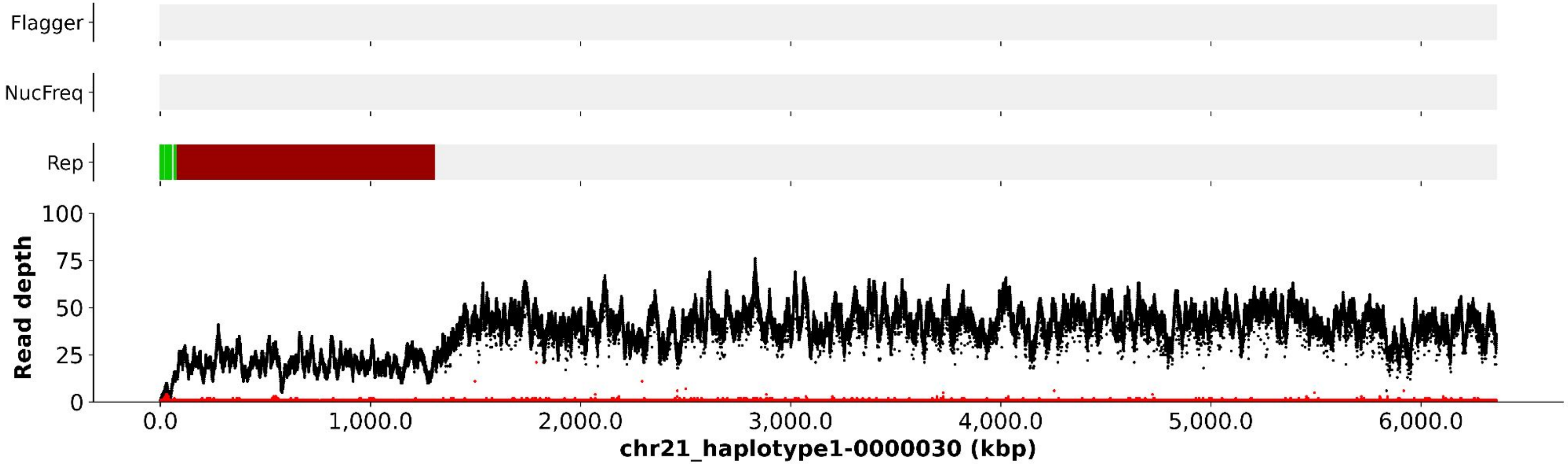
Flagger
NucFreq
Rep
Read depth
100
75
50
25
0
0.0
1,000.0
2,000.0
3,000.0
4,000.0
5,000.0
6,000.0
chr21_haplotype1-0000030 (kbp)

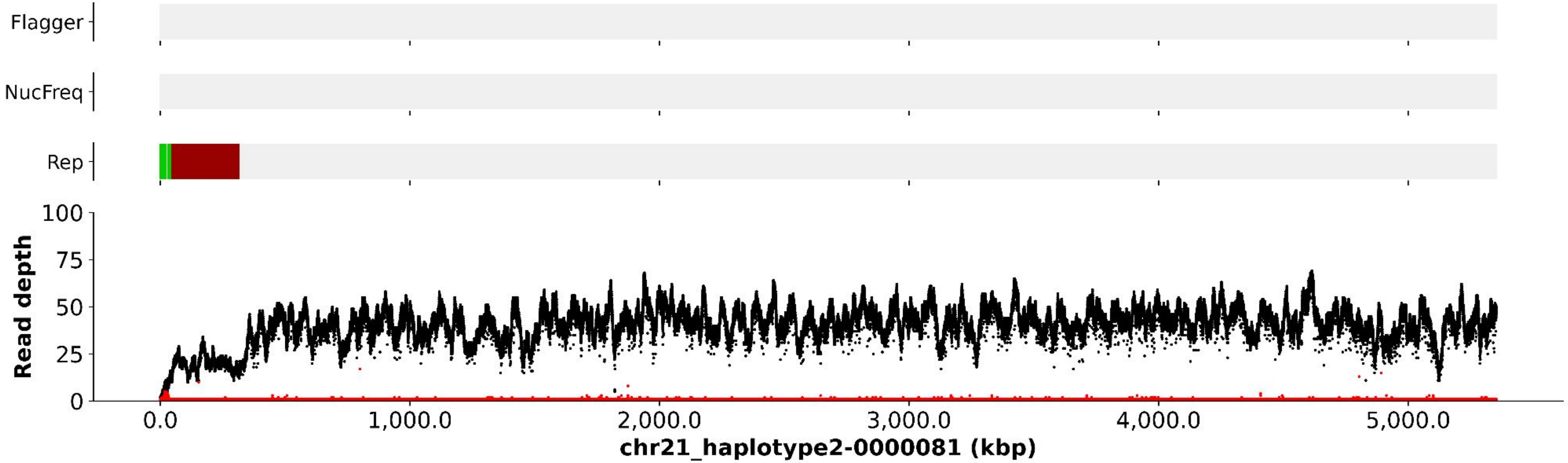
Flagger
NucFreq
Rep
Read depth
100
75
50
25
0
0.0
1,000.0
2,000.0
3,000.0
4,000.0
5,000.0
chr21_haplotype2-0000081 (kbp)

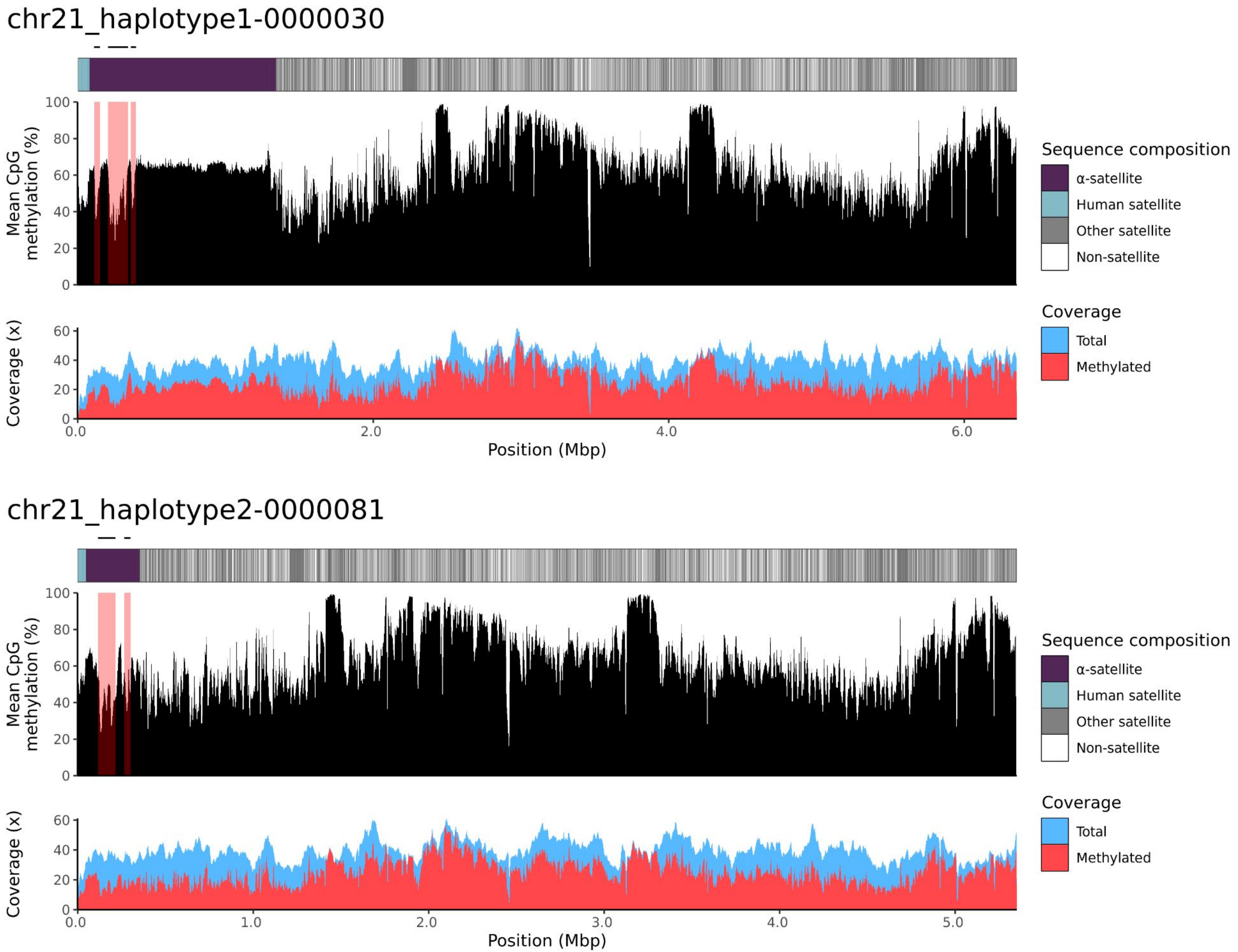

chr21_haplotype1-0000030
Mean CpG methylation (%)
100
80
60
40
20
0
Coverage (x)
60
40
20
0
0.0
2.0
4.0
6.0
Position (Mbp)
Sequence composition
α-satellite
Human satellite
Other satellite
Non-satellite
Coverage
Total
Methylated
chr21_haplotype2-0000081
Mean CpG methylation (%)
Coverage (x)
0.0
1.0
2.0
3.0
4.0
5.0
Position (Mbp)

### chr22

#### K200080_1_haplotype1-0000016_chr22

results/chr22_1_20711065/moddotplot/K200080_1/K200080_1_haplotype1-0000016_chr22

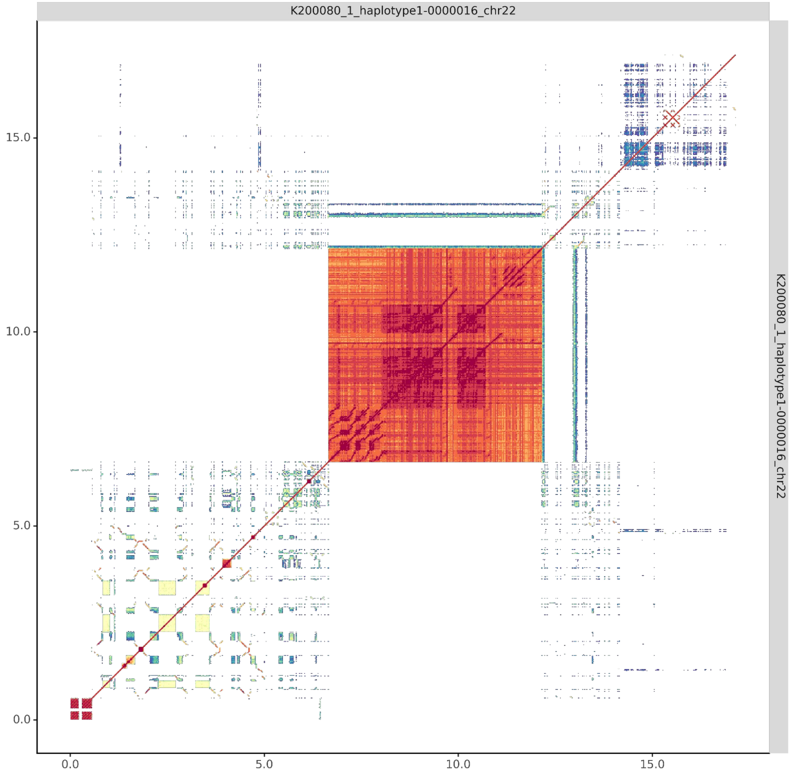


#### K200080_2_haplotype2-0000063_chr22

results/chr22_1_20711065/moddotplot/K200080_2/K200080_2_haplotype2-0000063_chr22

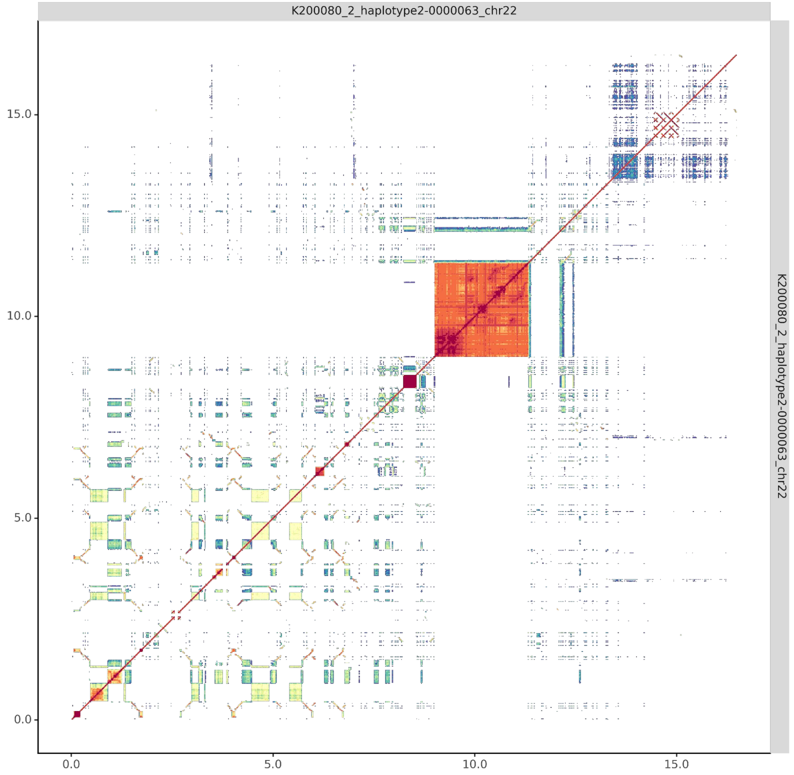

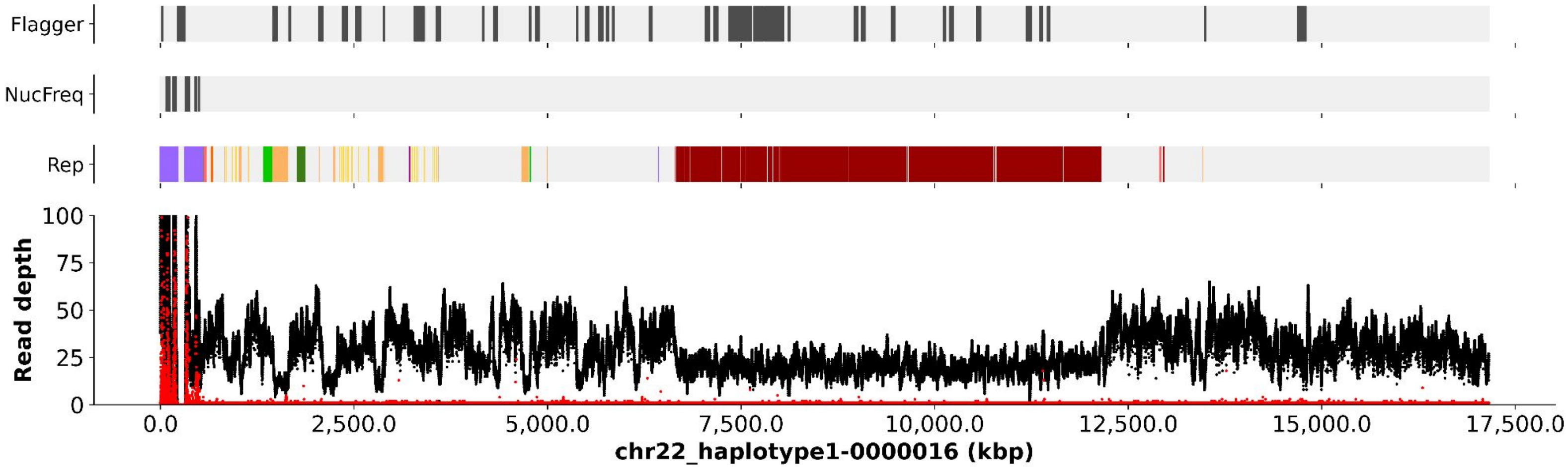
Flagger
NucFreq
Rep
Read depth
100
75
50
25
0
0.0
2,500.0
5,000.0
7,500.0
10,000.0
12,500.0
15,000.0
17,500.0
chr22_haplotype1-0000016 (kbp)

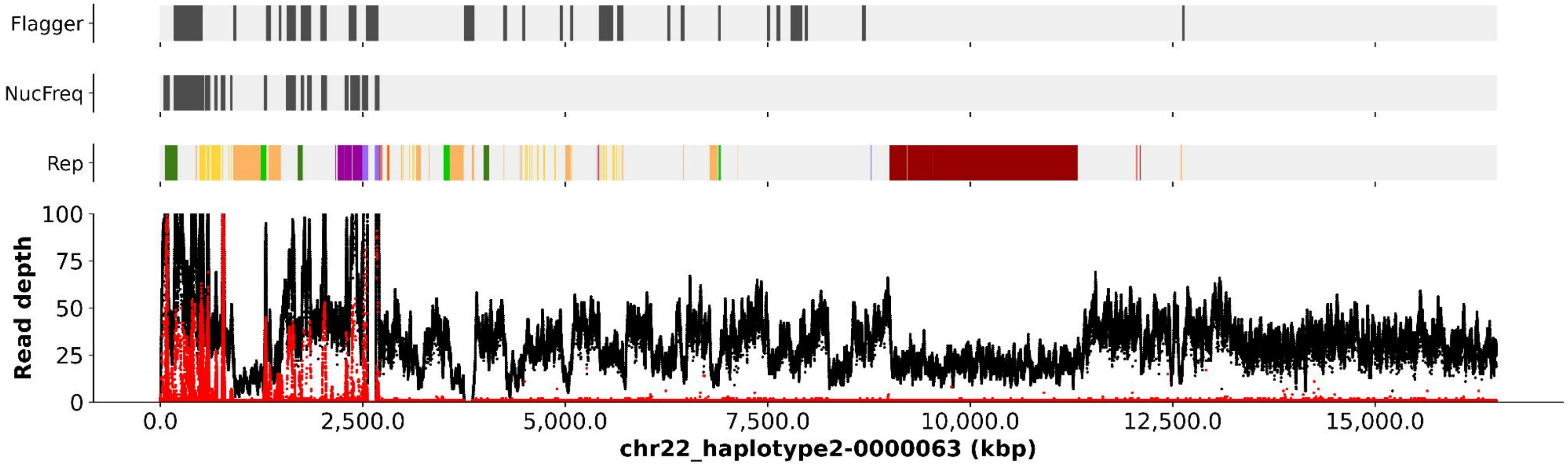
Flagger
NucFreq
Rep
Read depth
100
75
50
25
0
0.0
2,500.0
5,000.0
7,500.0
10,000.0
12,500.0
15,000.0
chr22_haplotype2-0000063 (kbp)

chr22_haplotype1-0000016

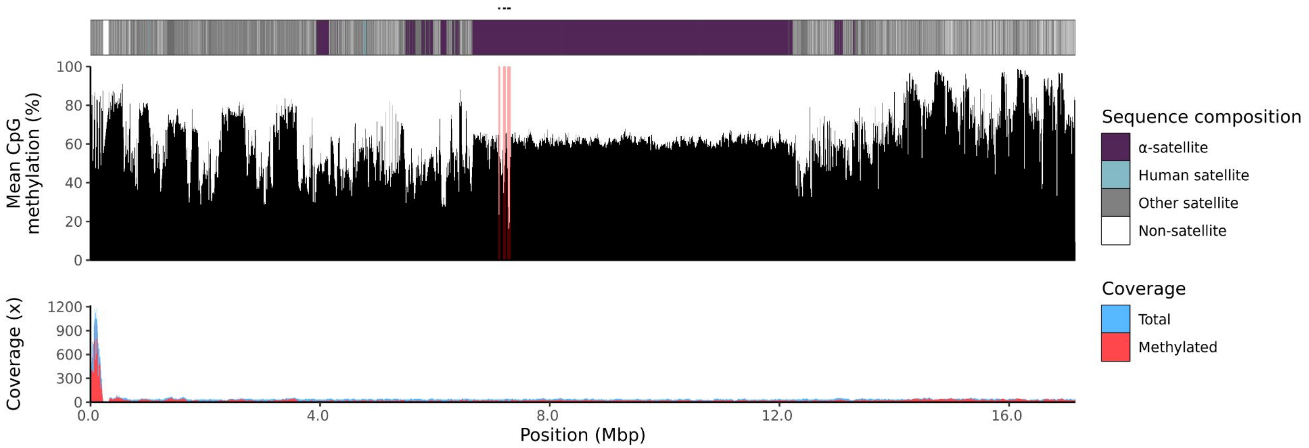


chr22_haplotype2-0000063

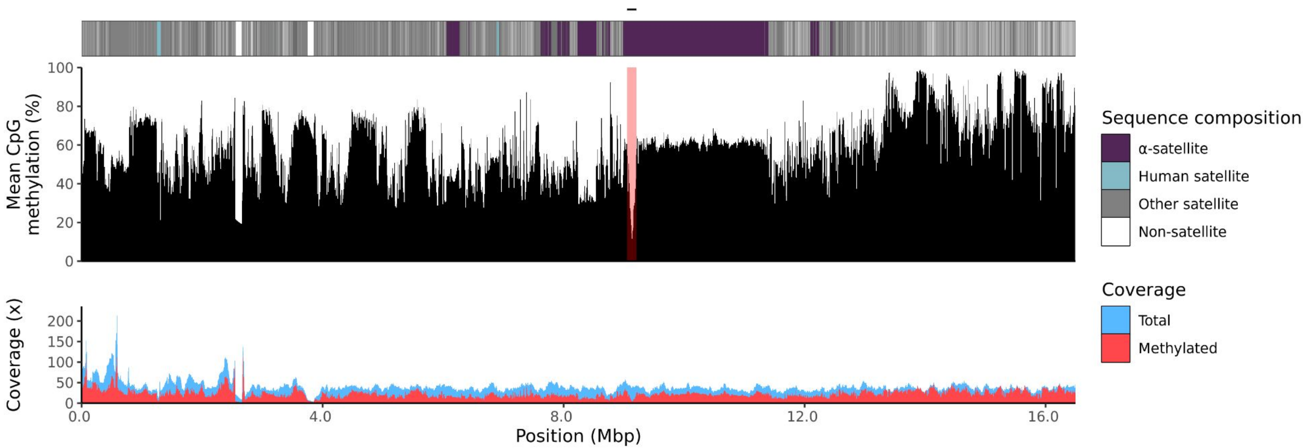

200081

### chr13

#### K200081_1_haplotype1-0000023_chr13

#### K200081_2_haplotype2-0000065_chr13

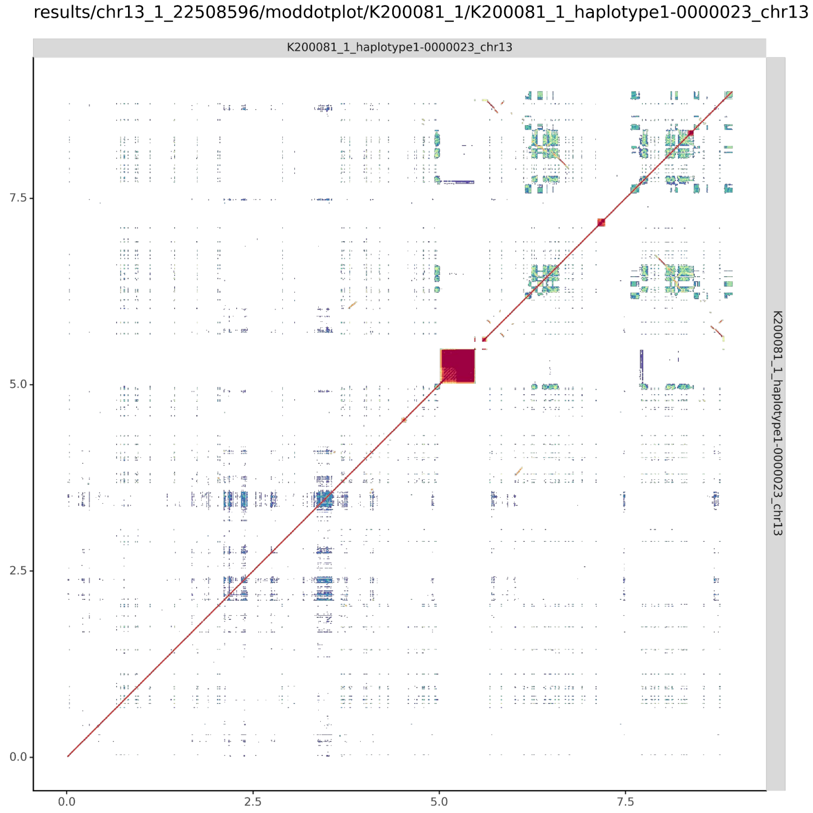


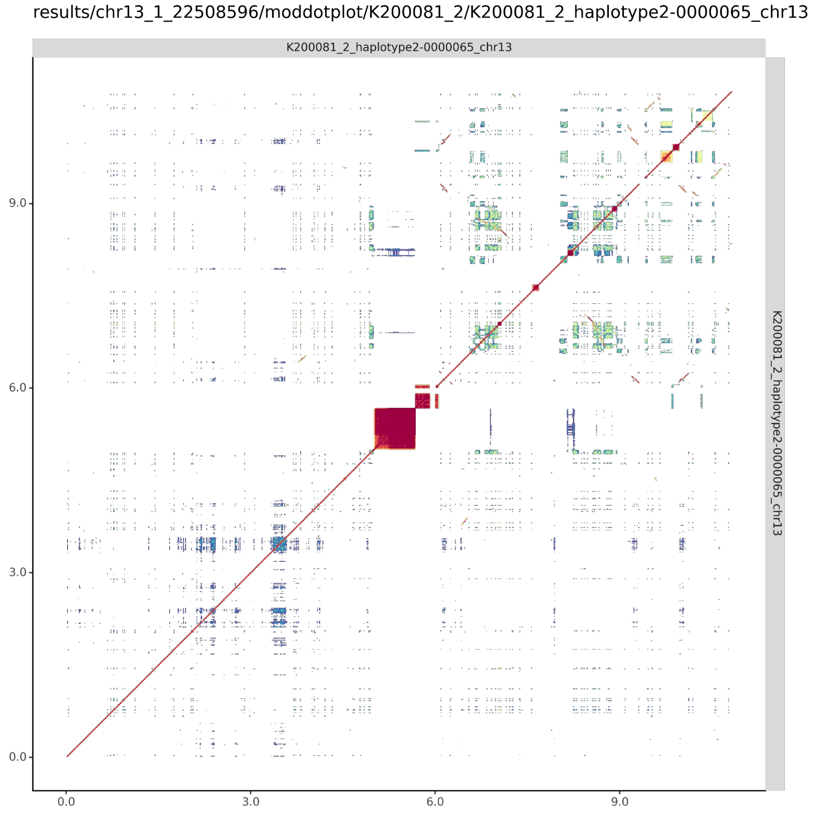

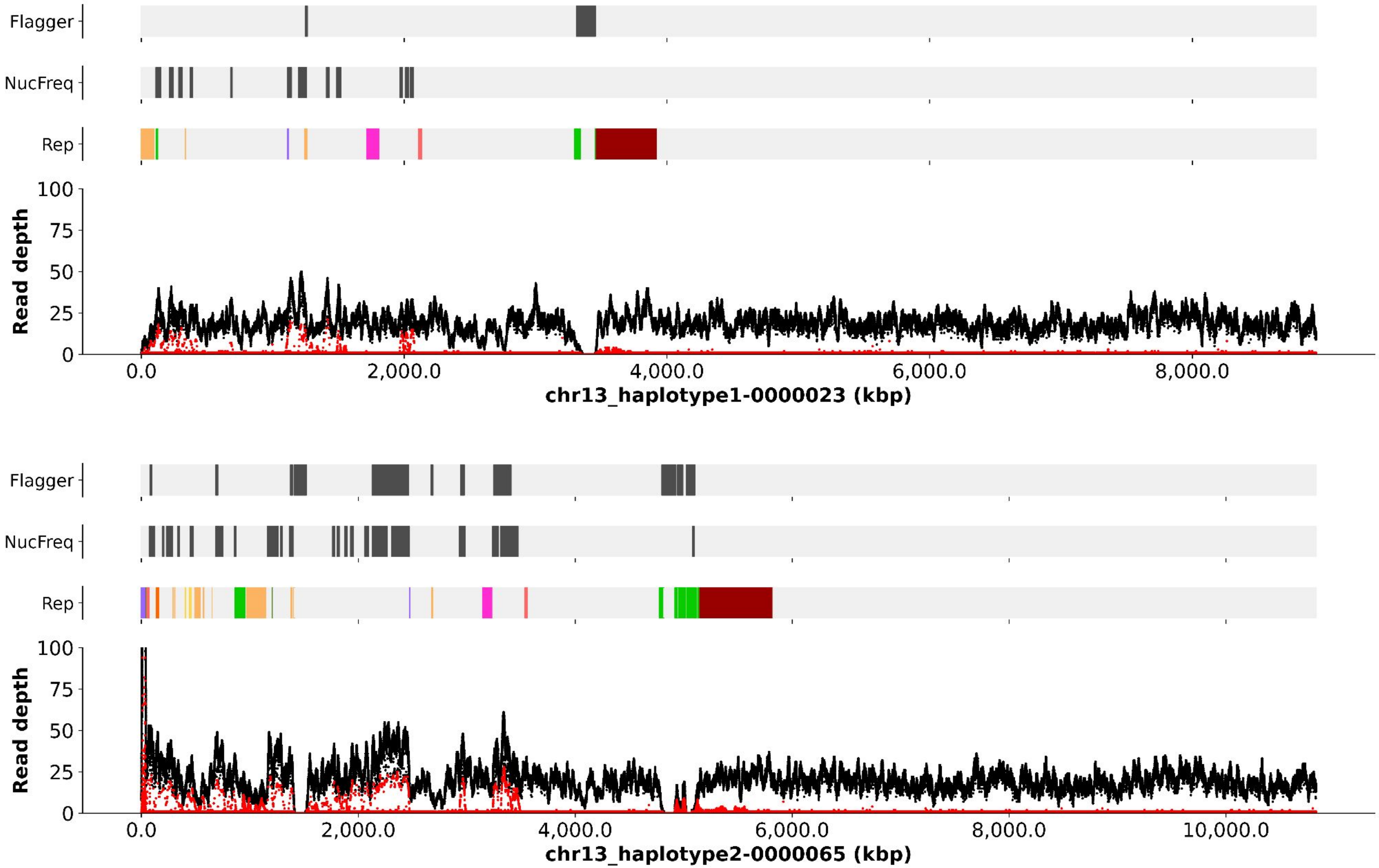

Flagger
NucFreq
Rep
Read depth
100
75
50
25
0
0.0
2,000.0
4,000.0
6,000.0
8,000.0
chr13_haplotype1-0000023 (kbp)
Flagger
NucFreq
Rep
Read depth
100
75
50
25
0
0.0
2,000.0
4,000.0
6,000.0
8,000.0
10,000.0
chr13_haplotype2-0000065 (kbp)

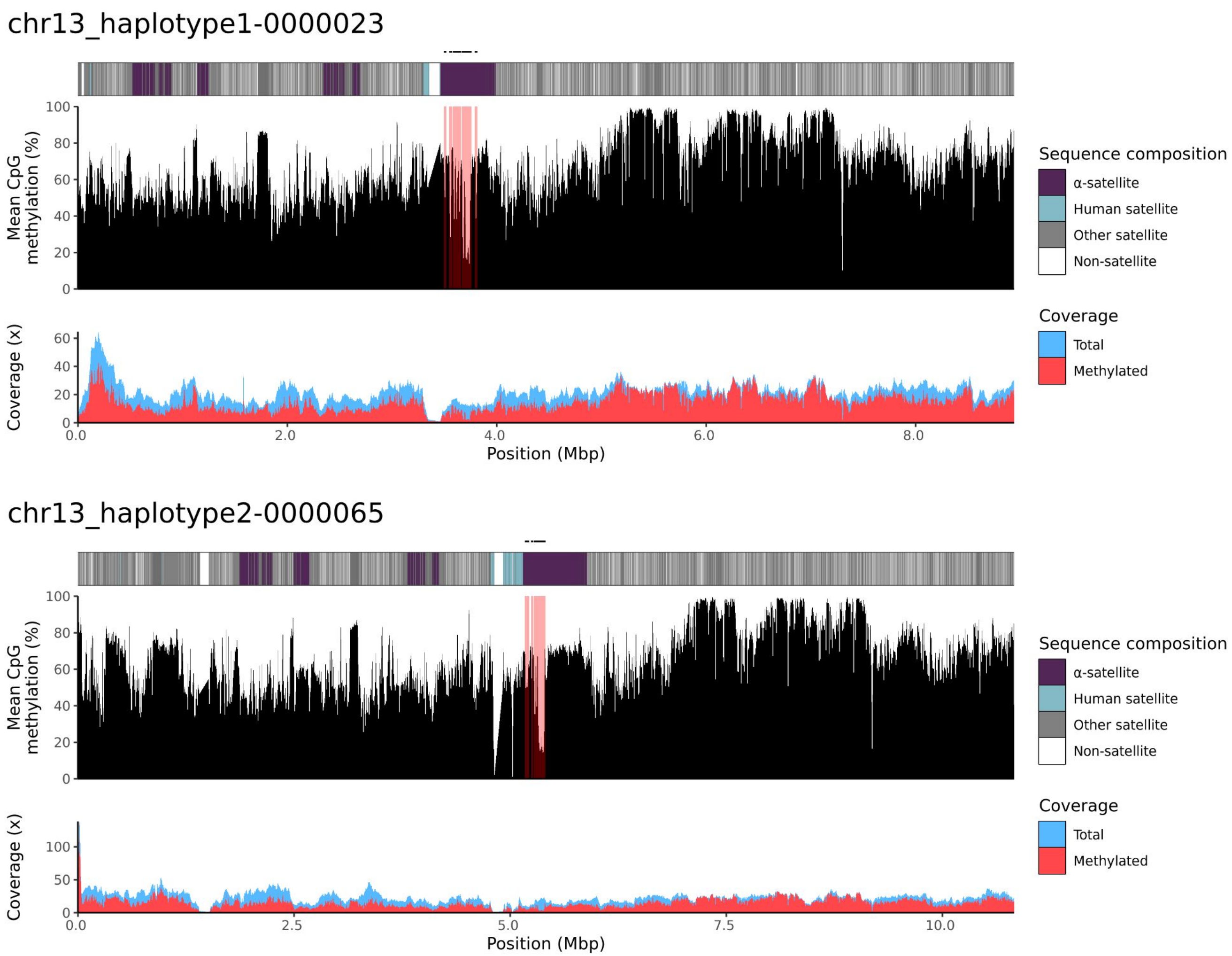
chr13_haplotype1-0000023
Mean CpG methylation (%)
Sequence composition
α-satellite
Human satellite
Other satellite
Non-satellite
Coverage (x)
Coverage
Total
Methylated
Position (Mbp)
chr13_haplotype2-0000065
Mean CpG methylation (%)
Sequence composition
α-satellite
Human satellite
Other satellite
Non-satellite
Coverage (x)
Coverage
Total
Methylated
Position (Mbp)

### chr14

#### K200081_1_haplotype1-0000027_chr14

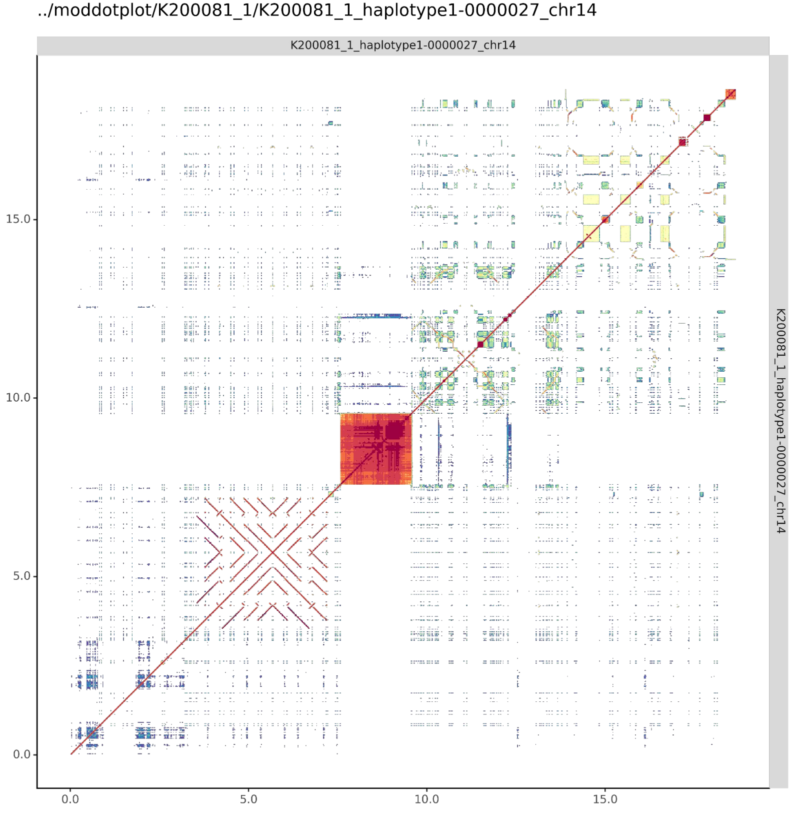


#### K200081_2_haplotype2-0000075_chr14

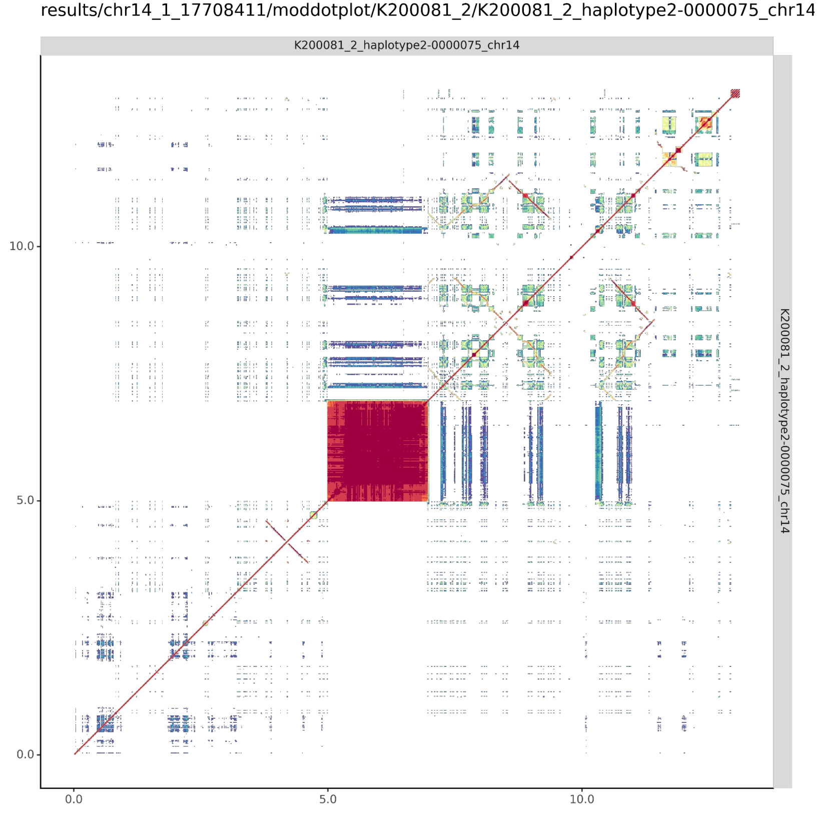

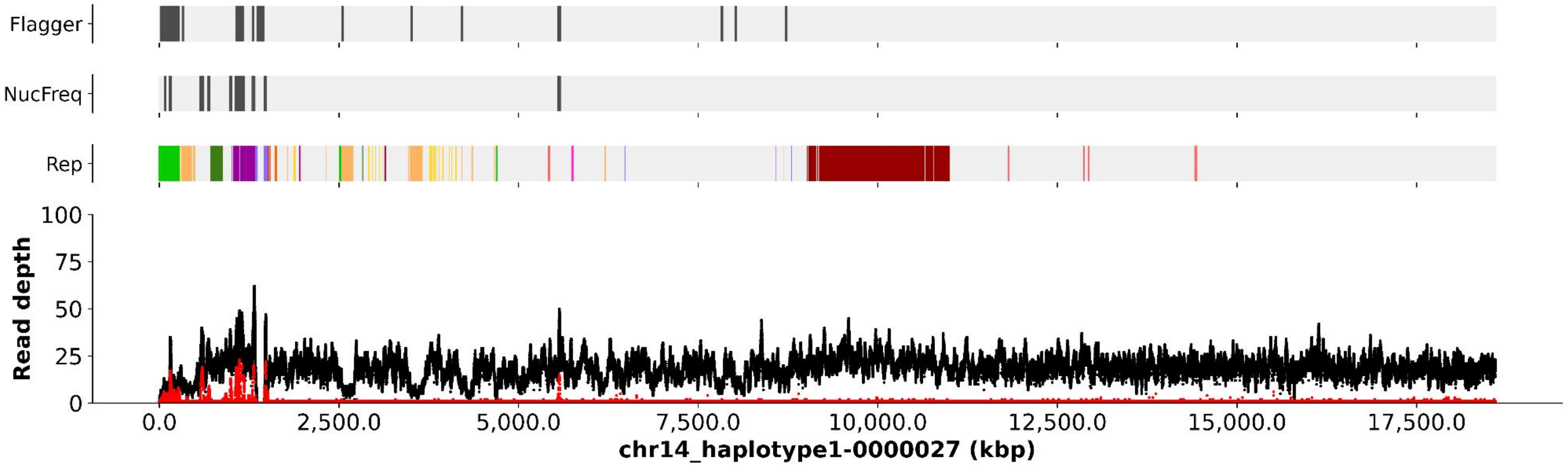

Flagger
NucFreq
Rep
Read depth
100
75
50
25
0
0.0
2,500.0
5,000.0
7,500.0
10,000.0
12,500.0
15,000.0
17,500.0
chr14_haplotype1-0000027 (kbp)


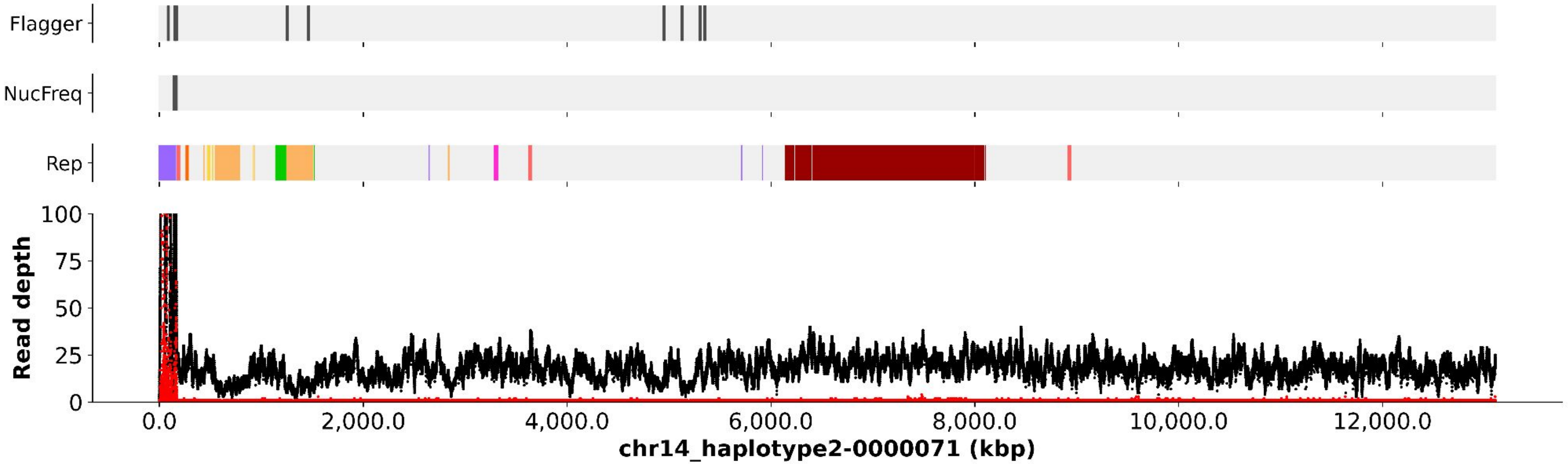

Flagger
NucFreq
Rep
Read depth
100
75
50
25
0
0.0
2,000.0
4,000.0
6,000.0
8,000.0
10,000.0
12,000.0
chr14_haplotype2-0000071 (kbp)

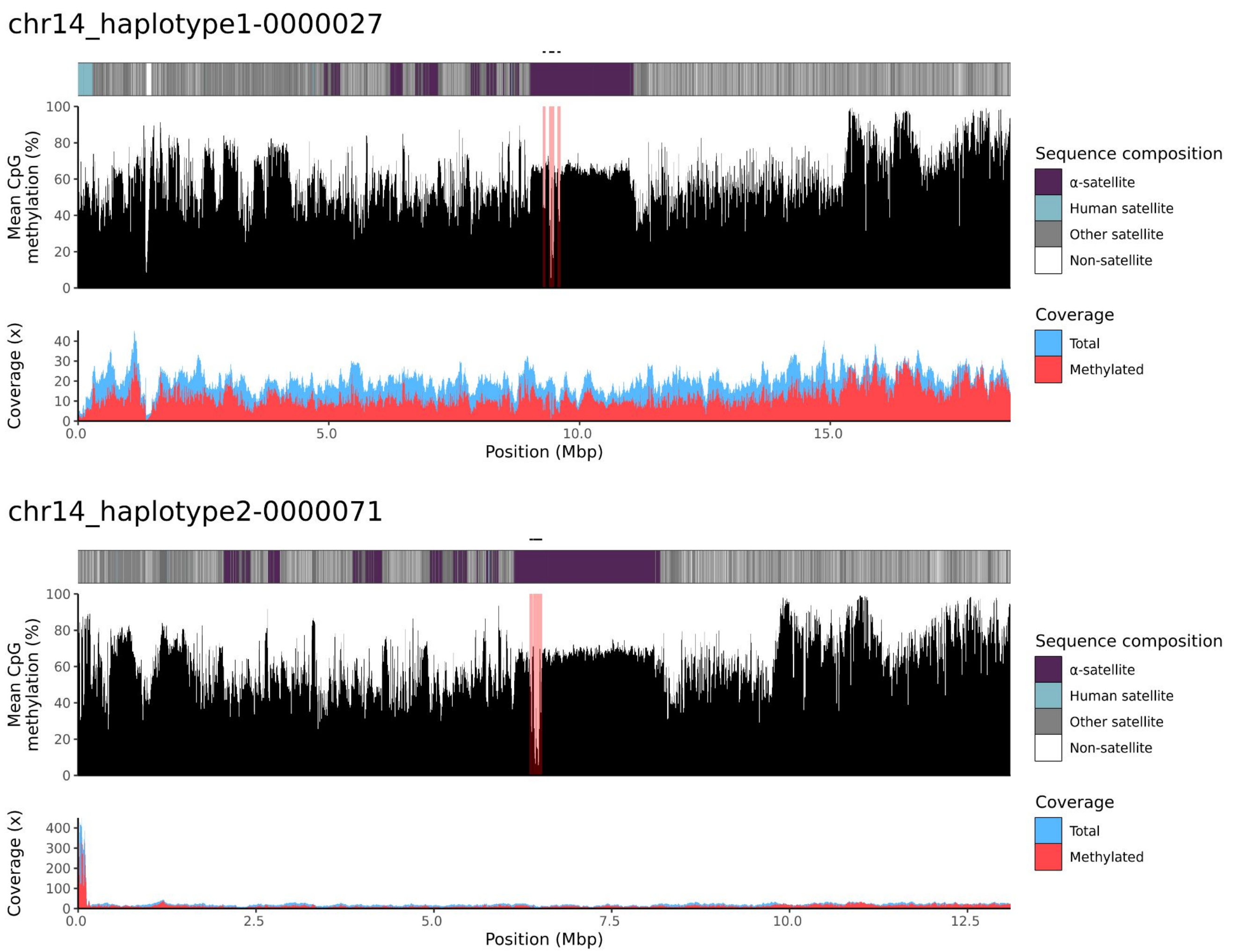

chr14_haplotype1-0000027
Mean CpG methylation (%)
100
80
60
40
20
0
Coverage (x)
40
30
20
10
0
0.0
5.0
10.0
15.0
Position (Mbp)
Sequence composition
α-satellite
Human satellite
Other satellite
Non-satellite
Coverage
Total
Methylated
chr14_haplotype2-0000071
Mean CpG methylation (%)
100
80
60
40
20
0
Coverage (x)
400
300
200
100
0
0.0
2.5
5.0
7.5
10.0
12.5
Position (Mbp)
Sequence composition
α-satellite
Human satellite
Other satellite
Non-satellite
Coverage
Total
Methylated

### chr15

#### K200081_1_haplotype1-0000009_chr15

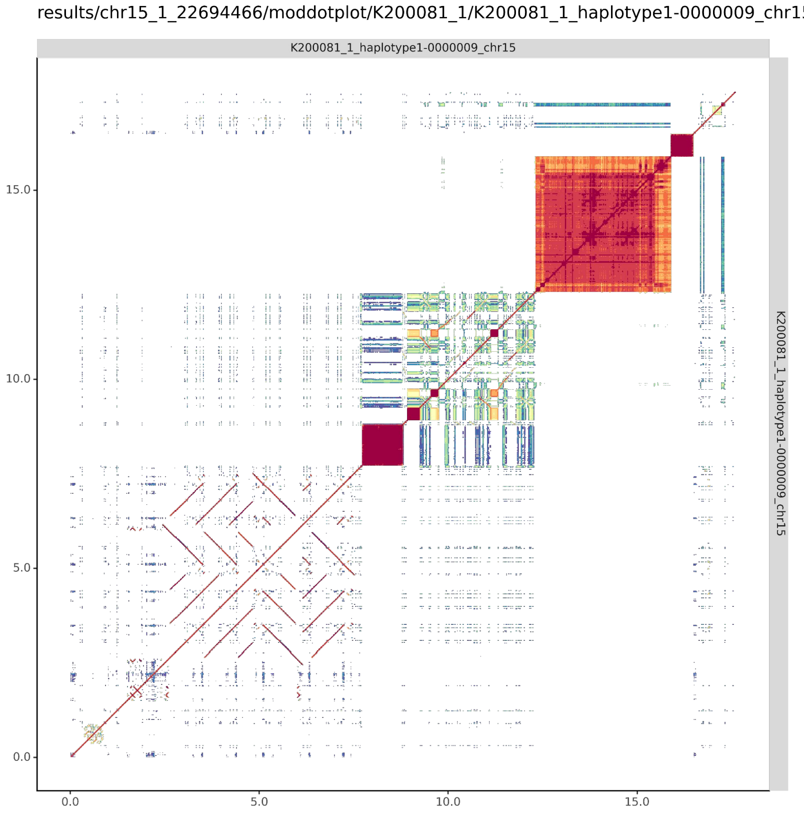


#### K200081_2_haplotype2-0000056_chr15

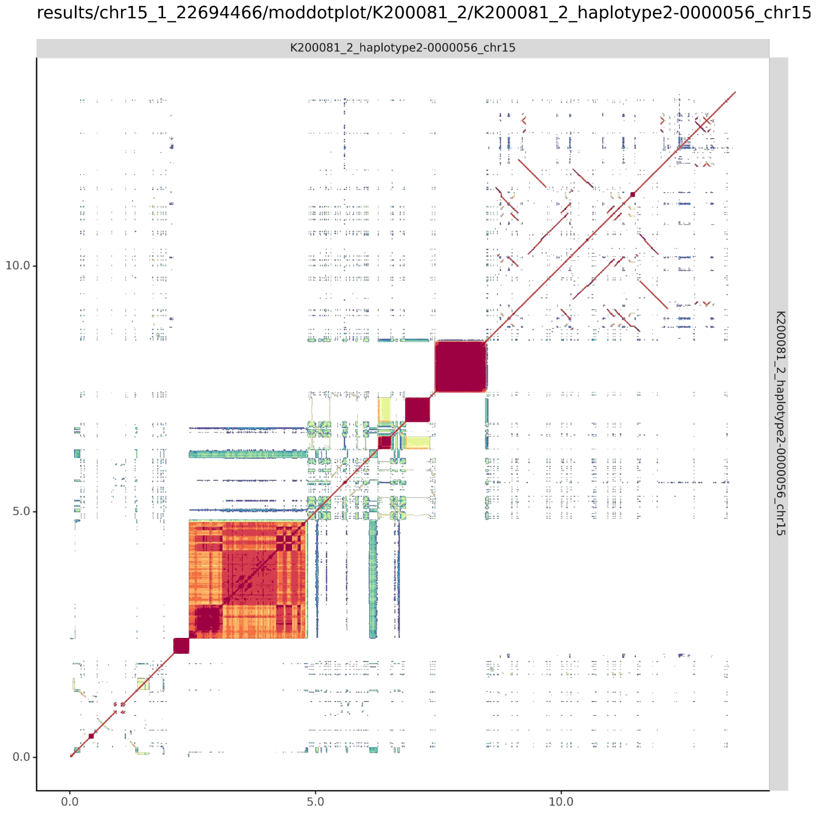

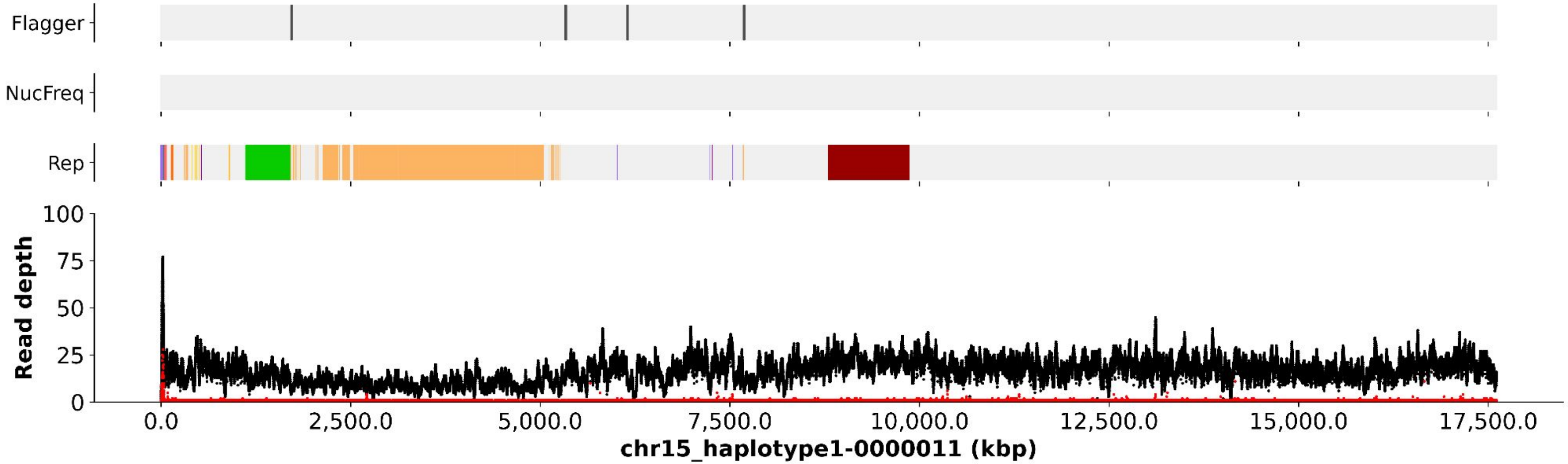
Flagger
NucFreq
Rep
Read depth
100
75
50
25
0
0.0
2,500.0
5,000.0
7,500.0
10,000.0
12,500.0
15,000.0
17,500.0
chr15_haplotype1-0000011 (kbp)

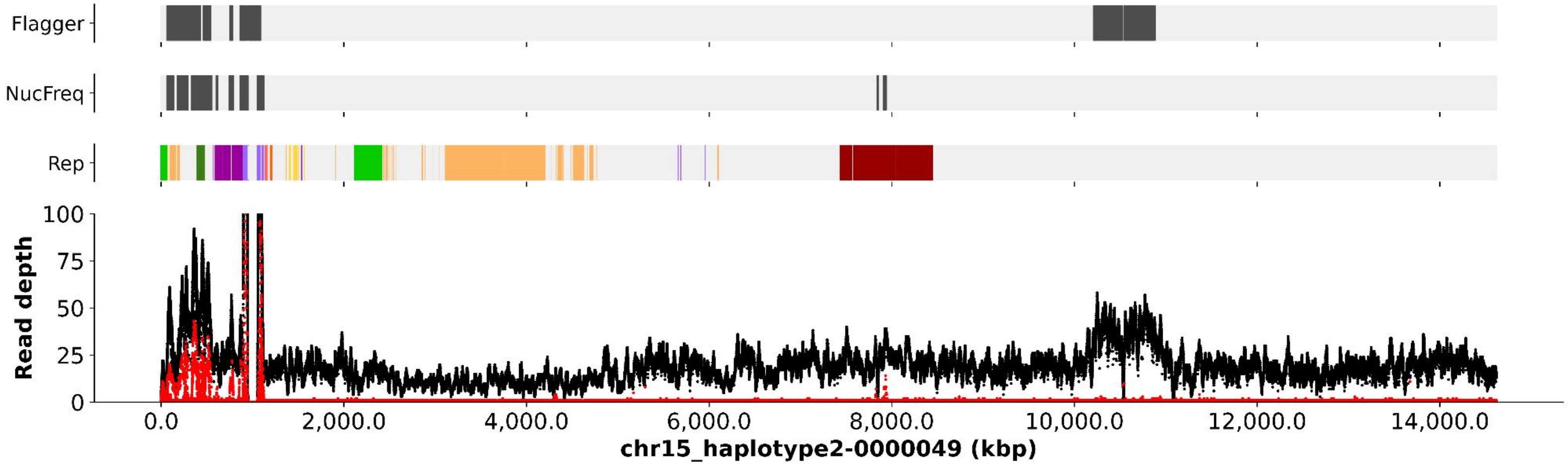
Flagger
NucFreq
Rep
Read depth
100
75
50
25
0
0.0
2,000.0
4,000.0
6,000.0
8,000.0
10,000.0
12,000.0
14,000.0
chr15_haplotype2-0000049 (kbp)

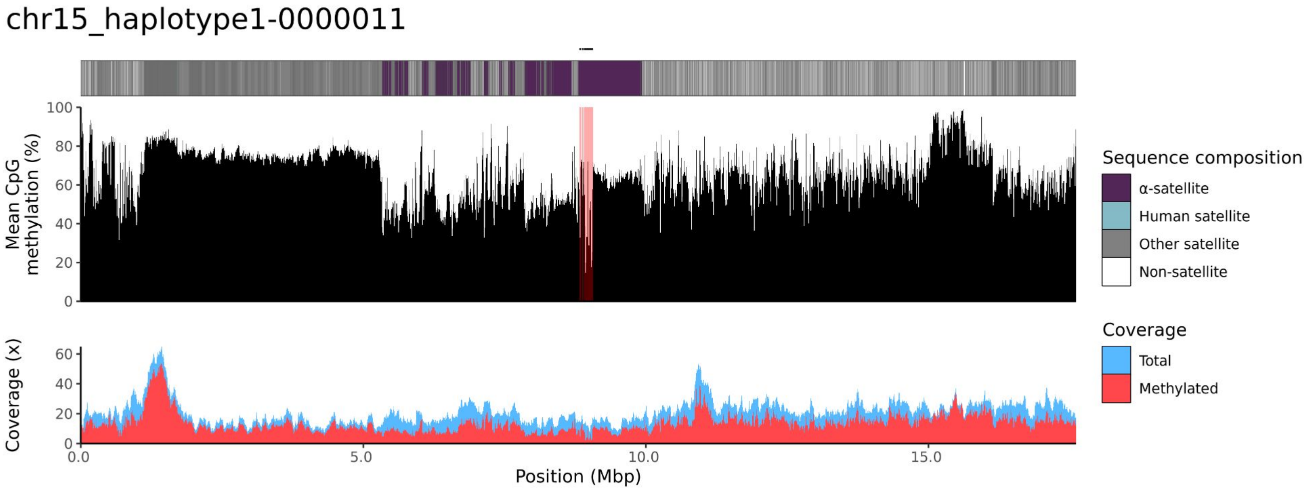
chr15_haplotype1-0000011
Mean CpG methylation (%)
100
80
60
40
20
0
Coverage (x)
60
40
20
0
0.0
5.0
10.0
15.0
Position (Mbp)
Sequence composition
α-satellite
Human satellite
Other satellite
Non-satellite
Coverage
Total
Methylated

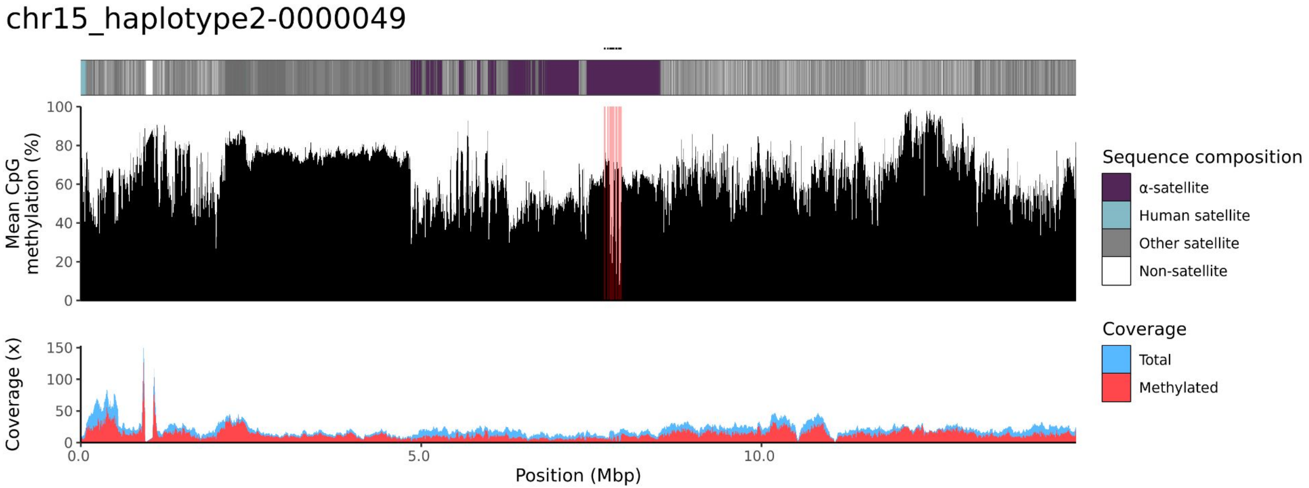
chr15_haplotype2-0000049
Mean CpG methylation (%)
100
80
60
40
20
0
Coverage (x)
150
100
50
0
0.0
5.0
10.0
Position (Mbp)
Sequence composition
α-satellite
Human satellite
Other satellite
Non-satellite
Coverage
Total
Methylated

### chr21

#### K200081_1_haplotype1-0000015_chr21

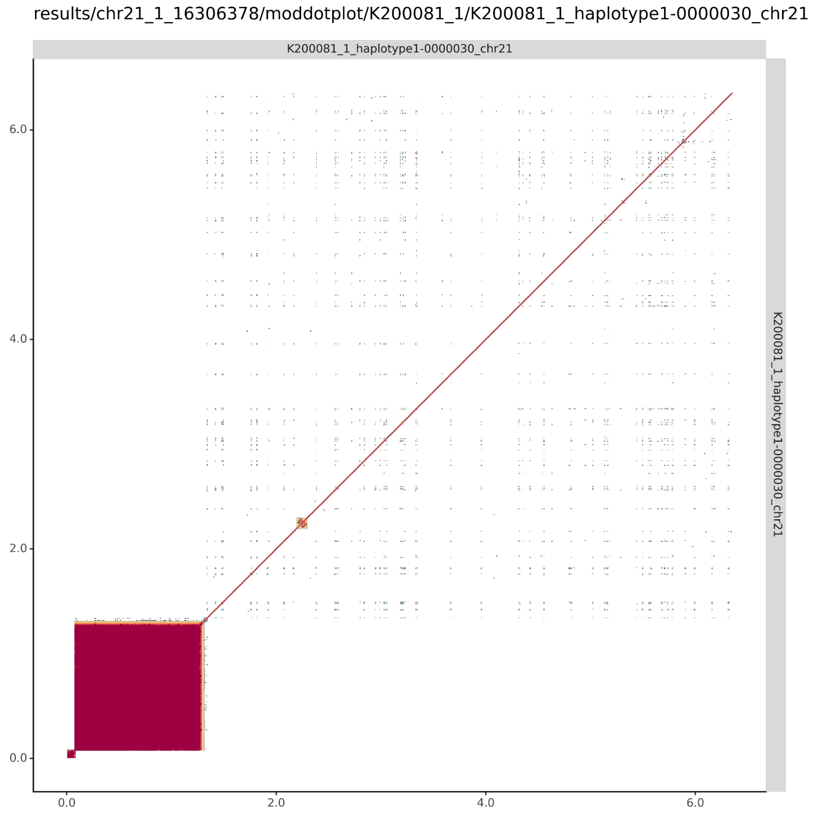


#### K200081_2_haplotype2-0000059_chr21

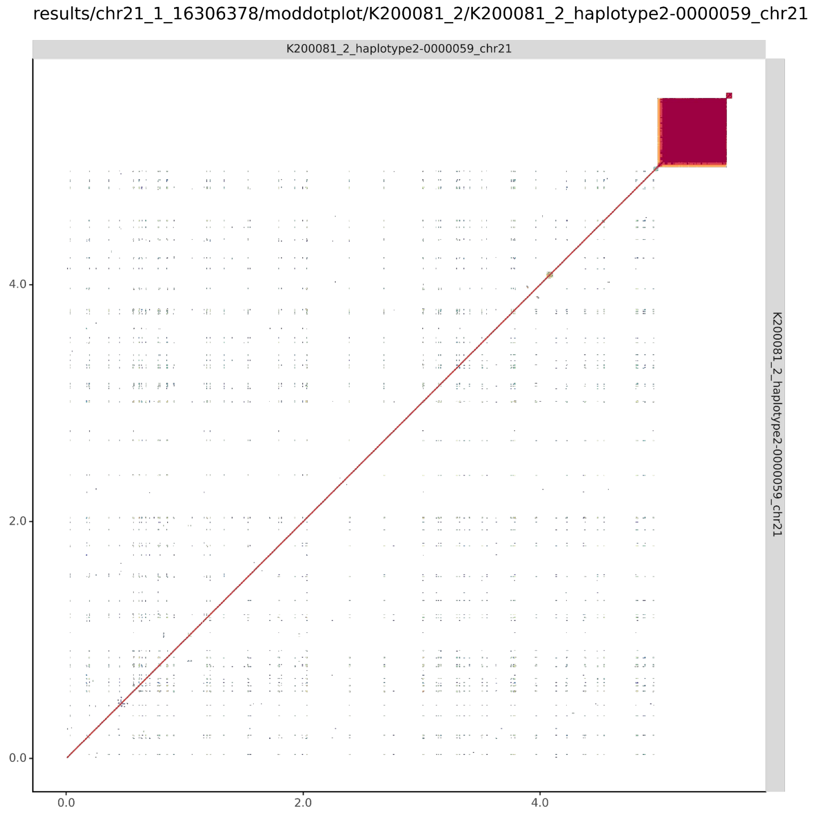

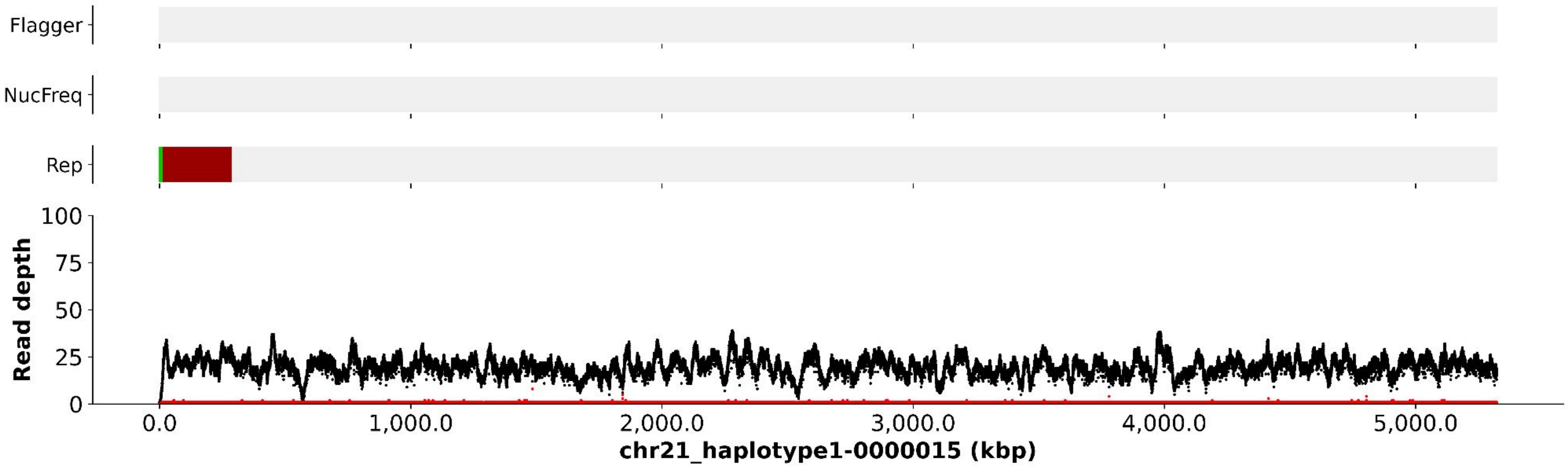

Flagger
NucFreq
Rep
Read depth
100
75
50
25
0
0.0
1,000.0
2,000.0
3,000.0
4,000.0
5,000.0
chr21_haplotype1-0000015 (kbp)


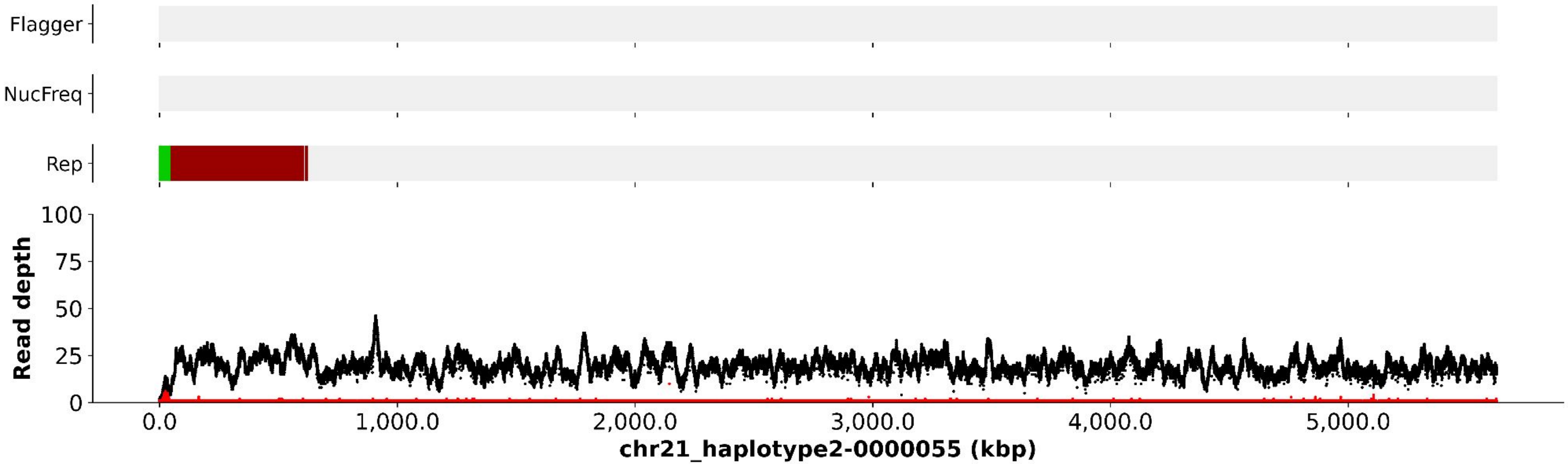

Flagger
NucFreq
Rep
Read depth
100
75
50
25
0
0.0
1,000.0
2,000.0
3,000.0
4,000.0
5,000.0
chr21_haplotype2-0000055 (kbp)

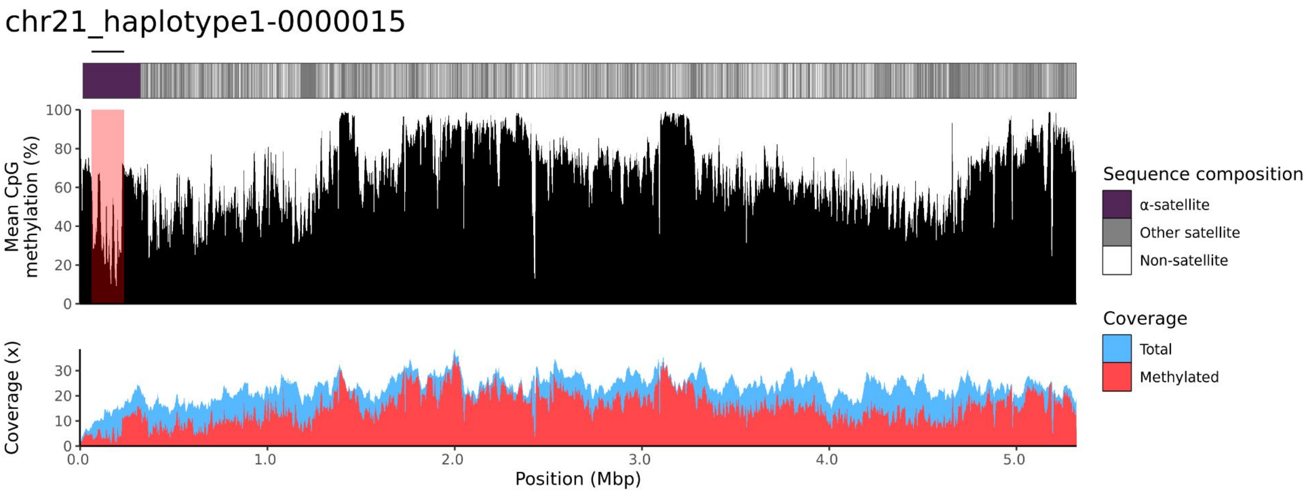
chr21_haplotype1-0000015
Mean CpG methylation (%)
100
80
60
40
20
0
Coverage (x)
30
20
10
0
0.0
1.0
2.0
3.0
4.0
5.0
Position (Mbp)
Sequence composition
α-satellite
Other satellite
Non-satellite
Coverage
Total
Methylated

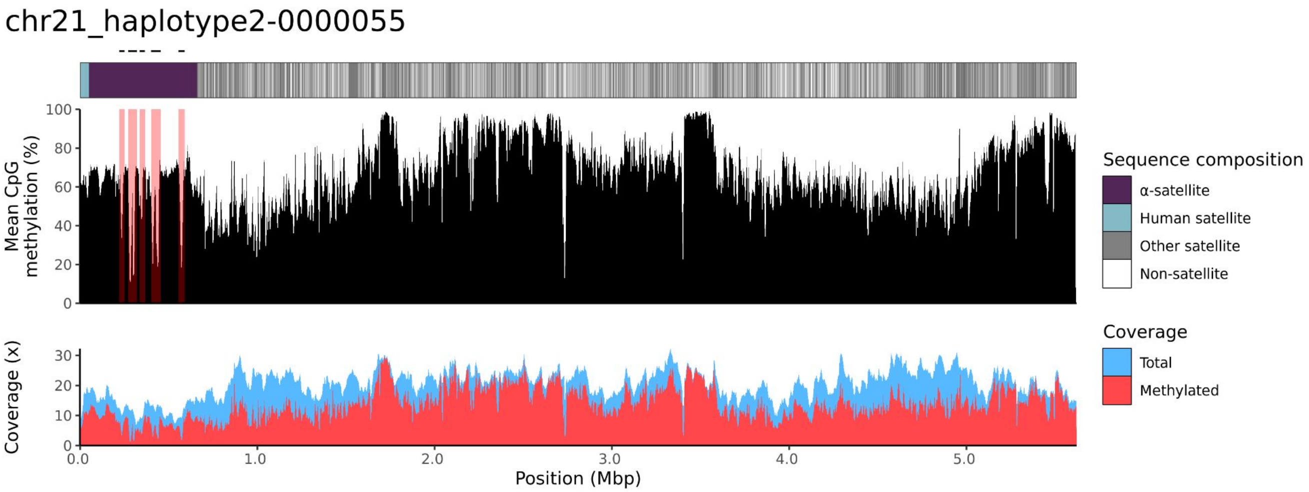
chr21_haplotype2-0000055
Mean CpG methylation (%)
100
80
60
40
20
0
Coverage (x)
30
20
10
0
0.0
1.0
2.0
3.0
4.0
5.0
Position (Mbp)
Sequence composition
α-satellite
Human satellite
Other satellite
Non-satellite
Coverage
Total
Methylated

### chr22

#### K200081_1_haplotype1-0000024_chr22

results/chr22_1_20711065/moddotplot/K200081_1/K200081_1_haplotype1-0000024_chr22

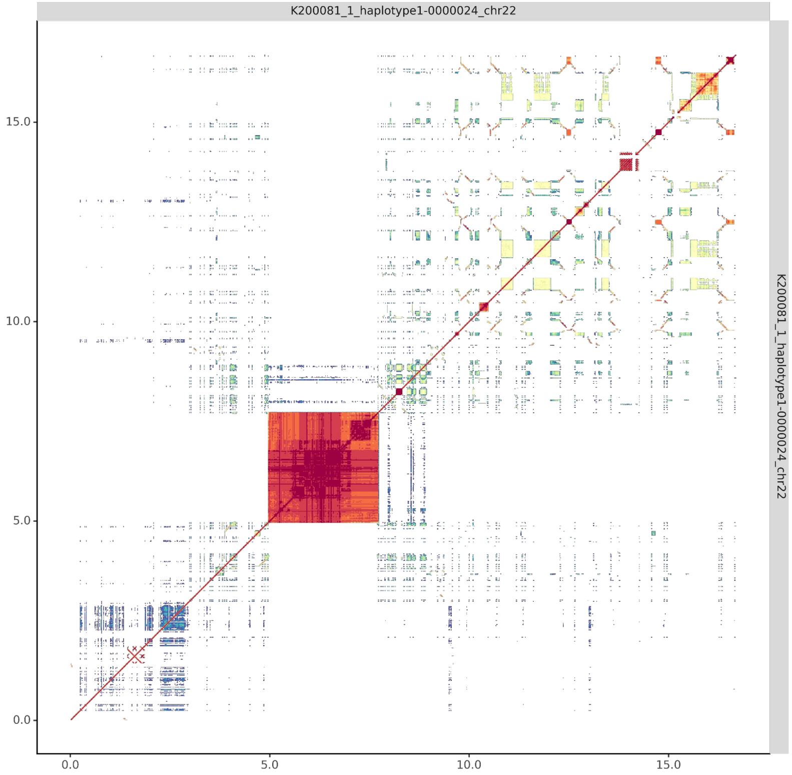


#### K200081_2_haplotype2-0000072_chr22

results/chr22_1_20711065/moddotplot/K200081_2/K200081_2_haplotype2-0000072_chr22

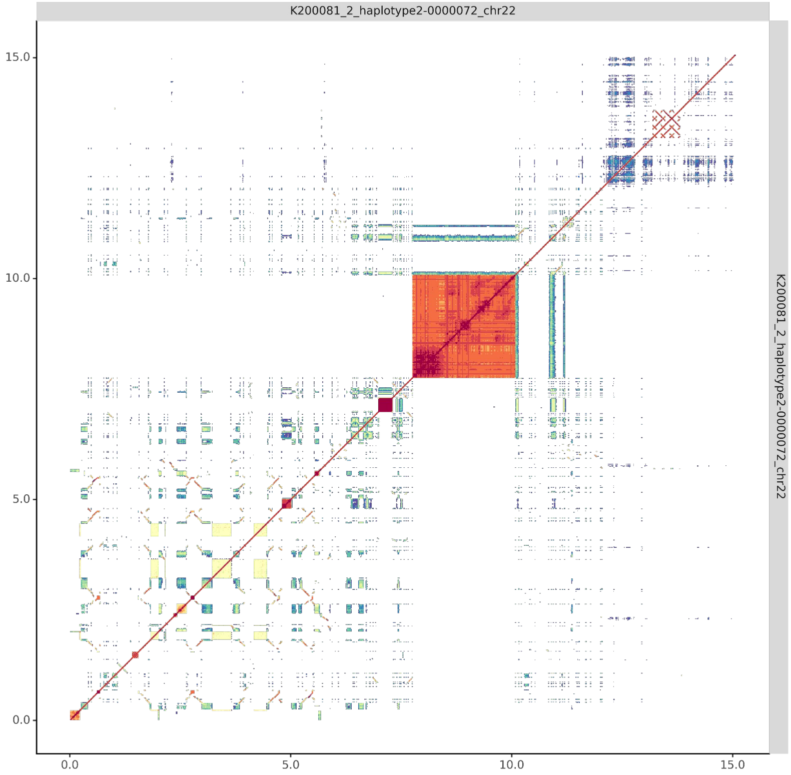

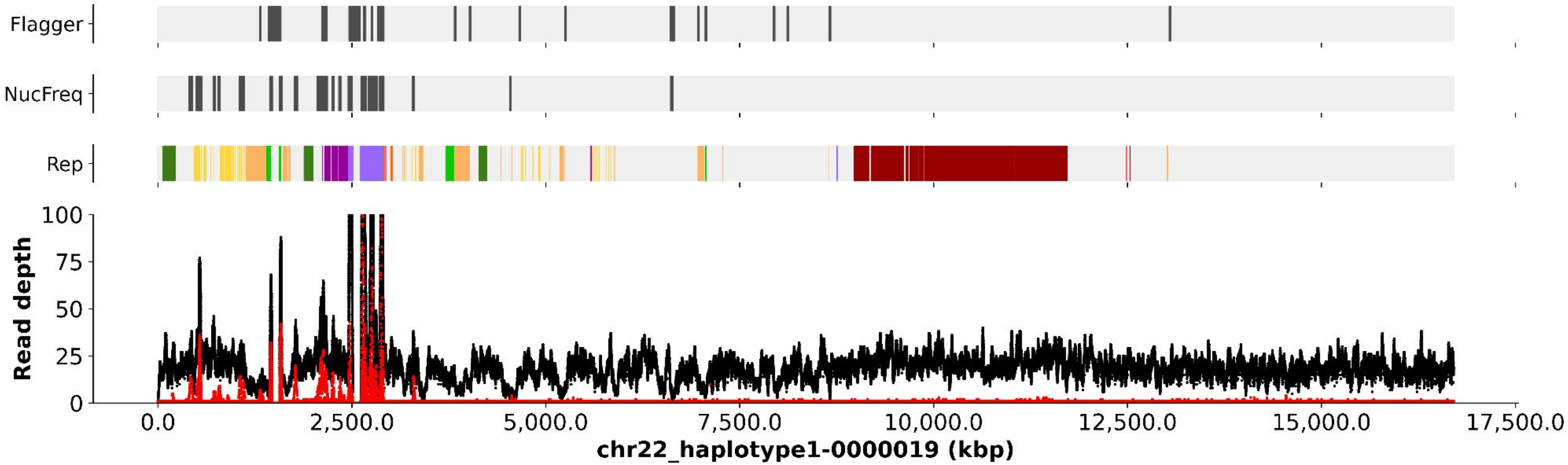
Flagger
NucFreq
Rep
Read depth
100
75
50
25
0
0.0
2,500.0
5,000.0
7,500.0
10,000.0
12,500.0
15,000.0
17,500.0
chr22_haplotype1-0000019 (kbp)

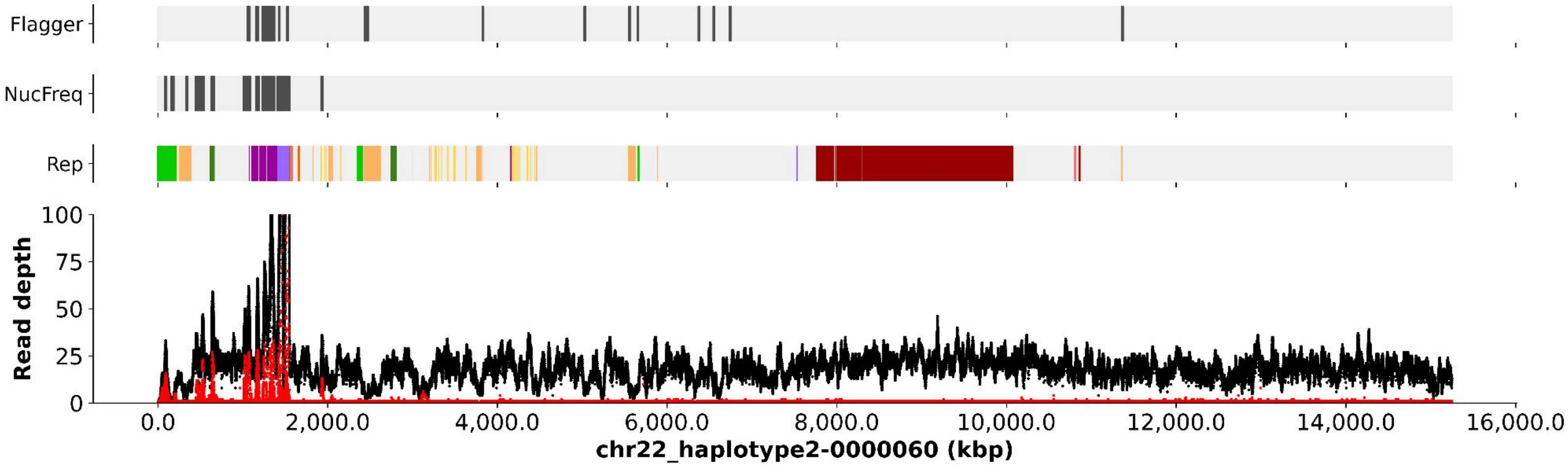
Flagger
NucFreq
Rep
Read depth
100
75
50
25
0
0.0
2,000.0
4,000.0
6,000.0
8,000.0
10,000.0
12,000.0
14,000.0
16,000.0
chr22_haplotype2-0000060 (kbp)

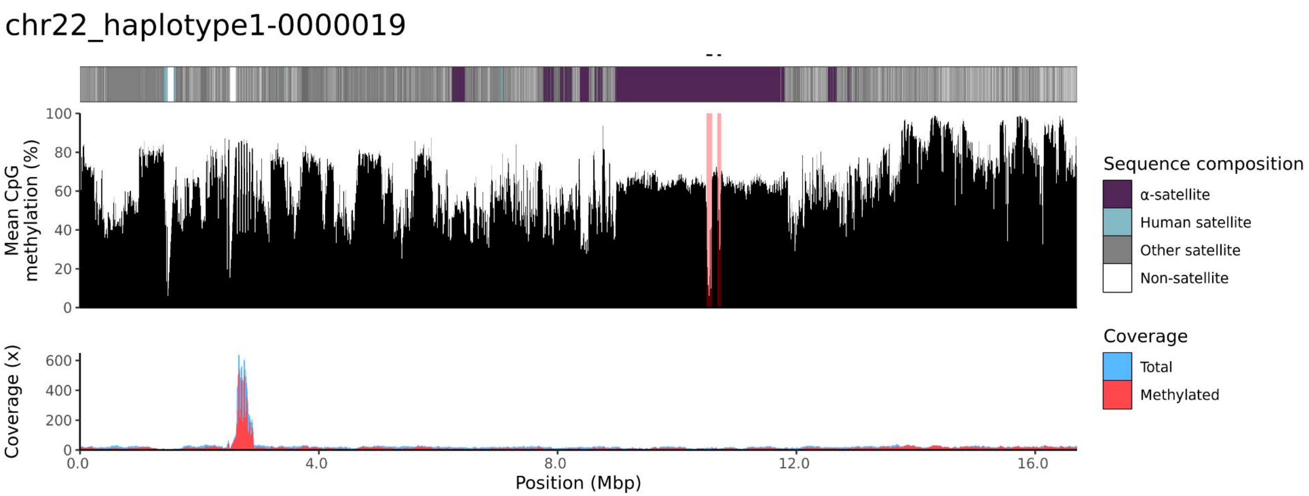

chr22_haplotype1-0000019
Mean CpG methylation (%)
Coverage (x)
Position (Mbp)
Sequence composition
α-satellite
Human satellite
Other satellite
Non-satellite
Coverage
Total
Methylated


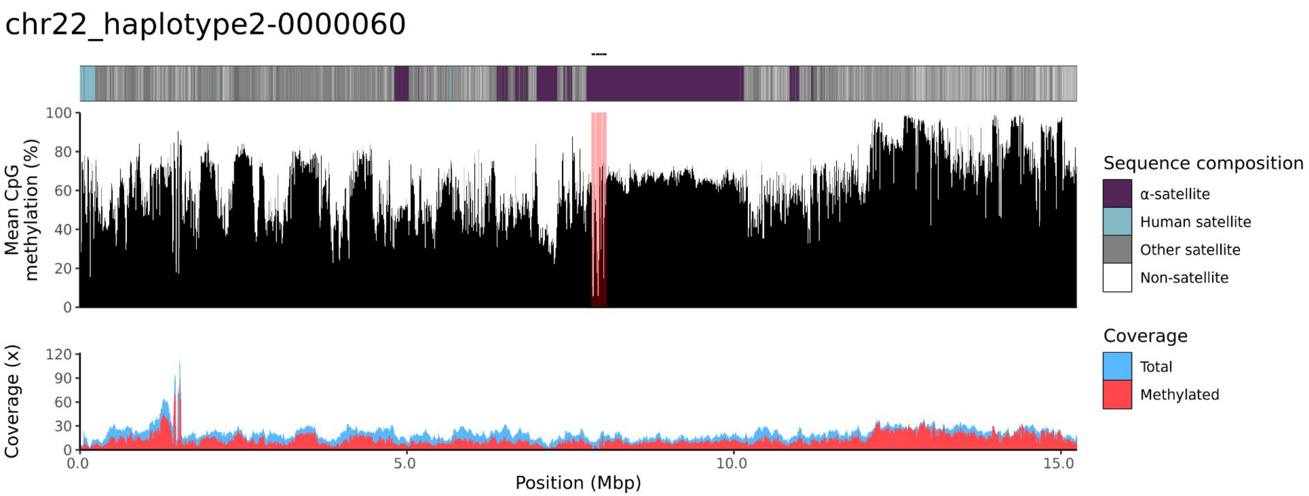

chr22_haplotype2-0000060
Mean CpG methylation (%)
Coverage (x)
Position (Mbp)
Sequence composition
α-satellite
Human satellite
Other satellite
Non-satellite
Coverage
Total
Methylated

200082

### chr14

#### K200082_1_haplotype1-0000010_chr14

results/chr14_1_17708411/moddotplot/K200082_1/K200082_1_haplotype1-0000010_chr14

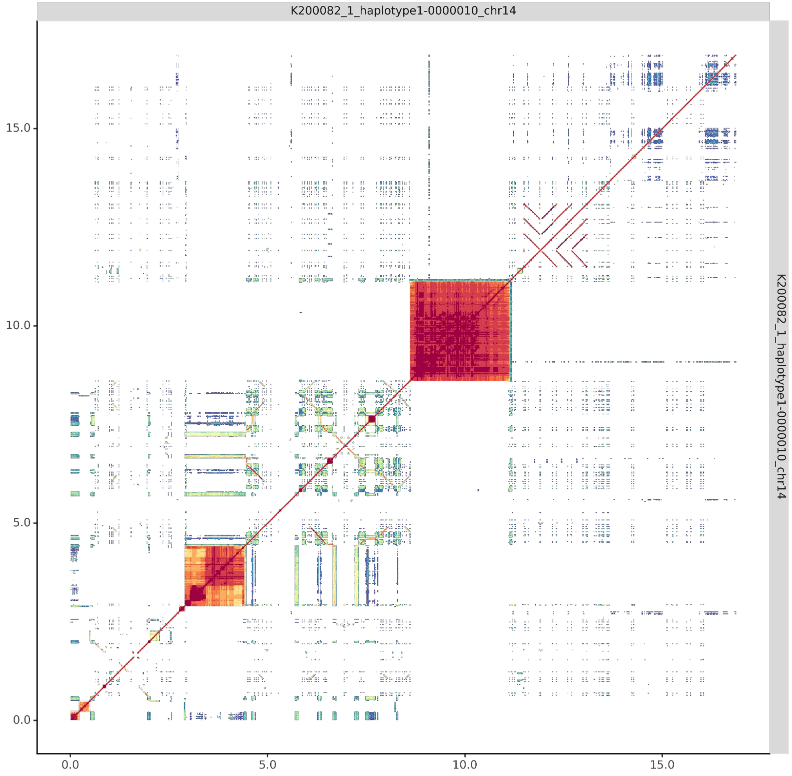


#### K200082_2_haplotype2-0000128_chr14

results/chr14_1_17708411/moddotplot/K200082_2/K200082_2_haplotype2-0000128_chr14

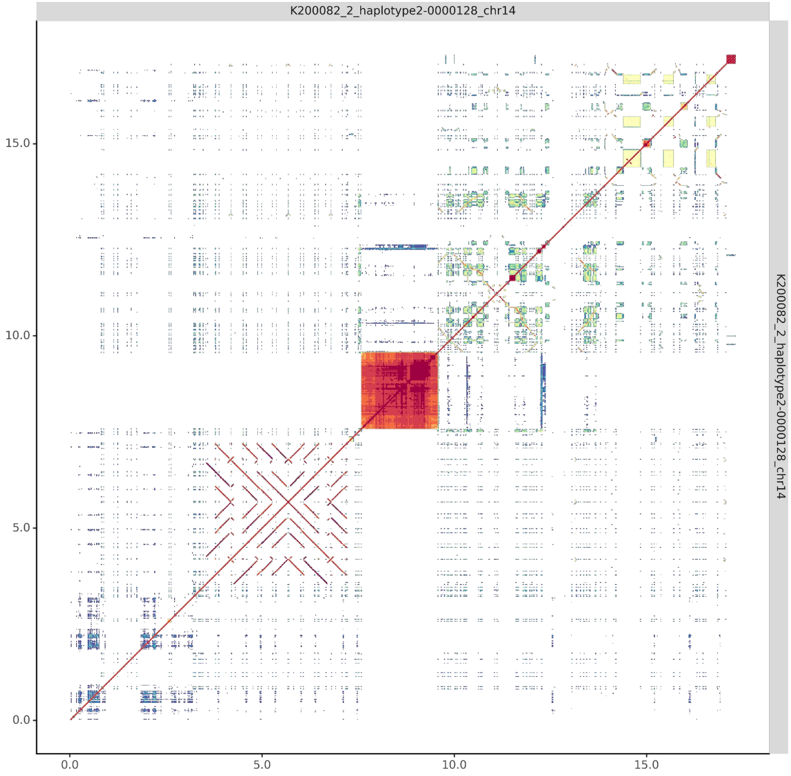

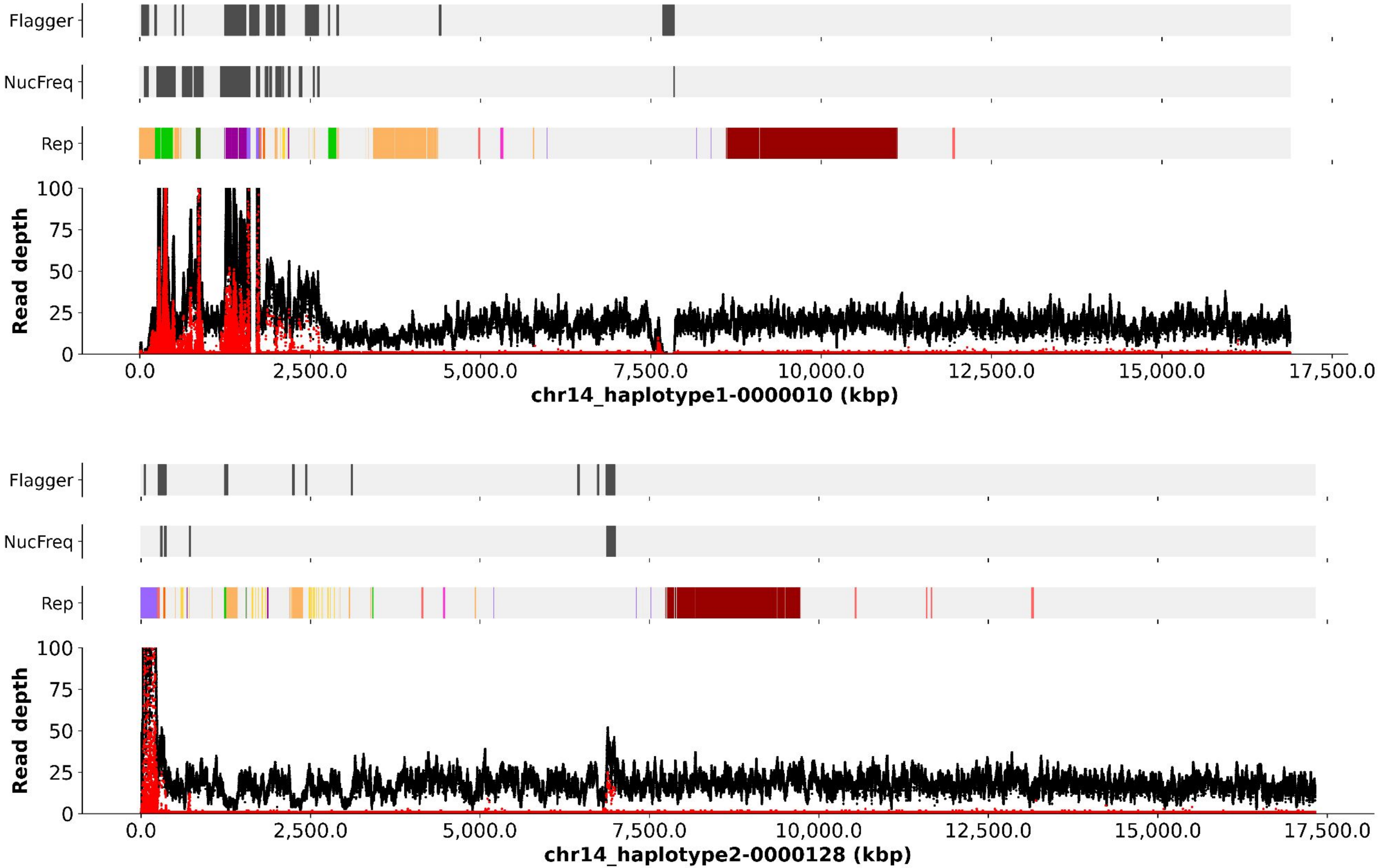
Flagger
NucFreq
Rep
Read depth
100
75
50
25
0
0.0
2,500.0
5,000.0
7,500.0
10,000.0
12,500.0
15,000.0
17,500.0
chr14_haplotype1-0000010 (kbp)
Flagger
NucFreq
Rep
Read depth
chr14_haplotype2-0000128 (kbp)

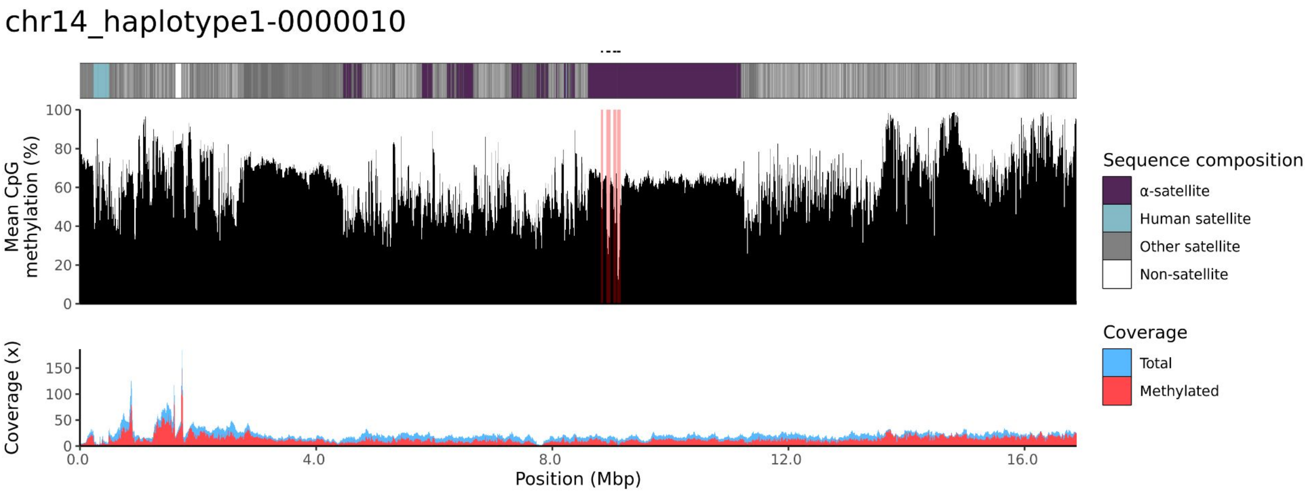
chr14_haplotype1-0000010
Mean CpG methylation (%)
100
80
60
40
20
0
Coverage (x)
150
100
50
0
0.0
4.0
8.0
12.0
16.0
Position (Mbp)
Sequence composition
α-satellite
Human satellite
Other satellite
Non-satellite
Coverage
Total
Methylated

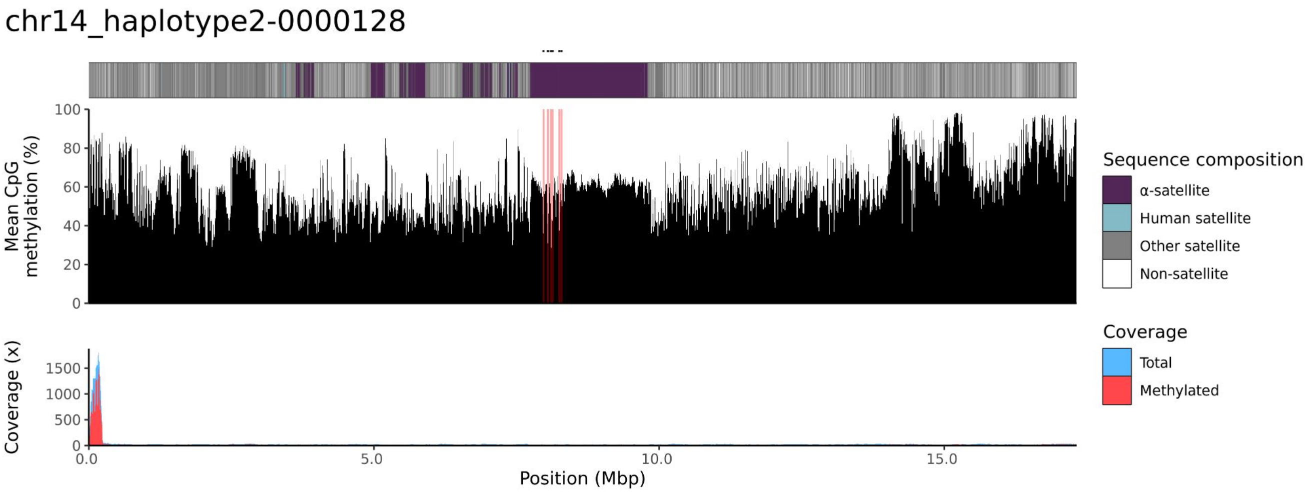
chr14_haplotype2-0000128
Mean CpG methylation (%)
100
80
60
40
20
0
Coverage (x)
1500
1000
500
0
0.0
5.0
10.0
15.0
Position (Mbp)
Sequence composition
α-satellite
Human satellite
Other satellite
Non-satellite
Coverage
Total
Methylated

### chr15

#### K200082_1_haplotype1-0000028_chr15

#### K200082_2_haplotype2-0000146_chr15

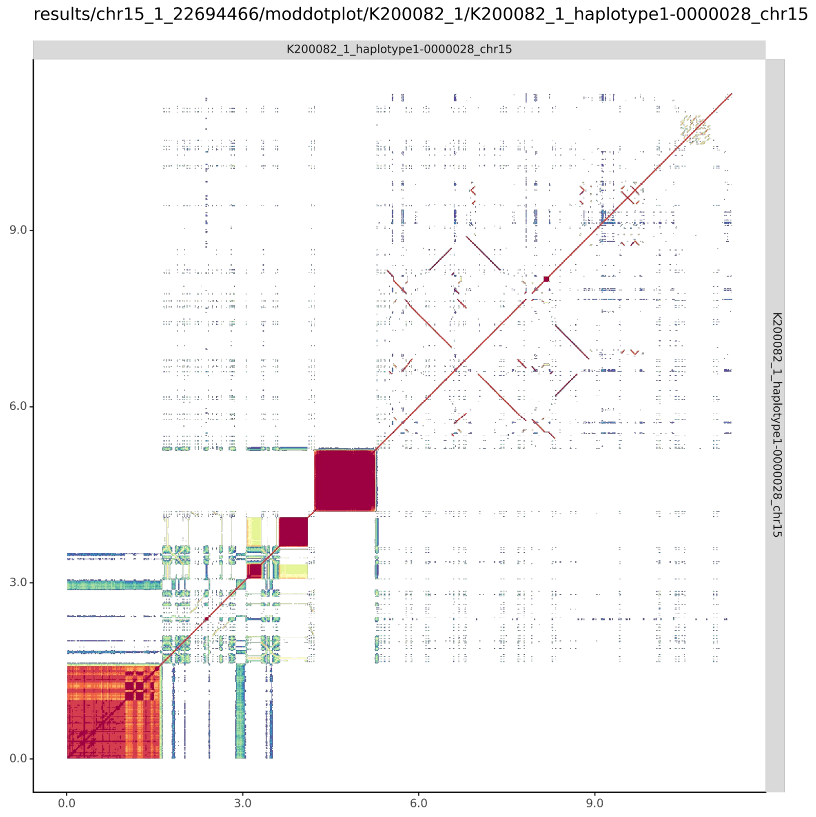


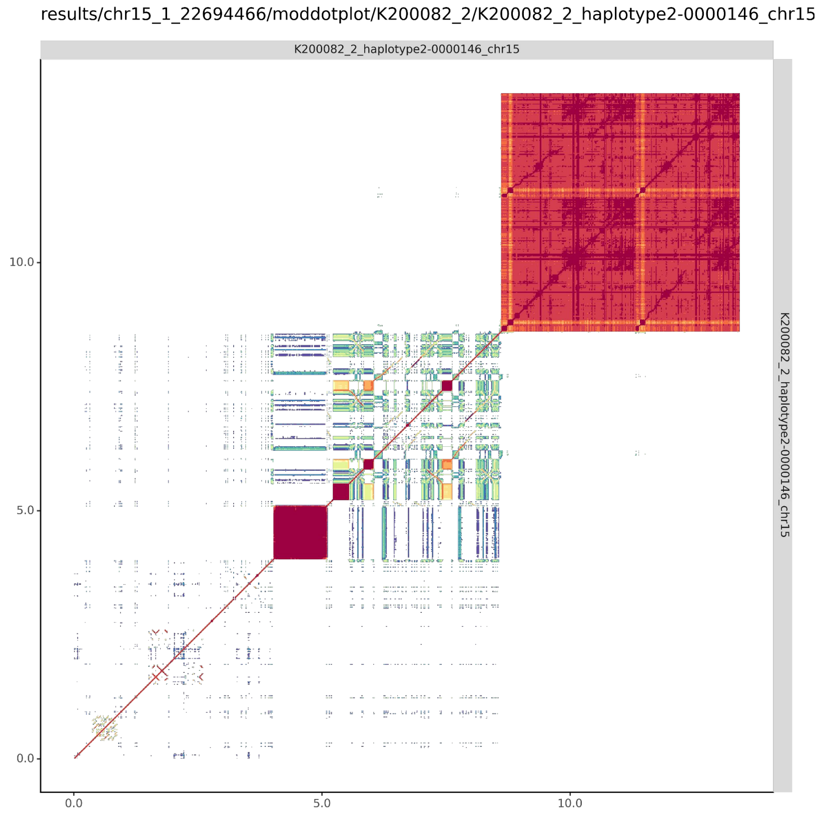

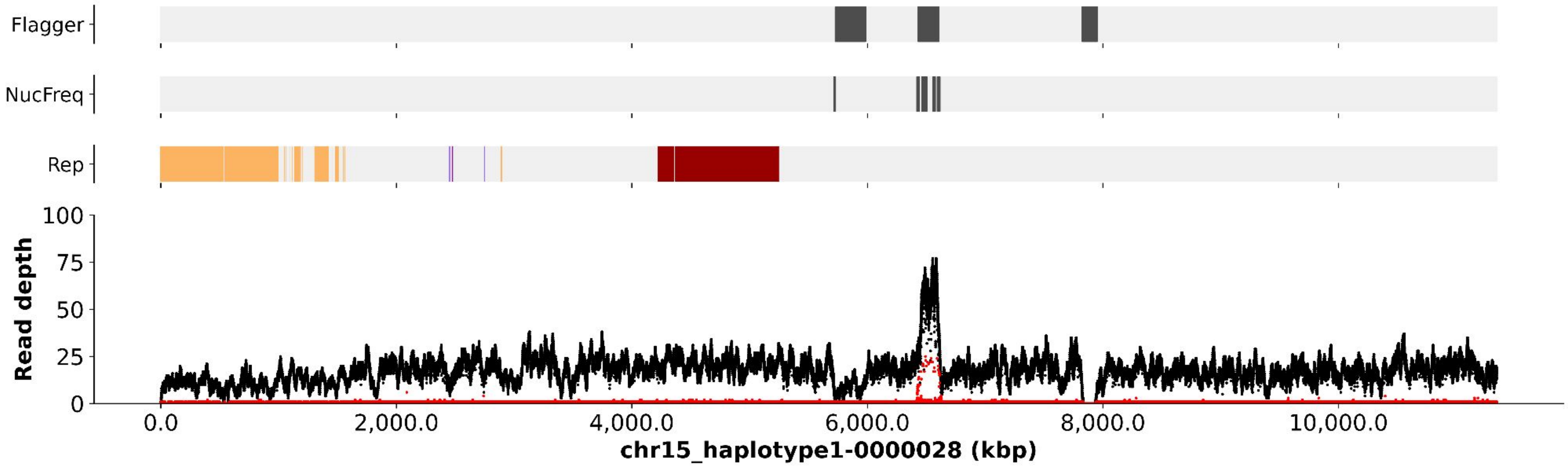

Flagger
NucFreq
Rep
Read depth
100
75
50
25
0
0.0
2,000.0
4,000.0
6,000.0
8,000.0
10,000.0
chr15_haplotype1-0000028 (kbp)


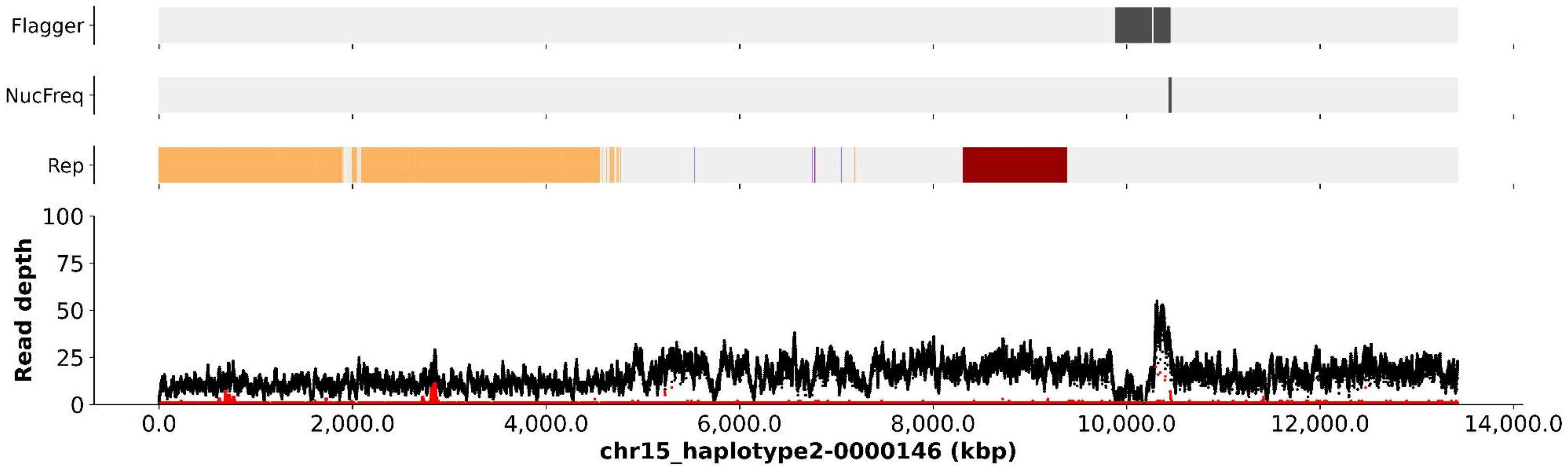

Flagger
NucFreq
Rep
Read depth
100
75
50
25
0
0.0
2,000.0
4,000.0
6,000.0
8,000.0
10,000.0
12,000.0
14,000.0
chr15_haplotype2-0000146 (kbp)

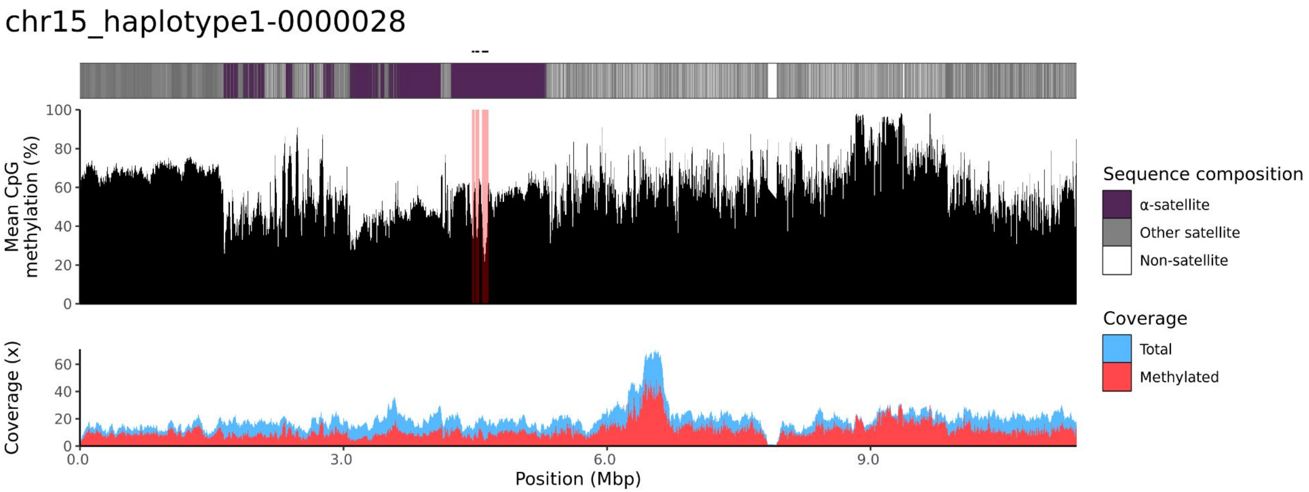

chr15_haplotype1-0000028
Mean CpG methylation (%)
100
80
60
40
20
0
Coverage (x)
60
40
20
0
0.0
3.0
6.0
9.0
Position (Mbp)
Sequence composition
α-satellite
Other satellite
Non-satellite
Coverage
Total
Methylated


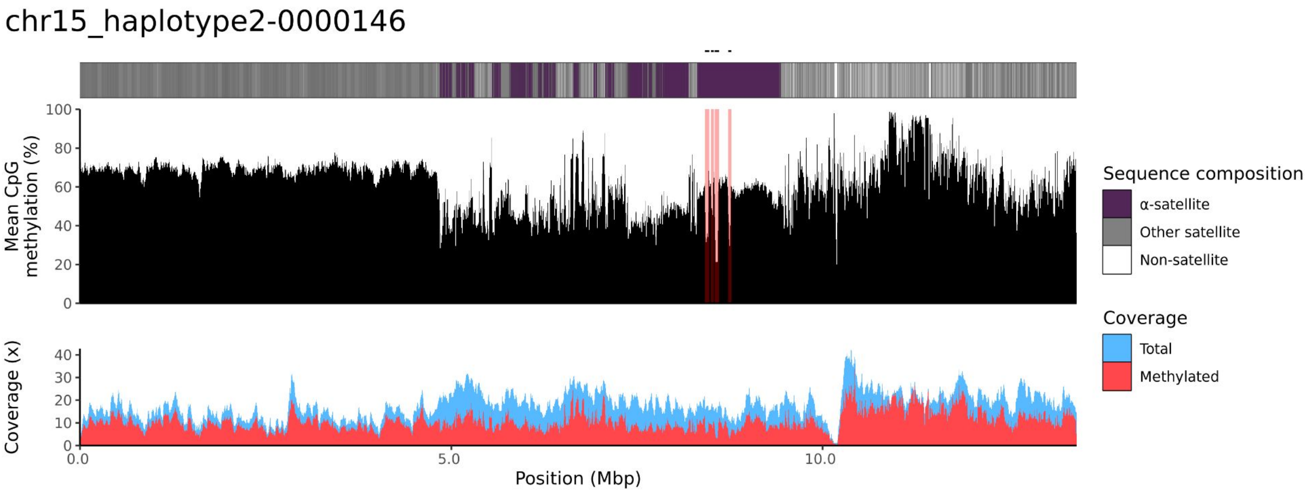

chr15_haplotype2-0000146
Mean CpG methylation (%)
100
80
60
40
20
0
Coverage (x)
40
30
20
10
0
0.0
5.0
10.0
Position (Mbp)
Sequence composition
α-satellite
Other satellite
Non-satellite
Coverage
Total
Methylated

### chr21

#### K200082_1_haplotype1-0000020_chr21

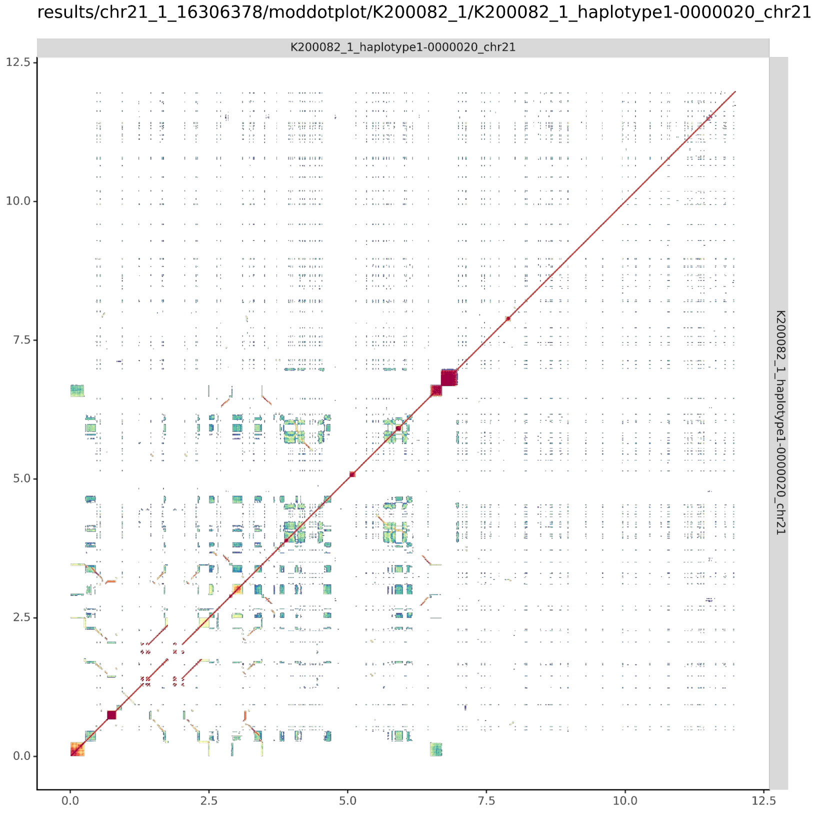


#### K200082_2_haplotype2-0000136_chr21

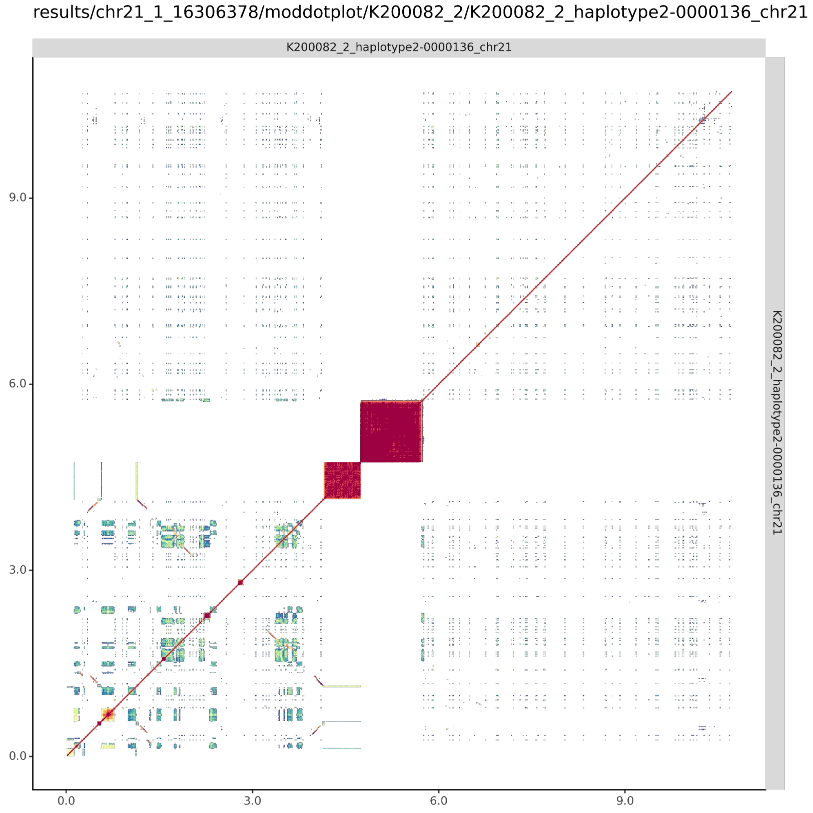

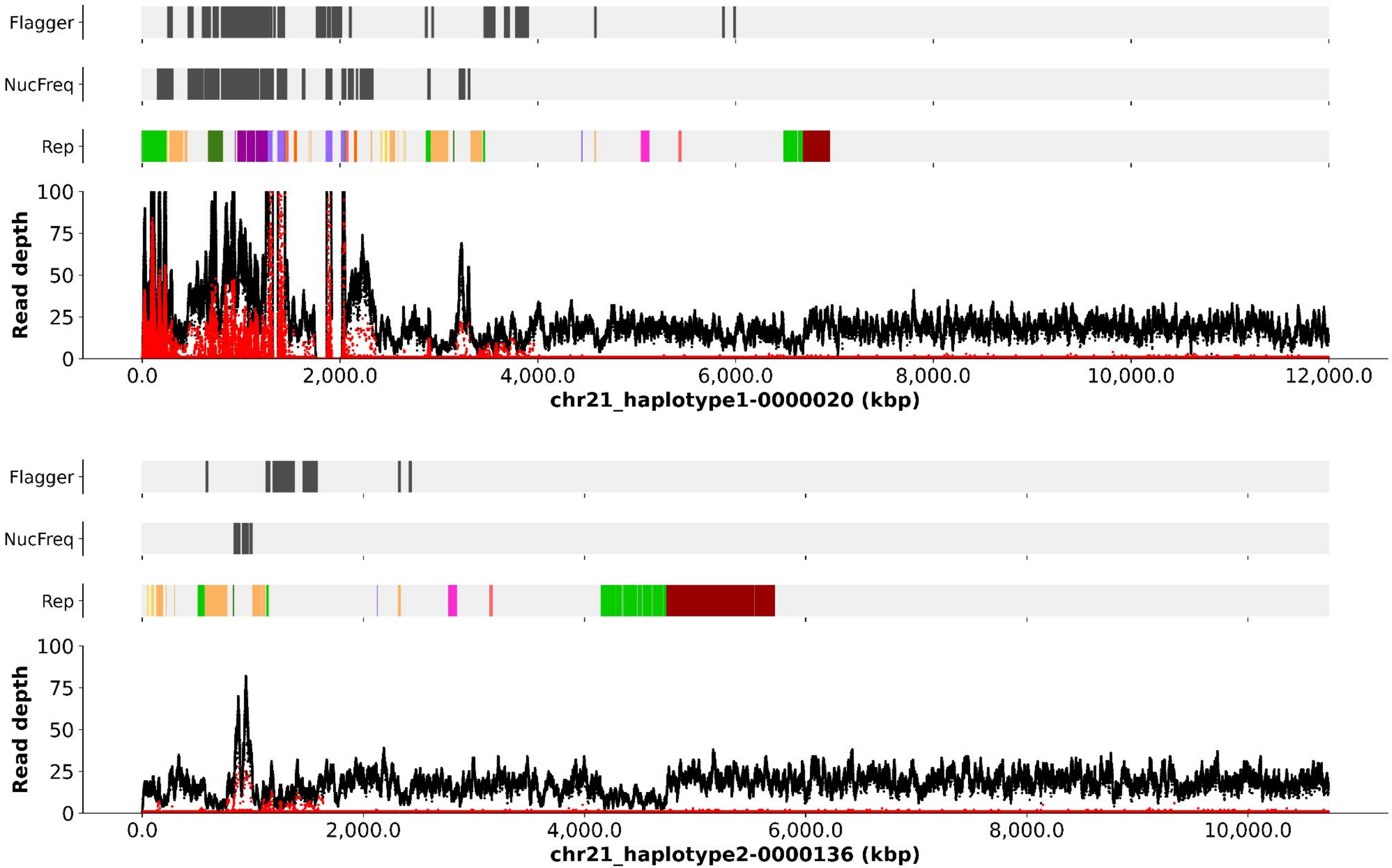

Flagger
NucFreq
Rep
Read depth
100
75
50
25
0
0.0
2,000.0
4,000.0
6,000.0
8,000.0
10,000.0
12,000.0
chr21_haplotype1-0000020 (kbp)
Flagger
NucFreq
Rep
Read depth
100
75
50
25
0
0.0
2,000.0
4,000.0
6,000.0
8,000.0
10,000.0
chr21_haplotype2-0000136 (kbp)

#### chr21_haplotype1-0000020

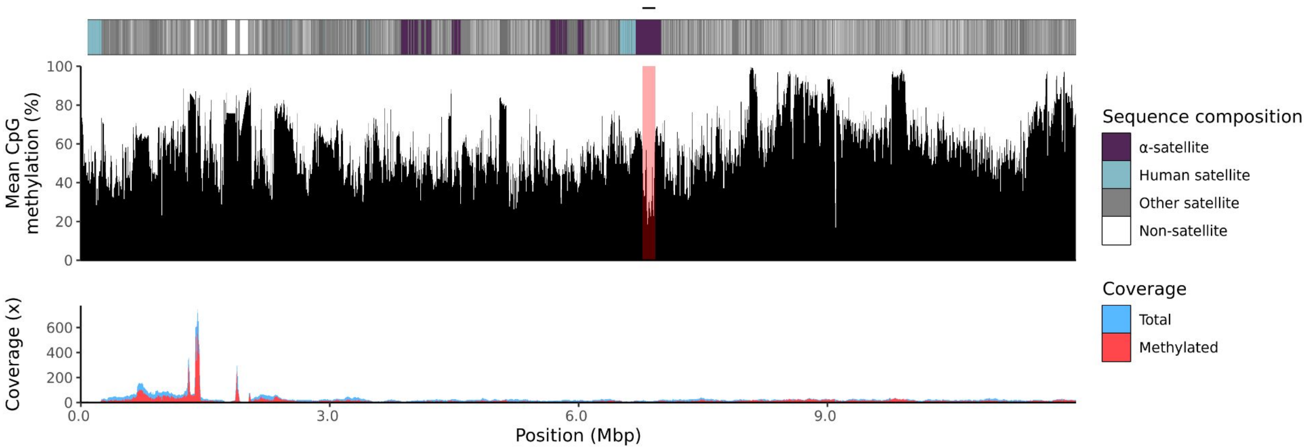


#### chr21_haplotype2-0000136

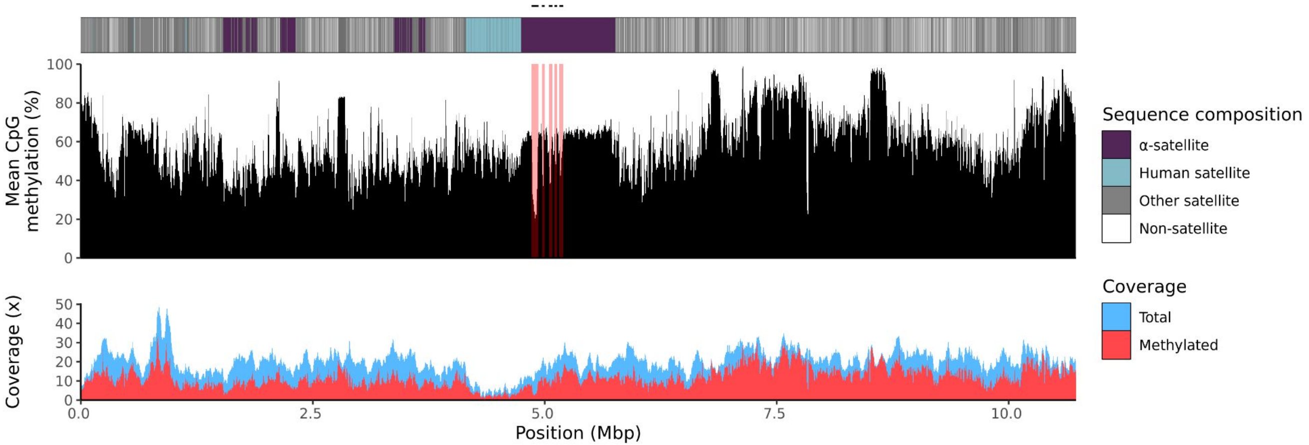

### chr22

#### K200082_1_haplotype1-0000003_chr22

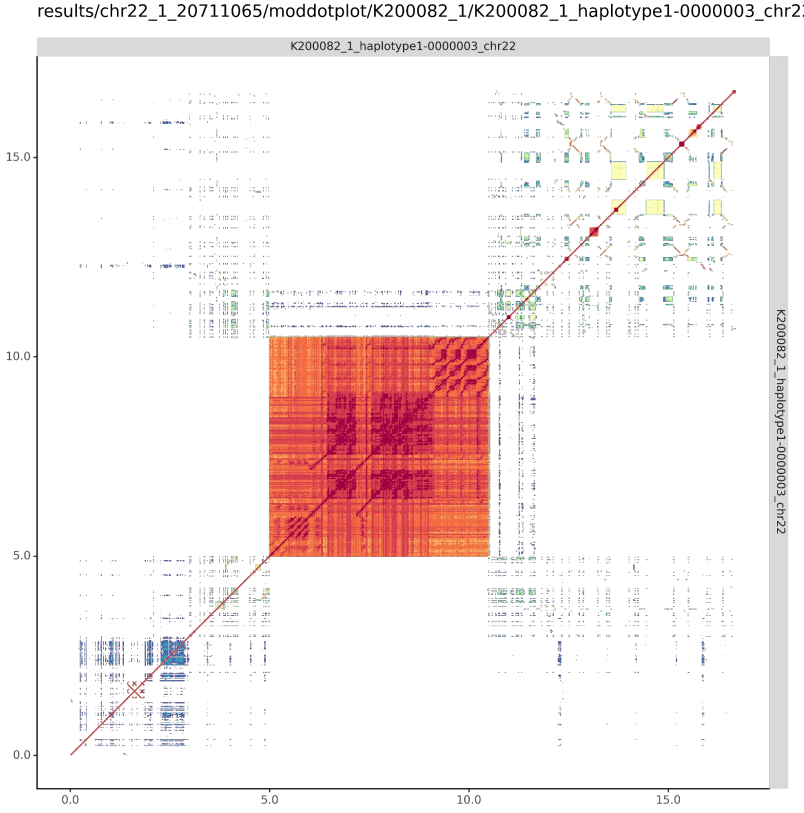


#### K200082_2_haplotype2-0000121_chr22

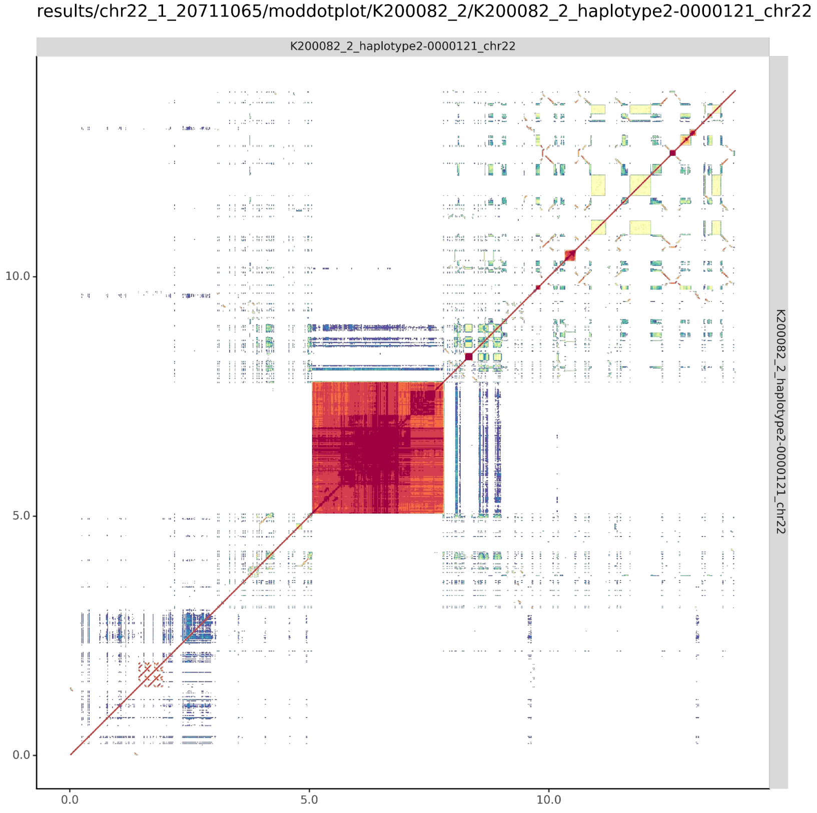

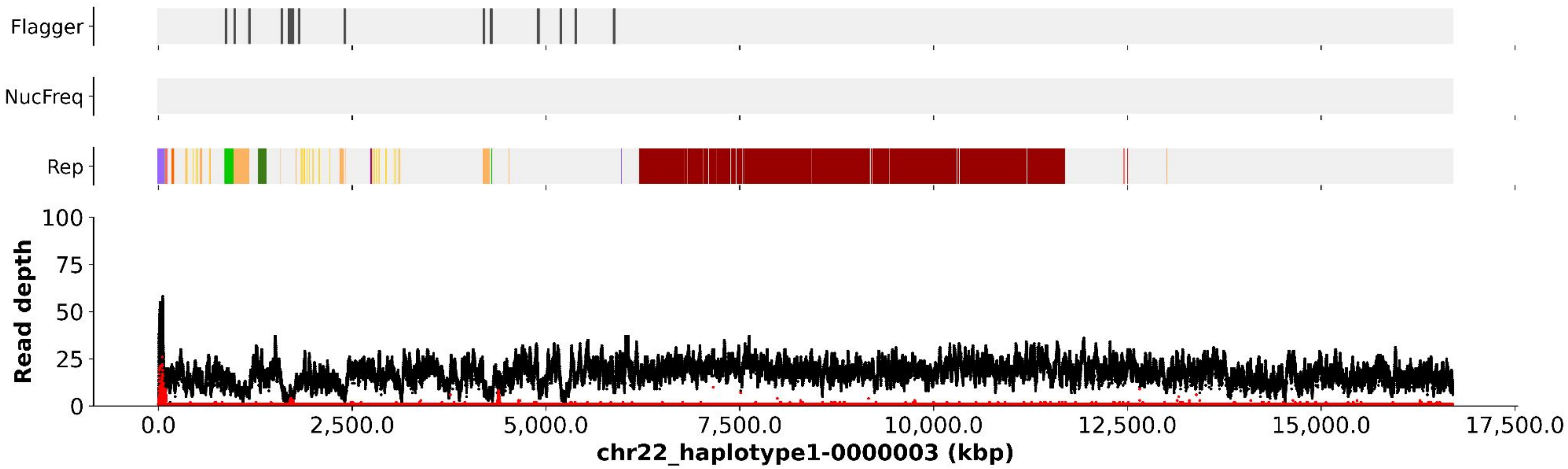
Flagger
NucFreq
Rep
Read depth
100
75
50
25
0
0.0
2,500.0
5,000.0
7,500.0
10,000.0
12,500.0
15,000.0
17,500.0
chr22_haplotype1-0000003 (kbp)

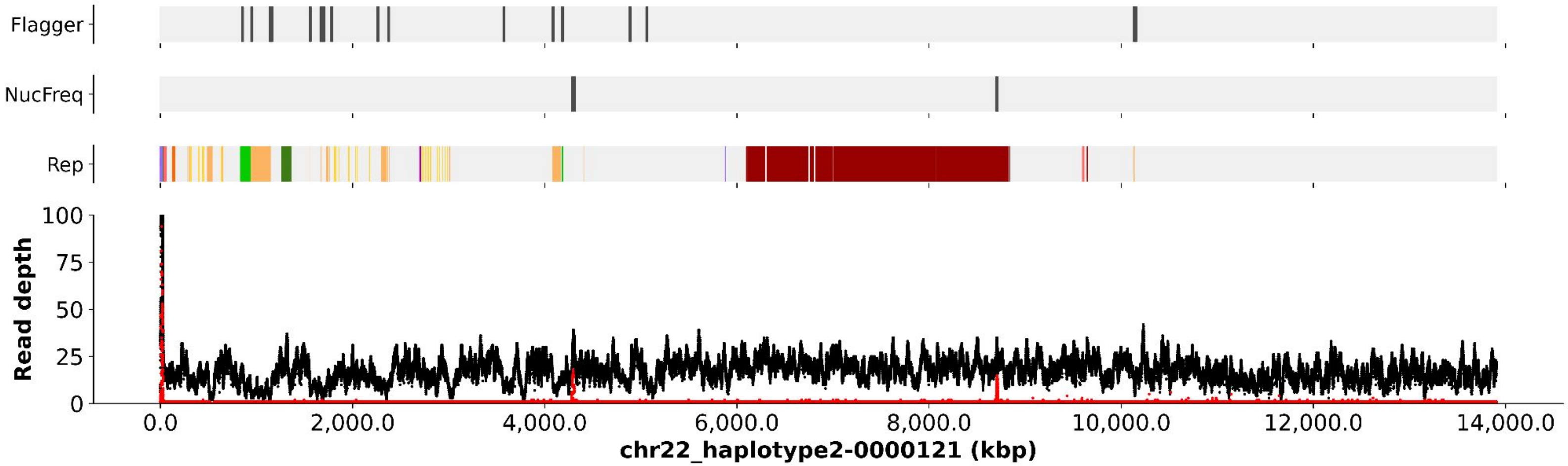
Flagger
NucFreq
Rep
Read depth
100
75
50
25
0
0.0
2,000.0
4,000.0
6,000.0
8,000.0
10,000.0
12,000.0
14,000.0
chr22_haplotype2-0000121 (kbp)

chr22_haplotype1-0000003

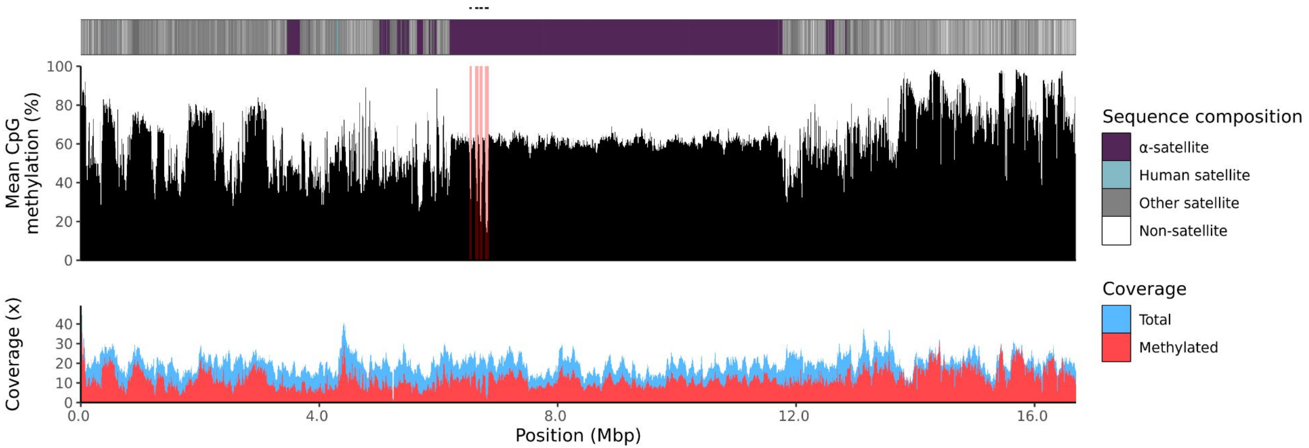


chr22_haplotype2-0000121

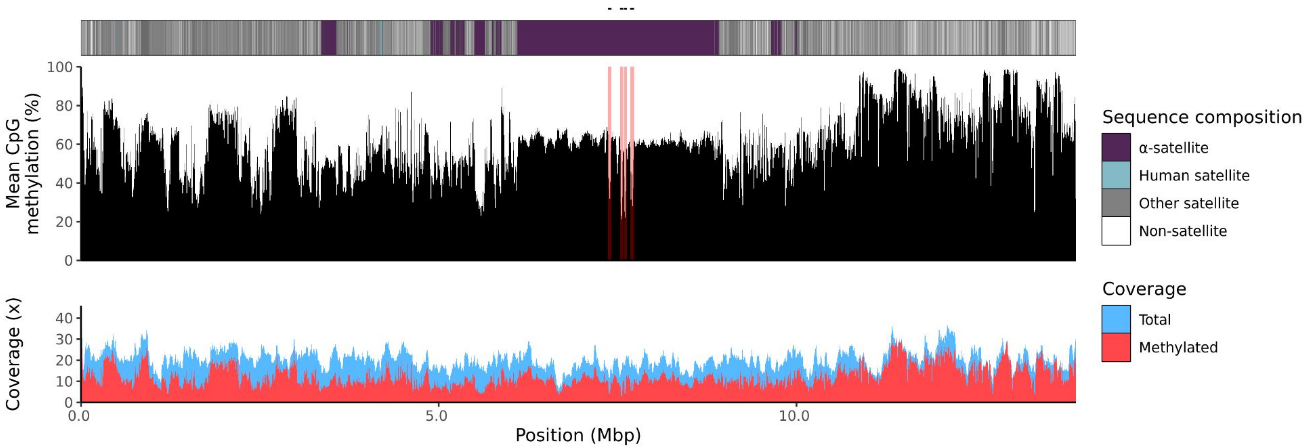

200084

### chr13

#### K200084_1_haplotype1-0000019_chr13

#### K200084_2_haplotype2-0000083_chr13

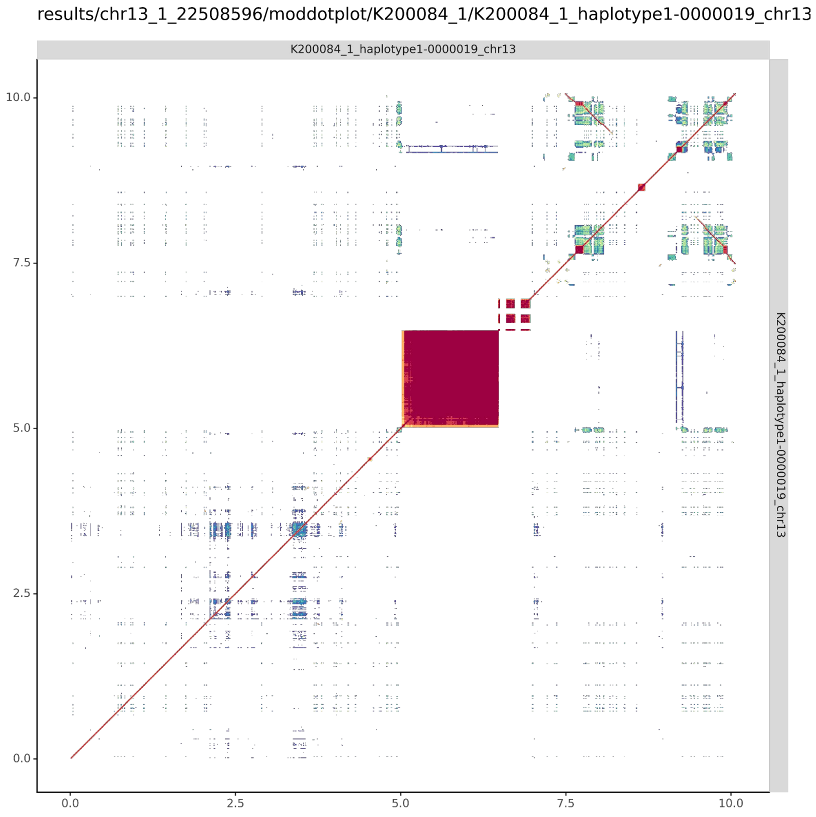


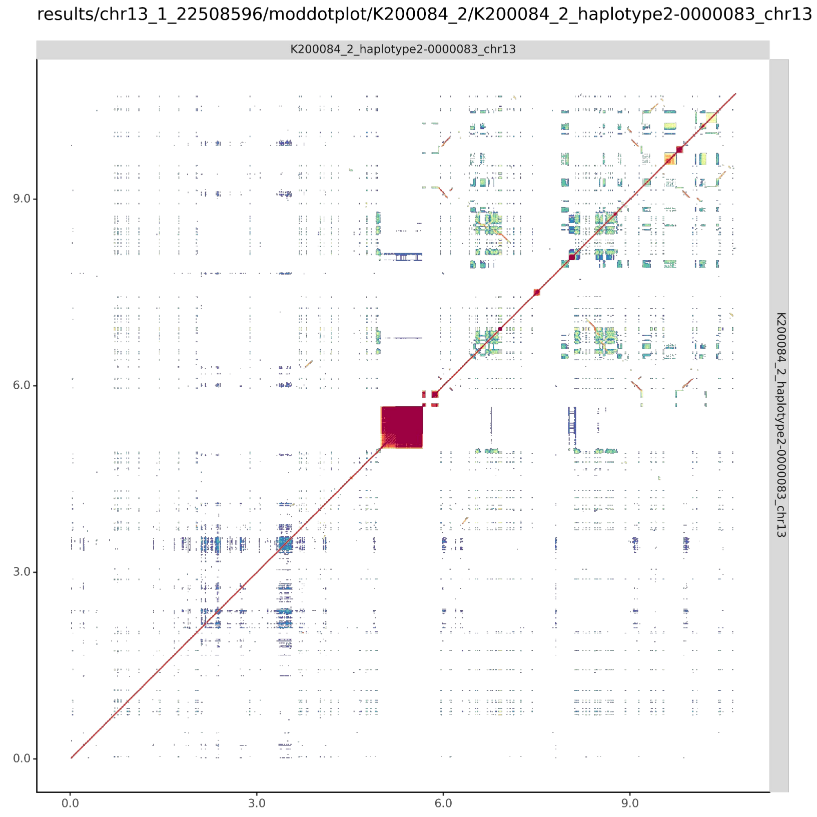

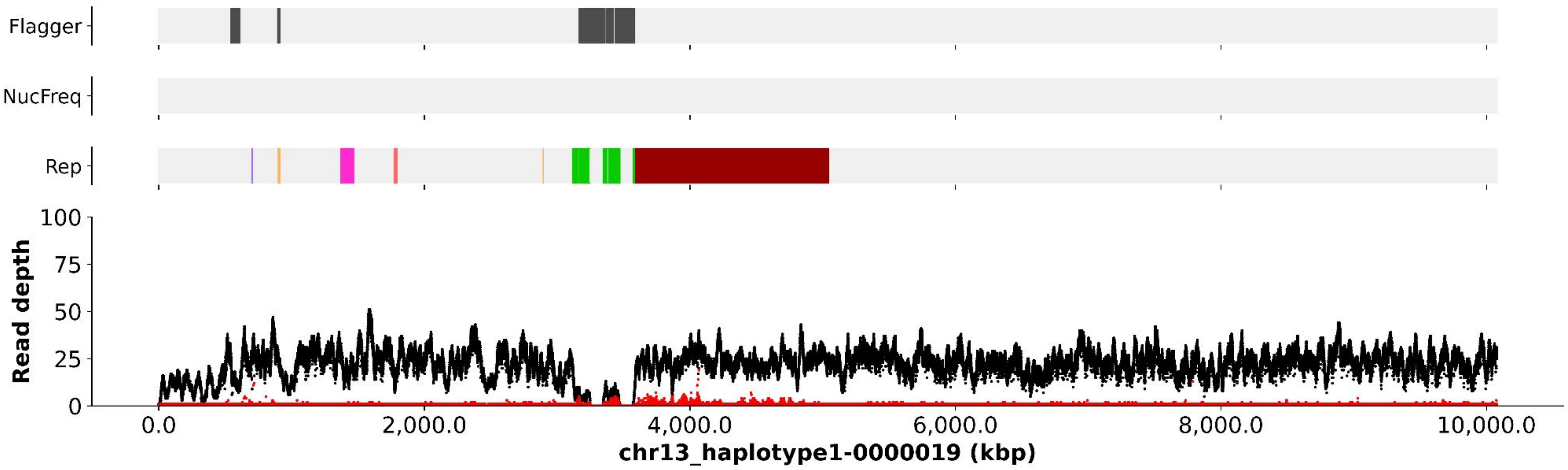
Flagger
NucFreq
Rep
100
75
50
25
0
Read depth
0.0
2,000.0
4,000.0
6,000.0
8,000.0
10,000.0
chr13_haplotype1-0000019 (kbp)

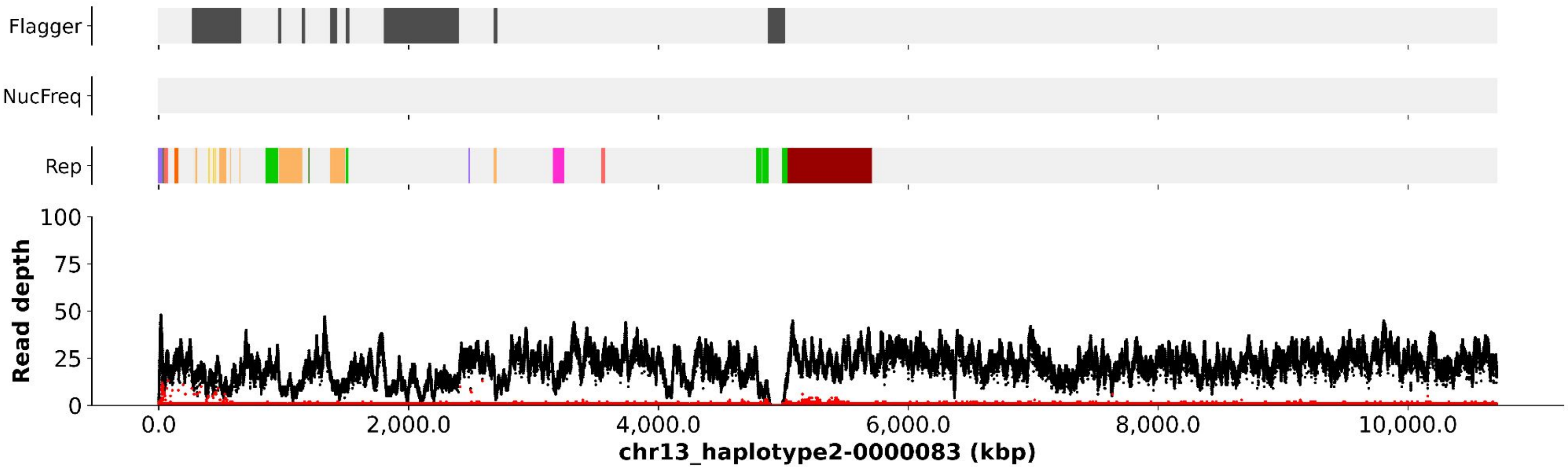
Flagger
NucFreq
Rep
100
75
50
25
0
Read depth
0.0
2,000.0
4,000.0
6,000.0
8,000.0
10,000.0
chr13_haplotype2-0000083 (kbp)

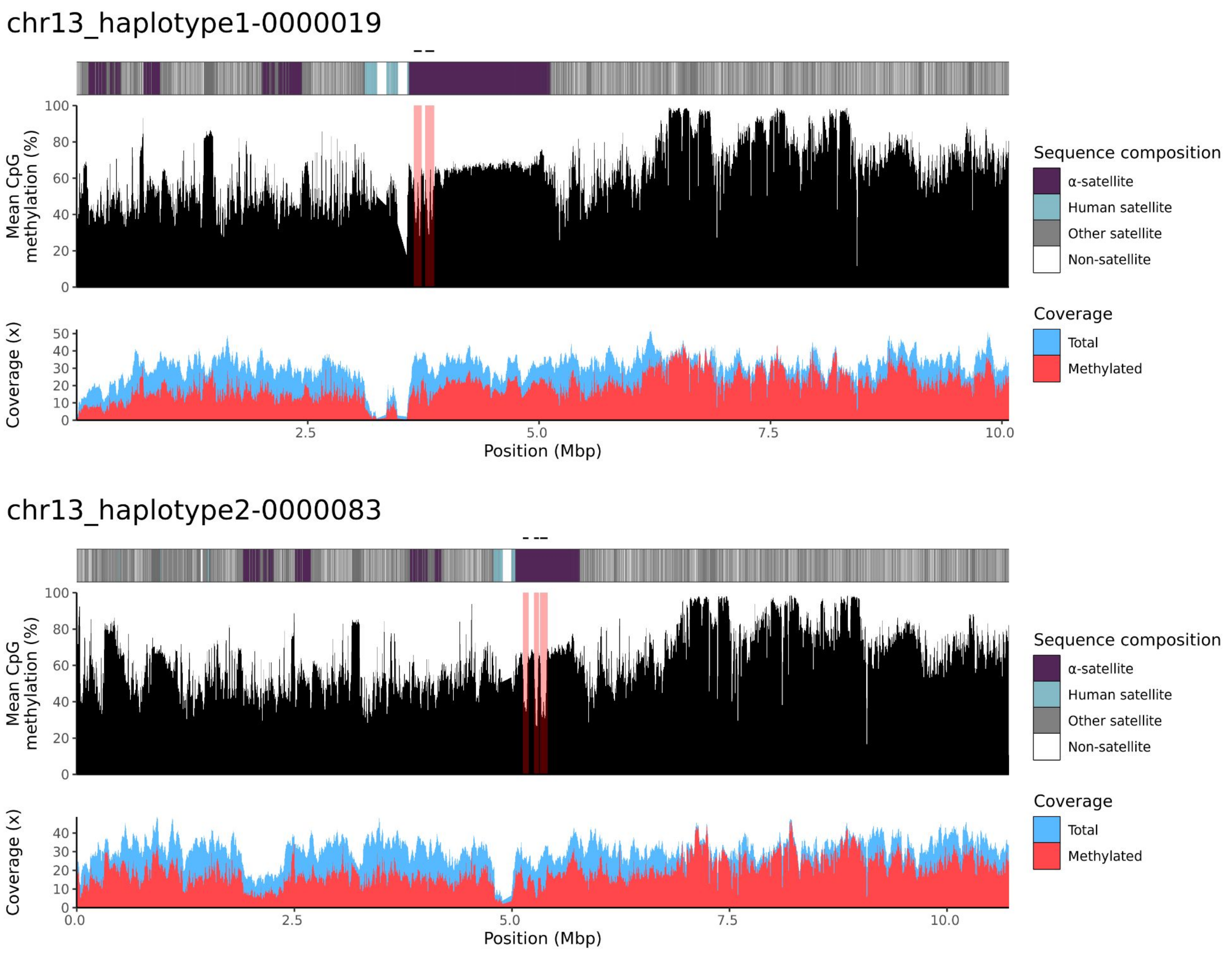

chr13_haplotype1-0000019
Mean CpG methylation (%)
Coverage (x)
Position (Mbp)
Sequence composition
α-satellite
Human satellite
Other satellite
Non-satellite
Coverage
Total
Methylated
chr13_haplotype2-0000083
Mean CpG methylation (%)
Coverage (x)
Position (Mbp)
Sequence composition
α-satellite
Human satellite
Other satellite
Non-satellite
Coverage
Total
Methylated

### chr14

#### K200084_1_haplotype1-0000018_chr14

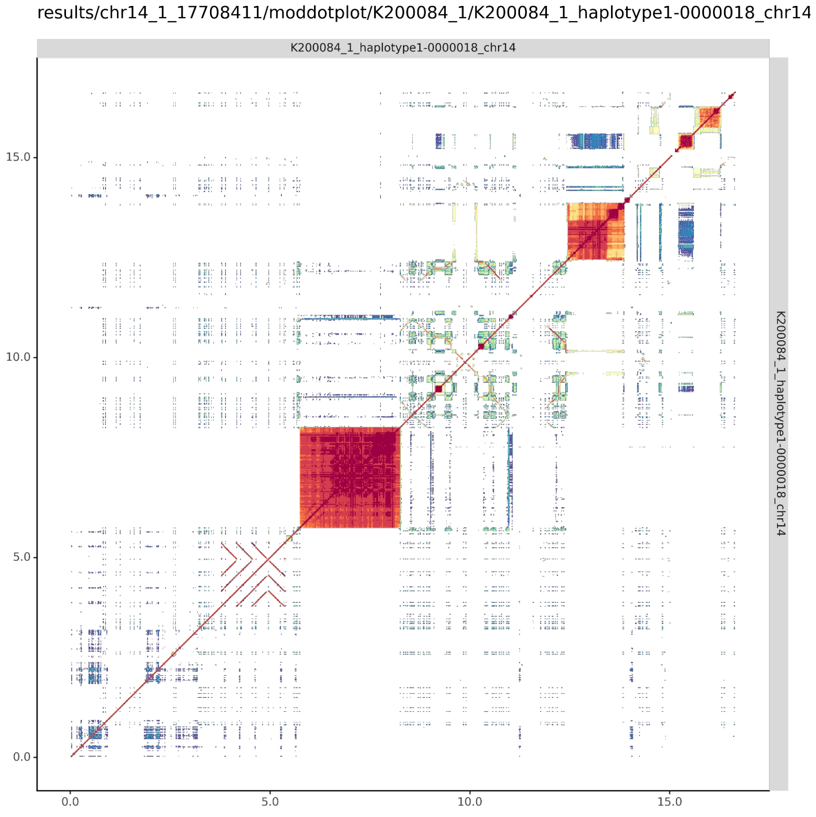


#### K200084_2_haplotype2-0000078_chr14

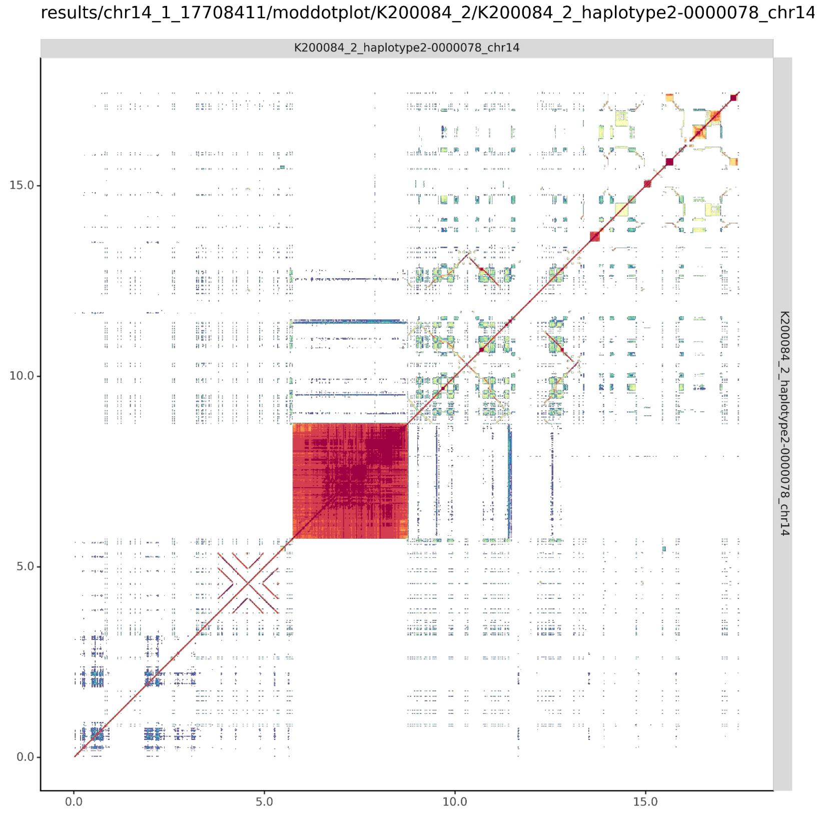

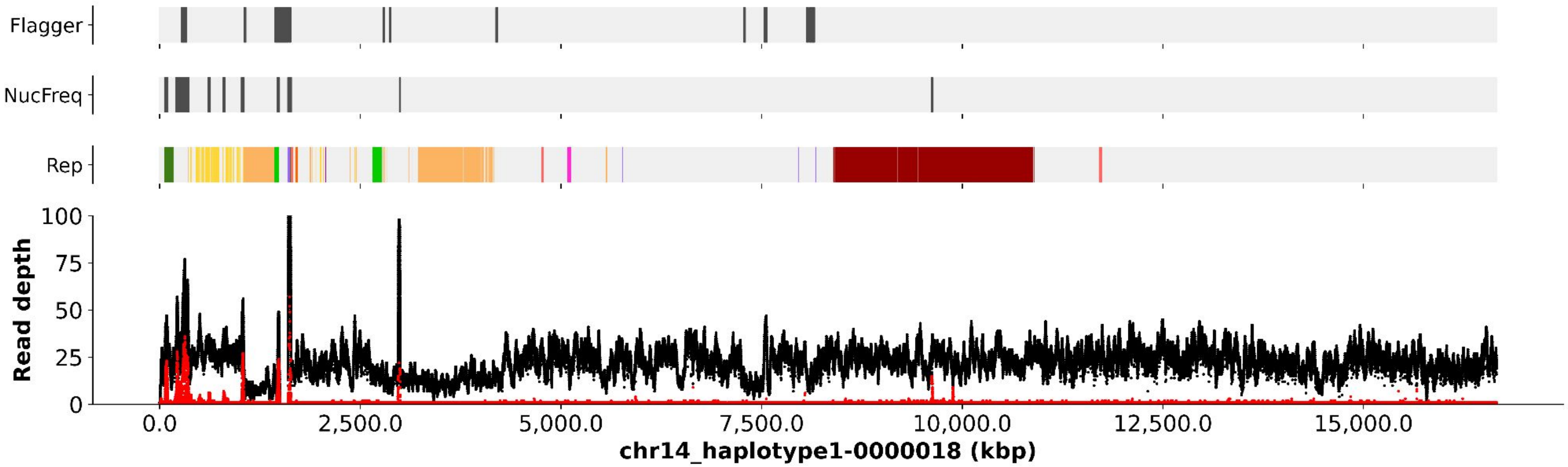
Flagger
NucFreq
Rep
Read depth
100
75
50
25
0
0.0
2,500.0
5,000.0
7,500.0
10,000.0
12,500.0
15,000.0
chr14_haplotype1-0000018 (kbp)

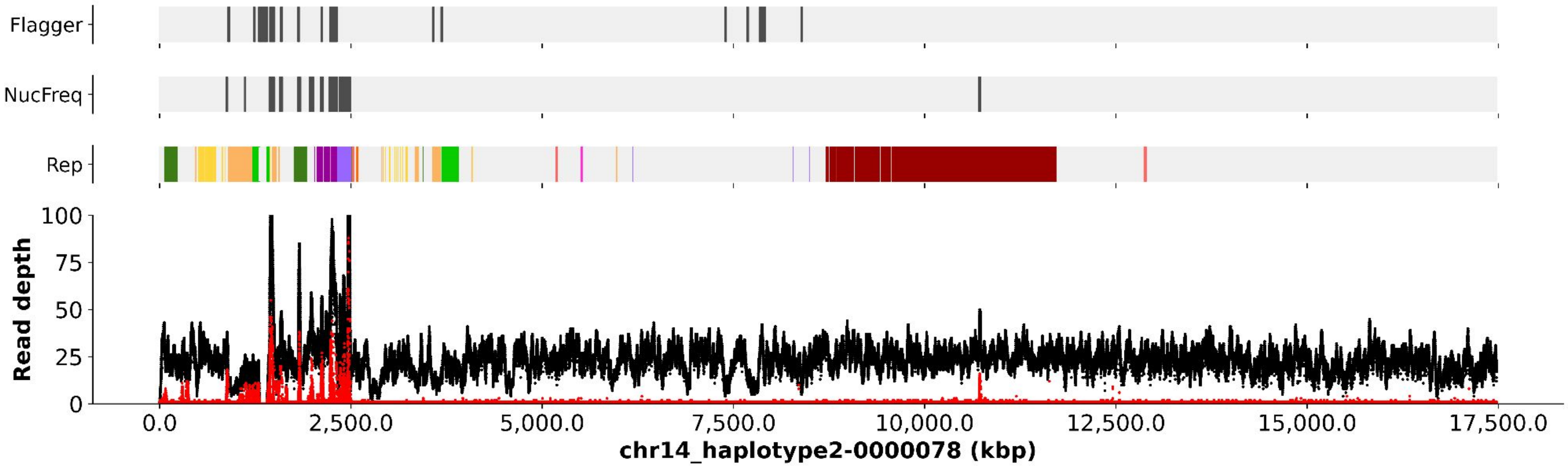
Flagger
NucFreq
Rep
Read depth
100
75
50
25
0
0.0
2,500.0
5,000.0
7,500.0
10,000.0
12,500.0
15,000.0
17,500.0
chr14_haplotype2-0000078 (kbp)

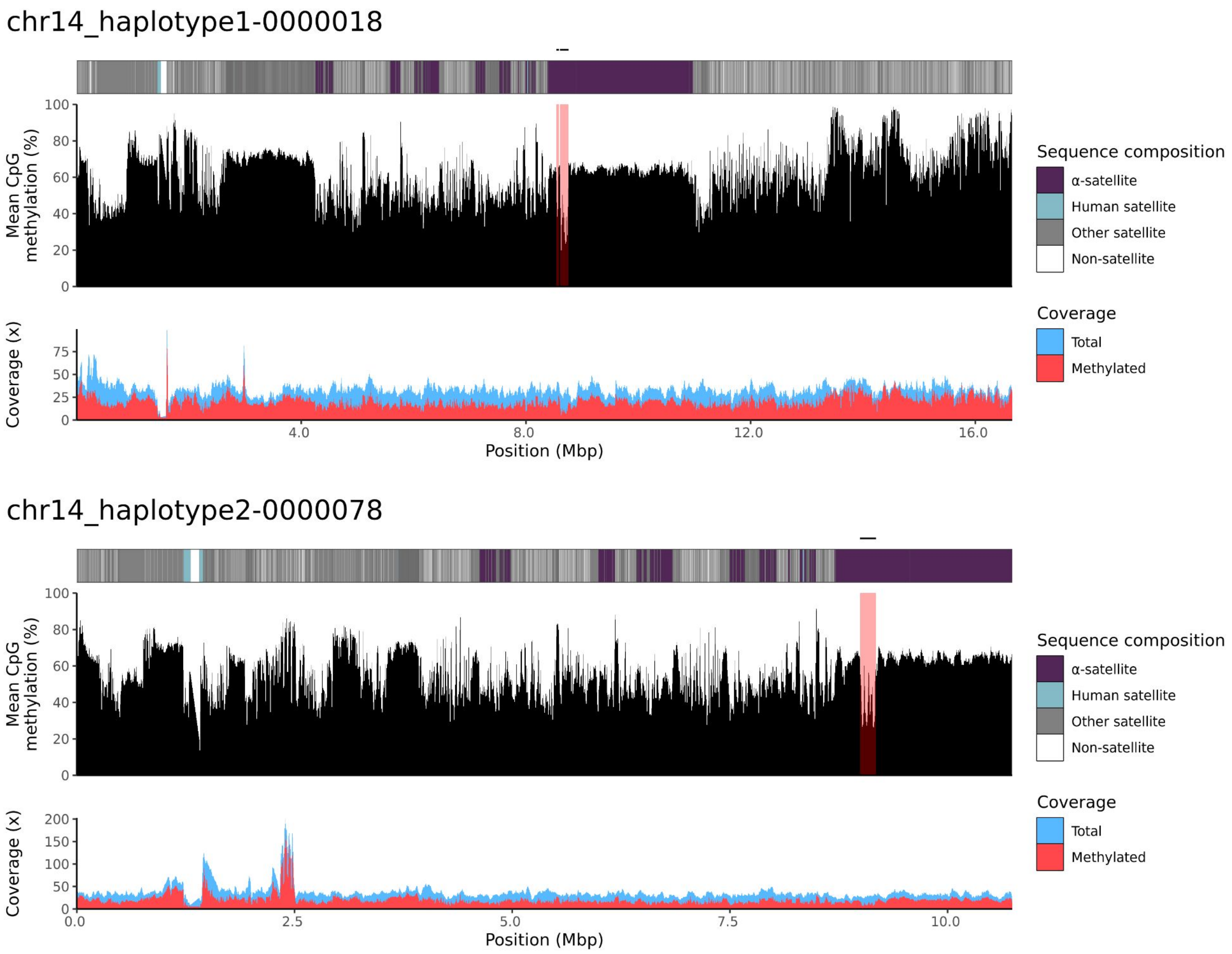
chr14_haplotype1-0000018
Mean CpG methylation (%)
100
80
60
40
20
0
Coverage (x)
75
50
25
0
4.0
8.0
12.0
16.0
Position (Mbp)
Sequence composition
α-satellite
Human satellite
Other satellite
Non-satellite
Coverage
Total
Methylated
chr14_haplotype2-0000078
Mean CpG methylation (%)
100
80
60
40
20
0
Coverage (x)
200
150
100
50
0
0.0
2.5
5.0
7.5
10.0
Position (Mbp)
Sequence composition
α-satellite
Human satellite
Other satellite
Non-satellite
Coverage
Total
Methylated

### chr15

#### K200084_1_haplotype1-0000007_chr15

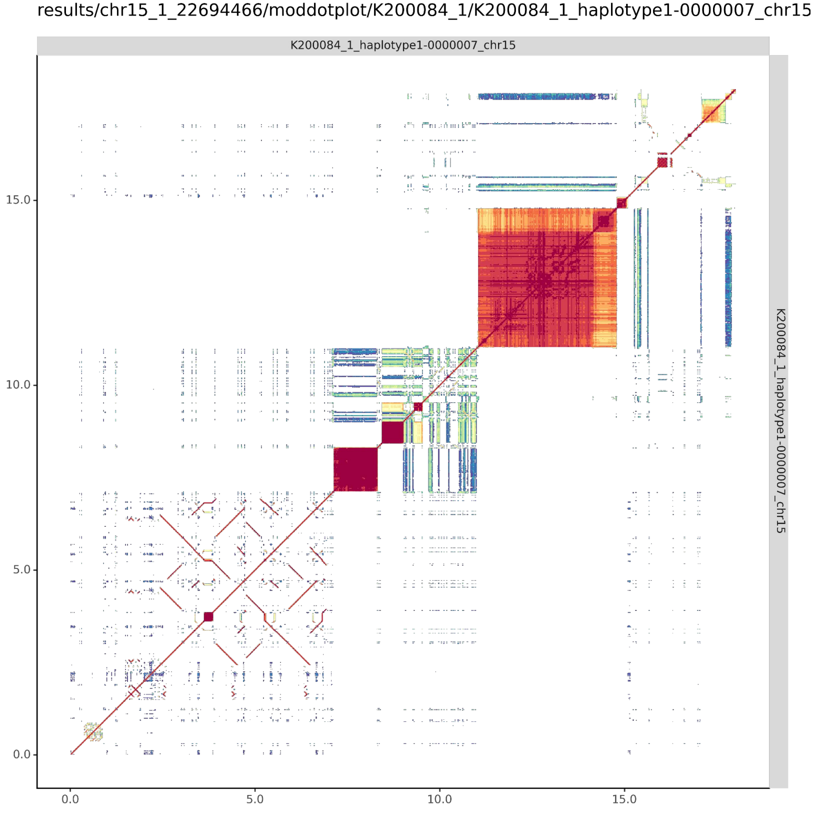


#### K200084_2_haplotype2-0000070_chr15

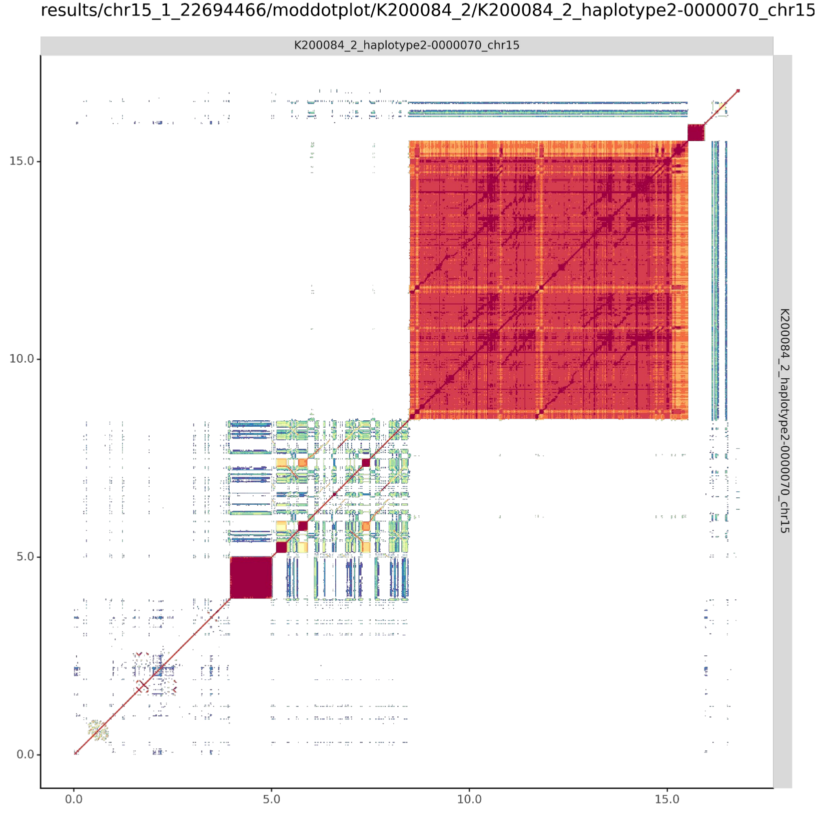

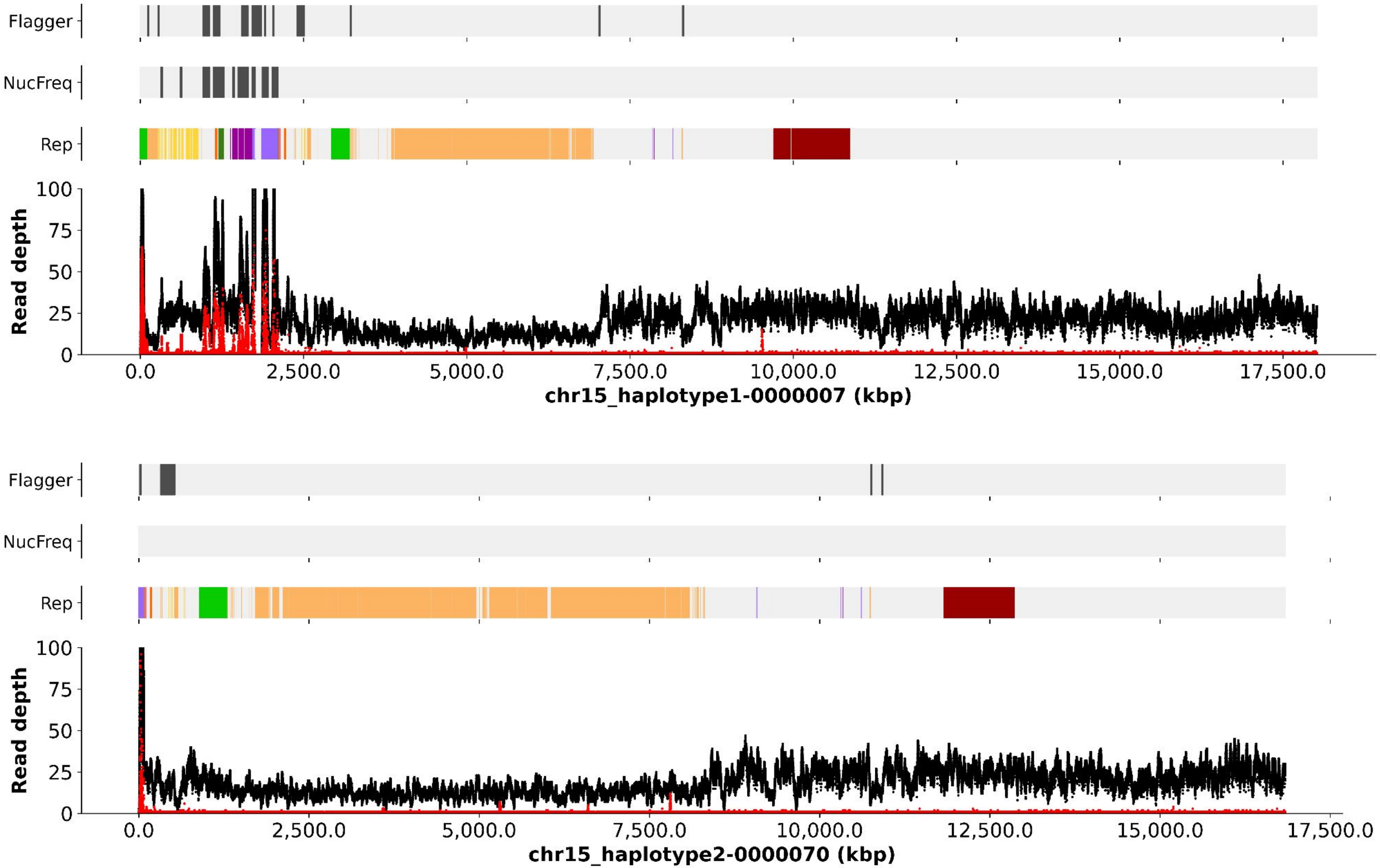

Flagger
NucFreq
Rep
Read depth
100
75
50
25
0
0.0
2,500.0
5,000.0
7,500.0
10,000.0
12,500.0
15,000.0
17,500.0
chr15_haplotype1-0000007 (kbp)
Flagger
NucFreq
Rep
Read depth
100
75
50
25
0
0.0
2,500.0
5,000.0
7,500.0
10,000.0
12,500.0
15,000.0
17,500.0
chr15_haplotype2-0000070 (kbp)

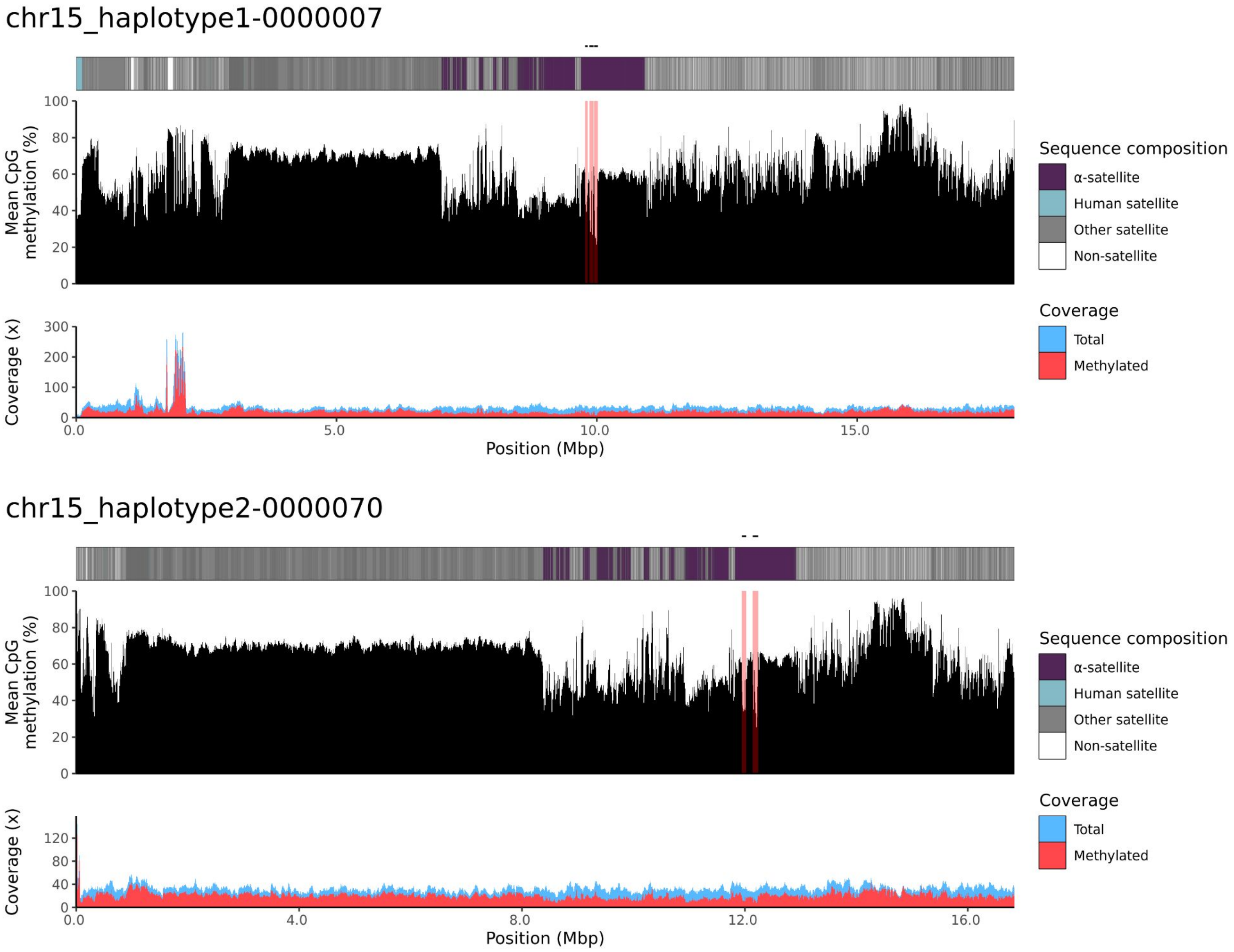
chr15_haplotype1-0000007
Mean CpG methylation (%)
Coverage (x)
Position (Mbp)
0.0
5.0
10.0
15.0
Sequence composition
α-satellite
Human satellite
Other satellite
Non-satellite
Coverage
Total
Methylated
chr15_haplotype2-0000070
Mean CpG methylation (%)
Coverage (x)
Position (Mbp)
0.0
4.0
8.0
12.0
16.0
Sequence composition
α-satellite
Human satellite
Other satellite
Non-satellite
Coverage
Total
Methylated

### chr21

#### K200084_1_haplotype1-0000012_chr21

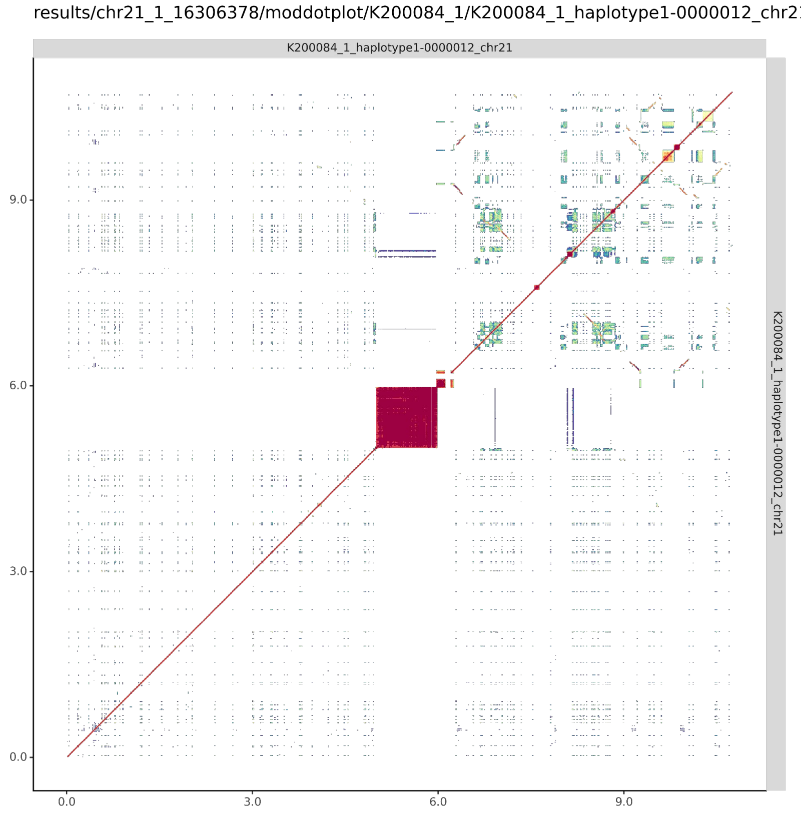


#### K200084_2_haplotype2-0000074_chr21

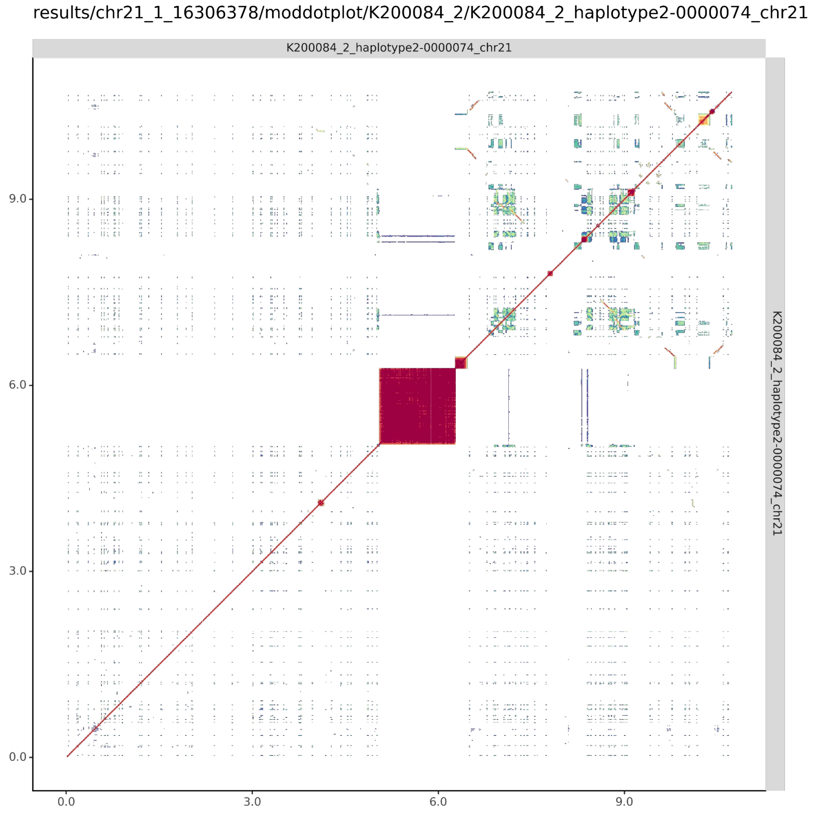

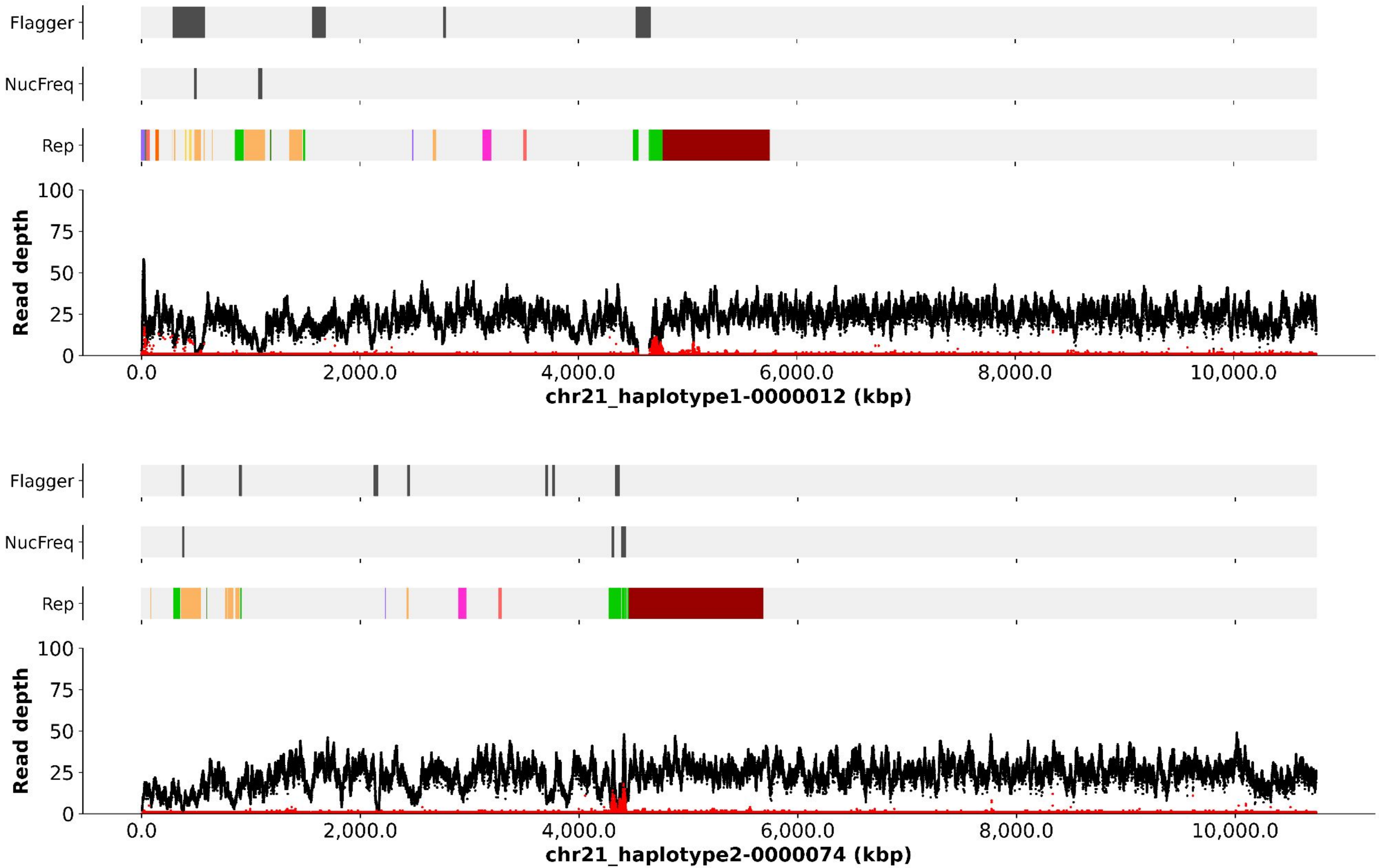
Flagger
NucFreq
Rep
Read depth
100
75
50
25
0
0.0
2,000.0
4,000.0
6,000.0
8,000.0
10,000.0
chr21_haplotype1-0000012 (kbp)
Flagger
NucFreq
Rep
Read depth
100
75
50
25
0
0.0
2,000.0
4,000.0
6,000.0
8,000.0
10,000.0
chr21_haplotype2-0000074 (kbp)

#### chr21_haplotype1-0000012

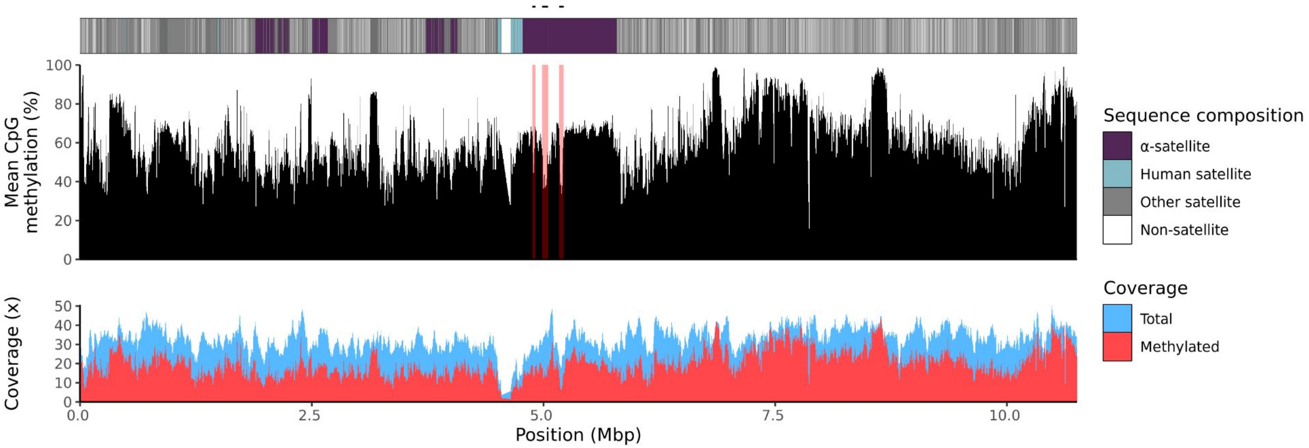


#### chr21_haplotype2-0000074

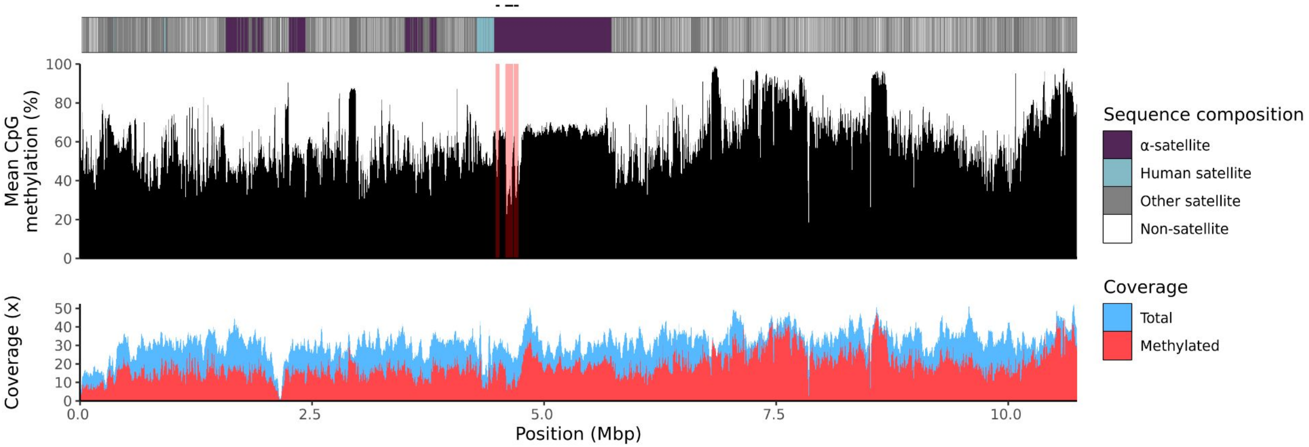

### chr22

#### K200084_1_haplotype1-0000026_chr22

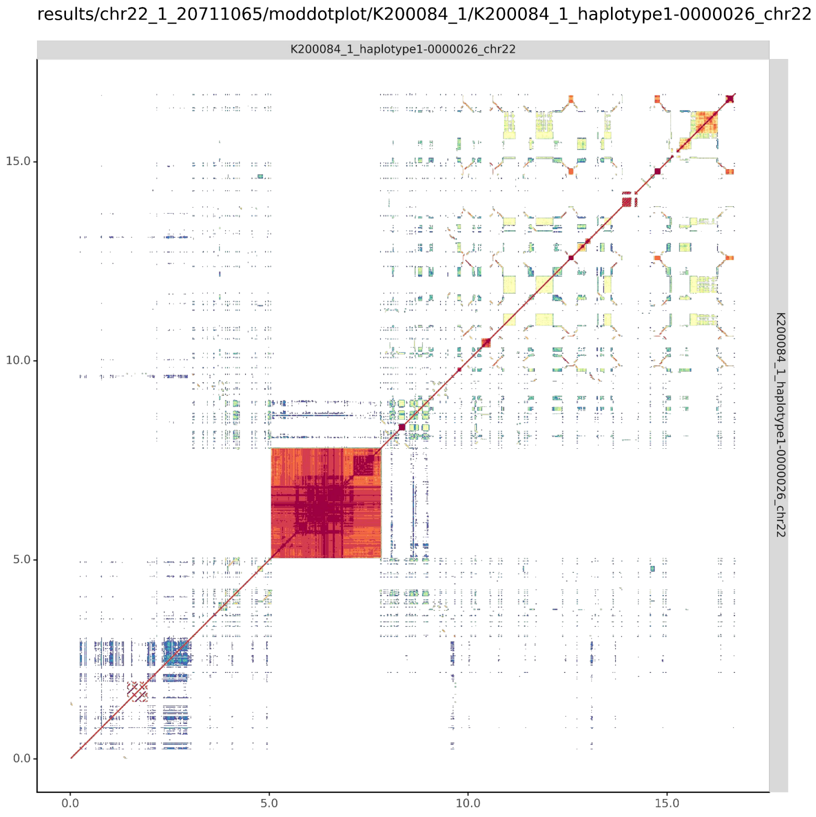


#### K200084_2_haplotype2-0000088_chr22

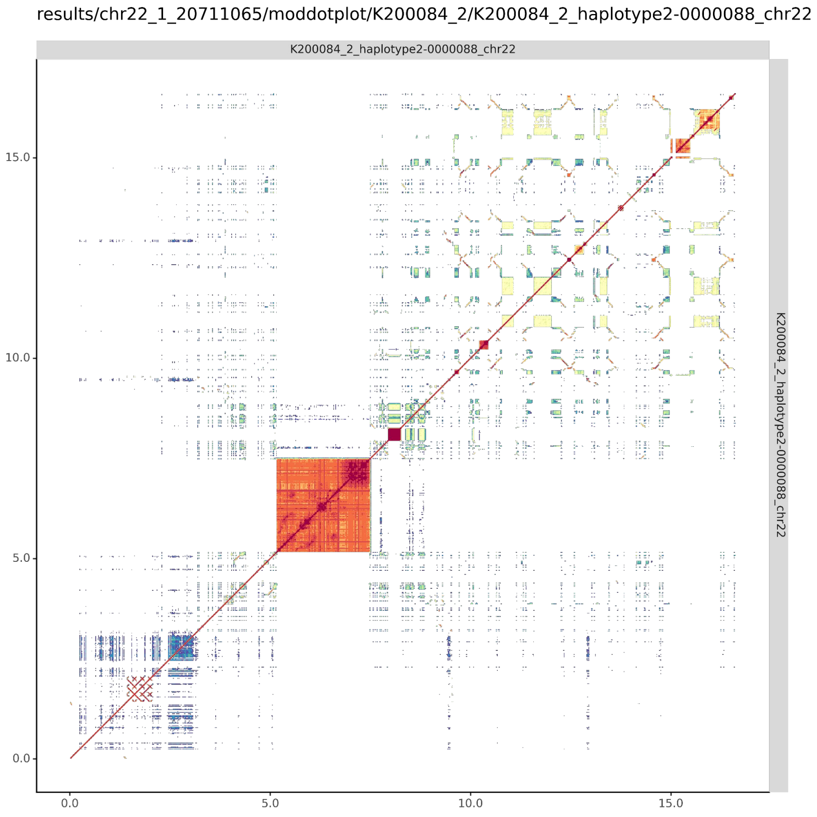

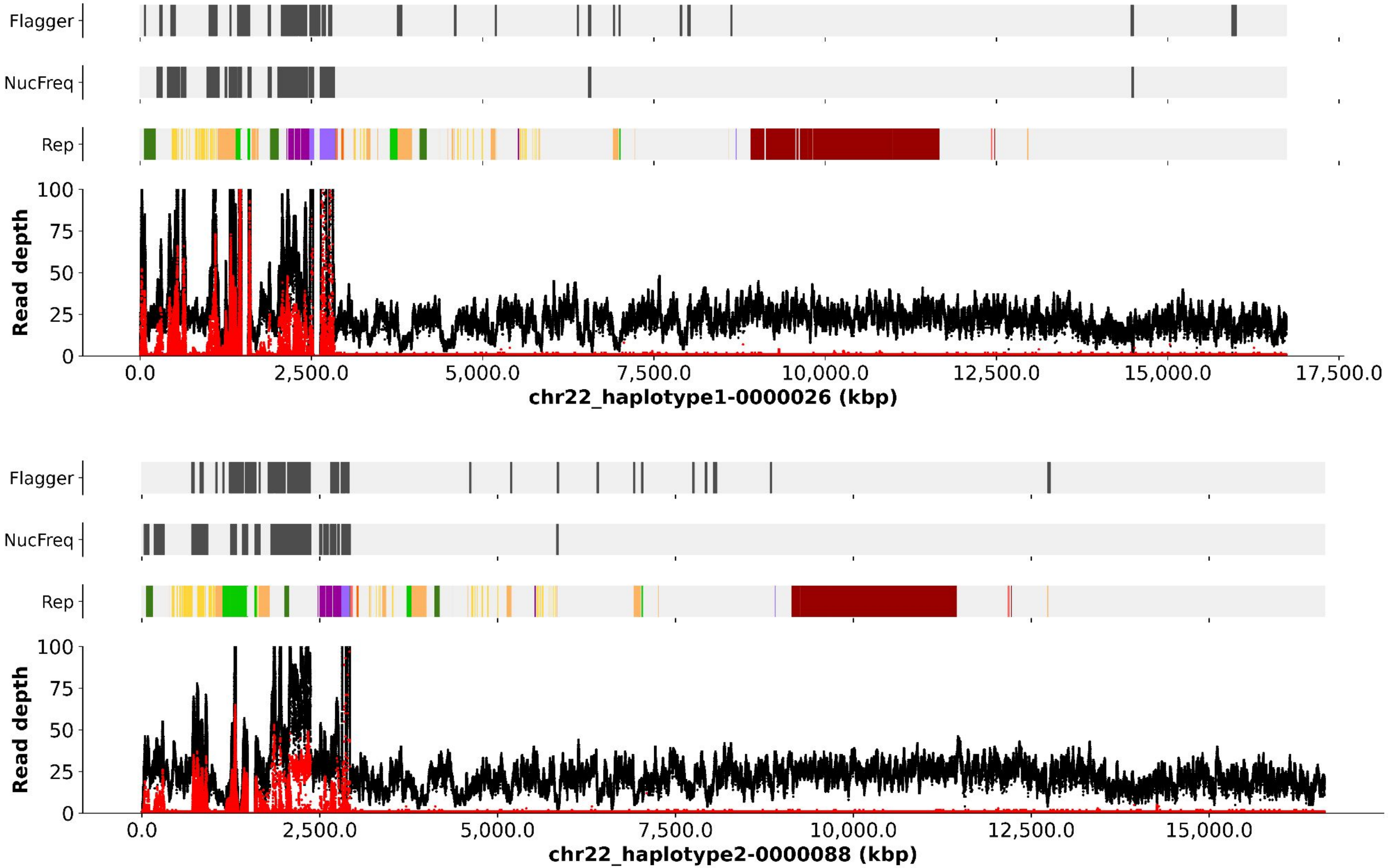

Flagger
NucFreq
Rep
Read depth
100
75
50
25
0
0.0
2,500.0
5,000.0
7,500.0
10,000.0
12,500.0
15,000.0
17,500.0
chr22_haplotype1-0000026 (kbp)
Flagger
NucFreq
Rep
Read depth
chr22_haplotype2-0000088 (kbp)

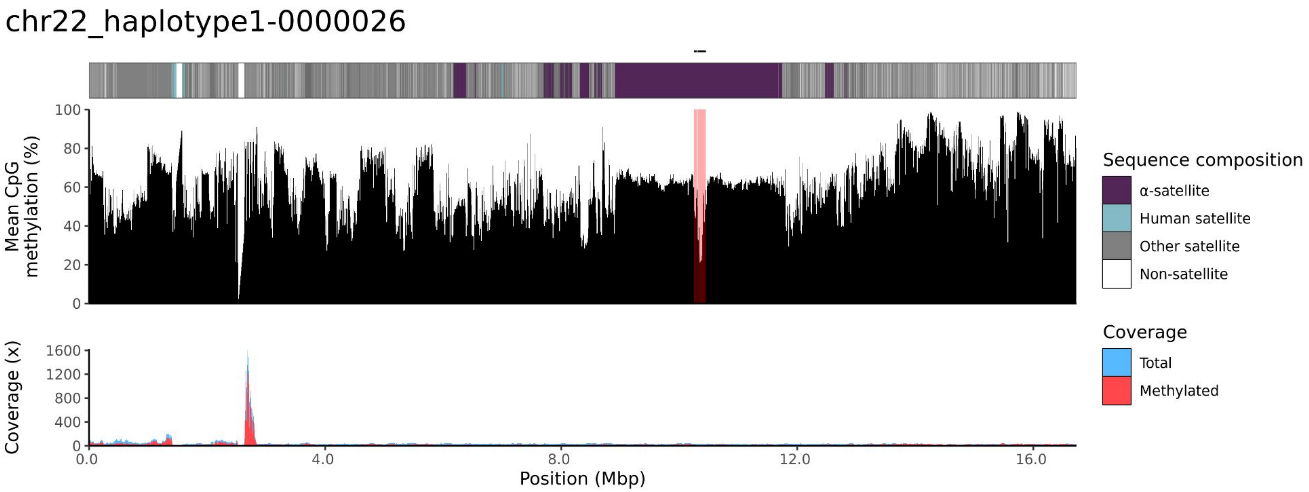
chr22_haplotype1-0000026
Mean CpG methylation (%)
100
80
60
40
20
0
Coverage (x)
1600
1200
800
400
0
0.0
4.0
8.0
12.0
16.0
Position (Mbp)
Sequence composition
α-satellite
Human satellite
Other satellite
Non-satellite
Coverage
Total
Methylated

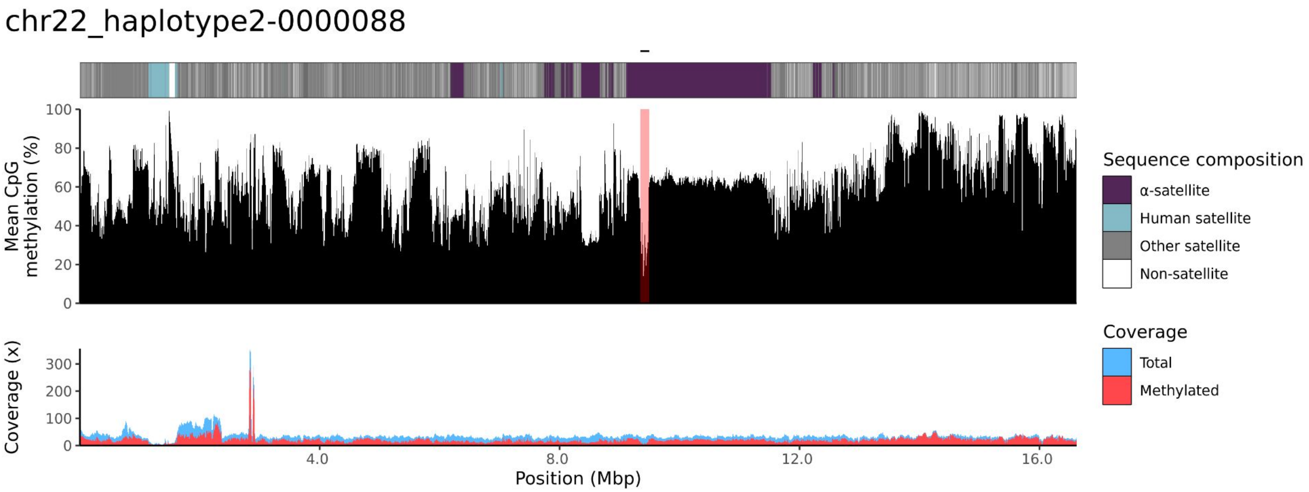
chr22_haplotype2-0000088
Mean CpG methylation (%)
100
80
60
40
20
0
Coverage (x)
300
200
100
0
4.0
8.0
12.0
16.0
Position (Mbp)
Sequence composition
α-satellite
Human satellite
Other satellite
Non-satellite
Coverage
Total
Methylated

200085

### chr13

#### K200085_1_haplotype1-0000015_chr13

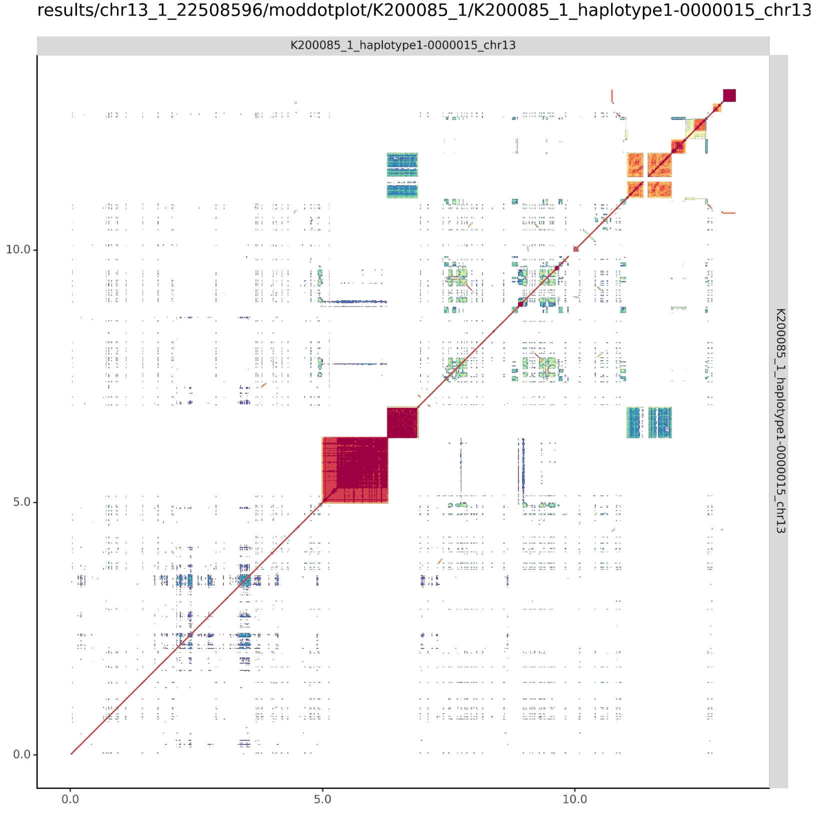


#### K200085_2_haplotype2-0000056_chr13

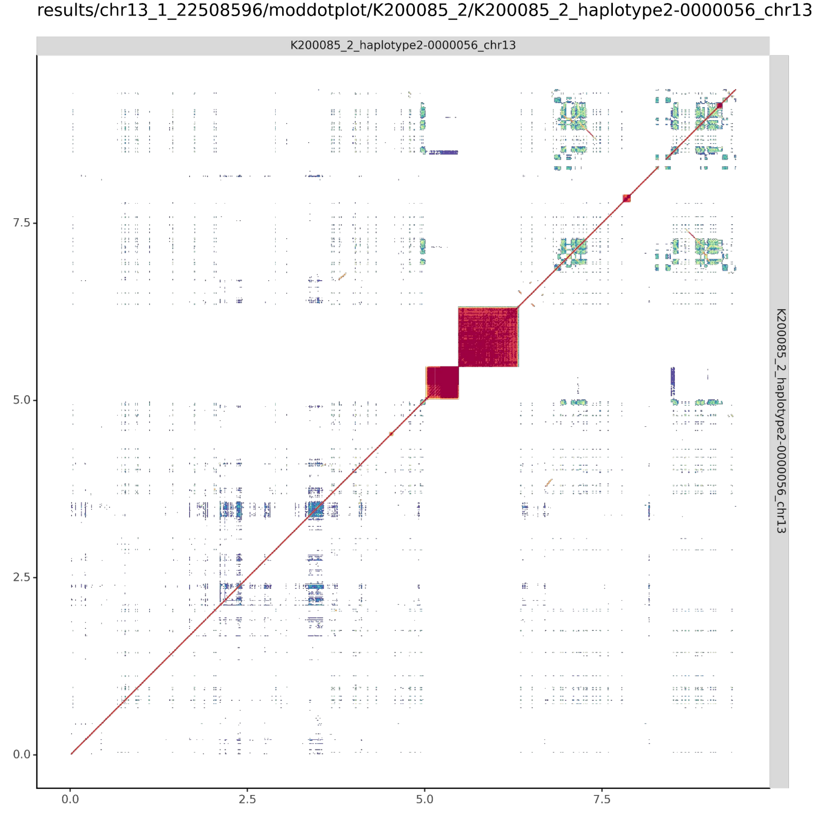

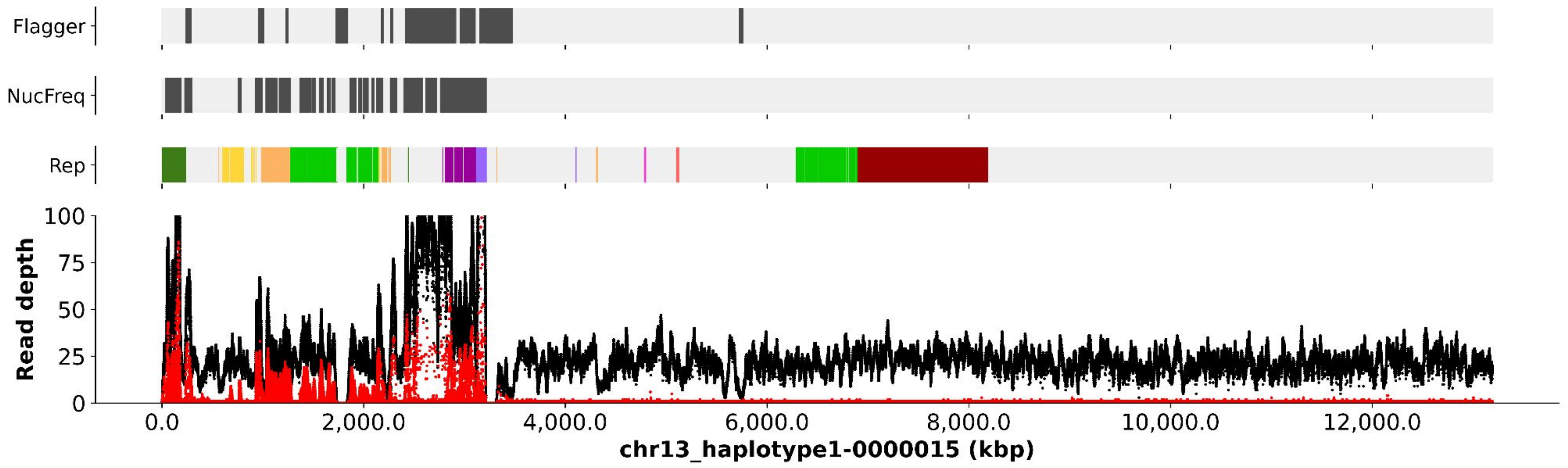

Flagger
NucFreq
Rep
Read depth
100
75
50
25
0
0.0
2,000.0
4,000.0
6,000.0
8,000.0
10,000.0
12,000.0
chr13_haplotype1-0000015 (kbp)


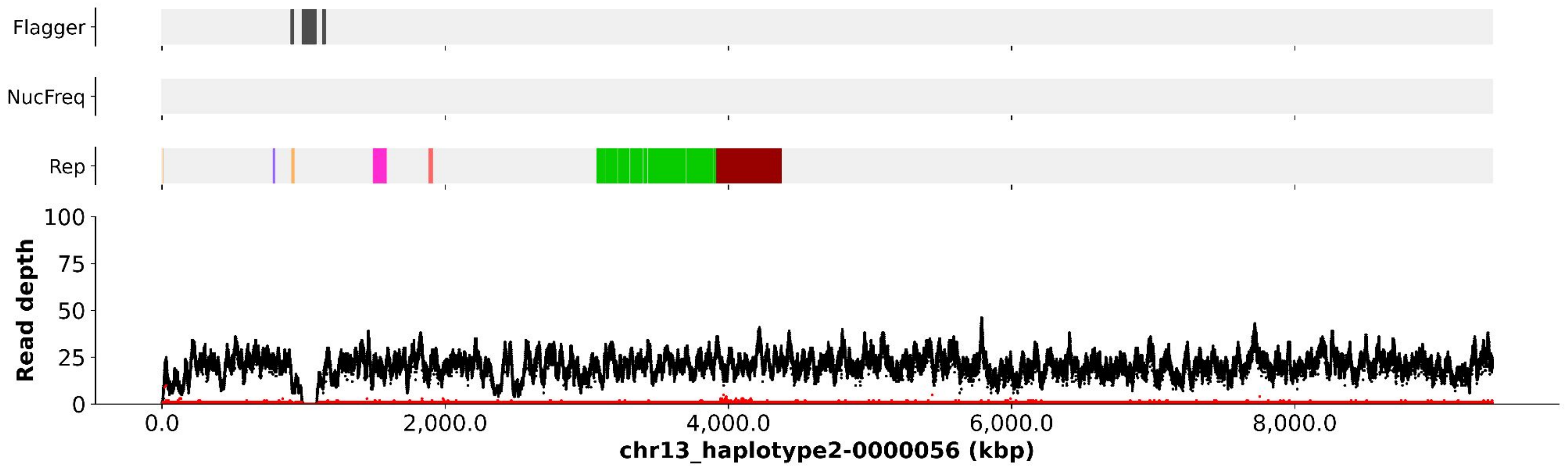

Flagger
NucFreq
Rep
Read depth
100
75
50
25
0
0.0
2,000.0
4,000.0
6,000.0
8,000.0
chr13_haplotype2-0000056 (kbp)

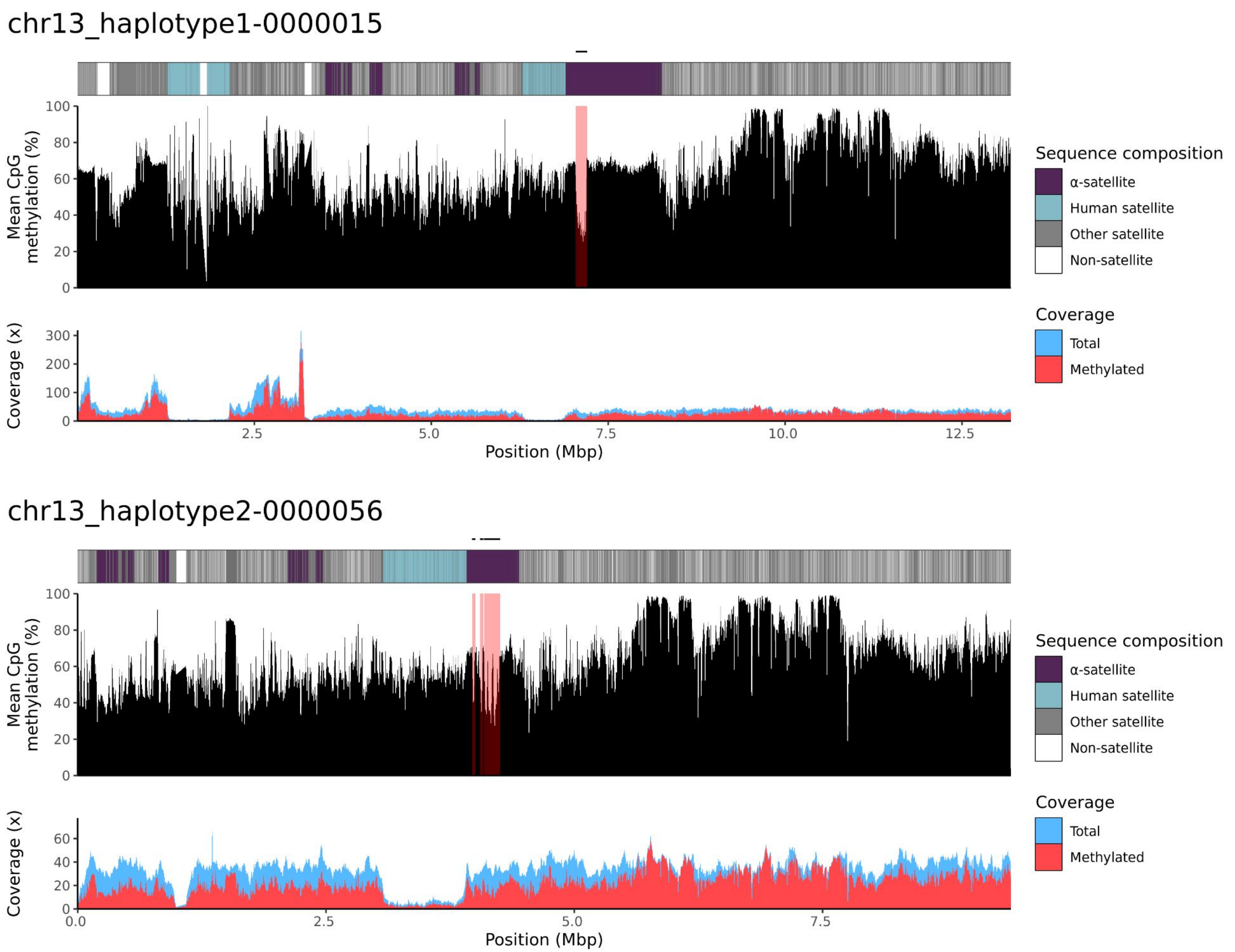
chr13_haplotype1-0000015
Mean CpG methylation (%)
Coverage (x)
Position (Mbp)
Sequence composition
α-satellite
Human satellite
Other satellite
Non-satellite
Coverage
Total
Methylated
chr13_haplotype2-0000056
Mean CpG methylation (%)
Coverage (x)
Position (Mbp)
Sequence composition
α-satellite
Human satellite
Other satellite
Non-satellite
Coverage
Total
Methylated

### chr14

#### K200085_1_haplotype1-0000026_chr14

results/chr14_1_17708411/moddotplot/K200085_1/K200085_1_haplotype1-0000026_chr14

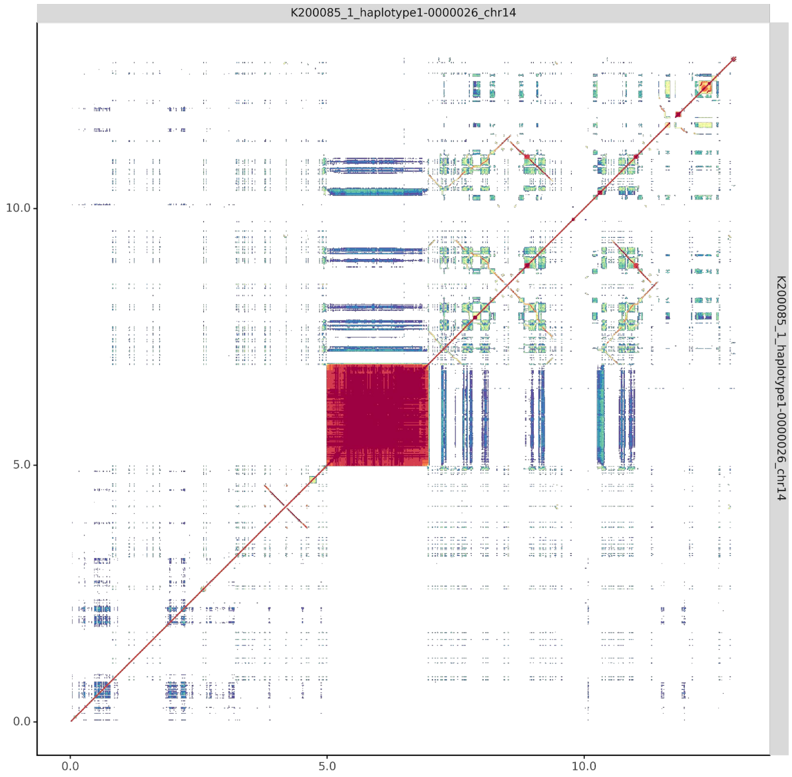


#### K200085_2_haplotype2-0000064_chr14

results/chr14_1_17708411/moddotplot/K200085_2/K200085_2_haplotype2-0000064_chr14

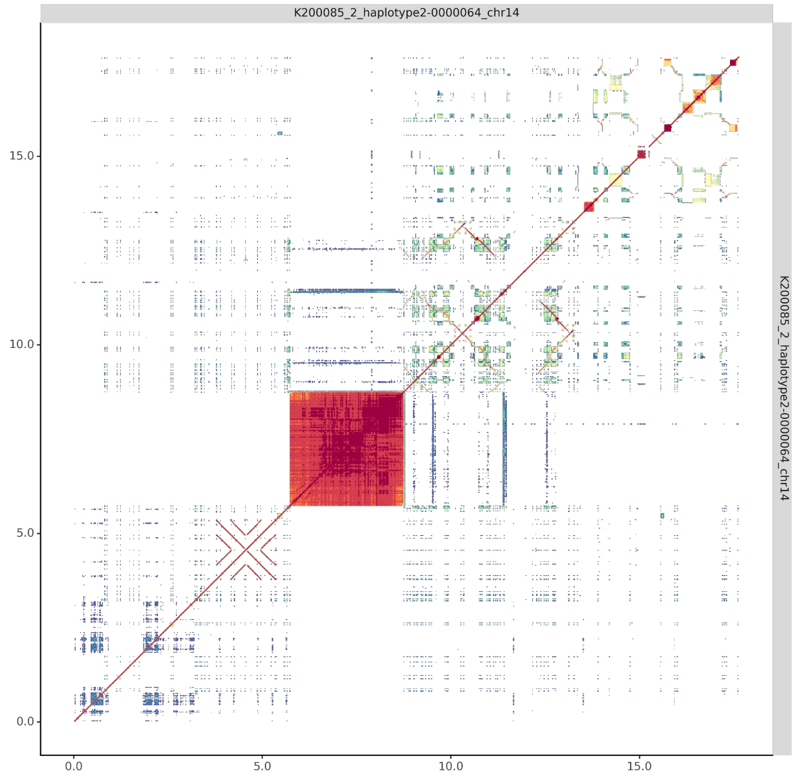

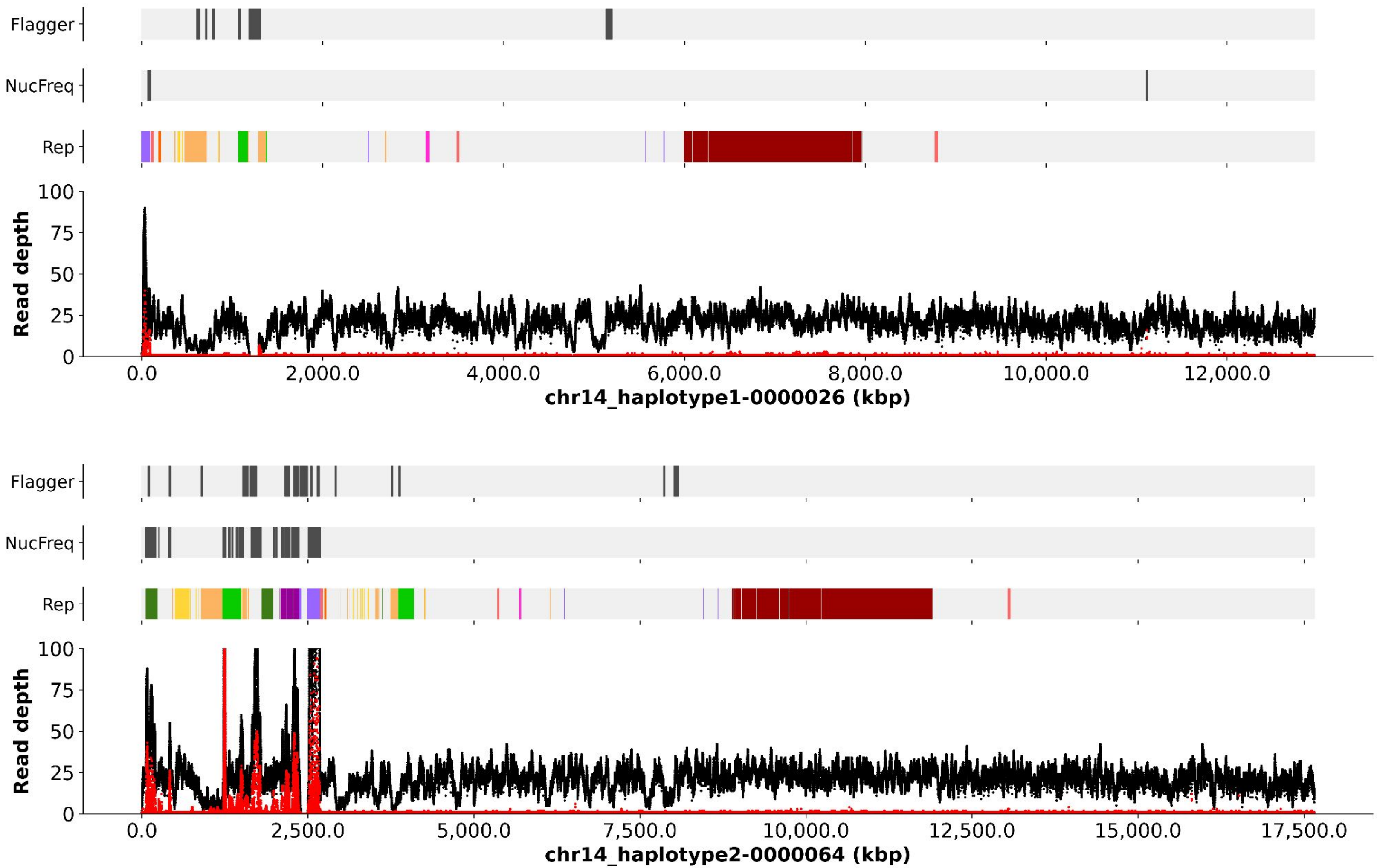

Flagger
NucFreq
Rep
Read depth
100
75
50
25
0
0.0
2,000.0
4,000.0
6,000.0
8,000.0
10,000.0
12,000.0
chr14_haplotype1-0000026 (kbp)
Flagger
NucFreq
Rep
Read depth
100
75
50
25
0
0.0
2,500.0
5,000.0
7,500.0
10,000.0
12,500.0
15,000.0
17,500.0
chr14_haplotype2-0000064 (kbp)

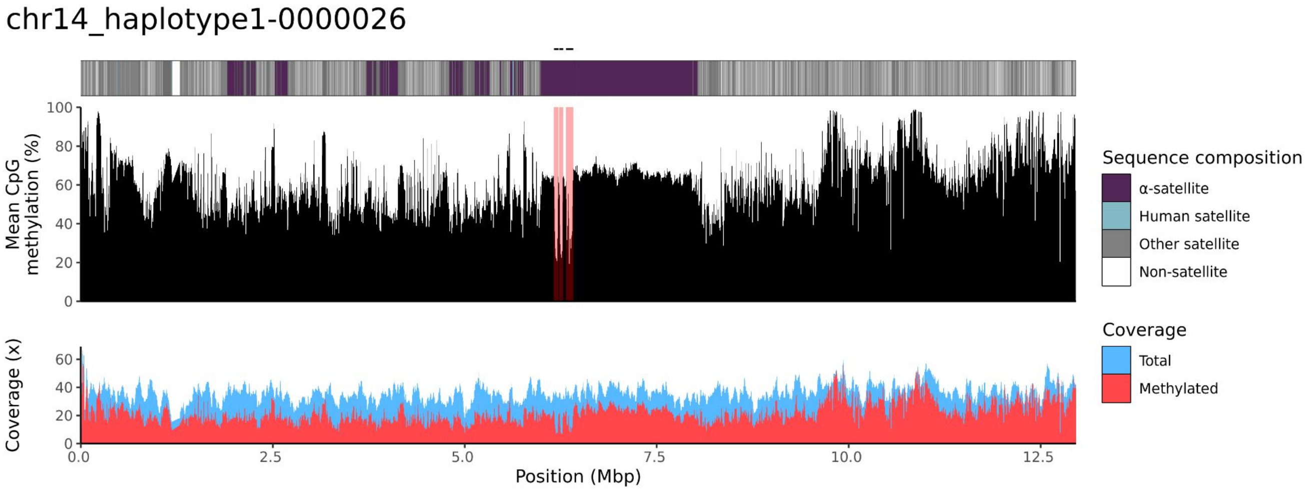

chr14_haplotype1-0000026
Mean CpG methylation (%)
Coverage (x)
Position (Mbp)
Sequence composition
α-satellite
Human satellite
Other satellite
Non-satellite
Coverage
Total
Methylated


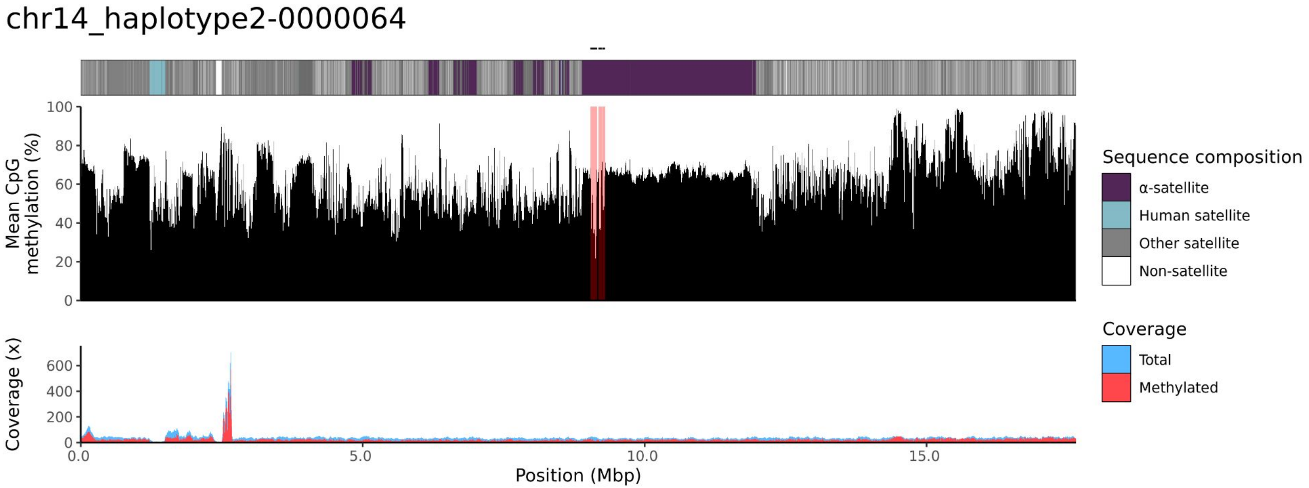

chr14_haplotype2-0000064
Mean CpG methylation (%)
Coverage (x)
Position (Mbp)
Sequence composition
α-satellite
Human satellite
Other satellite
Non-satellite
Coverage
Total
Methylated

### chr15

#### K200085_1_haplotype1-0000025_chr15

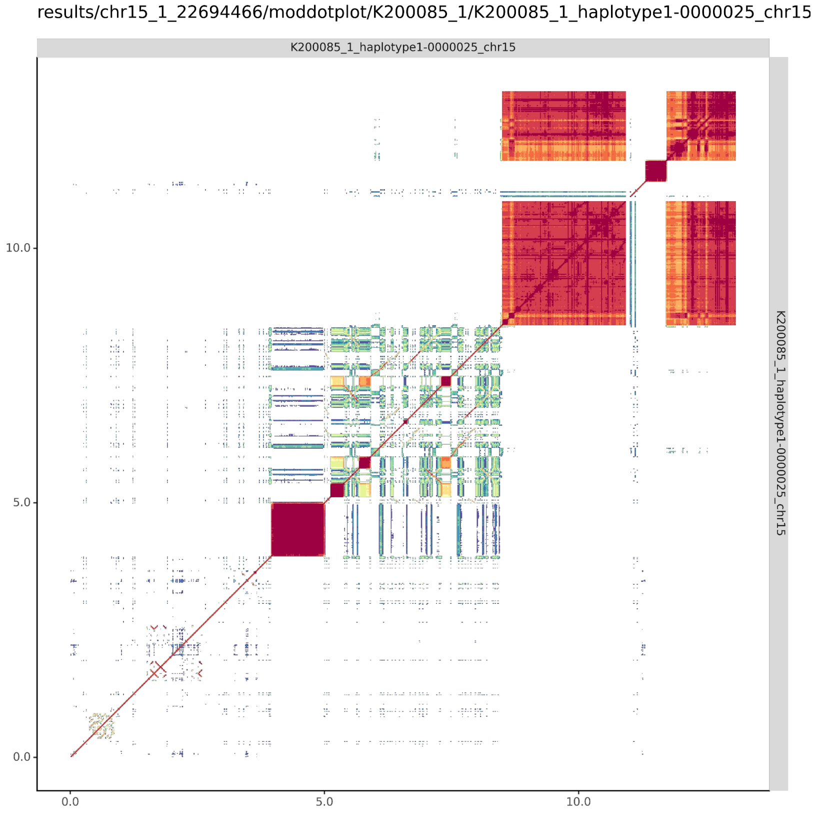


#### K200085_2_haplotype2-0000063_chr15

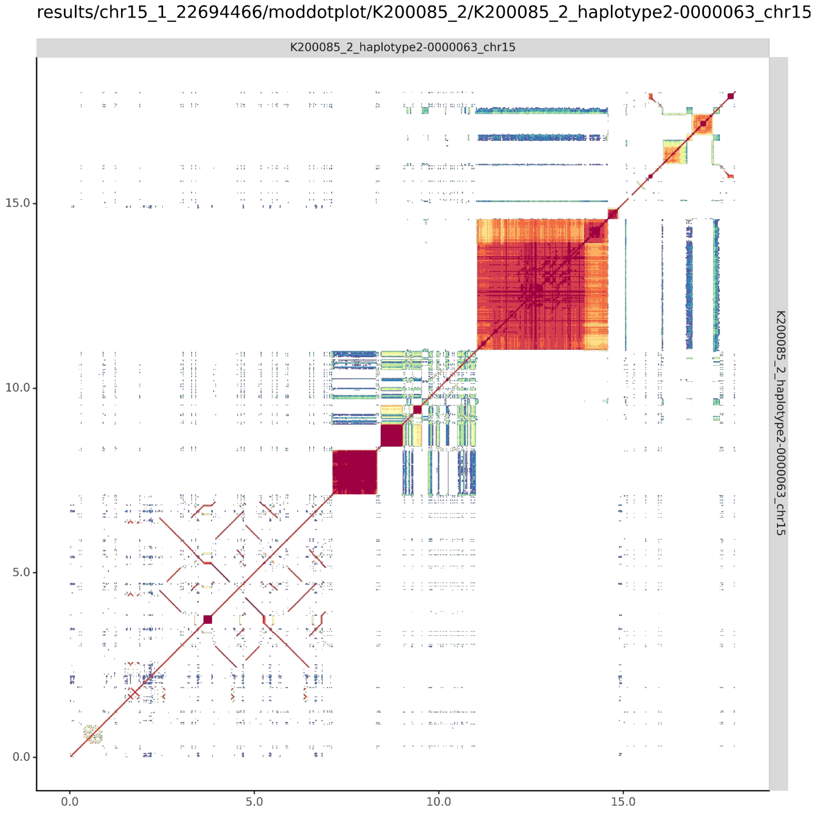

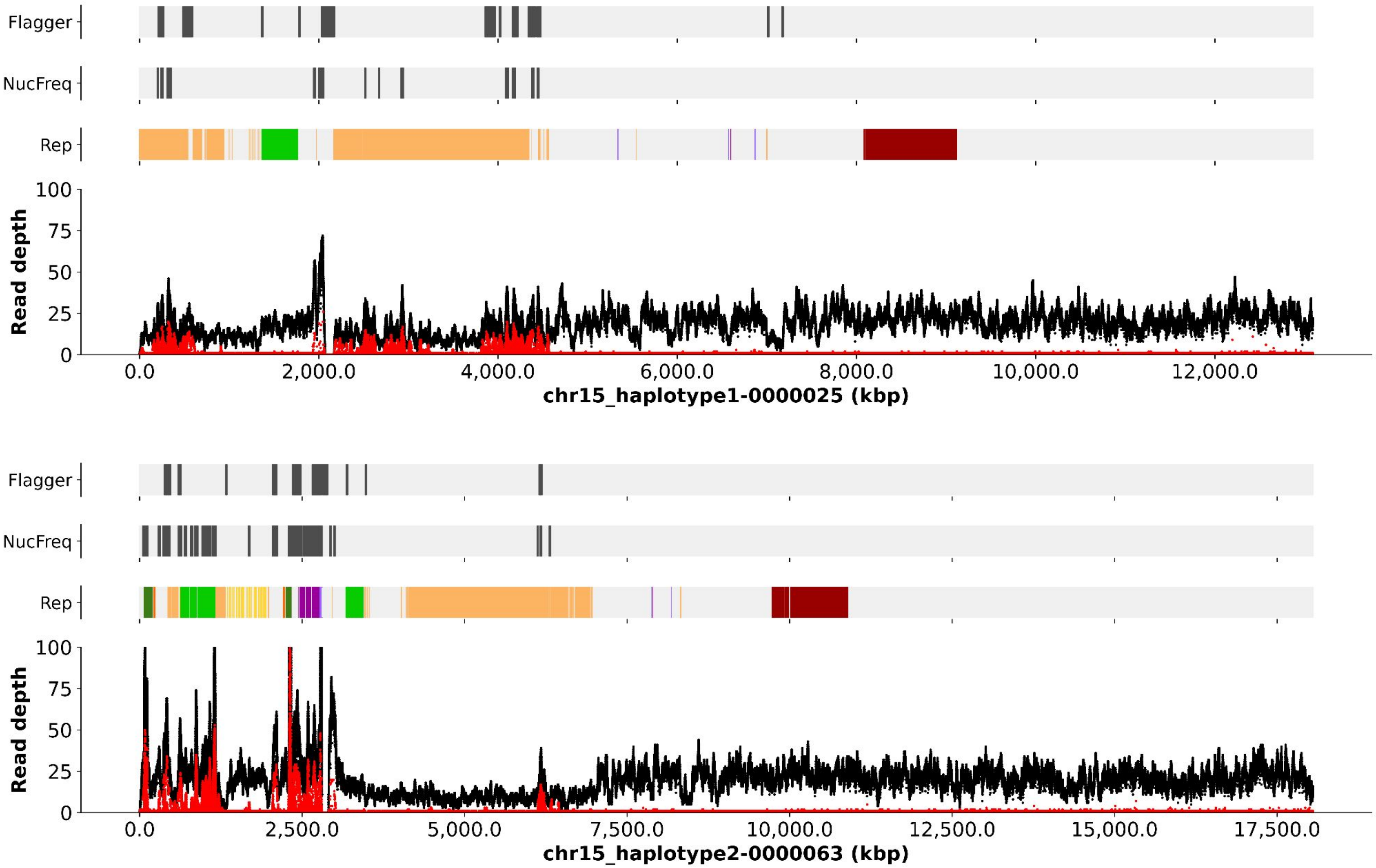

Flagger
NucFreq
Rep
Read depth
100
75
50
25
0
0.0
2,000.0
4,000.0
6,000.0
8,000.0
10,000.0
12,000.0
chr15_haplotype1-0000025 (kbp)
Flagger
NucFreq
Rep
Read depth
100
75
50
25
0
0.0
2,500.0
5,000.0
7,500.0
10,000.0
12,500.0
15,000.0
17,500.0
chr15_haplotype2-0000063 (kbp)

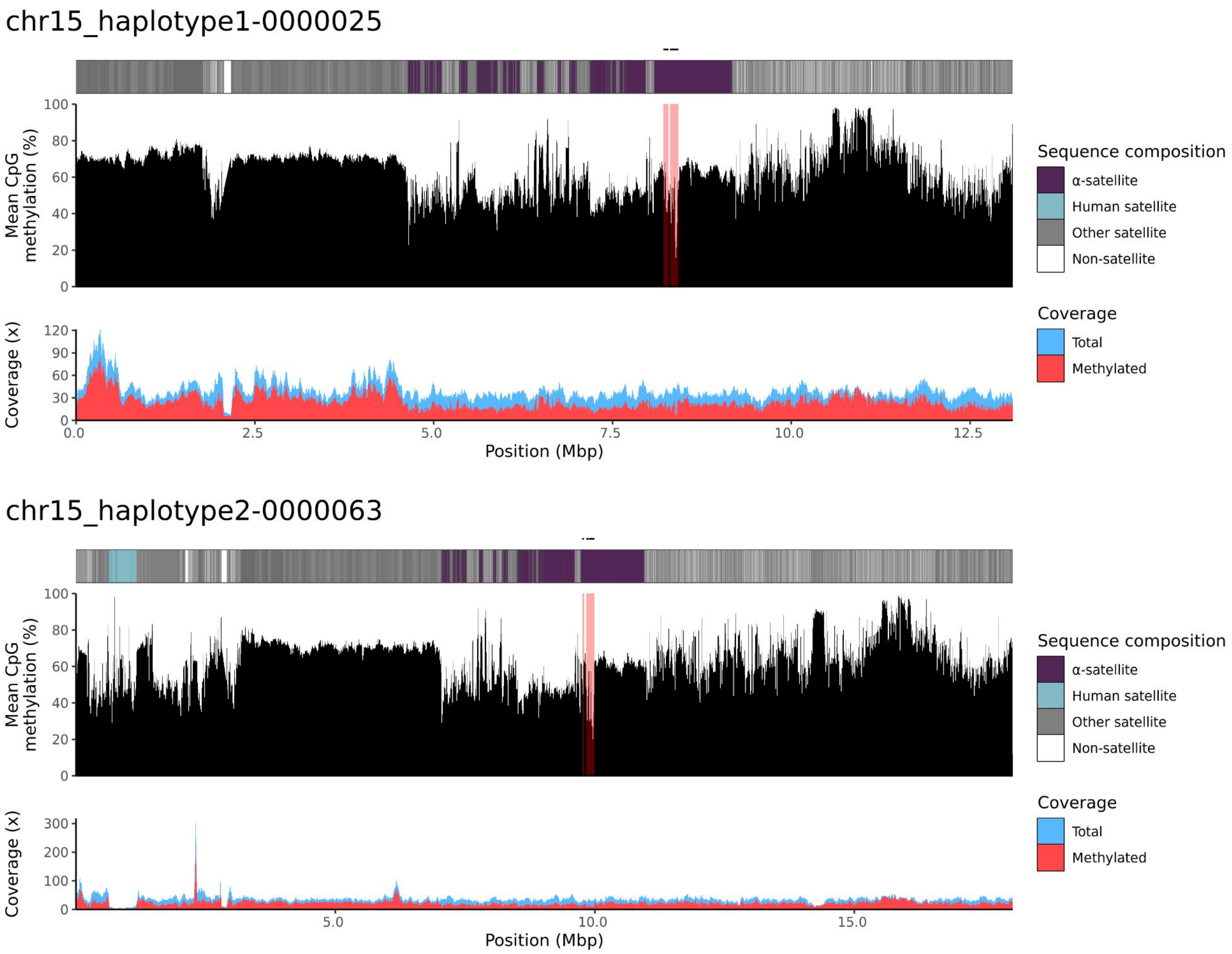
chr15_haplotype1-0000025
Mean CpG methylation (%)
Coverage (x)
Position (Mbp)
Sequence composition
α-satellite
Human satellite
Other satellite
Non-satellite
Coverage
Total
Methylated
chr15_haplotype2-0000063
Mean CpG methylation (%)
Coverage (x)
Position (Mbp)
Sequence composition
α-satellite
Human satellite
Other satellite
Non-satellite
Coverage
Total
Methylated

### chr21

#### K200085_1_haplotype1-0000012_chr21

results/chr21_1_16306378/moddotplot/K200085_1/K200085_1_haplotype1-0000012_chr21

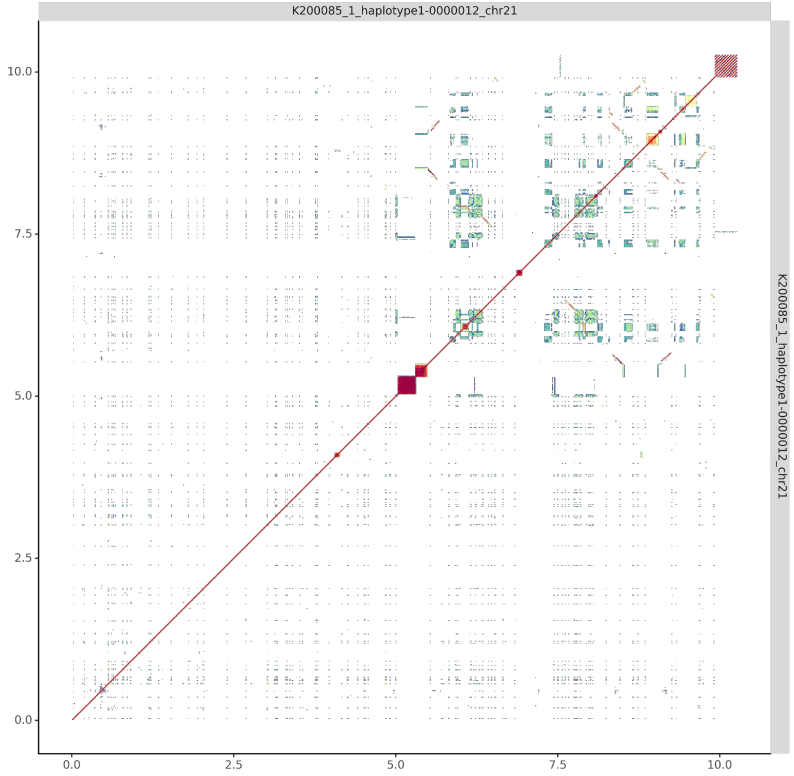


#### K200085_2_haplotype2-0000053_chr21

results/chr21_1_16306378/moddotplot/K200085_2/K200085_2_haplotype2-0000053_chr21

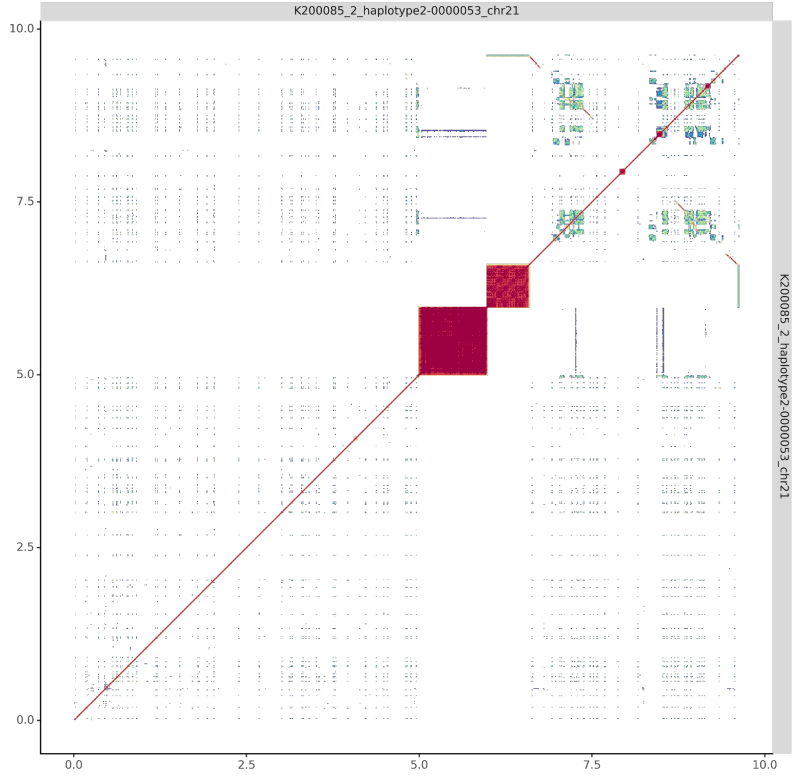

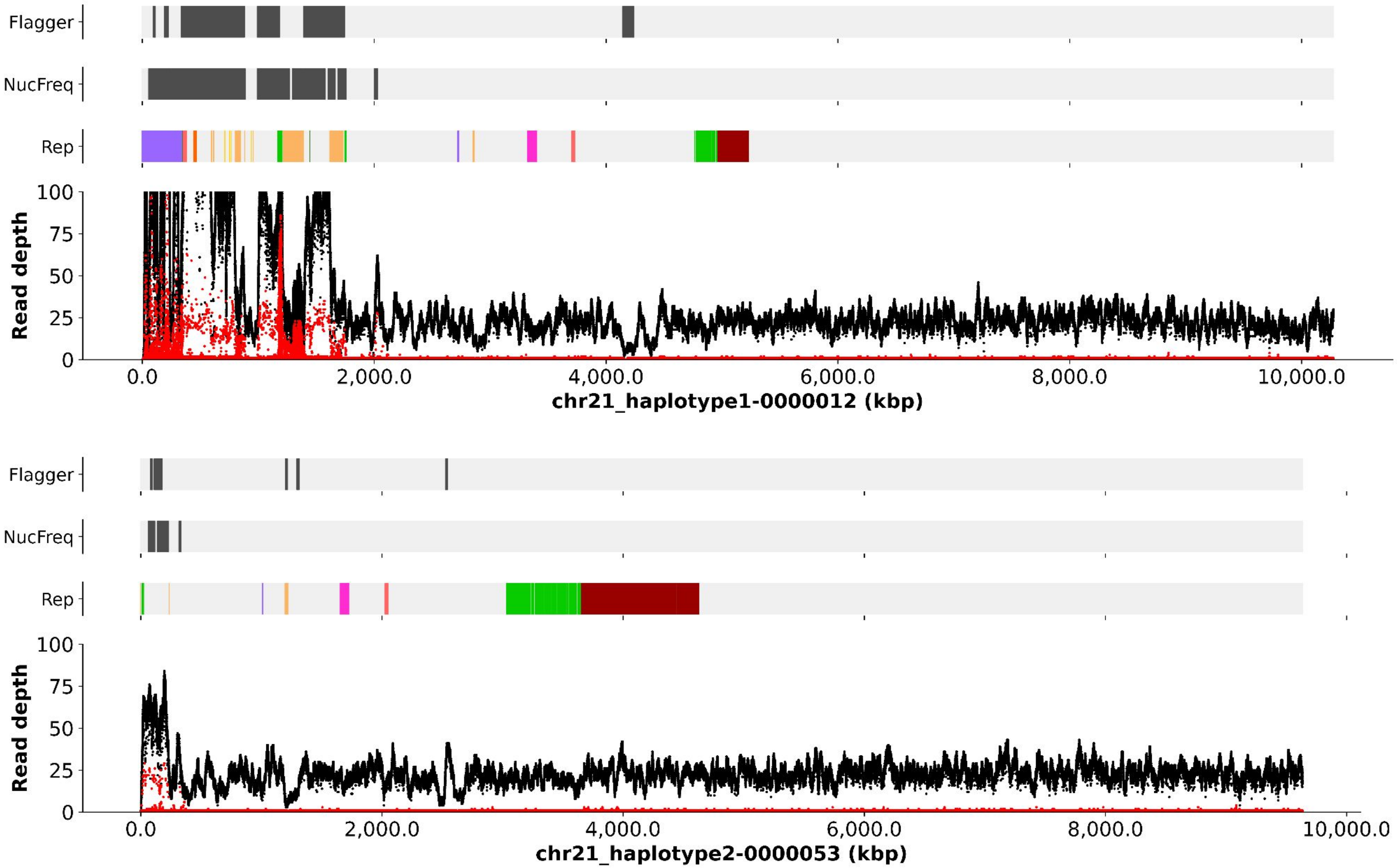
Flagger
NucFreq
Rep
Read depth
100
75
50
25
0
0.0
2,000.0
4,000.0
6,000.0
8,000.0
10,000.0
chr21_haplotype1-0000012 (kbp)
Flagger
NucFreq
Rep
Read depth
100
75
50
25
0
0.0
2,000.0
4,000.0
6,000.0
8,000.0
10,000.0
chr21_haplotype2-0000053 (kbp)

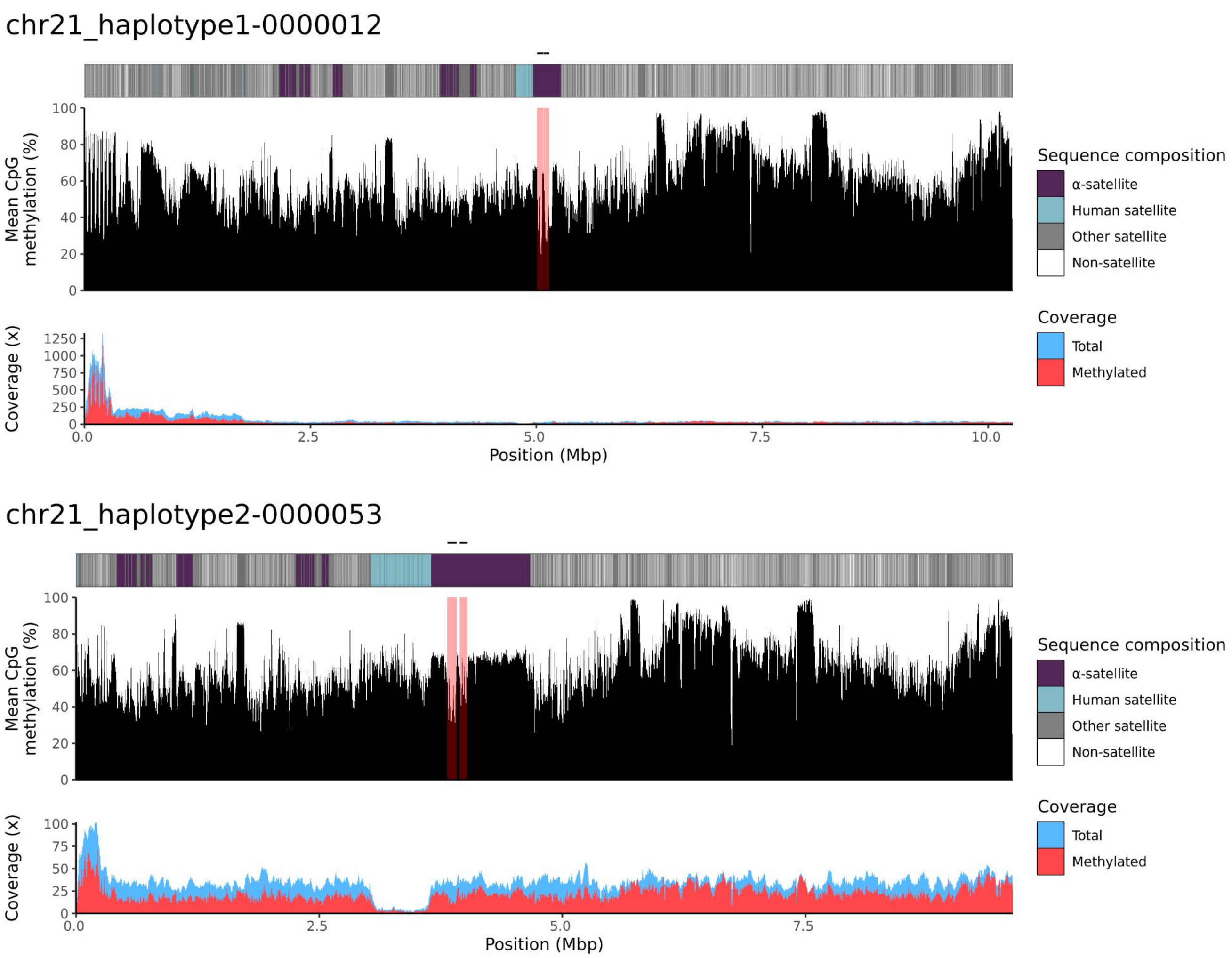

chr21_haplotype1-0000012
Mean CpG methylation (%)
100
80
60
40
20
0
Coverage (x)
1250
1000
750
500
250
0
0.0
2.5
5.0
7.5
10.0
Position (Mbp)
Sequence composition
α-satellite
Human satellite
Other satellite
Non-satellite
Coverage
Total
Methylated
chr21_haplotype2-0000053
Mean CpG methylation (%)
100
80
60
40
20
0
Coverage (x)
100
75
50
25
0
0.0
2.5
5.0
7.5
Position (Mbp)
Sequence composition
α-satellite
Human satellite
Other satellite
Non-satellite
Coverage
Total
Methylated

### chr22

#### K200085_1_haplotype1-0000001_chr22

results/chr22_1_20711065/moddotplot/K200085_1/K200085_1_haplotype1-0000001_chr22

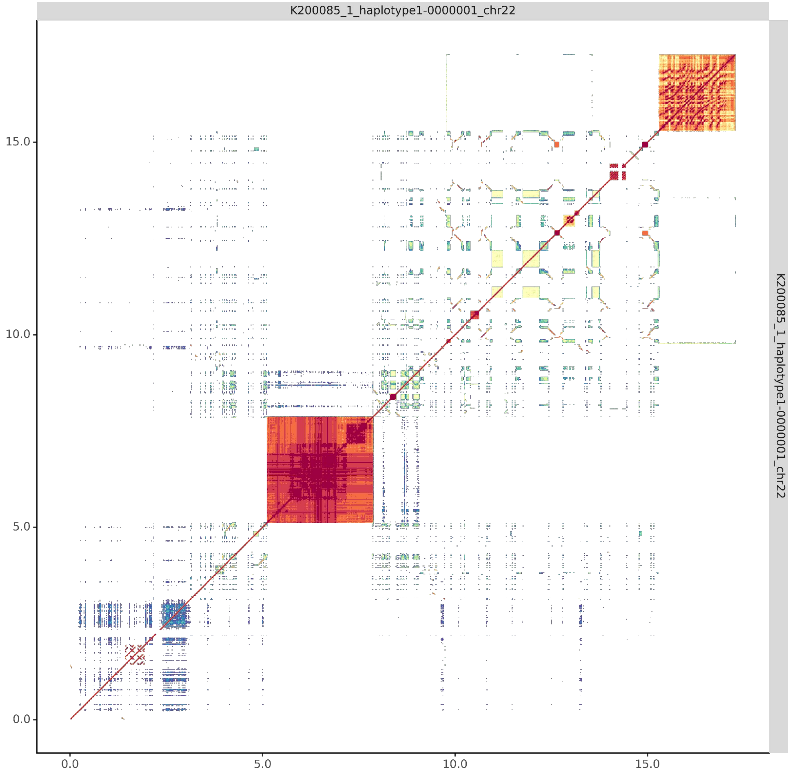


#### K200085_2_haplotype2-0000042_chr22

results/chr22_1_20711065/moddotplot/K200085_2/K200085_2_haplotype2-0000042_chr22

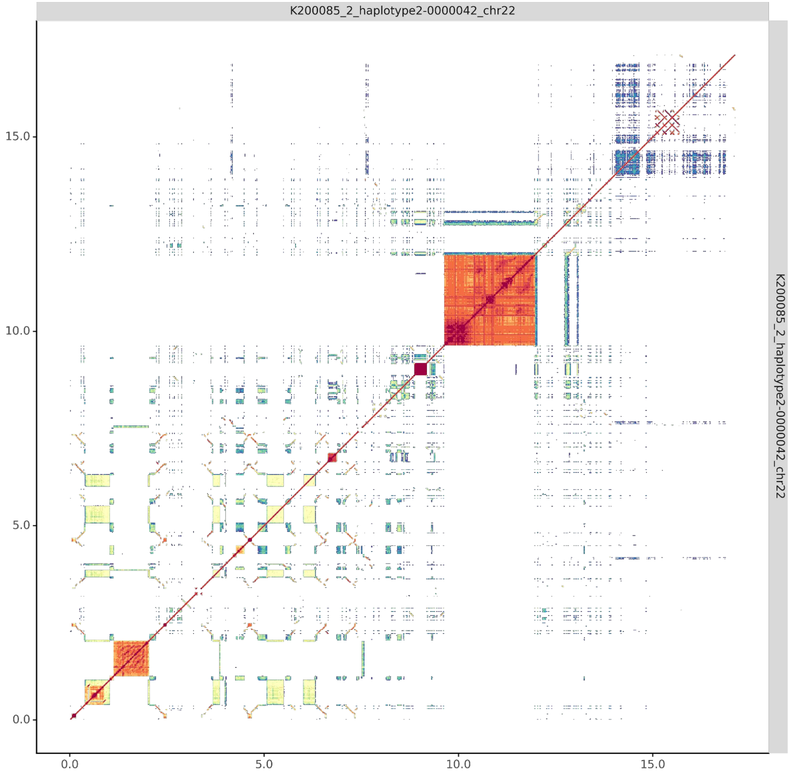

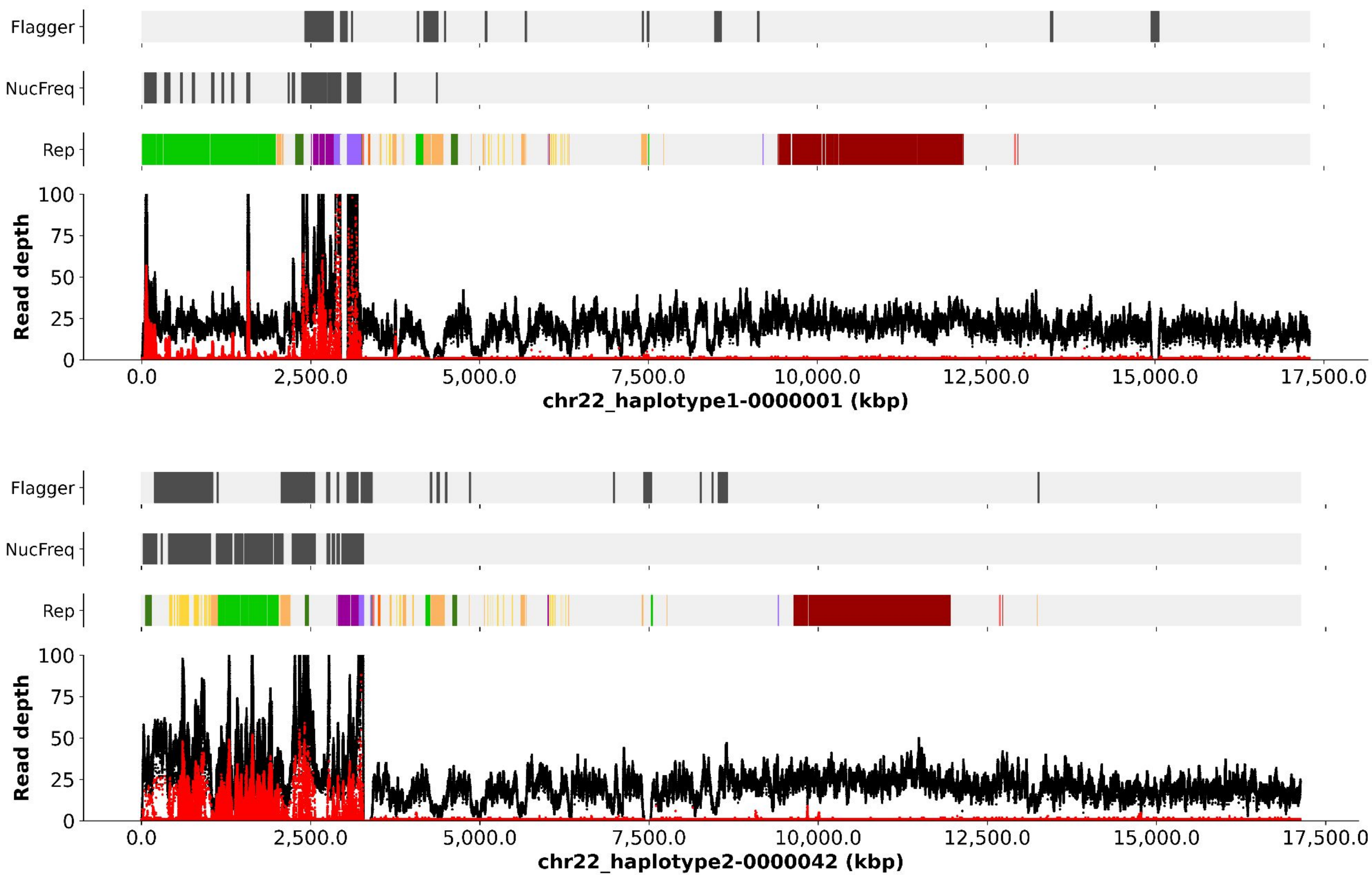

Flagger
NucFreq
Rep
Read depth
100
75
50
25
0
0.0
2,500.0
5,000.0
7,500.0
10,000.0
12,500.0
15,000.0
17,500.0
chr22_haplotype1-0000001 (kbp)
Flagger
NucFreq
Rep
Read depth
100
75
50
25
0
0.0
2,500.0
5,000.0
7,500.0
10,000.0
12,500.0
15,000.0
17,500.0
chr22_haplotype2-0000042 (kbp)

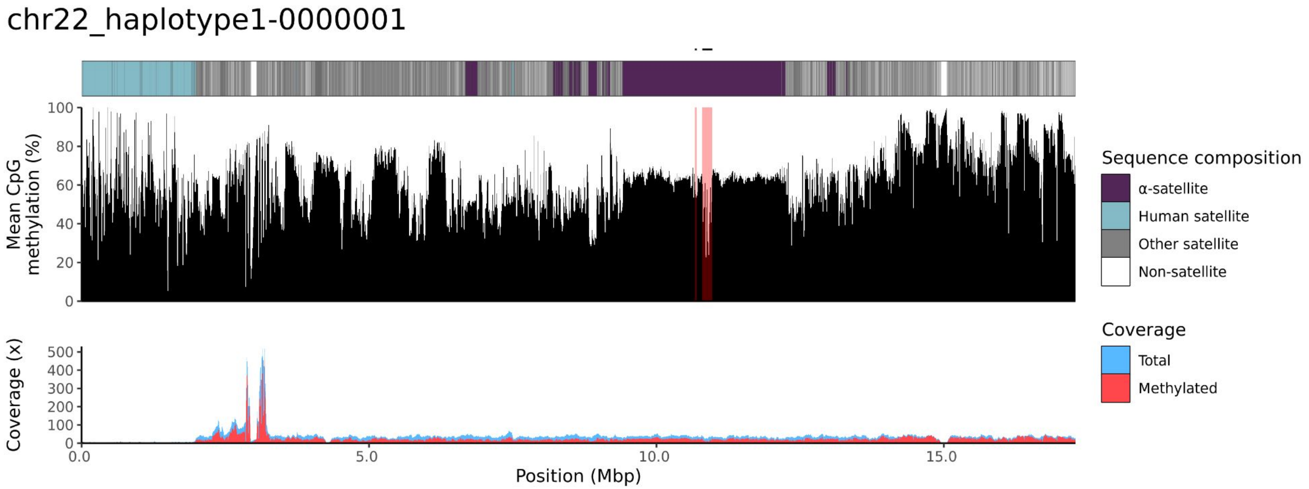
chr22_haplotype1-0000001
Mean CpG methylation (%)
0
20
40
60
80
100
Coverage (x)
0
100
200
300
400
500
0.0
5.0
10.0
15.0
Position (Mbp)
Sequence composition
α-satellite
Human satellite
Other satellite
Non-satellite
Coverage
Total
Methylated

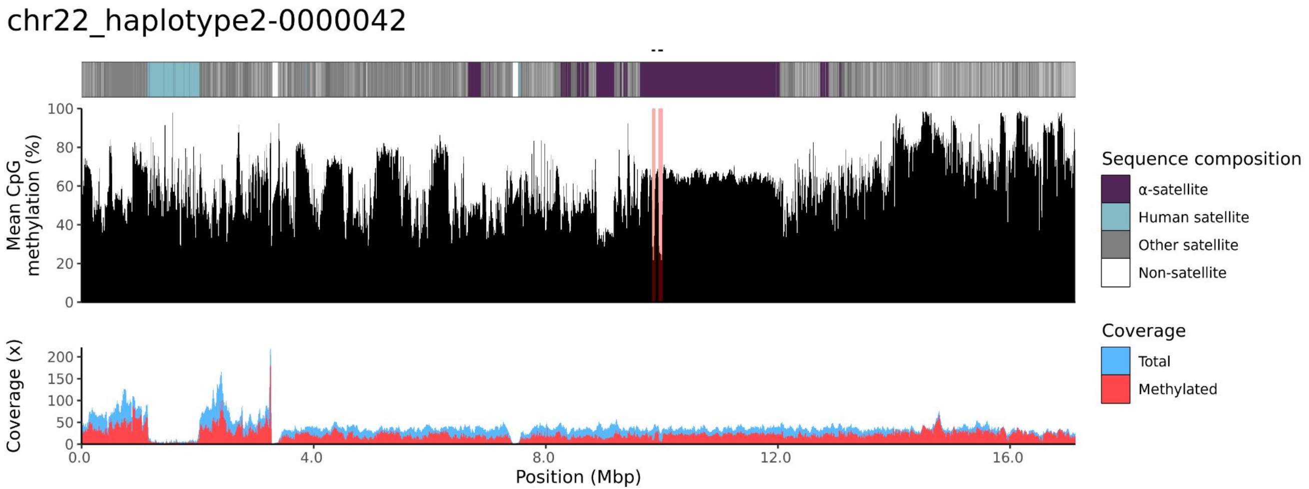
chr22_haplotype2-0000042
Mean CpG methylation (%)
0
20
40
60
80
100
Coverage (x)
0
50
100
150
200
0.0
4.0
8.0
12.0
16.0
Position (Mbp)
Sequence composition
α-satellite
Human satellite
Other satellite
Non-satellite
Coverage
Total
Methylated

200087

### chr13

#### K200087_2_haplotype2-0000205_chr13

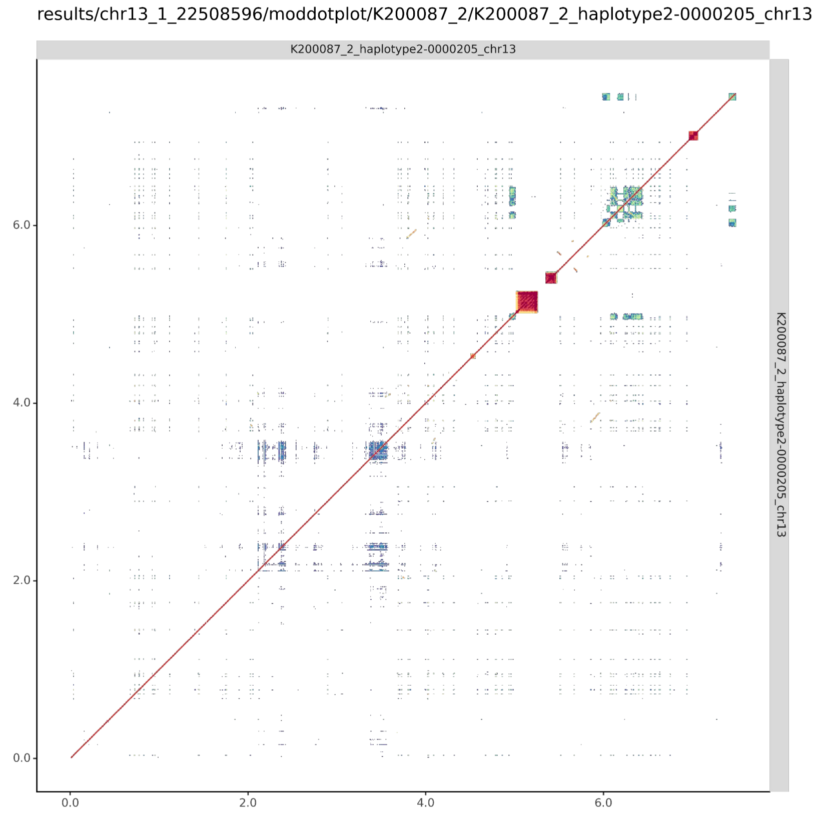

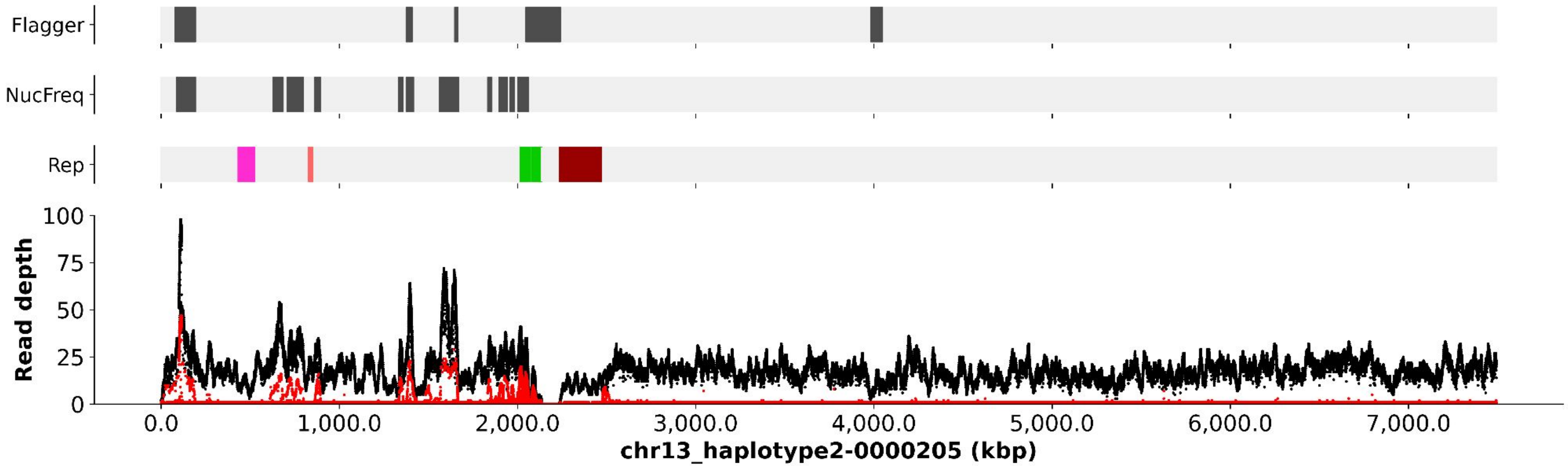
Flagger
NucFreq
Rep
100
75
50
25
0
Read depth
0.0
1,000.0
2,000.0
3,000.0
4,000.0
5,000.0
6,000.0
7,000.0
chr13_haplotype2-0000205 (kbp)

### chr13_haplotype2-0000205

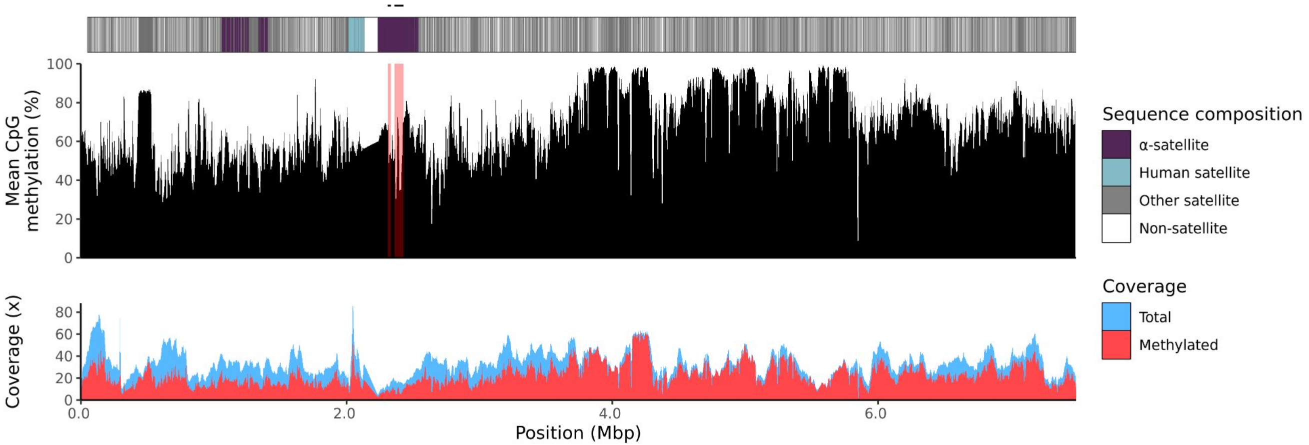

### chr14

#### K200087_1_haplotype1-0000002_chr14

results/chr14_1_17708411/moddotplot/K200087_1/K200087_1_haplotype1-0000002_chr14

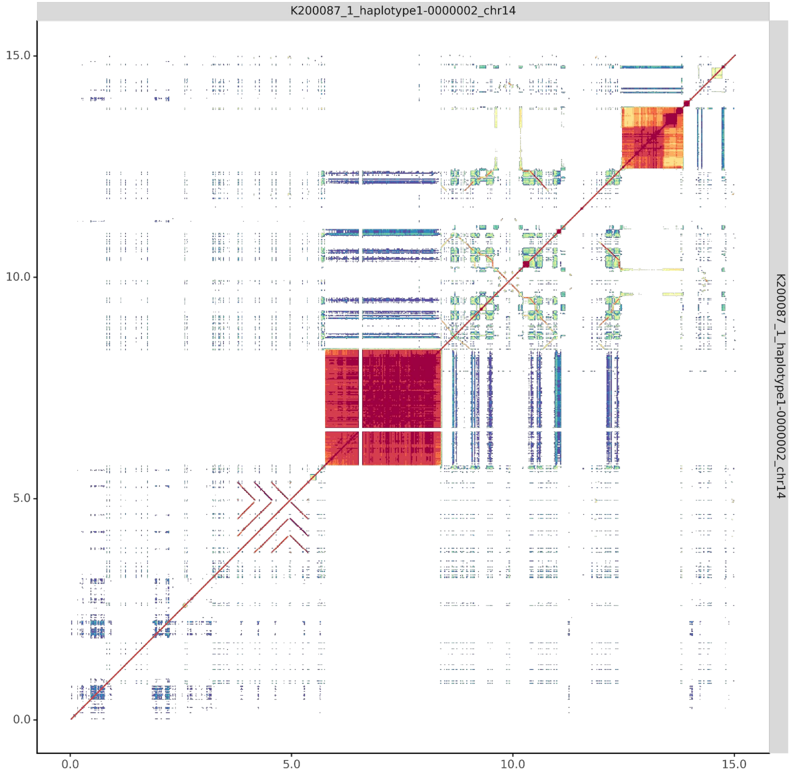


#### K200087_2_haplotype2-0000194_chr14

results/chr14_1_17708411/moddotplot/K200087_2/K200087_2_haplotype2-0000194_chr14

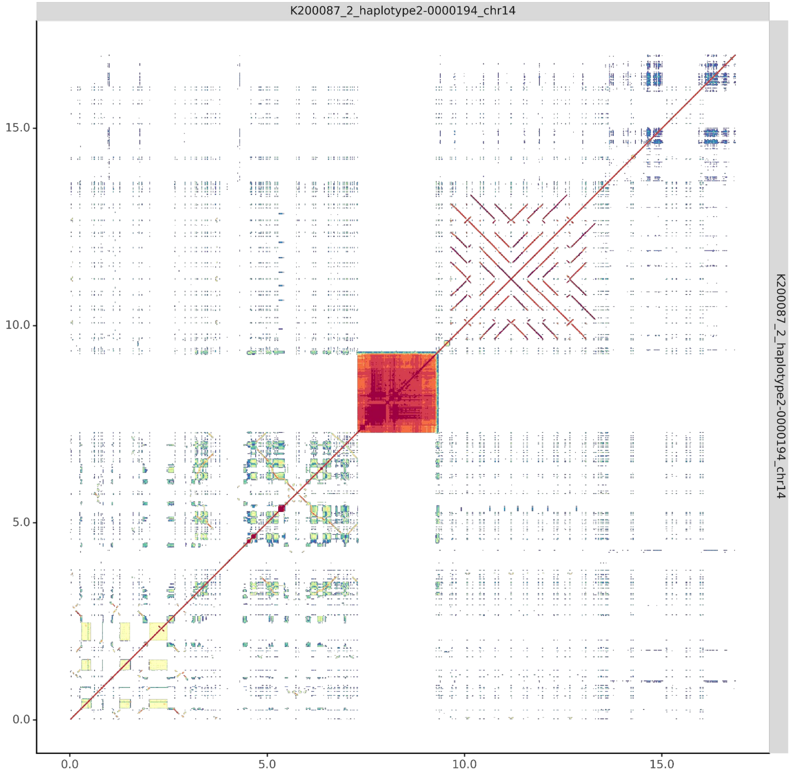

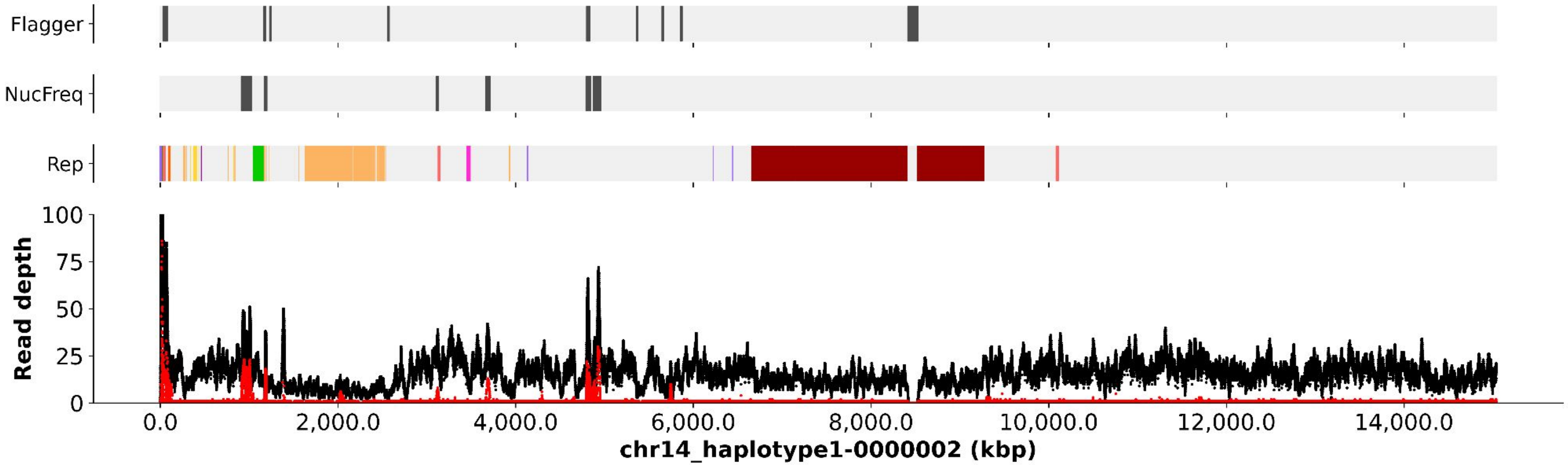

Flagger
NucFreq
Rep
Read depth
100
75
50
25
0
0.0
2,000.0
4,000.0
6,000.0
8,000.0
10,000.0
12,000.0
14,000.0
chr14_haplotype1-0000002 (kbp)


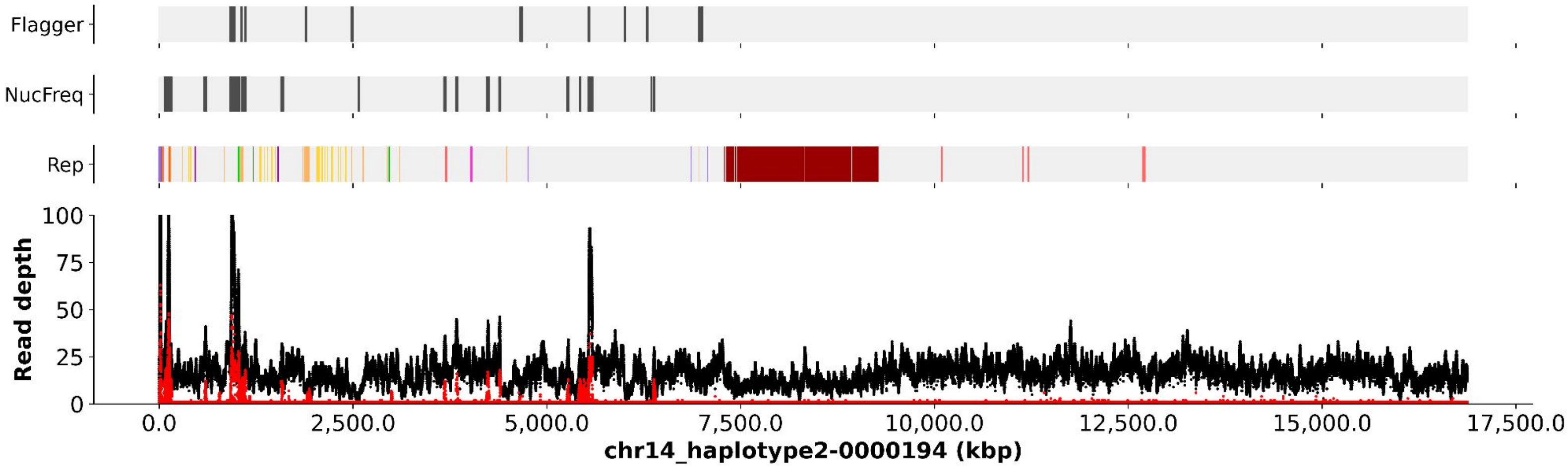

Flagger
NucFreq
Rep
Read depth
100
75
50
25
0
0.0
2,500.0
5,000.0
7,500.0
10,000.0
12,500.0
15,000.0
17,500.0
chr14_haplotype2-0000194 (kbp)

chr14_haplotype1-0000002

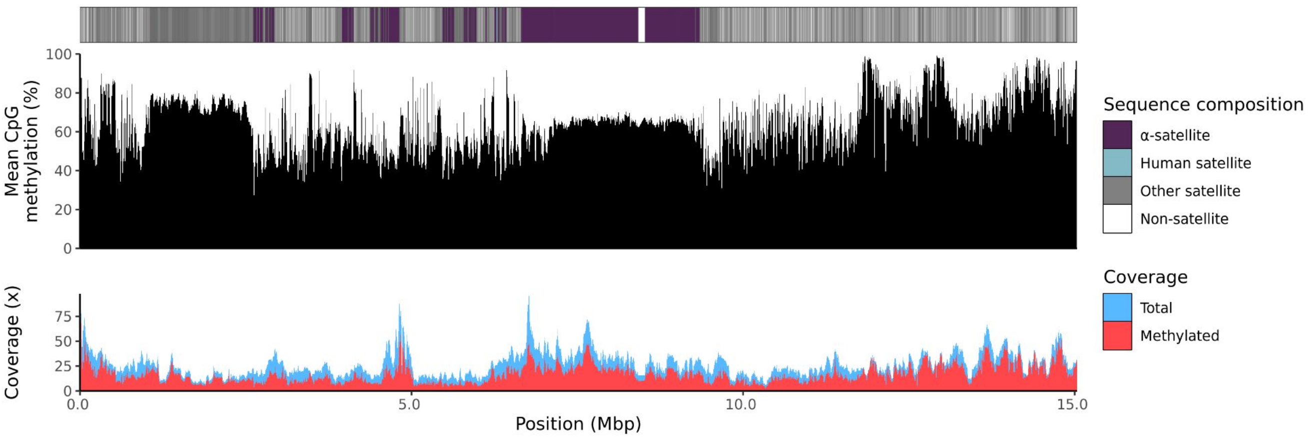


chr14_haplotype2-0000194

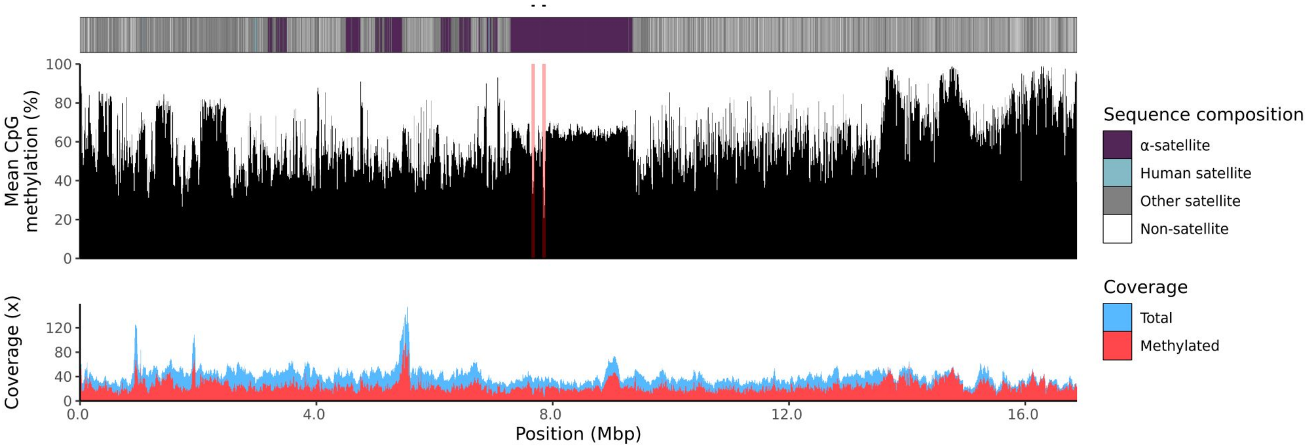

### chr15

#### K200087_1_haplotype1-0000014_chr15

results/chr15_1_22694466/moddotplot/K200087_1/K200087_1_haplotype1-0000014_chr15

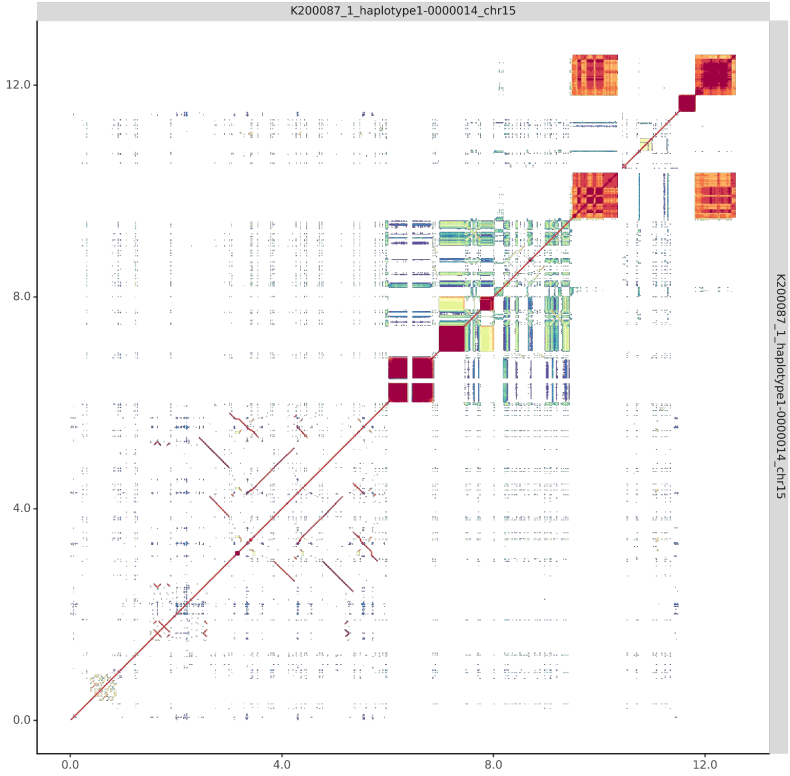


#### K200087_2_haplotype2-0000206_chr15

results/chr15_1_22694466/moddotplot/K200087_2/K200087_2_haplotype2-0000206_chr15

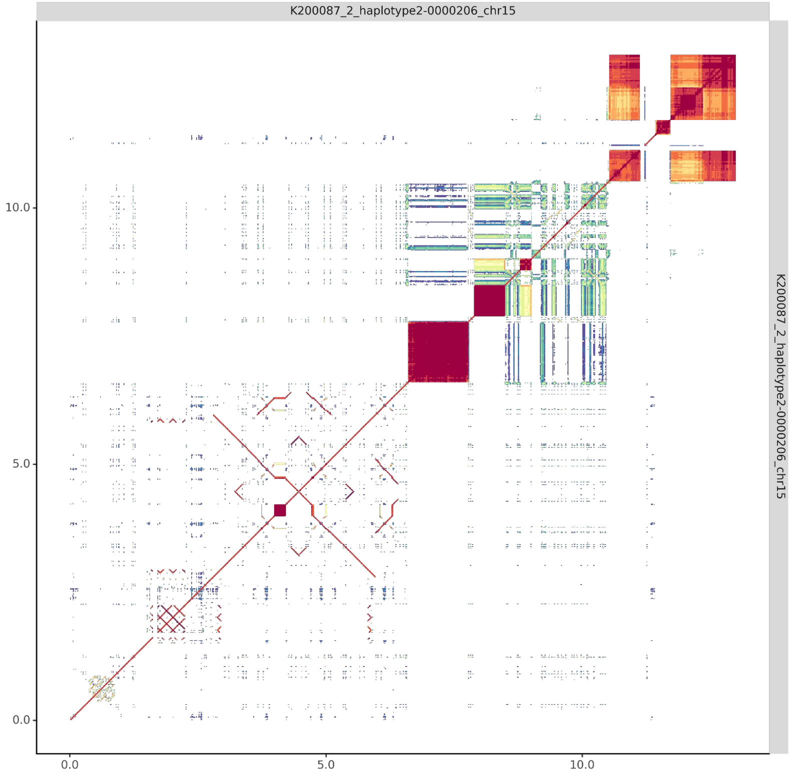

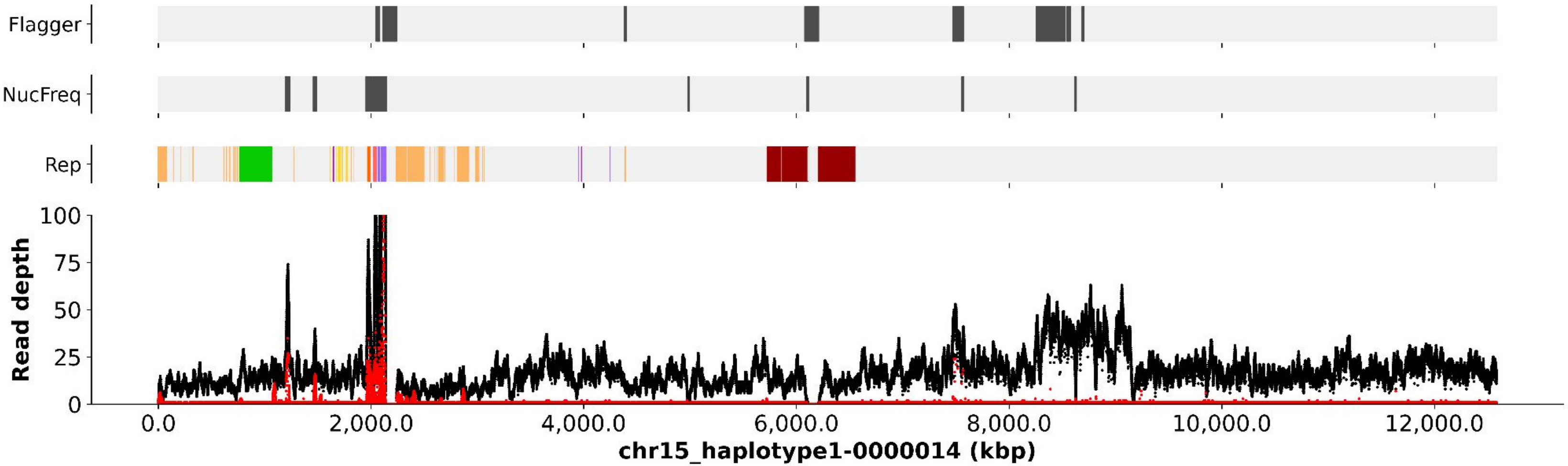
Flagger
NucFreq
Rep
Read depth
0.0 2,000.0 4,000.0 6,000.0 8,000.0 10,000.0 12,000.0
chr15_haplotype1-0000014 (kbp)

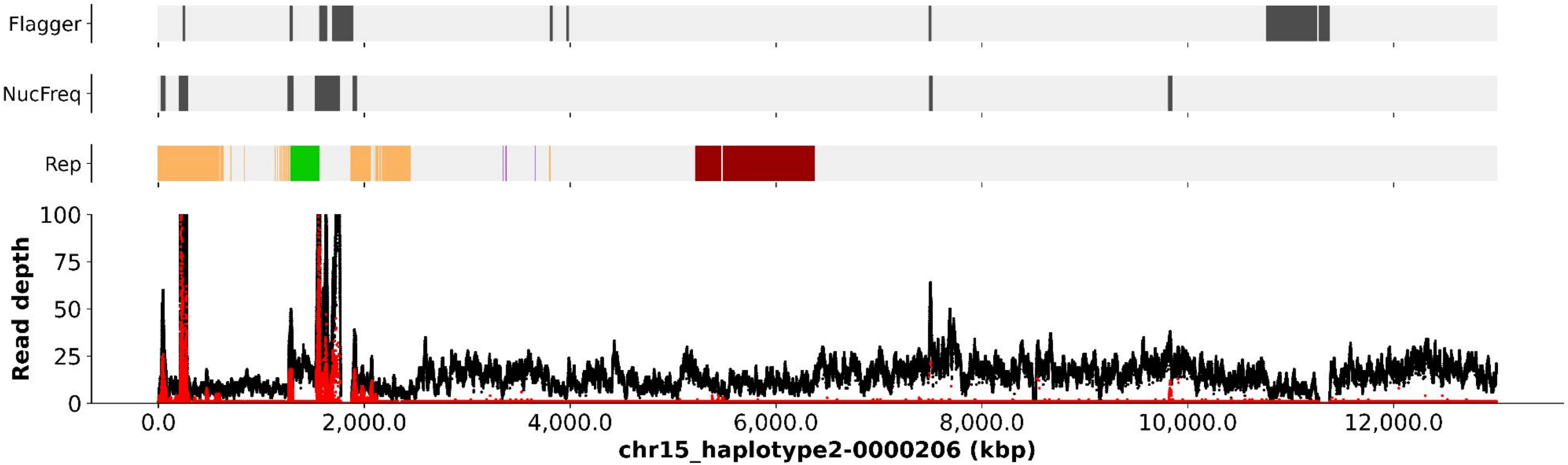
Flagger
NucFreq
Rep
Read depth
0.0 2,000.0 4,000.0 6,000.0 8,000.0 10,000.0 12,000.0
chr15_haplotype2-0000206 (kbp)

chr15_haplotype1-0000014

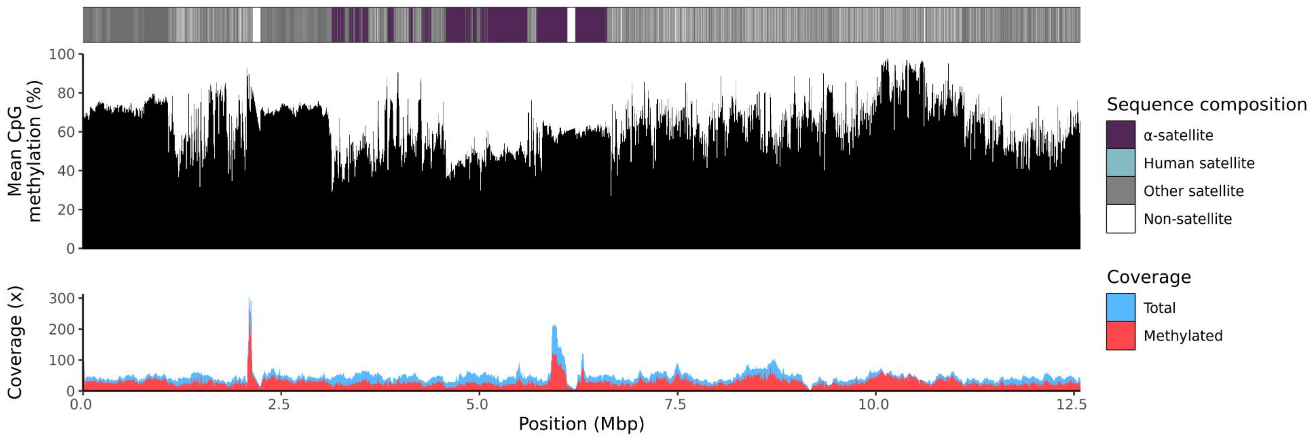


chr15_haplotype2-0000206

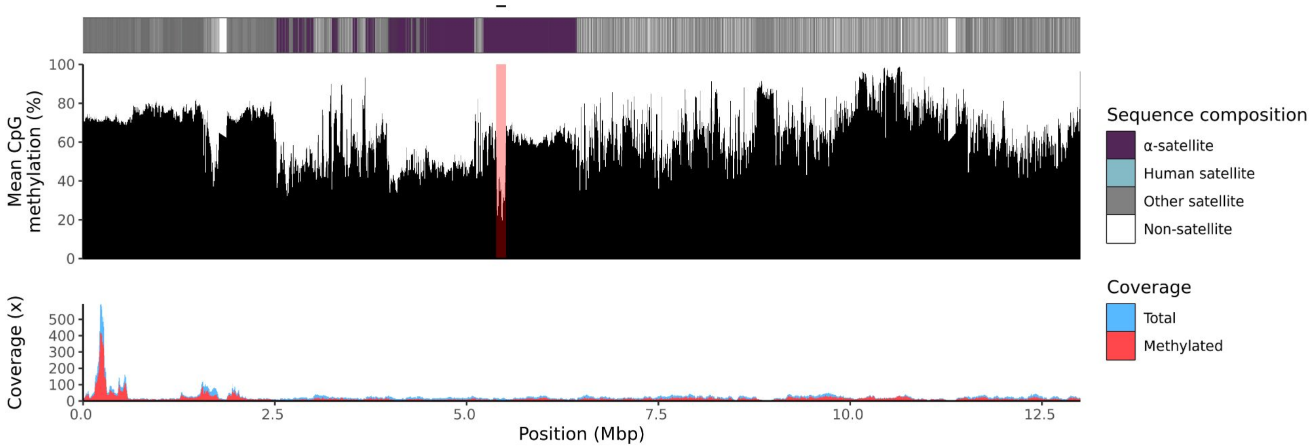

### chr21

#### K200087_1_haplotype1-0000020_chr21

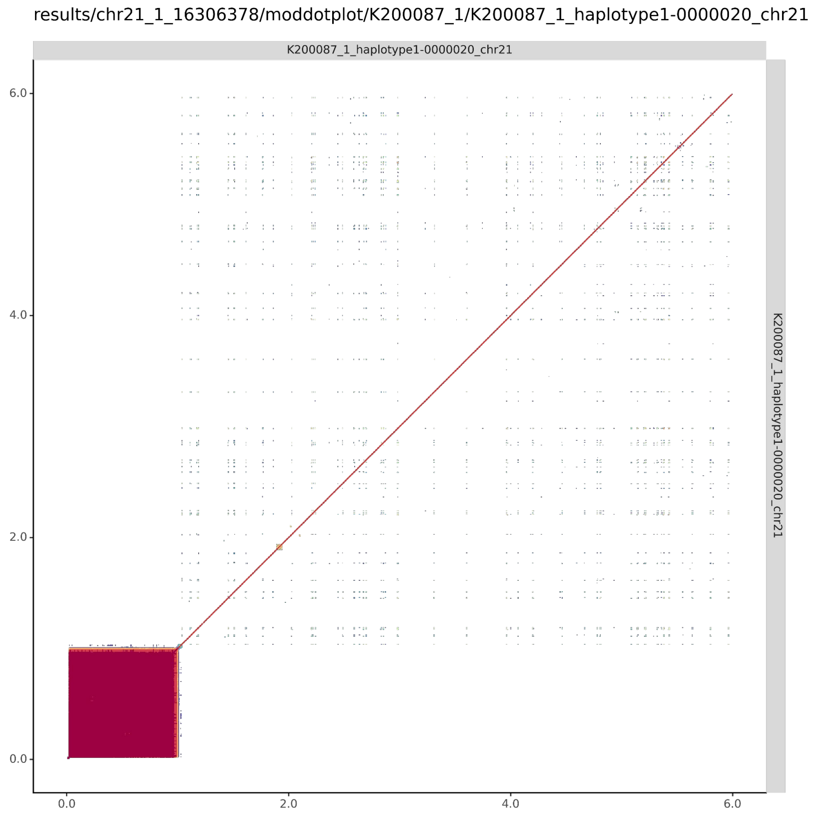


#### K200087_2_haplotype2-0000214_chr21

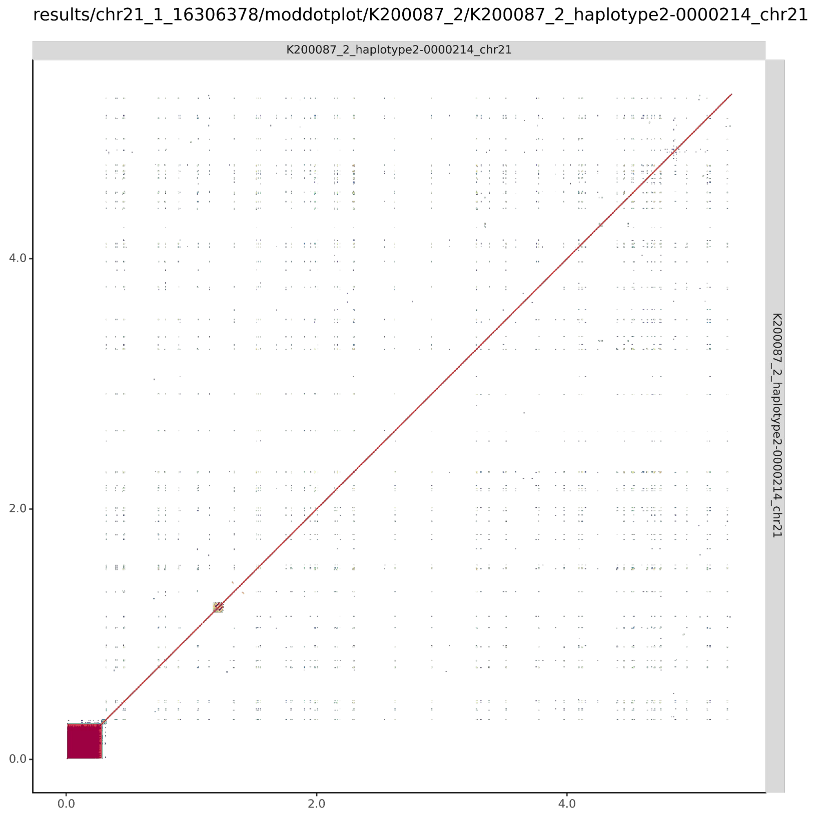

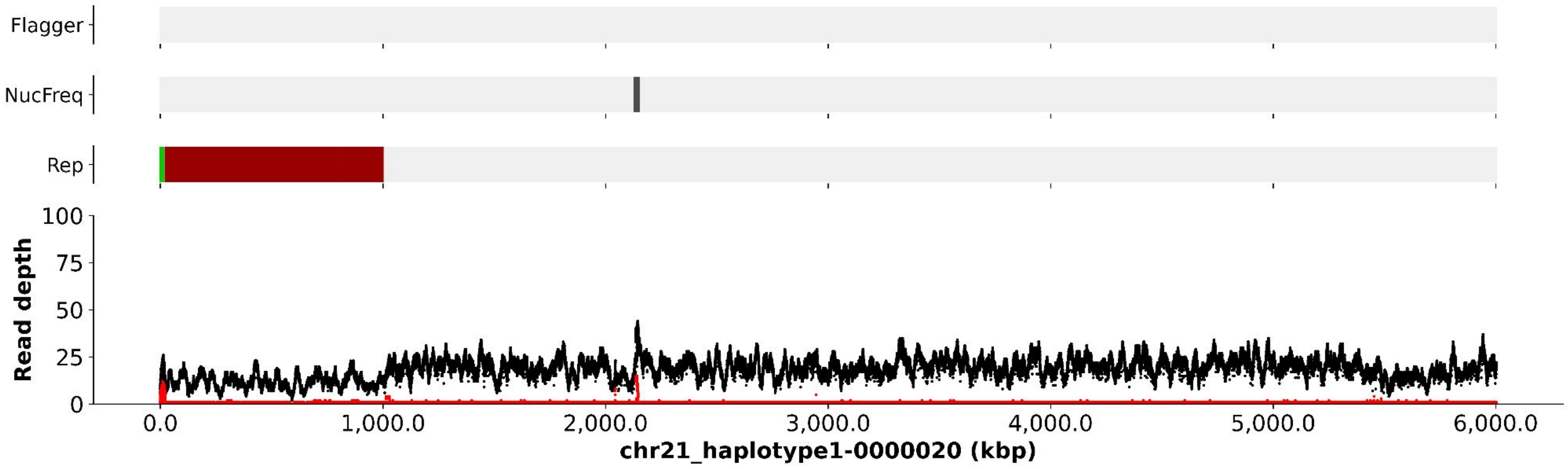
Flagger
NucFreq
Rep
Read depth
100
75
50
25
0
0.0
1,000.0
2,000.0
3,000.0
4,000.0
5,000.0
6,000.0
chr21_haplotype1-0000020 (kbp)

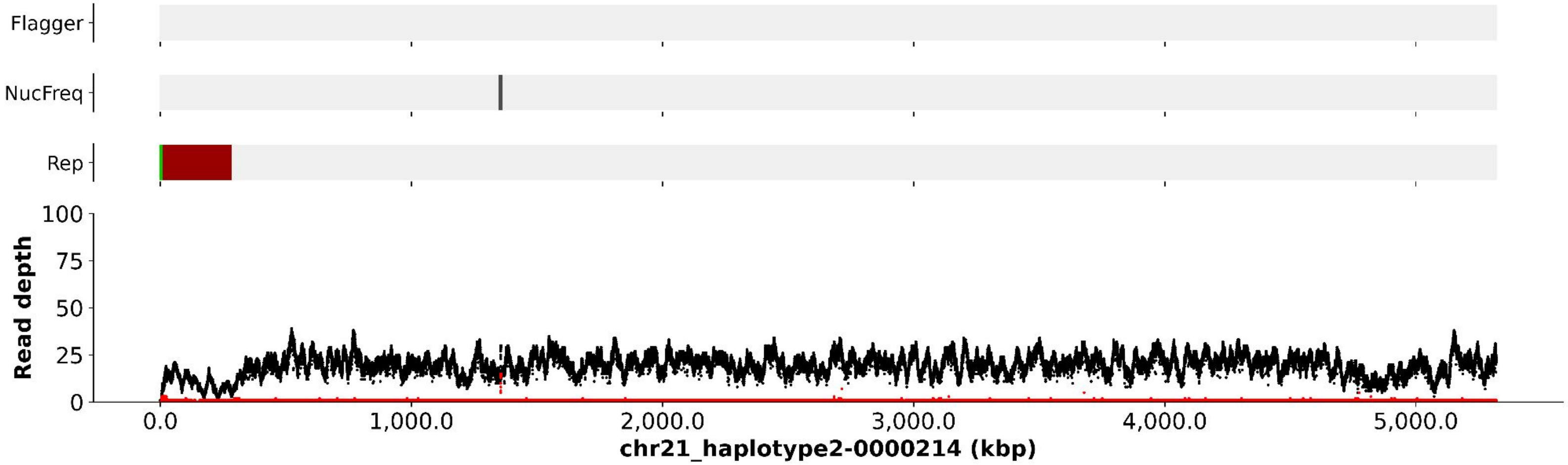
Flagger
NucFreq
Rep
Read depth
100
75
50
25
0
0.0
1,000.0
2,000.0
3,000.0
4,000.0
5,000.0
chr21_haplotype2-0000214 (kbp)

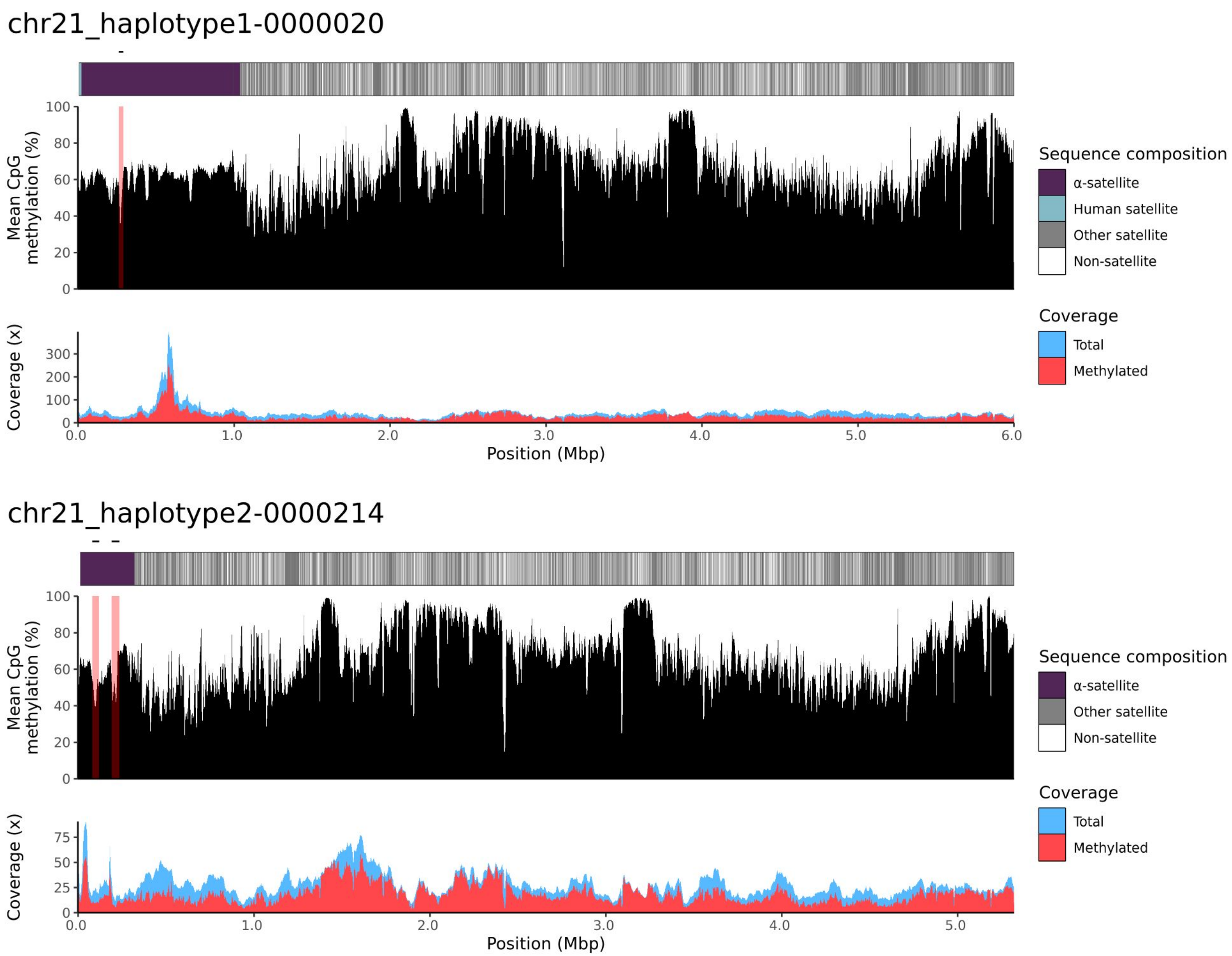
chr21_haplotype1-0000020
Mean CpG methylation (%)
Coverage (x)
Position (Mbp)
Sequence composition
α-satellite
Human satellite
Other satellite
Non-satellite
Coverage
Total
Methylated
chr21_haplotype2-0000214
Mean CpG methylation (%)
Coverage (x)
Position (Mbp)
Sequence composition
α-satellite
Other satellite
Non-satellite
Coverage
Total
Methylated

### chr22

#### K200087_1_haplotype1-0000031_chr22

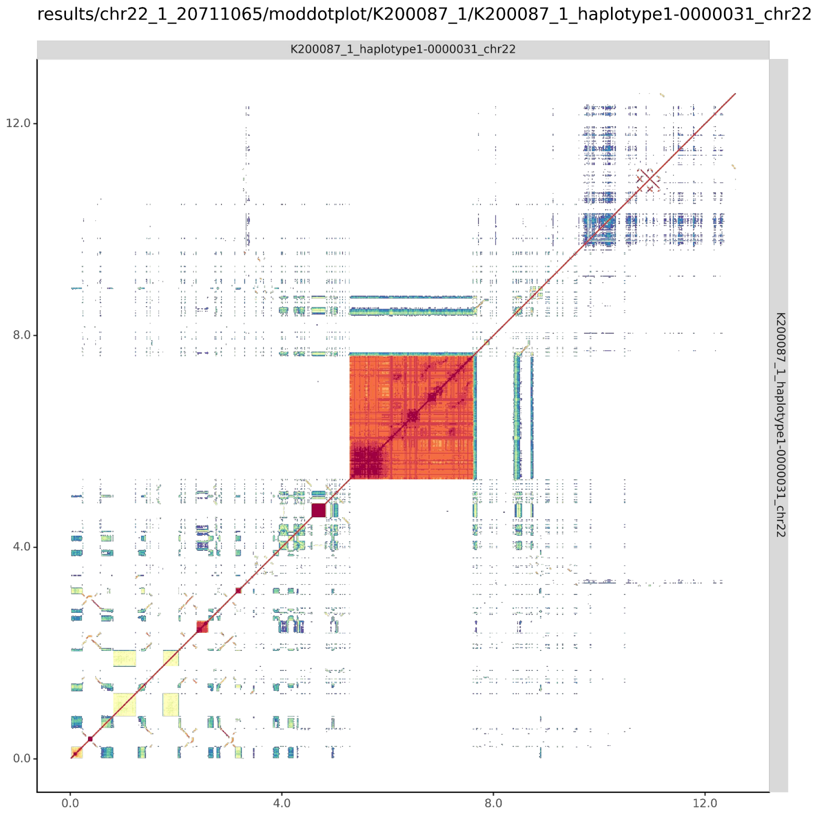


#### K200087_2_haplotype2-0000227_chr22

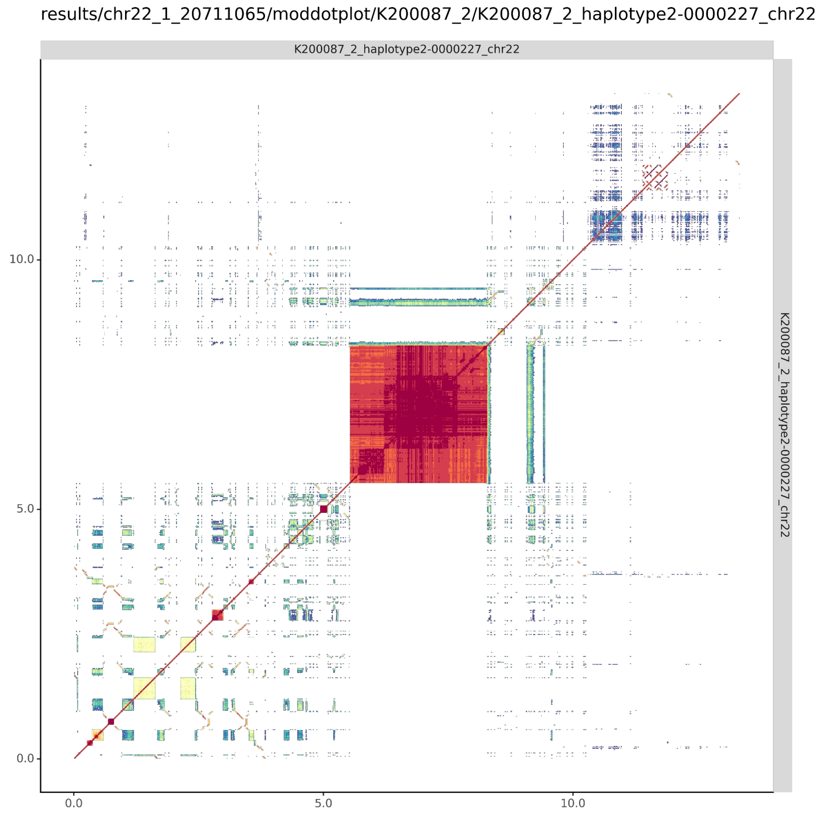

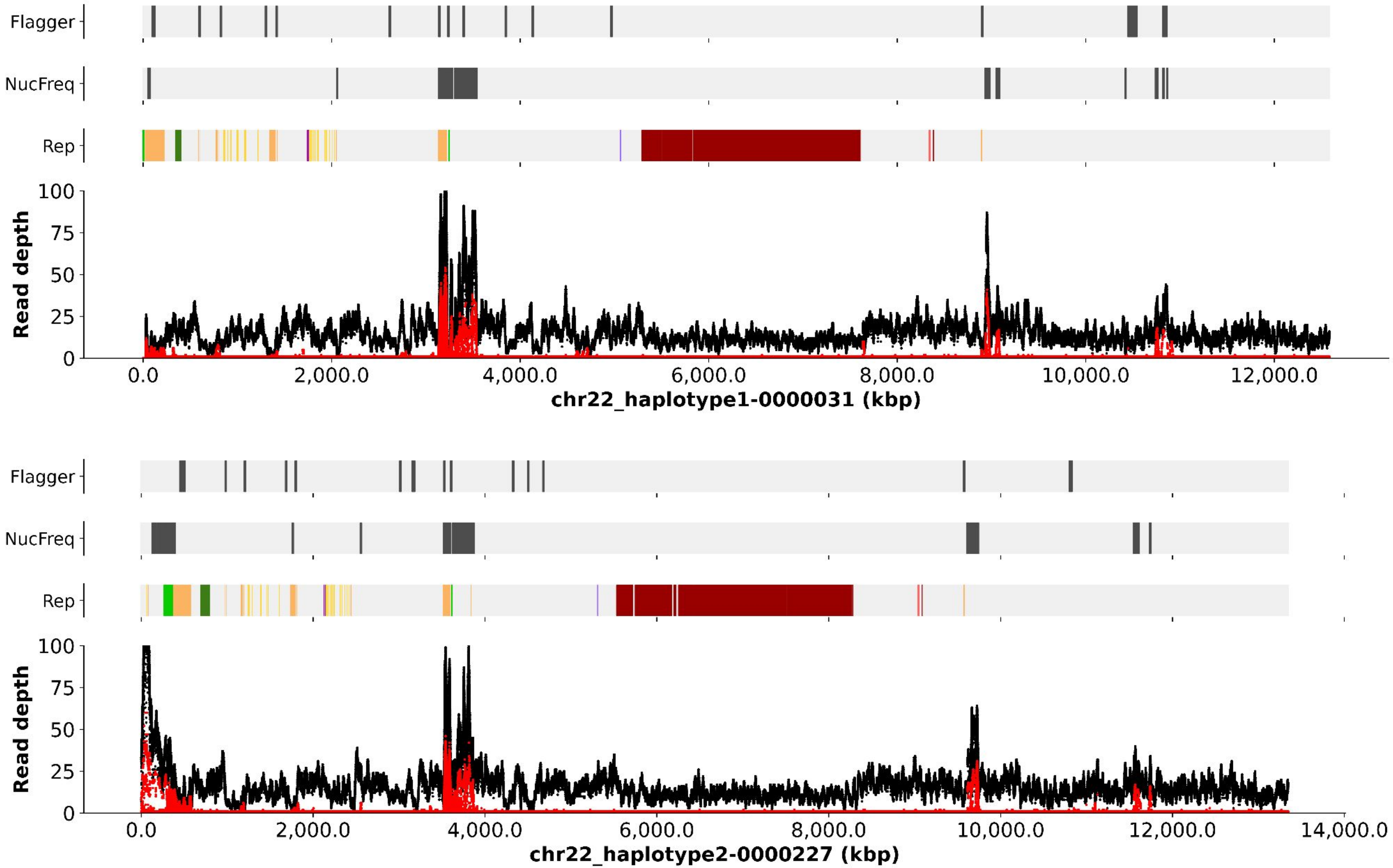
Flagger
NucFreq
Rep
Read depth
100
75
50
25
0
0.0
2,000.0
4,000.0
6,000.0
8,000.0
10,000.0
12,000.0
chr22_haplotype1-0000031 (kbp)
Flagger
NucFreq
Rep
Read depth
100
75
50
25
0
0.0
2,000.0
4,000.0
6,000.0
8,000.0
10,000.0
12,000.0
14,000.0
chr22_haplotype2-0000227 (kbp)

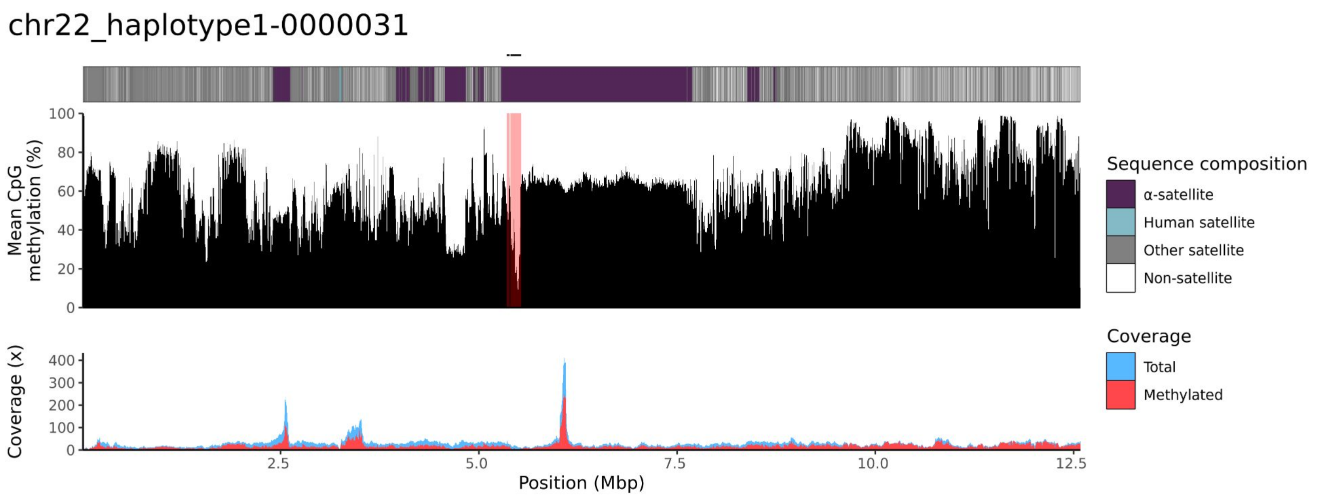
chr22_haplotype1-0000031
Mean CpG methylation (%)
0
20
40
60
80
100
Coverage (x)
0
100
200
300
400
2.5
5.0
7.5
10.0
12.5
Position (Mbp)
Sequence composition
α-satellite
Human satellite
Other satellite
Non-satellite
Coverage
Total
Methylated

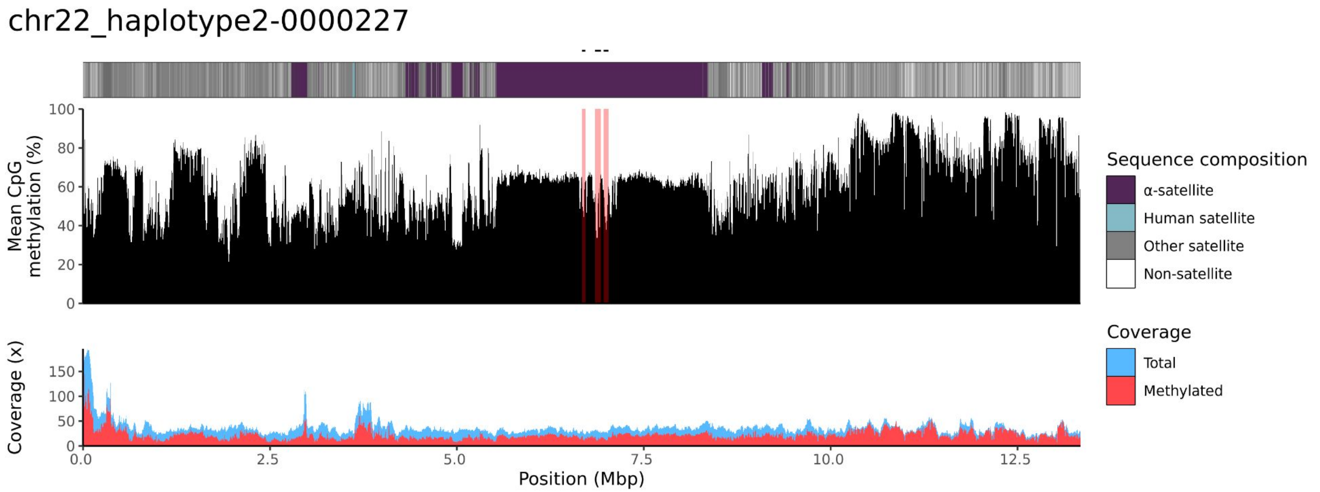
chr22_haplotype2-0000227
Mean CpG methylation (%)
0
20
40
60
80
100
Coverage (x)
0
50
100
150
0.0
2.5
5.0
7.5
10.0
12.5
Position (Mbp)
Sequence composition
α-satellite
Human satellite
Other satellite
Non-satellite
Coverage
Total
Methylated
