## Supplementary material for "Human acrocentric chromosome short arm *de novo* mutation and recombination": Data S3

- This file includes all IGV screenshots of DNMs listed in Table S3.
- IGV screenshots show the alignment of offspring and parental reads aligned to the parental reference sequence.
- For each IGV screenshot, the tracks from top to bottom are alignments from child’s HiFi, child’s UL-ONT, parent’s HiFi and parent’s UL-ONT.

### Single-nucleotide variants

### NA12884_NA12878_1_haplotype1-0000005_chr13_423702_T_C

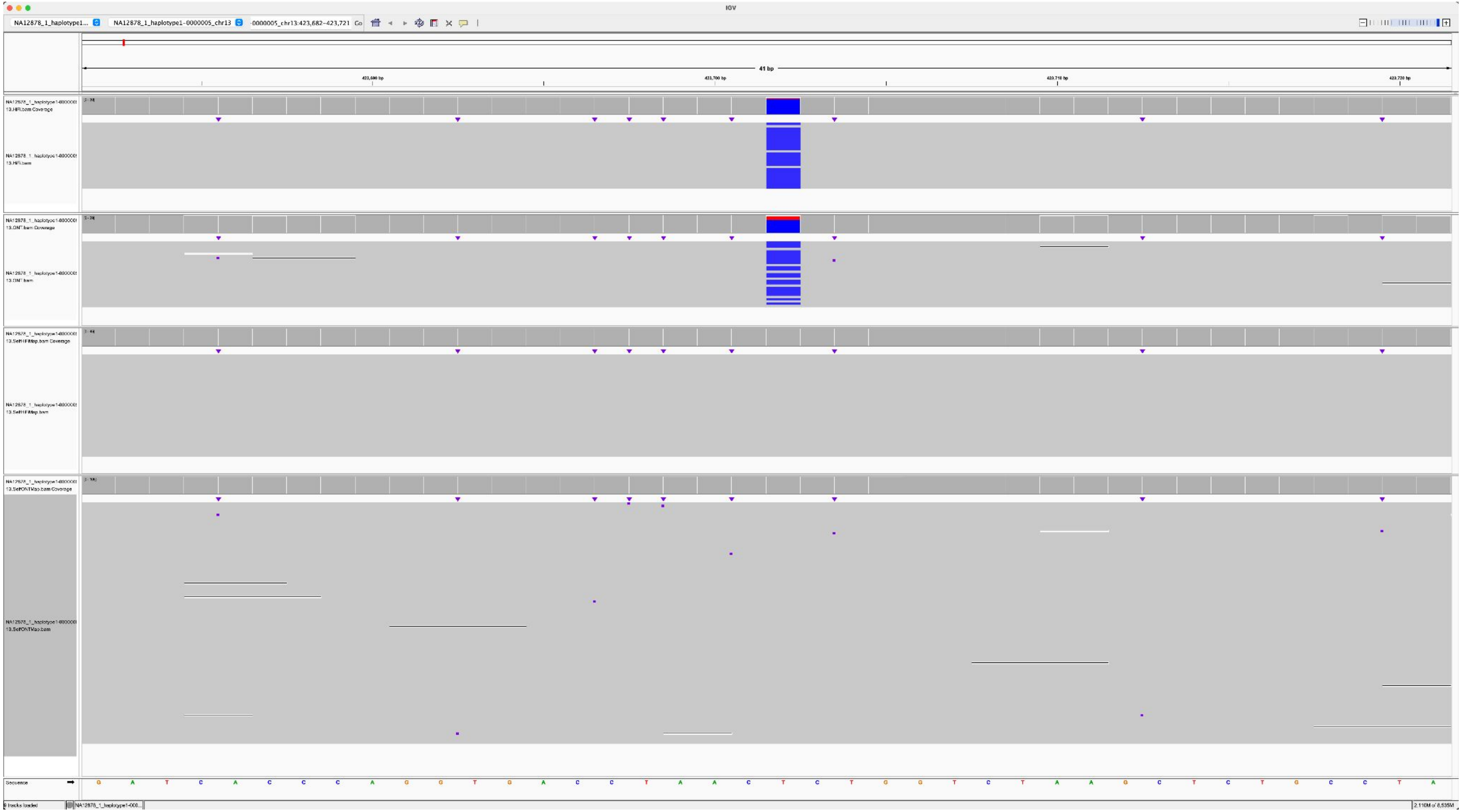

### NA12883_NA12878_1_haplotype1-0000008_chr22_13251153_A_T

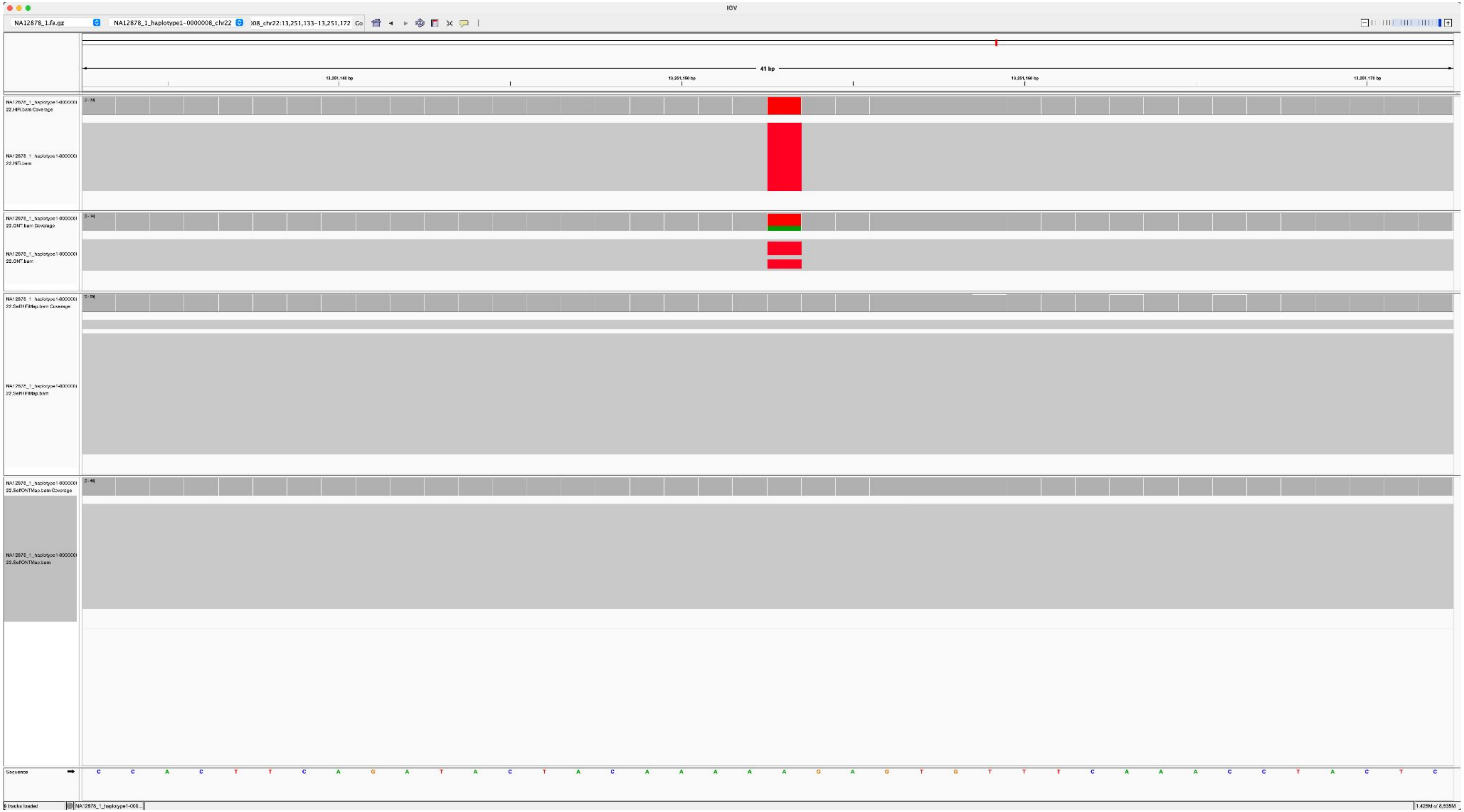

### NA12881_NA12878_2_haplotype2-0000030_chr13_4561495_T_C

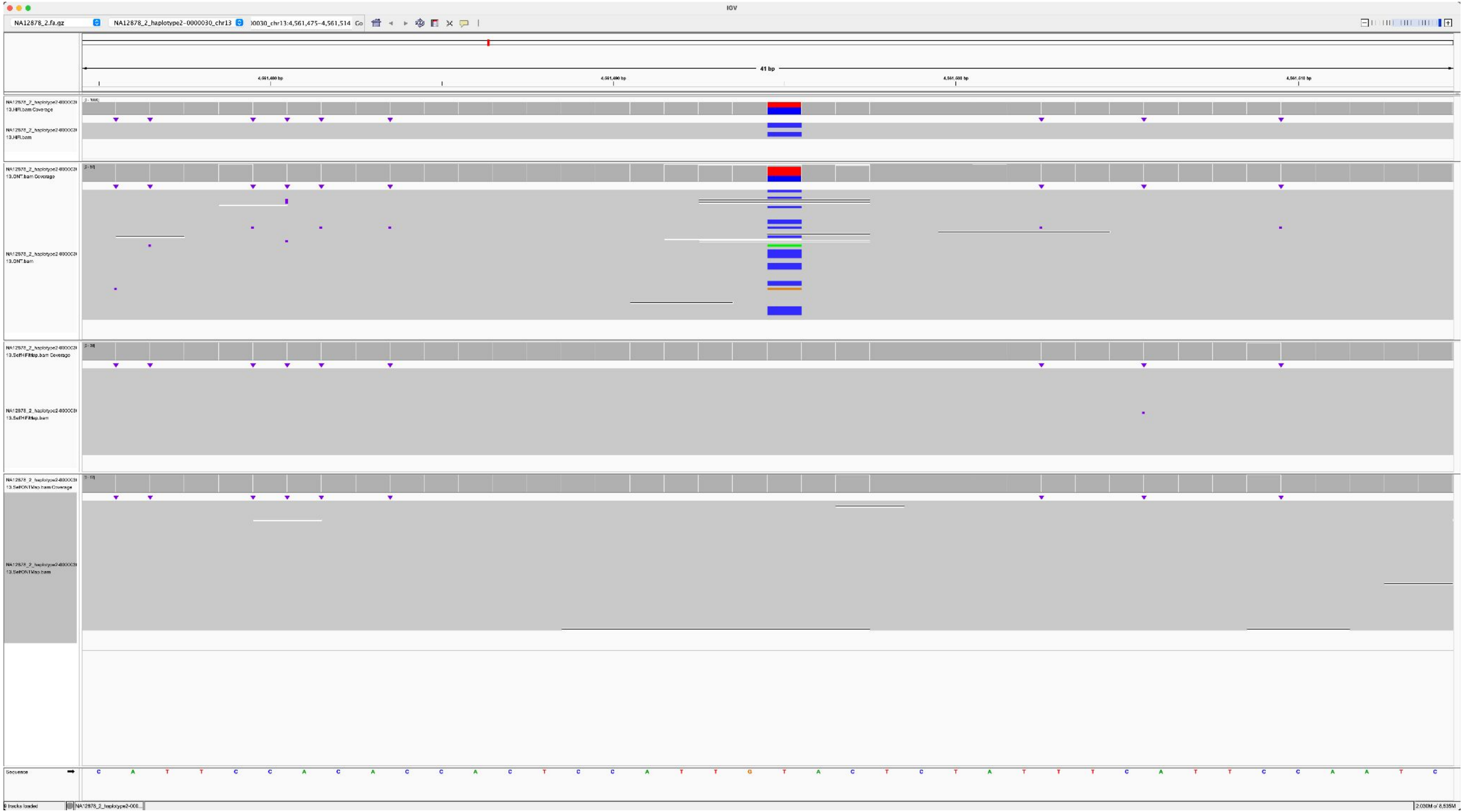

### NA12882_NA12878_2_haplotype2-0000030_chr13_4192721_C_G

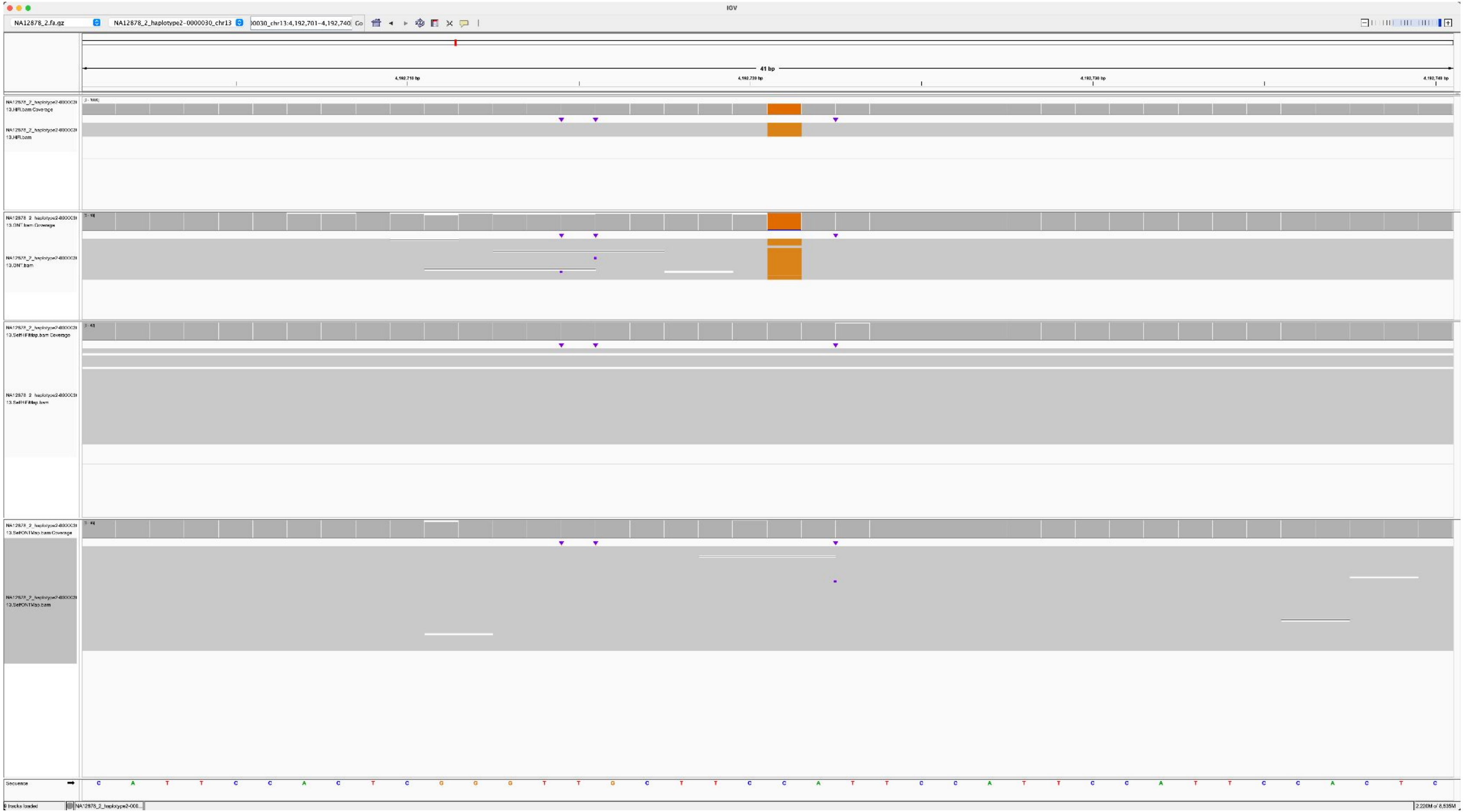

### NA12885_NA12878_2_haplotype2-0000041_chr14_11092901_A_T

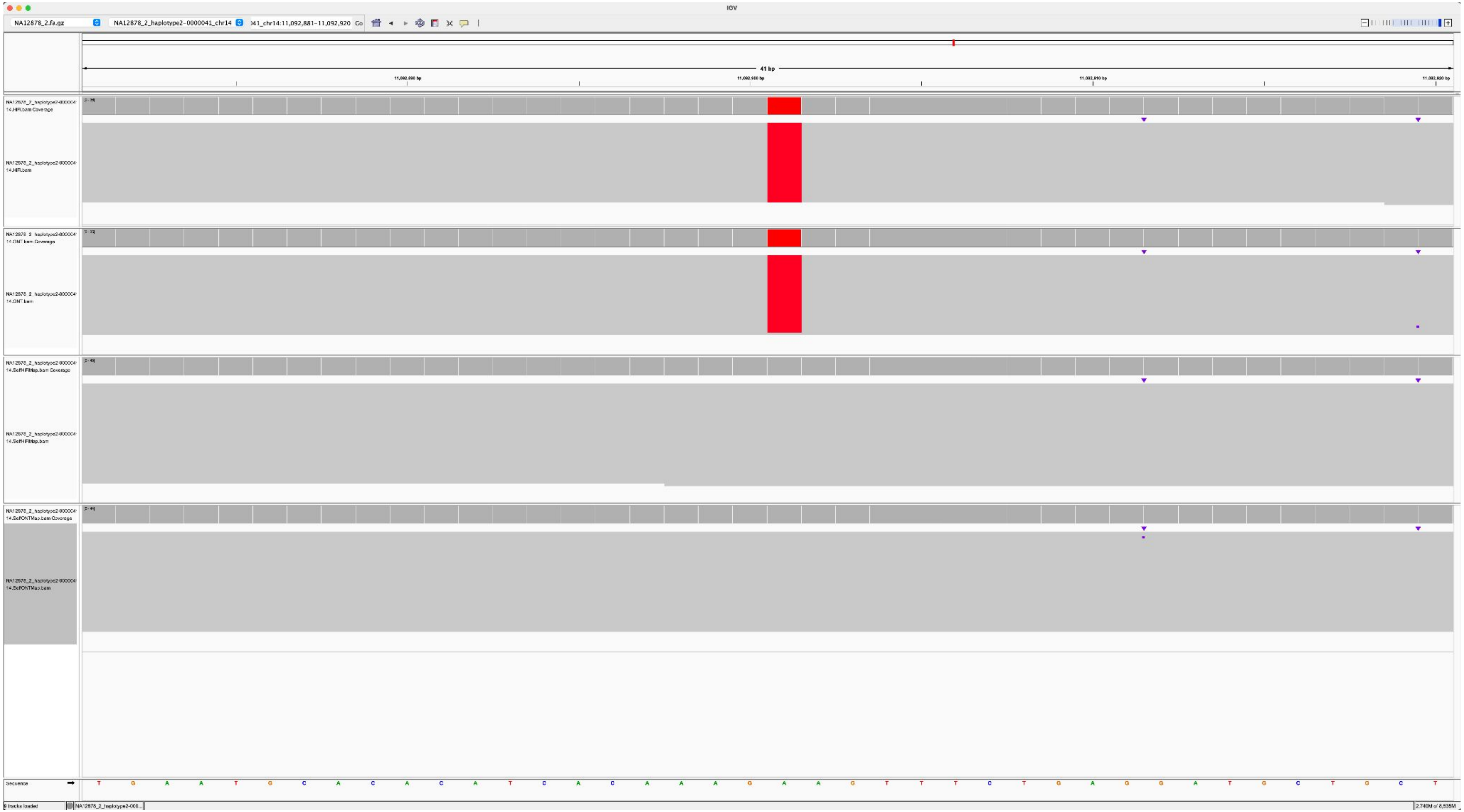

### NA12887_NA12878_2_haplotype2-0000041_chr14_9548449_G_A

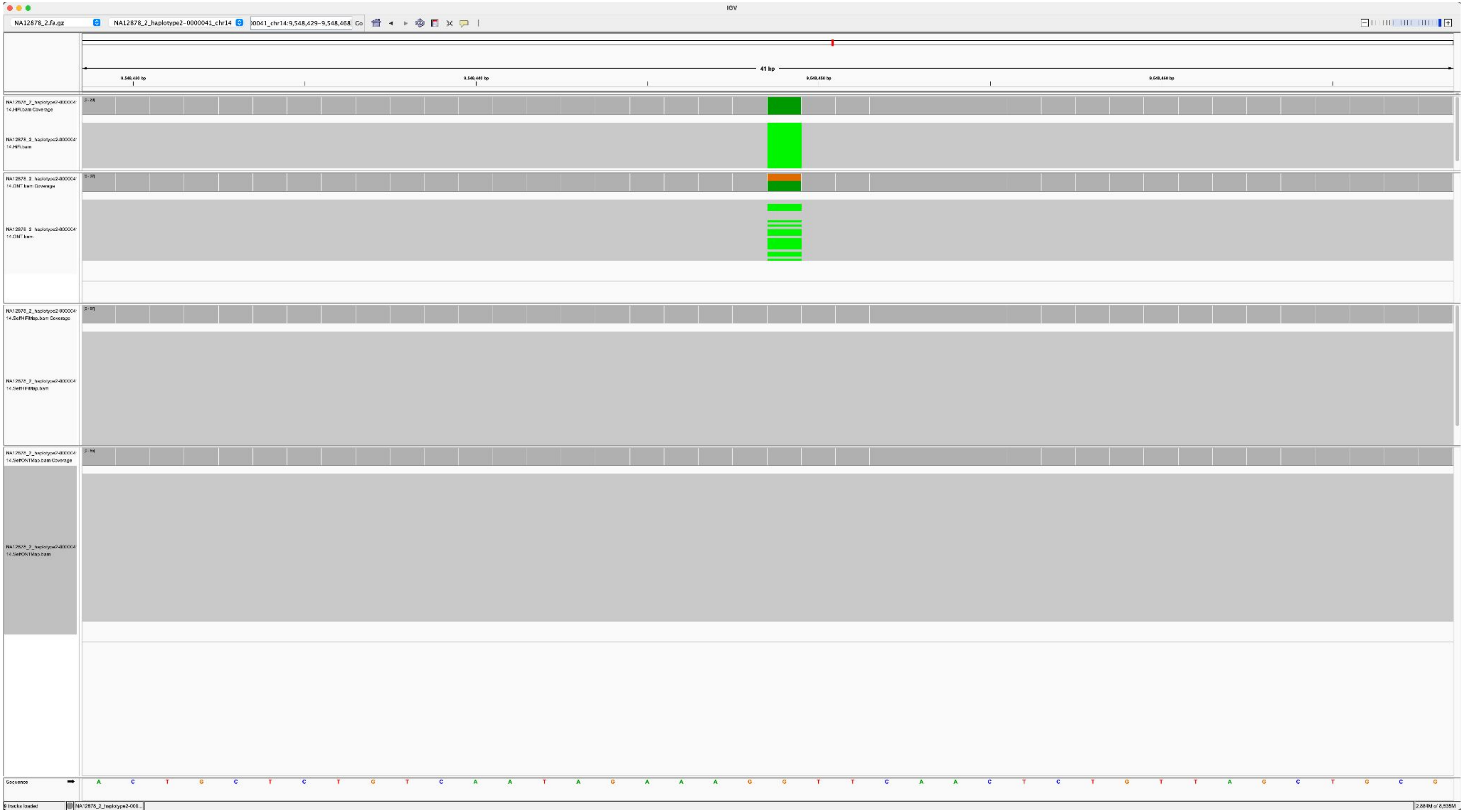

### NA12887_NA12878_2_haplotype2-0000041_chr14_11092598_A_G

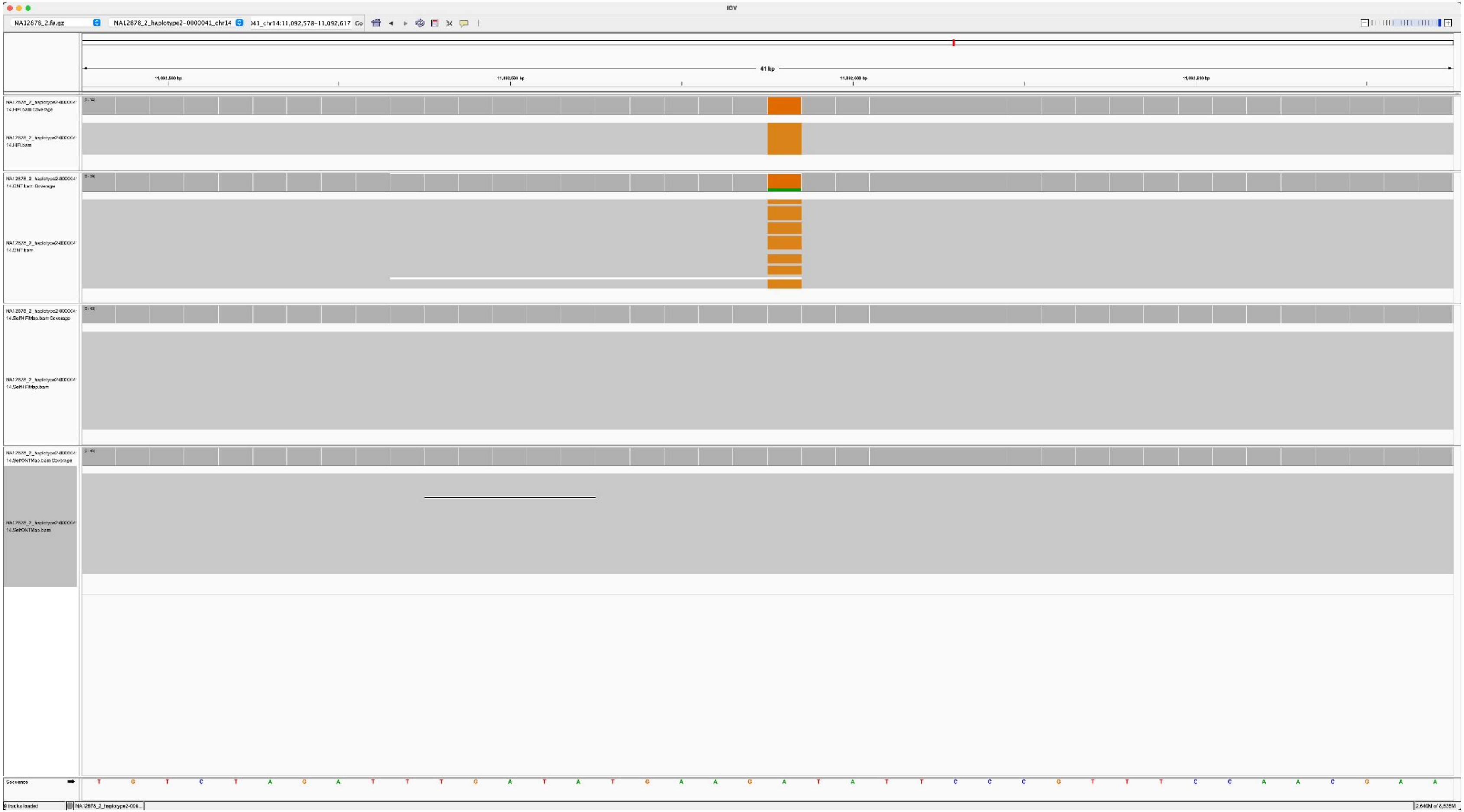

### NA12882_NA12878_2_haplotype2-0000044_chr15_7014450_A_G

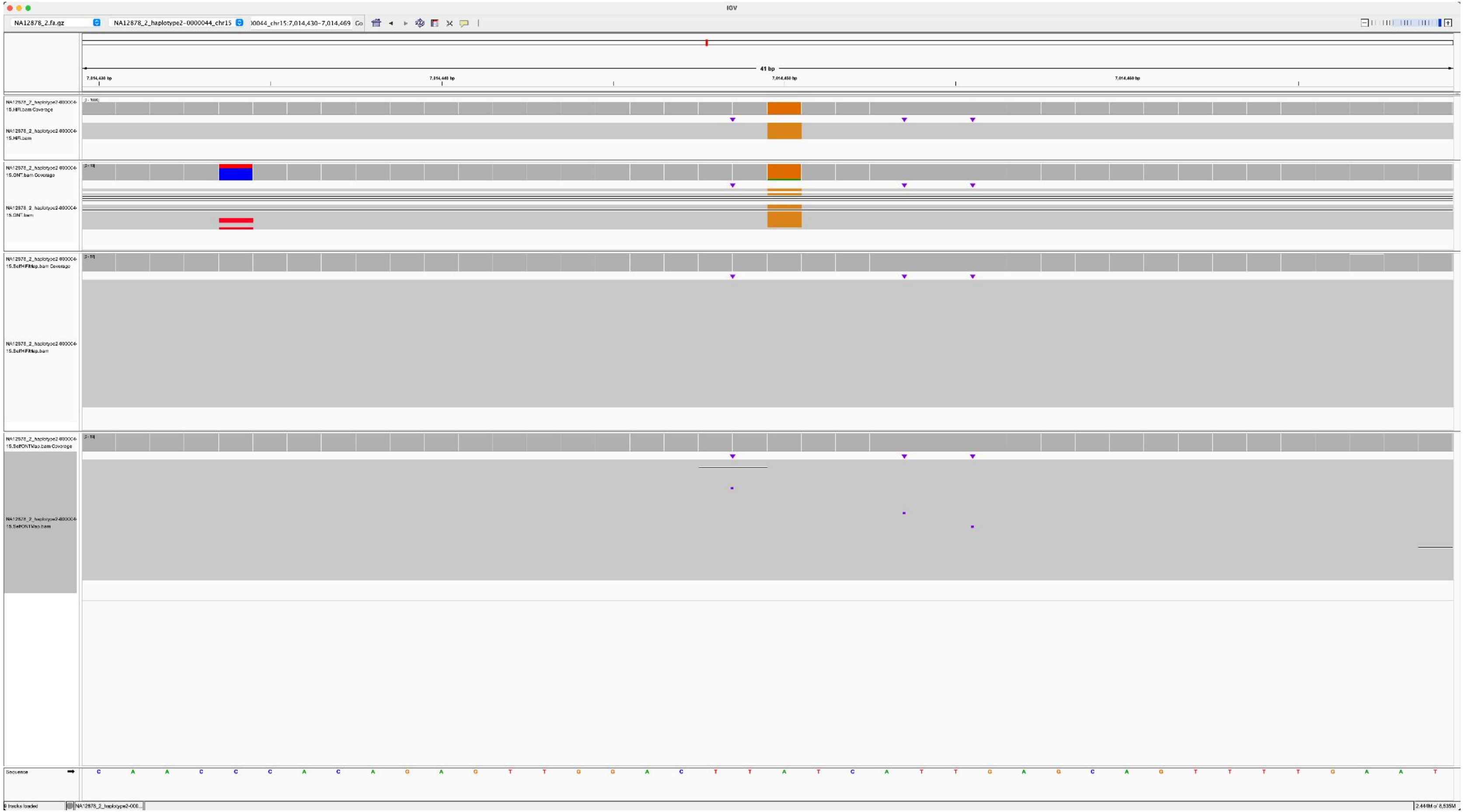

### NA12883_NA12878_2_haplotype2-0000044_chr15_9110709_A_T

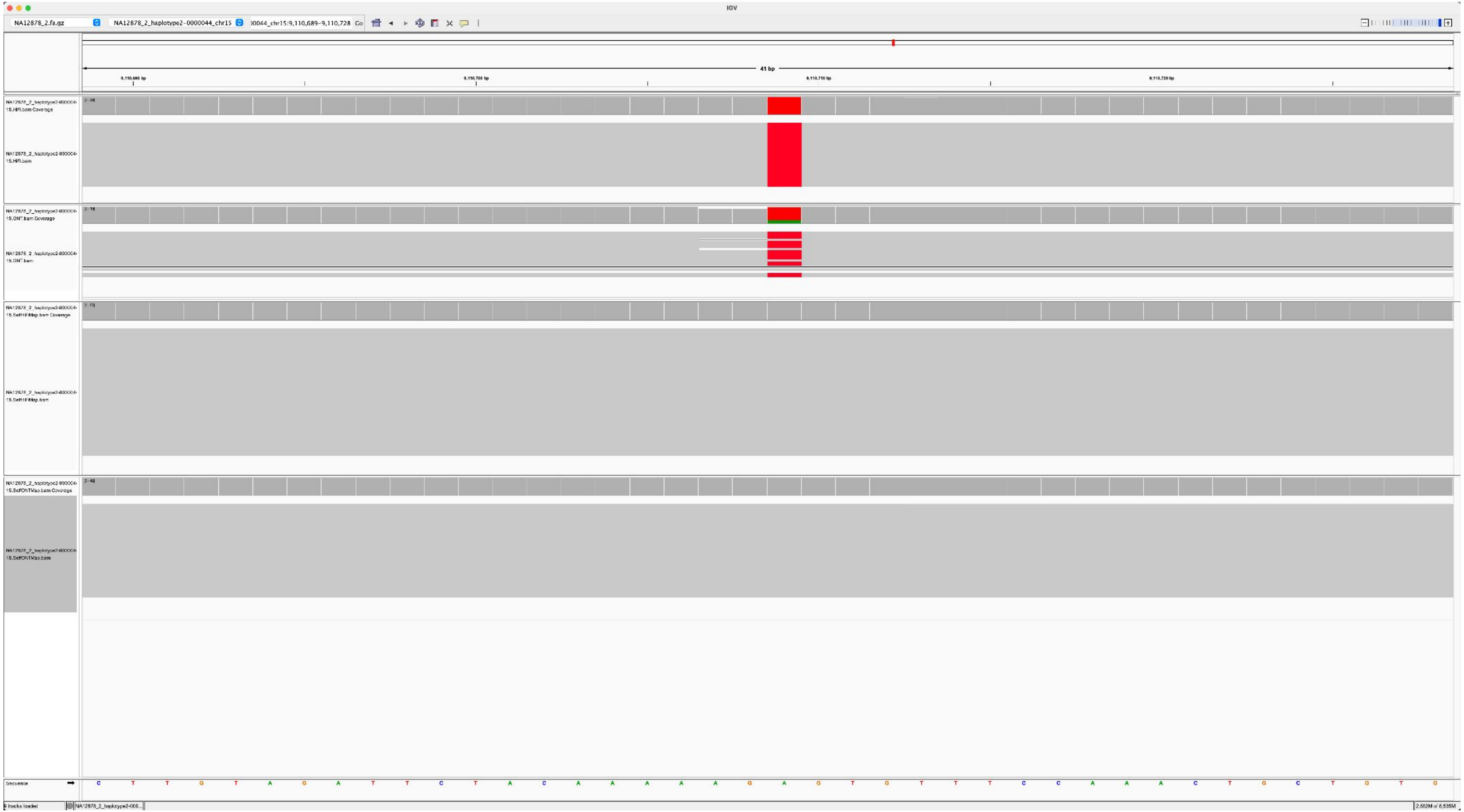

### NA12886_NA12878_2_haplotype2-0000044_chr15_7677712_T_C

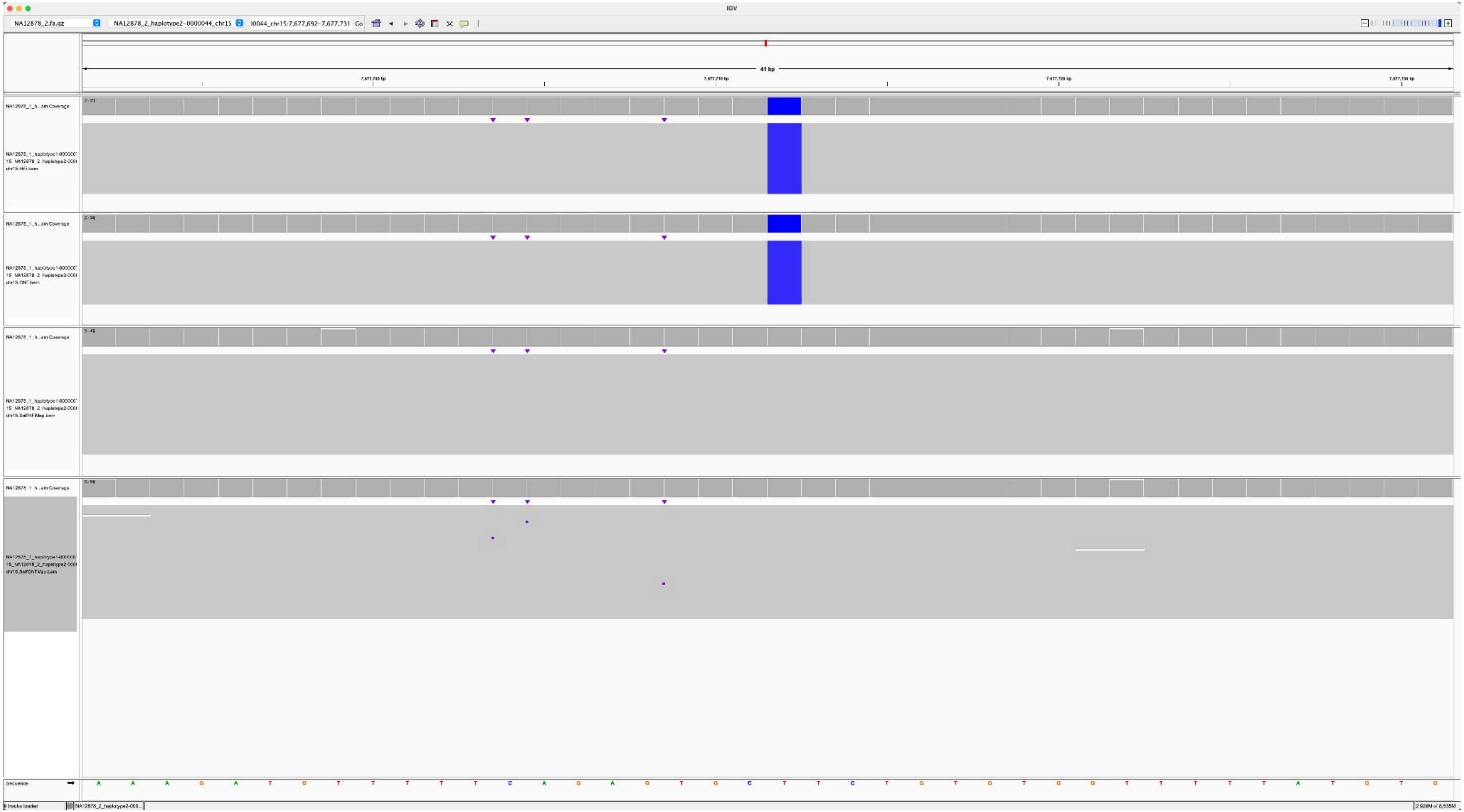

### NA12884_NA12878_2_haplotype2-0000032_chr21_3619981_G_A

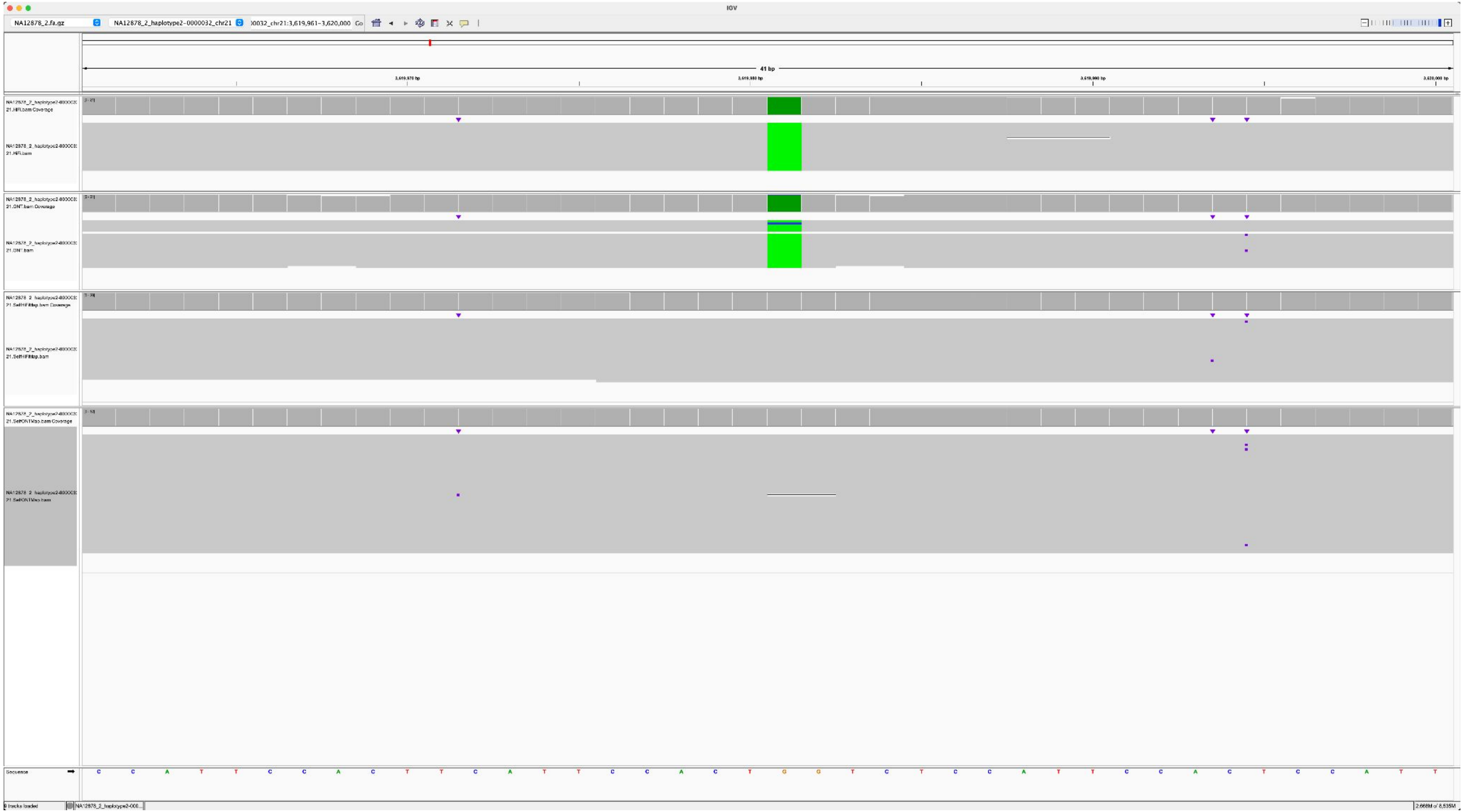

### NA12885_NA12878_2_haplotype2-0000032_chr21_3790057_G_A

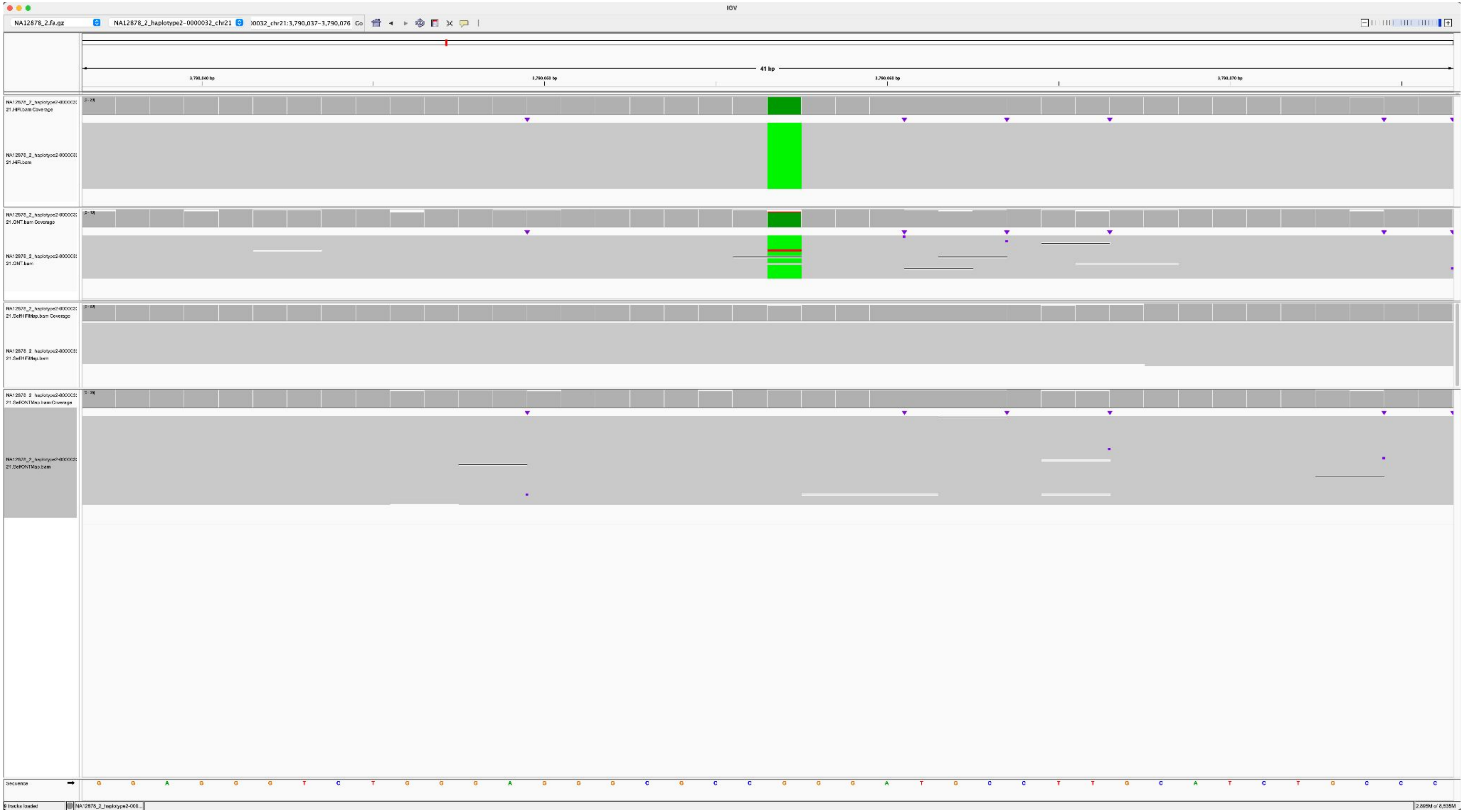

NA12887_NA12878_2_haplotype2-0000032_chr21_6951927_T_C

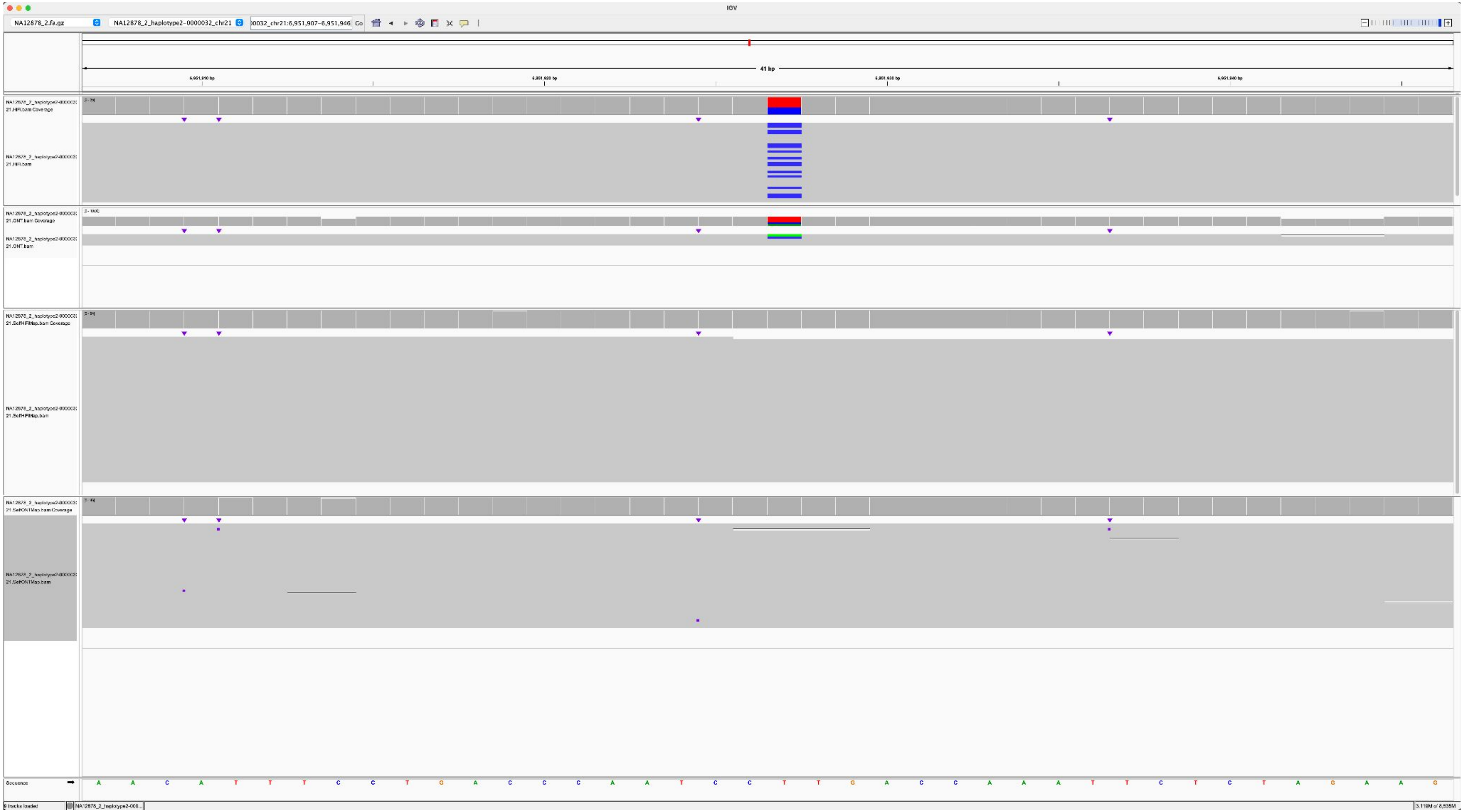

### NA12885_NA12877_1_haplotype1-0000001_chr13_7731652_A_C

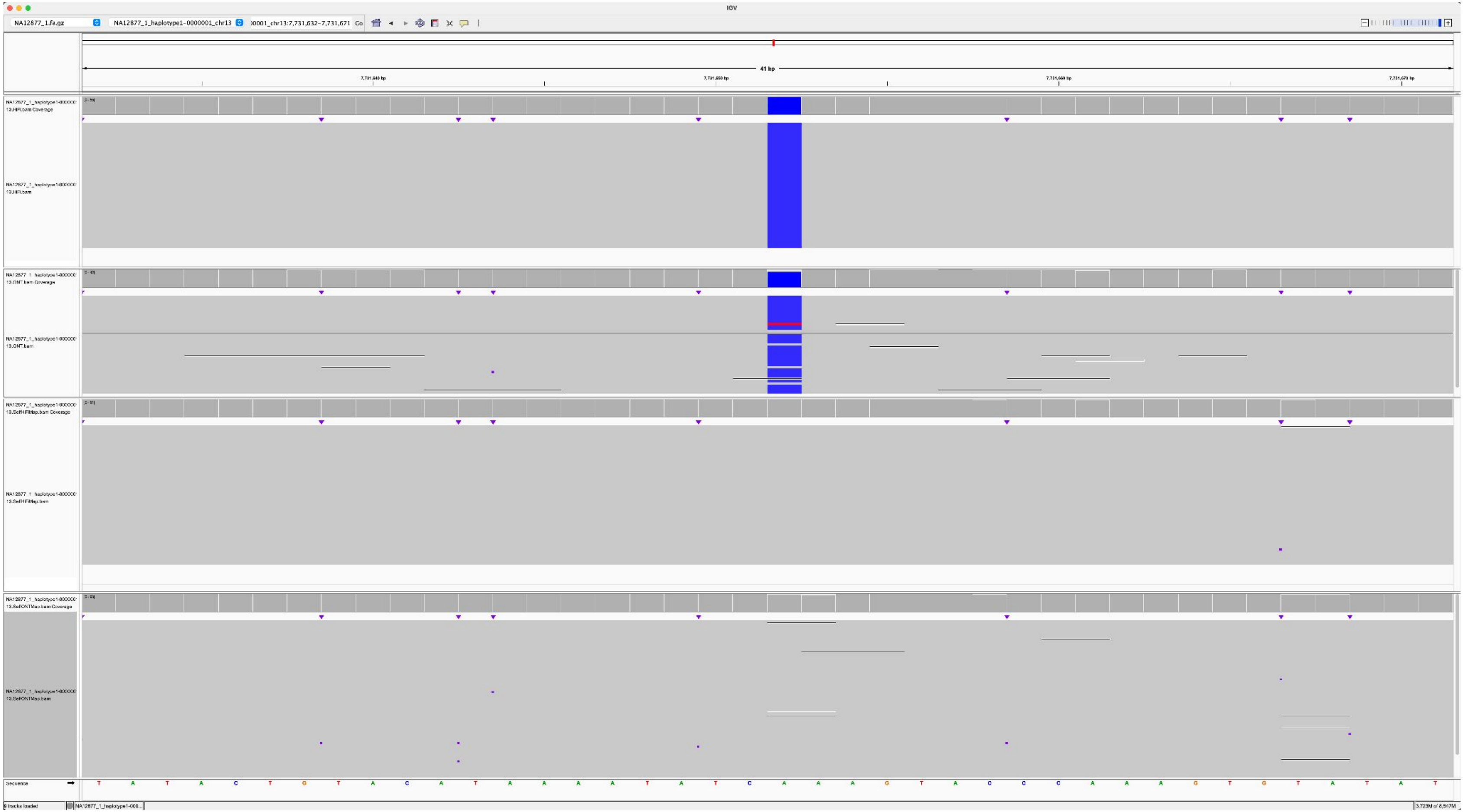

NA12885_NA12877_1_haplotype1-0000001_chr13_7819035_C_G

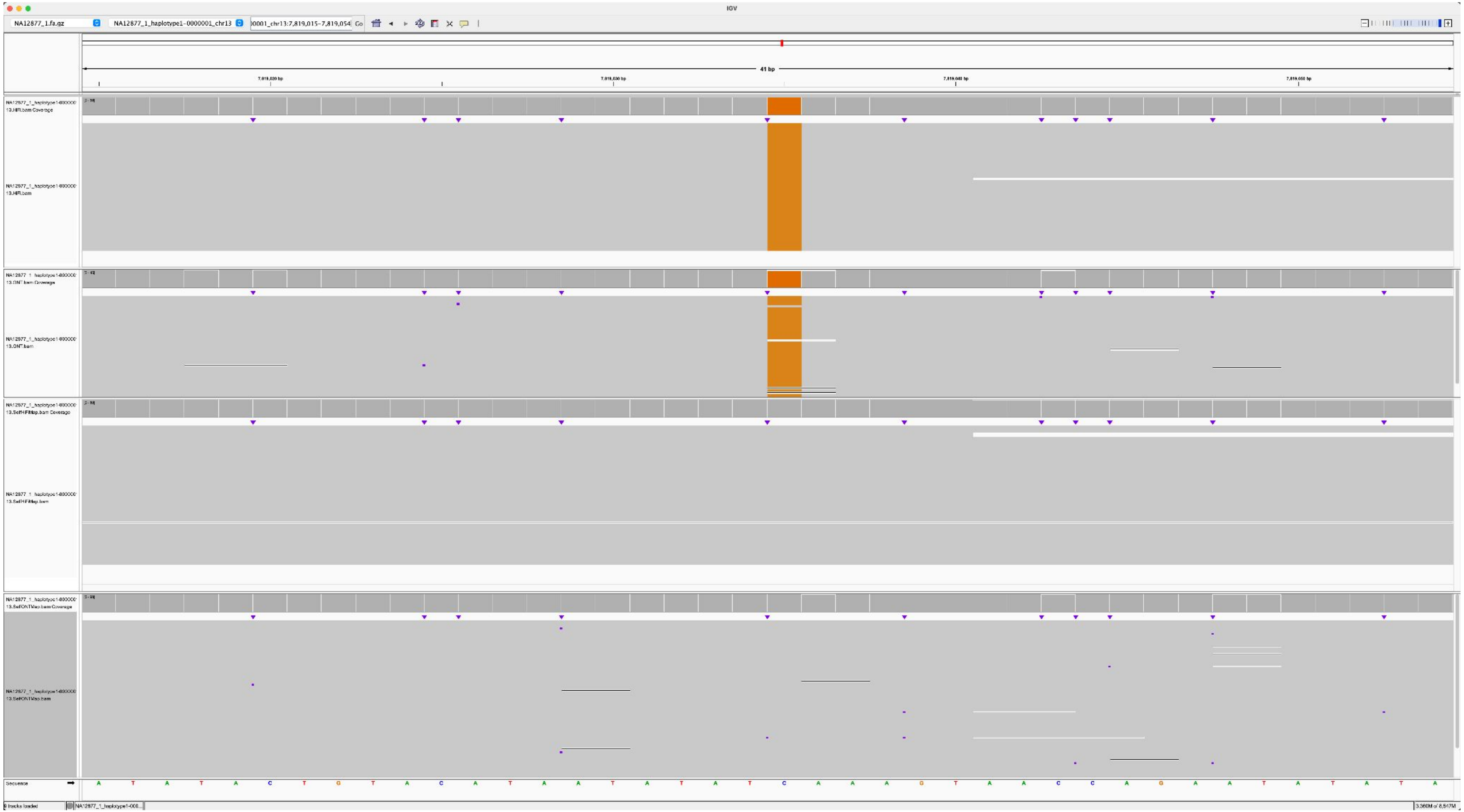

### NA12885_NA12877_1_haplotype1-0000001_chr13_8313063_A_C

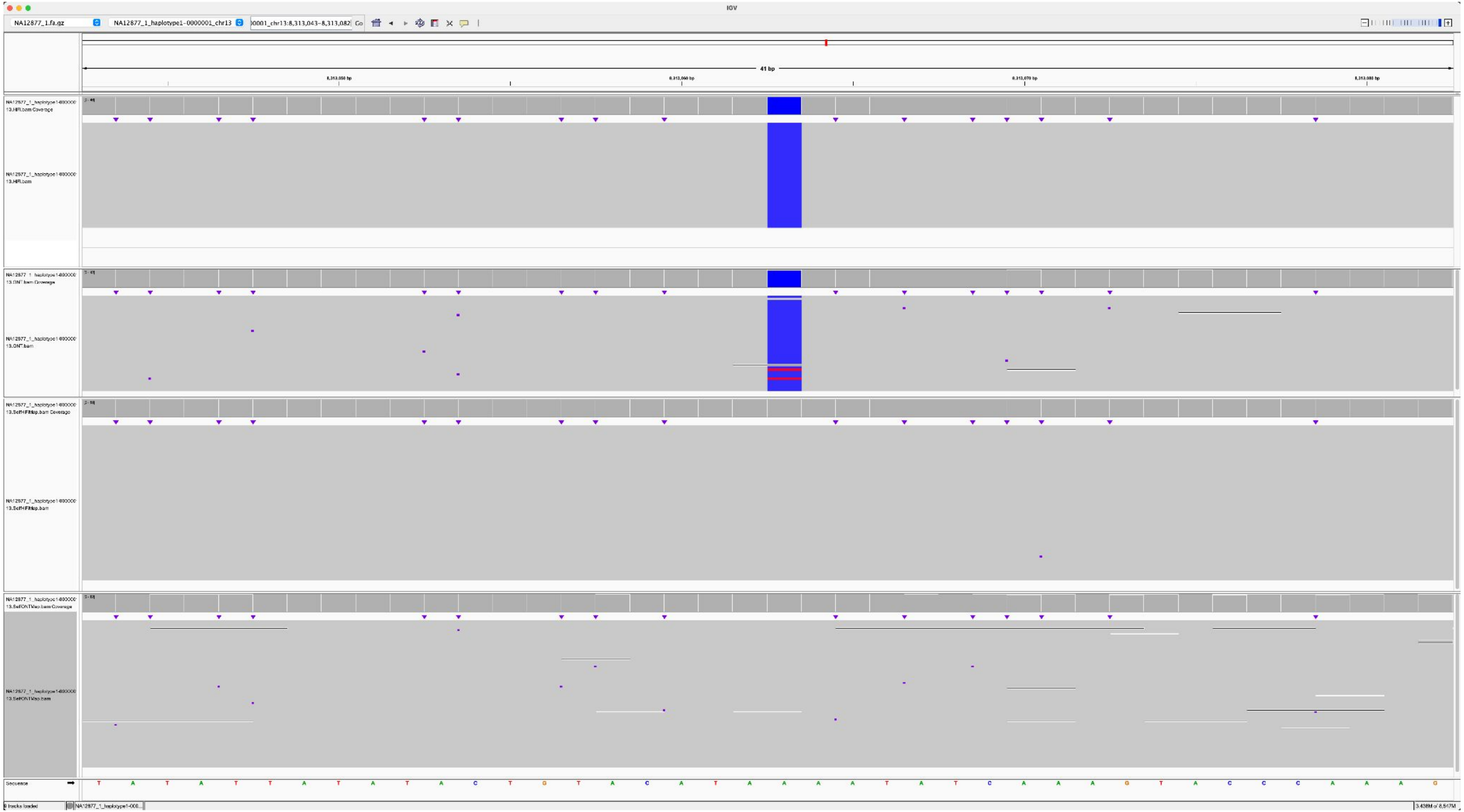

### NA12885_NA12877_1_haplotype1-0000001_chr13_161049_T_G

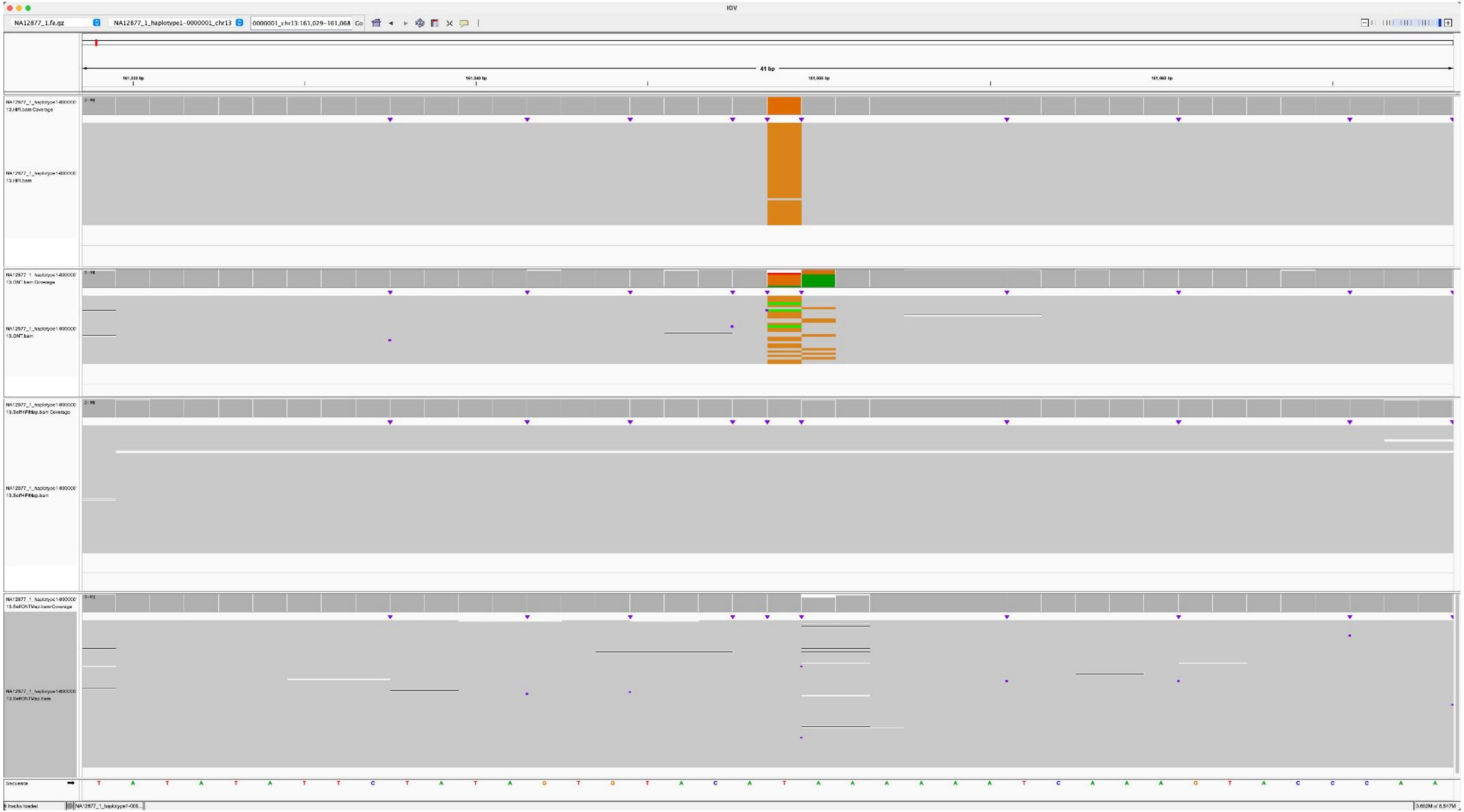

### NA12886_NA12877_1_haplotype1-0000001_chr13_9391278_C_T

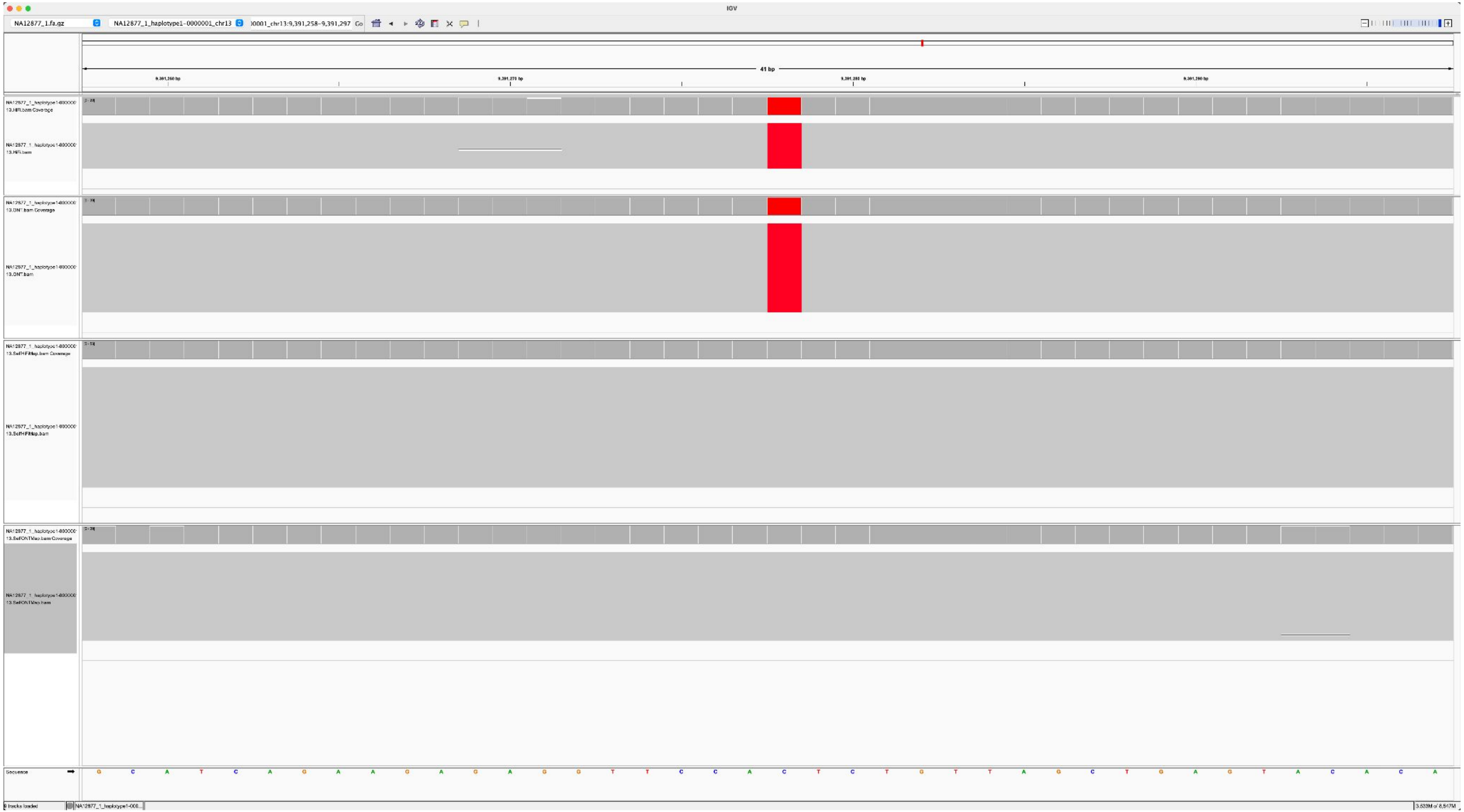

NA12883_NA12877_1_haplotype1-0000013_chr14_10882872_G_A

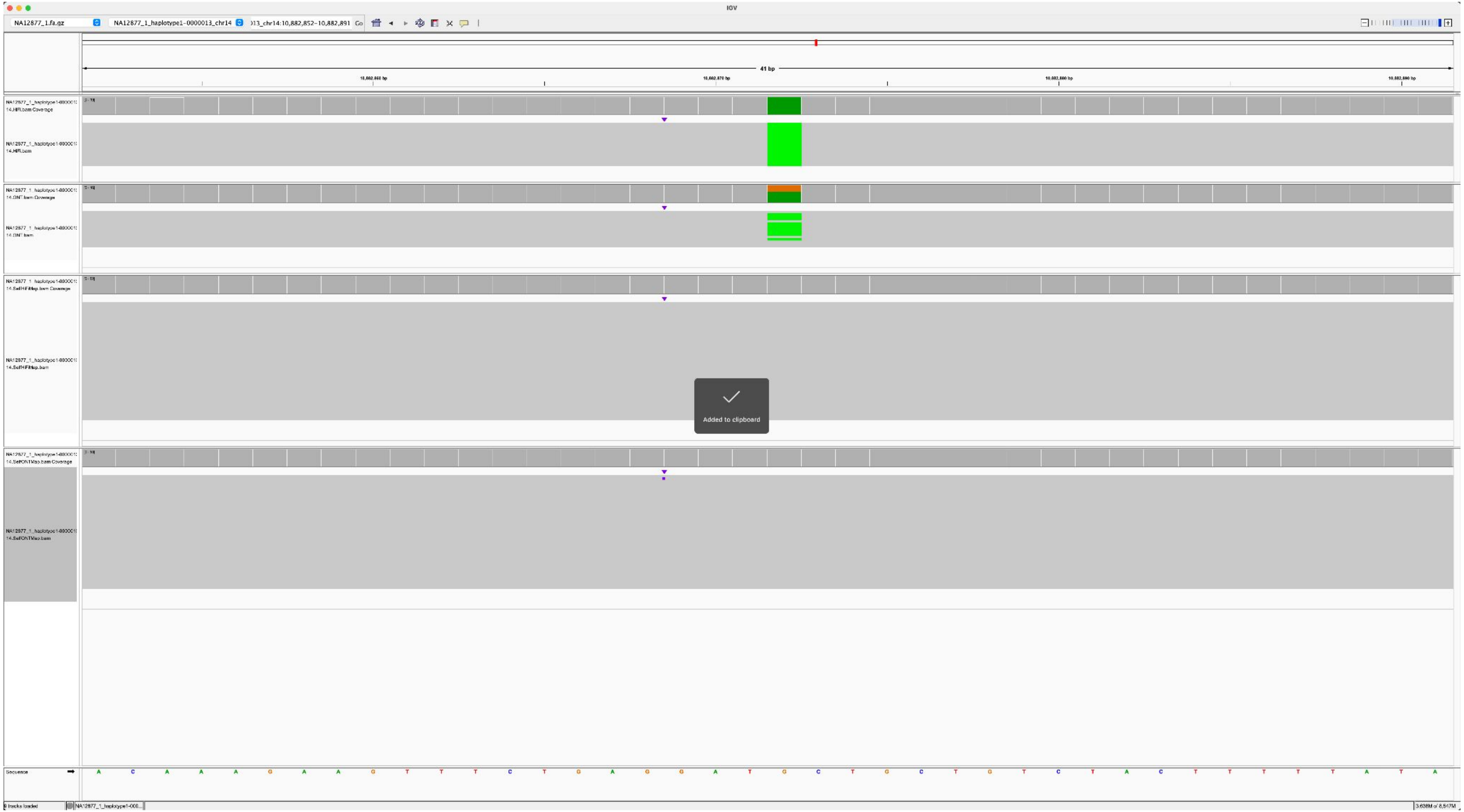

### NA12883_NA12877_1_haplotype1-0000013_chr14_5545018_T_C

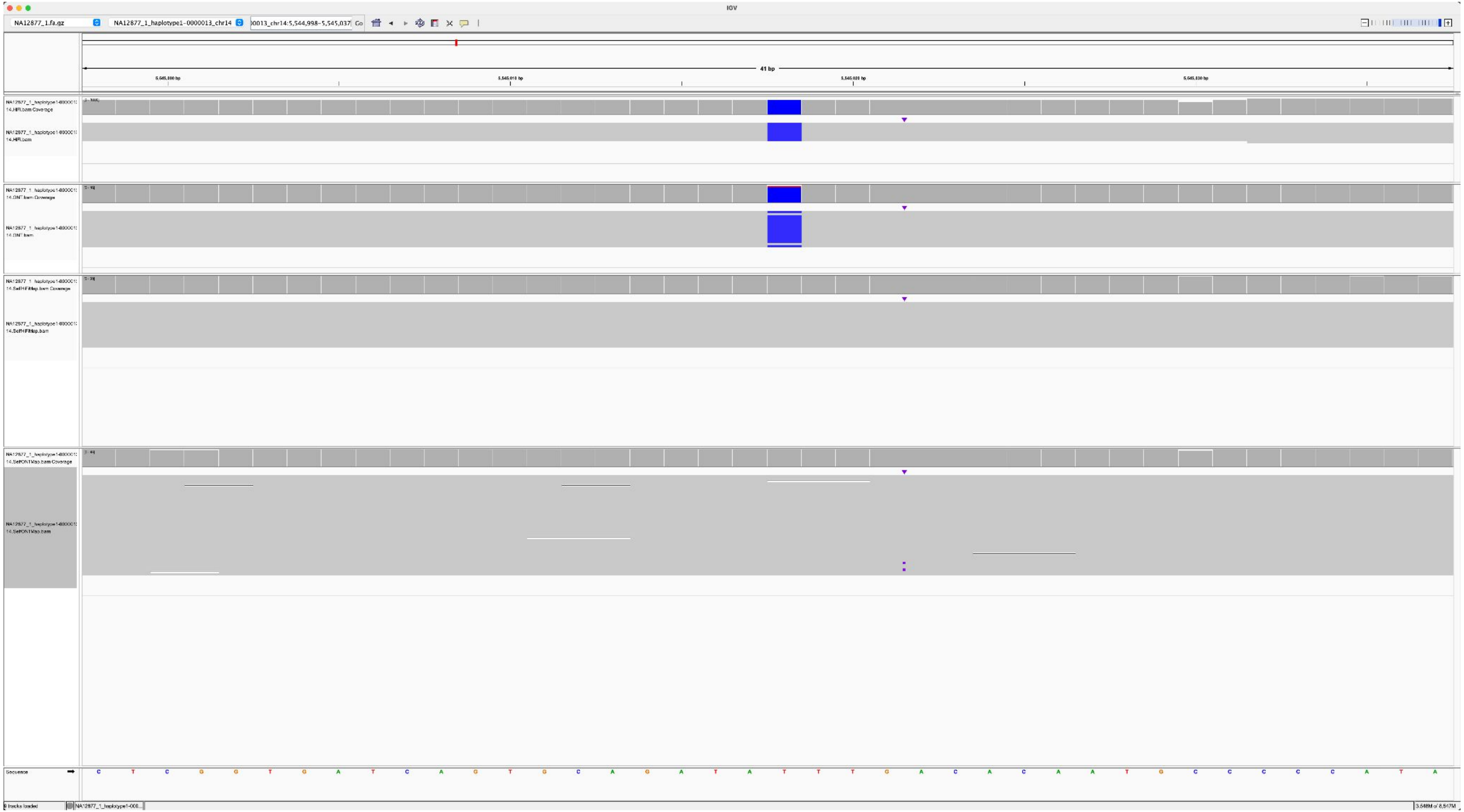

### NA12883_NA12877_1_haplotype1-0000013_chr14_5809047_C_A

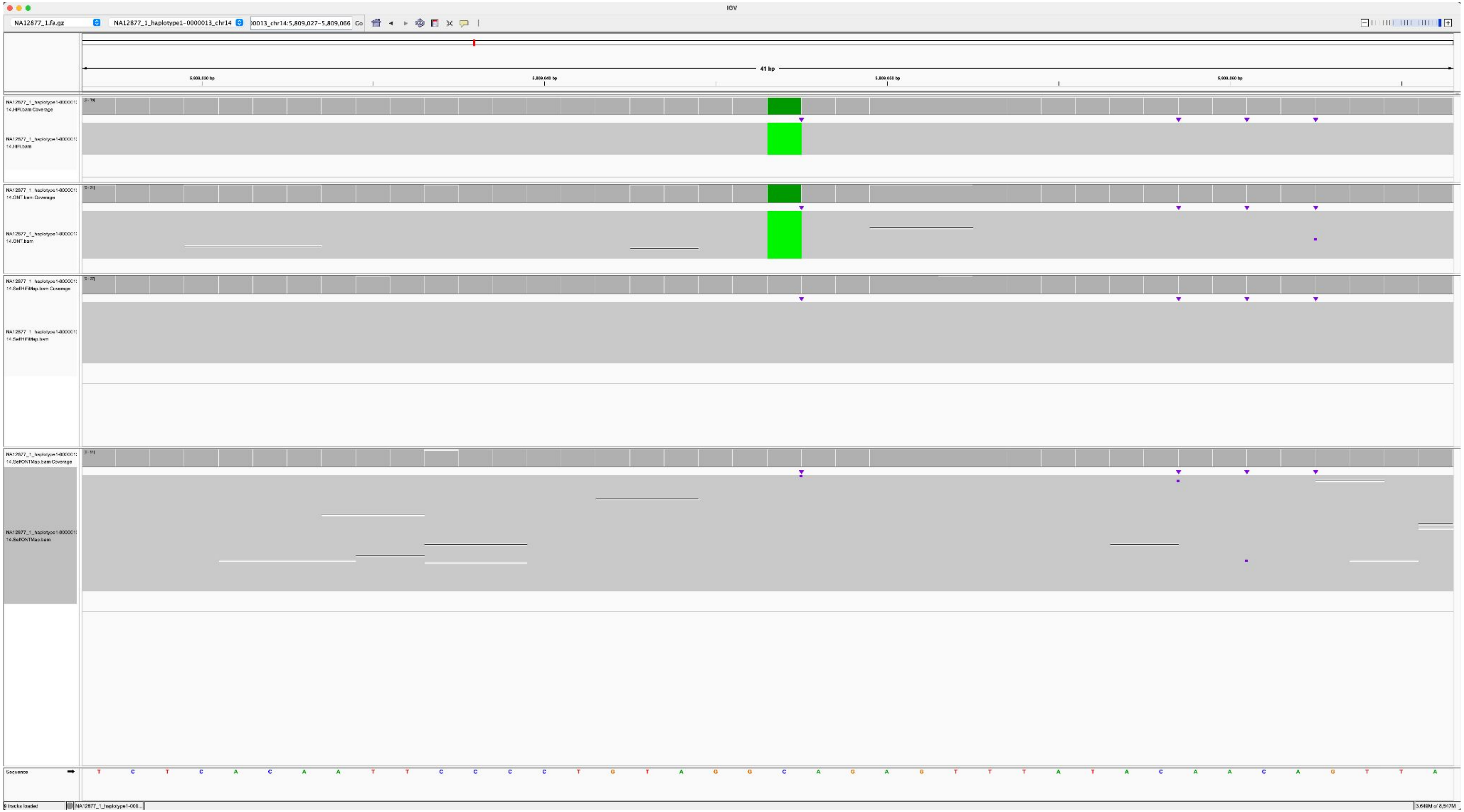

NA12883_NA12877_1_haplotype1-0000013_chr14_555107_T_C

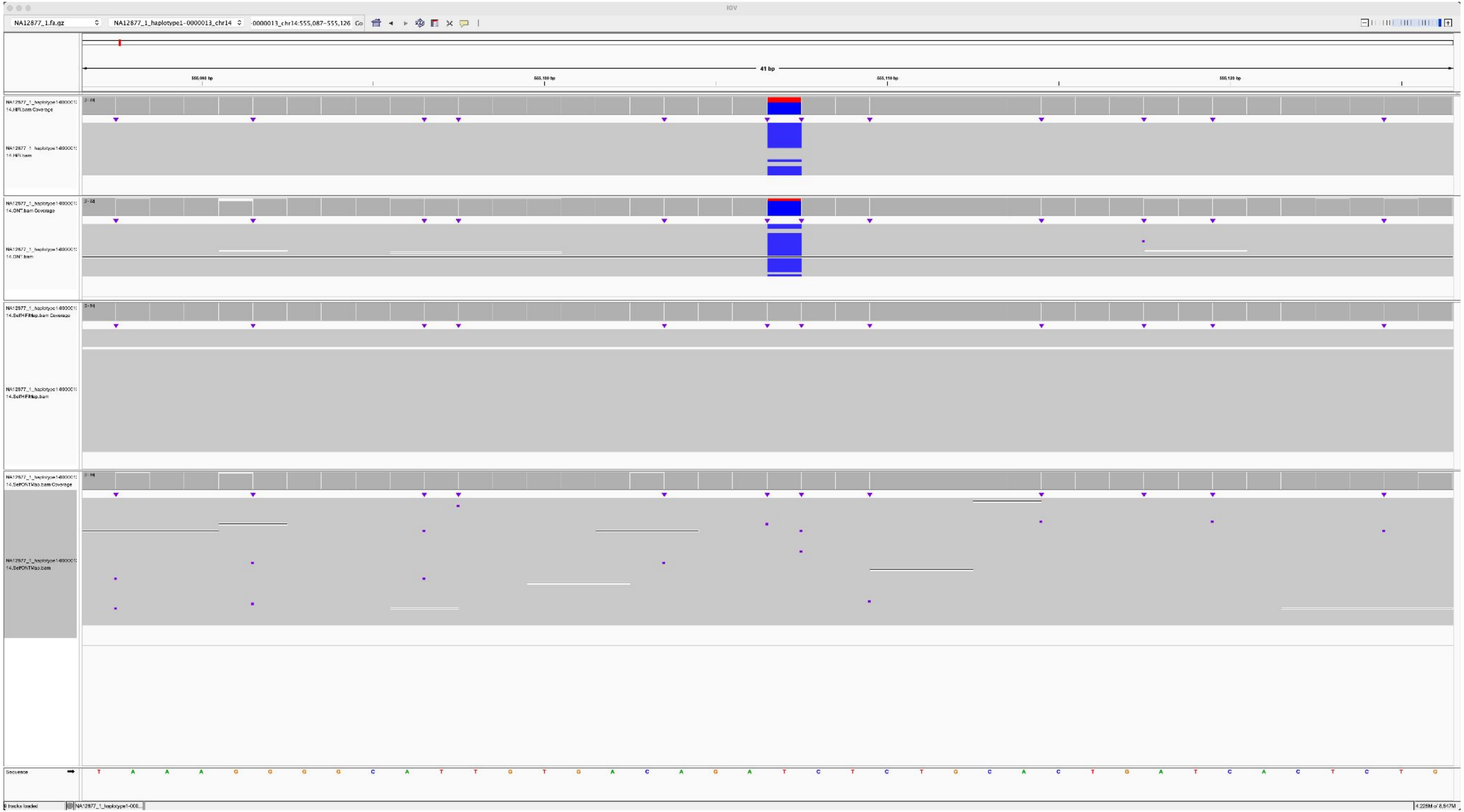

### NA12884_NA12877_1_haplotype1-0000013_chr14_12333960_A_G

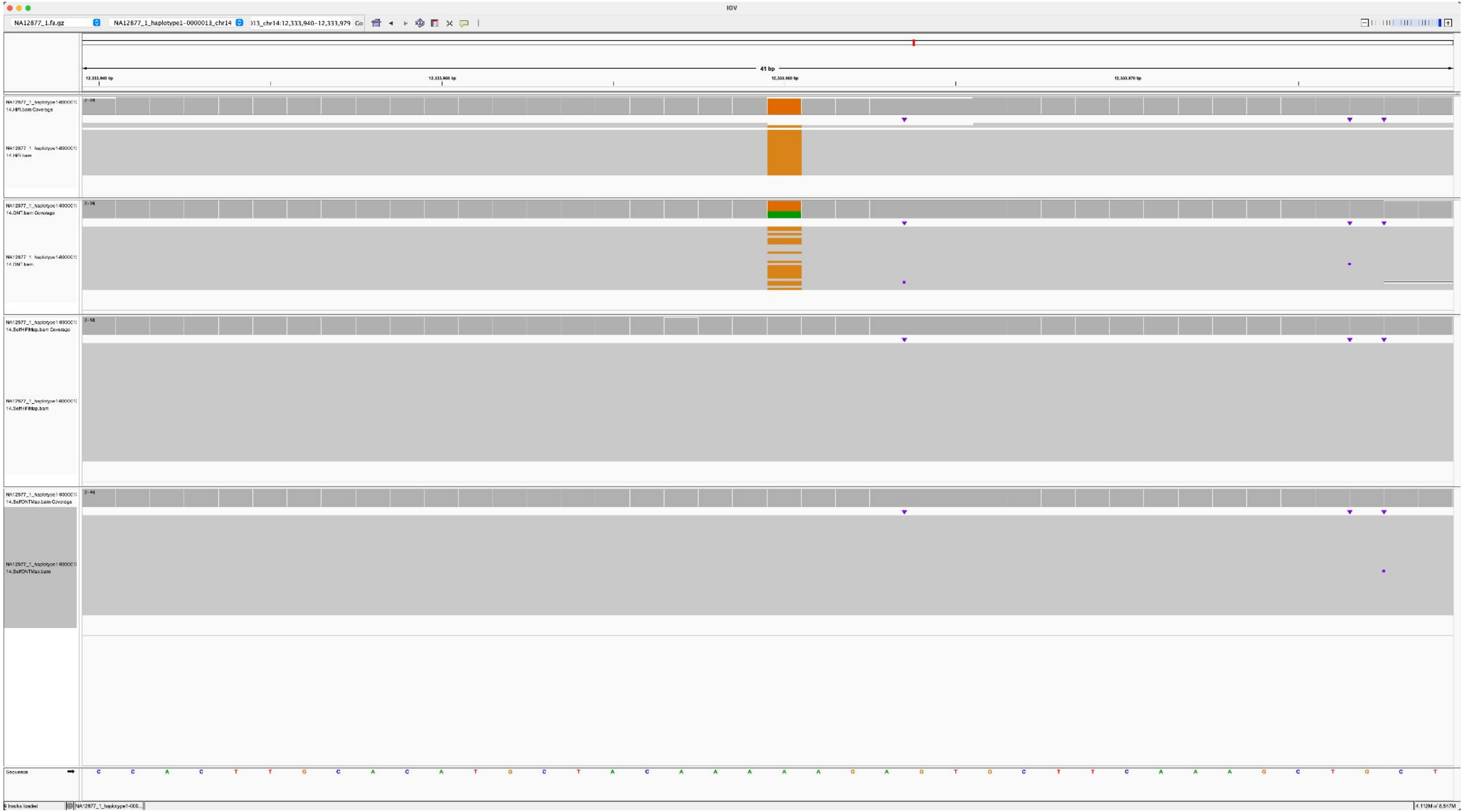

### NA12879_NA12877_1_haplotype1-0000014_chr15_5017549_A_T

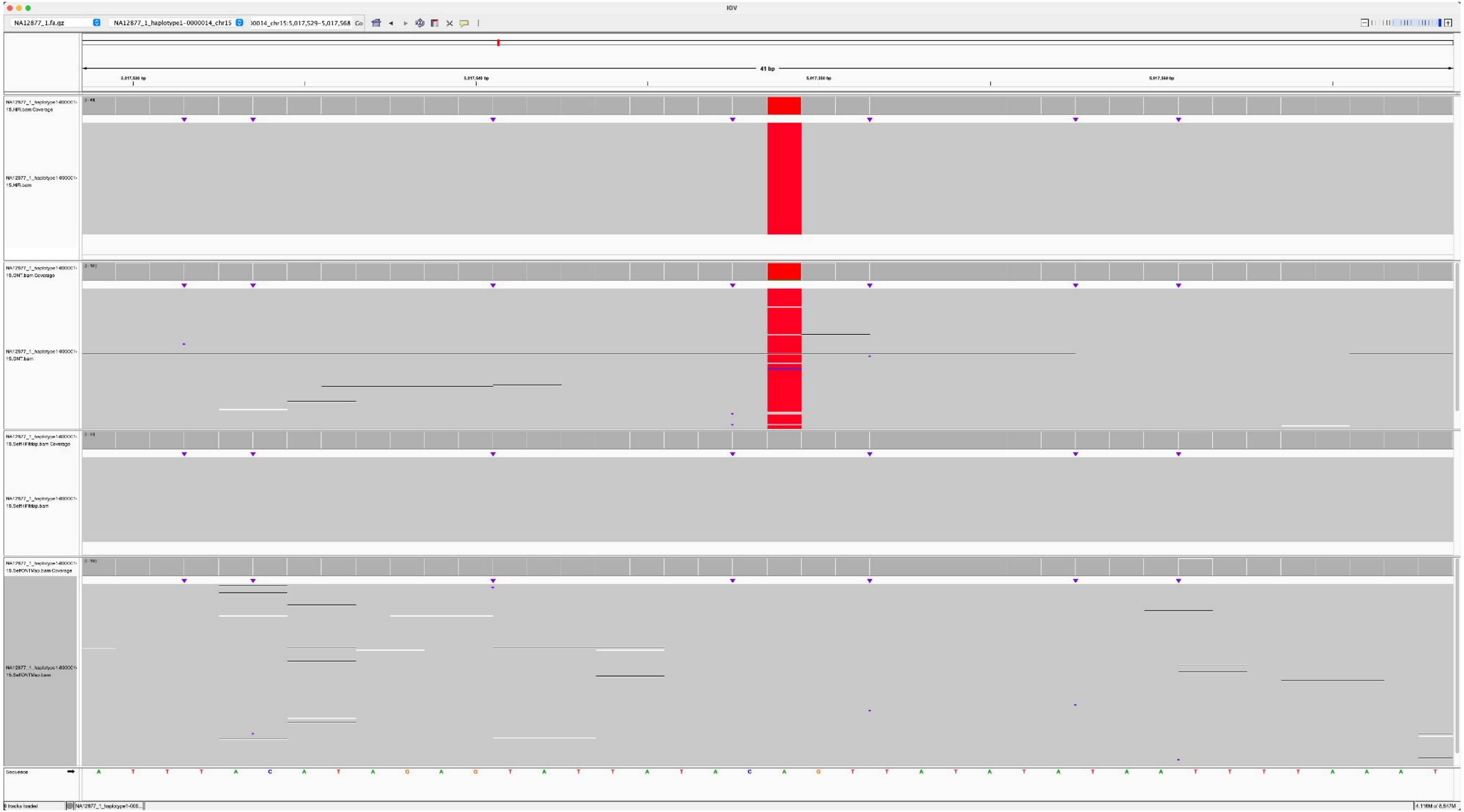

### NA12879_NA12877_1_haplotype1-0000014_chr15_6829060_T_G

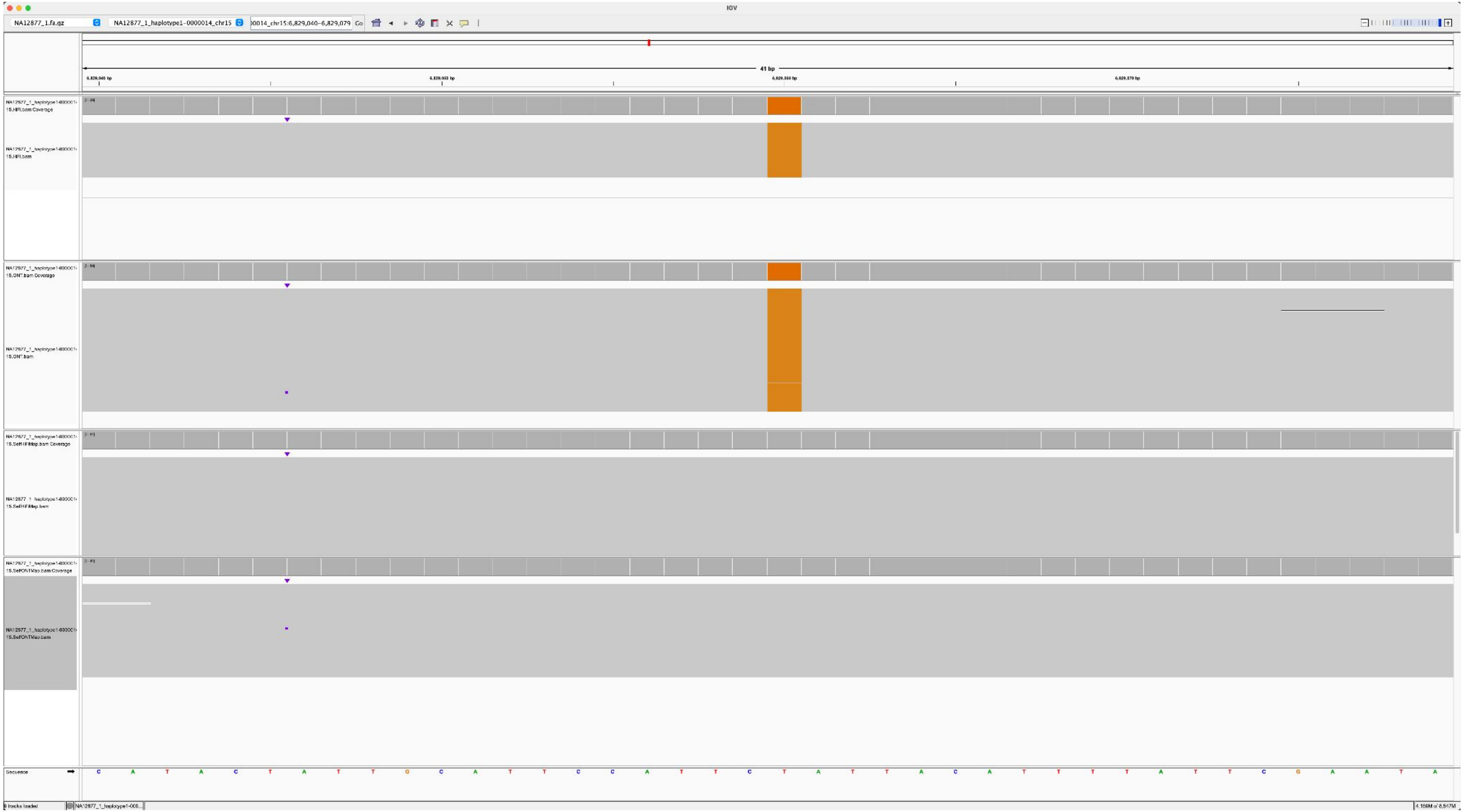

### NA12879_NA12877_1_haplotype1-0000014_chr15_10124692_A_C

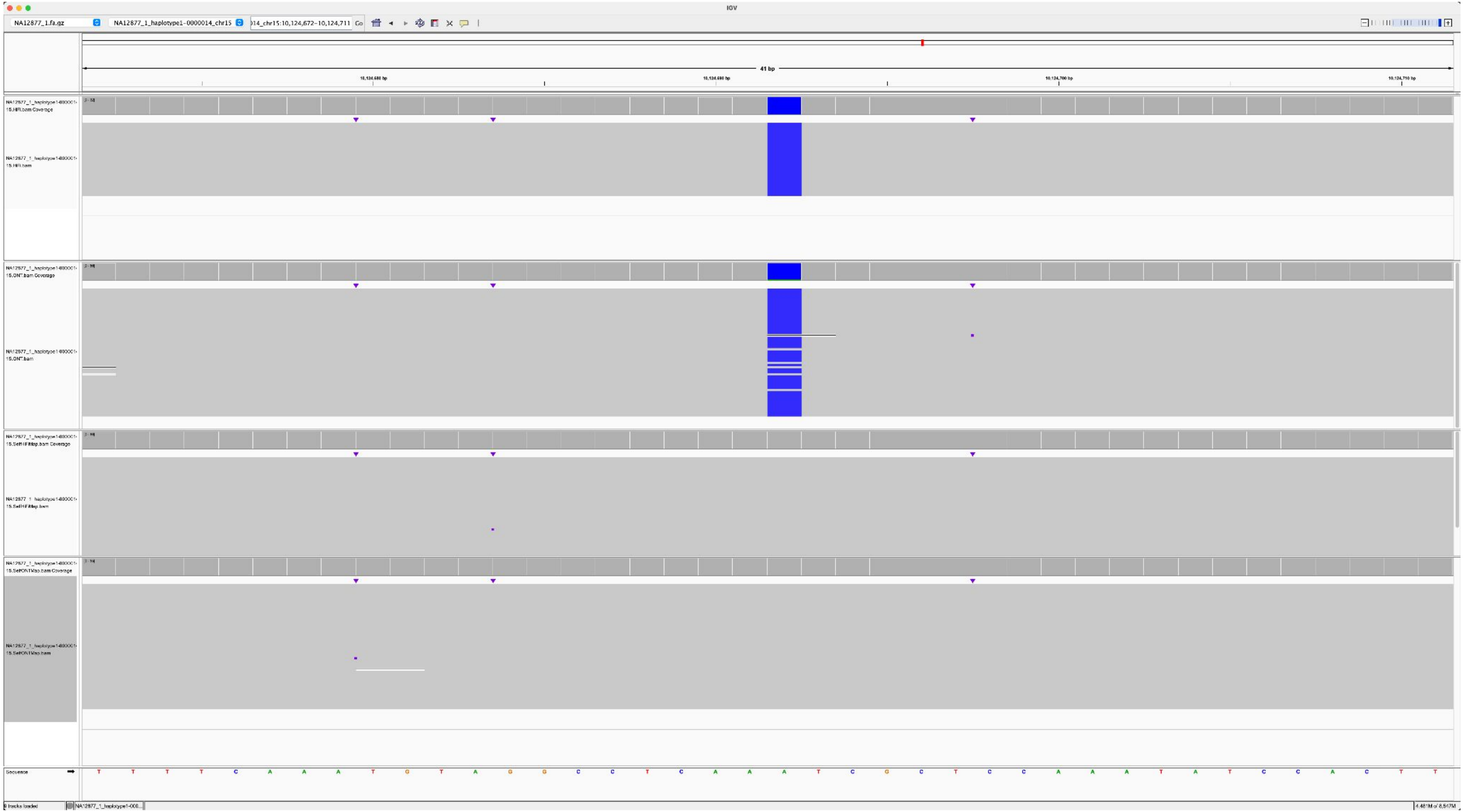

### NA12882_NA12877_1_haplotype1-0000014_chr15_5929145_C_G

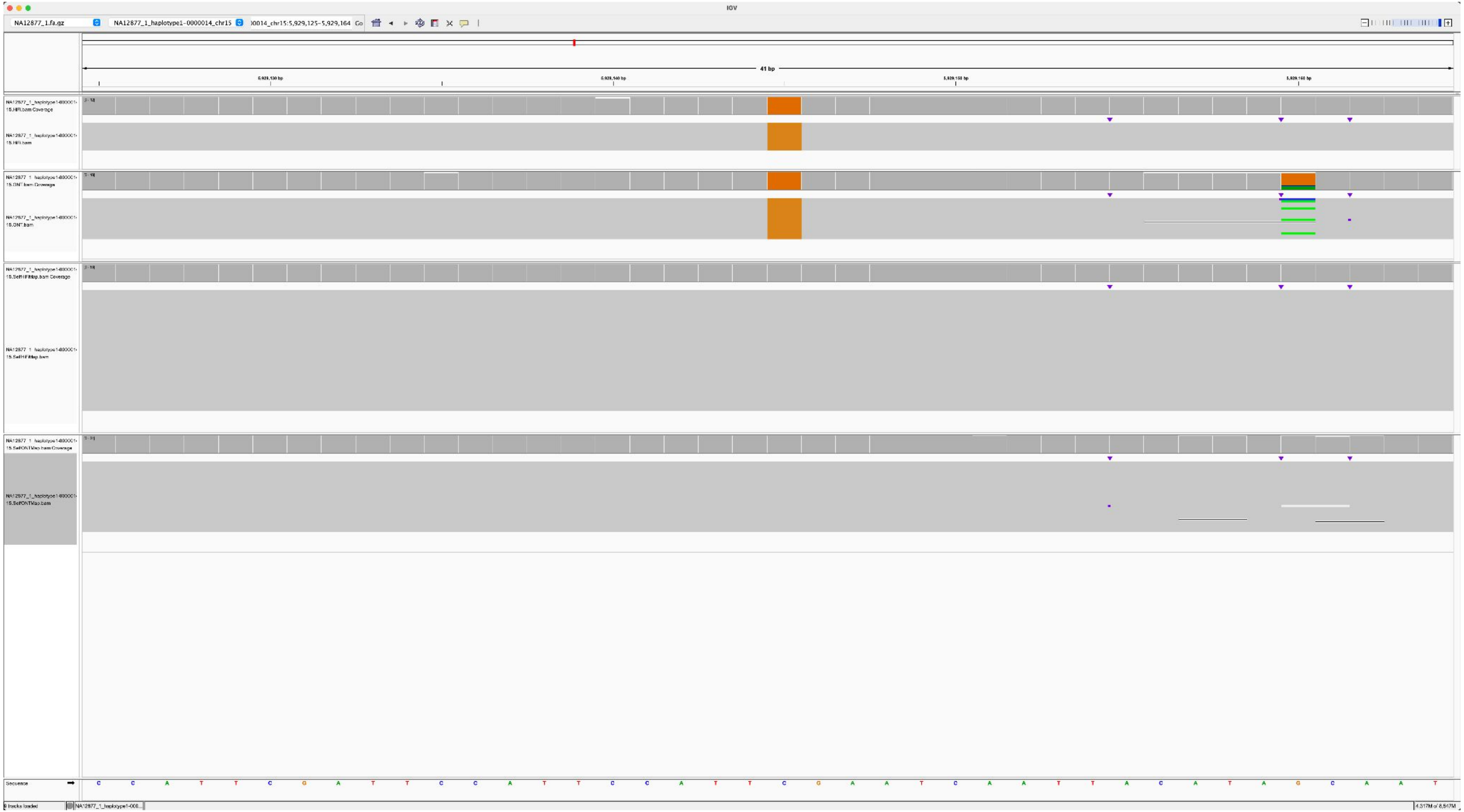

### NA12885_NA12877_1_haplotype1-0000014_chr15_5314406_A_C

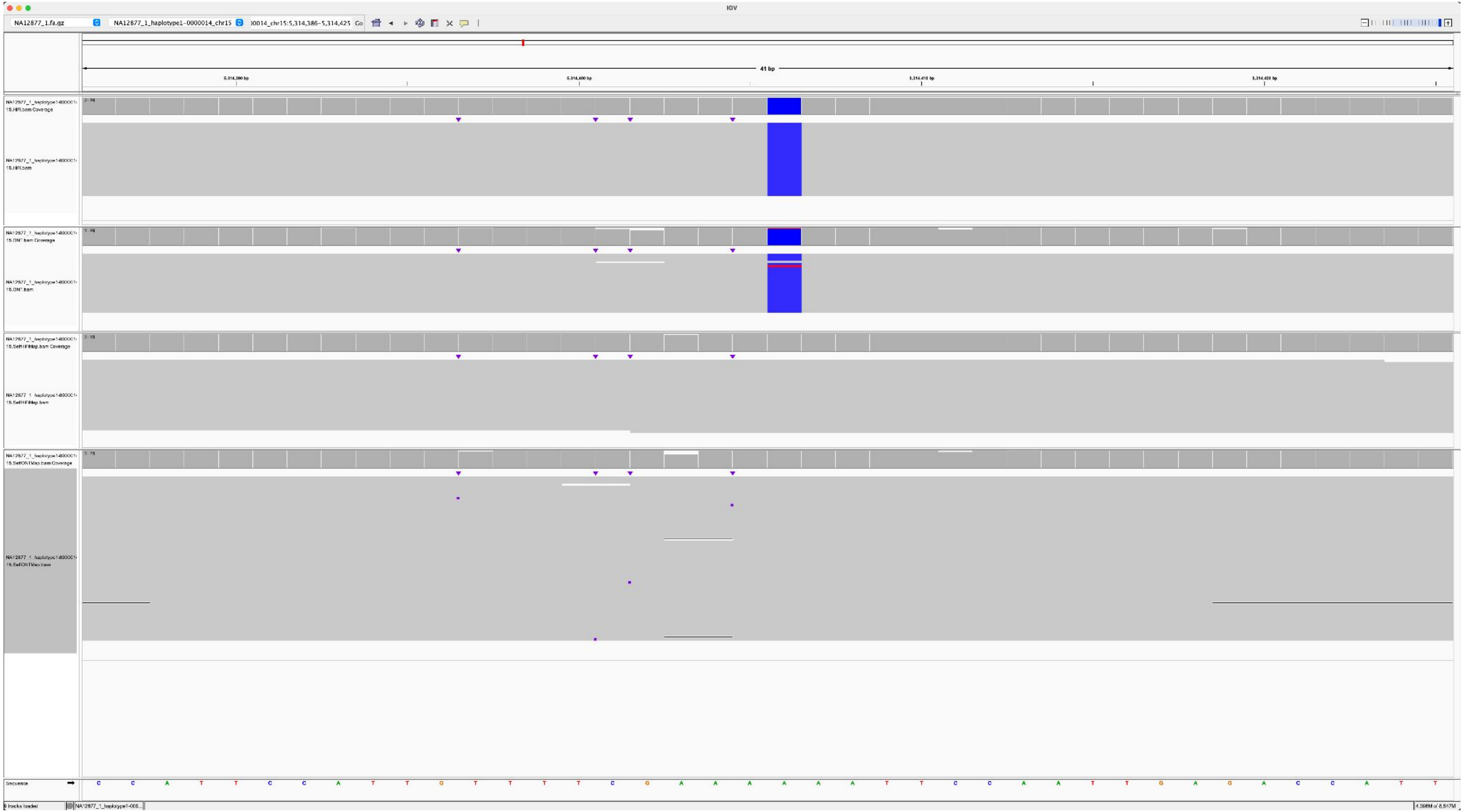

### NA12886_NA12877_1_haplotype1-0000014_chr15_6512838_A_C

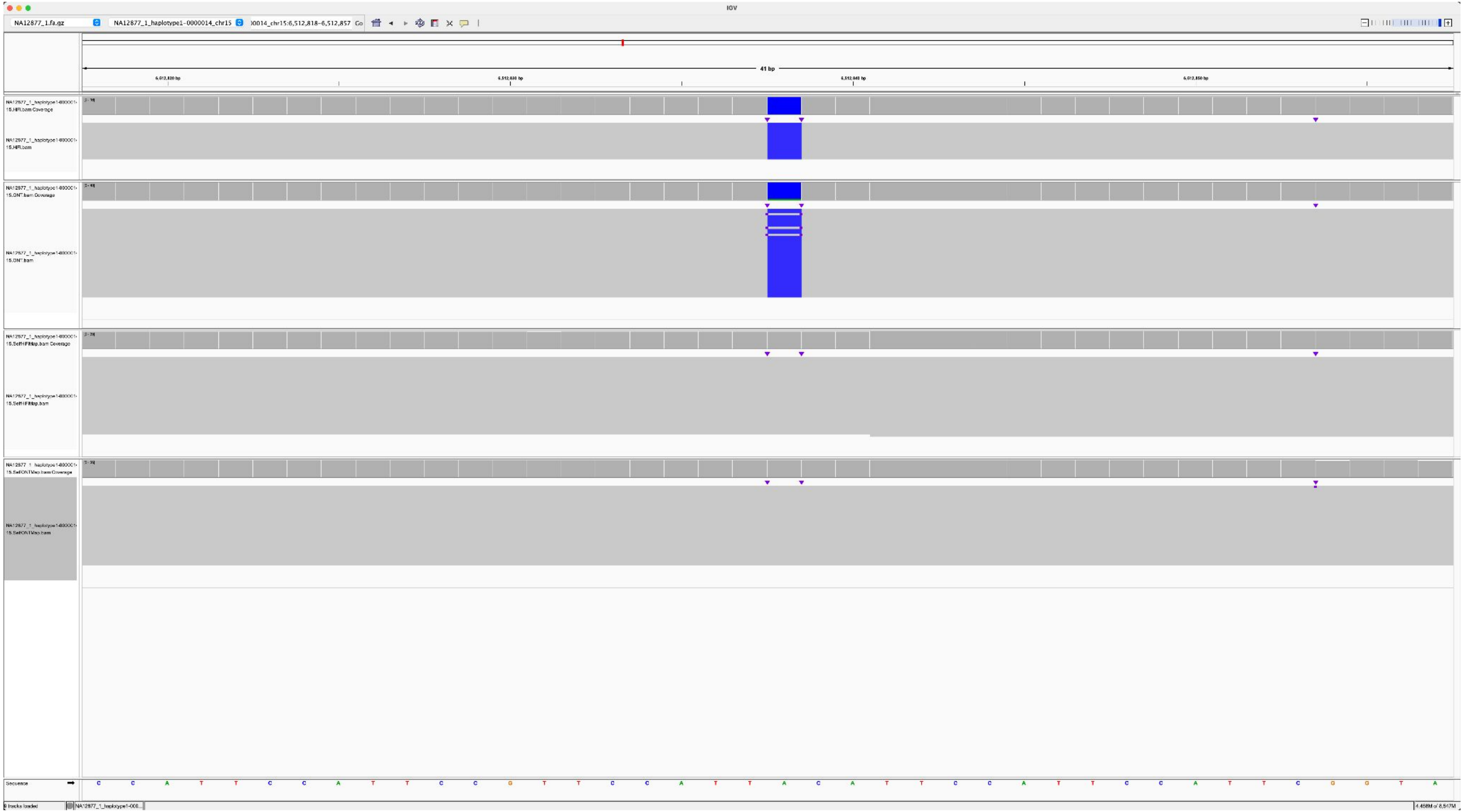

### NA12886_NA12877_1_haplotype1-0000014_chr15_1028029_G_C

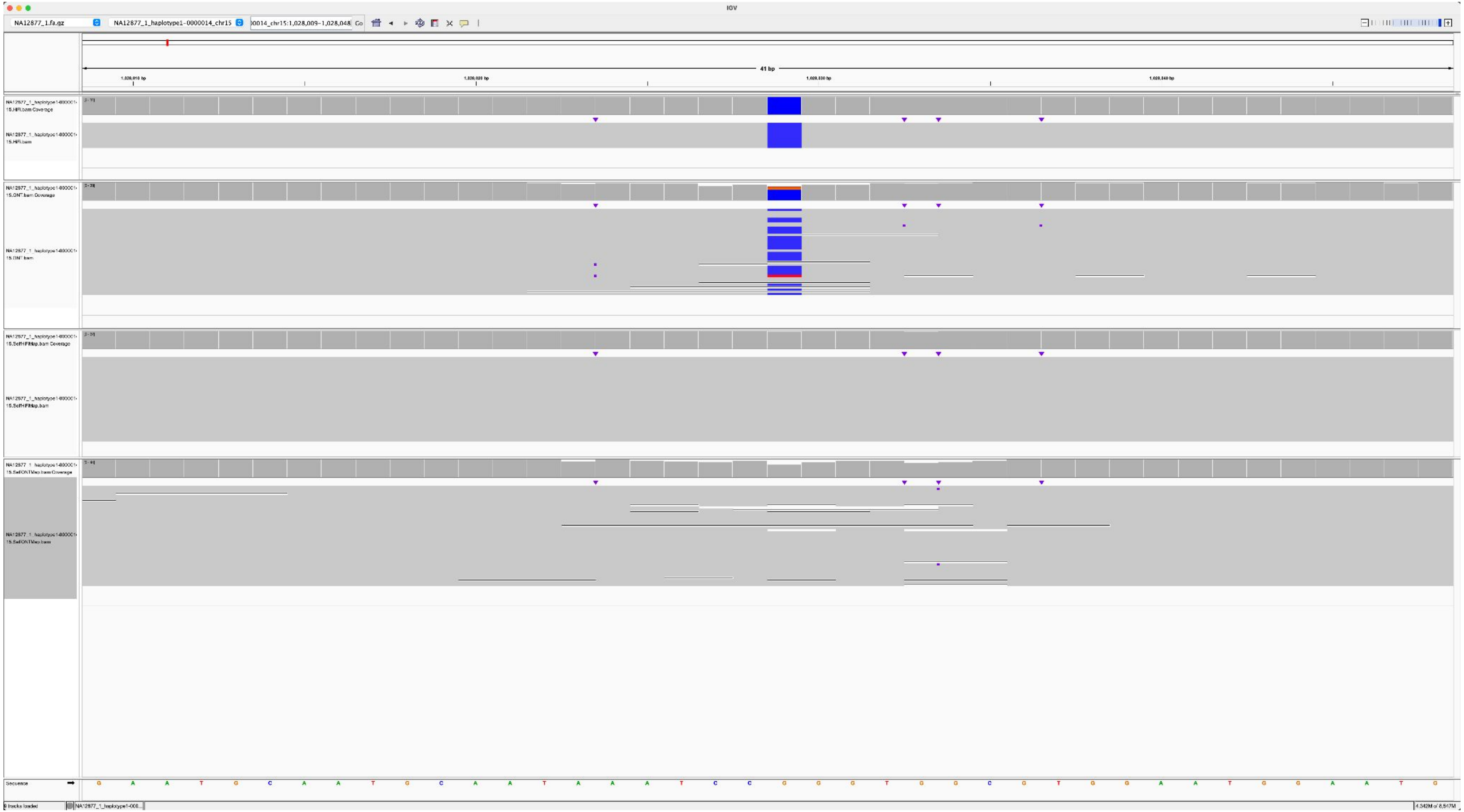

NA12886_NA12877_1_haplotype1-0000014_chr15_2316825_C_A

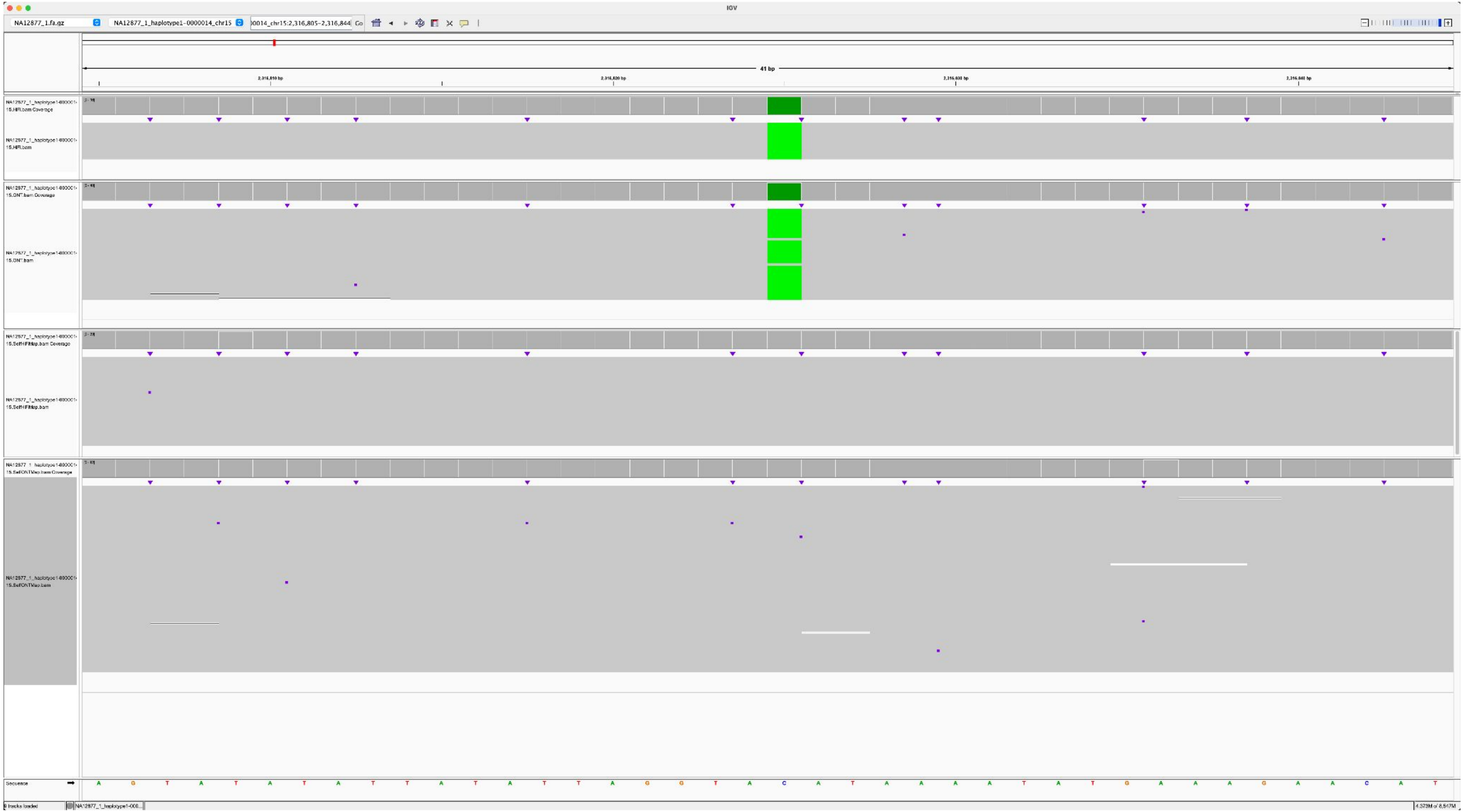

### NA12887_NA12877_1_haplotype1-0000014_chr15_7039843_T_G

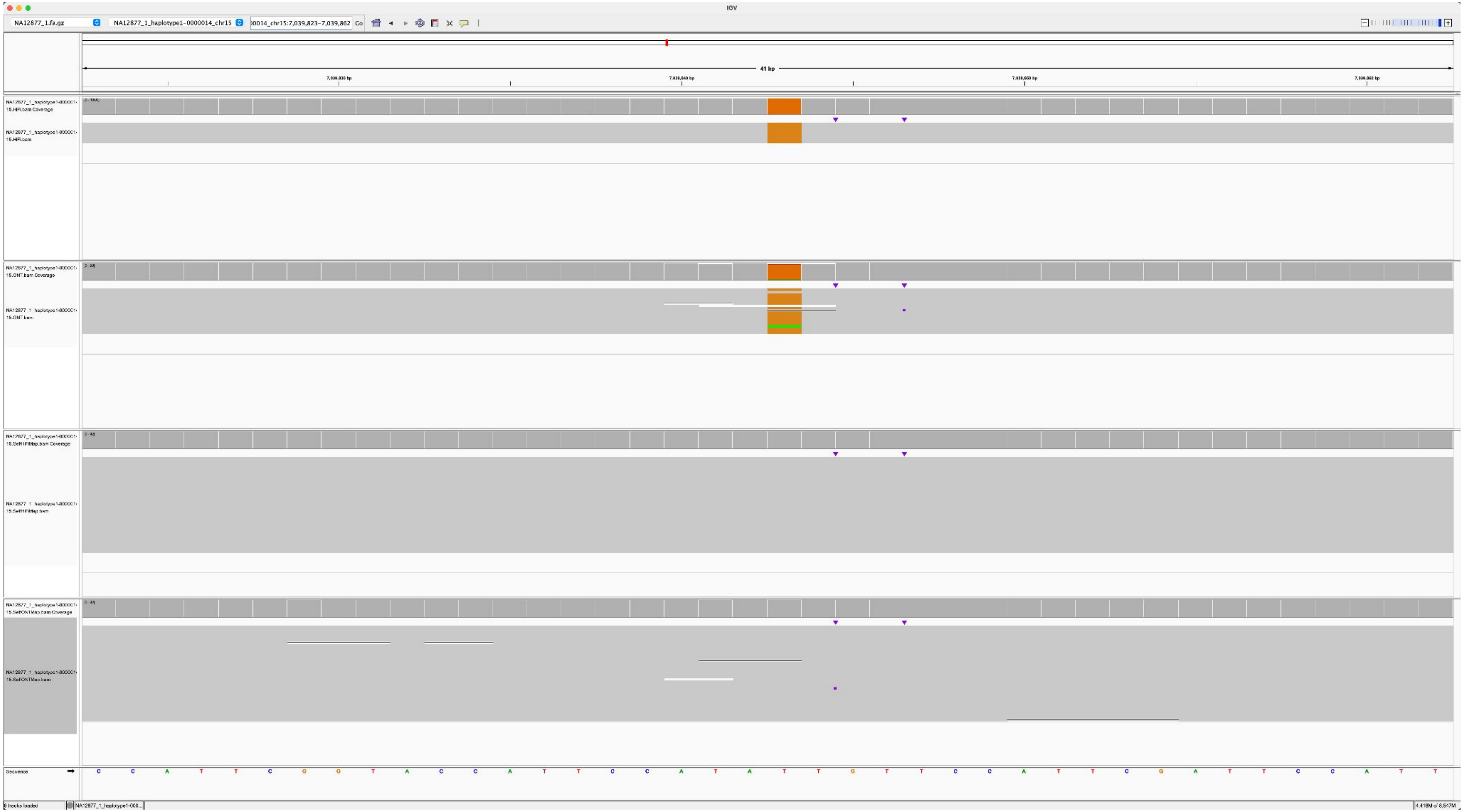

### NA12887_NA12877_1_haplotype1-0000014_chr15_7782498_G_A

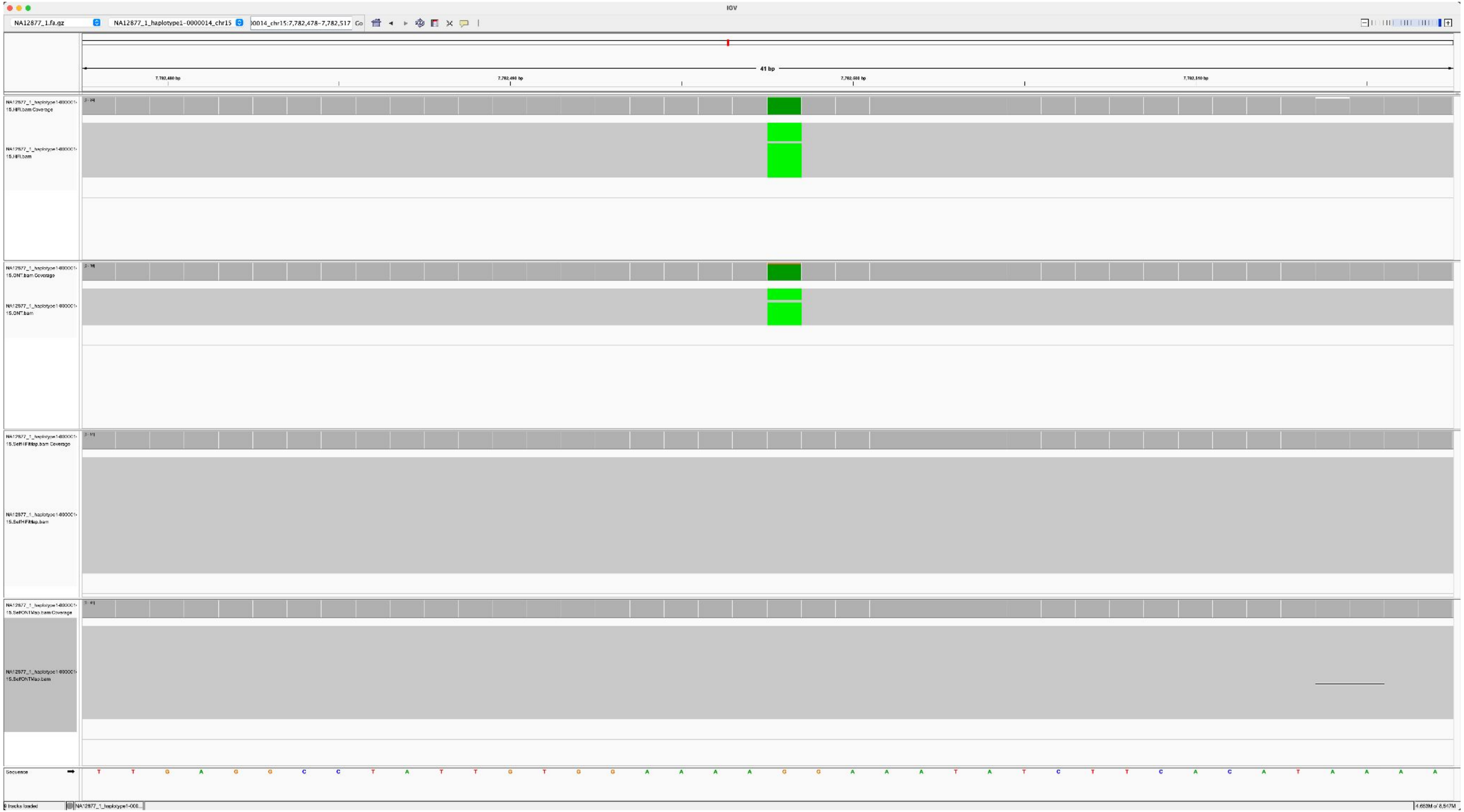

NA12887_NA12877_1_haplotype1-0000014_chr15_5124976_A_T

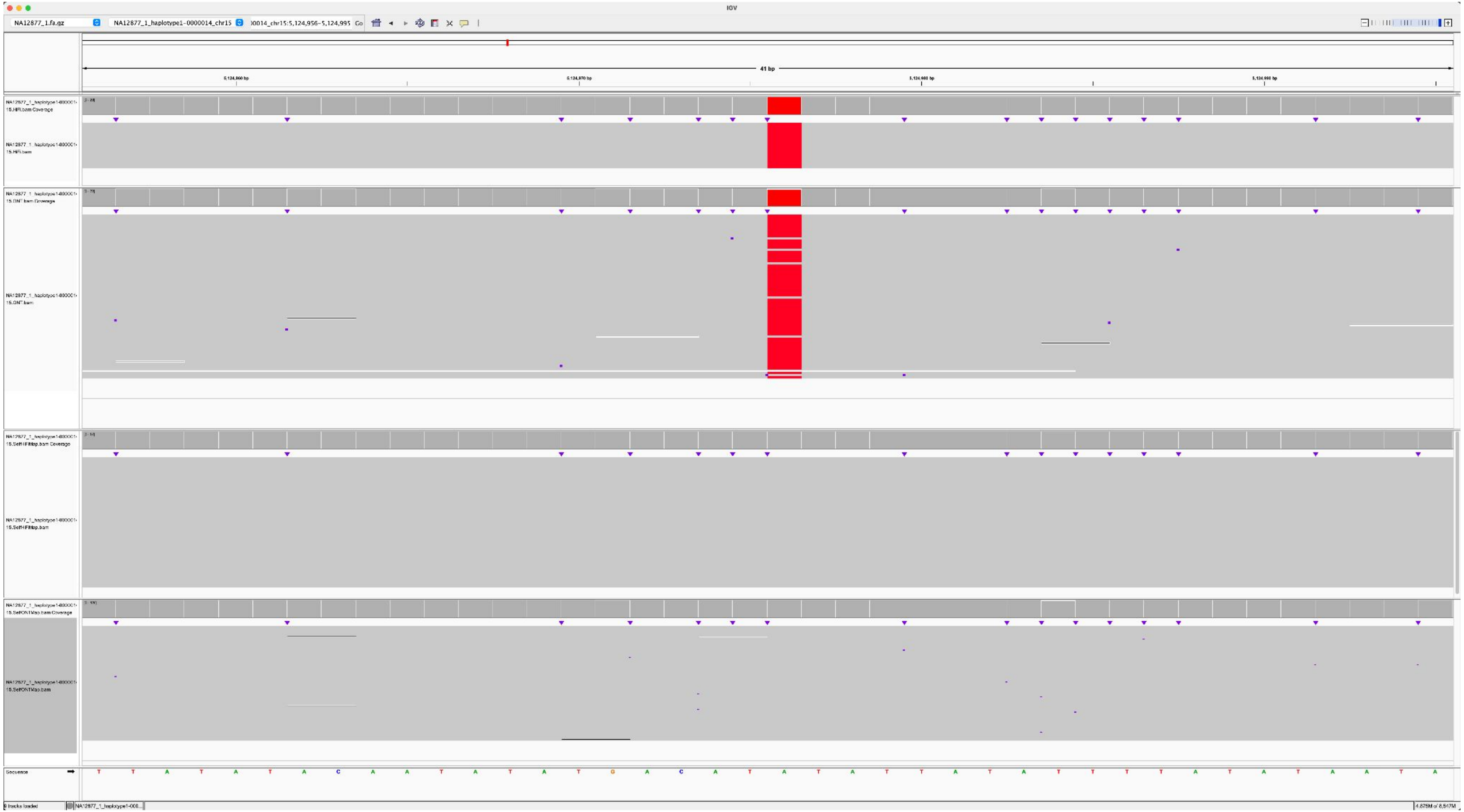

### NA12885_NA12877_1_haplotype1-0000023_chr21_5867616_C_A

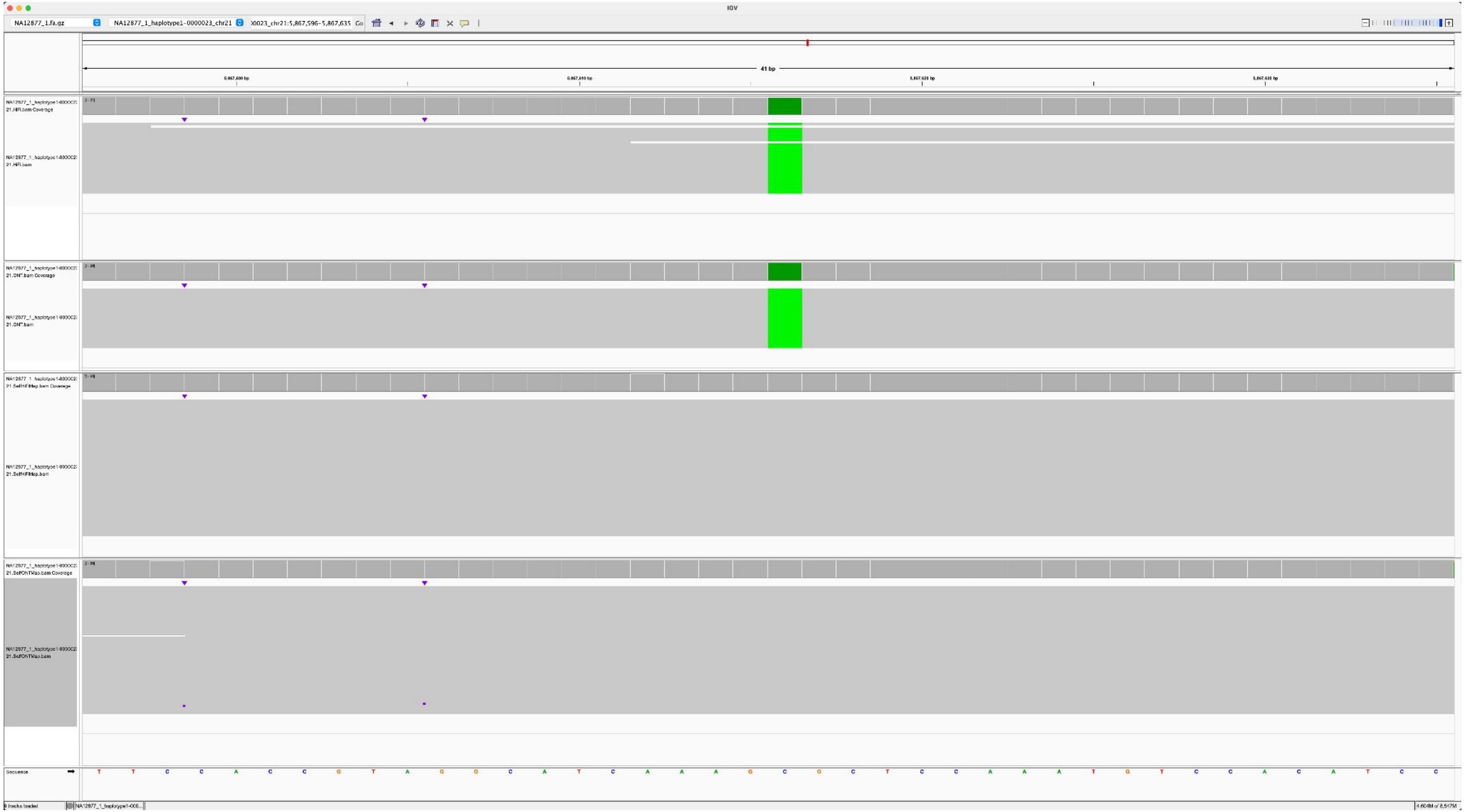

### NA12886_NA12877_1_haplotype1-0000023_chr21_3423589_C_G

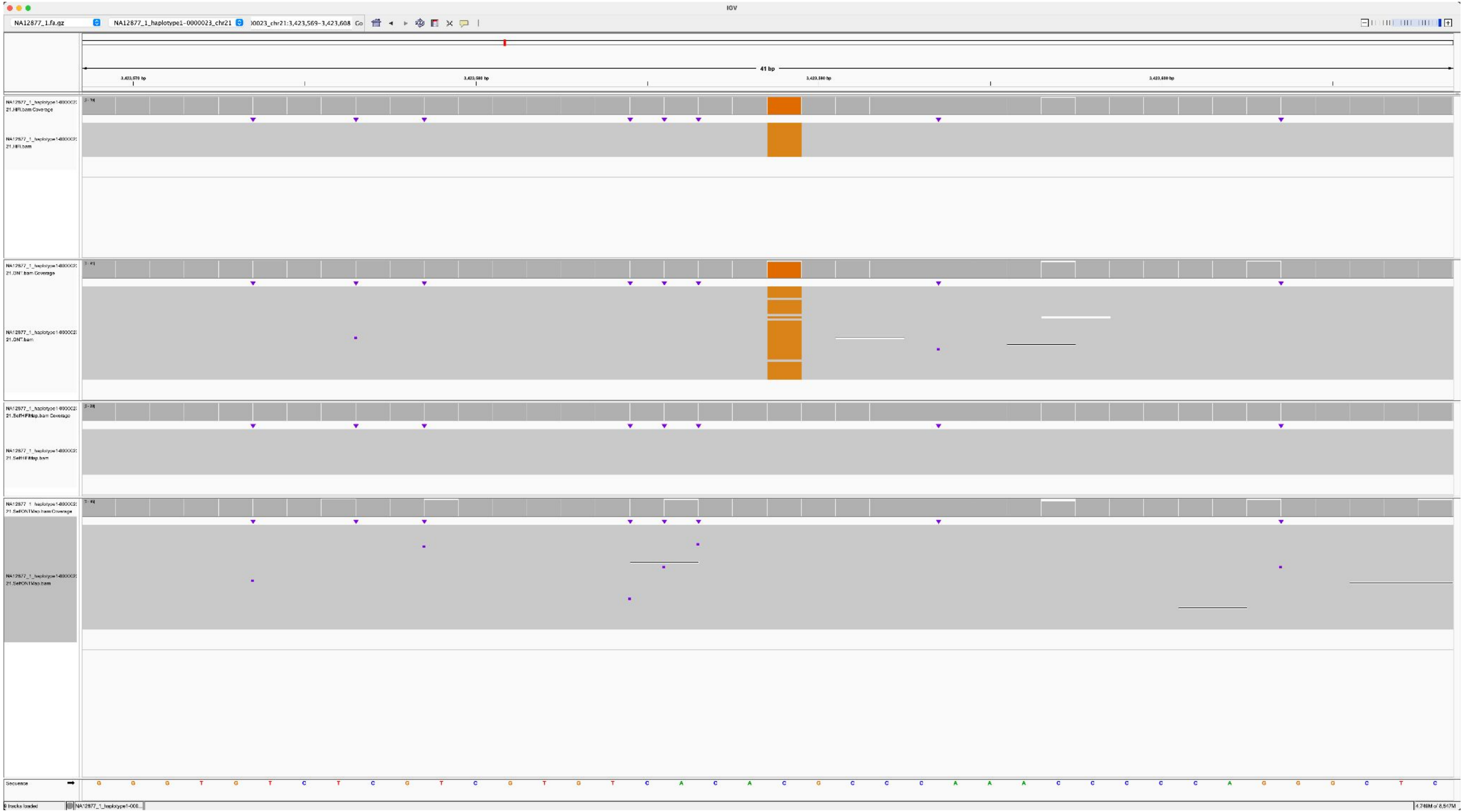

### NA12879_NA12877_1_haplotype1-0000020_chr22_7604871_G_T

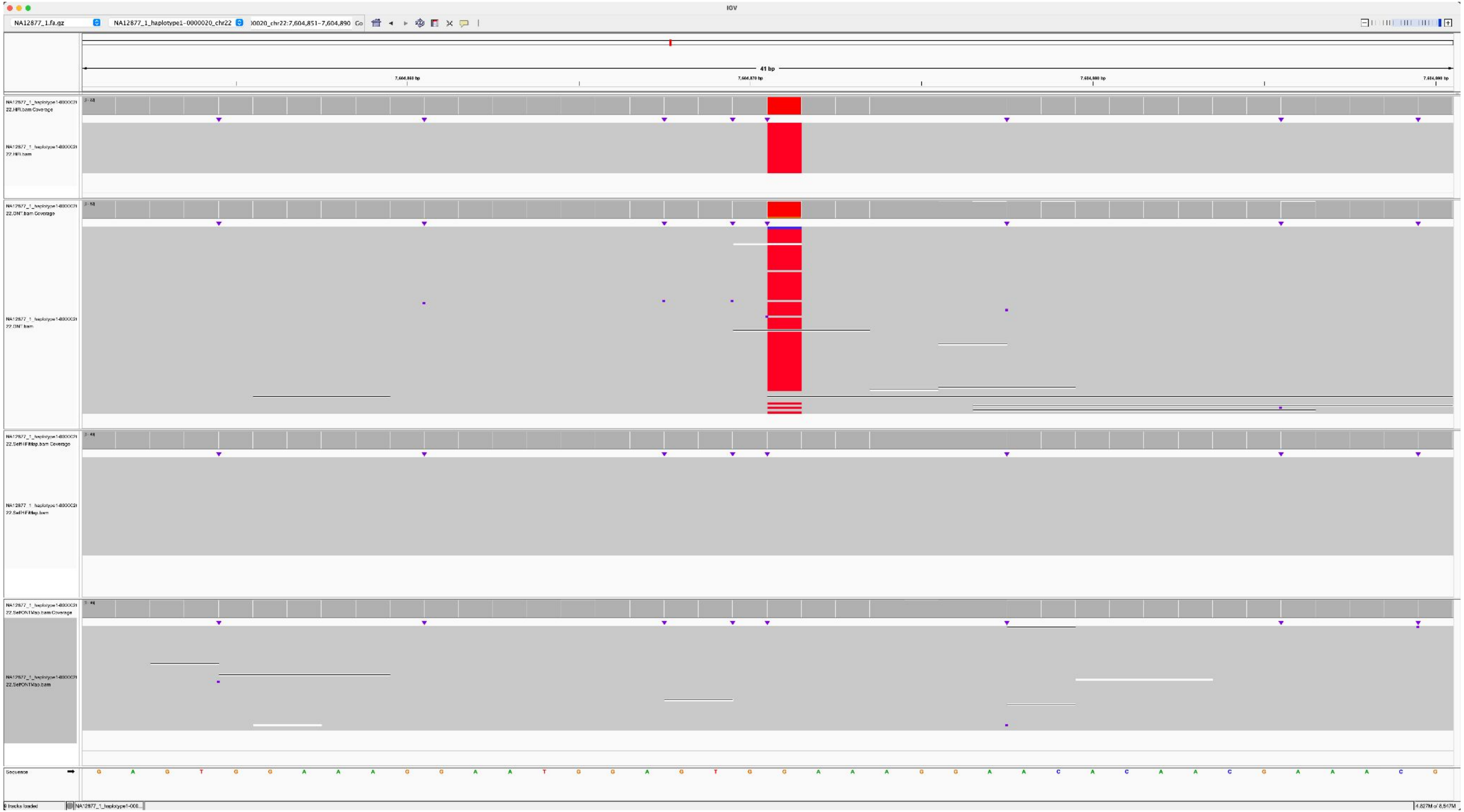

### NA12879_NA12877_1_haplotype1-0000020_chr22_1427257_T_G

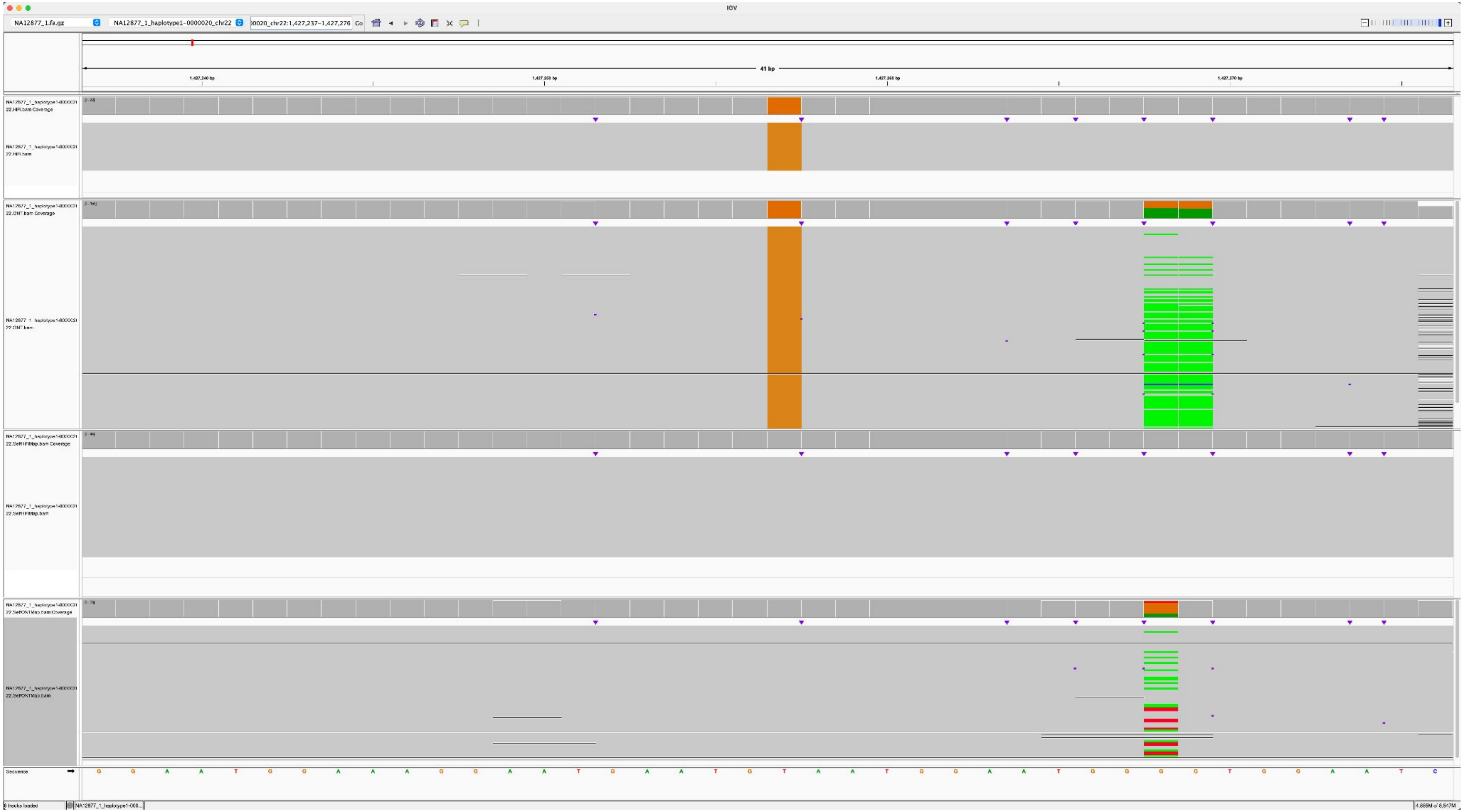

### NA12882_NA12877_1_haplotype1-0000020_chr22_10320157_G_C

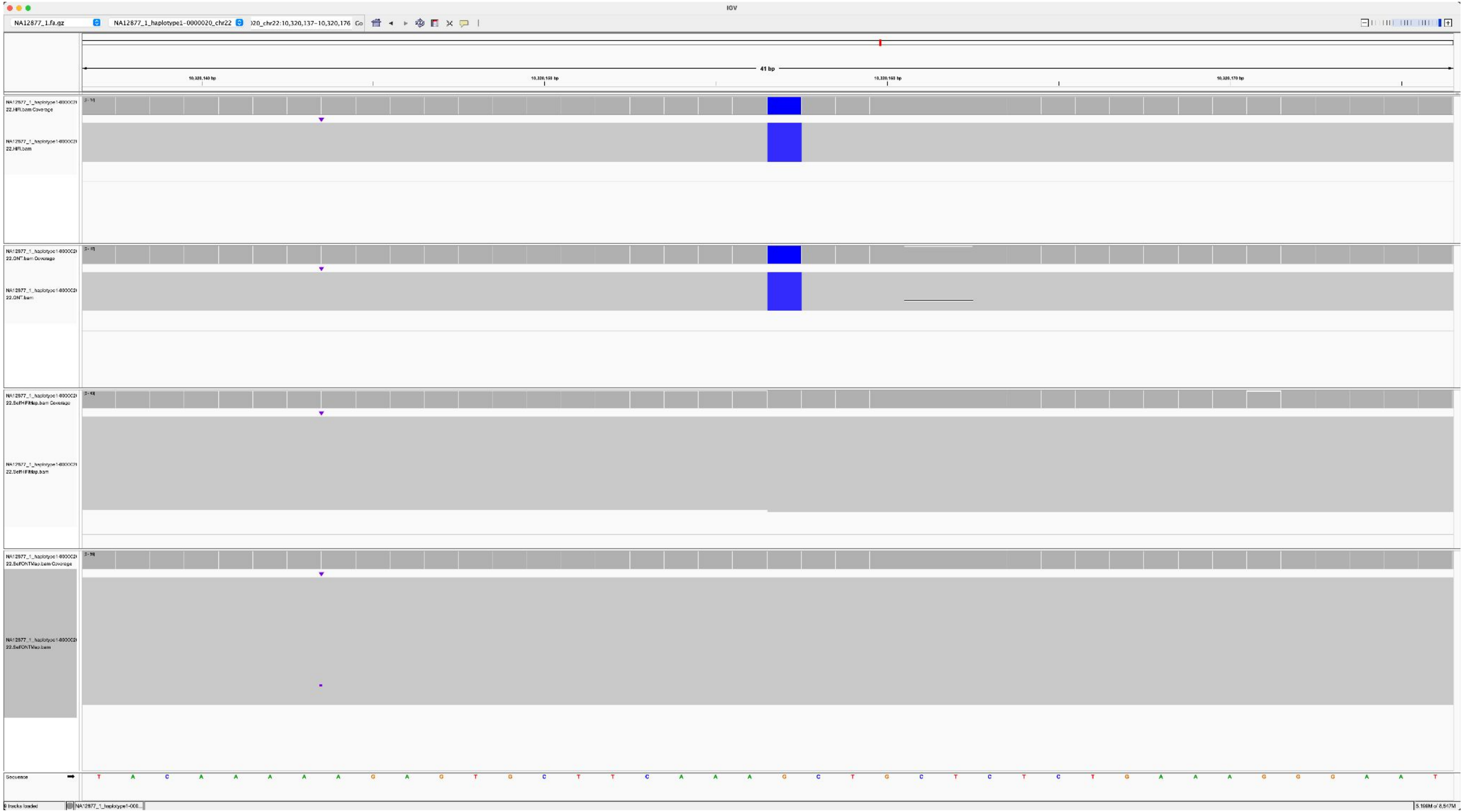

### NA12883_NA12877_1_haplotype1-0000020_chr22_6909898_G_T

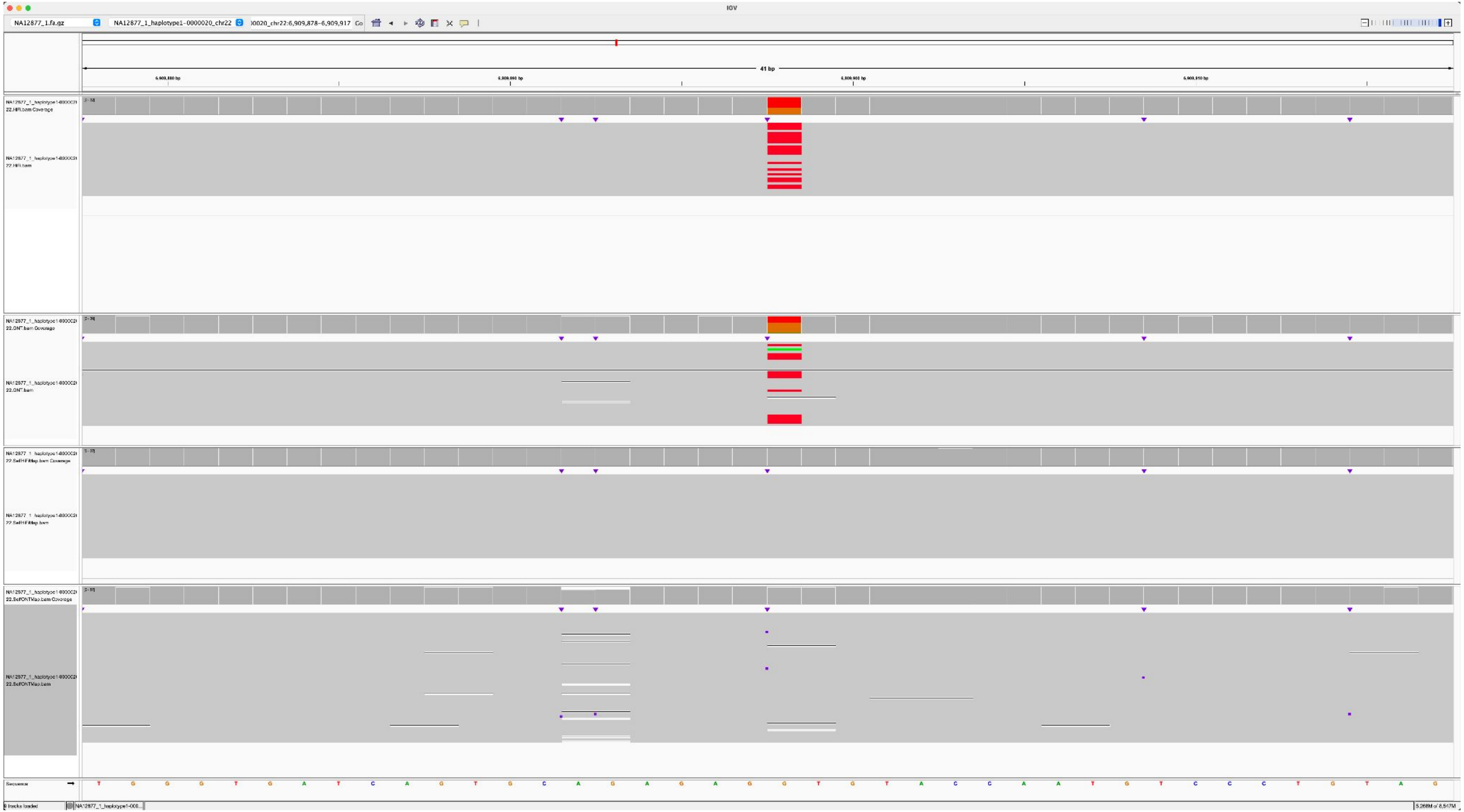

NA12883_NA12877_1_haplotype1-0000020_chr22_6919775_G_T

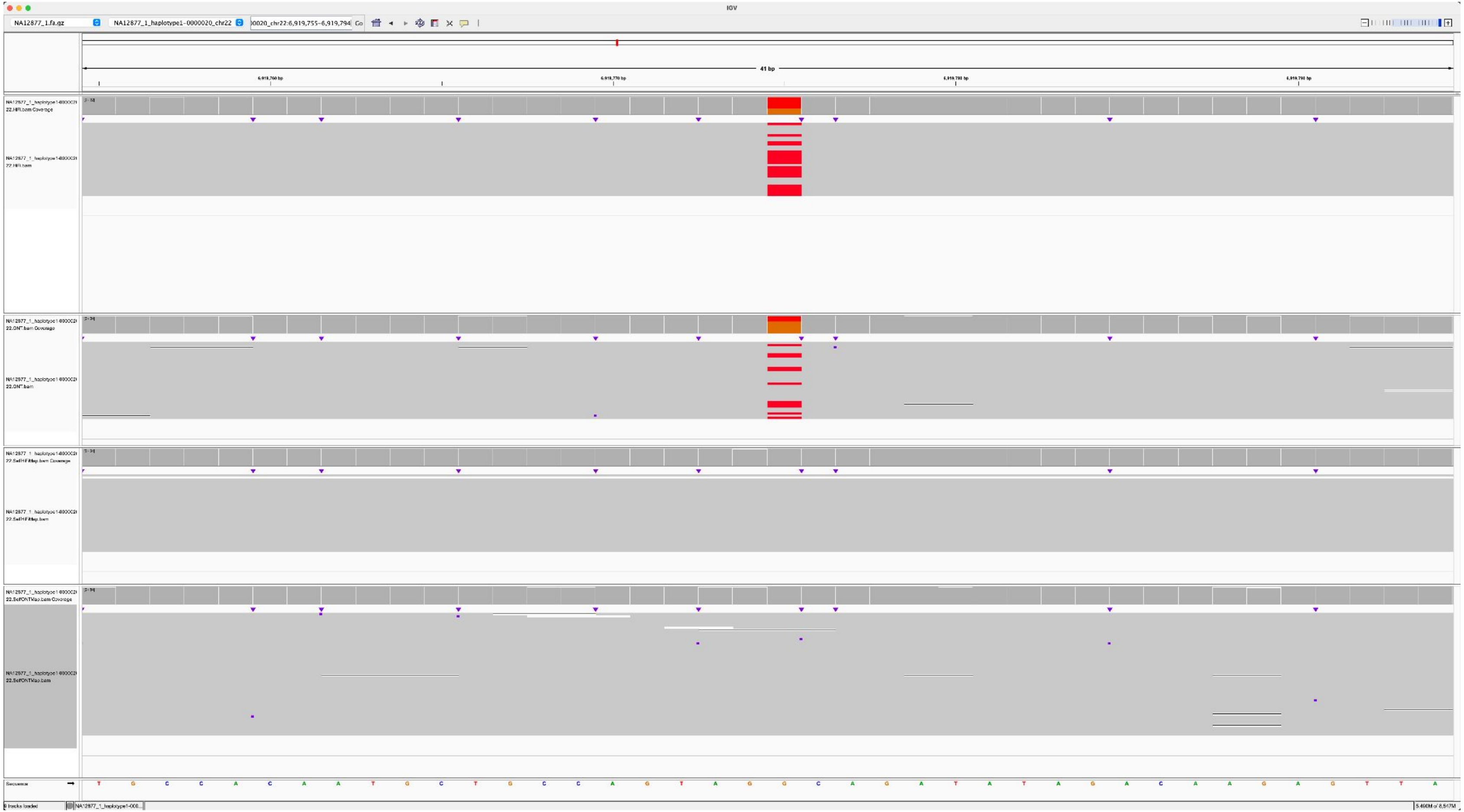

NA12883_NA12877_1_haplotype1-0000020_chr22_6929157_T_C

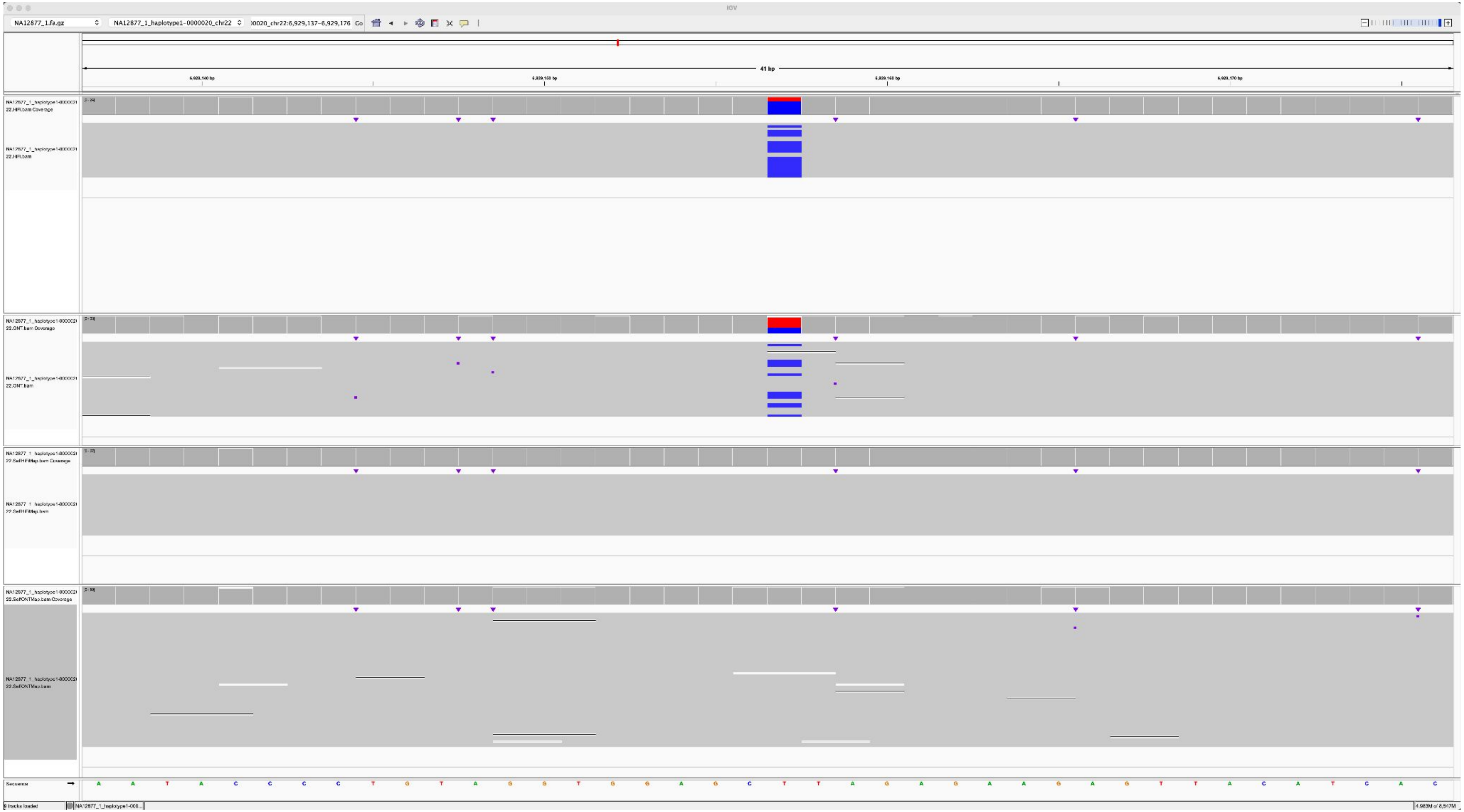

### NA12883_NA12877_1_haplotype1-0000020_chr22_5456040_A_G

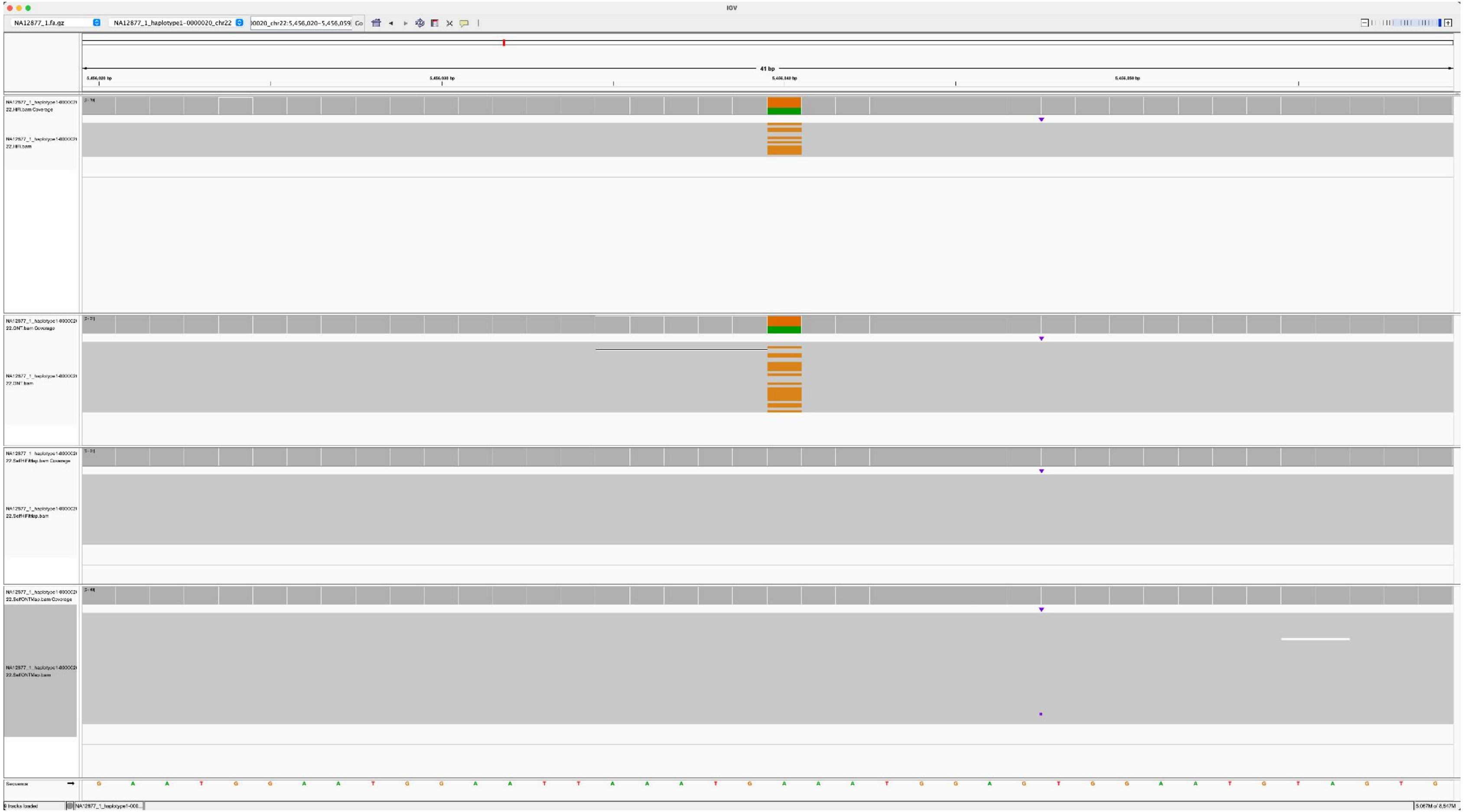

### NA12884_NA12877_1_haplotype1-0000020_chr22_8755692_C_T

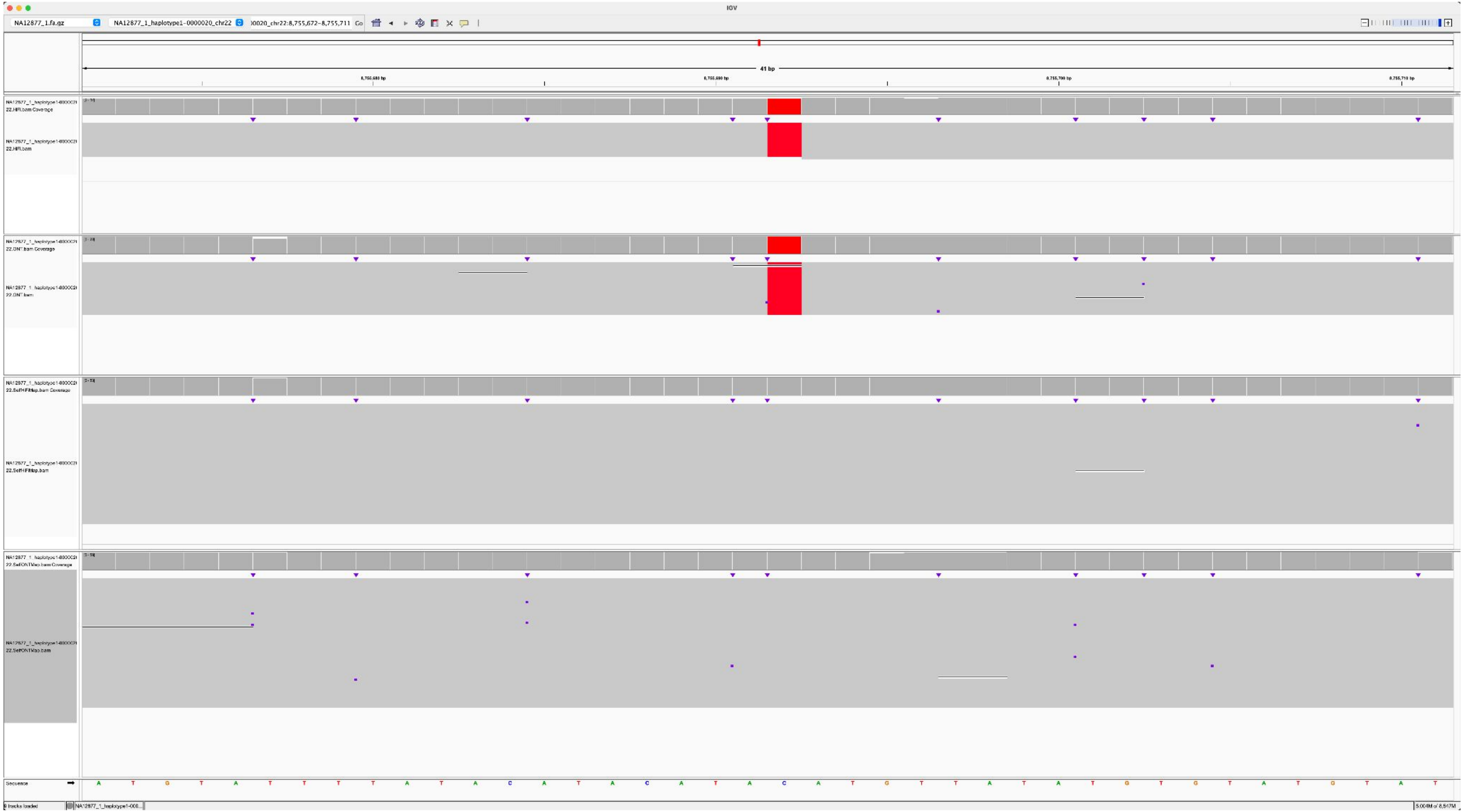

### NA12884_NA12877_1_haplotype1-0000020_chr22_12720441_A_T

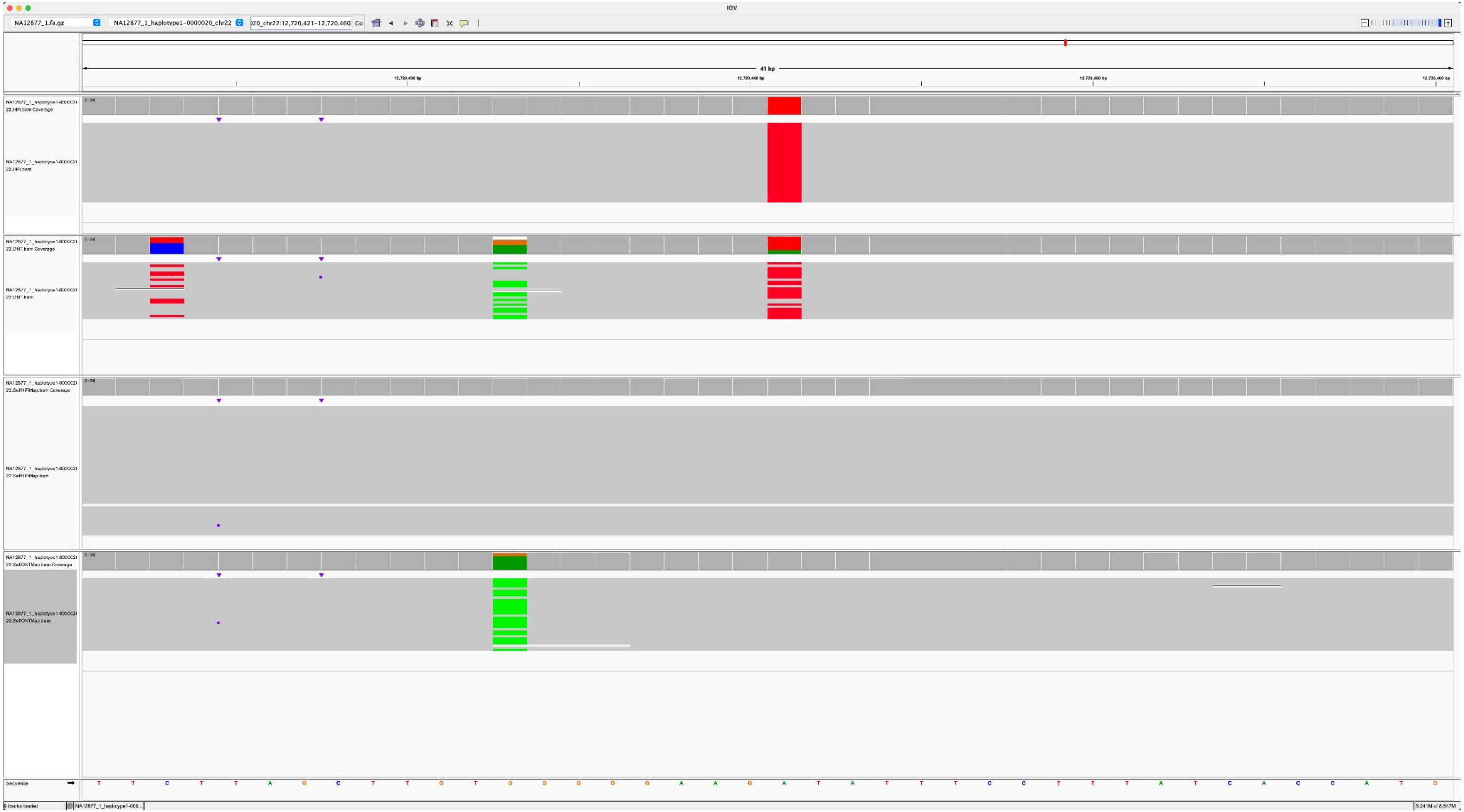

NA12884_NA12877_1_haplotype1-0000020_chr22_1826537_G_A

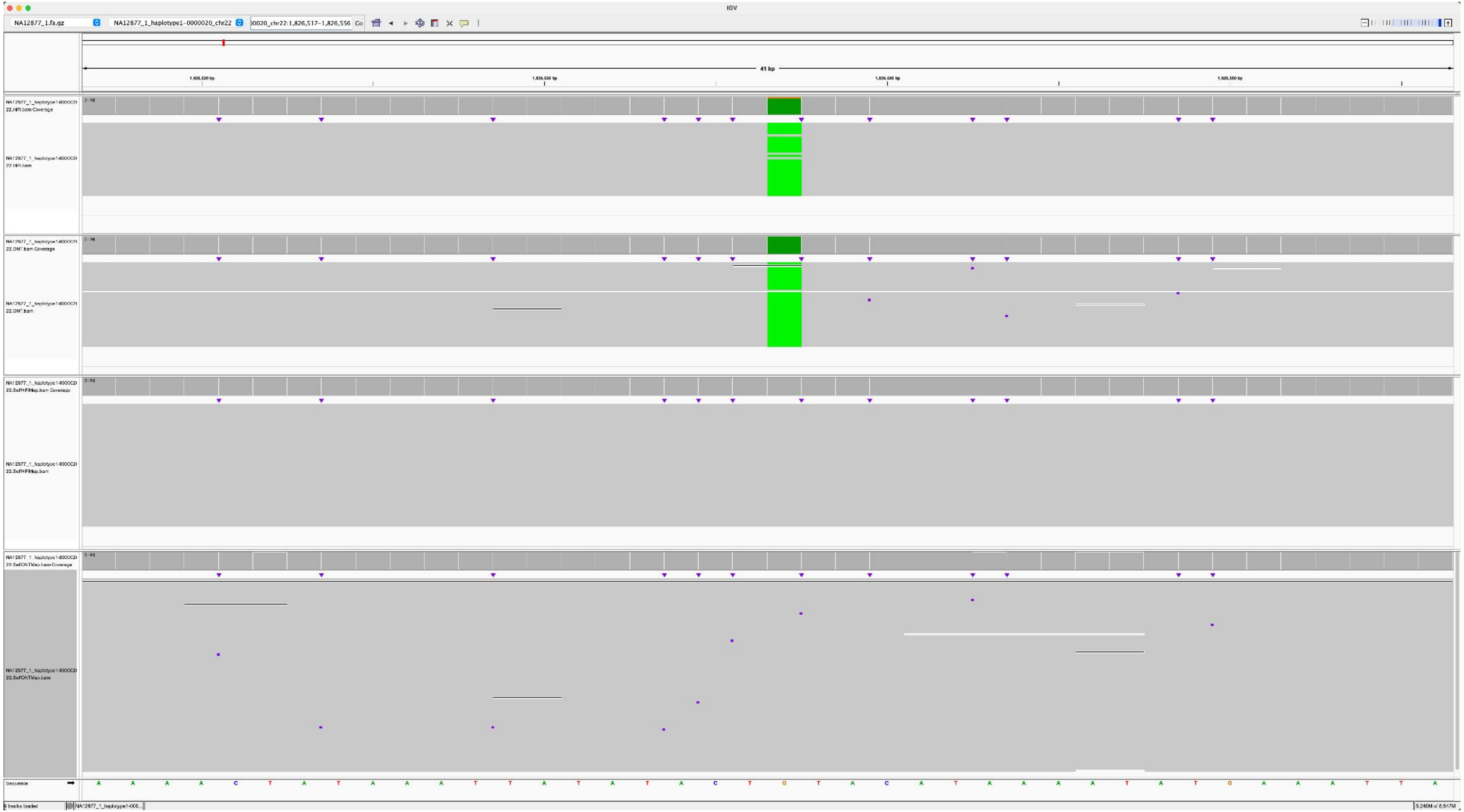

### NA12886_NA12877_1_haplotype1-0000020_chr22_7371972_T_G

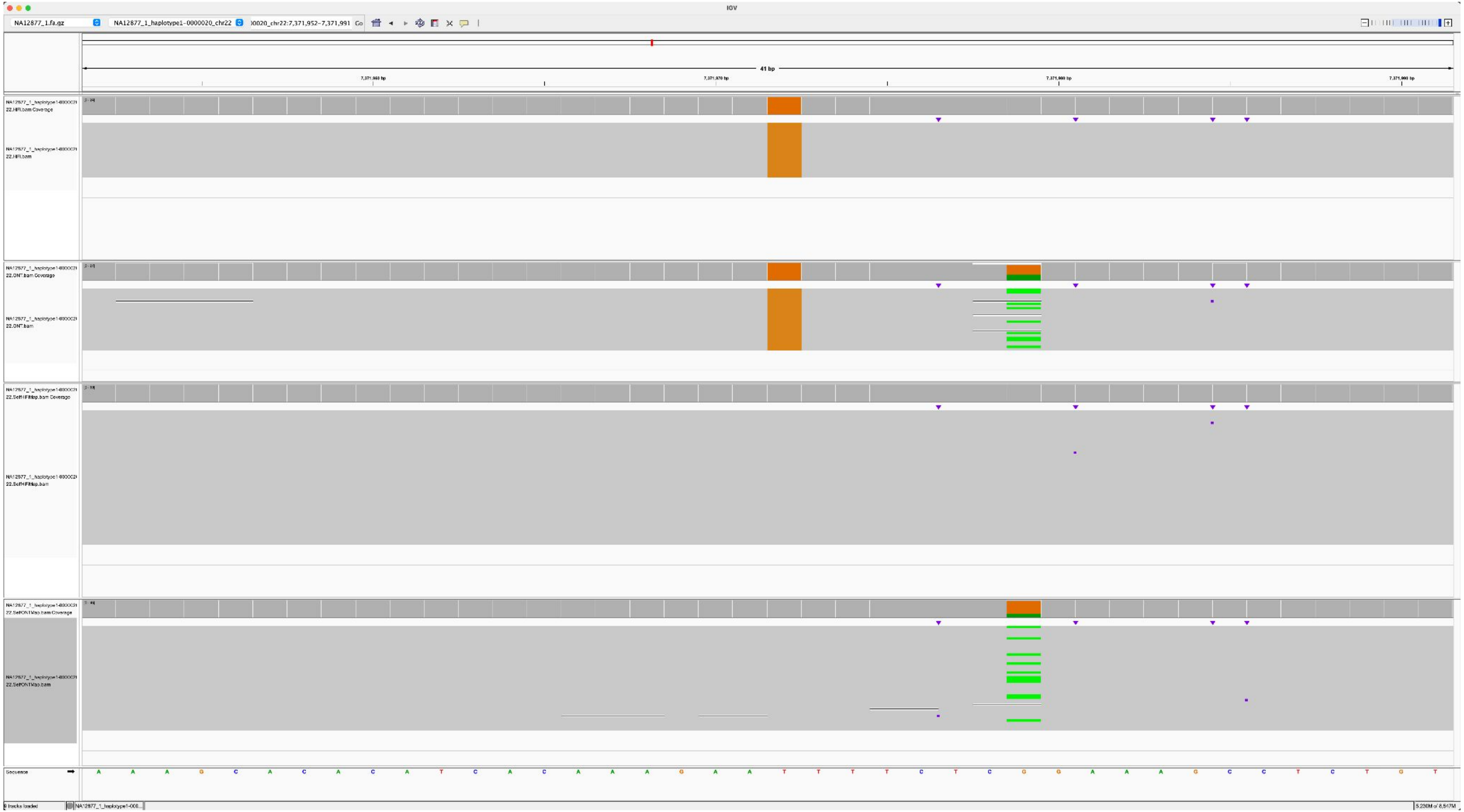

### NA12886_NA12877_1_haplotype1-0000020_chr22_7587735_T_A

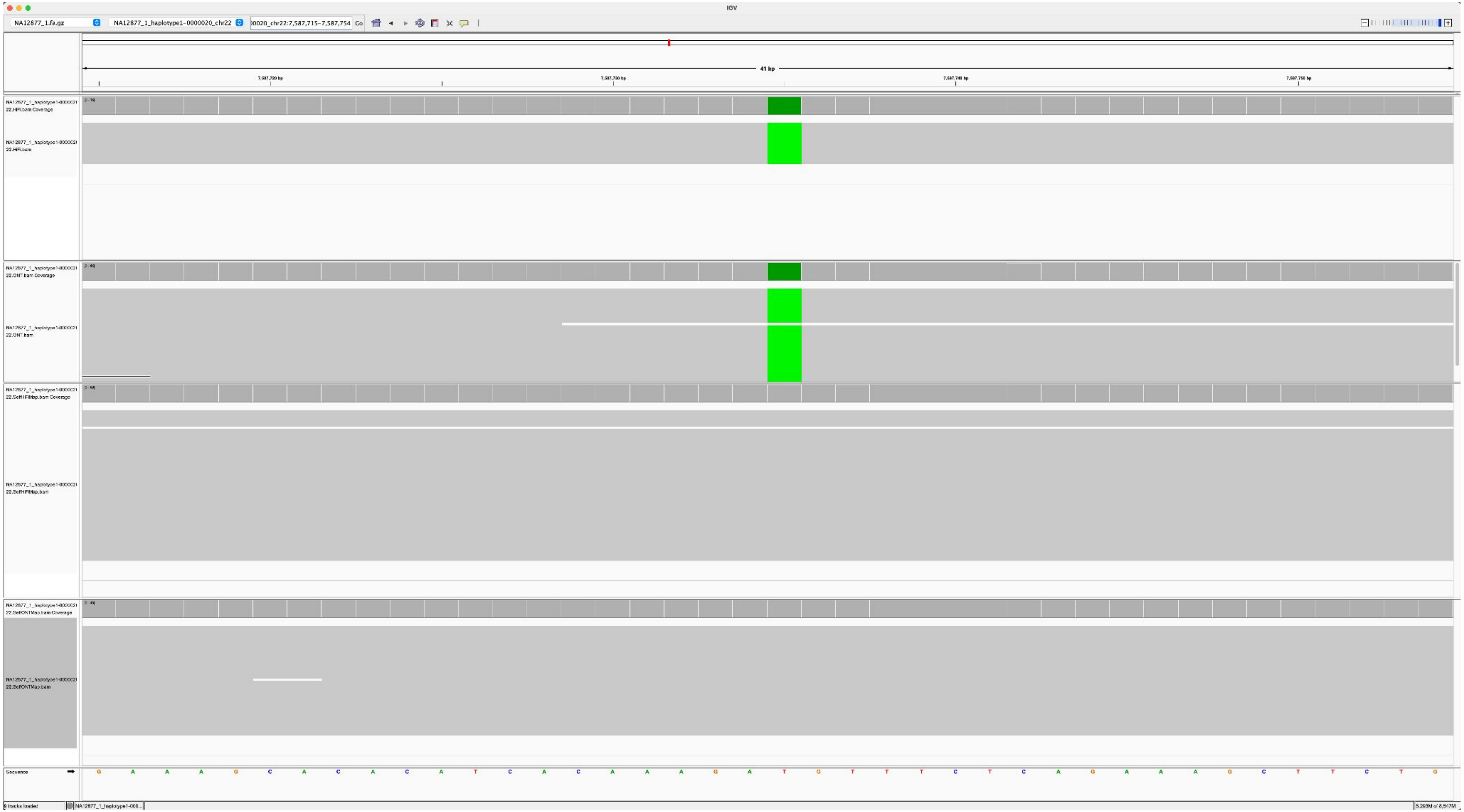

### NA12886_NA12877_1_haplotype1-0000020_chr22_11066082_G_T

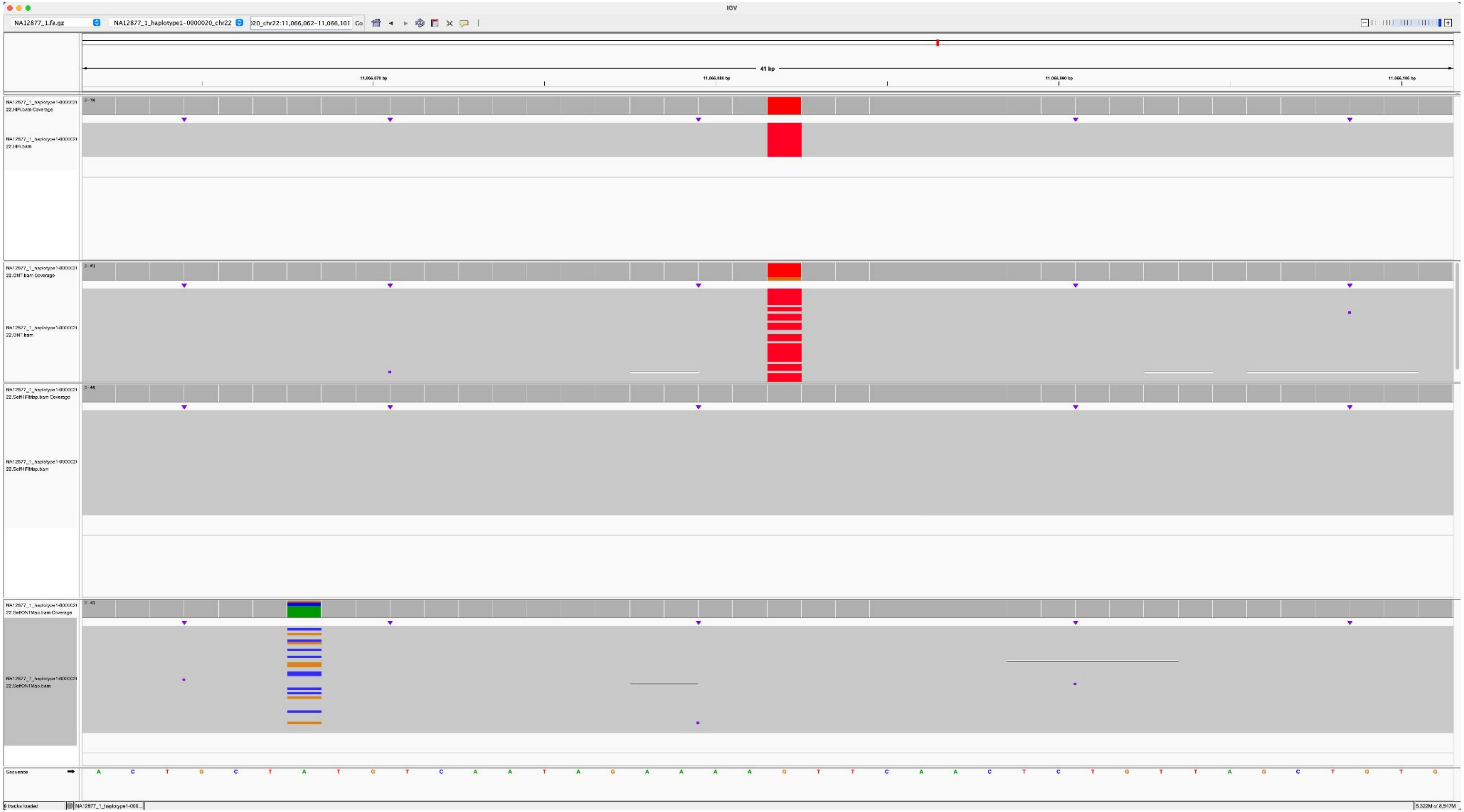

### NA12886_NA12877_1_haplotype1-0000020_chr22_12036821_G_T

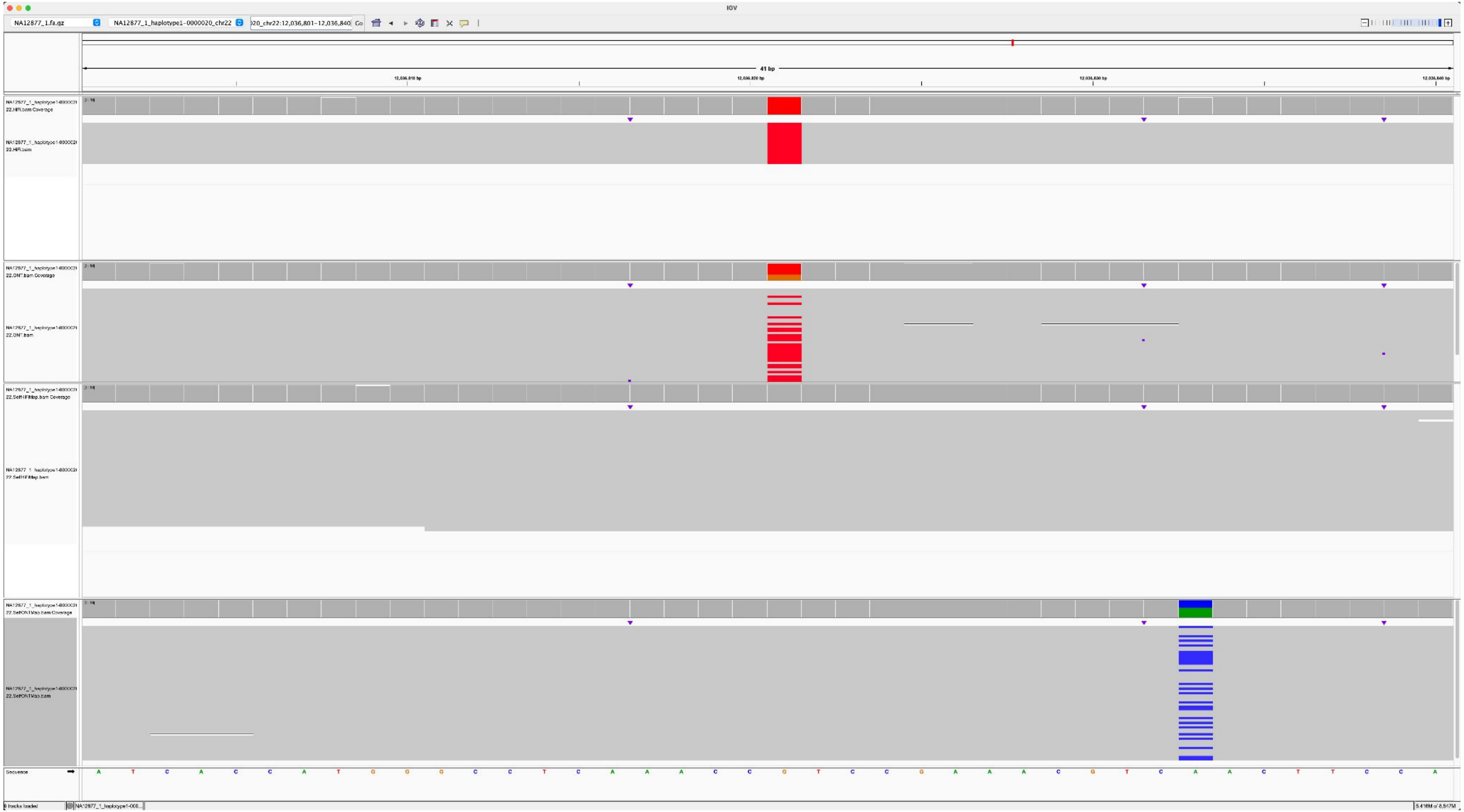

### NA12886_NA12877_1_haplotype1-0000020_chr22_1672446_G_T

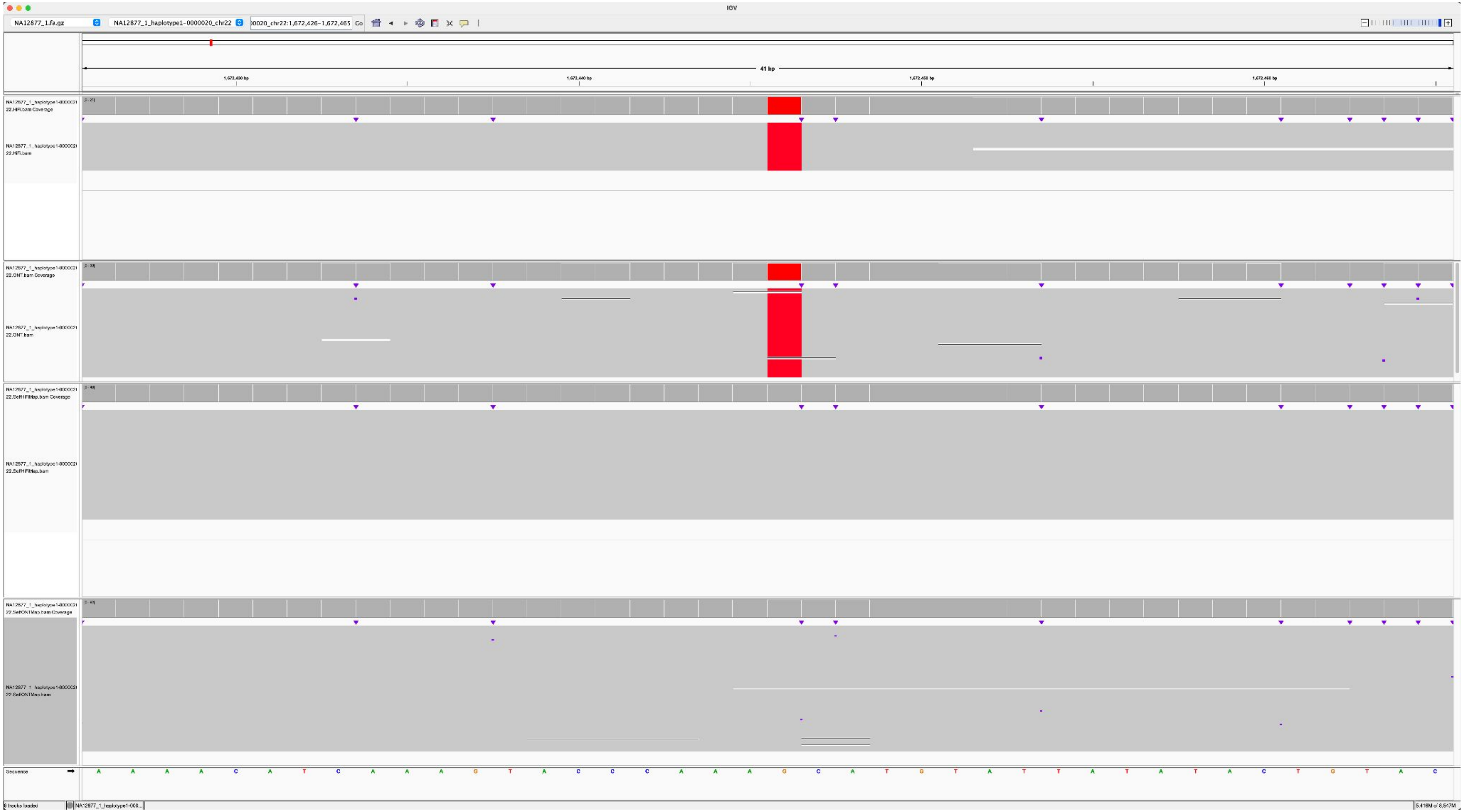

### NA12887_NA12877_1_haplotype1-0000020_chr22_11220963_C_G

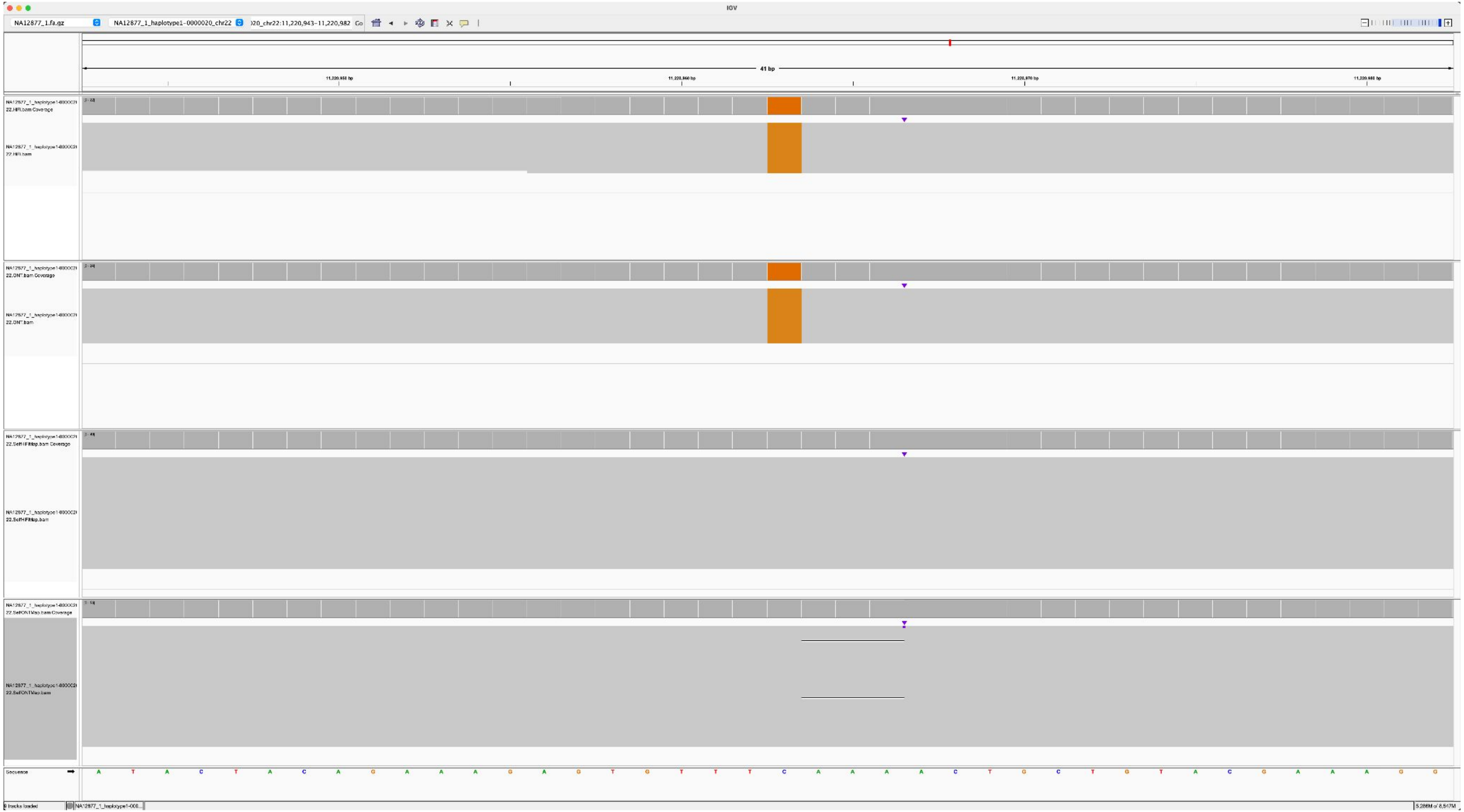

### NA12885_NA12877_2_haplotype2-0000070_chr22_8410373_G_A

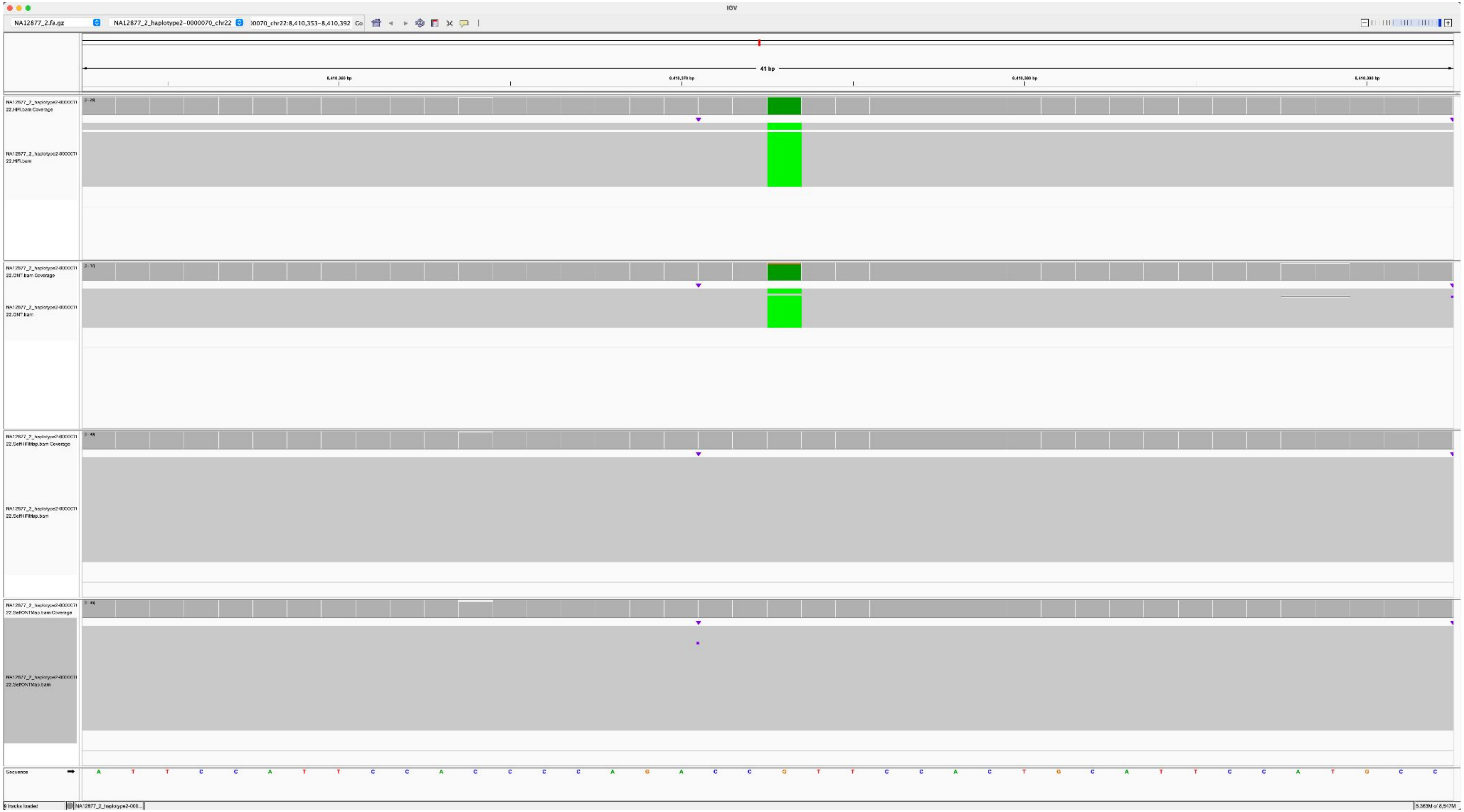

### NA12885_NA12877_2_haplotype2-0000070_chr22_1205955_A_G

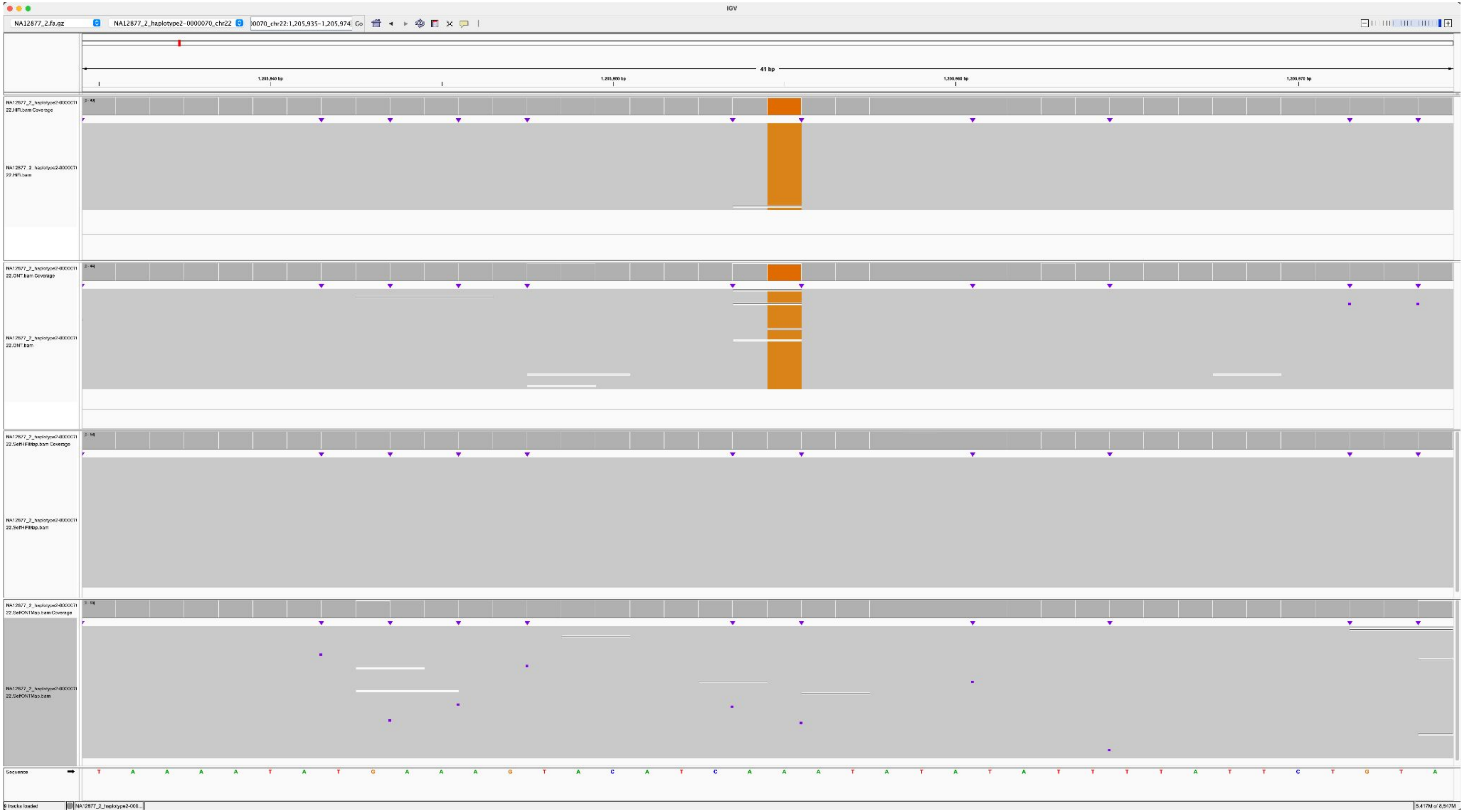

### NA12884_NA12877_2_haplotype2-0000064_chr21_8262862_A_G

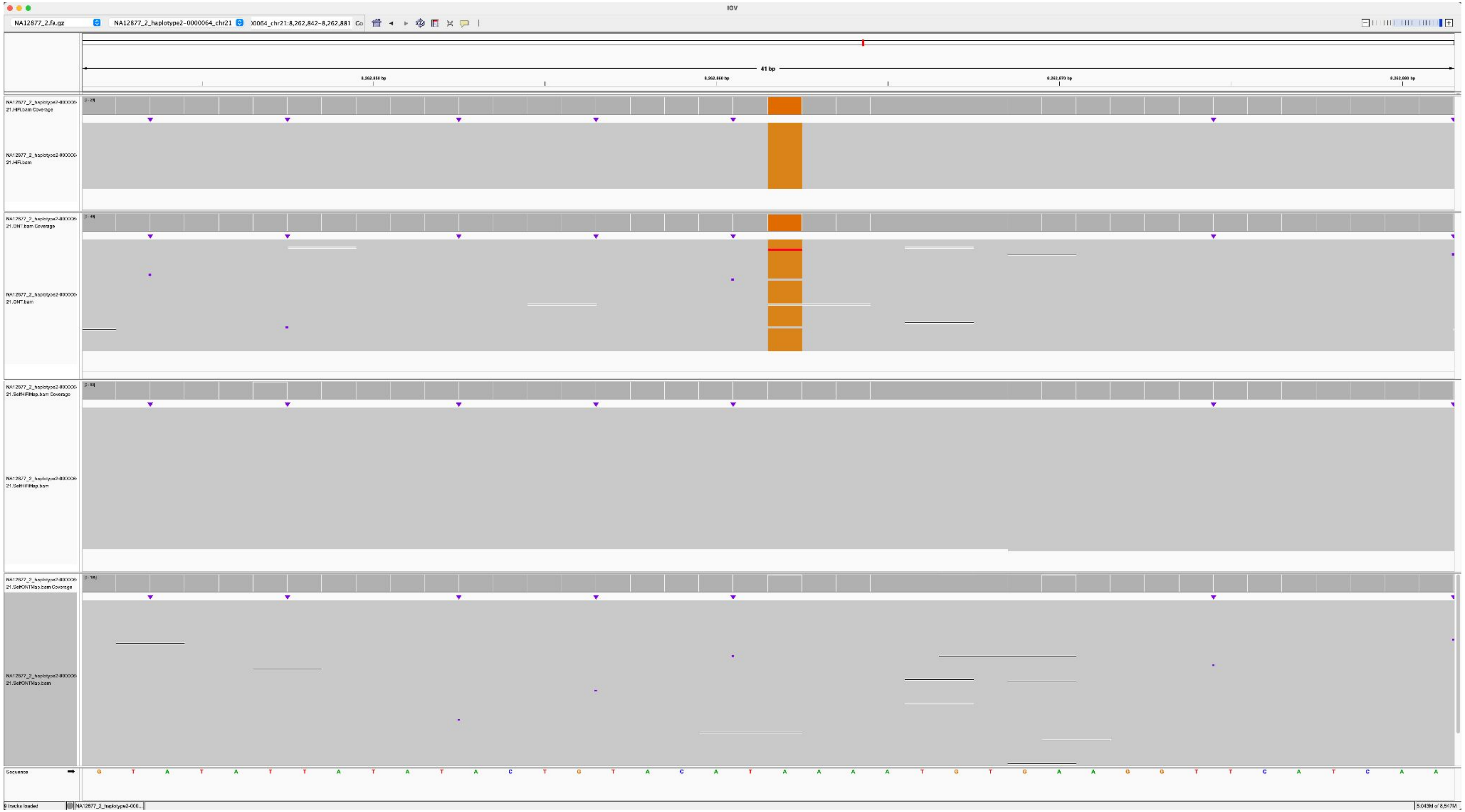

### NA12884_NA12877_2_haplotype2-0000064_chr21_4875376_A_G

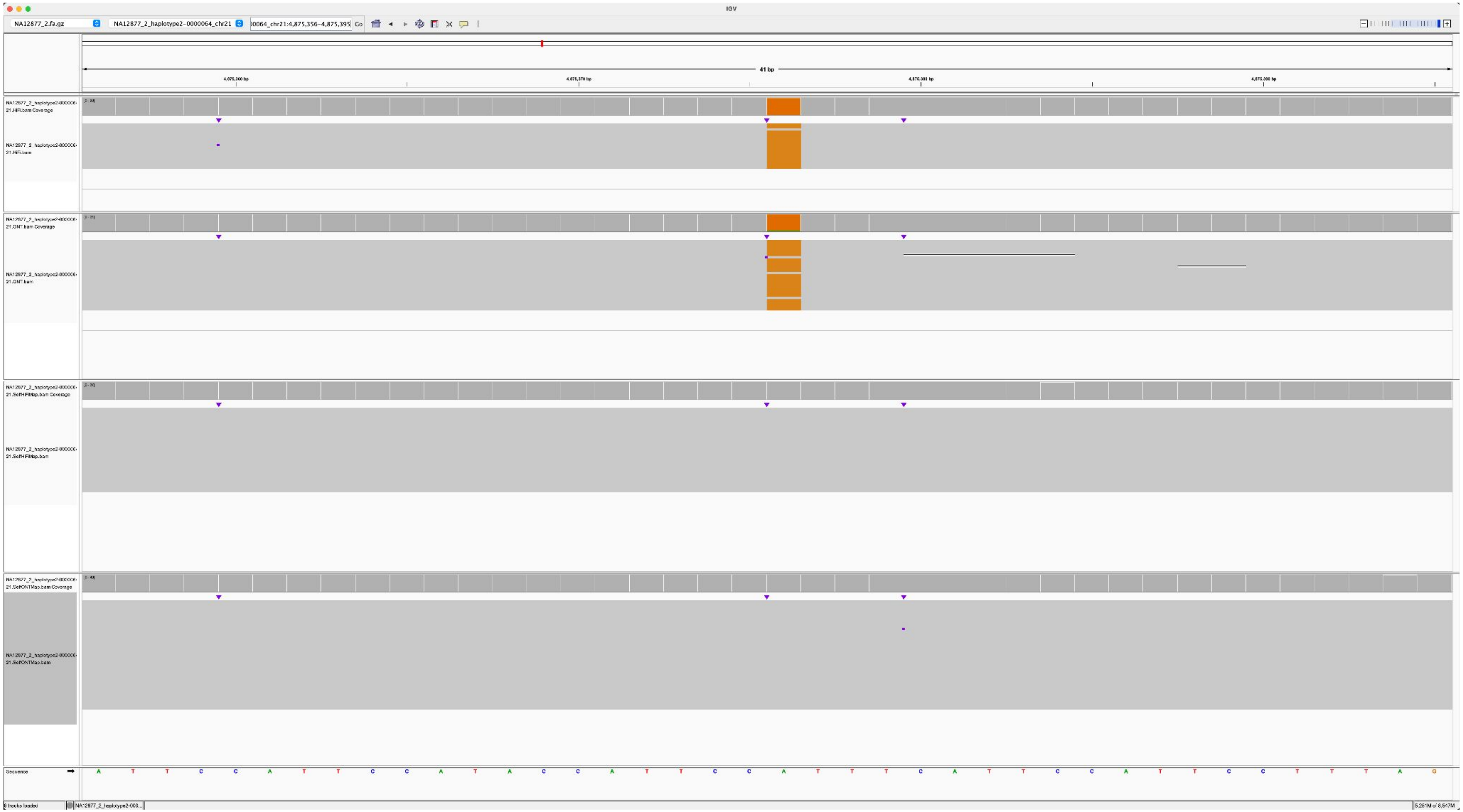

### NA12884_NA12877_2_haplotype2-0000064_chr21_1992089_A_C

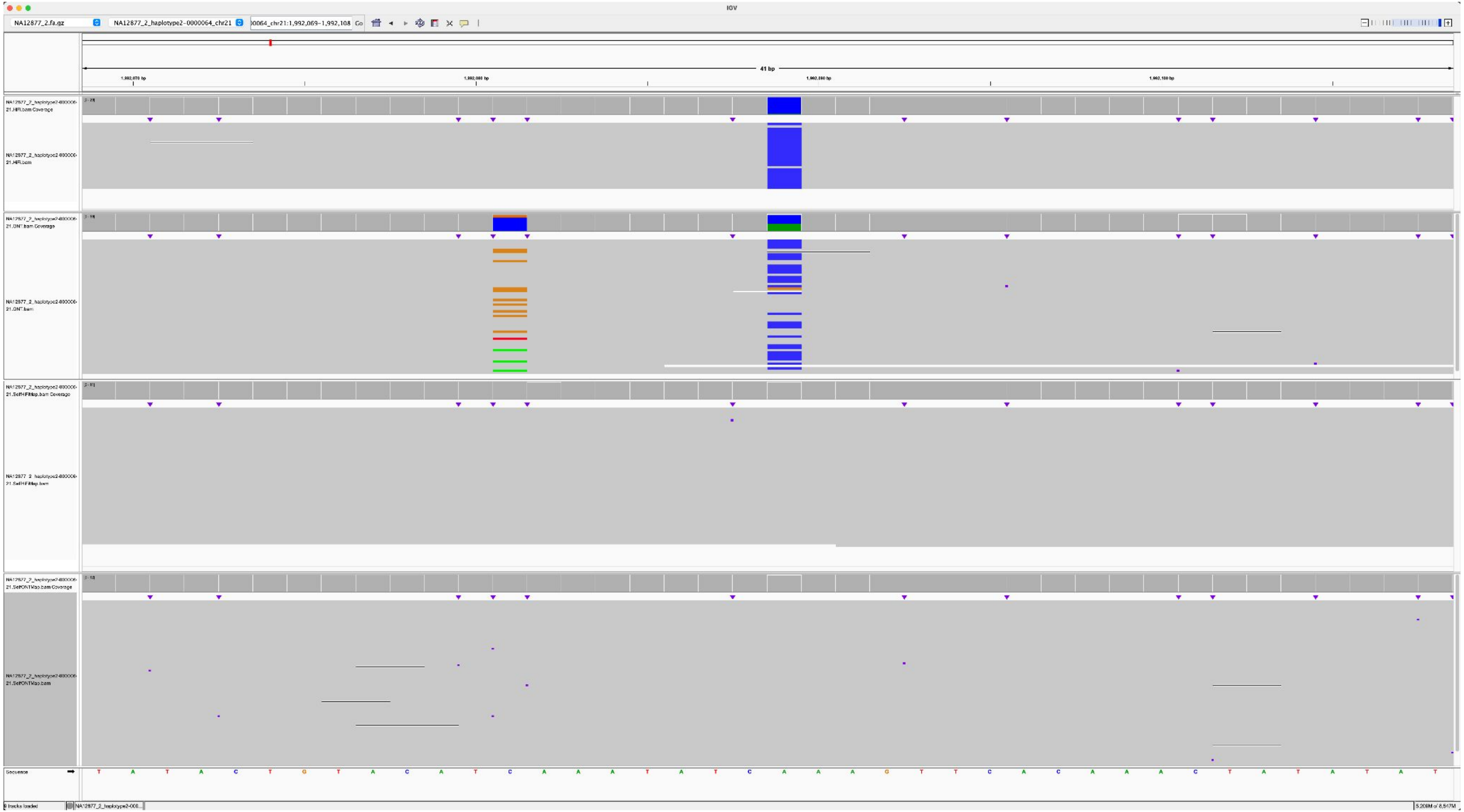

### NA12884_NA12877_2_haplotype2-0000057_chr15_4057950_T_A

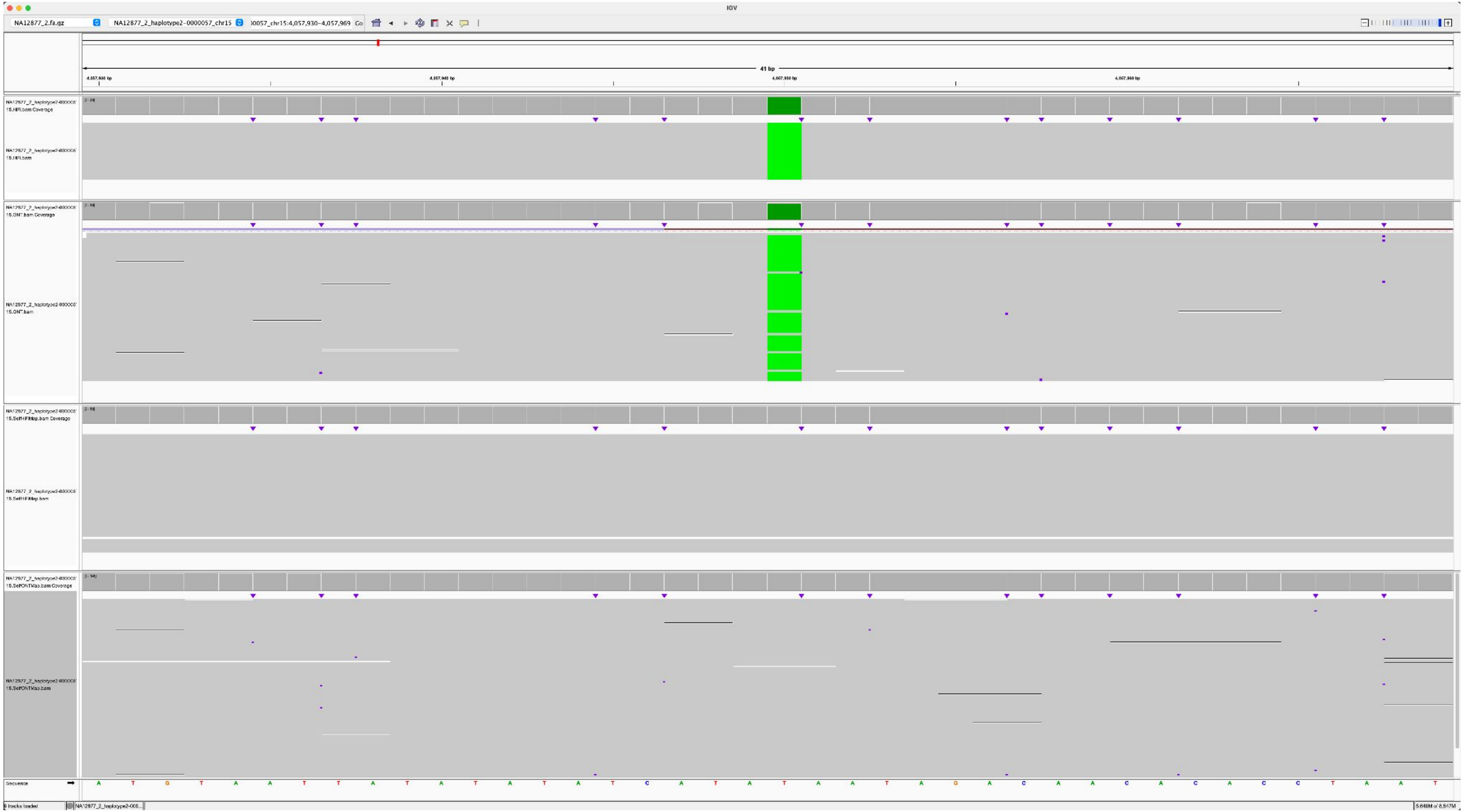

### NA12884_NA12877_2_haplotype2-0000057_chr15_5483966_C_G

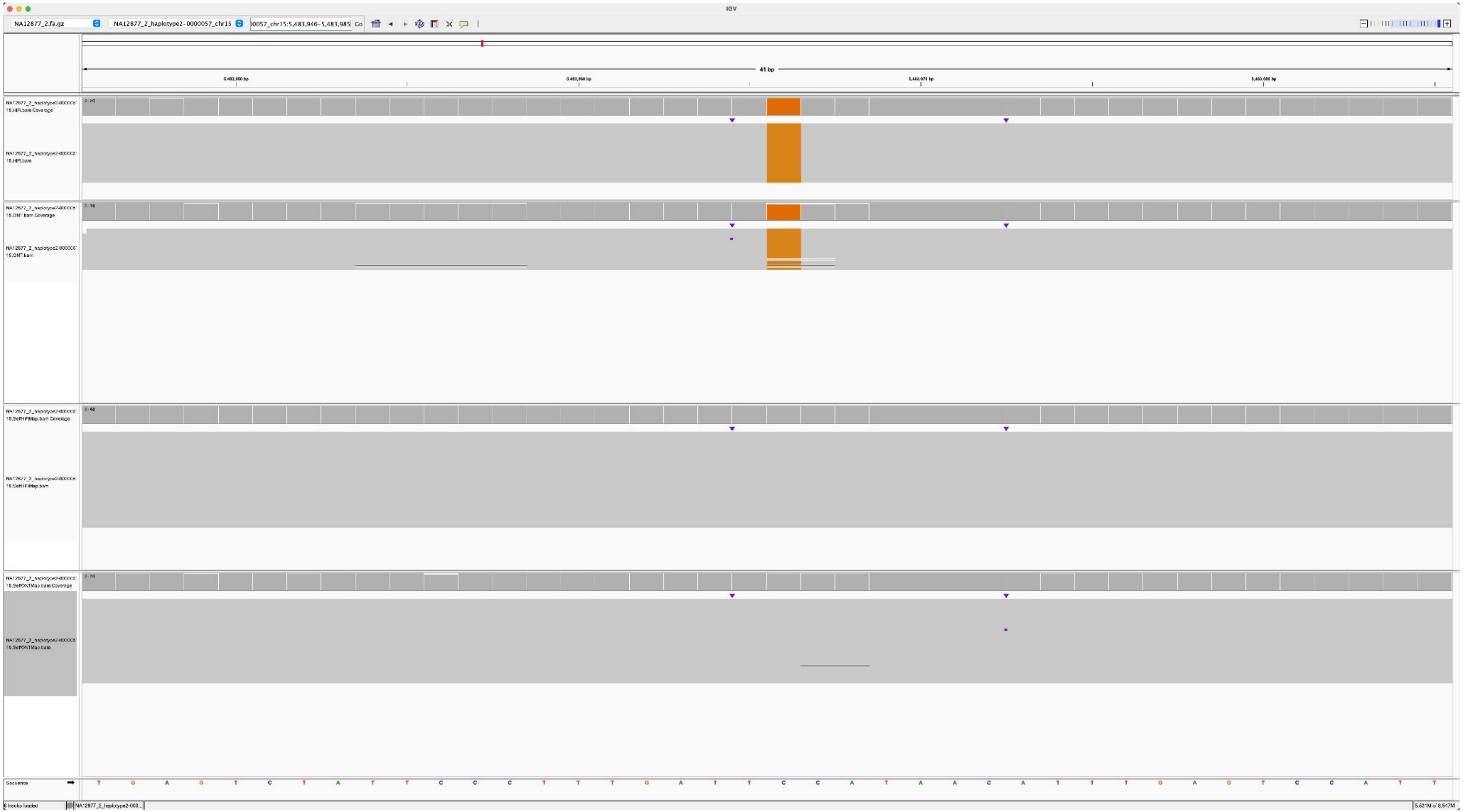

### NA12884_NA12877_2_haplotype2-0000057_chr15_8131749_C_A

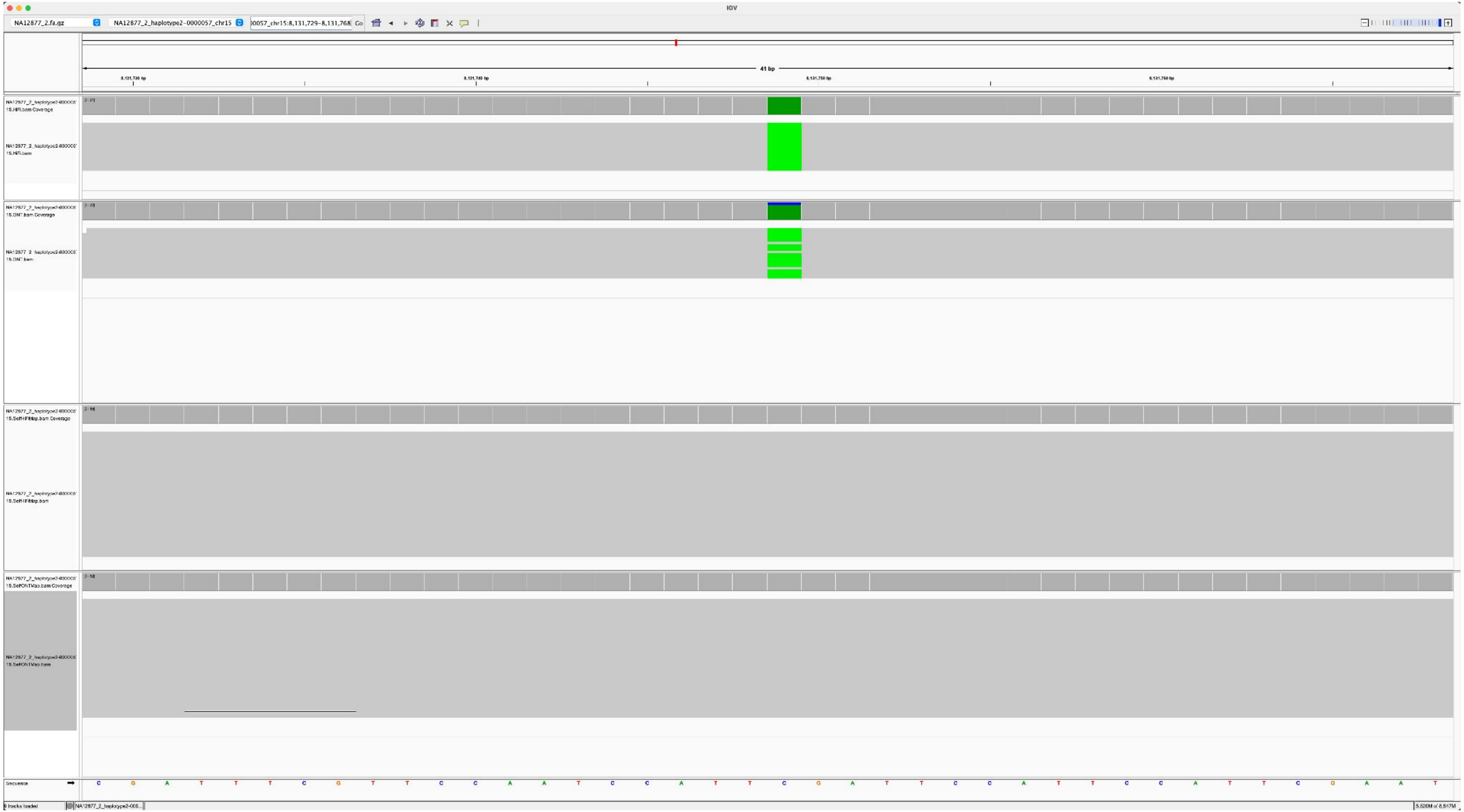

### NA12882_NA12877_2_haplotype2-0000056_chr14_1795388_T_C

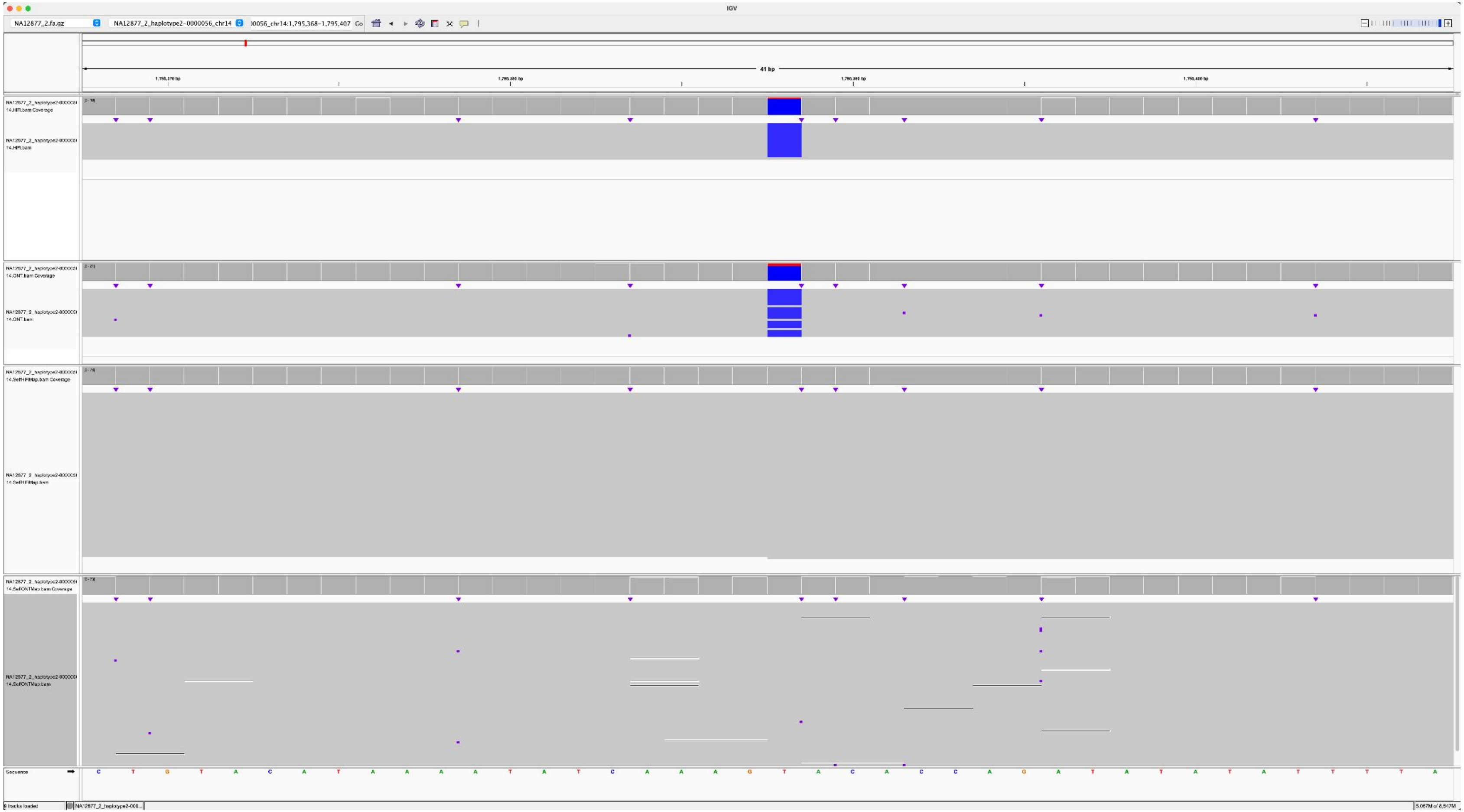

### NA12885_NA12877_2_haplotype2-0000056_chr14_1157105_A_G

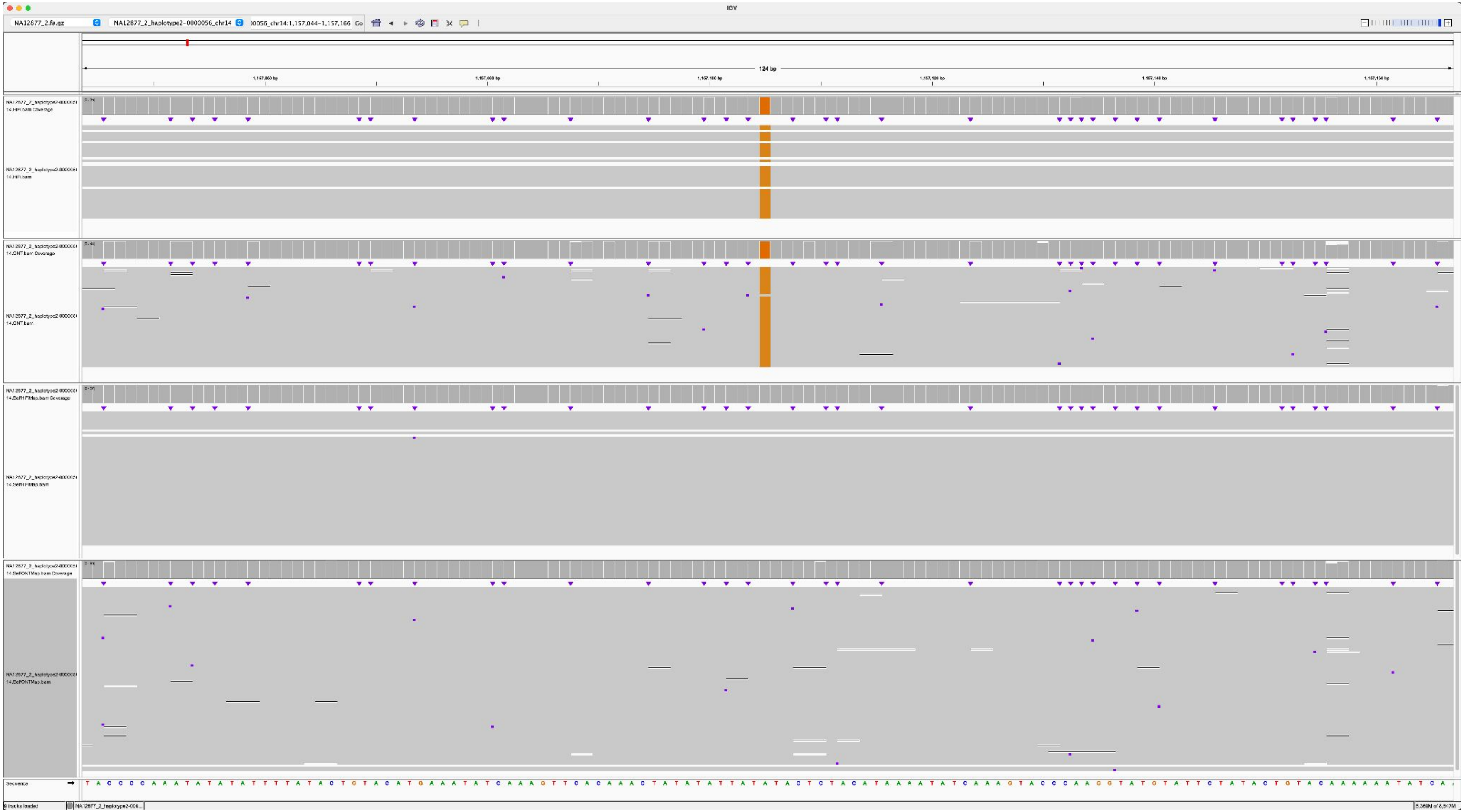

### NA12887_NA12877_2_haplotype2-0000056_chr14_3547549_T_C

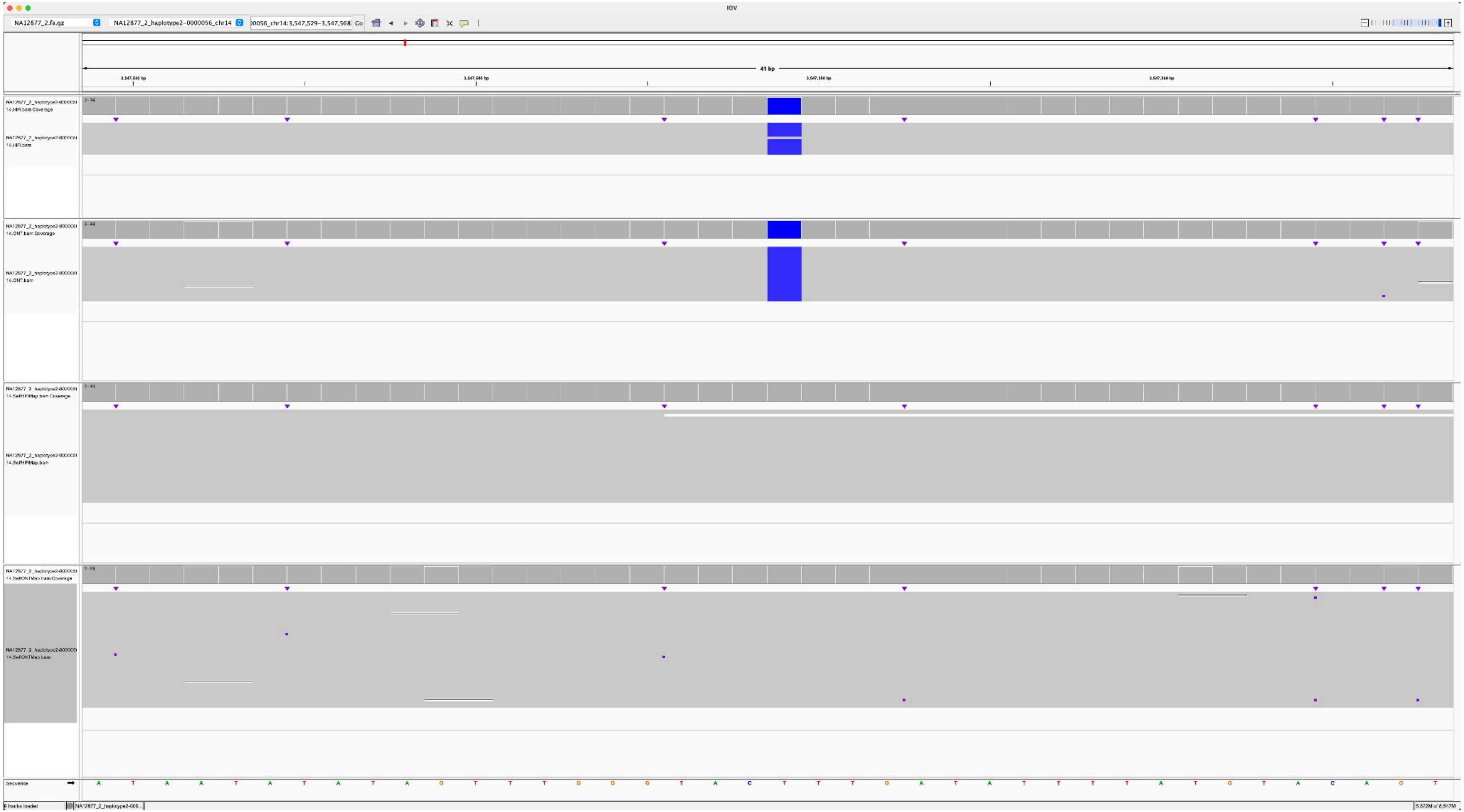

NA12883_NA12877_2_haplotype2-0000046_chr13_8253115_C_G

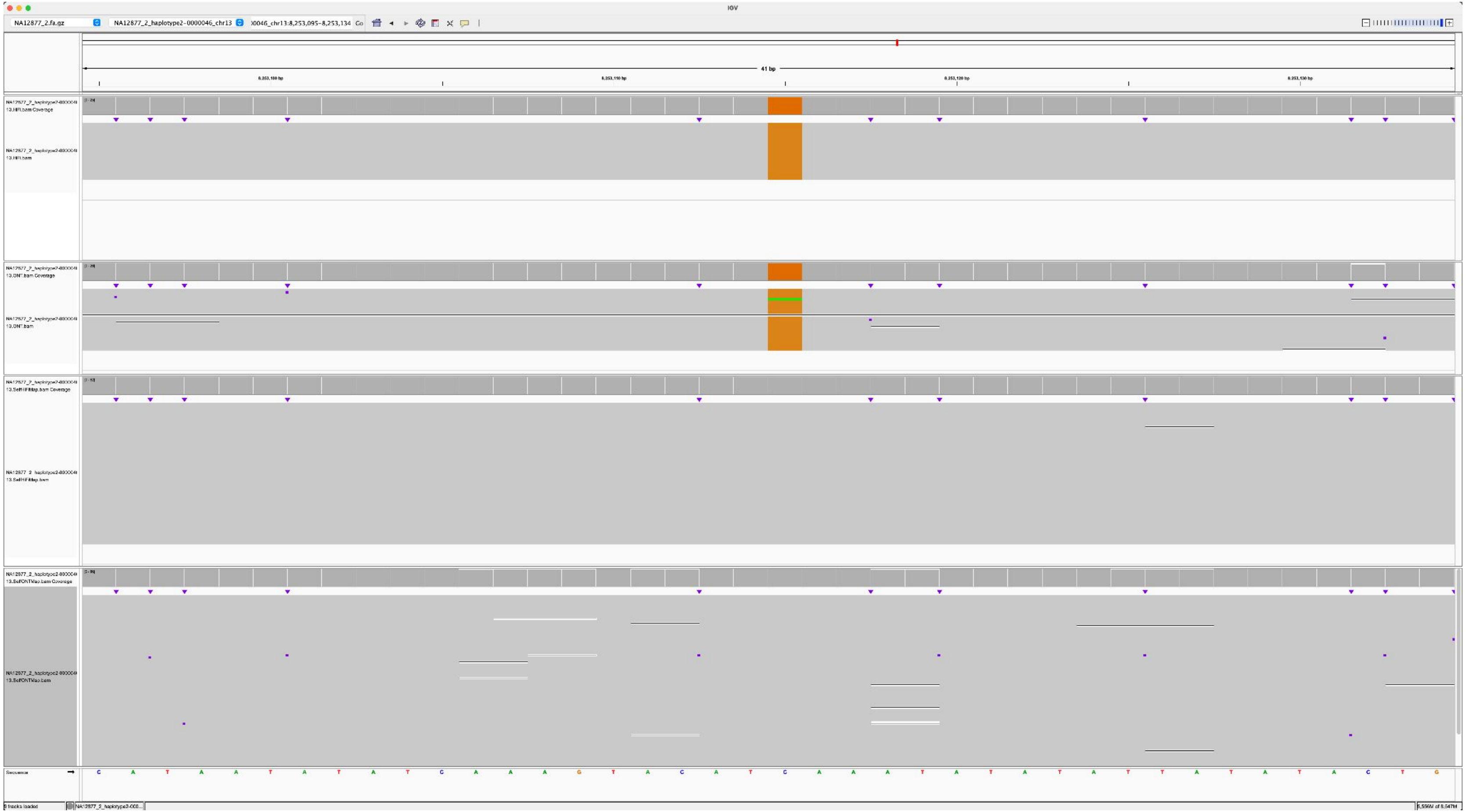

### NA12883_NA12877_2_haplotype2-0000046_chr13_407411_A_G

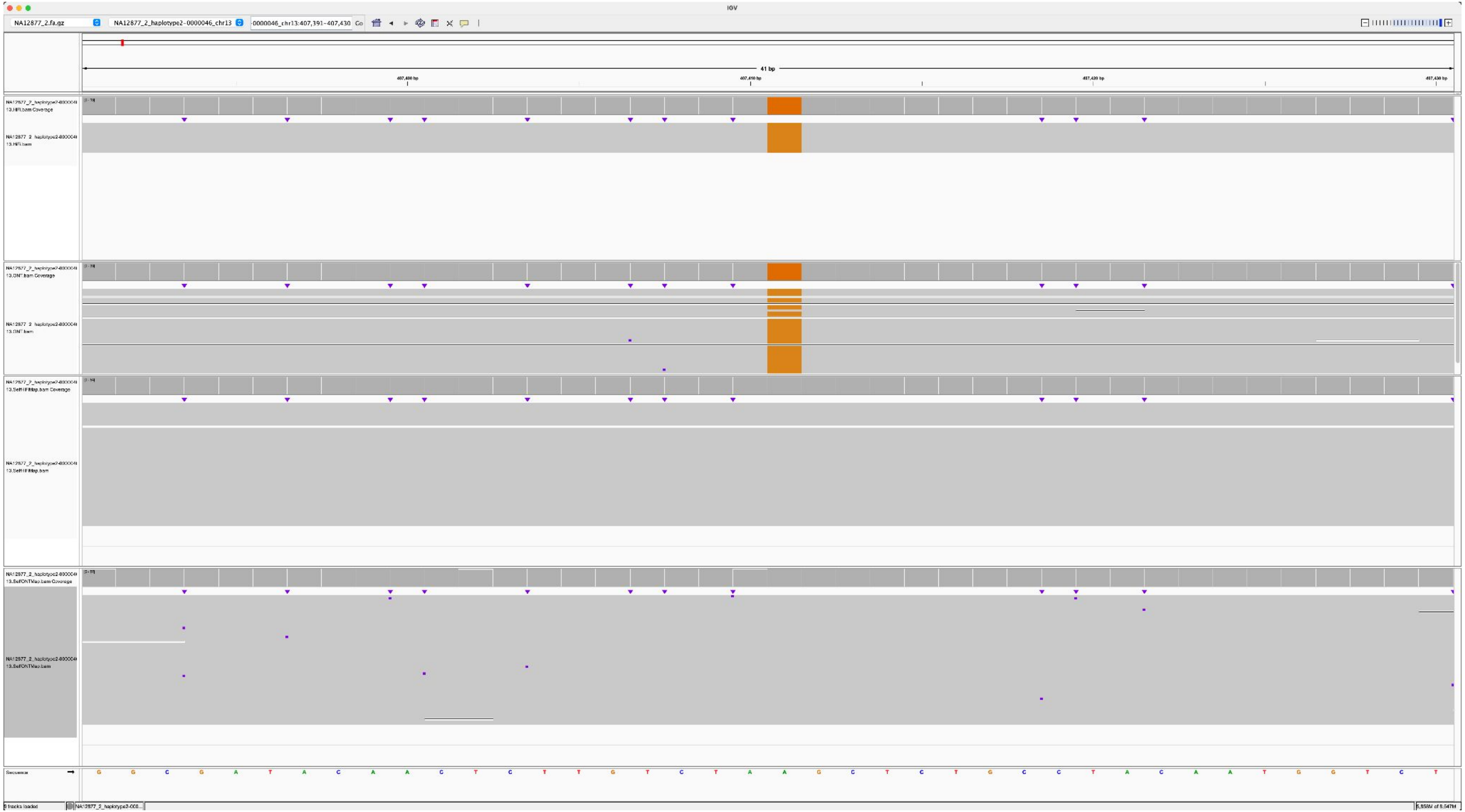

### NA12887_NA12878_2_haplotype2-0000041_chr14_11623273_C_T

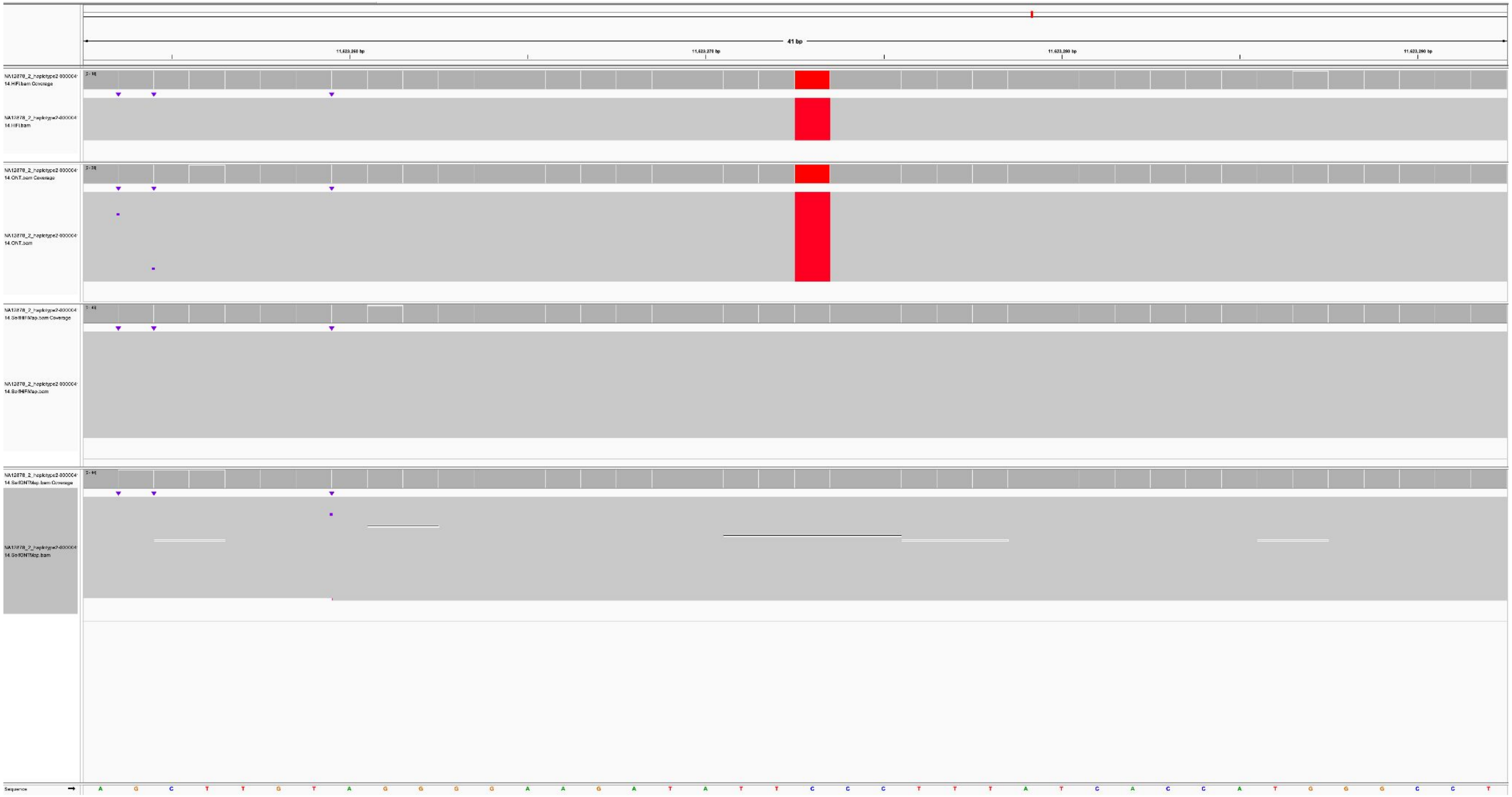

K200081_NA12879_2_haplotype2-0000092_chr13_4505530_C_G

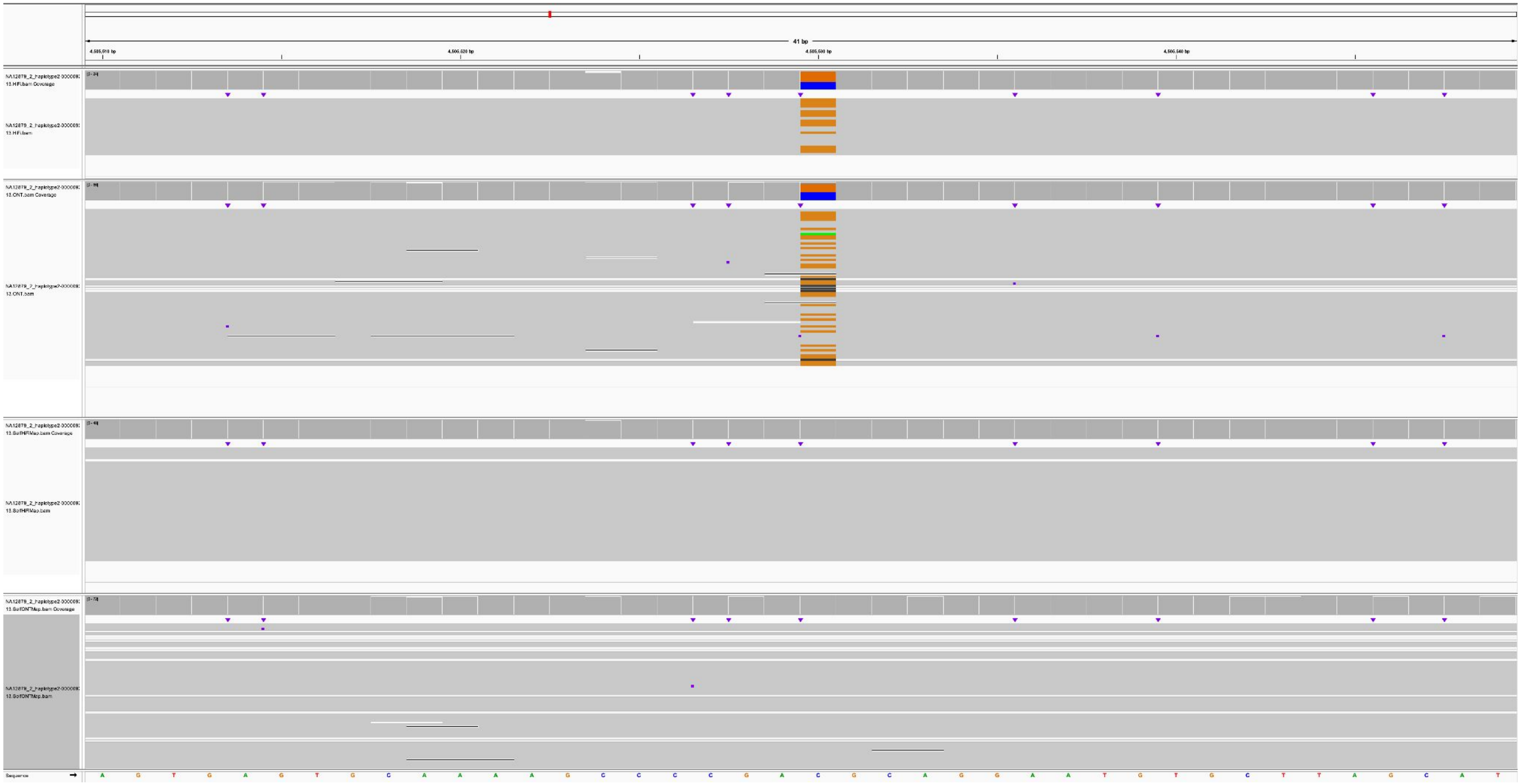

### K200081_NA12879_1_haplotype1-0000015_chr14_8548143_A_C

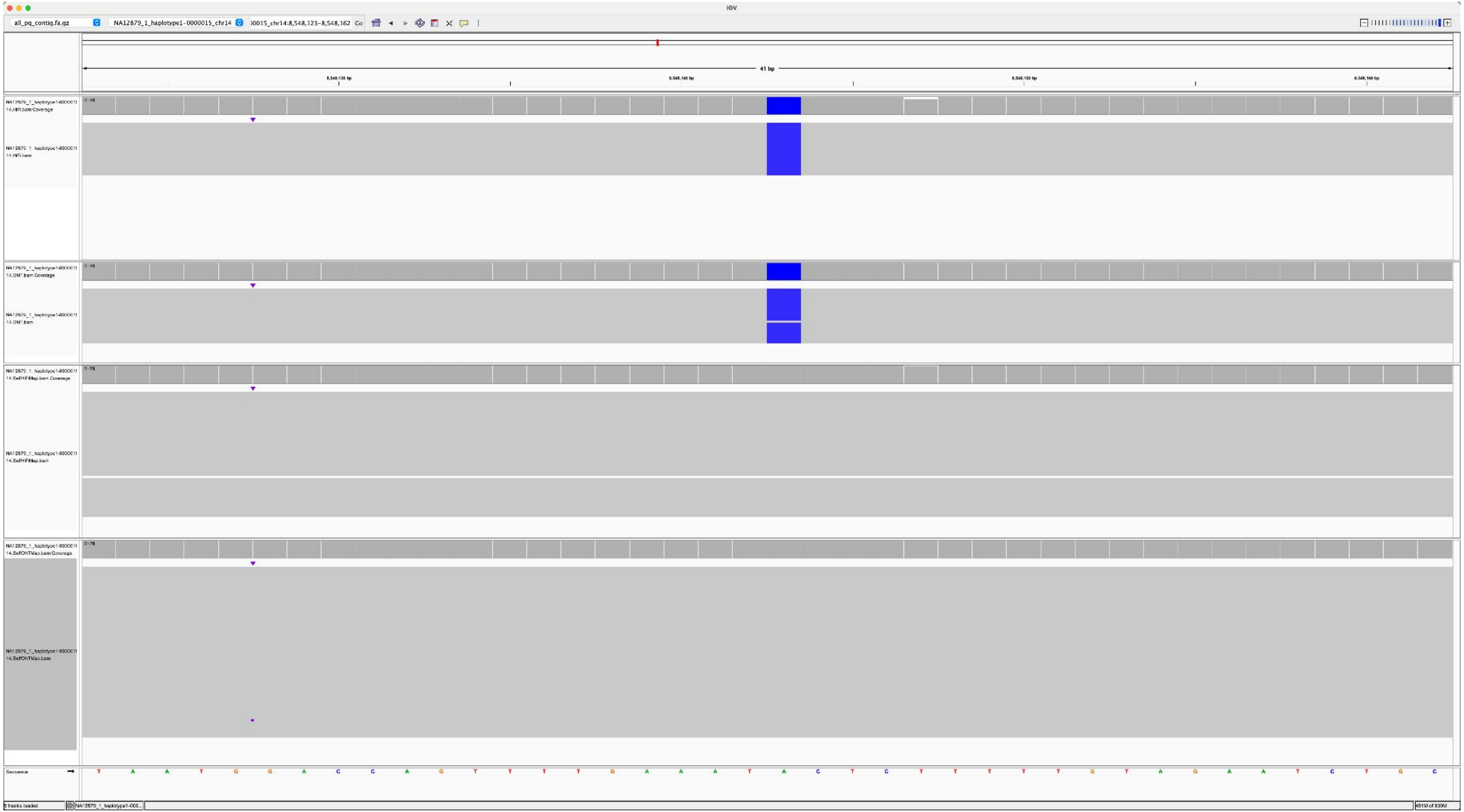

K200081_NA12879_2_haplotype2-0000086_chr22_7632468_C_T

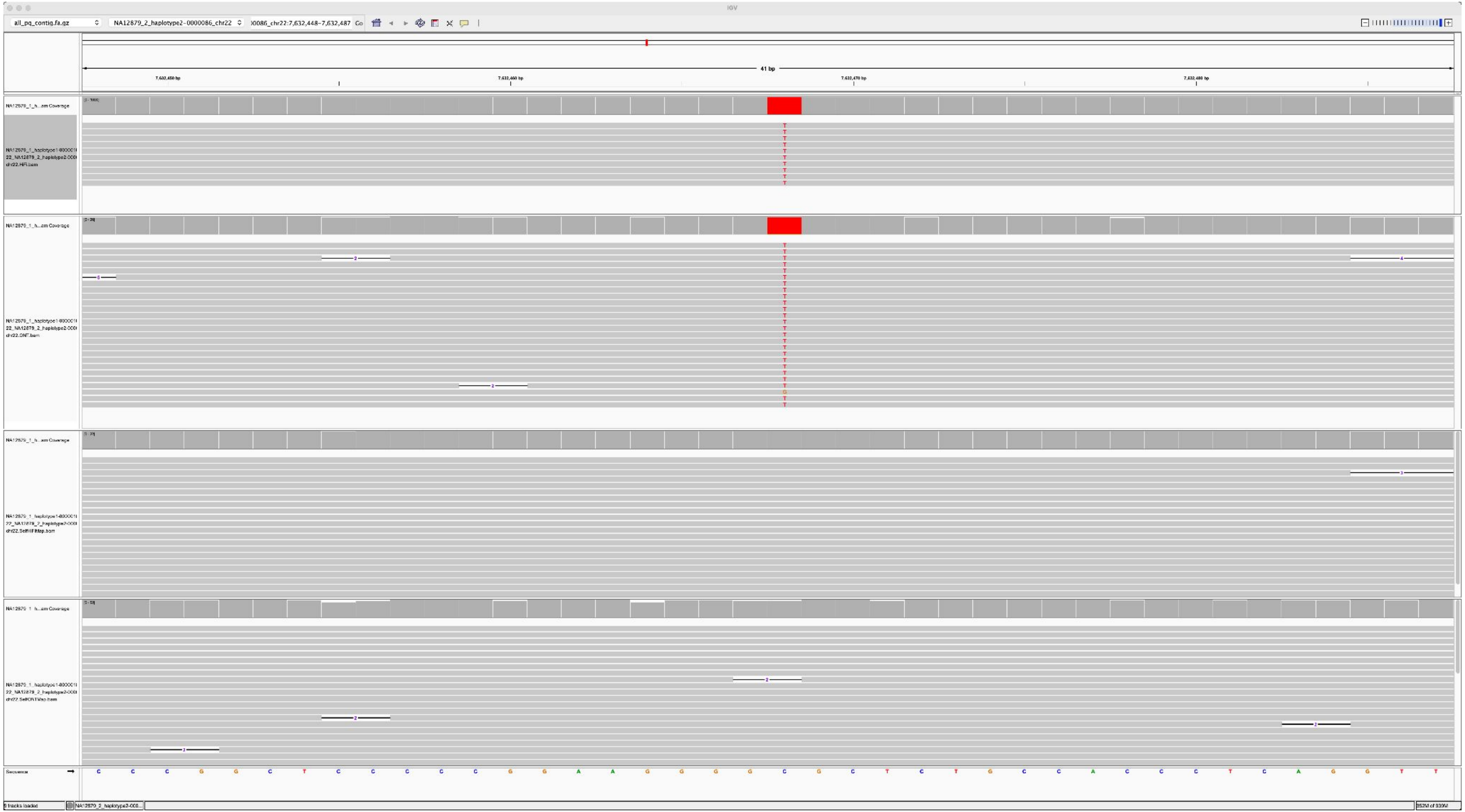

K200081_NA12879_2_haplotype2-0000086_chr22_8422020_T_G

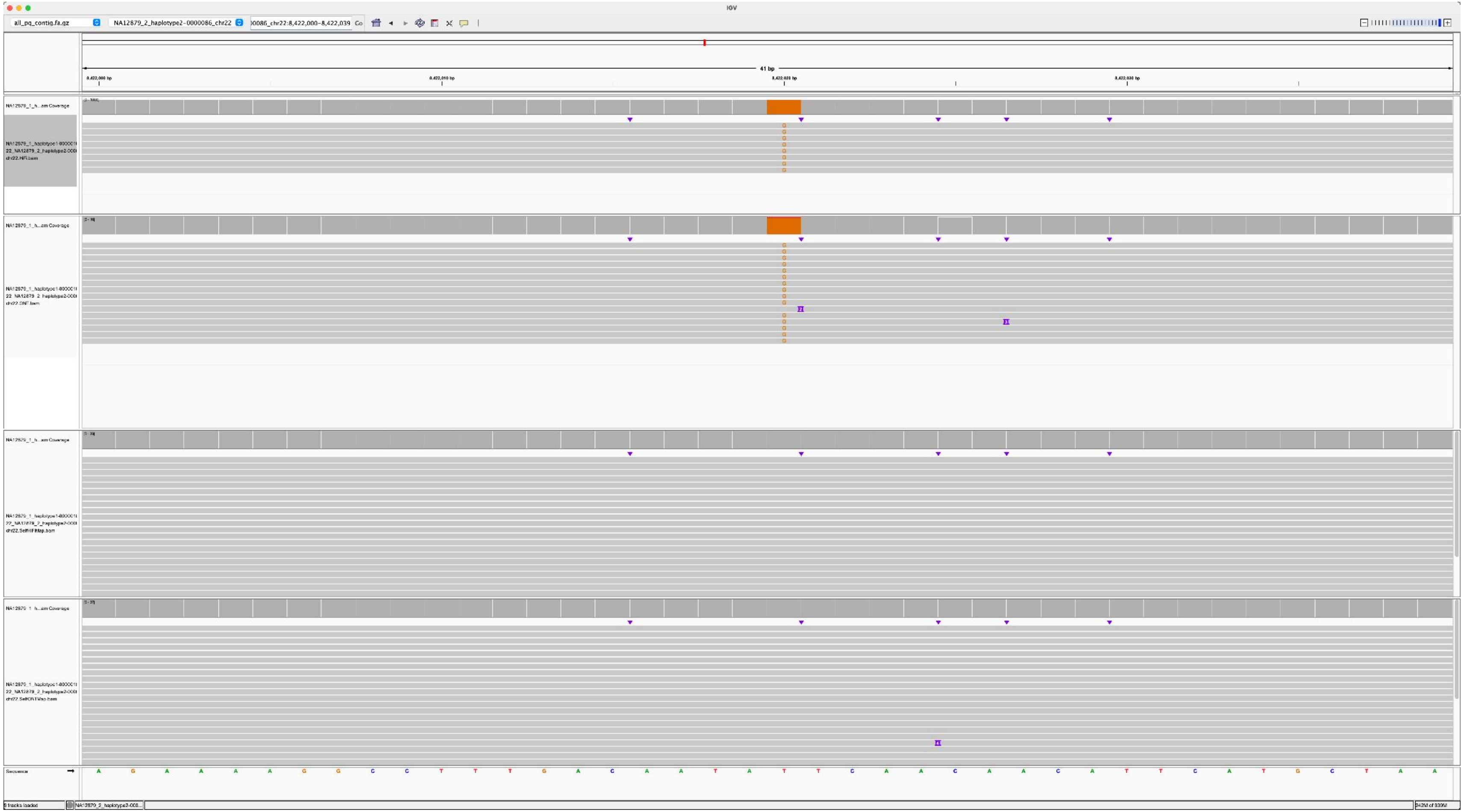

K200082_NA12879_2_haplotype2-0000086_chr22_11966149_C_A

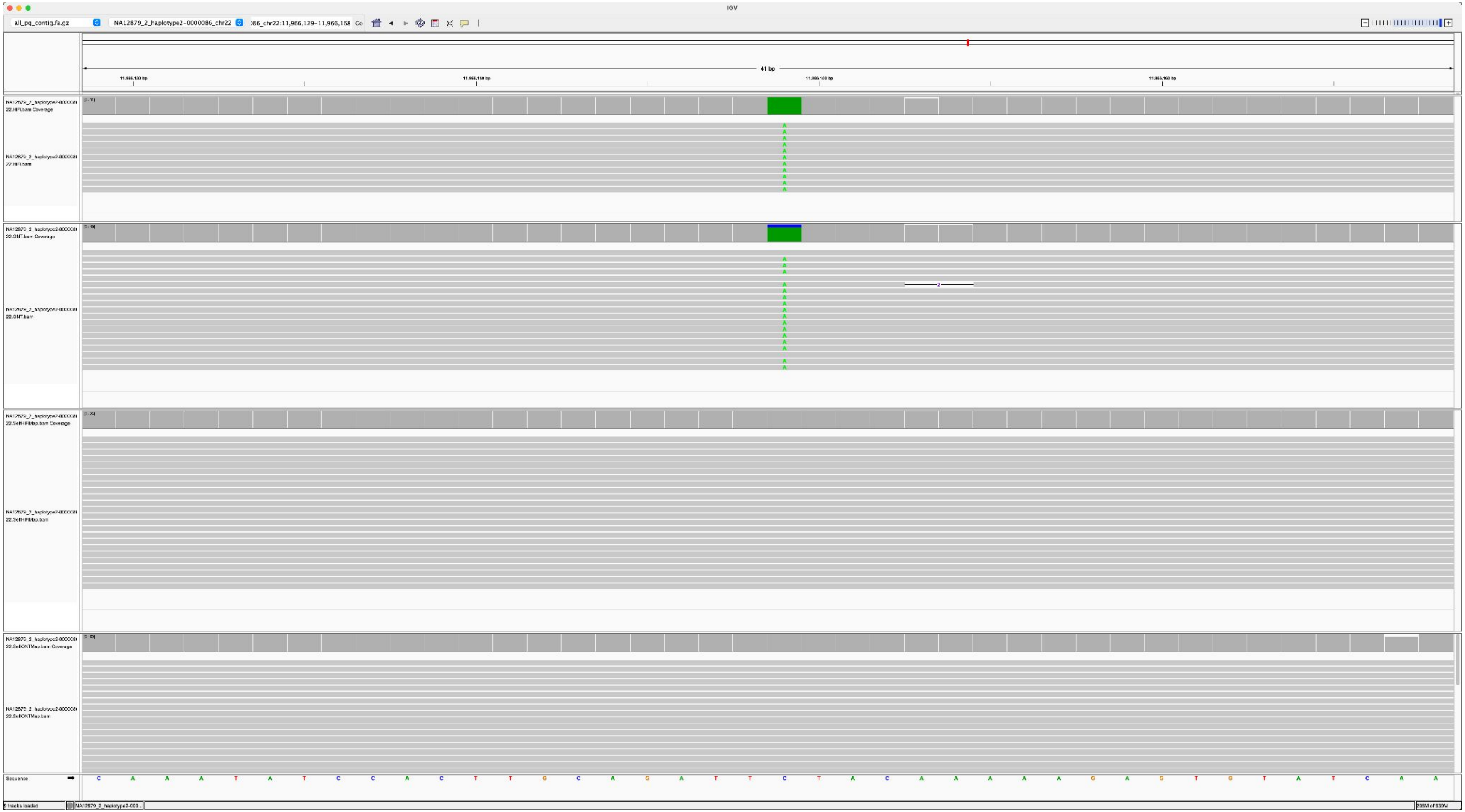

K200084_NA12879_2_haplotype2-0000083_chr14_11683995_G_A

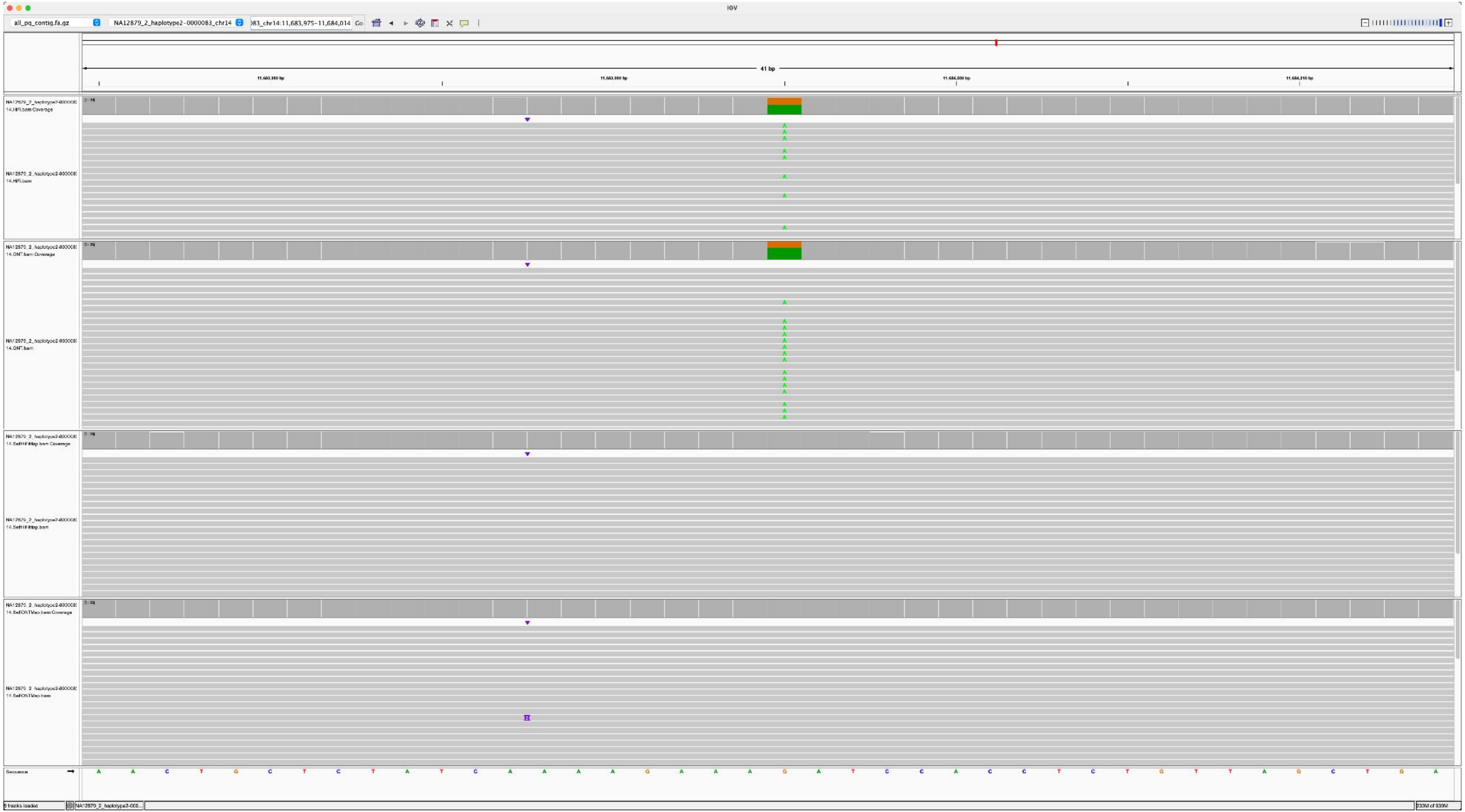

K200084_NA12879_2_haplotype2-0000075_chr15_3760599_G_C

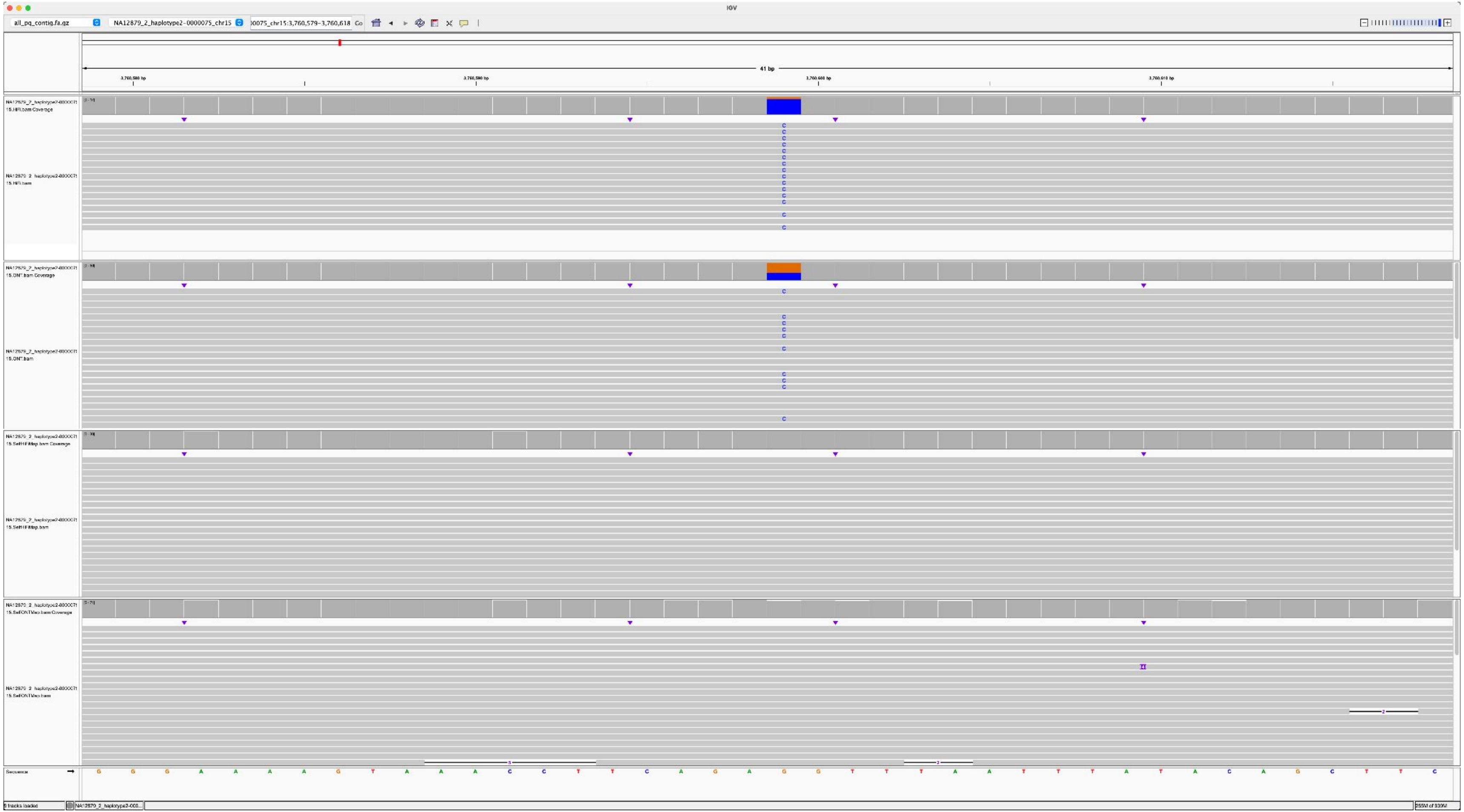

K200084_NA12879_2_haplotype2-0000075_chr15_13275536_C_T

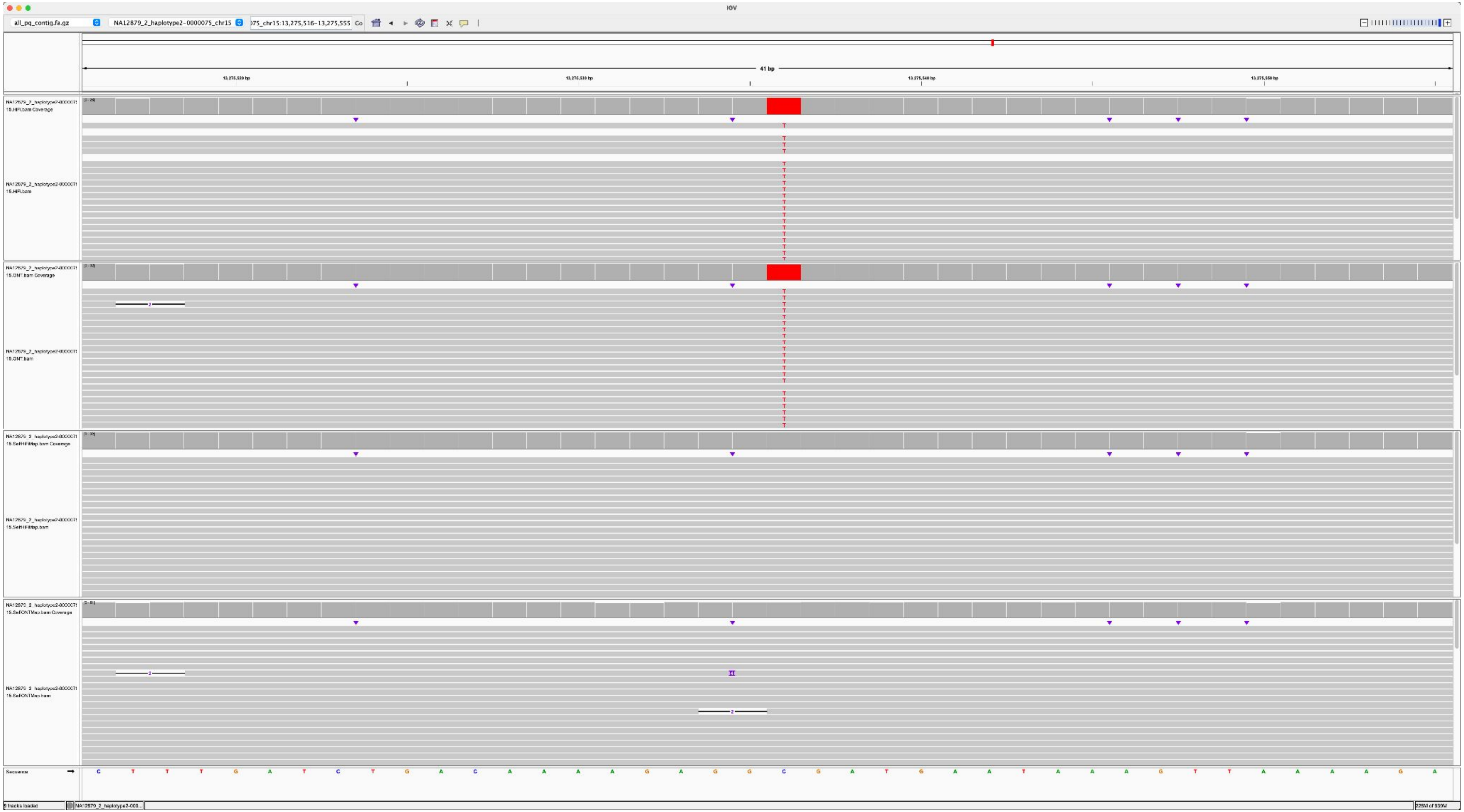

### K200085_NA12879_2_haplotype2-0000086_chr22_12864909_G_C

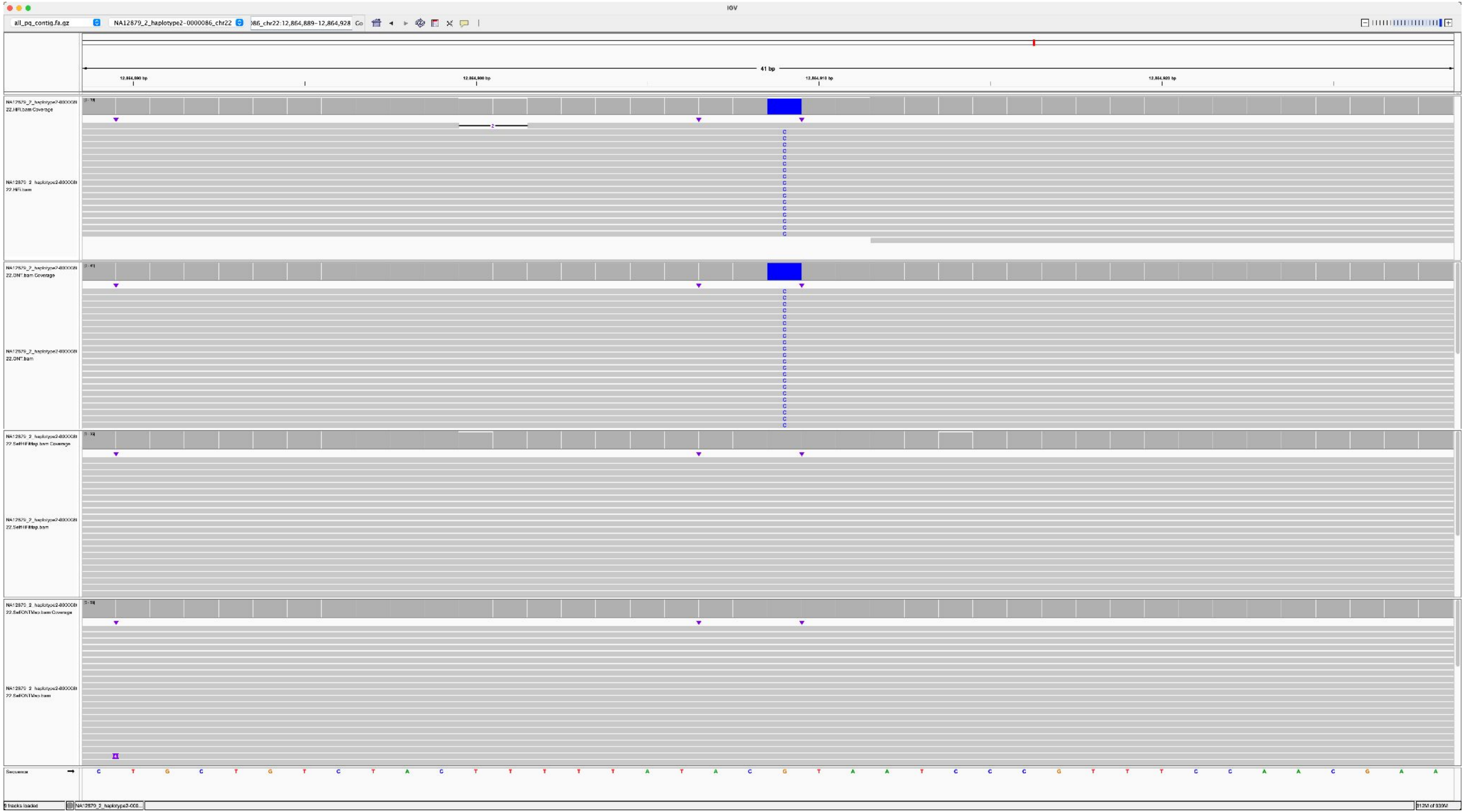

### K200087_NA12879_1_haplotype1-0000015_chr14_11265763_A_G

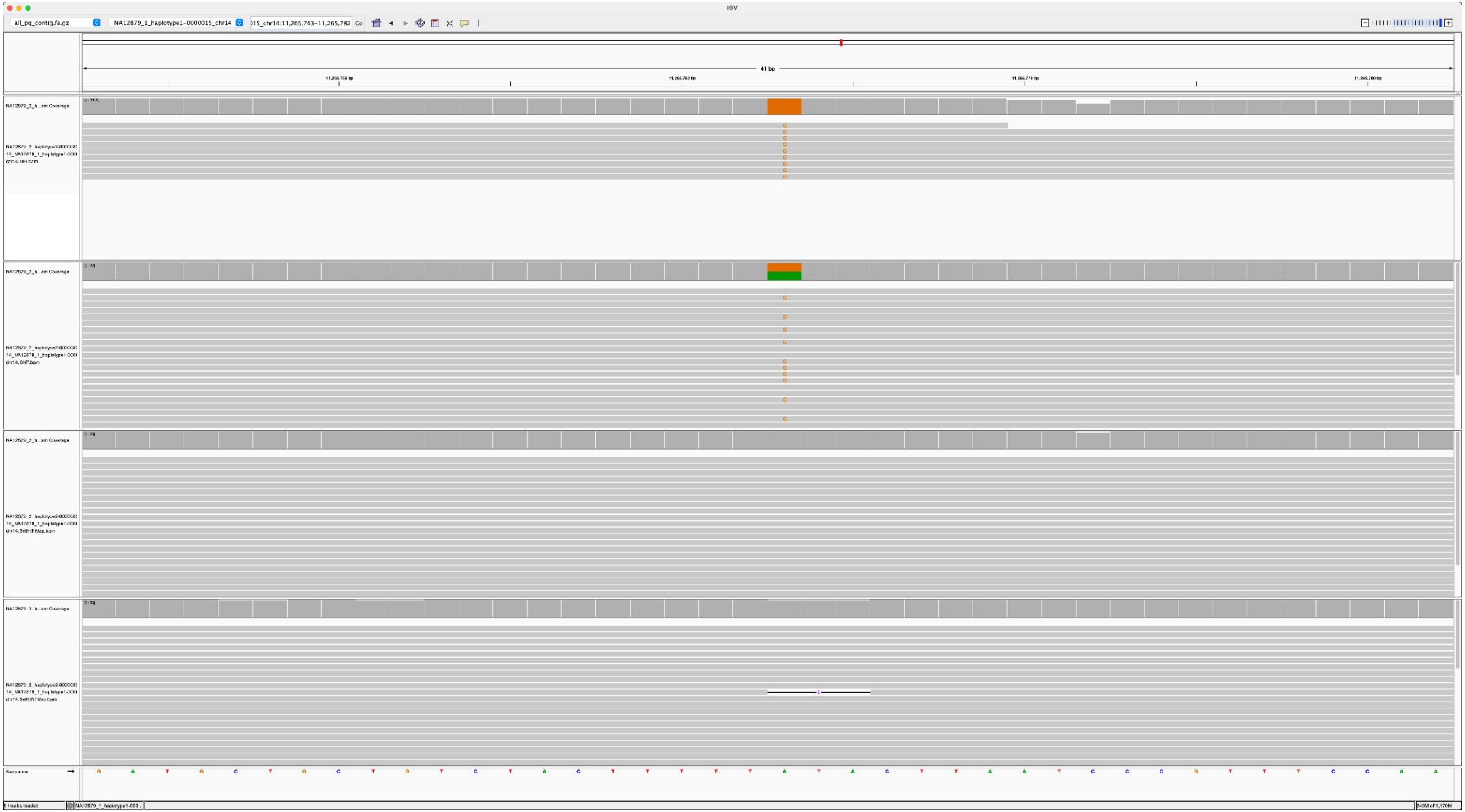

K200081_K200080_2_haplotype2-0000055_chr14_4984822_C_T

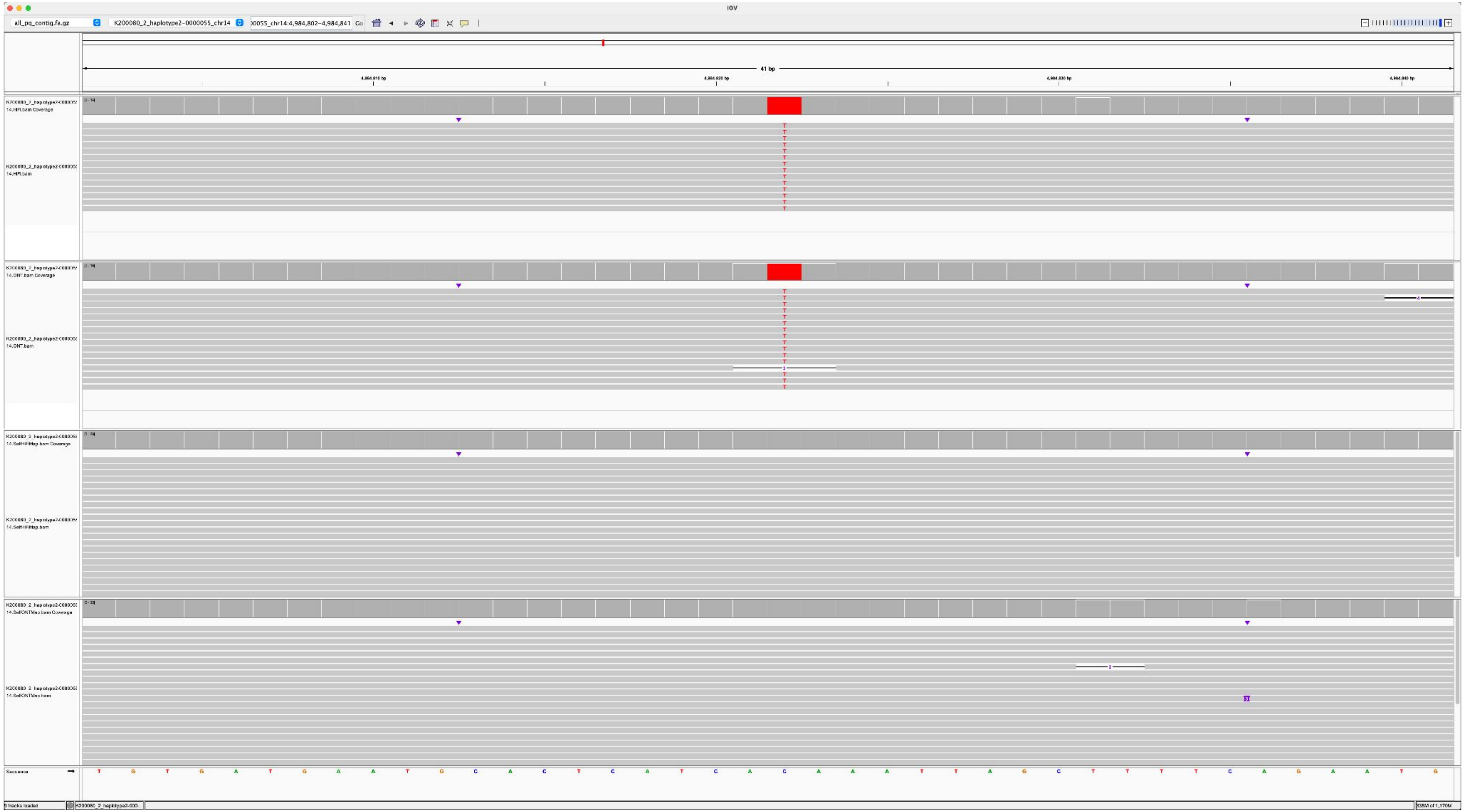

### K200081_K200080_2_haplotype2-0000071_chr15_7970241_A_C

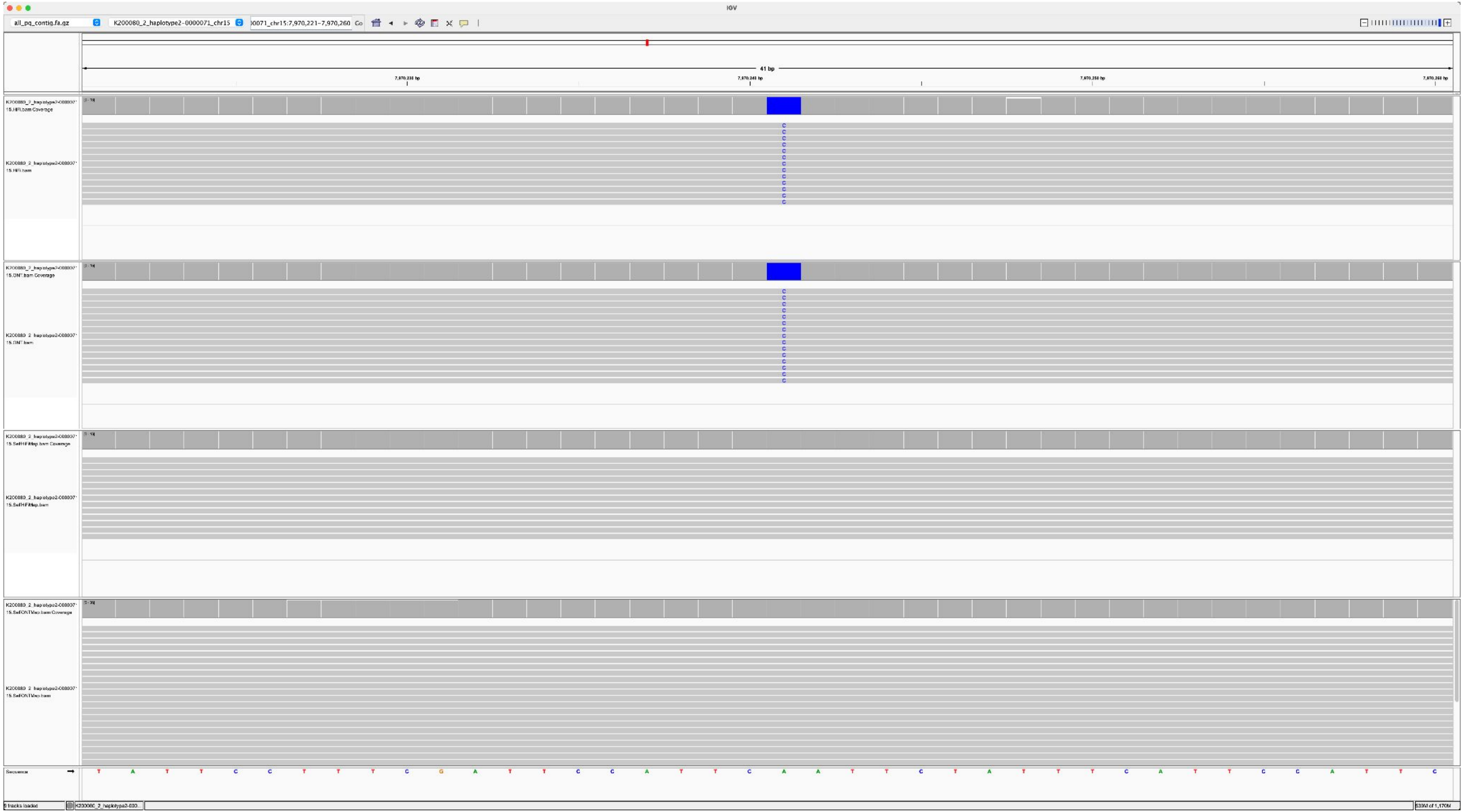

### K200081_K200080_2_haplotype2-0000071_chr15_2059154_A_C

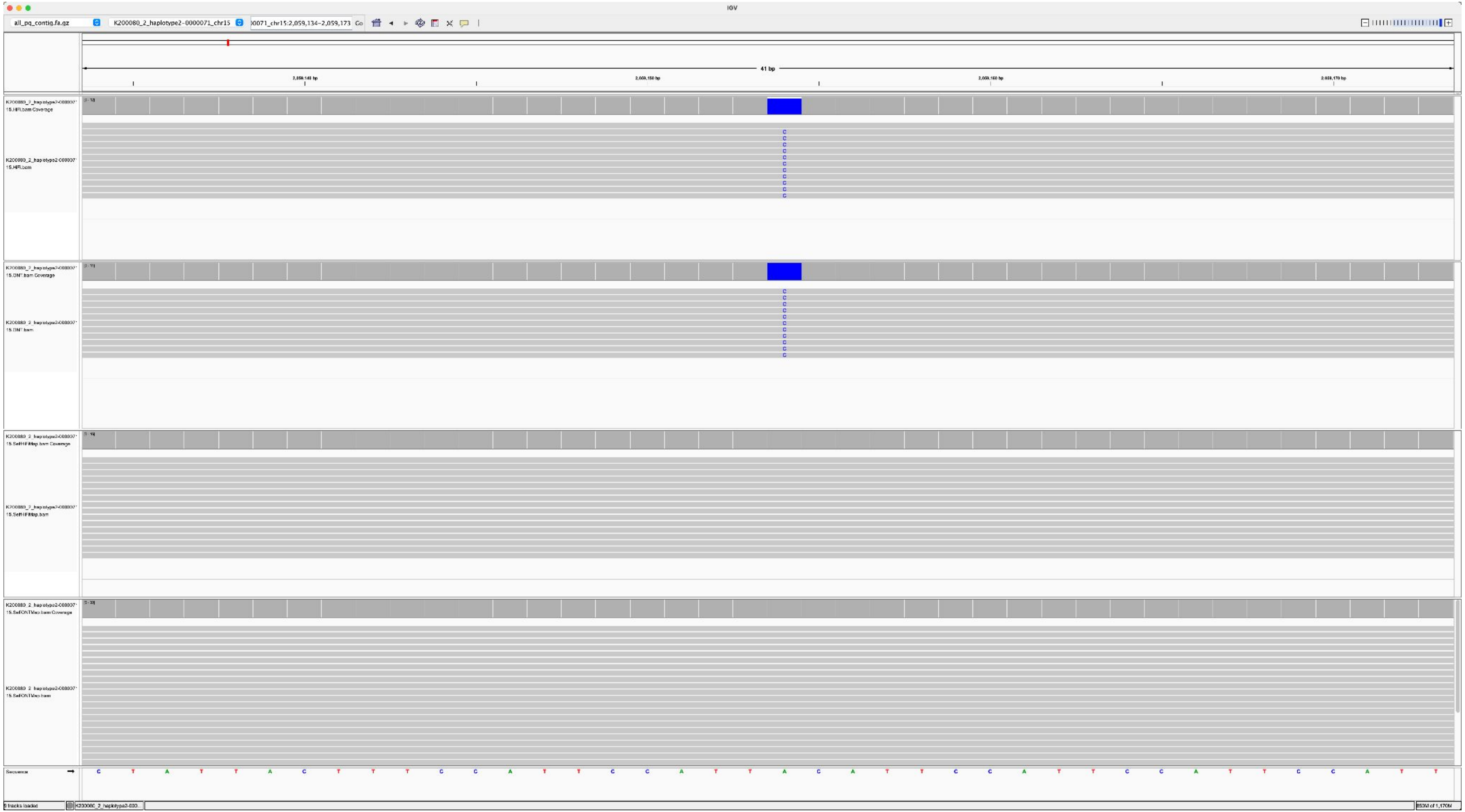

K200082_K200080_1_haplotype1-0000006_chr14_7674507_A_G

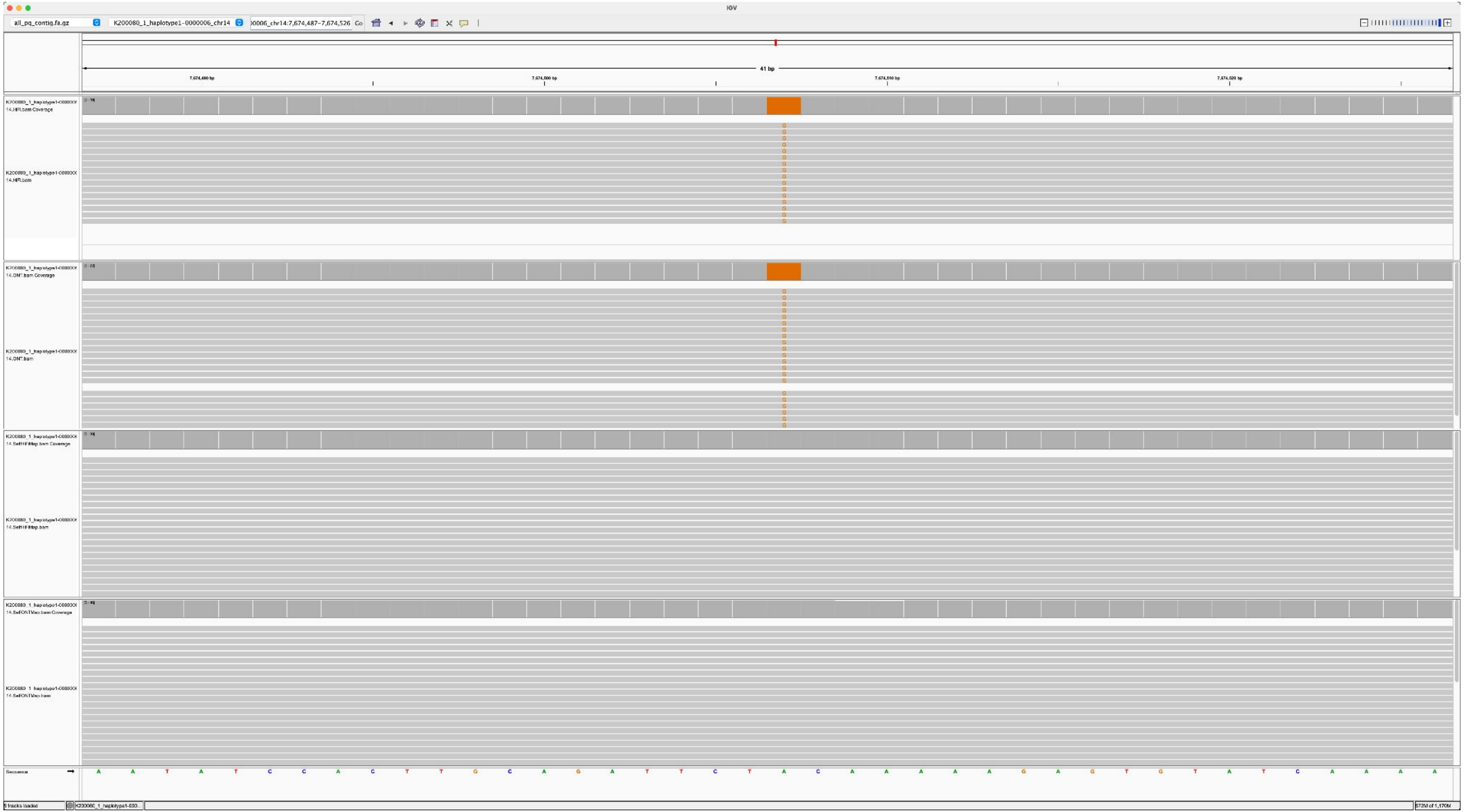

K200082_K200080_2_haplotype2-0000071_chr15_4958436_C_T

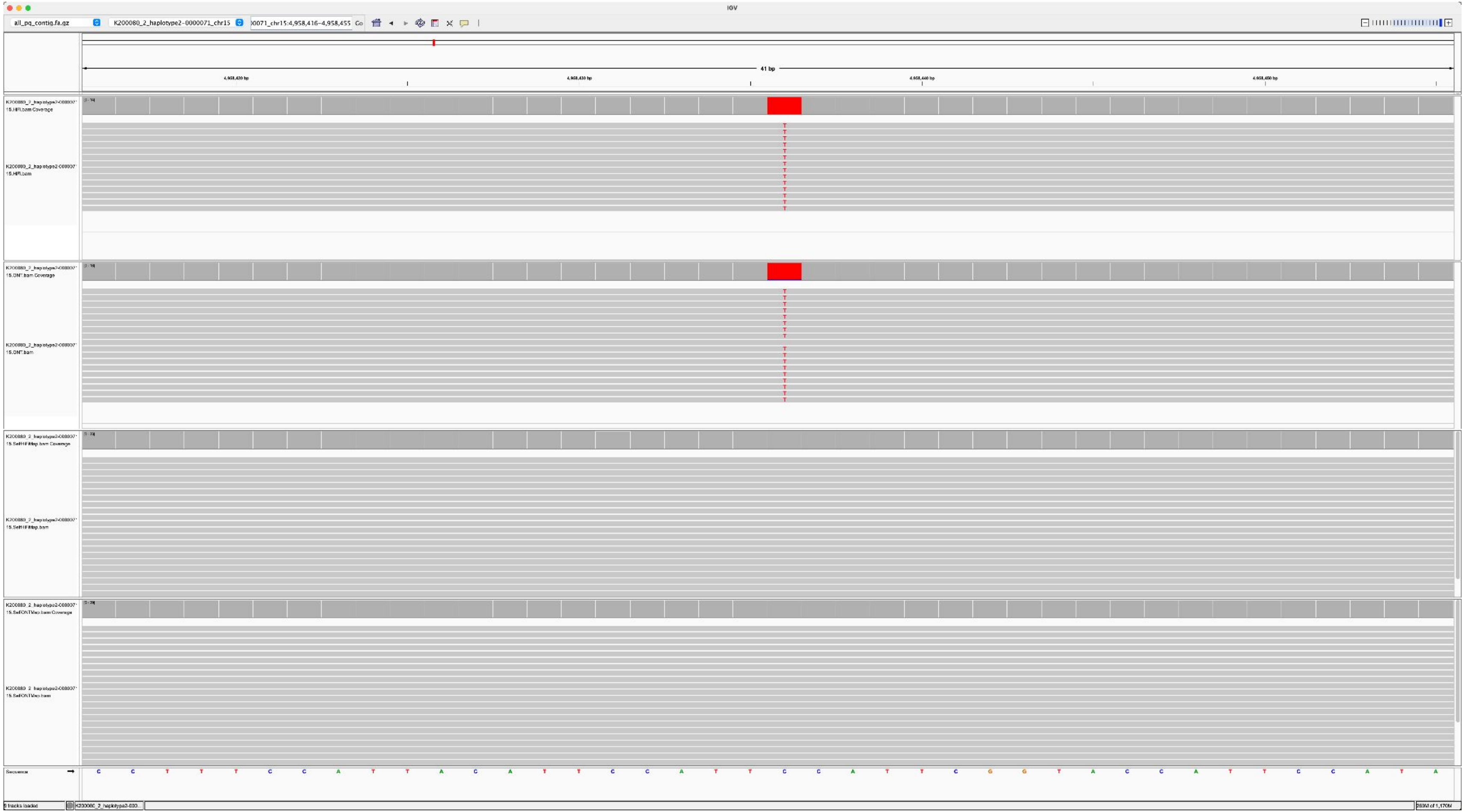

K200082_K200080_1_haplotype1-0000016_chr22_7243982_C_T

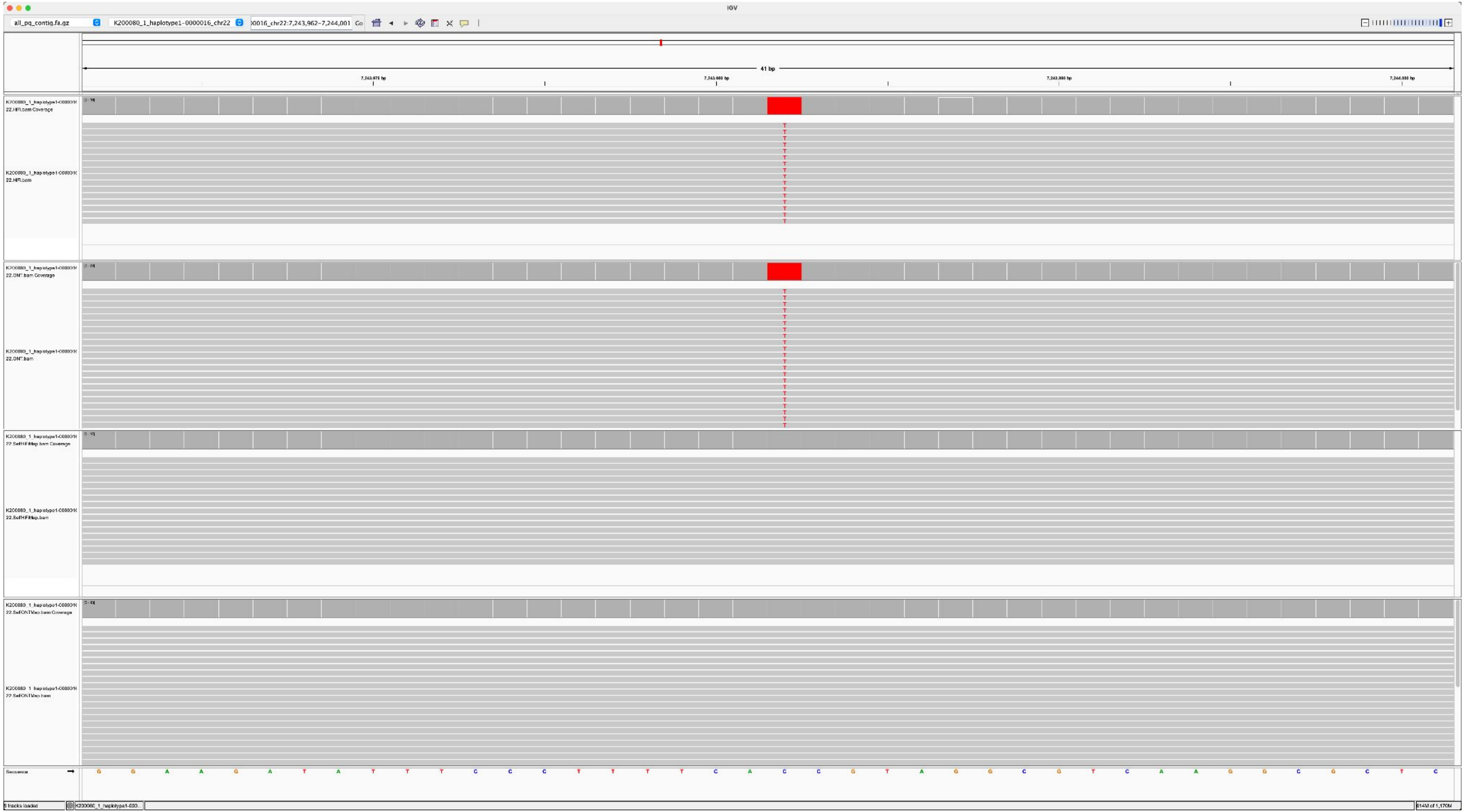

K200082_K200080_1_haplotype1-0000016_chr22_9027391_G_C

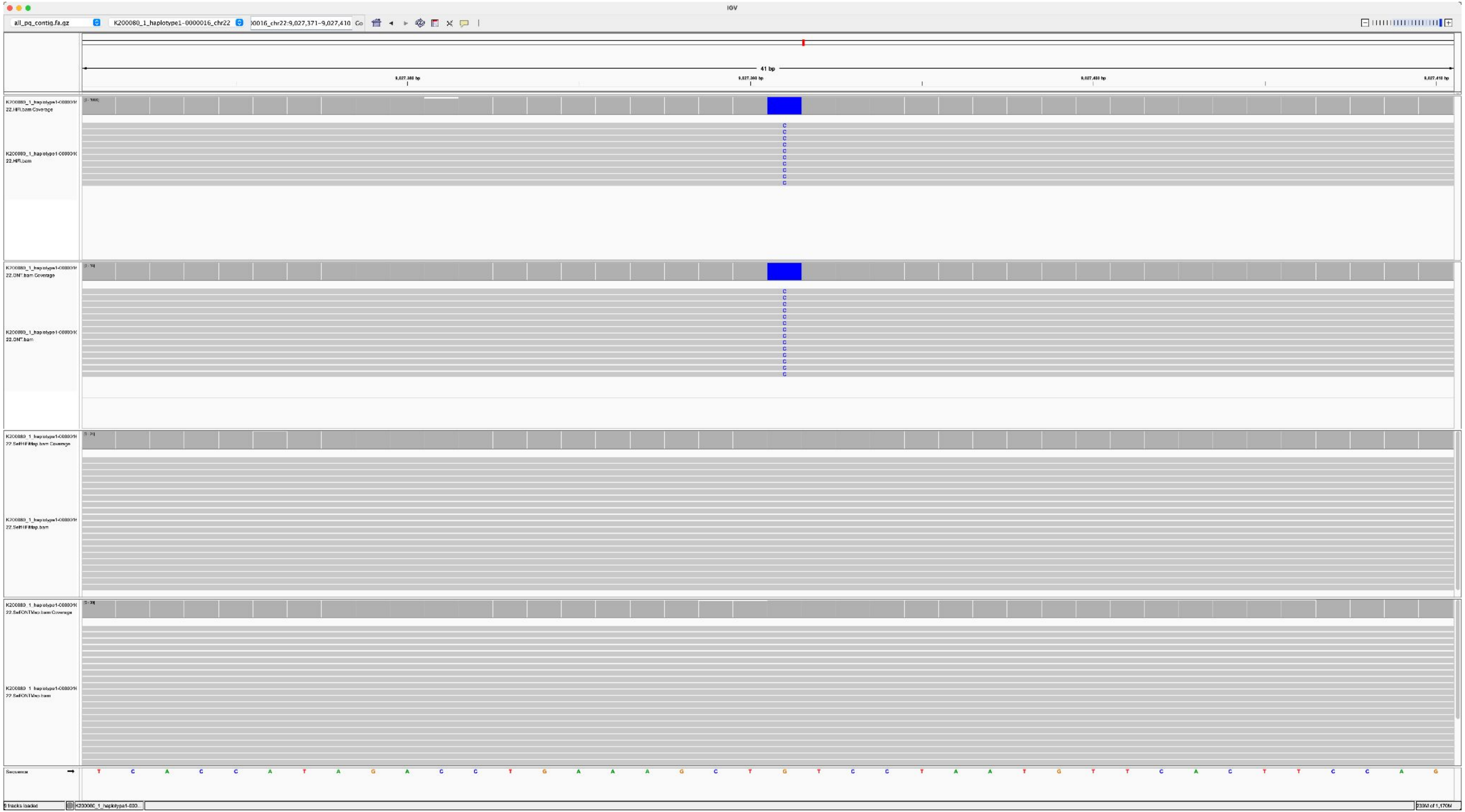

K200082_K200080_1_haplotype1-0000016_chr22_9135907_A_G

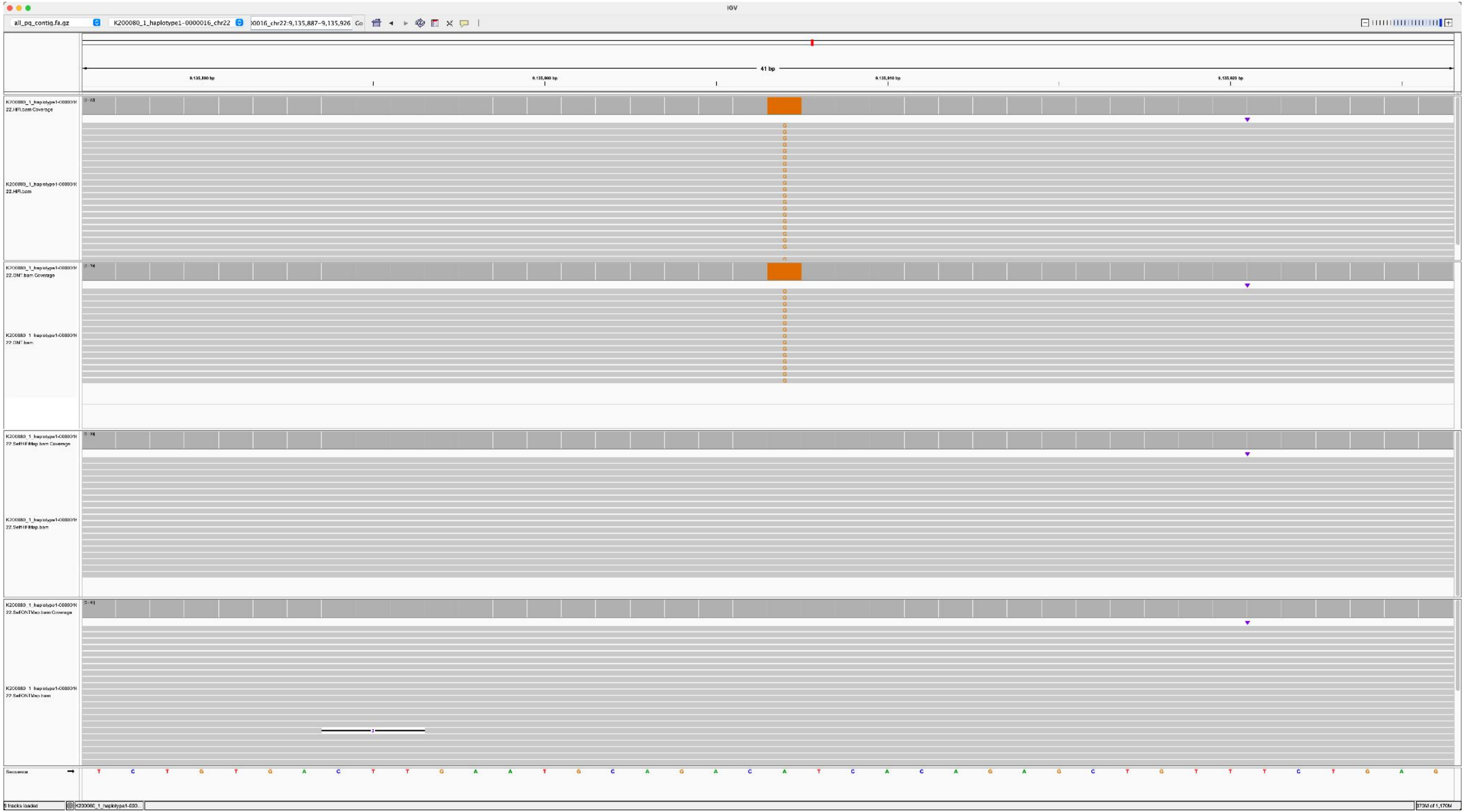

K200082_K200080_1_haplotype1-0000016_chr22_9703280_A_T

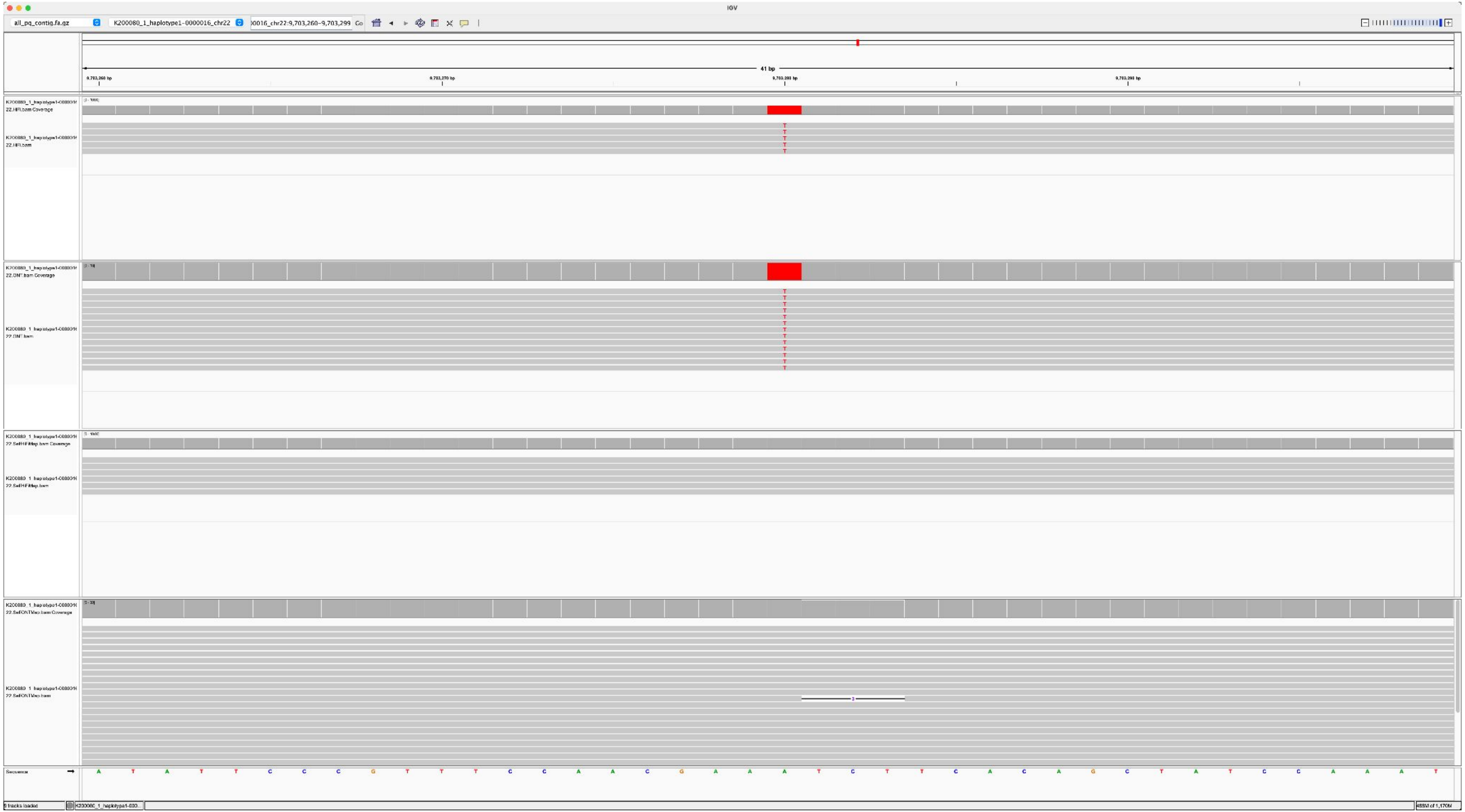

K200084_K200080_1_haplotype1-0000024_chr13_2804996_C_T

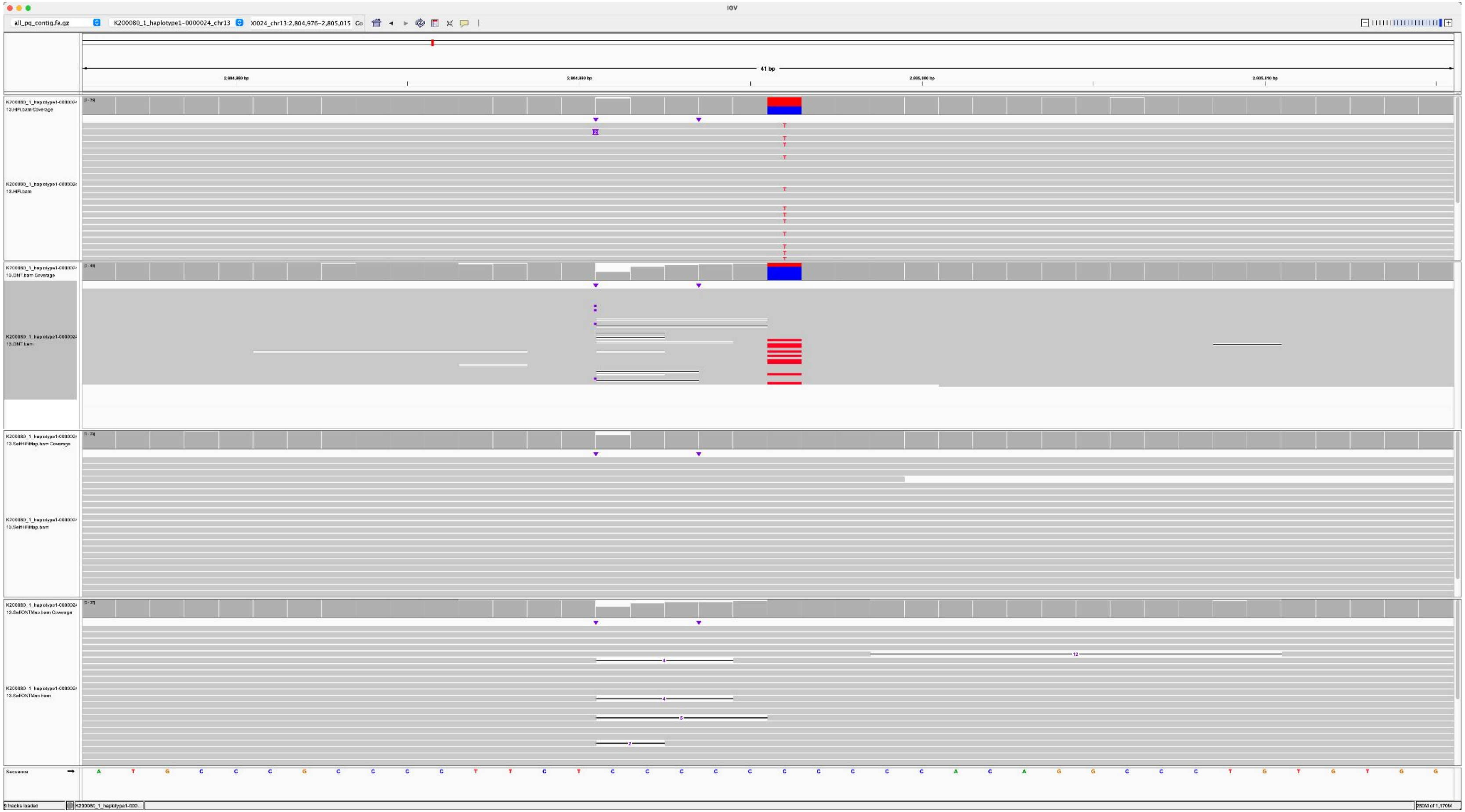

K200084_K200080_1_haplotype1-0000026_chr15_4730006_C_T

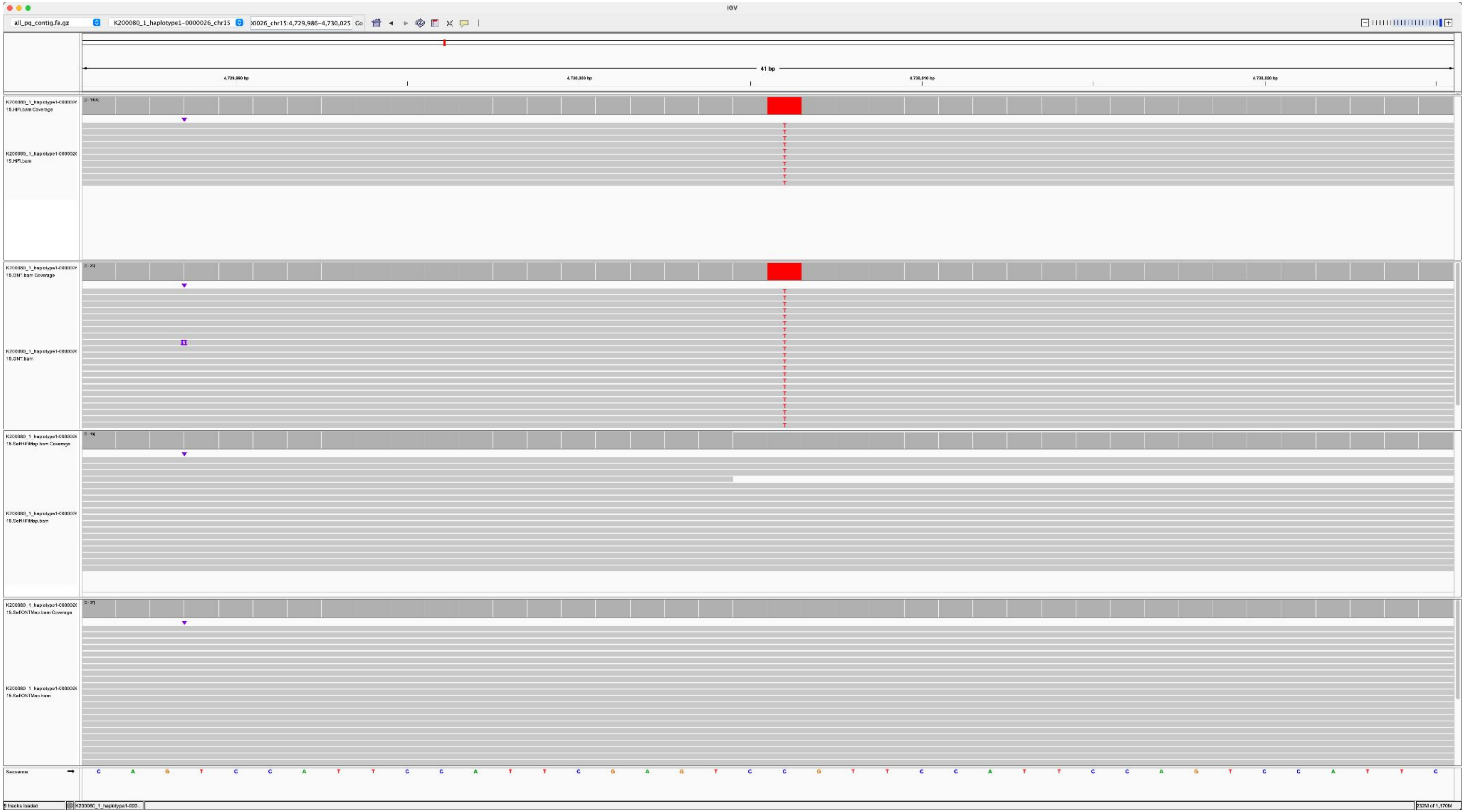

K200084_K200080_1_haplotype1-0000026_chr15_9313874_A_C

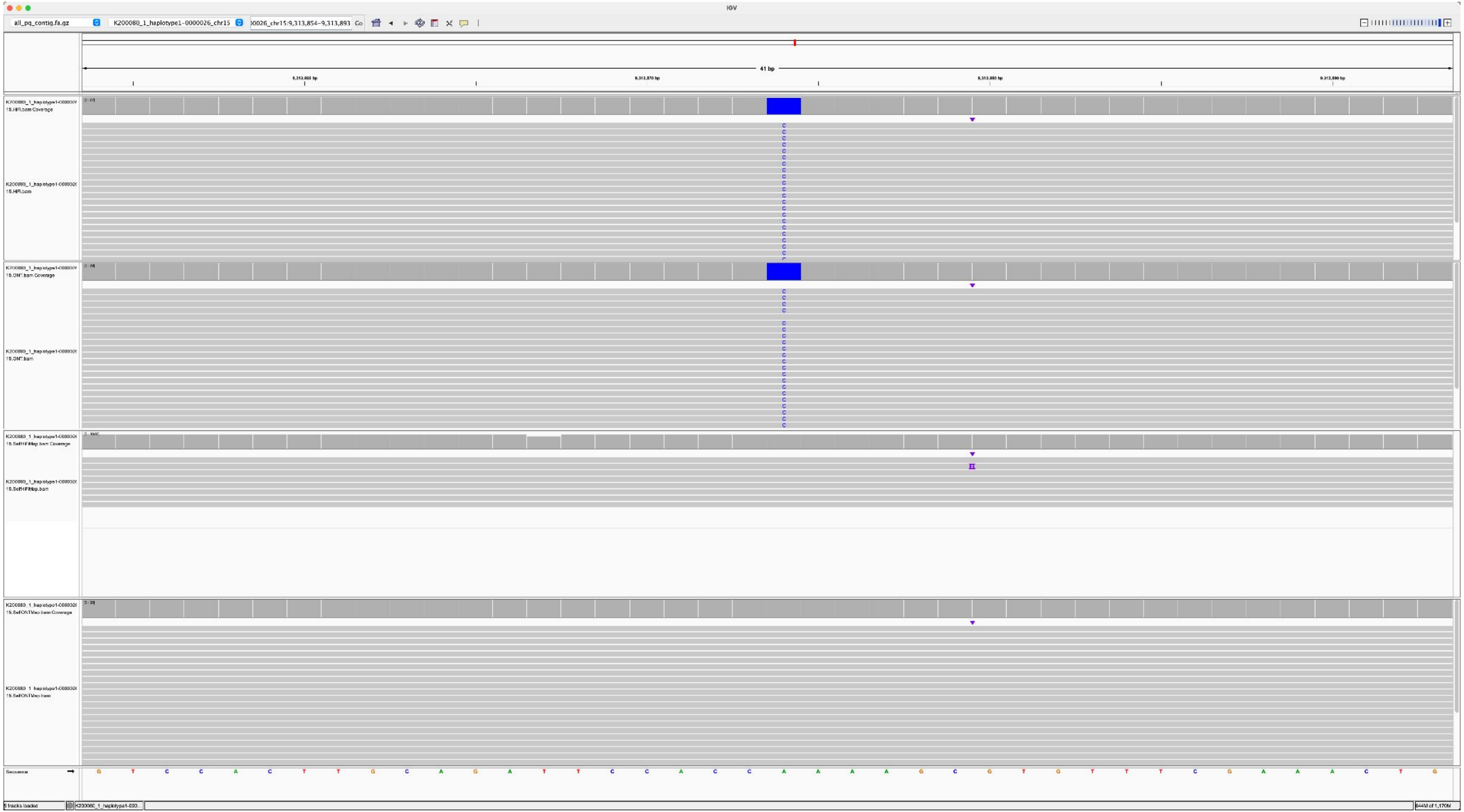

### K200084_K200080_2_haplotype2-0000063_chr22_4755532_C_T

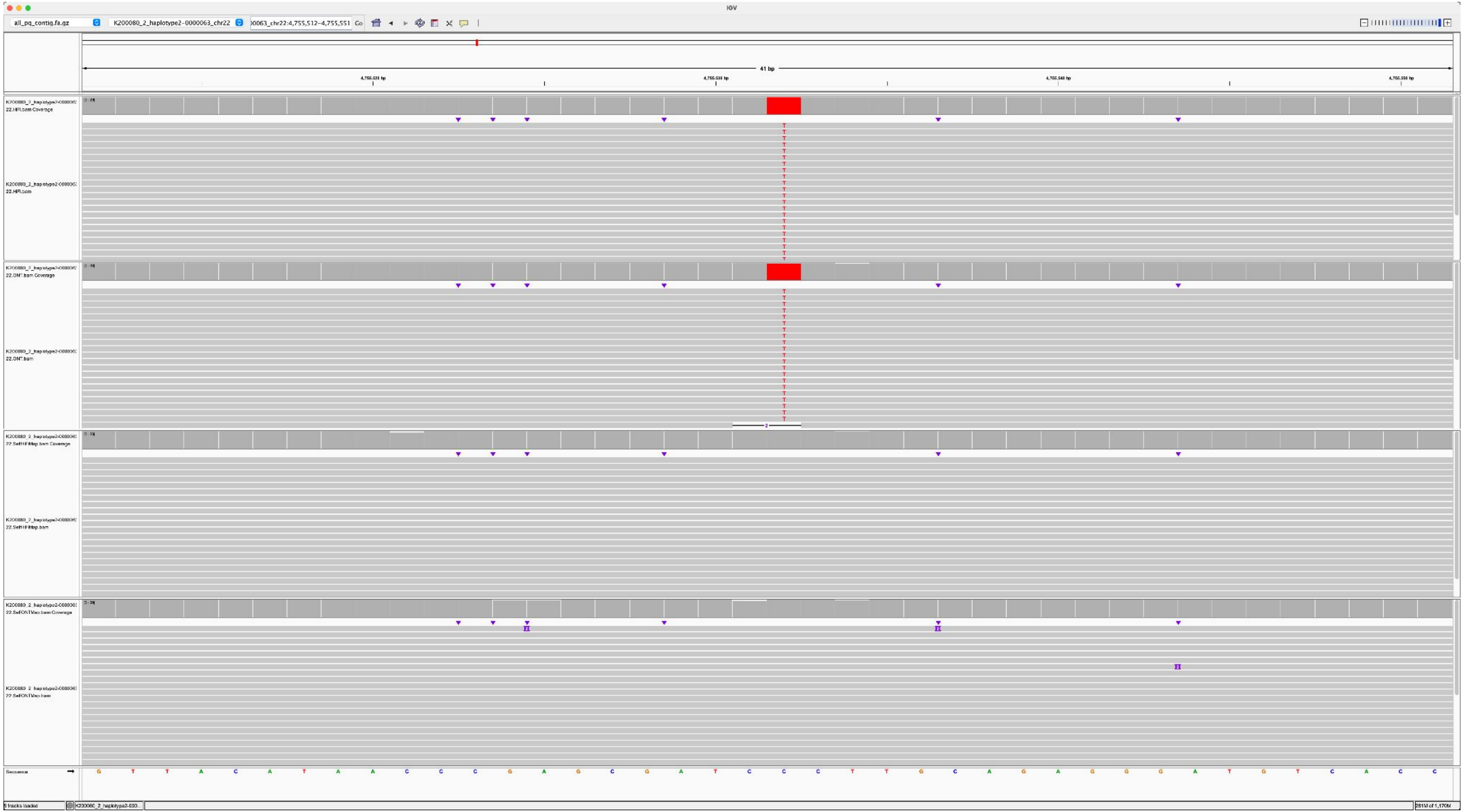

K200084_K200080_2_haplotype2-0000063_chr22_6305935_G_C

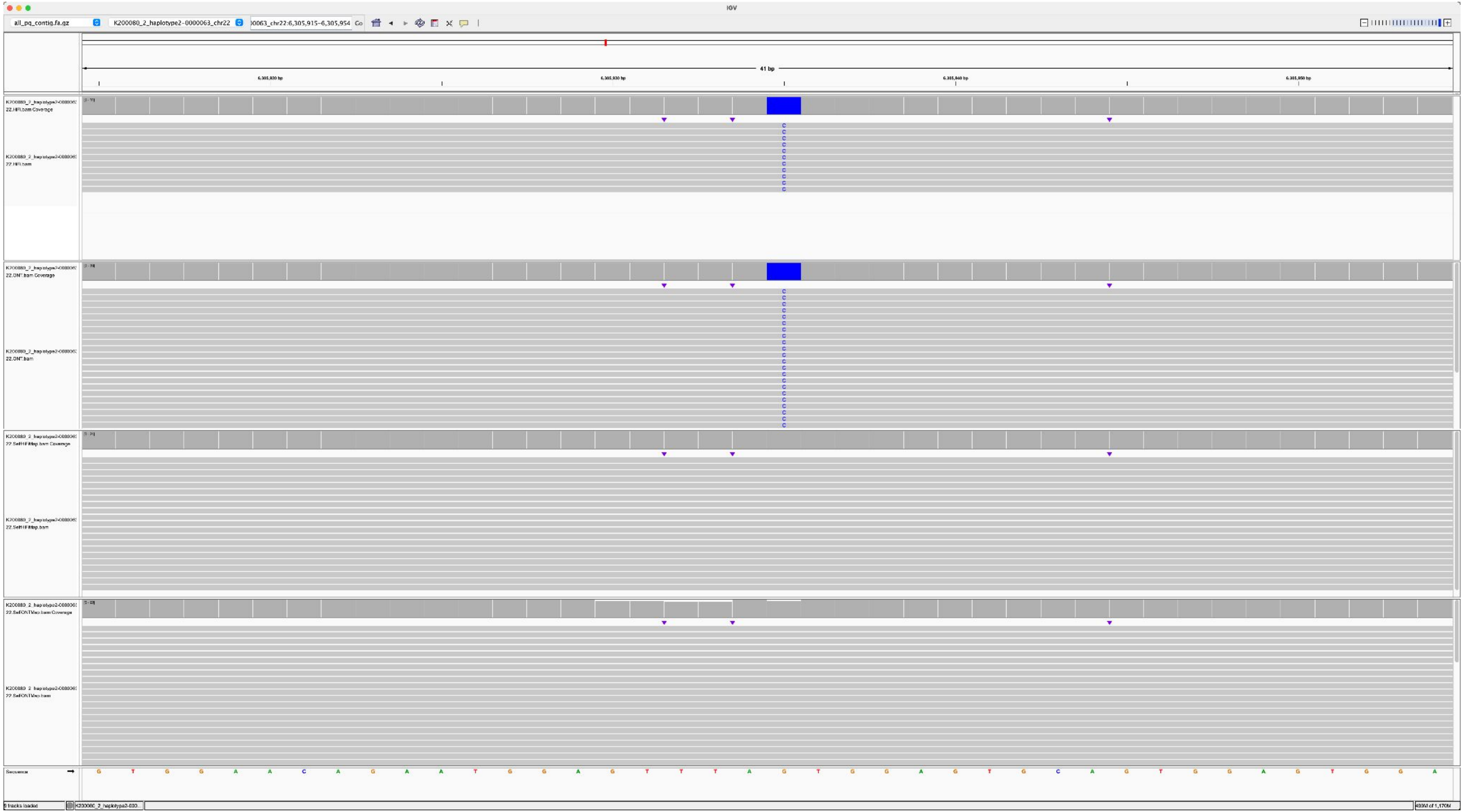

K200084_K200080_2_haplotype2-0000063_chr22_10560366_G_C

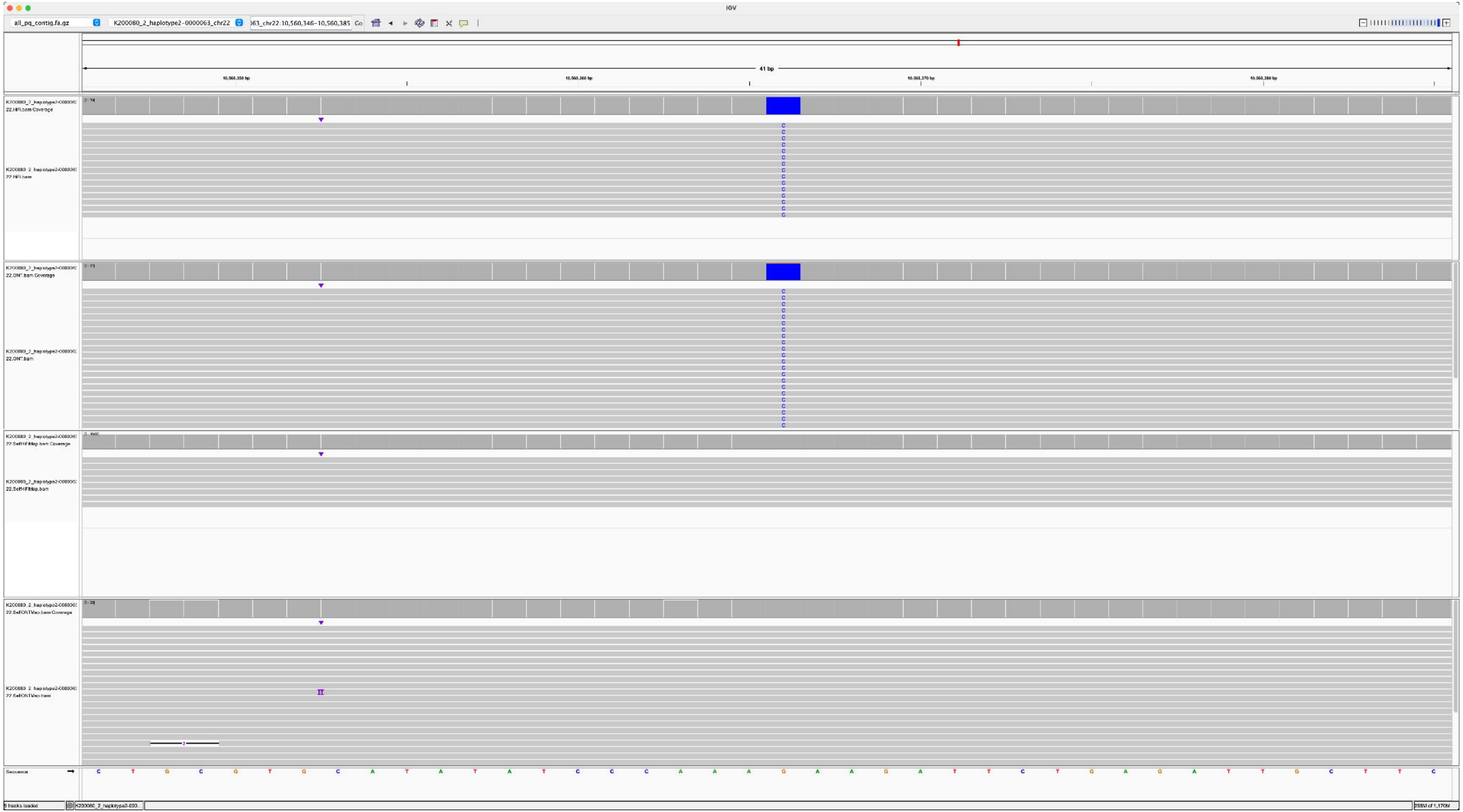

K200084_K200080_2_haplotype2-0000063_chr22_3155000_C_G

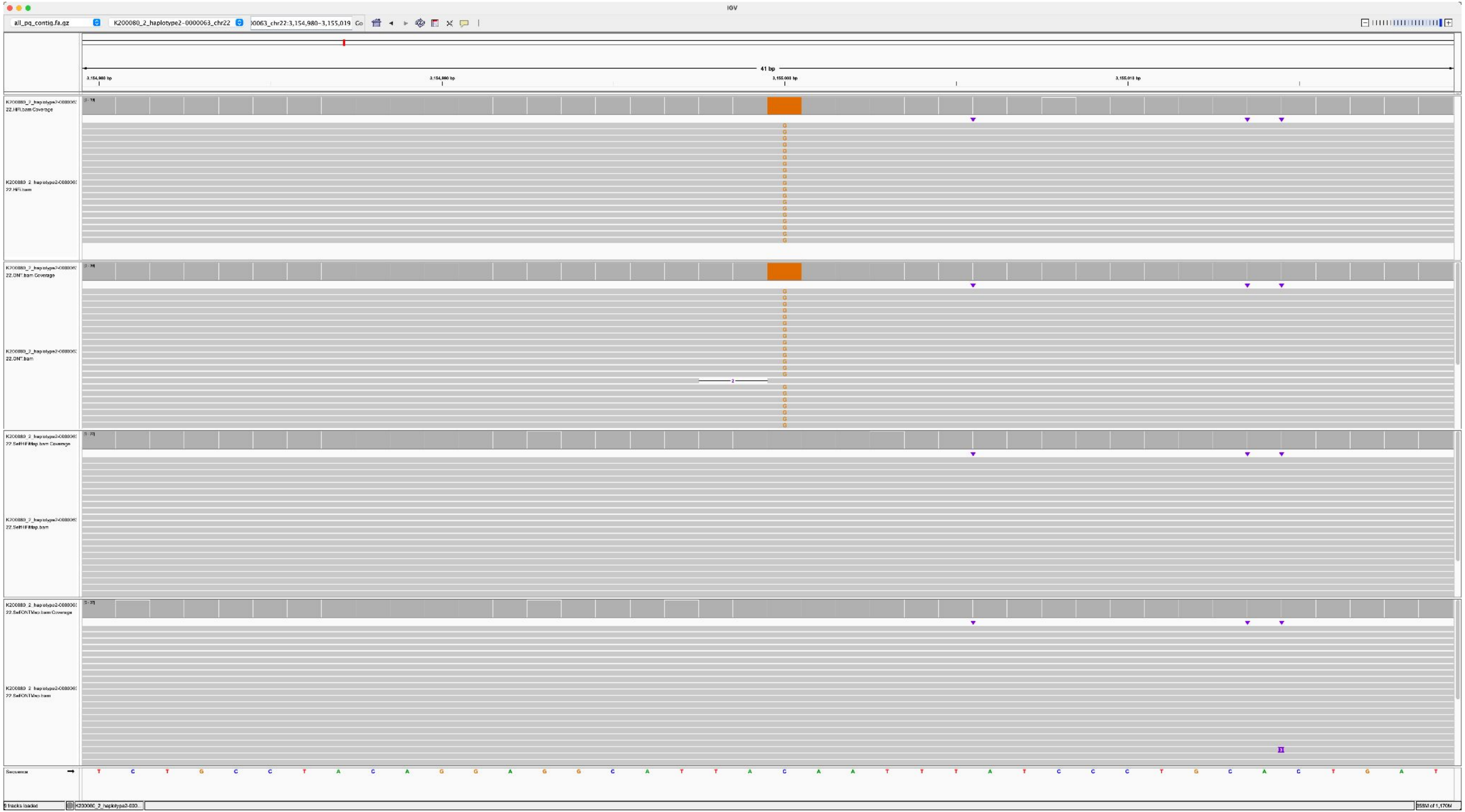

### K200085_K200080_2_haplotype2-0000075_chr13_4998628_A_C

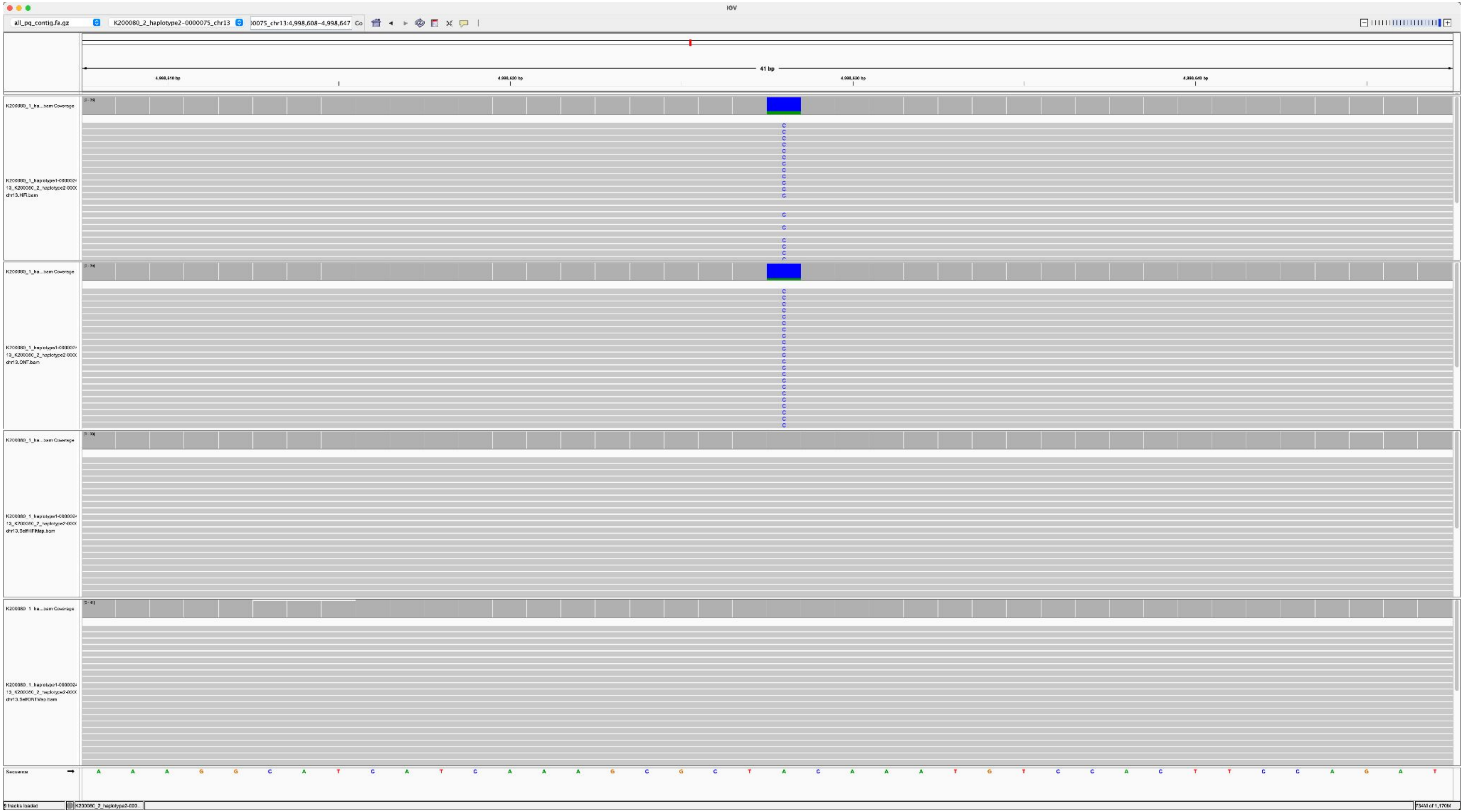

K200085_K200080_2_haplotype2-0000075_chr13_5922694_G_A

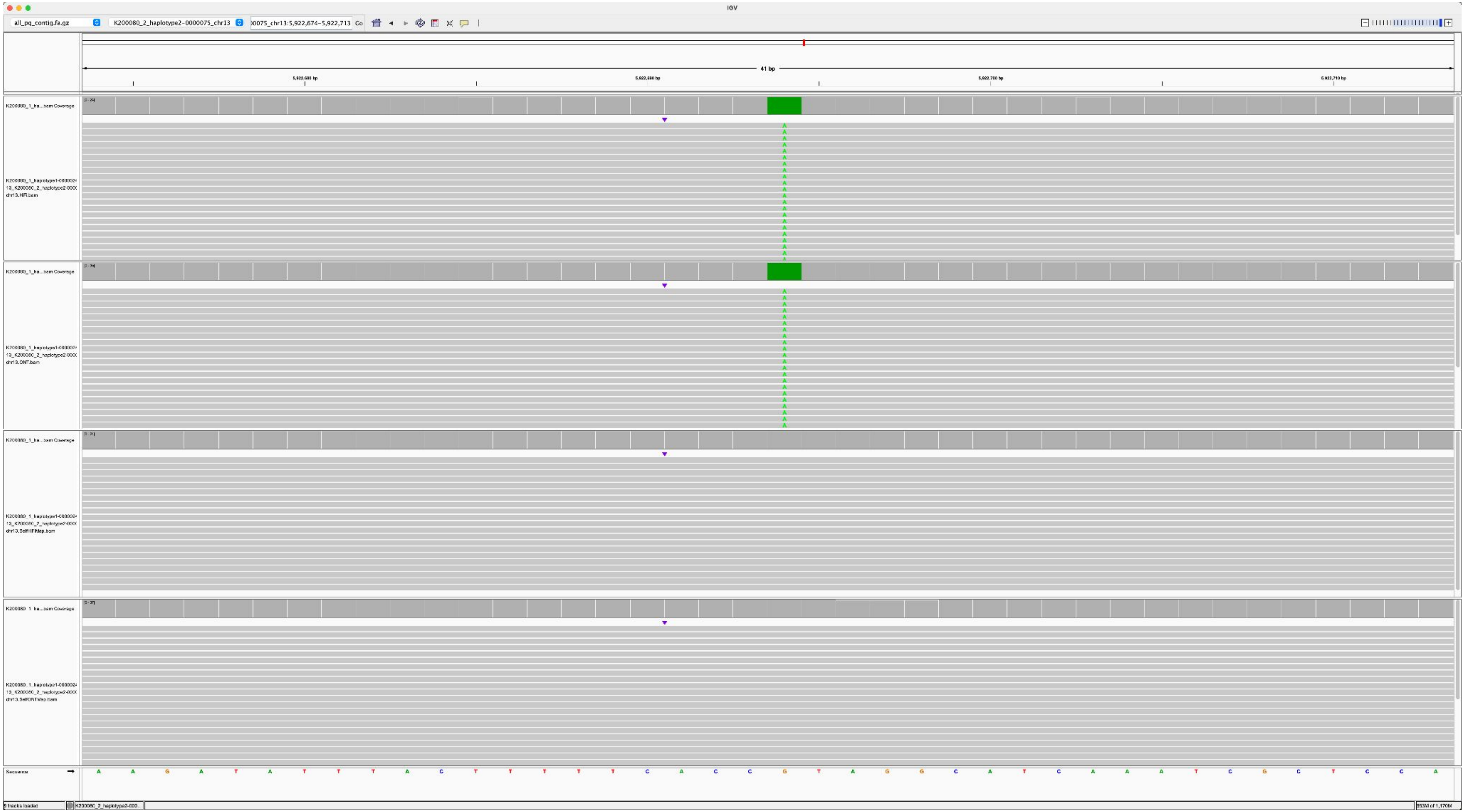

K200085_K200080_2_haplotype2-0000075_chr13_4497595_A_G

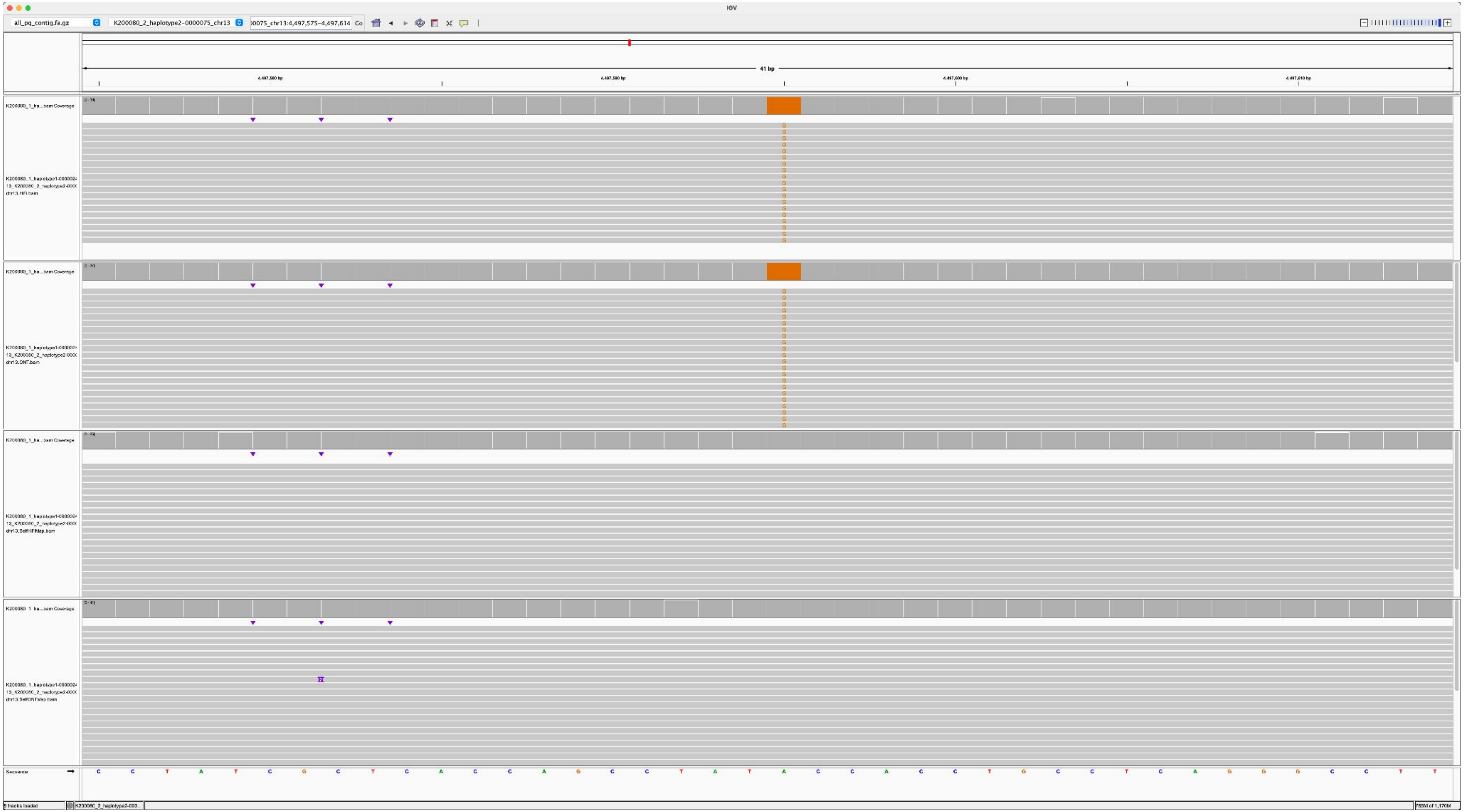

K200085_K200080_1_haplotype1-0000026_chr15_2926150_A_G

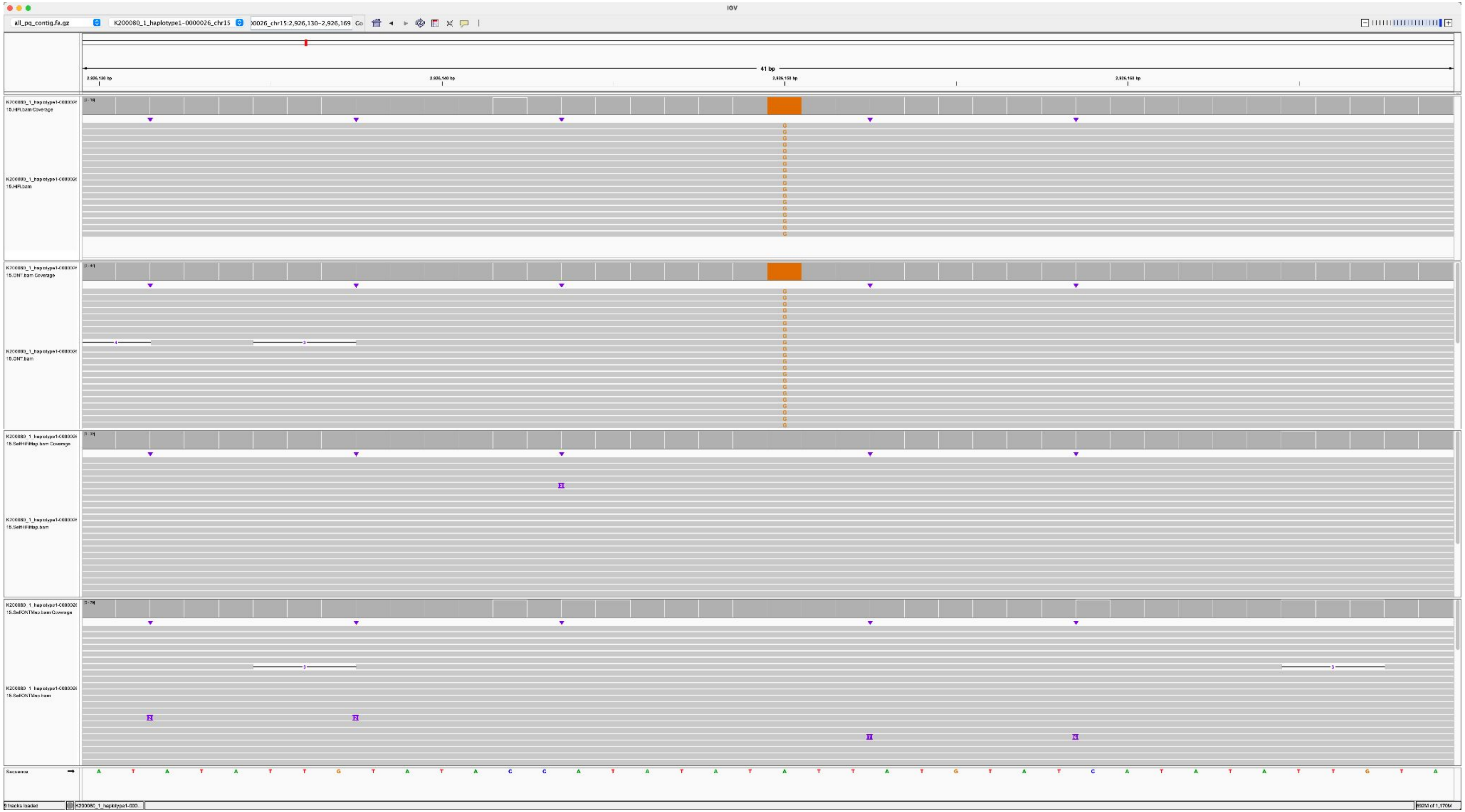

K200085_K200080_2_haplotype2-0000063_chr22_10560366_G_C

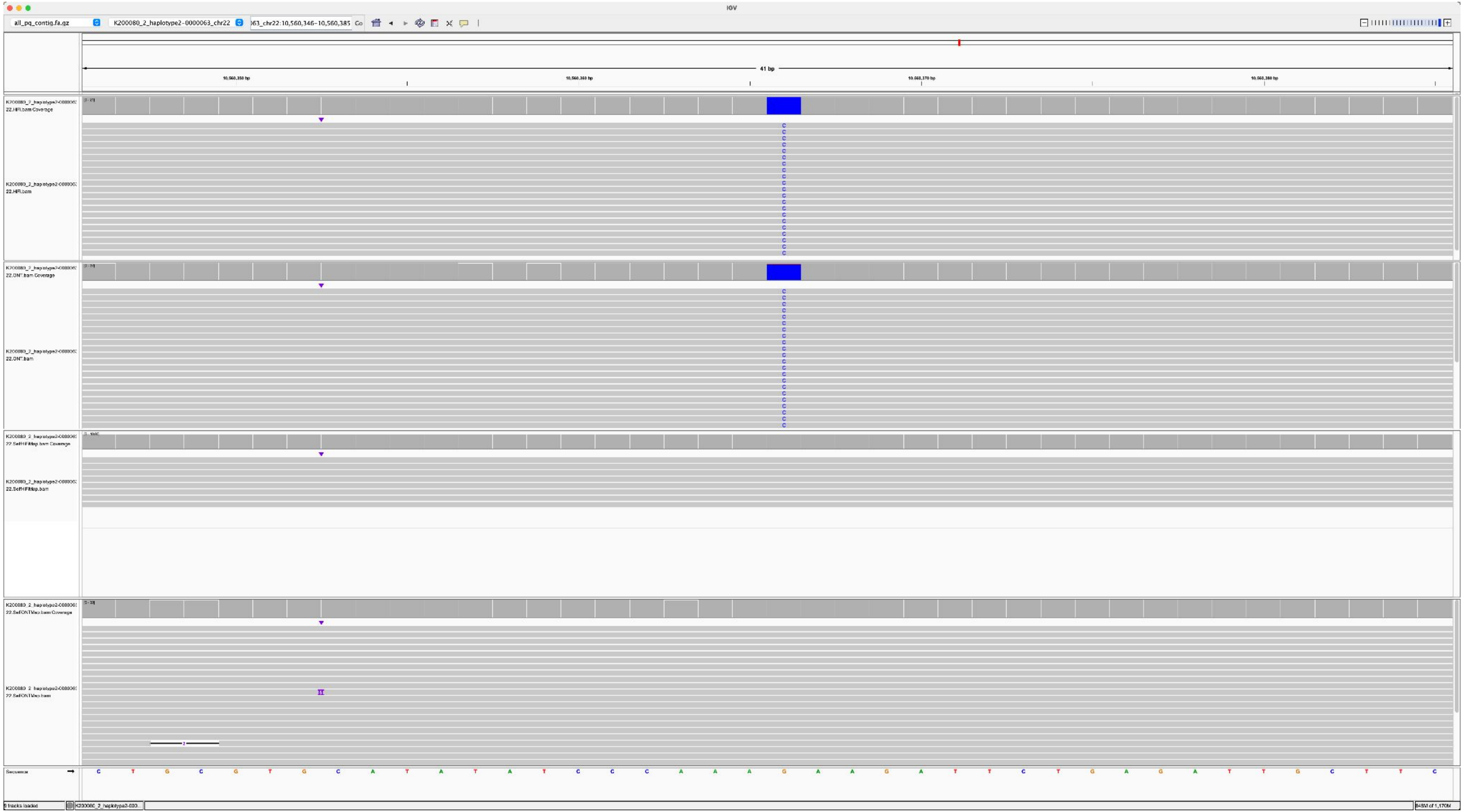

K200085_K200080_2_haplotype2-0000063_chr22_3556890_G_A

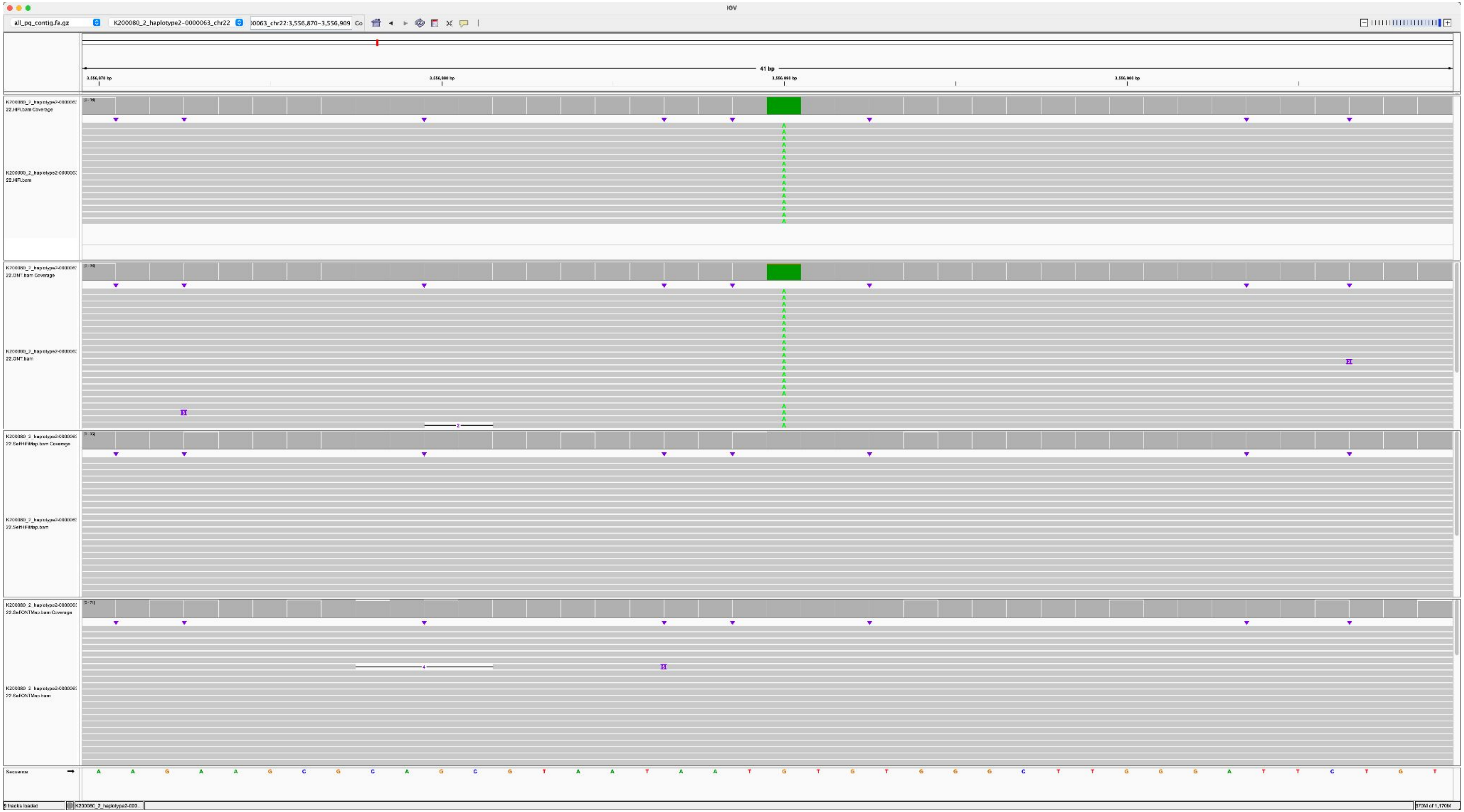

### K200087_K200080_1_haplotype1-0000006_chr14_7032154_G_C

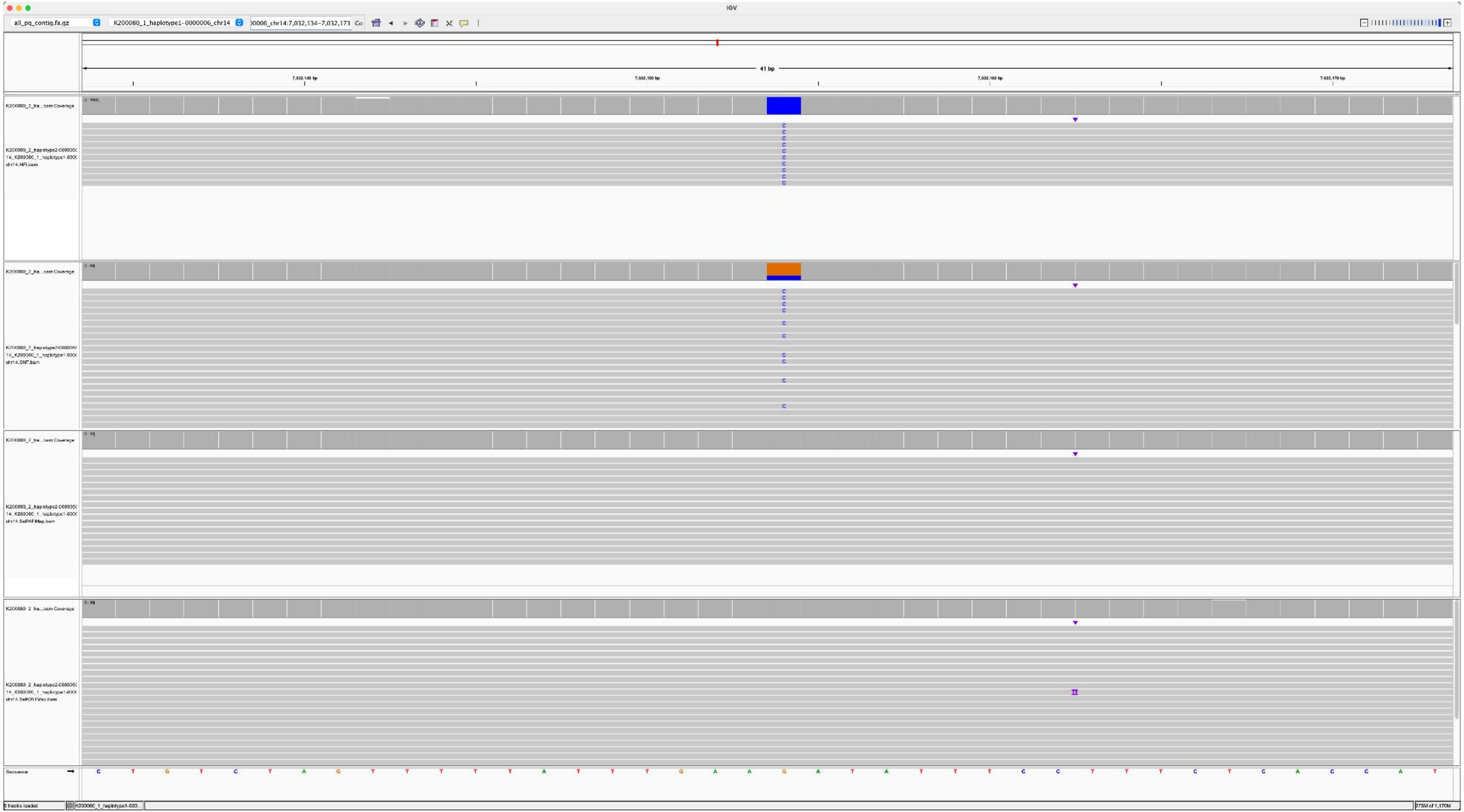

K200087_K200080_1_haplotype1-0000006_chr14_7441600_C_G

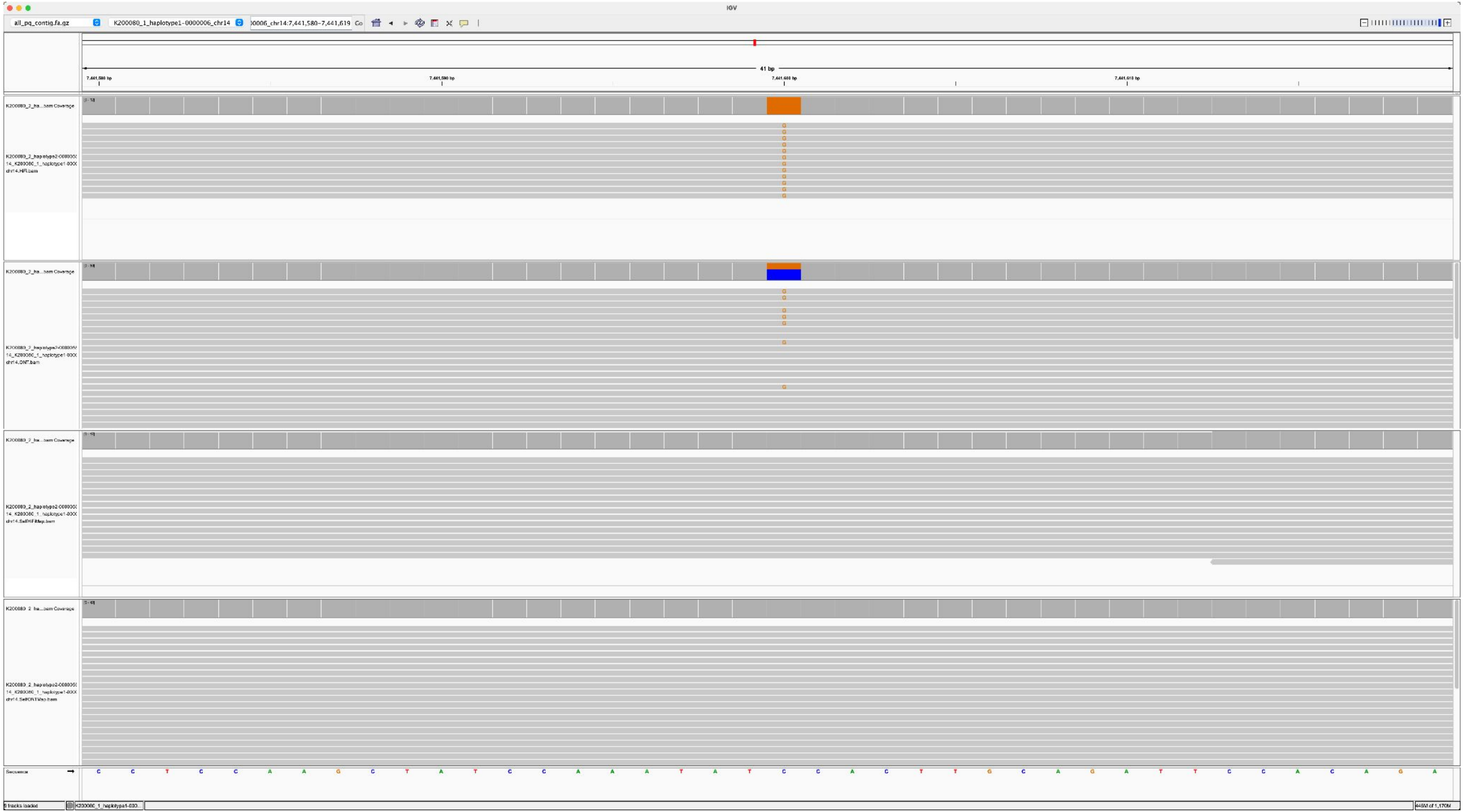

K200087_K200080_1_haplotype1-0000006_chr14_7449611_T_G

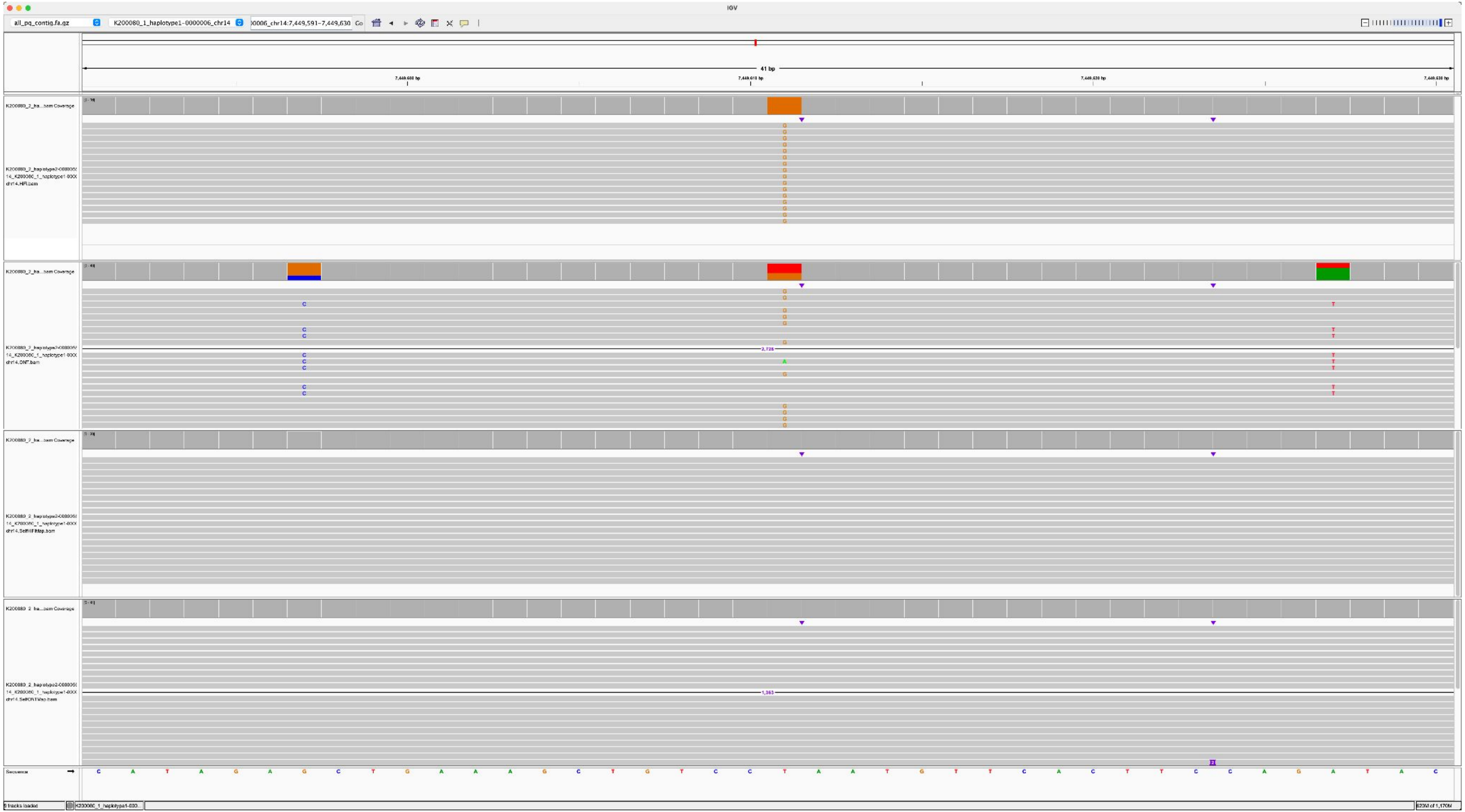

### K200087_K200080_1_haplotype1-0000006_chr14_7529401_A_T

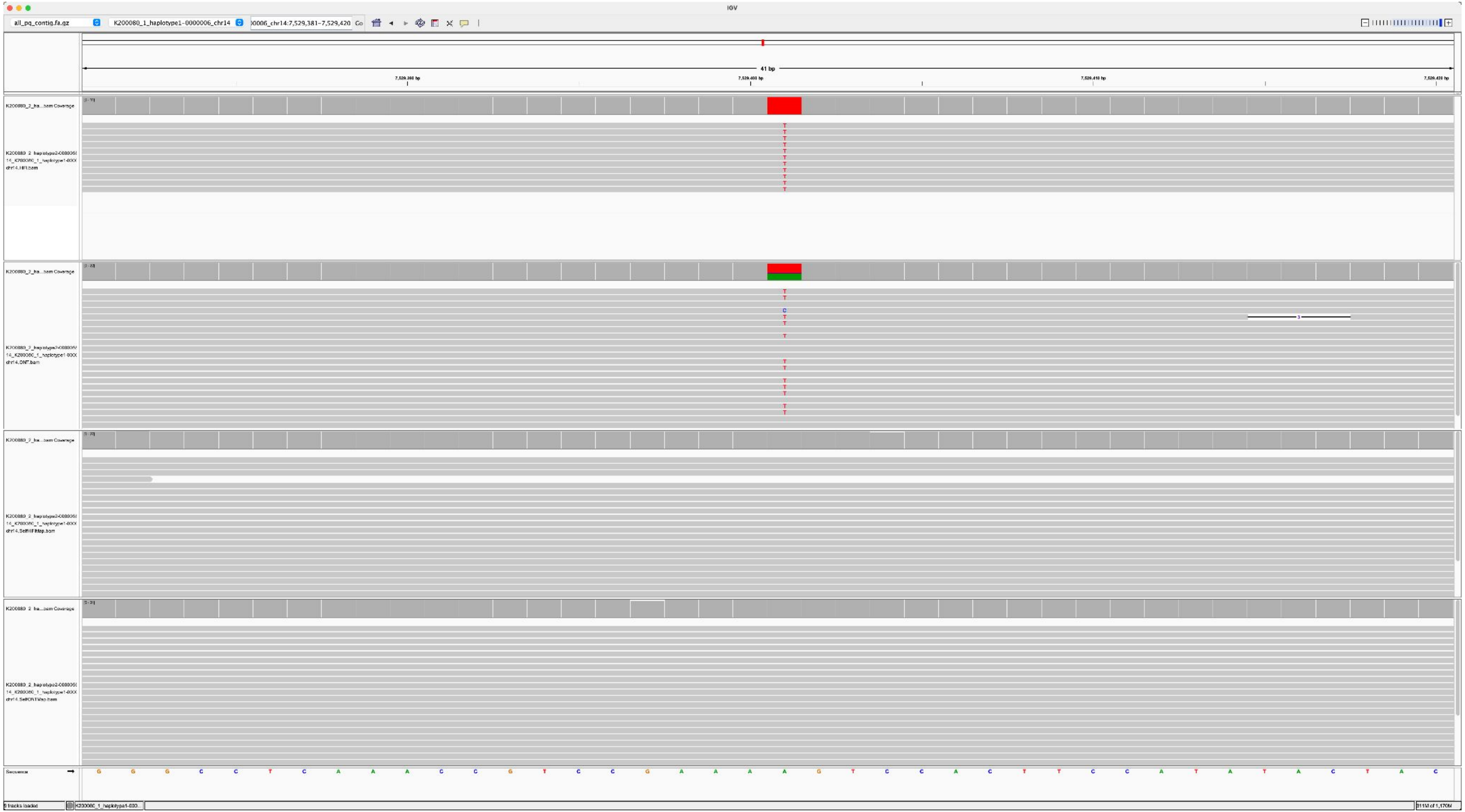

### K200087_K200080_1_haplotype1-0000006_chr14_3578074_A_G

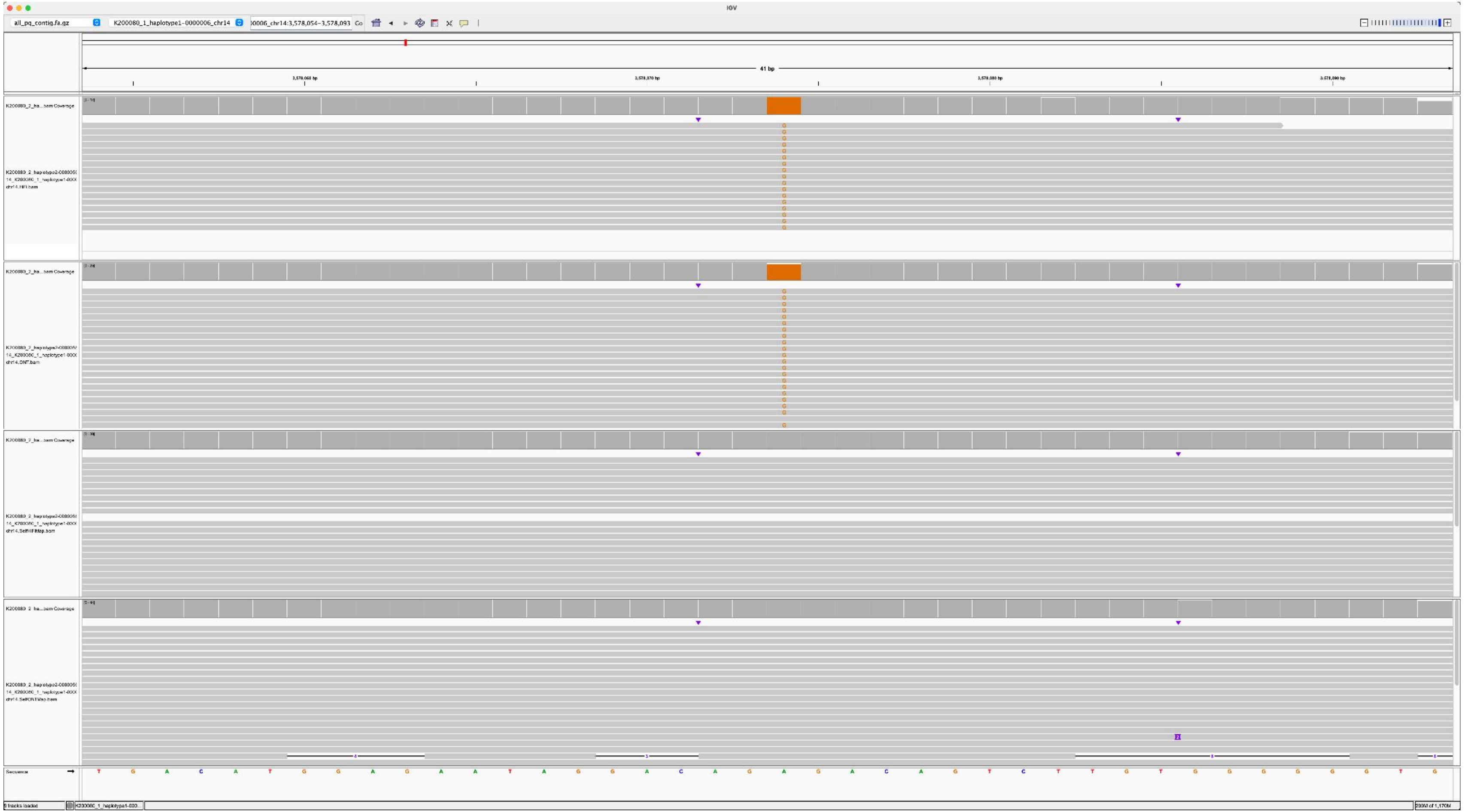

### Structural variants

### NA12878_2_haplotype2-0000041_chr14, NA12884_1_haplotype1-0000003_chr14, 751-DEL, HSat1A

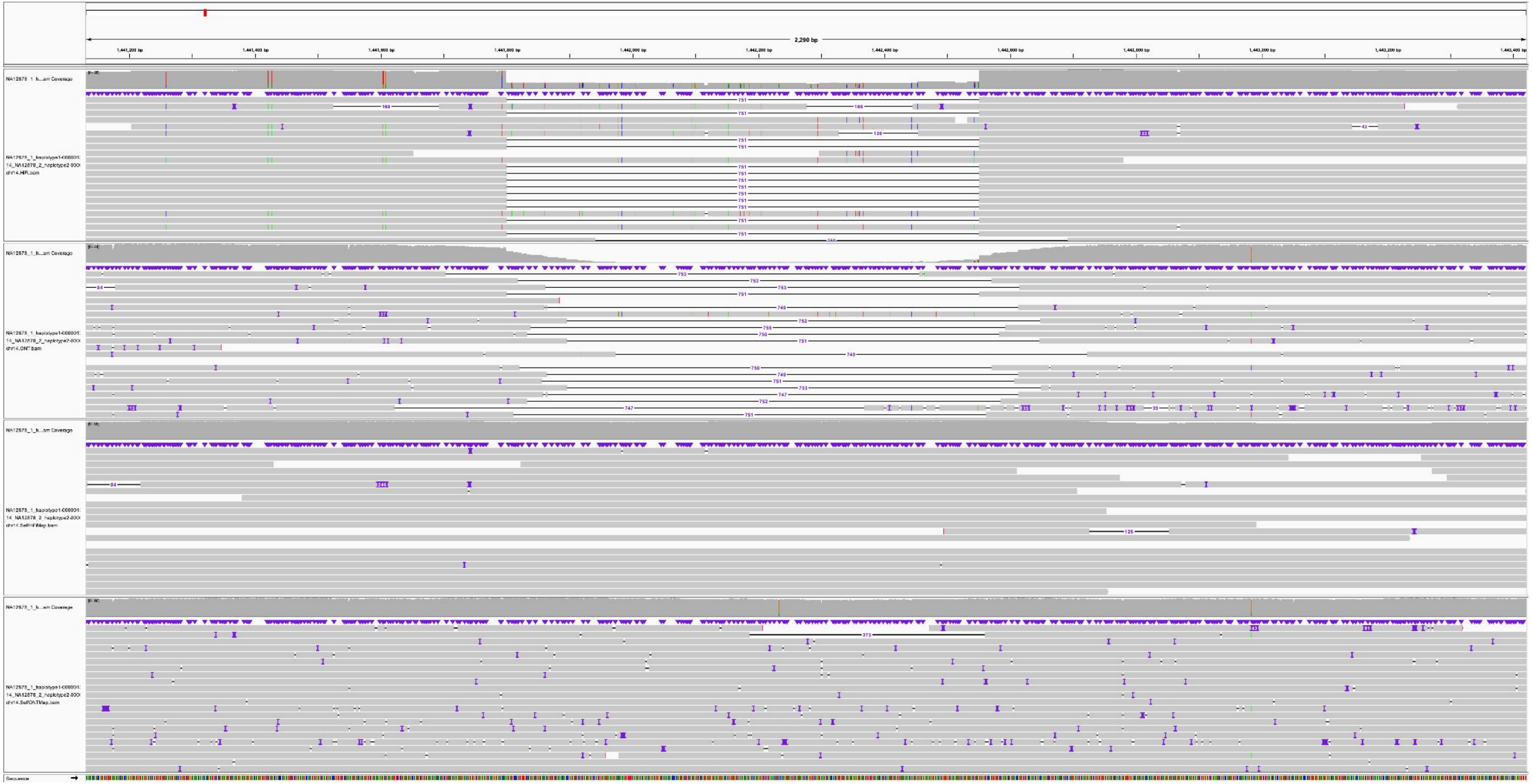

NA12878_1_haplotype1-0000009_chr21, NA12886_1_haplotype1-0000005_chr21, 680-INS, aSat_HOR

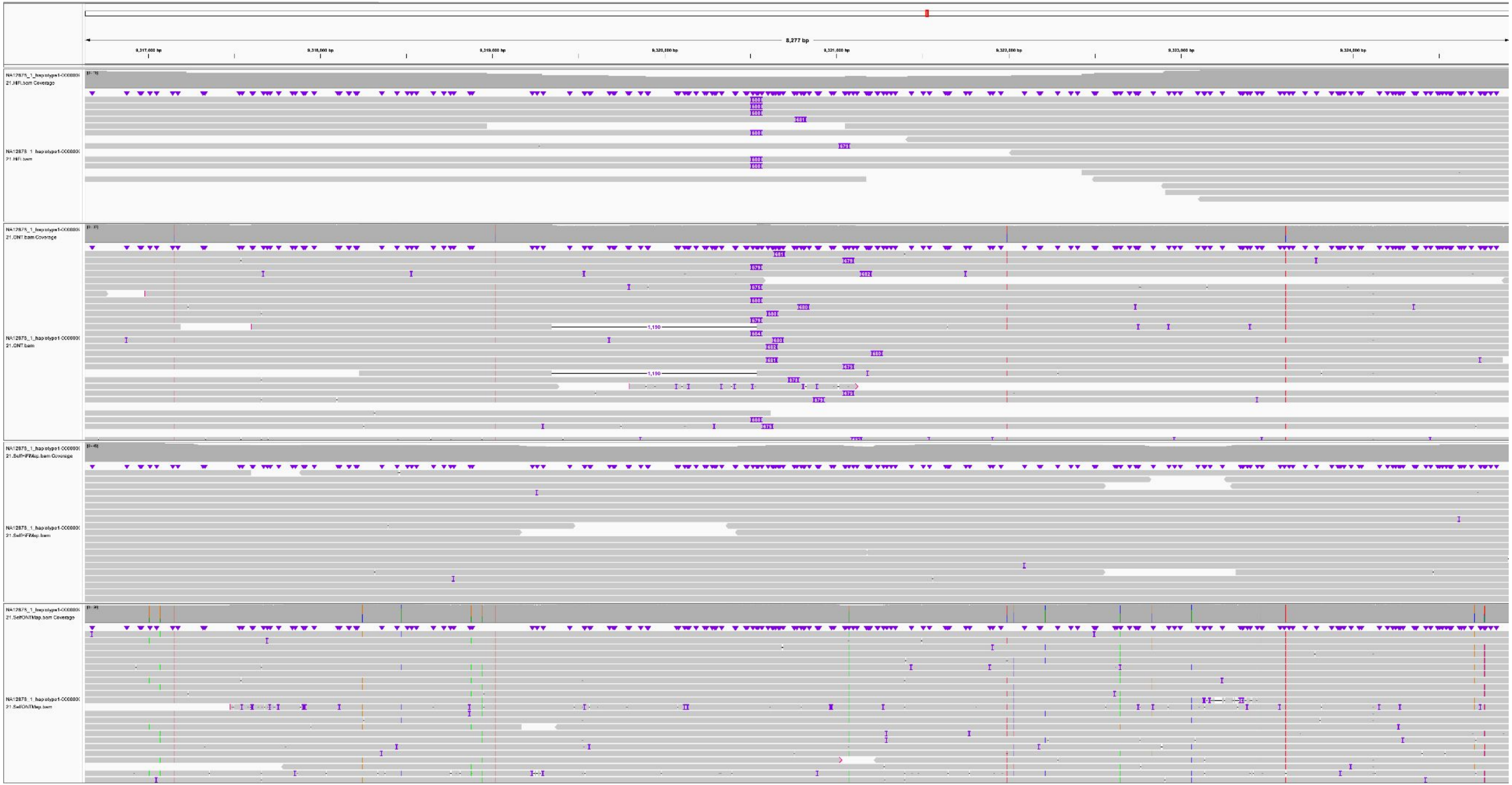

NA12877_1_haplotype1-0000014_chr15, NA12886_2_haplotype2-0000095_chr15, 1794-DEL, HSat3

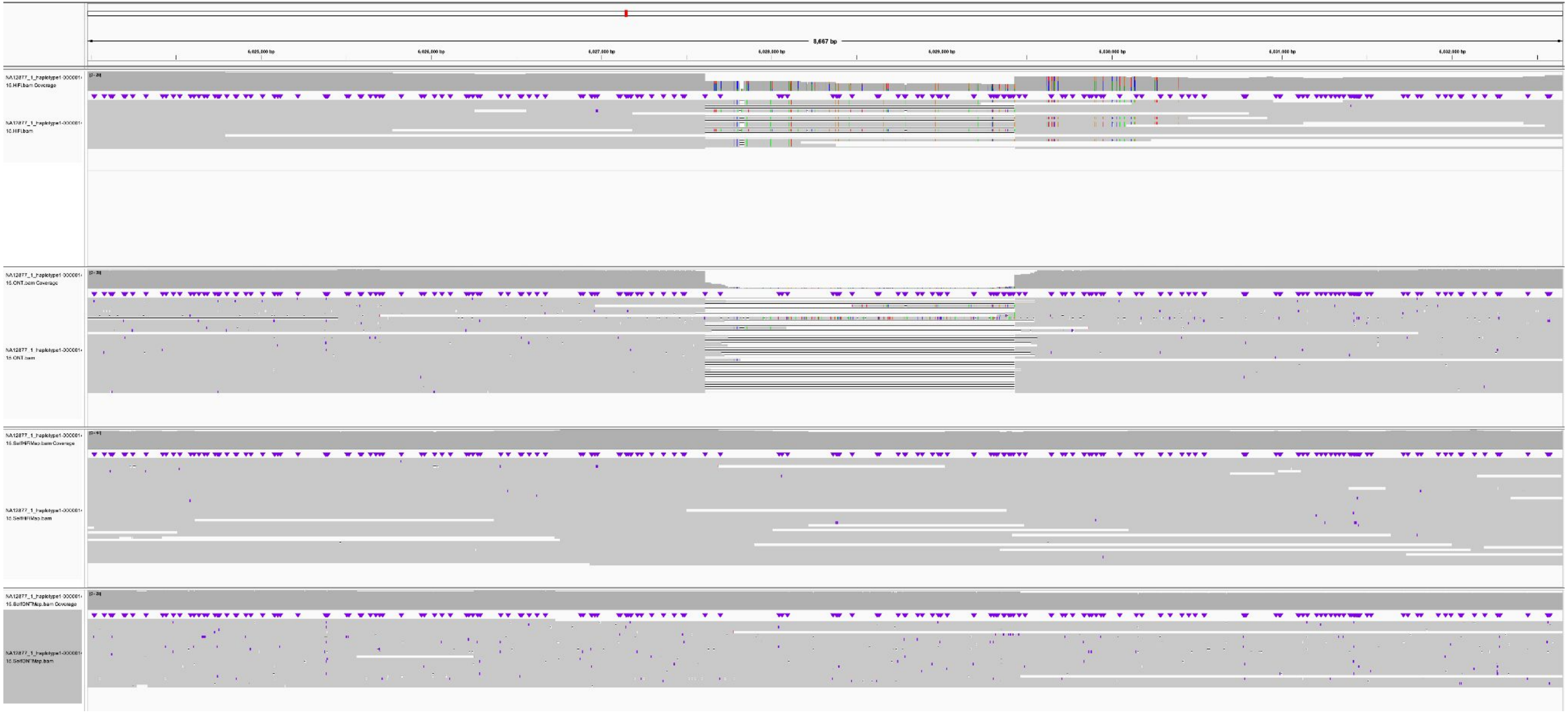

NA12877_2_haplotype2-0000070_chr22, NA12881_1_haplotype1-0000024_chr22, 6763-DEL, HSat1A

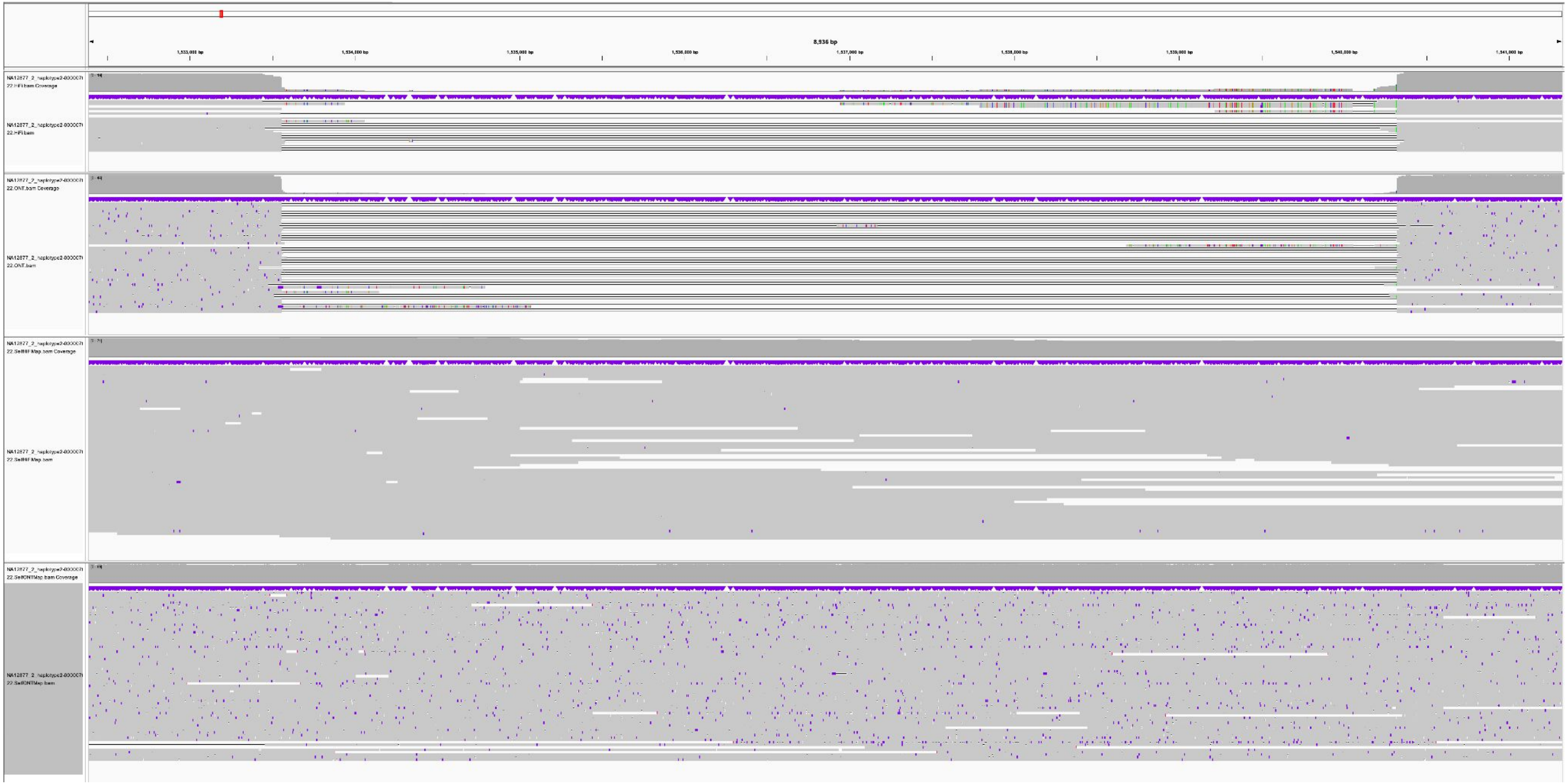

### NA12877_1_haplotype1-0000020_chr22, NA12884_2_haplotype2-0000071_chr22, 1362-DEL, aSat_HOR

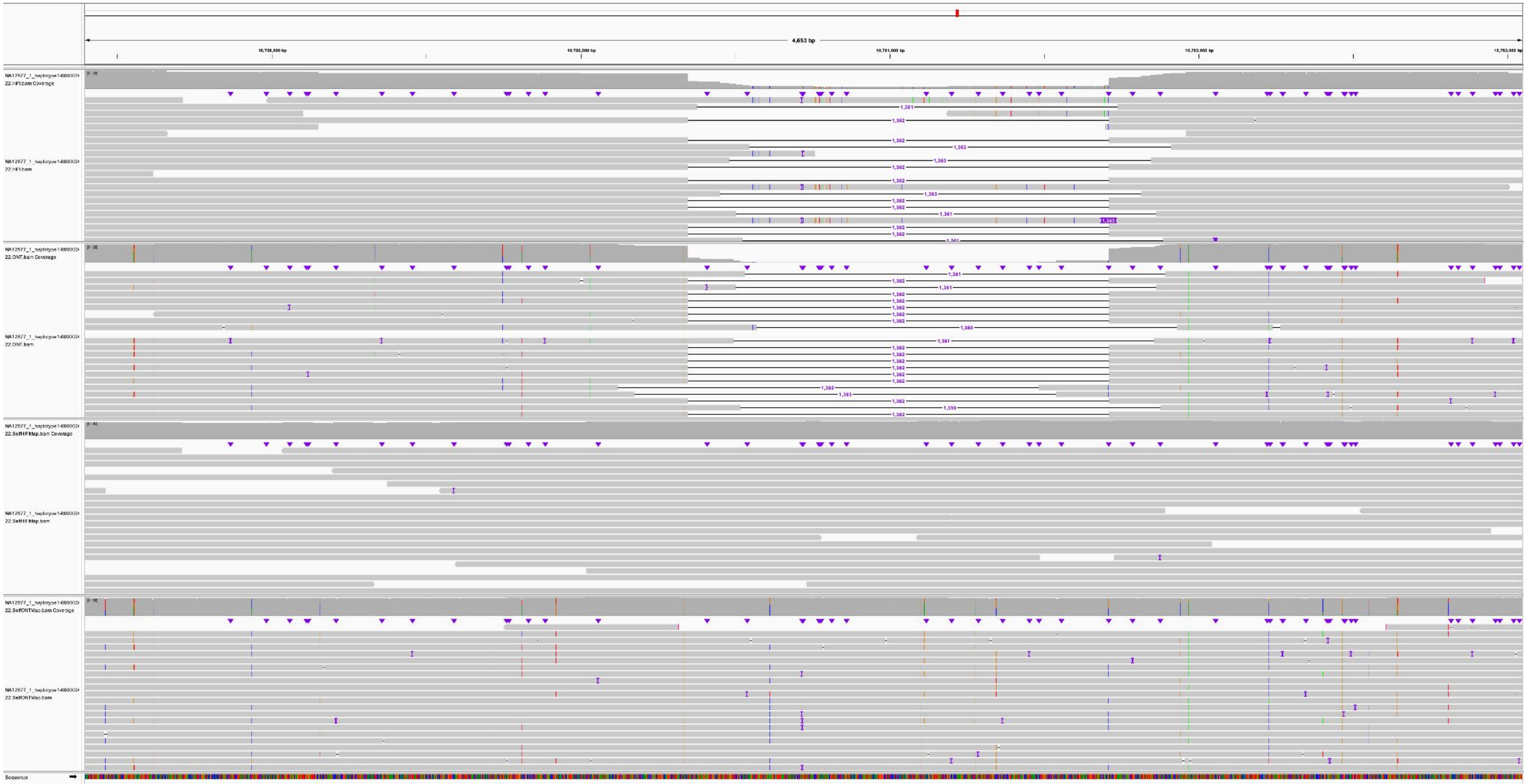

### NA12878_1_haplotype1-0000008_chr22, NA12887_2_haplotype2-0000080_chr22, 1364-DEL, aSat_HOR

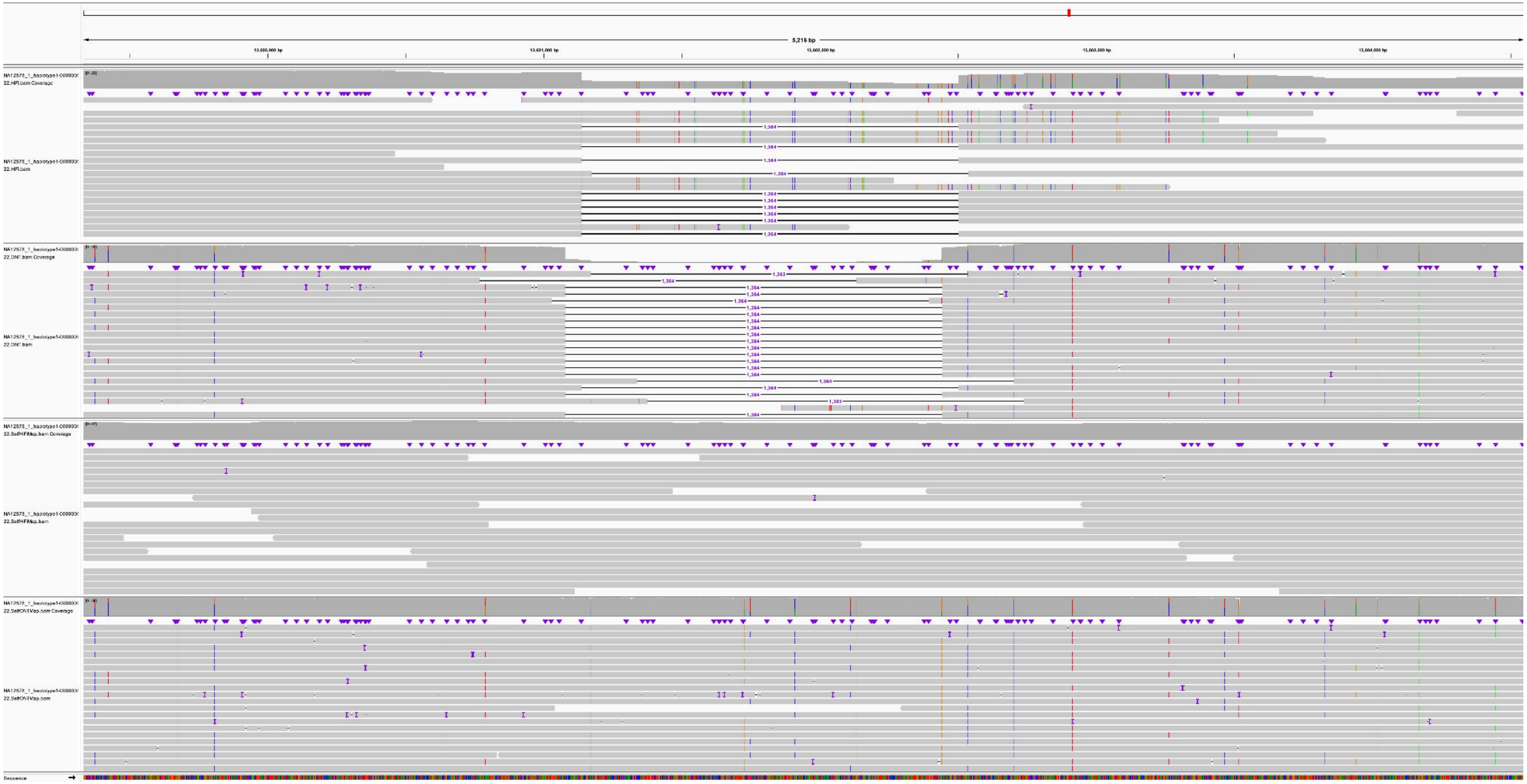

NA12879_2_haplotype2-0000083_chr14, K200084_2_haplotype2-0000078_chr14, 135-INS, HSat3

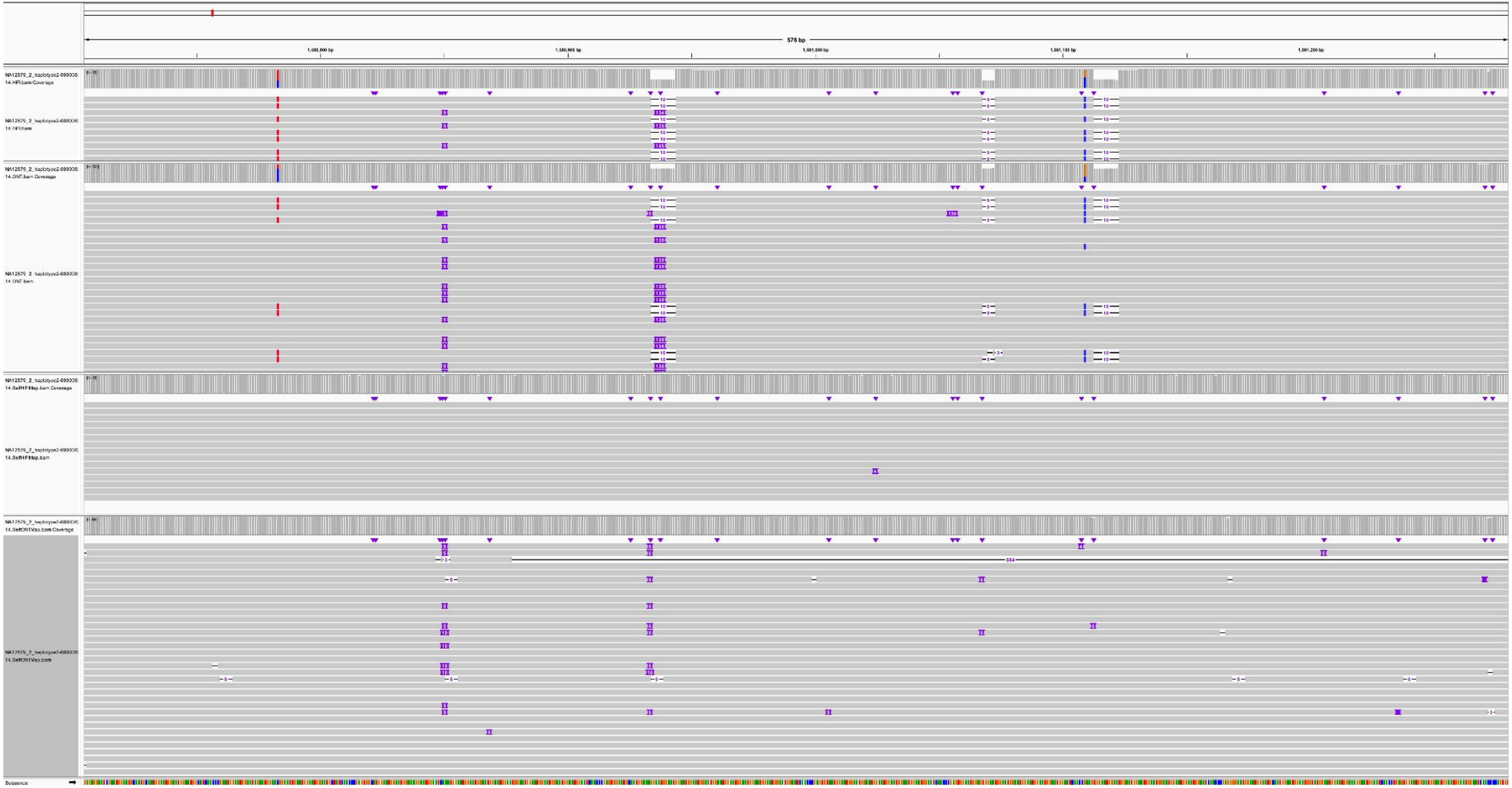

K200080_1_haplotype1-0000024_chr13, K200081_2_haplotype2-0000065_chr13, 1407-DEL, SST1

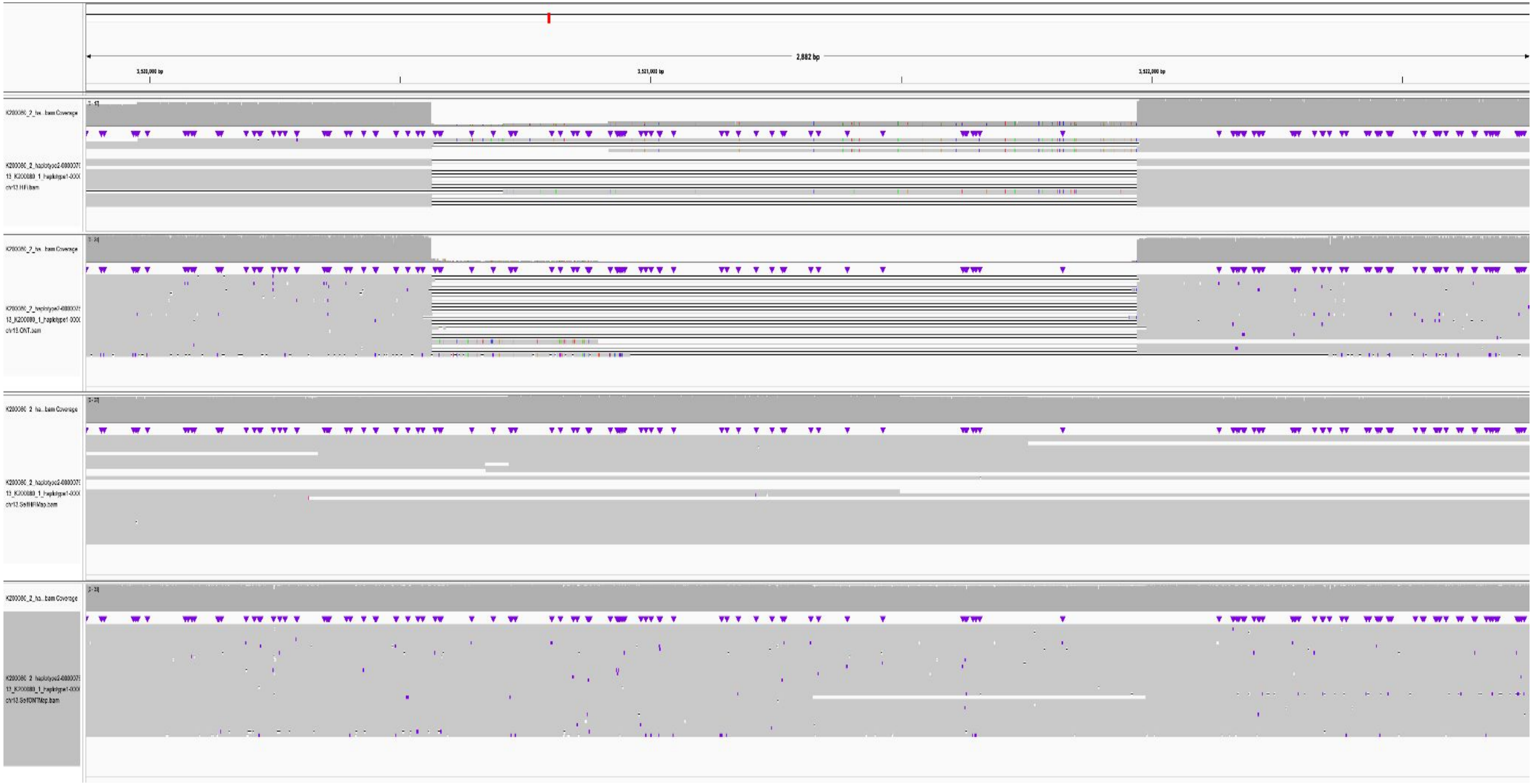
